# A living biobank of canine mammary tumor organoids as a comparative model for human breast cancer

**DOI:** 10.1101/2022.09.02.505845

**Authors:** Marine Inglebert, Martina Dettwiler, Kerstin Hahn, Anna Letko, Cord Drögemüller, John Doench, Adam Brown, Yasin Memari, Helen Davies, Andrea Degasperi, Serena Nik-Zainal, Sven Rottenberg

## Abstract

Mammary tumors in dogs hold great potential as naturally occurring breast cancer models in translational oncology, as they share the same environmental risk factors, key histological features, hormone receptor expression patterns, prognostic factors, and genetic characteristics as their human counterparts. We aimed to develop *in vitro* tools that allow functional analysis of canine mammary tumors (CMT), as we have a poor understanding of the underlying biology that drives the growth of these heterogeneous tumors. We established the long-term culture of 24 organoid lines from 16 patients, including organoids derived from normal mammary epithelium or benign lesions. CMT organoids recapitulated key morphological and immunohistological features of the primary tissue from which they were derived, including hormone receptor status. Furthermore, genetic characteristics (driver gene mutations, DNA copy number variations, and single-nucleotide variants) were conserved within tumor-organoid pairs. We show how CMT organoids are a suitable model for *in vitro* drug assays and can be used to investigate whether specific mutations predict therapy outcomes. In addition, we could genetically modify the CMT organoids and use them to perform pooled CRISPR/Cas9 screening, where library representation was accurately maintained. In summary, we present a robust 3D *in vitro* preclinical model that can be used in translational research, where organoids from normal, benign as well as malignant mammary tissues can be propagated from the same patient to study tumorigenesis.

## Introduction

Human breast cancer (HBC) is the most frequent and deadly cancer in women worldwide^1^. Numerous laboratory animal models are available to study HBC progression and develop novel targeted therapies, but translating results to patients remains challenging^2,3^. Companion animals are increasingly considered in translational cancer research, as they face the same environmental risk factors as their owners, have an intact immune system, and are closer to humans than rodents when assessing a drug’s toxicity and efficacy^4^. Companion animals such as dogs and cats often develop mammary tumors spontaneously in their lifetime. Canine mammary tumors (CMT) are the second most diagnosed cancer in dogs, affecting mainly older female dogs^5^. They share various molecular and clinical aspects (natural history, prognosis) with HBC^6,7^. CMT represent a highly heterogeneous disease with many histological subtypes, and about half of them are classified as malignant^8^. Interestingly, individual dog patients frequently develop multiple CMT with different histological subtypes located in different mammary complexes^9^. Contrary to HBC mainly arising from epithelial cells, malignant CMT comprise either one neoplastic cell type (simple carcinomas) or two cell types (complex carcinomas defined by a malignant epithelial component and a benign myoepithelial one).

To complement the molecular and pathological analysis of CMT at the functional level, representative CMT models that can be generated efficiently are required. Few CMT cell lines are available and do not capture the diversity of the disease^10–12^. Recently, the development of 3D organoids (ORG) allowed the modeling of many diseases, including cancer (colon, ovarian, HBC’…), where the organoids recapitulate the epithelial architecture and physiology of their organs of origin^13–16^. In dogs, organoid cultures have been established from the liver, mammary tissue, prostate cancer, epidermis, intestines, kidney, bladder cancer, and thyroid follicular carcinoma^17–24^. Here, we aimed to establish a biobank of CMT, including stable organoid lines derived from different epithelial origins. In particular, our goal was to develop new tools using CMT-derived organoids for functional analyses, study specific mutations, and investigate the differences between healthy and carcinoma tissues. In this context, genome engineering techniques such as CRISPR-based gene modification allow the study of specific mutations in biological processes and can be applied to organoid cultures^25–28^.

In this study, we present a protocol enabling the efficient derivation and long-term expansion of CMT organoids, including organoids derived from normal mammary epithelium or benign lesions. We comprehensively characterize our representative resource of CMT and organoids derived thereof in terms of morphology, histology, and genomic features, and we propose a novel preclinical model for HBC that can be used in translational research to study tumorigenesis.

## Results

### Patient cohort

We collected 78 CMT from 49 patients: 40 malignant tumors from 32 patients and 38 benign ones (adenomas, mixed tumors) from 30 patients (17 patients presenting exclusively benign tumors, 13 patients presenting both malignant and benign tumors). Supplementary Table S1 summarizes the clinical information and histopathological diagnosis of all patients. The mean age at diagnosis was 10.8 years. Most dogs were intact at the time of diagnosis (37/49), in line with the protective effect of castration on CMT development^6^. There was no breed predisposition in this cohort. We focused on the 32 patients affected with malignant tumors for further analysis and characterization (Supplementary Table S2). Many histological subtypes were represented (including tubular, anaplastic, comedocarcinoma, and complex carcinomas), matching the diversity of CMT^8^. The grading system yielded an average grade of 1.62^8^, and the TNM classification (based on tumor size, regional lymph nodes’ invasion, and distant metastasis) gave an average of 2.1^29^ When we analyzed our collection of CMT using the Nielsen classification, we found that 24% (8/33) of the tumors are triple-negative basal-like, and 25/33 tumors (76%) are luminal A^30,31^.

### Establishment of a living biobank of CMT organoids

Cryopreserved CMT tissues were thawed to generate organoids. CMT organoid proliferation rates varied strongly between different organoid lines (passaging intervals varied from two to four weeks and split ratios from 1:1 to 1:3). We generated multiple organoid lines from different epithelial origins (Figure 1A), summarized in Supplementary Table S3. In total, 24 organoid lines derived from carcinoma, adenoma, or non-neoplastic mammary tissues from 16 patients were successfully generated (Figure 1B), representing the diversity of CMT. Organoids could be passaged for an extended period (thirteen lines were passaged more than eight times and two lines more than 20 times without losing proliferation capacity; Figure 1C) and efficiently recovered following cryopreservation (Supplementary Table S3).

**Figure 1:**
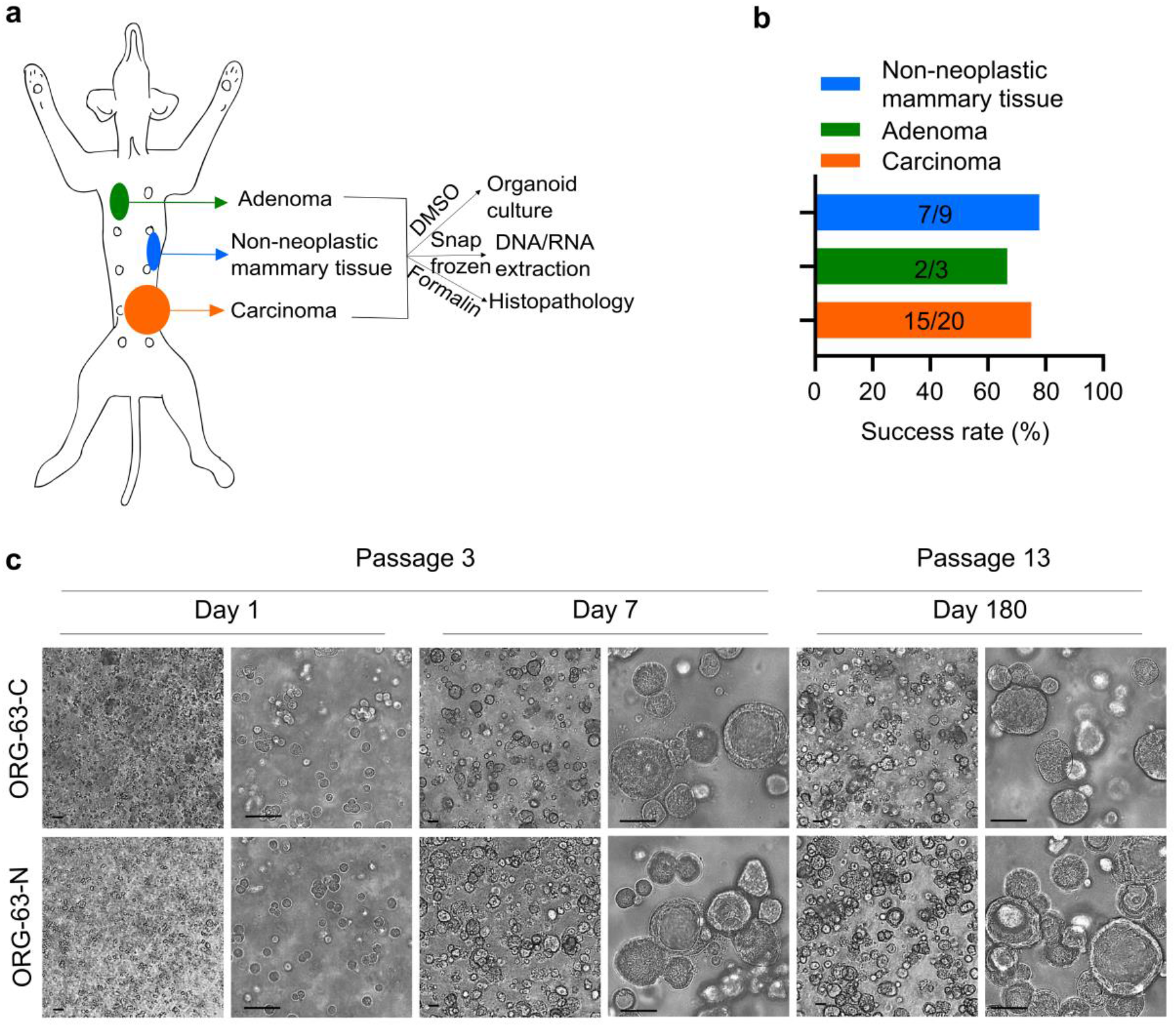
Establishment of a living biobank of CMT organoids. a. Sampling and generation of organoids from different primary canine epithelial mammary tissues from the same patient: malignant (carcinomas), benign (adenomas), or non-neoplastic mammary tissues. b. Success rates for establishing 3D *in vitro* organoids from the indicated mammary tumors and normal mammary epithelium. Values indicate the number of donor tumors from which models were successfully derived versus the total number of donor tumors for a total of 16 patients. c. Brightfield images of 3D organoids of mammary tumor (ORG-63-C) and of normal mammary epithelium (ORG-63-N) grown in Basement Membrane Extract 1 day, 7 days (passage 3), and 180 days (passage 13) following isolation. Both organoid lines are derived from patient 63. Scale bar, 50 μm.

### Morphological and histological characterization of CMT organoids

To compare the organoids with their tissue of origin, we performed morphologic phenotyping of hematoxylin and eosin (H&E)-stained tissues. CMT organoids showed a heterogeneous morphology displaying both compact (Figure 2A, green arrows) and cystic (Figure 2A, orange arrows) organoids recapitulating primary tissue structures. Acini structures were conserved (single- or double-layered). Organoids presented tumor characteristics such as cellular atypia, pleomorphism, and vacuolization, and some presented squamous differentiation. Some organoids derived from non-neoplastic mammary tissue showed secretory activity (Figure 2A, blue arrows), consistent with the physiology of their tissue of origin.

**Figure 2:**
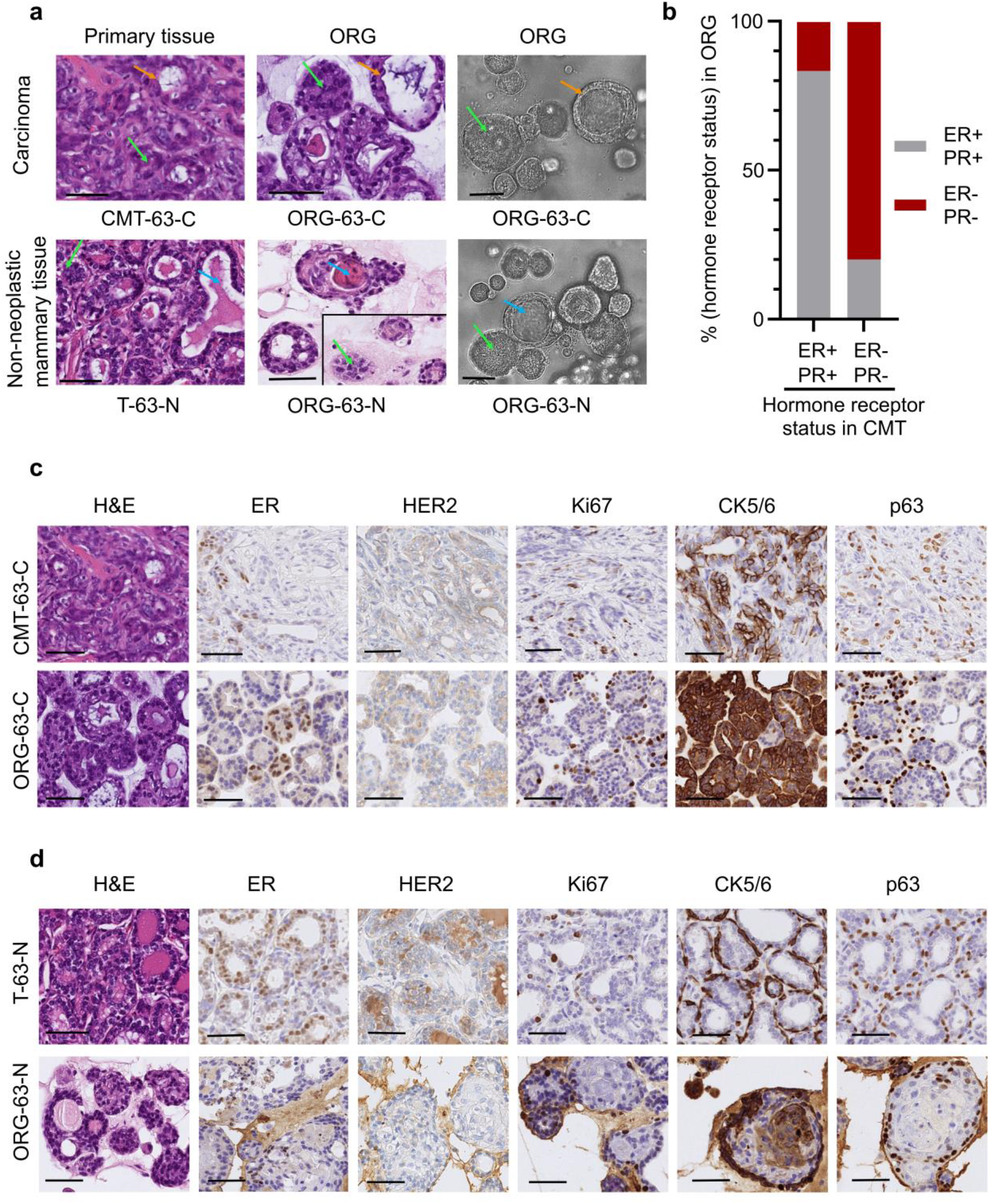
CMT organoids share important morphological characteristics and hormonal status with their primary tissue. a. Representative images of H&E stainings of primary tissue (first column), patient-derived organoids (second column), and brightfield images (third column) of 3D organoids of normal mammary epithelium (bottom line) and of mammary tumor (top line) grown in Basement Membrane Extract. The green arrows indicate solid organoids, and the orange ones indicate cystic organoids; the blue arrows indicate luminal secretion. For comparison purposes, two different areas of the same scanned slide are represented for the H&E image of ORG-63-N. Scale bar, 50 μm. b. Histogram showing the distribution of organoids that are hormone receptor-positive (brown) and negative (grey) grouped per original carcinoma receptor status. c. Representative images of H&E stainings and immunohistochemical analyses of estrogen receptor (ER), human epidermal growth factor receptor 2 (HER2), Ki67 (marker of cell proliferation), cytokeratin (CK) 5/6 (luminal epithelial cells), and p63 (basal cells) in CMT-63-C tumor and tumor-derived organoids ORG-63-C. Scale bar, 50 μm. d. Representative images of H&E stainings and immunohistochemical analyses of estrogen receptor (ER), human epidermal growth factor receptor 2 (HER2), Ki67 (marker of cell proliferation), cytokeratin (CK) 5/6 (luminal epithelial cells), and p63 (basal cells) in normal mammary epithelium (T-63-N) and tissue-derived organoids ORG-63-N. Scale bar, 50 μm.

Next, we evaluated known HBC and CMT markers for myoepithelial cells (vimentin (VIM) and p63), cytokeratin (CK) (including luminal and basal markers CK7/8, CK5/6, CK14), proliferation index (Ki67), human epidermal growth factor receptor 2 (HER2), and hormone receptors estrogen (ER), and progesterone (PR) according to standard guidelines^32^ (Supplementary Table S3). In 82% of the cases (14/17 tumors), the organoids recapitulated the Nielsen classification of their primary tumors with high fidelity. No HER2-positive (*i.e*., HER2 scoring of 3+) tumor/tissue was collected, which was conserved in the organoids. Hormone receptor status is of fundamental importance in HBC as well as in CMT^33^. CMT positive for ER/PR led to organoids positive for ER/PR in 83% of the cases; CMT negative for ER/PR led to organoids negative for these markers in 80% of the cases (Figure 2B; Supplementary Table S3).

A matching score between tissues and organoids was established: poor (1) when the Nielsen classification differed between primary tissues and organoids or more than three biomarkers showed substantial differences in the percentage of cell positivity, medium (2) when the classification was conserved but more than two biomarkers showed substantial differences in the percentage of cell positivity, good (3) when the Nielsen classification was conserved, and most biomarkers showed the same trend. 7/24 organoid lines (29%) scored 3, 10/24 organoid lines (42%) scored 2, and 7/24 (29%) organoid lines scored 1. Tumor heterogeneity and the fact that only a tumor fragment was used for organoid generation may account for the differences. Ki67 proliferation marker varied greatly between organoid lines (from 1% to 87.7%), underlining CMT’s cell cycle regulation heterogeneity^34^. Moreover, it suggests that different CMT subtypes may require individualized optimization of the culture medium.

In summary, we found that most organoids (derived from malignant (Figure 2C), benign or non-neoplastic (Figure 2D) epithelial tissues) match their primary tissue regarding morphology, histopathology, and important biomarkers such as HER2 and hormone receptor status.

### Genetic characterization of CMT

As the genomic data concerning CMT is scarce compared to HBC, we set out to characterize part of our CMT cohort with Whole Genome Sequencing (WGS). When we had 21 malignant CMT available in the project, we performed WGS (coverage of 30X). By comparing the tumor sample and matched normal sample, a total of 15550 single-nucleotide substitutions, 6035 short insertions/deletions (indels), and 118 rearrangements were identified. For these CMT, there was a median of 603.5 substitutions (mean of 706.8), 268 indels (mean of 274.3), and 4 rearrangements (mean of 6.5). In comparison, HBC shows a much higher number of substitutions and rearrangements. For example, in a repository of 560 HBC, a median of 3491.5 substitutions (mean of 6213.7), 204.5 indels (mean of 664.3), and 85 rearrangements (mean of 138.8) were found^35^. We then investigated commonly mutated genes in CMT and HBC. The mutational landscape of the 21 CMT is represented in Figure 3A. Mutations of *PIK3CA* are the most frequent (6/21), and they occur at known hot spots (H1047R, G118D, R108Q) in HBC and CMT, leading to activation of the PI3K-AKT pathway^36^. Further downstream in this signaling pathway, we found mutations of *AKT1* with the E17K hotspot shared both by humans and dogs^37,38^. Of note, only complex carcinomas displayed mutations of *AKT1* (0/13 mutations in simple carcinomas and 5/8 mutations in complex carcinomas; P=0.0028, Fisher’s exact test), consistent with previously described data^37^. This indicates that the activation of the PI3K-AKT pathway plays a fundamental role in CMT tumorigenesis, similar to part of the HBC. *KRAS* mutations were found in 2 cases in known hotspots (G12D/R) shared by CMT and HBC^37^. Mutations in *TP53*, a common tumor suppressor gene mutated in various human cancers, were found only in one case at the location of the DNA binding domain (P268T). Moreover, mutations affecting intronic regions present in more than 40% of the cases for the simple carcinomas but not found in the Cancer Mutation Census (Catalogue Of Somatic Mutations In Cancer = COSMIC) for HBC, were identified for *ZNF511, DIAPH2, SORCS3, CNTNAP5, NFIA, ADGRL3, DGKB* (Figure 3A, genes specifically mutated in CMT).

**Figure 3:**
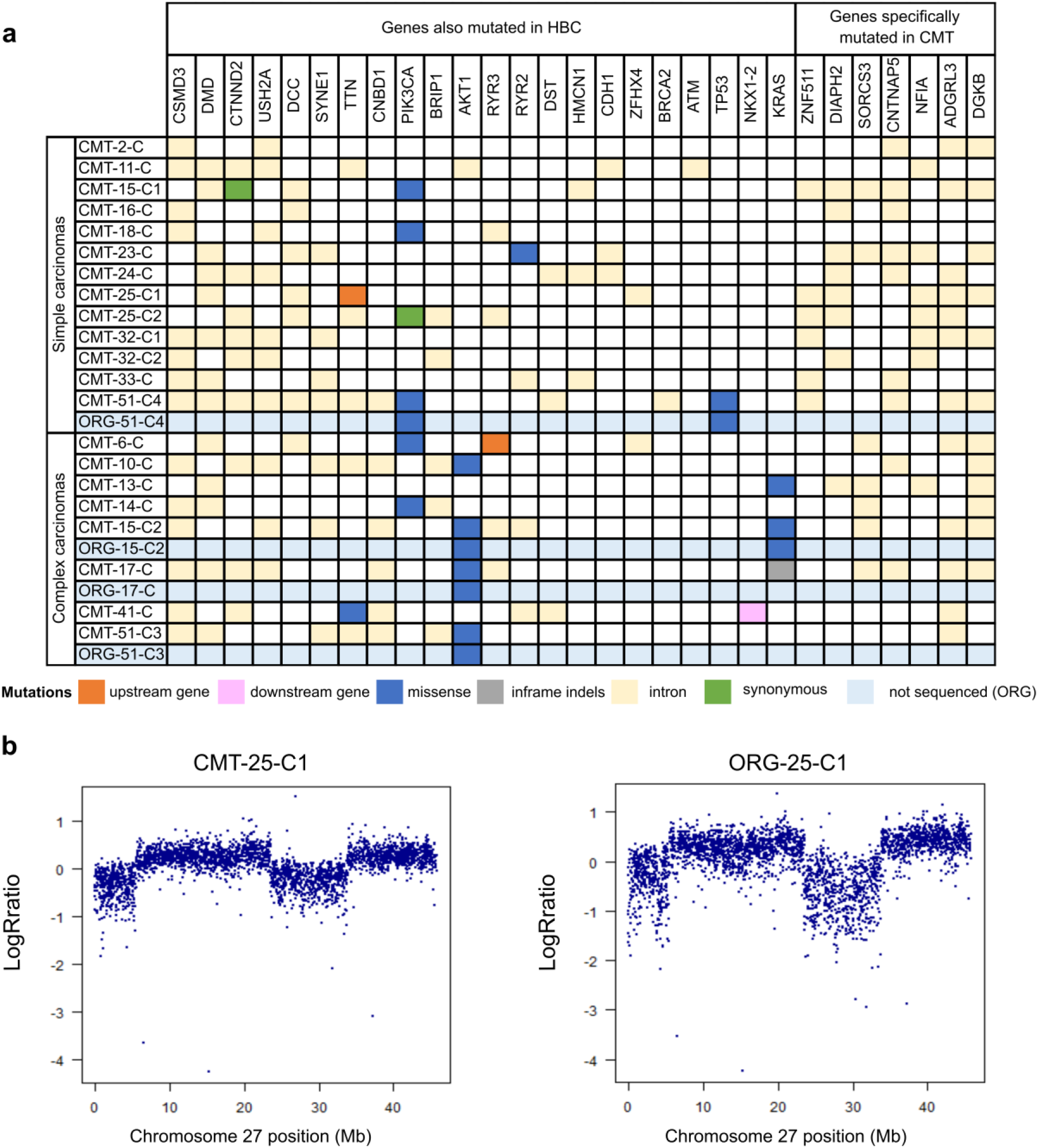
The genetic landscape of CMT is conserved in patient-derived organoids. a. Overview of somatic mutations of 21 CMTs for 29 mutated genes, with differentiation between simple and complex carcinomas. Six mutation types are represented according to the color legend. Point mutations of the corresponding organoids for the mentioned gene (*PIK3CA, AKT1, TP53, KRAS*) are represented (light blue lines). b. Log R ratios markers on chromosome 27 for matched pair CMT-25-C1 / ORG-25-C1. Note the conservation of the copy number variation in the organoid line. See also Supplementary Figure S2A.

We then examined single base substitution mutational signatures (SBS) present in this cohort^39^, which revealed SBS1 in nearly all CMT (17/21, Supplementary Table S4). This SBS, related to the age at diagnosis, is also present in HBC and many other cancer types. However, other SBS usually found in most HBC (associated with APOBEC cytidine deaminases or homologous recombination repair deficiency) were only sporadically found in this cohort (Supplementary Table S4). Finally, using a combined fit and extraction algorithm, an unknown dog signature was isolated and appeared similar to SBS57 (Supplementary Figure S1). With the available data, no other novel dog-specific SBS was detected. Hence, our WGS analysis revealed substantial differences in the mutational landscape of CMT compared to HBC. Although the number of our malignant CMT is limited (n=21), it clearly shows that the number of mutations is much lower in CMT. Moreover, HER2 amplifications were not observed, and several SBS frequently present in HBC are rare in CMT. Instead, age-related deamination of methyl cytosines seems to be the dominant genotype.

### Genomic characterization of CMT organoids

To assess whether organoids conserve genetic characteristics of their primary tissue, we next performed an single-nucleotide polymorphism (SNP) array analysis and genotyped a subset of 15 organoid lines (which could be expanded for more than eight passages, yielding enough cells for DNA extraction), and their matched primary tissues (Supplementary Table S3). Most SNP genotypes present in the primary tissue were conserved in the derived organoids (less than 0,5% difference in shared genotypes for most of the lines, Supplementary Figure S2G). Two lines (ORG-33-C and ORG-51-C4) showed around 7,5% genotypic differences with their primary tissue. Those two lines proliferate much faster than the other organoid lines, which might lead to a higher frequency of mutations arising during cell division and therefore explain this difference. This analysis also allowed for the identification of outliers, *i.e*., swapped samples (which happened in the case of ORG-05-C, easily noticed as it shares only around 60% identical genotypes with its supposed tissue of origin). This reinforces the importance of regular controls when establishing primary cultures and simultaneously handling samples from many different patients.

The copy number variation profile obtained after calculating the log R ratio (LRR) revealed similar patterns between organoid/tissue pairs. For example, the detail of chromosome 27 of the matched pair CMT-25-C1/ORG-25-C1 shows an increase in LRR values (Figure 3B) and a splitting of heterozygous genotype clusters into two clusters in the B-allele frequency (BAF) plots, indicating two larger-sized (>10 Mb) duplication events (Supplementary Figure S2A). These events are conserved in the ORG-25-C1. Overall, the organoid lines conserved the genomic heterogeneity of their different epithelial origins (for an overview, see Supplementary Figure S2D,E,F). Moreover, SNP array genotyping was also performed for organoids of later passages, revealing the conservation of the genomic landscape after a prolonged time in culture (Supplementary Figure S2B,C).

To further characterize the genetic landscape of CMT organoids, we performed targeted sequencing and analyzed somatic single-nucleotide variants of interest. Driver gene point mutations in CMT were maintained in the derived organoid lines (Figure 3A, blue columns). A more careful analysis of the chromatograms revealed a mixed population of the normal reference allele and the mutated base in the tumor tissue, whereas the organoid population showed enrichment in the mutated base (Supplementary Figure S2H). This is in accordance with the fact that tumor pieces also contain non-tumorous cells (*i.e*., stromal cells), and the organoid culture selects for the epithelial cells.

In summary, we show that organoids derived from CMT and non-neoplastic mammary tissues recapitulate the genetic characteristics of the primary epithelial tissue, even after extended passaging.

### Drug testing in CMT organoids

To investigate whether this model may be helpful to test anti-cancer therapies, we tested different drugs *in vitro* commonly used for HBC and canine cancer treatment. The drug panel included platinum drugs (cisplatin, carboplatin), doxorubicin^40^, and an inhibitor of the PI3K/AKT pathway, alpelisib, shown to prolong progression-free survival among patients with *PIK3CA-mutated*, ER/PR-positive, HER2-negative HBC^41^. Cell viability assays generated reproducible dose-response curves. IC50s for cisplatin, carboplatin and doxorubicin in all organoid lines tested were compatible with drug concentrations tolerated in patients (Supplementary Figure S3A,B,C)^42^. As expected, *PIK3CA*-mutated organoid lines (ORG-51-C4, ORG-MCF7) were more sensitive to alpelisib than wild-type organoid line ORG-51-C3 (Figure 4A). Since the *TP53* gene is an important driver gene in HBC as well as in CMT, we tested whether nutlin-3a, an inhibitor of the MDM2-TP53 interaction, can distinguish between wild-type and *TP53-*mutated organoids. As expected, all organoid lines were sensitive to nutlin-3a treatment (Figure 4B), except for ORG-51-C4, a *TP53-*mutated organoid line (Supplementary Figure S2H). Hence, CMT organoids can be used to test various drugs and thereby investigate whether specific mutations predict therapy outcomes.

**Figure 4:**
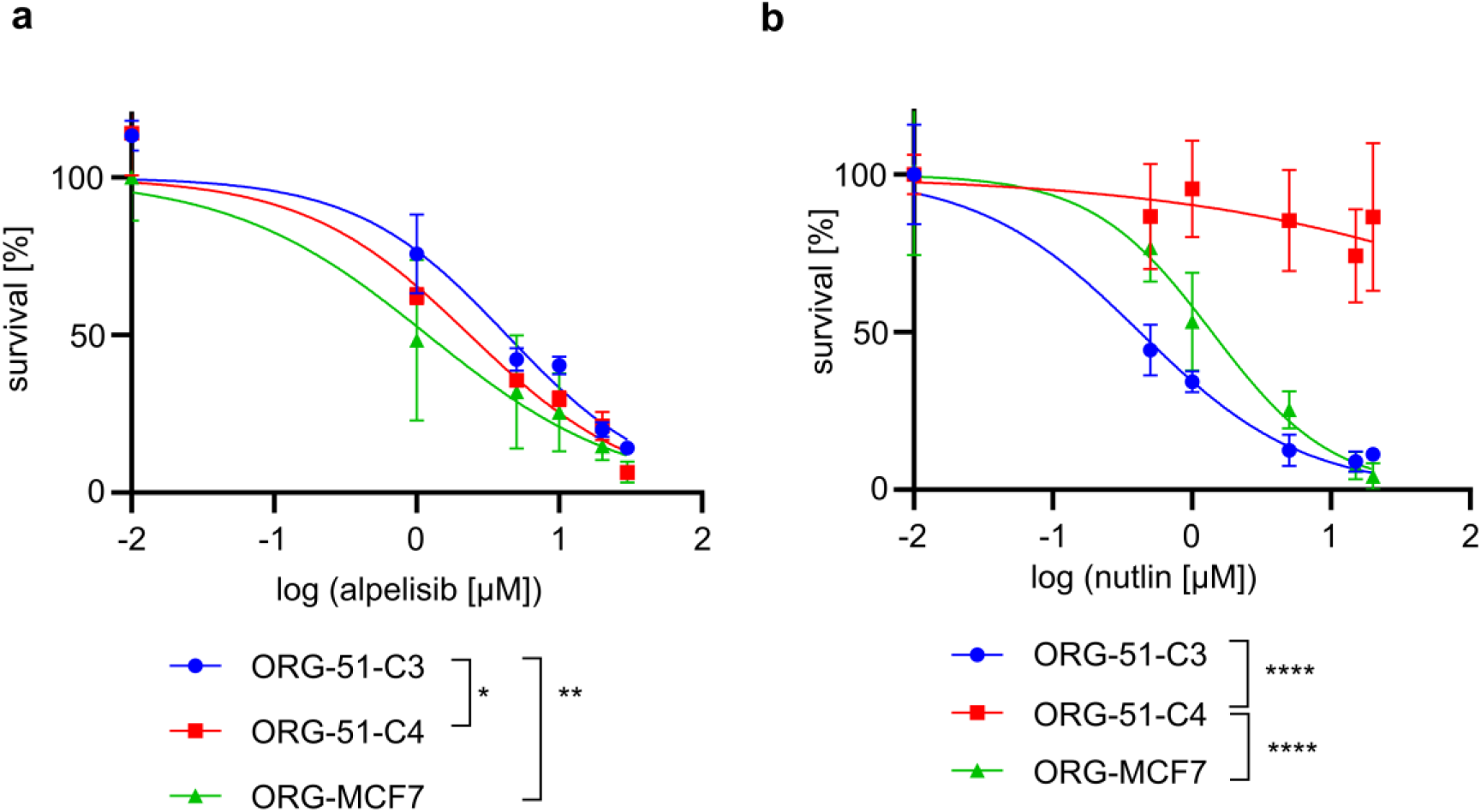
CMT organoids allow *in vitro* drug testing. a. Dose-response curves indicating viability 8 days after treatment with alpelisib for *PIK3CA*-mutated organoid lines (ORG-51-C4, ORG-MCF7) and wild-type organoid line ORG-51-C3. Error bars represent the standard deviation (SD) of three independent experiments. P-values are calculated by one-way ANOVA followed by Tukey’s multiple comparisons test for the log(IC50) values of the survival curves, * P=0.0363, ** P= 0.0092 As a control, the HBC *PIK3CA*-mutated, *TP53* wild-type MCF-7 cell line was used and grown in BME to form organoids (ORG-MCF7) following the same conditions as the CMT organoids. b. Dose-response curves indicating viability 8 days after treatment with nutlin-3a for *TP53*-mutated organoid line ORG-51-C4 and wild-type organoid lines (ORG-51-C3, ORG-MCF7). Error bars represent SD of three independent experiments. P-values are calculated by one-way ANOVA followed by Tukey’s multiple comparisons test for the log(IC50) values of the survival curves, **** P<0.0001 As a control, the HBC *PIK3CA*-mutated, *TP53* wild-type MCF-7 cell line was used and grown in BME to form organoids (ORG-MCF7) following the same conditions as the CMT organoids.

### Gene editing and CRISPR/Cas9 screening of CMT organoids

To examine the experimental potential of organoids to study mechanisms of tumorigenesis or therapy response, we tested gene modification techniques for organoids derived from CMT and non-neoplastic mammary tissues. Using a green fluorescent protein (GFP) encoding lentivirus, we found that a multiplicity of infection (MOI, representing the number of viral particles per cell) of 1 resulted in 12,6% of cells expressing GFP (Figure 5A). The transduction efficacy increased to 35,3% with an MOI of 4 (Figure 5A). To demonstrate that the organoids can be genetically modified in a stable manner, we spinoculated the ORG-25-C1 line with a lentiviral vector into which we cloned a guide RNA targeting the *VIM* gene (gVIM-1, gVIM-2). Vimentin is not essential and can therefore be knocked out without impeding organoid growth. After selection, the surviving organoids were expanded and analyzed for *VIM* mutations (Figure 5B). More than 90% of the polyclonal population showed frameshift mutations in the *VIM* gene, and almost no protein was detected (Figure 5C).

**Figure 5:**
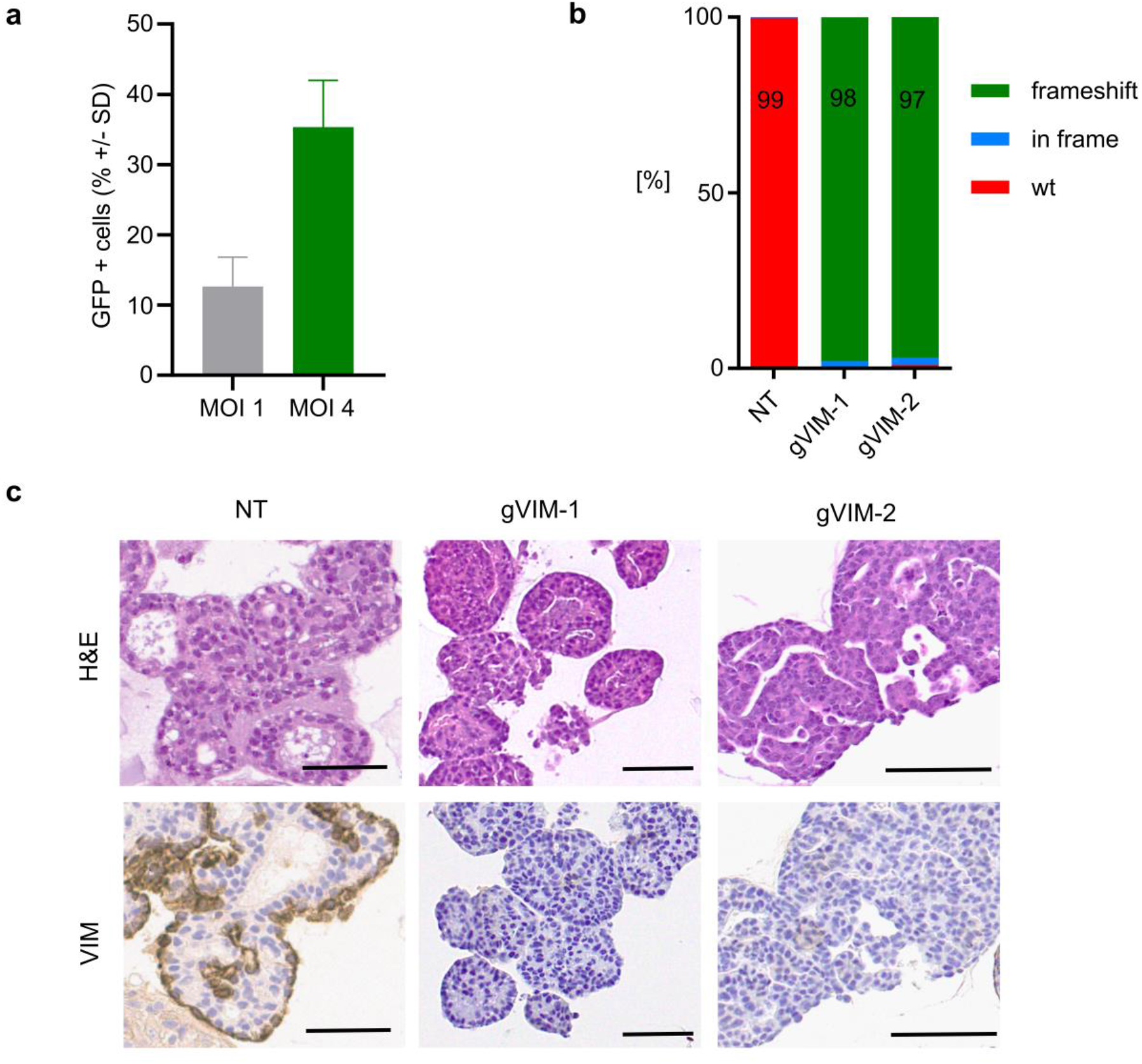
CMT organoids can be genetically modified with the CRISPR/Cas9 system. a. GFP was introduced into ORG by lentiviral transduction at the indicated multiplicities of infection (MOI). GFP expression was analyzed by flow cytometry 3 days after transduction. For MOI 1, Mean ± SD = 12.6 ± 4.2%, for MOI 4, Mean ± SD = 35.3 ± 6.6%. Two independent experiments that were conducted with three different organoid lines are presented, demonstrating robust transduction efficiency. b. Frequency of frameshift indels in organoids ORG-25-C1 modified by CRISPR/Cas9 using a gRNA targeting vimentin (gVIM-1 and gVIM-2), compared to the non-targeting (NT) gRNA. TIDE analysis two passages after transduction. c. Representative images of H&E stainings and immunohistochemistry of VIM in organoids ORG-25-C1 modified by CRISPR/Cas9 using a gRNA targeting vimentin (gVIM-1 and gVIM-2), compared to the non-targeting (NT) gRNA. Embedding four passages after transduction. Scale bar, 50 μm.

To test whether functional genetic screens could be performed at a larger scale, we transduced two organoid lines from the same patient (ORG-63-N and ORG-63-C) with a customized canine CRISPR/Cas9 sublibrary that we designed to target druggable genes, following standard guidelines for single-guide RNA (gRNA) optimization (Figure 6A)^43^. This library contains 6004 gRNAs, targeting 834 genes (six gRNAs/gene, see Supplementary Table S5) and was cloned into the lentiCRISPRv2 (pXPR_023) one vector system containing Cas9. We aimed to express the library in CMT-derived organoids at a coverage of 500 cells per gRNA. After twelve days of puromycin selection (necessary for the organoids to recover from the transduction and survive the following trypsinization), the distribution of the gRNA counts remained similar between the plasmid DNA (pDNA) and D0 (Figure 6B,C). In addition, when comparing gRNAs enrichment between D0 and pDNA, some gRNAs were clearly enriched for both organoid lines, matching genes known to bring a survival benefit for the cells when the gene is knocked-out like *TP53* and *CDKN2A* (Figure 6D,E). On the other hand, genes involved in essential processes, such as *SMC3*, involved in chromosome cohesion during mitosis, or *TET1*, involved in DNA methylation, were depleted (Figure 6D,E). Hence, we show that the CMT organoids can be transduced with customized CRISPR libraries, and the gRNAs’ representation is sufficient to investigate which genes are essential during growth.

**Figure 6:**
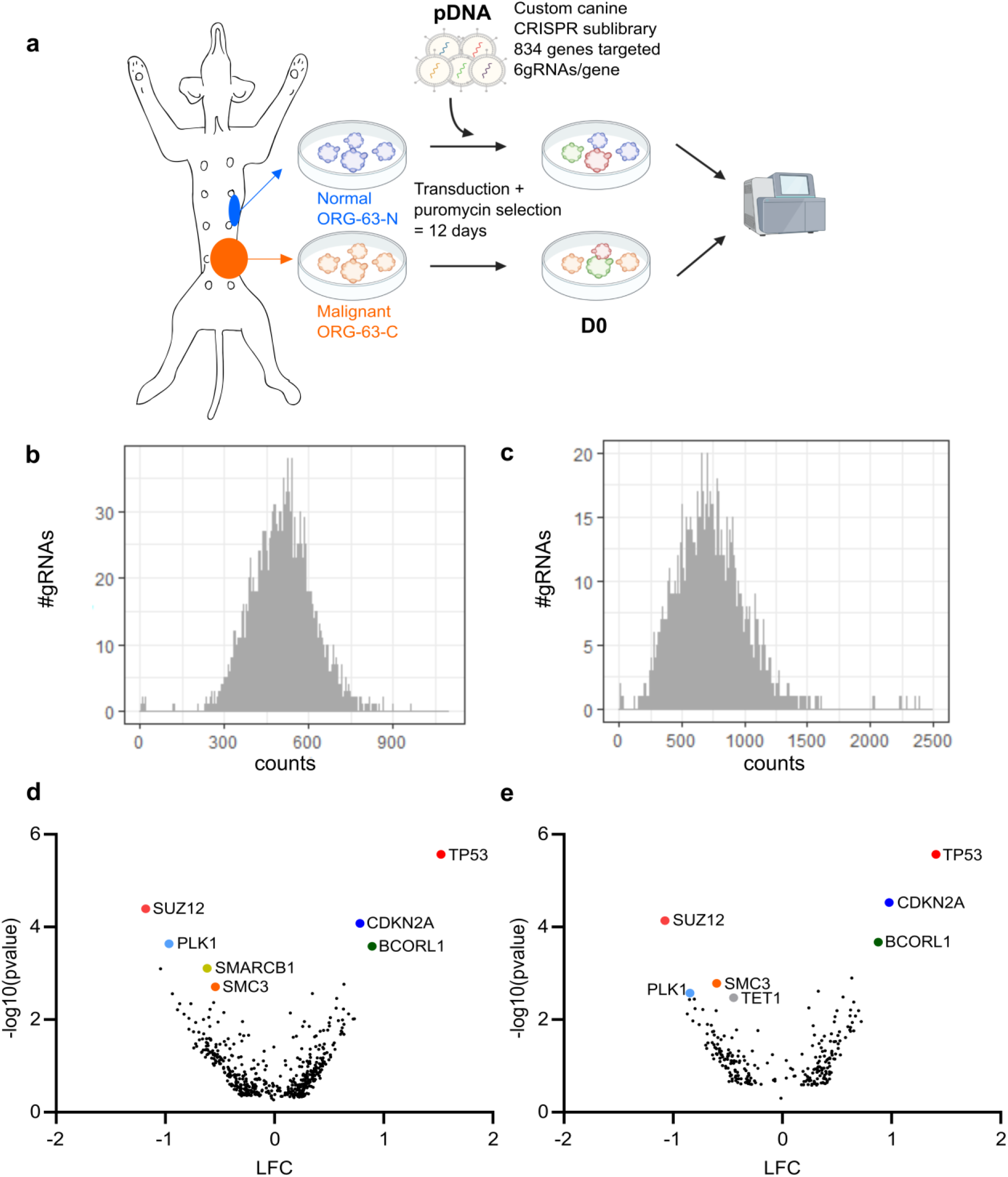
CMT organoids can be used to perform large-scale CRISPR/Cas9 screening. a. Outline of the screen performed with a custom CRISPR/Cas9 library. b. Histogram representing the distribution of the gRNA counts for the plasmid DNA (pDNA). c. Histogram representing the distribution of the gRNA counts for ORG-63-N at D0 (replicate 1). d. Volcano plot representing depleted (Log Fold Change = LFC<0) and enriched (LFC>0) genes for ORG-63-N twelve days after transduction (D0 vs. pDNA). LFC and P-values were calculated from two independent replicates with MAGeCK analysis. Each dot represents one gene for which at least three gRNAs (out of 6) were significant. Top hits are labeled with gene names. e. Volcano plot representing depleted (Log Fold Change = LFC<0) and enriched (LFC>0) genes for ORG-63-C twelve days after transduction (D0 vs. pDNA). LFC and P-values were calculated from two independent replicates with MAGeCK analysis. Each dot represents one gene for which at least three gRNAs (out of 6) were significant. Top hits are labeled with gene names.

## Discussion

Despite numerous *in vitro* and *in vivo* models established to study HBC, translating the research results to patients remains challenging^2,3^. Therefore, new models are warranted, and focusing on spontaneous models of HBC offers a new angle. Here, we established long-term 3D organoid cultures from CMT as a tool to functionally study tumorigenesis and therapy response in a spontaneously developing mammary tumor model for HBC.

In contrast to human samples, using different naturally occurring CMT from the same patient overcomes the bias of inter-individual genetic variability. Moreover, we can derive organoid lines from normal mammary tissue of the same dog, for which we also have tumor samples. This is usually difficult with human samples, where normal mammary tissues are obtained from preventive surgery and originate from another patient. Genetic editing of tumor organoids represents an opportunity to study carcinogenesis, as was performed in colon organoids^27,44,45^. To the best of our knowledge, we show the first use of a custom CRISPR library in patient-derived organoids from both healthy and neoplastic mammary tissue. Using those unique organoids from individual patients will allow us to investigate which druggable genes are essential in the malignant tumor cells but not in the non-neoplastic epithelium and thereby design individualized therapeutic strategies. Moreover, the possibility of performing pooled CRISPR/Cas9 screening in both normal, benign, and malignant organoids from the same patient represents a unique way to study the differences between those tissues and specifically investigate carcinogenesis, following stepwise evolution from adenoma to carcinoma^9^.

Our comprehensive analysis shows that CMT organoids maintain tumor morphological characteristics and biomarker expressions, such as hormone receptor and HER2 status. Hormone receptor positivity and lack of HER2 overexpression/amplification is the most common feature in HBC (around 2/3 of the cases)^46^. CMT organoids showing the same features are therefore an invaluable preclinical model. At the genomic level, organoid/tissue pairs remain similar, and mutations in driver genes involved in CMT and HBC development (*PIK3CA, TP53*) are conserved in organoids.

There are, however, some limitations when using CMT as a model for HBC. We show in our cohort that the mutational burden in CMT is substantially lower than in HBC. This can be explained by a shorter disease development time as dogs usually develop CMT around the age of ten, leaving less time for the cells to accumulate mutations. Another striking difference between CMT and HBC lies in their SBS profiles. Although SBS1 is present in more than 75% of HBC and prevails in CMT, other prominent HBC signatures are rare in our cohort. We do not think this is due to the small number of CMT we sequenced (n=21), as different SBS specific to HBC were initially found in a small cohort (n=21)^47^. Our data conclude that typical HBC mutational signatures, such as those resulting from defects in APOBEC cytidine deaminases or homologous recombination repair deficiency, do not play a significant role in CMT carcinogenesis. Despite the lower number of mutations and SBS, substantial heterogeneity is seen in CMT phenotypes. This suggests that additional modifications at the epigenetic or posttranslational level play an important role. It may also explain why many CMT have co-evolved as complex carcinomas with a strong myoepithelial component.

CMT may offer a valuable model for specific HBC types, notably the *PIK3CA* mutated ER+ tumors. We currently lack solid genetically engineered mouse models to mimic ER+ HBC^48^. As around 2/3 of HBC are ER+, this is a severe limitation to our available model systems. This may also contribute to the observation that many preclinical studies in rodents only poorly predict outcomes of human clinical trials^49^. Here, the ER+ CMT and organoids may prove to be helpful. The Comparative Oncology Trials Consortium (within the US National Cancer Institute) executes multi-institutional clinical trials in companion animals with naturally occurring cancer (for a review, see^50^). The shorter lifespan of dogs allows for a quicker clinical trial completion than in humans. One could envision preclinical studies based on CMT organoids implemented in a canine clinical trial. For example, a combination of alpelisib and paclitaxel, which shows promising results *in vitro* and *in vivo* for *PIK3CA* mutated human gastric cancer^51^, could benefit *PIK3CA* mutated ER+ HBC patients not responding to endocrine therapy. Testing this combination, first in *PIK3CA* mutated ER+ CMT organoids to predict efficacy, secondly in canine patients presenting the same subtype to assess the safety and validate efficacy, would lead to translatable results benefiting both canine and human patients.

In summary, this unique model relying on stable patient-derived organoids developed from spontaneous CMT opens many possibilities in translational research. Specifically, performing genetic screening on multiple organoid lines from different epithelial origins derived from the same patient will bring the understanding of mammary tumorigenesis to another level. In addition, specific CMT subtypes, such as the *PIK3CA* mutated ER+ ones, can be of value to study not only *in vitro* but also to set up a preclinical and clinical model highly relevant to HBC research.

## Methods

### Sample collection and tissue processing

Spontaneous CMT and non-neoplastic mammary tissue were obtained from excess tissues that were collected while client-owned dogs were undergoing standard of care surgical removal of mammary gland tumors in small animal clinics in Switzerland and at the Vetsuisse faculty (University of Bern, Switzerland) between 2018 and 2020. In accordance with relevant Swiss guidelines and regulations, an informed consent was obtained from the authorized welfare advocate of each participating dog to receive standard of care veterinary diagnostics and treatment and use of excess tissues for research purposes. All animals in this study were handled according to the ethical standards in Switzerland. We followed the ARRIVE guidelines (https://arriveguidelines.org) where relevant to our study. Ethical review and approval were waived by the Cantonal Veterinary Office for the investigation of the mammary tissues of the affected dogs in this study, as the sampling was performed during clinical and pathological veterinary diagnostics. The “Cantonal Committee for Animal Experiments” approved the collection of blood samples (Canton of Bern; permit 71/19). EDTA blood samples were stored at −80°C for future DNA isolation. For standard histopathological analysis, half of the CMT was formalin-fixed paraffin-embedded (FFPE). The rest was dissected into 1-2mm^3^ pieces. Randomly selected pieces were snap-frozen and stored at −80°C for future DNA isolation and the rest was routinely frozen in freezing medium (45% Dulbecco’s Modified Eagle Medium, 45% Fetal Calf Serum (ThermoFisher, Massachusetts, USA)) and 10% DMSO.

### Establishment and maintenance of CMT organoid cultures

Cryopreserved CMT tissues were thawed and processed to obtain a cell suspension following routine procedures^28^. The pellet was resuspended in cold CMT organoid medium (Supplementary Table S6) and mixed at a 1:1 ratio with Cultrex^®^ PathClear Reduced Growth Factors Basement Membrane Extract (BME) Type 2 (Amsbio, Abingdon, England). The BME-cell suspension was seeded as 30 μL drops on prewarmed 24-wells suspension culture plates (Greiner Bio-One, Kremsmünster, Austria) and cultured following standard protocols^28^.

### Histology, imaging, and immunohistochemistry

Organoids were washed and centrifuged at 2°C at 300 rcf for 5 min. The pellet was resuspended in formalin for 2 h, after which it was embedded in 2.5% low-melting agarose and followed by paraffin embedding. Hematoxylin and eosin (H&E)-stained FFPE sections of both primary tumors and organoids were analyzed by a board-certified veterinary pathologist (M.D.). Primary tumors were classified and graded according to the gold standard classification and grading system, as well as the Nielsen classification^30,8^. Immunohistochemistry was performed on FFPE sections using antibodies against different molecular markers (detailed in Supplementary Table S7). Slides were subsequently scanned on NanoZoomer S360 Digital slide scanner (C13220-01, Hamamatsu) and analyzed with QuPath software^52^. Scoring was performed following classical guidelines^32^.

### gDNA isolation, amplification, and Tracking of Indels by DEcomposition (TIDE) analysis

Organoids were trypsinized, and genomic DNA (gDNA) was extracted using the standard chloroform extraction protocol. Target loci were amplified following standard procedures^53^, and target modifications were confirmed using the TIDE algorithm^54^. Primers used in this PCR are mentioned in Supplementary Table S8.

### Whole Genome Sequencing (WGS)

gDNA extraction from tissue samples and blood was performed with QIAamp^®^ DNA Mini and Blood Mini kit (Qiagen). 1 mg per tumor sample and matched normal was used to generate DNA libraries for Illumina WGS using standard protocols. Individual lanelets (pairs of fastqs) were mapped to canine canFam3.1 genome, coordinate sorted merged, and duplicates-marked using dockerised cgpmap v3.0.4: https://github.com/cancerit/dockstore-cgpmap. Copy-number variants were called using ASCAT, which was executed as part of the dockerised cgpwgs v2.0.1: https://github.com/cancerit/dockstore-cgpwgs. Somatic singlenucleotide and indel variants were called using Strelka v2.9.10. Only variants flagged with SomaticEVS scores >=16 were carried forward for further analysis. Structural variants were called using Manta v1.6.0. Only variants where PR (paired-read coverage) >= 8 in the tumor were used in the analysis. To check if the mutations found in dogs were also present in HBC, we checked if commonly mutated genes (*i.e*., in more than 40% of the cases for either simple or complex carcinomas) were found in The Cancer Mutation Census accessed in July 2021 (https://cancer.sanger.ac.uk/cmc/home).

### Mutational signatures analysis

SBS were classified according to the type of mutation and trinucleotide context, *e.g*., C>T at ACA (mutated base underlined)^39^. A mutational catalog for each patient sample was constructed by counting the number of mutations in each of the 96 mutational classes. We then proceeded to identify the SBS that composed each of the samples’ mutational catalog. We first followed a signature fit approach, where an *a priori* set of SBS (identified in the Genomics England dataset^39^) is used, and the number of mutations associated with each signature (the exposures) is estimated (Supplementary Table S4). After noticing that all samples seemed to contain frequent mutations in the A<u>C</u>A and T<u>C</u>T contexts, as well as frequent T>A mutations in the T<u>T</u>A context (an unusual pattern for HBC), we attempted to isolate this pattern by using a combined fit and extraction algorithm that allowed for estimating the shape of one mutational signature alongside fitting our *a priori* set. The R package NNLM was used^55^.

### SNP array analysis

1 mg per sample was used to generate DNA libraries for Illumina CanineHD BeadChip array (Neogen, Nebraska, USA), containing over 220,000 highly polymorphic SNP. Data were quality-filtered and analyzed with PLINK v1.9^56^. All samples had call rates higher than 90%. Marker filtering was based on missing call rate (>10%) and Hardy-Weinberg equilibrium (P<10^-6^), resulting in the final dataset containing 209,878 markers and 31 samples, which was used for sample identification. PLINK v2.0 software was used for the detection of pairwise sample discordances between organoids and tissue samples^56^. Subsequently, a combination of two SNP array signal intensity measures, BAF and LRR, were plotted using R^55^.

### Drug testing and cell viability assays

Organoids were dissociated into single cells, and 35,000 cells were seeded per well in 10 μL BME/CMT medium drops on a 24-wells plate. Organoids were cultured for eight days in different concentrations of carboplatin, cisplatin, doxorubicin (Teva Pharma AG, Basel, Switzerland), alpelisib, and nutlin-3a (Selleckchem). The growth medium was refreshed and replaced with growth medium after 48 h (carboplatin, cisplatin, doxorubicin) or replaced with drug-refreshed medium after 96 h (alpelisib, nutlin-3a). After eight days, cell viability was assessed using the resazurin-based Cell Titer Blue following the manufacturer’s instructions (Promega). In brief, 25 μL of the reagent was added to the culture medium and incubated for 4 h at 37°C. 200 μL of the medium was then pipetted to a 96-wells plate, and fluorescence intensity at 560Ex/590Em nm was determined with an Enspire Multimode Plate Reader (PerkinElmer, Waltham, USA). Three individual biological replicates were at least performed. Results were normalized to the untreated control. Inhibition cell growth was fitted to four-parameter logistic sigmoidal curve and led to the determination of IC50 values. As a control, the HBC estrogen-positive, *PIK3CA*-mutated, *TP53* wild-type MCF-7 cell line (provided by Andrea Morandi, Tumor Biochemistry Lab, Departement of Experimental and Clinical Biomedical Sciences, Italy) was used and grown to form organoids (named ORG-MCF7) following the same conditions as the CMT organoids.

### Lentivirus production, lentiviral transduction

Lentiviral stocks were generated following standard procedures^53^. Lentiviruses were produced by transient co-transfection of lentiviral packaging plasmids and the plentiCRISPRv2 vector containing the respective gRNA or a non-targeting gRNA (Supplementary Table S9). Organoids were transduced following a previously established spinoculation protocol^57^.

### Fluorescence-Activated Cell Sorting (FACS)

Transduction efficiency was determined by flow cytometry three days after transduction with a GFP-encoding lentivirus. After trypsinisation, at least 100,000 cells per condition were resuspended in phosphate-buffered saline and sorted on a BD Biosciences FACSCanto™ II Clinical Flow Cytometry System. The analysis was performed with the Flow Jo software, where the forward versus side scatter gating was used to identify cells of interest. Doublets were excluded using the forward scatter height versus forward scatter area density plot. Quantification of GFP-positive populations led to the determination of the transduction efficiency. Two independent experiments were conducted on three different organoid lines.

### CRISPR/Cas9 screening

Based on the canine CanFam3.1 assembly, we established a lentivirus-based CRISPR/Cas9 sublibrary targeting genes known to be druggable, containing 6004 gRNAs, targeting 834 genes (six gRNAs/gene) in addition to 500 non-targeting and 500 intergenic controls (Supplementary Table S5). Sixteen million ORG-63-C and ORG-63-N were collected and transduced with lentivirus at a multiplicity of infection (number of viral particles per cell) of four. The medium was replaced with puromycin-containing medium (3,5 μg/ml, GIBCO) 24 h later, and selection was performed for eleven days. Twelve days after transduction (day 0 = D0), organoids were trypsinized, and gDNA was isolated from 6 million cells. Two biological replicates were performed. For PCR amplification, gDNA was divided into 100□μL reactions such that each well had at most 10□μg of gDNA. Per 96-wells plate, a master mix consisted of 150□μL DNA Polymerase (Titanium Taq; Takara), 1 mL of 10x buffer, 800□μL of dNTPs (Takara), 50□μL of P5 stagger primer mix (100□μM), and water to bring the final volume to 4 mL. Each well consisted of 50□μL gDNA plus water, 40□μL PCR master mix, and 10□μL of a uniquely barcoded P7 primer (5□μM). PCR cycling conditions were as follows: (1) 95°C for 1 min; (2) 94°C for 30 s; (3) 52.5°C for 30 s; (4) 72°C for 30 s; (5) go to (2), x 27; (6) 72°C for 10 min. Primers used in this PCR are mentioned in Supplementary Table S8. PCR products were purified with Agencourt AMPure XP SPRI beads according to the manufacturer’s instructions (Beckman Coulter, A63880). Samples were sequenced on a HiSeq2500 HighOutput (Illumina) with a 5% spike-in of PhiX. MAGeCK (Model-based Analysis of Genome-wide CRISPR-Cas9 Knockout) algorithm was used for enrichment analysis^58^.

### Statistical analysis and data representation

Prism statistical software (v9.0; GraphPad Inc, San Diego, USA) was used for statistical analyses and data representation. Statistical tests and P-values are indicated in the text or the figures’ legends.

## Acknowledgements

We thank all owners who consented to participate in the study and all clinicians and nurses involved in the samples’ acquisition. We are grateful to Eve Tièche and Merve Mutlu for their experimental help. We want to thank Sara Soto for the help with the antibodies establishment and Pathologie Länggasse laboratory for technical assistance. We wish to thank Kerry Woods, Carmen Widmer, and Diego Dibittetto for their critical reading of the manuscript and Cédric Walker for his bioinformatical support. We thank the Interfaculty Bioinformatics Unit of the University of Bern for providing high-performance computing infrastructure. Finally, we are grateful to the pathology residents’ and technician’s team for the samples and sections’ processing. Parts of some figures were created with BioRender.com.

## Author contributions statement

S.R. supervised the study. M.I. and S.R. planned and designed the study. M.I. and K.H. collected samples and clinical data. M.I. carried out experiments and analyzed data. M.D. performed the pathological assessment. Y.M., H.D., and S.NZ. analyzed the WGS data, and A.D. performed the mutational signatures analysis. A.L. and C.D. analyzed the SNP data. J.D. and A.B. designed the canine CRISPR sublibrary. S.R. acquired financial support for the project. M.I. wrote the original draft manuscript. S.R., M.D., A.L., J.D., A.D. critically reviewed and edited the manuscript, and all authors approved its final version.

## Data availability statement

The whole-genome sequencing data generated in this study are available in the Sequence Read Archive (SRA) under accession number PRJNA839753 (https://www.ncbi.nlm.nih.gov/bioproject/PRJNA839753/). Raw data for the SNP array analysis can be found under DOI 10.17605/OSF.IO/N85Q9.

