## Supplementary information for "A living biobank of canine mammary tumor organoids as a comparative model for human breast cancer"

**Figure S1.** Comparison between unknown dog signature and SBS57 PCR-related sequencing artefact

**Figure S2.** Conservation of the genetic landscape of CMT in patient-derived ORG

**Figure S3.** CMT organoids allow *in vitr*o drug testing

**Table S1.** Clinical information of all patients

**Table S2.** Clinical information and characterization of patients presenting malignant tumors

**Table S3.** Detailed characterization of tissue/organoid pairs

**Table S4.** Estimation of the number of mutations associated with each Single Base Substitution Signatures (SBS) following signature fit approach and comparison of prevalence of SBS between CMT and HBC

**Table S5.** Genes targeted by the custom canine CRISPR library and gRNAs counts (plasmid DNA and day 0)

**Table S6.** Canine mammary tumor organoid medium components

**Table S7.** Primary antibodies used for immunohistochemistry

**Table S8.** Primers sequences used for Sanger sequencing and TIDE analysis

**Table S9.** Primers sequences used for gene editing

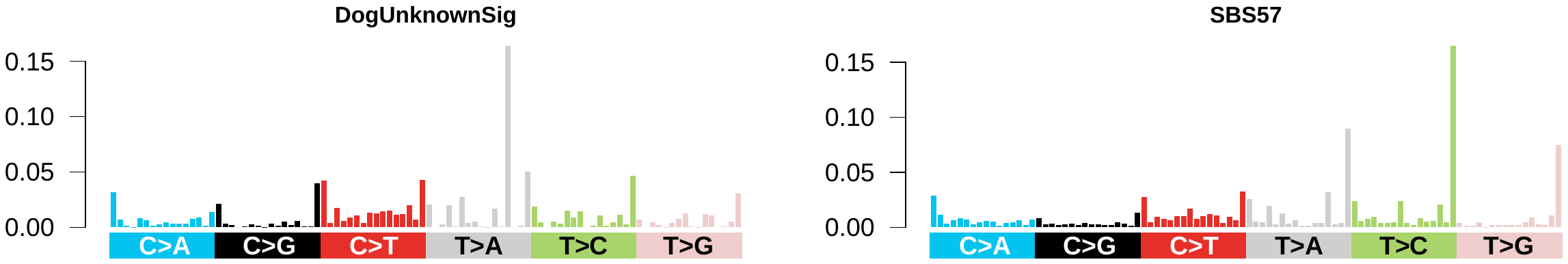

**Figure S1. Comparison between unknown dog signature and SBS57 PCR-related sequencing artefact**

The main difference between the two signatures seems to be the much higher frequency of T>A mutations at the TTA context in the dog signature.

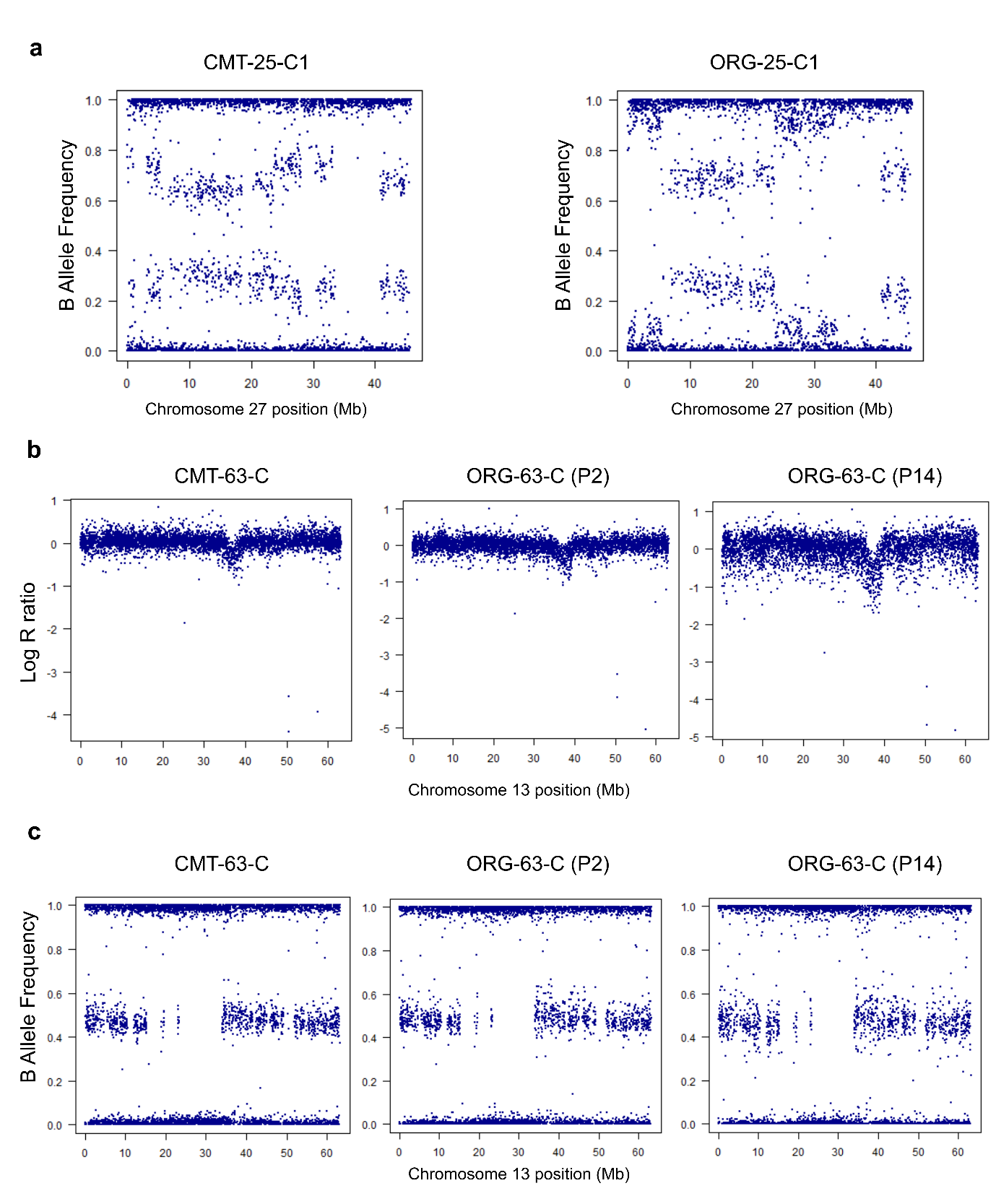

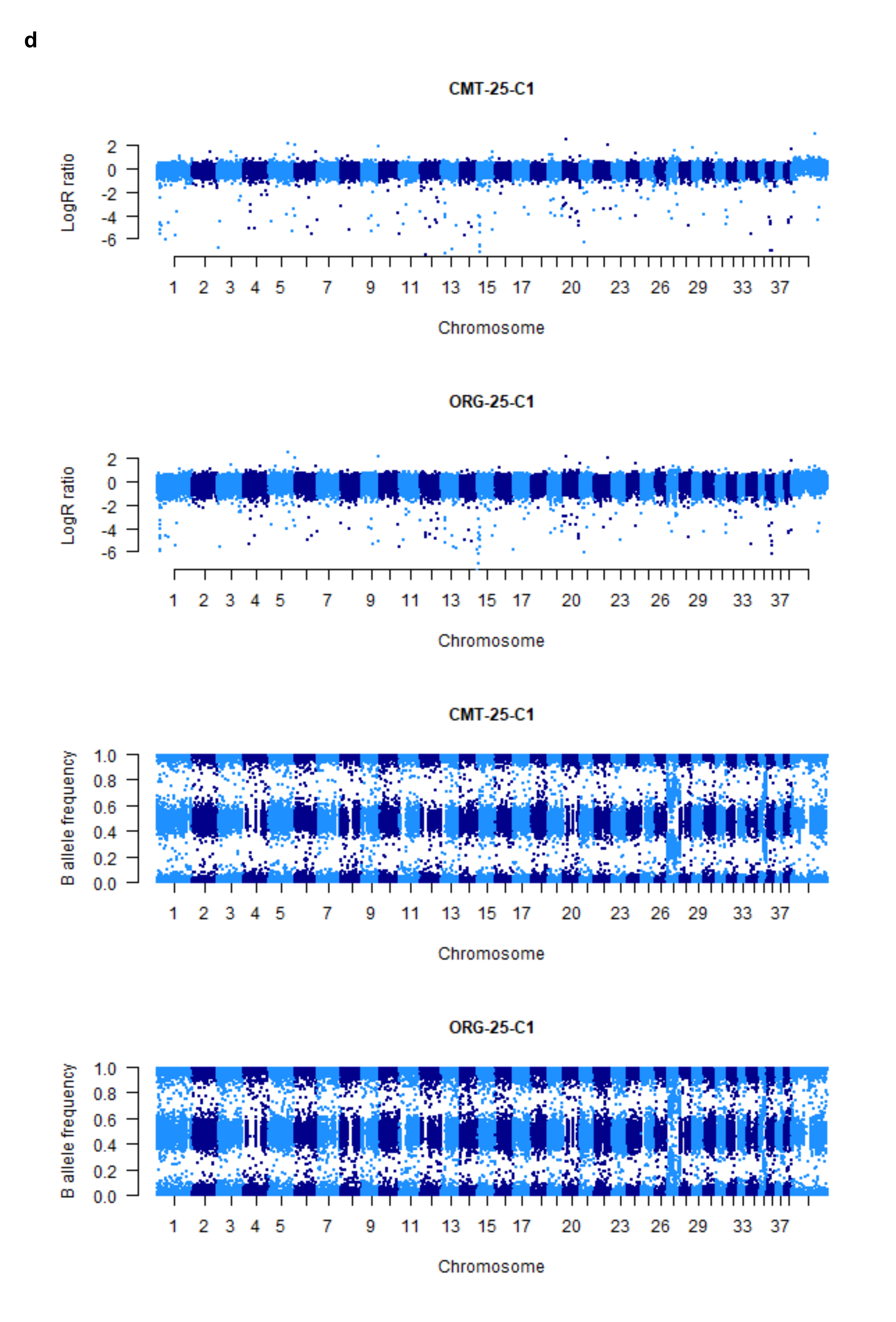

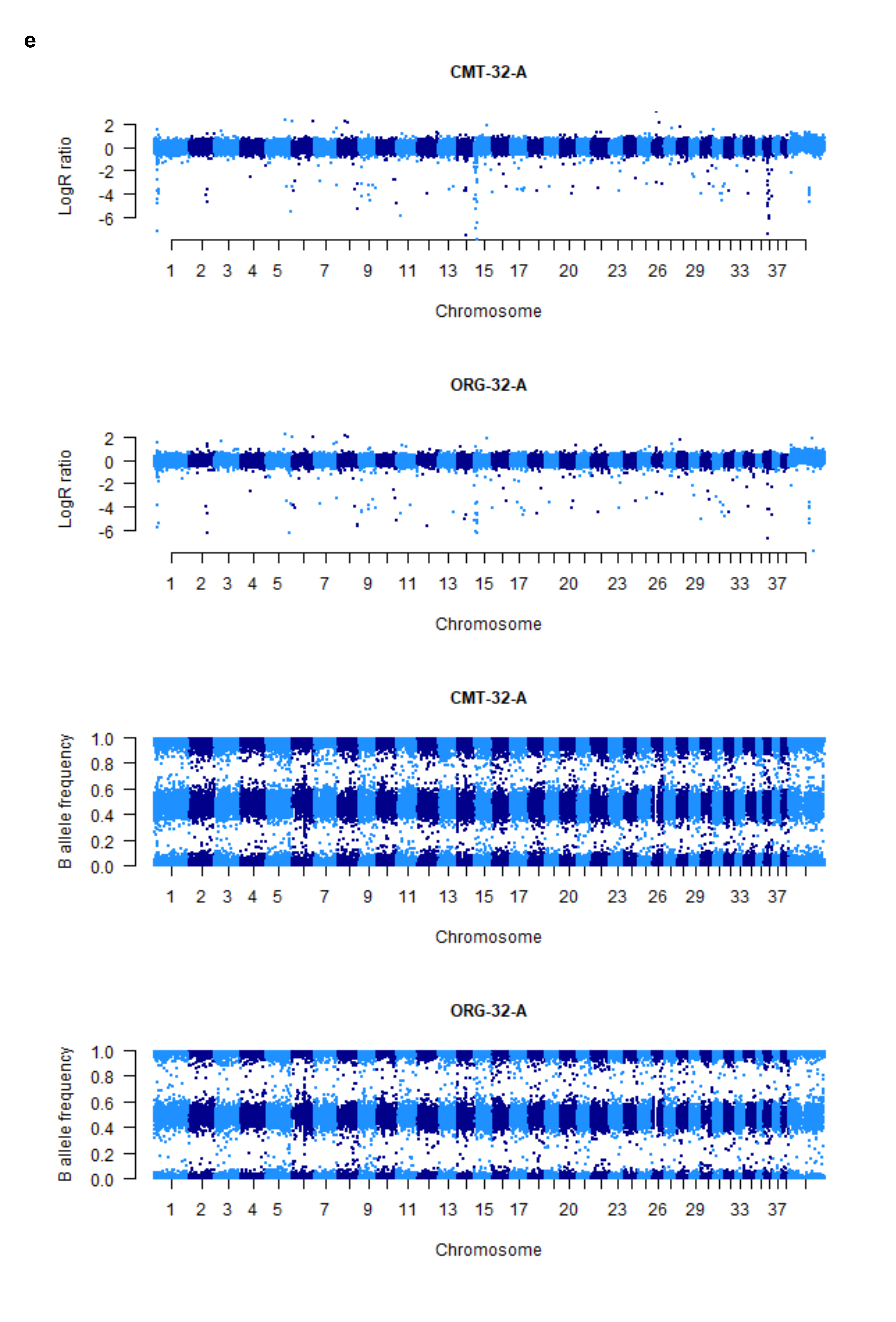

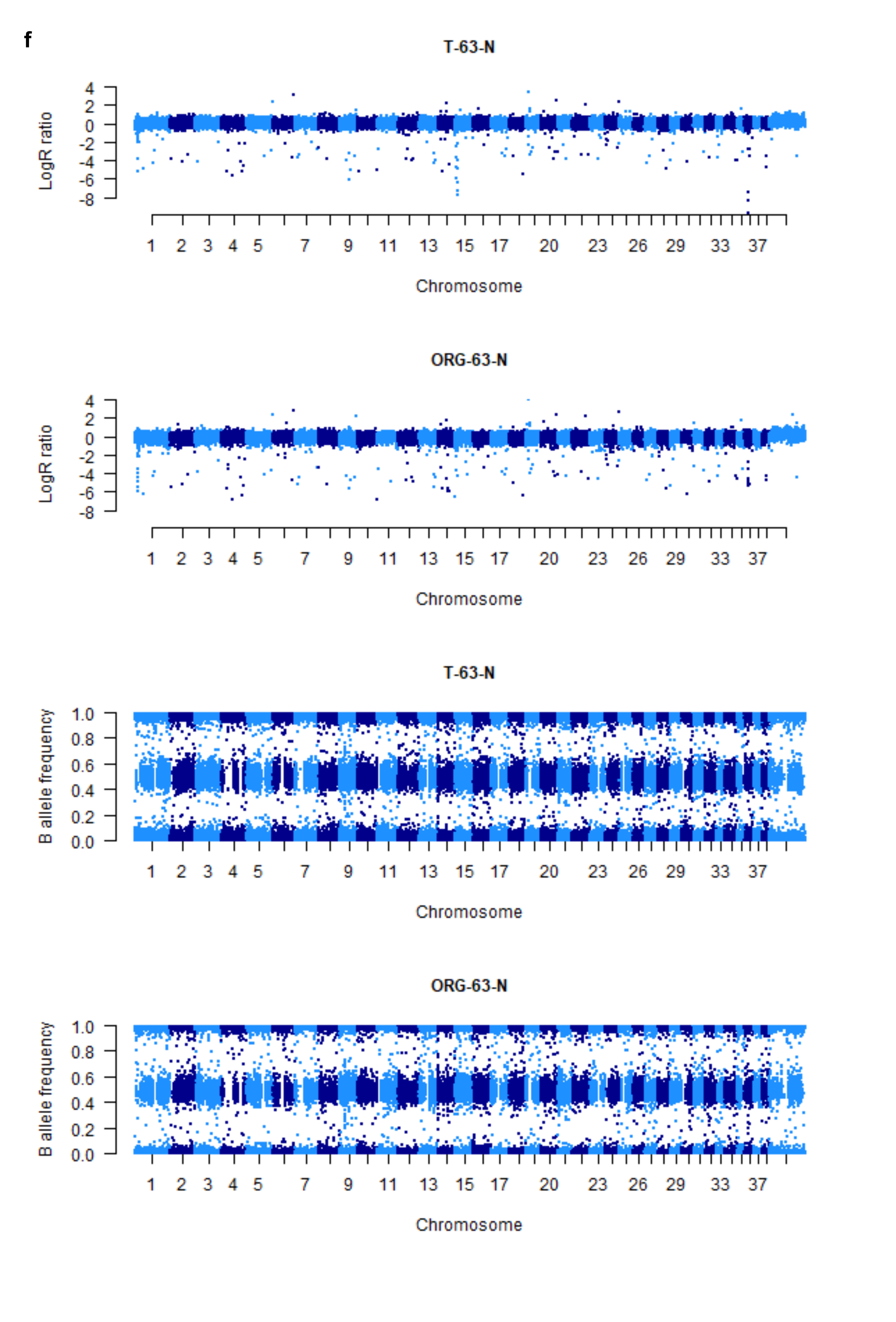

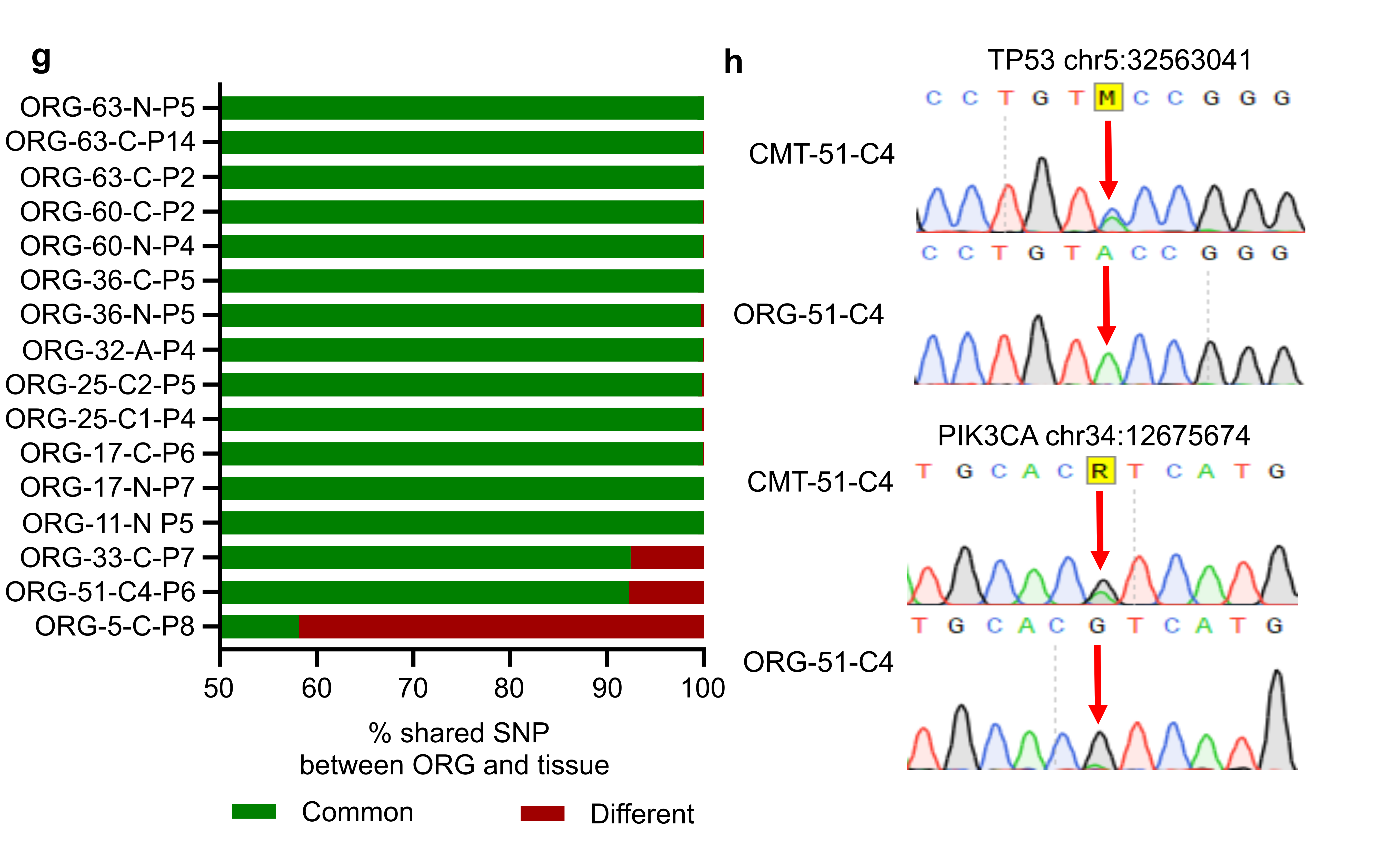

**Figure S2. Conservation of the genetic landscape of CMT in patient-derived ORG**

a. B-allele frequencies on chromosome 27 for matched pair CMT-25-C1 / ORG-25-C1. See also Figure 3B.
b. Log R ratios on chromosome 13 for matched pair CMT-63-C / ORG-63-C (P2) and later passage (P) ORG-63-C (P14). Note the conservation of the SNP distribution in the organoid line after extended time in culture.
c. B-allele frequencies on chromosome 13 for matched pair CMT-63-C / ORG-63-C (P2) and later passage ORG-63-C (P14). Note the conservation of the SNP distribution in the organoid line after extended time in culture.
d. Genome-wide SNP distribution of log R ratios on top part, B-allele frequencies on the bottom part for carcinoma-organoid pair CMT-25-C1 / ORG-25-C1.
e. Genome-wide SNP distribution of log R ratios on top part, B-allele frequencies on the bottom part for adenoma-organoid pair CMT-32-A / ORG-32-A.
f. Genome-wide SNP distribution of log R ratios on top part, B-allele frequencies on the bottom part for non-neoplastic mammary tissue-organoid pair T-63-N / ORG-63-N.
g. Proportion of shared SNP genotypes between primary tissues and matched organoids.
h. Chromatograms of point mutations for matched pair CMT-51-C4 / ORG-51-C4. Top panel: *TP53* chr5:32563041, bottom panel: *PIK3CA* chr34:12675674

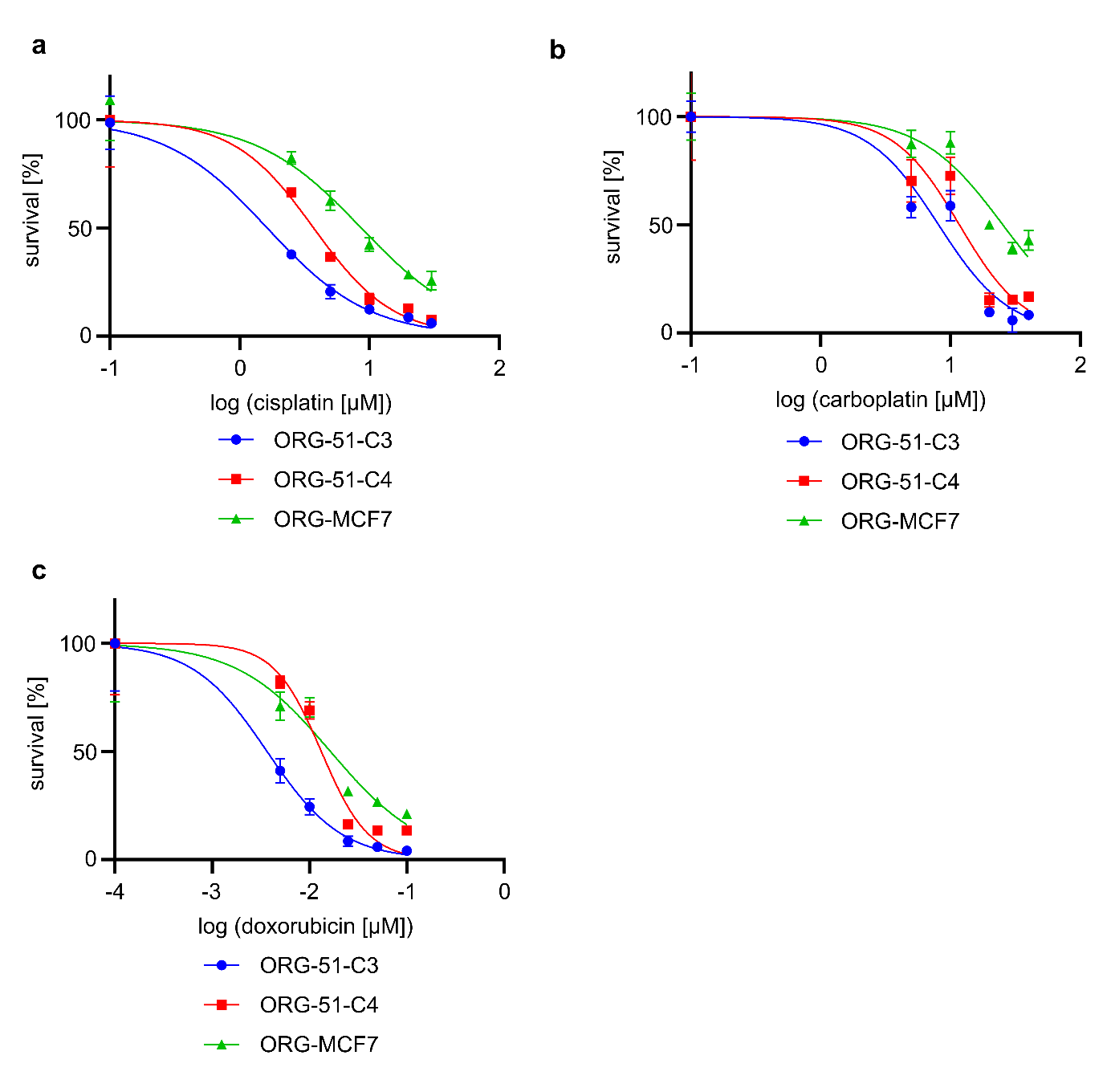

**Figure S3. CMT organoids allow *in vitr*o drug testing**

a. Dose response curves indicating viability 8 days after treatment with cisplatin. Error bars represent SD of three independent experiments.
b. Dose response curves indicating viability 8 days after treatment with carboplatin. Error bars represent SD of three independent experiments.
c. Dose response curves indicating viability 8 days after treatment with doxorubicin. Error bars represent SD of three independent experiments.

**Table S1.** Clinical information of all patients

| Patient number | Breed | Owner consent signed: yes(y)/ no(n) | Age (yr), ?=unknown | Sex: female (f) | Neuter status | Histopathological diagnosis |
| --- | --- | --- | --- | --- | --- | --- |
| 1 | King Charles Spaniel | y | 15 | f | Spayed | Carcinoma - tubular |
| 2 | Yorkshire Terrier | y | 9 | f | Intact | Ductal carcinoma |
|  |  |  |  |  |  | Adenoma - complex |
|  |  |  |  |  |  | Mixed tumor |
| 3 | French Bulldog | y | 11 | f | Spayed | Ductal carcinoma |
| 4 | Tibet-Terrier | y | 11 | f | Intact | Mixed tumor |
|  |  |  |  |  |  | Adenoma - complex |
|  |  |  |  |  |  | Simple adenoma |
|  |  |  |  |  |  | Ductal adenoma |
| 5 | Italian wind chime | y | 13 | f | Intact | Carcinoma - anaplastic |
|  |  |  |  |  |  | Tubular adenoma |
|  |  |  |  |  |  | Mixed tumor |
| 6 | Mixed breed | y | 12 | f | Intact | Carcinoma arising in a mixed tumor |
|  |  |  |  |  |  | Tubular adenoma |
|  |  |  |  |  |  | Mixed tumor |
| 8 | Yorkshire Terrier | y | 15 | f | Spayed | Comedocarcinoma |
| 9 | English Cocker Spaniel | y | 7 | f | Intact | Tubular adenoma |
|  |  |  |  |  |  | Adenoma - complex |
| 10 | Jack Russell Terrier | y | 9 | f | Intact | Carcinoma arising in a complex adenoma |
| 11 | Yorkshire Terrier | y | 11.5 | f | Intact | In situ carcinoma arising in a simple tubular adenoma |
|  |  |  |  |  |  | Tubular adenoma |
| 13 | Sheltie | y | 6.7 | f | Spayed | Carcinoma - complex |
| 14 | Golden retriever | y | 10.2 | f | Intact | Carcinoma - complex |
| 15 | Golden retriever | y | 10.5 | f | Intact | Intraductal papillary carcinoma |
|  |  |  |  |  |  | Carcinoma - complex |
| 16 | Dalmatian | y | 8.5 | f | Intact | Carcinoma - tubulopapillary |
| 17 | Irish setter | y | 9.8 | f | Intact | Carcinoma and malign myoepithelioma |
| 18 | Parson russelTerrier | y | 11 | f | Intact | Intraductal papillary carcinoma |
| 19 | English Springer Spaniel | y | 7 | f | Intact | Mixed tumor |
| 21 | Havaneser | y | 8.8 | f | Spayed | Mixed tumor |
| 23 | Labrador | y | 11 | f | Spayed | Intraductal papillary carcinoma and ductal carcinoma |
| 24 | Dackel | y | 12 | f | Intact | Ductal carcinoma - solid |
|  |  |  |  |  |  | Adenoma - complex |
| 25 | Cocker Spaniel | y | 13 | f | Intact | Carcinoma - anaplastic |
|  |  |  |  |  |  | Comedocarcinoma |
| 26 | Jack Russell Terrier | y | ? | f | Spayed | Tubular adenoma |
|  |  |  |  |  |  | Adenoma - complex |
| 28 | Chihuahua | y | ? | f | Intact | Mixed tumor |
| 29 | German Sheperd mix | y | 12 | f | Spayed | Mixed tumor |
| 31 | Dackel | y | 11 | f | Intact | Intraductal adenoma |
| 32 | French Bulldog | y | 12 | f | Intact | Carcinoma - micropapillar |
|  |  |  |  |  |  | Carcinoma - tubular |
|  |  |  |  |  |  | Simple adenoma |
| 33 | Labrador | y | 13 | f | Intact | Intraductal tubulopapillary carcinoma |
| 36 | Doberman | y | 7 | f | Intact | Metastasis from a carcinoma |
| 37 | Jack Russell Terrier | y | ? | f | Intact | Intraductal adenoma |
| 38 | Yorkshire Terrier | y | 12 | f | Intact | Mixed tumor |
| 41 | Icelandic Shepherd Dog | y | 9.5 | f | Intact | Carcinoma and malignant myoepithelioma |
| 43 | Beauceron | y | 13 | f | Spayed | Carcinoma - complex |
|  |  |  |  |  |  | Mixed tumor |
| 44 | Border collie | y | 10 | f | Intact | Tubular adenoma |
| 45 | Mixed breed | y | 9 | f | Intact | Adenoma - complex |
| 46 | Gordon Setter | y | 9 | f | Intact | Mixed tumor |
| 47 | Puddle | y | 11 | f | Intact | Carcinoma - complex |
|  |  |  |  |  |  | Adenoma - complex |
| 48 | Puddle | y | 13 | f | Spayed | Carcinoma - complex |
|  |  |  |  |  |  | Tubular adenoma |
| 49 | Briard | y | 9.7 | f | Intact | Ductal adenoma |
| 50 | American Staffordshire Terrier | y | 5.5 | f | Intact | Mixed tumor |
| 51 | Appenzell Mountain dog | y | 14 | f | Intact | Carcinoma - complex |
|  |  |  |  |  |  | Carcinoma - solid |
|  |  |  |  |  |  | Mixed tumor |
| 54 | Chihuahua | y | 10 | f | Spayed | Simple adenoma |
| 55 | Hovawart | y | 7 | f | Intact | Carcinoma - tubulopapillar |
| 56 | Nova Scotia Duck Tolling Retriever | y | 4 | f | Intact | Adenoma - complex |
| 57 | Mixed breed | y | 6 | f | Spayed | Carcinoma - micropapillar |
|  |  |  |  |  |  | Carcinoma - complex |
|  |  |  |  |  |  | Carcinoma - complex |
|  |  |  |  |  |  | Mixed tumor |
| 59 | Border collie | y | 10 | f | Intact | Carcinoma - complex |
| 60 | German Sheperd | y | 9 | f | Intact | Carcinoma - complex |
| 61 | Irish setter | y | 9 | f | Intact | Comedocarcinoma |
| 62 | English Springer Spaniel | y | ? | f | Intact | Carcinoma - complex |
|  |  |  |  |  |  | Mixed tumor |
| 63 | Cocker Spaniel | y | 9 | f | Intact | Intraductal papillar carcinoma |
|  |  |  |  |  |  | Complex carcinoma |
|  |  |  |  |  |  | Complex carcinoma |
|  |  |  |  |  |  | Mixed tumor |

**Table S2.** Clinical information and characterization of patients presenting malignant tumors

| Patient number | Sample Name | Breed | Age (yr) | Sex: female (f) | Neuter status | Histopathological diagnosis | Grade | TNM staging | Lymphatic vessel invasion | Adenoma (A) or mixed tumors (M) affect other complexes | ER | PR | HER2 | Nielsen classification | Organoids established success(s) / failure (f) / not tried (nt) |
| --- | --- | --- | --- | --- | --- | --- | --- | --- | --- | --- | --- | --- | --- | --- | --- |
| 1 | CMT-01-C | King Charles Spaniel | 15 | f | Spayed | Carcinoma - tubular | III | 3 | No | No | - | - | 1+ | Triple negative basal like | s |
| 2 | CMT-02-C | Yorkshire Terrier | 9 | f | Intact | Ductal carcinoma | I | 2 | No | A + M | - | - | - | Triple negative basal like | s |
| 3 | CMT-03-C | French Bulldog | 11 | f | Spayed | Ductal carcinoma | II | 1 | No | No | - | - | - | Triple negative basal like | s |
| 5 | CMT-05-C | Italian wind chime | 13 | f | Intact | Carcinoma - anaplastic | III | 4 | Yes | A + M | + | - | - | Luminal A | s |
| 6 | CMT-06-C | Mixed breed | 12 | f | Intact | Carcinoma arising in a mixed tumor | I | 1 | No | A + M | + | + | 1+ | Luminal A | s |
| 8 | CMT-08-C | Yorkshire Terrier | 15 | f | Spayed | Comedocarcinoma | III | 4 | Yes | No | - | - | 2+ | Triple negative basal like | s |
| 10 | CMT-10-C | Jack Russell Terrier | 9 | f | Intact | Carcinoma arising in a complex adenoma | I | 2 | No | No |  |  |  |  | nt |
| 11 | CMT-11-C | Yorkshire Terrier | 11,5 | f | Intact | In situ carcinoma arising in a simple tubular adenoma | I | 1 | No | A | + | + | 1+ | Luminal A | f |
| 13 | CMT-13-C | Sheltie | 6,7 | f | Spayed | Carcinoma - complex | II | 1 | No | No | + | + | 1+ | Luminal A | f |
| 14 | CMT-14-C | Golden retriever | 10,2 | f | Intact | Carcinoma - complex | II | 1 | No | No | + | + | 1+ | Luminal A | nt |
| 15 | CMT-15-C1 | Golden retriever | 10,5 | f | Intact | Intraductal papillary carcinoma | I | 2 | No | No | + | + | 1+ | Luminal A | nt |
|  | CMT-15-C2 |  |  |  |  | Carcinoma - complex | I |  |  |  | + | + | 1+ | Luminal A | s |
| 16 | CMT-16-C | Dalmatian | 8.5 | f | Intact | Carcinoma - tubulopapillary | I | 3 | No | No | + | + | 2+ | Luminal A | s |
| 17 | CMT-17-C | Irish setter | 9.8 | f | Intact | Carcinoma and malign myoepithelioma | II | 3 | No | No | + | - | 1+ | Luminal A | s |
| 18 | CMT-18-C | Parson russelTerrier | 11 | f | Intact | Intraductal papillary carcinoma | II | 2 | No | No | - | - | 1+ | Triple negative basal like | nt |
| 23 | CMT-23-C | Labrador | 11 | f | Spayed | Intraductal papillary carcinoma and ductal carcinoma | II | 1 | No | No | + | - | 2+ | Luminal A | f |
| 24 | CMT-24-C | Dackel | 12 | f | Intact | Ductal carcinoma - solid | III | 2 | No | A | + | - | 1+ | Luminal A | nt |
| 25 | CMT-25-C1 | Cocker Spaniel | 13 | f | Intact | Carcinoma - anaplastic | II | 2 | No | No | + | - | 0 | Luminal A | s |
|  | CMT-25-C2 |  |  |  |  | Comedocarcinoma | II | 4 | Yes |  | + | + | 1+ | Luminal A | s |
| 32 | CMT-32-C1 | French Bulldog | 12 | f | Intact | Carcinoma - micropapillar | I | 2 | No | A | + | - | 1+ | Luminal A | f |
|  | CMT-32-C2 |  |  |  |  | Carcinoma - tubular | I | 2 |  |  | + | + | 1+ | Luminal A | f |
| 33 | CMT-33-C | Labrador | 13 | f | Intact | Intraductal tubulopapillary carcinoma | II | 1 | No | No | + | + | 1+ | Luminal A | s |
| 36 | CMT-36-C | Doberman | 7 | f | Intact | Metastasis from a carcinoma |  | 4 | Yes | No | + | - | 1+ | Luminal A | s |
| 41 | CMT-41-C | Icelandic Shepherd Dog | 9,5 | f | Intact | Carcinoma and malignant myoepithelioma | I | 3 | No | No | + | + | 1+ | Luminal A | nt |
| 43 | CMT-43-C | Beauceron | 13 | f | Spayed | Carcinoma - complex | I | 2 | No | M |  |  |  |  | nt |
| 47 | CMT-47-C | Puddle | 11 | f | Intact | Carcinoma - complex | I | 1 | No | A |  |  |  |  | nt |
| 48 | CMT-48-C | Puddle | 13 | f | Spayed | Carcinoma - complex | I | 1 | No | A |  |  |  |  | nt |
| 51 | CMT-51-C3 | Appenzeller Sennenhund | 14 | f | Intact | Carcinoma - complex | II | 4 | Yes | M | + | + | 1+ | Luminal A | s |
|  | CMT-51-C4 |  |  |  |  | Carcinoma - solid | II |  |  |  | - | - | 0 | Triple negative basal like | s |
| 55 | CMT-55-C | Hovawart | 7 | f | Intact | Carcinoma - tubulopapillar | I | 1 | No | No |  |  |  |  | nt |
| 57 | CMT-57-C1 | Mixed breed | 6 | f | Spayed | Carcinoma - micropapillar | II | 1 | No | M | - | - | 1+ | Triple negative basal like | nt |
|  | CMT-57-C2 |  |  |  |  | Carcinoma - complex | I | 1 |  |  | - | - | 1+ | Triple negative basal like | nt |
|  | CMT-57-C3 |  |  |  |  | Carcinoma - complex | I | 1 |  |  | + | - | 1+ | Luminal A | nt |
| 59 | CMT-59-C | Border collie | 10 | f | Intact | Carcinoma - complex | I | 1 | No | No | + | + | 1+ | Luminal A | nt |
| 60 | CMT-60-C | German Sheperd | 9 | f | Intact | Carcinoma - complex | I | 2 | No | No | + | + | 2+ | Luminal A | s |
| 61 | CMT-61-C | Irish setter | 9 | f | Intact | Comedocarcinoma | III | 4 | Yes | No | + | - | 1+ | Luminal A | nt |
| 62 | CMT-62-C | English Springer Spaniel | ? | f | Intact | Carcinoma - complex | I | 2 | No | M | + | + | 1+ | Luminal A | nt |
| 63 | CMT-63-C1 | Cocker Spaniel | 9 | f | Intact | Intraductal papillar carcinoma | II | 4 | Yes | M |  |  |  |  | nt |
|  | CMT-63-C2 |  |  |  |  | Complex carcinoma | I |  | No |  |  |  |  |  | nt |
|  | CMT-63-C |  |  |  |  | Complex carcinoma | I |  | No |  | + | + | 1+ | Luminal A | s |

**Table S3.** Detailed characterization of tissue/organoid pairs

|  | Tissues | | | | | | | | | | | | | | | |
| --- | --- | --- | --- | --- | --- | --- | --- | --- | --- | --- | --- | --- | --- | --- | --- | --- |
| Patient number | Sample Name | Normal / Adenoma / Carcinoma | Histological grade (for carcinomas only) | HE | Vimentin | CK7/8 = Cam5.2 | CK5/6 | CK14 | p63 | Ki67 (% positive cells) | ER (% positive cells) | ER (Allred) | PR (% positive cells) | PR (Allred score) | HER2 (Pena) | Nielsen classification (for tumors only, not normal tissue) |
| 1 | CMT-01-C | C | II | carcinoma and malignant myoepithelioma | around 10% display week signal, mainly in tumor periphery | 20-25%, forming tubuli/acini embedded in negative cells. | >98% (tubuli and surrounding cells) | 5-10% in tumor periphery and near necrotic center (not tubuli) | 75-80%, not tubuli | 21.40% | 0.00% | 0 | 0 | 0 | 1+ | Triple negative basal like |
| 2 | CMT-02-C | C | II | intraductal papillary carcinoma | 5-10% (basal cells only) | 80-90% (not basal cells) | 25-30% (basal cells only) | 25-30% (basal cells only) | around 10% (basal cells only) | 39% | <1% | 2 | 0 | 0 | 0 | Triple negative basal like |
| 3 | CMT-03-C | C | II | carcinoma and malignant myoepithelioma | 40-50% basal cells or fibroblasts | 5-10% forming small tubuli embedded in negative cells. | >98% positive | >98% positive | around 90% positive (small tubuli negative) | 21.80% | <1% | 2 | 0 | 0 | 1+ | Triple negative basal like |
| 5 | CMT-05-C | C | I | carcinoma arising from a complex adenoma | 75% of all cells, mainly solid areas with spindeloid cells, and basal cells | around 25% of all cells, tubular and luminal cells; spindeloid cells negative. | around 25-30% of all cells, tubular and luminal cells. | around 25-30% of all cells, tubular and luminal cells. | around 50% of all cells, parts of spindeloid cells and basal layer of tubuli. | 15.20% | Epithelial cells: 10-20% strong. Spindeloid cells 90% strong. | Epithelial cells: 5 ; Spindeloid cells: 8 | 0 | 0 | 0 | Luminal A |
|  | CMT-05-A | A |  | complex adenoma | around 60% of all cells: basal cells and fibroblasts, and solid islands of spindeloid cells. | around 40% of all cells: tubular und luminal cells. Spindeloid cells negative. | around 50% of all cells: basal and 30% of luminal cells positive, spindeloid and 70% of luminal cells negative. | around 30-40% of all cells: basal cells positive, luminal and spindeloid cells negative. | around 25-30% of all cells: basal cells positive. Luminal and spindeloid cells negative. | 13.10% | Epithelial cells: 33-66% strong, Spindeloid cells 90% strong | Epithelial cells: 6 ; Spindeloid cells: 8 | Epithelial cells: 50% strong, Spindeloid cells <1% moderate | Epithelial cells: 7, Spindeloid cells 3 | 1+ | Luminal A |
| 6 | CMT-06-C | C | I | carcinoma and malignant myoepithelioma | Interstitial fibroblasts, around 40% of tumor cells (basal/vacuolated cells). | 60-70% of all cells (luminal) | 10-20% of all cells, mainly basal cells, but very patchy/intermingled. | around of all cells 20% (basal cells, patchy distribution) | 40-50% of all cells: single row of basal cells, but also more solid areas completely positive. | 6.60% | 69.70% | 8 | 42% | 7 | 1+ | Luminal A |
|  | T-6-N | N |  | lobular hyperplasia | Interstital fibroblasts and rare basal cells positive | Acini and ducts with one layer of luminal cells positive. | Acini: one layer of basal cells positive. Ducts with luminal and basal cells positive. | Acini: one layer of basal cells positive. Ducts with luminal and basal cells positive. | Acini and ducts with one layer of luminal cells positive. | 1.10% | 53.80% | 7 | 21.90% | 5 | 3+ |  |
| 8 | CMT-08-C | C | III | comedocarcinoma | 75% of neoplastic cells | 80% of neoplastic cells | 50% (patchy distribution, even within microproliferations) | 1% (individual cells in neoplastic proliferations) | 1% (individual cells in neoplastic proliferations) | 38.40% | 0% | 0 | 0% | 0 | 2+ | Triple negative basal like |
| 11 | T-11-N | N |  | normal to hyperplastic mammary tissue | 40% (basal cells) |  | 40-50% (basal cells of acini), 100% in collecting ducts | 30-40% (basal cells of acini), 50% in collecting ducts (basal cells mainly) | 10-20% in acini, around 40% in ducts | 15% | 81% | 8 | 7.80% | 4 | 2+ |  |
| 16 | CMT-16-C | C | I | simple tubular carcinoma | around 10% of tubular cells positive. Stromal fibroblasts positive. | All tubular cells positive. Few basal cells of proliferations in larger lumina negative. | Almost 100% of neoplastic cells positive. | >95% of neoplastic cells positive. Negative cells among larger lumina lining cells. | around 10% of neoplastic cells positive. Larger lumina lined by one layer of positive. cells, luminal layer negative. | 5.40% | 3.90% | 4 | 15% | 5 | 2+ | Luminal A |
| 17 | CMT-17-C | C | II | carcinoma and malignant myoepithelioma | Spindeloid cells and basal cells of tubulo-acinar structures positive. Luminal cells negative. | Tubulo-acinar cells positive. Spindeloid cells negative. (around 10% of all neoplastic cells positive). | Tubulo-acinar cells both layers positive. Ducts only basal layer positive. Spindeloid cells negative. (around 15-20% of all neoplastic cells positive) | About 50% of tubulo-acinar structures with only basal layer positive, other half with both layers positive. Ducts only basal layer positive. Spindeloid cells negative. (around 10-15% of all neoplastic cells positive) | Basal cells of tubulo-acinar structures and ducts and about 20% of spindeloid cells. | 12.25% | 32.50% | 5 | 0.40% | 3 | 1+ | Luminal A |
|  | T-17-N | N |  | lobular hyperplasia with secretory activity | Basal layer of acini and ducts positive. | Luminal cells of ducts positive, luminal cells of acinar structures only faintly positive. | Basal cells of acini and ducts,and luminal duct epithelium positive. | Basal cells of acini and ducts in normal tissue, but only around 20% of basal cells in hyperplastic nodules | Basal cells of acini (only around 20%) and ducts. | 37.90% | 82.90% | 8 | 30.30% | 6 | 2+ |  |
| 25 | CMT-25-C1 | C | II | ductal carcinoma | islands and cords with 1-3 peripheral positive layers, interspersed individual positive cells in center of islands. | Luminal cells moderately positive. around 30-40% of all cells. | around 80-90% strongly positive. (peripheral layer plus luminal cells). Some cells in between weakly to moderately positive. | around 60% of cells positive (peripheral always positive, intermediate and luminal cells variably positive) | 1-5 peripheral layers of positive cells, around 30-40% of all cells positive. Larger islands also with interspersed positive cells. | 15.45% | 1.50% | 3 | 0.57% | 3 | 0 | Luminal A |
|  | CMT-25-C2 | C | II | complex carcinoma | Around 98% of spindeloid cells are positive, as are basal cells of tubular structures. Luminal cells of tubular structures and small cell nests with polygonal to spindeloid cells are negative. | Luminal cells of about 50% of tubular structures are positve. Spindeloid cells and basal epithelial cells are negative. | Most luminal and basal cells of epithelial structures as well as small cell nests are positive. Spindeloid cells are negative. | Most basal cells of epithelial structures and small cell nests are positive. Luminal cells and spindeloid cells are negative. | 25-40% of spindeloid cells, often surrounding small tubular structures, and outer epithelial cells of tubular structures are positive. Remaining spindeloid cells are negative. | 9% | 4% | 5 | 5.70% | 4 | 1+ | Luminal A |
| 32 | CMT-32-A | A |  | Complex adenoma | Myoepithelial cells and basal cells of tubuli positive. | Luminal cells of double-layered tubuli and small single-layered tubuli positive. | Single-layered tubuli, basal cells of double-layered tubuli and about 75% of luminal cells positive. around 20-30% of spindeloid cells (myoepithelial cells) positive. | Basal cells of larger tubuli and around 25% of single-layerd tubuli positive. around 20% of spindeloid cells positive (scattered). | Basal cells of double-layerd tubuli and scattered cells in areas with small tubuli and spindeloid cells (around 10-20%). | 6.40% | 37.80% | 7 | 0.60% | 2 | 2+ | Luminal A |
|  | CMT-32-N | N |  | non neoplastic mammary tissue, mild to moderate interstitial fibrosis, mild lymphocytic infiltration. | Ducts and occasional acini with basal cells positive. | Luminal cells of acini and ducts. | Basal cells of acini and both basal and luminal cells of ducts positive. | Basal cells of acini and ducts positive. | Basal cells of acini and ducts positive. | 8.70% | 54.10% | 7 | 24.20% | 5 | 3+ |  |
| 33 | CMT-33-C | C | II | intraductal papillary carcinoma |  |  | In 25% of the tumor almost all neoplastic cells with moderate to strong signal. In 75% of the tumor, only superficial row of cells plus interspersed cells positive (around 10% of neoplastic cells). |  | Basal cells of papillary proliferations. | 3.40% | 1.50% | 5 | 12% | 4 | 1+ | Luminal A |
| 36 | CMT-36-C | C | II | Inflammatory carcinoma (only intralymphatic, no primary tumor, thus carcinoma type unknown) | around 75% of intralymphatic cells positive. | >99% intralymphatic cells positive. | Around 60% of intralymphatic cells positive. | >99% intralymphatic cells positive. | Intralymphatic cells negative. | 15% | 15% | 5 | <1% | 2 | 1+ | Luminal A |
|  | T-36-N | N |  | fibroadenomatous hyperplasia | Luminal cells negative, basal cells positive. | Luminal cells of acini and ducts positive. | Basal cells of acini and ducts, about 50% of ductal luminal cells positive. | Basal cells of acini and ducts, about 10% of ductal luminal cells positive. | Basal cells of acini and ducts positive. | 20% | 64% | 5 | 23% | 5 | 3+ |  |
| 51 | CMT-51-C4 | C | III | simple carcinoma, solid |  |  | >98% positive |  | 5-10% of neoplastic cells, frequently peripheral layer of islands and cords. | 44% | <1% | 3 | <1% | 3 | 0 | Triple negative basal like |
| 60 | CMT-60-C | C | I | complex carcinoma |  |  | >95% of basal cells of neoplastic tubuli, 75% of luminal cells of neoplastic tubuli (membranous signal) |  | >98% of basal cells of neoplastic tubuli, indivdual cells in between spindle cell proliferations (<5%) | 3.70% | 25% | 6 | 20% | 6 | 2+ | Luminal A |
|  | T-60-N | N |  | multifocal lobular hyperplasia |  |  | >95% of basal cells of ducts and acini, > 95% of luminal cells of ducts. Luminal cells of acini negative. |  | Basal cells of acini and ducts positive. | 8% | 68% | 8 | 18% | 6 | 3+ |  |
| 63 | CMT-63-C | C | II | complex carcinoma |  |  | 95% of basal cells and 80% of luminal cells of neoplastic tubuli |  | basal cells of neoplastic tubuli and 10-20% of spindle cells positive. | 6.80% | 12% | 5 | 20% | 5 | 1+ | Luminal A |
|  | T-63-N | N |  | lobular hyperplasia |  |  | almost 100% of basal cells of acini and ducts, around 10% of luminal cells of acini, 50% of luminal cells of ducts. |  | Basal cells of acini and ducts positive. | 10.00% | 60% | 7 | 13% | 5 | 2+ |  |

| Organoids | | | | | | | | | | | | | | | | | | |
| --- | --- | --- | --- | --- | --- | --- | --- | --- | --- | --- | --- | --- | --- | --- | --- | --- | --- | --- |
| Sample name | Normal / Adenoma / Carcinoma | HE | Vimentin | CK7/8 = Cam5.2 | CK5/6 | CK14 | p63 | Ki67 | ER (% positive cells) | ER (Allred score) | PR (% positive cells) | PR (Allred score) | HER2 (Pena) | Nielsen classification | IHC: match tissue/ organoid: good (3), medium (2), poor (1) | Passage reached | Thawing test success(s) / failure (f) / not tried (nt) | SNP assay: match tissue / organoids: Yes/no/not tried |
| ORG-01-C | C | Organoids with multiple small tubuli (cribriform), other organoids more solid with vacuolated poygonal cells. | almost all organoids surrounded by single layer of positive cells (20% of all cells) | 40-50% of all cells. Positive cells frequently arranged in tubuli and nests enclosed by negative cells. Some tubular organoids fully positive. | 80-90% positive. Negative cells encircled by positive cells. | around 50% positive, usually a single peripheral cell layer plus interspersed individual cells. Few organoids fully positive. | around 30-40% of the cells, in almost all organoids as a peripheral layer. Single organoids completely positive. | 25% | <1% | 2 | 0 | 0 | 2+ | Triple negative basal like | 3 | 2 | nt | nt |
| ORG-02-C | C | around 30% tubuloacini, other organoids with multiple irregular tubular lumina, or solid with vacuolated/pleomorphic cells. | around 5% (basal cells of 20% of all organoids) | 90% of all cells and organoids. Individual organoids completely negative. | 50-60% of all cells. Individual organoids fully positive, most organoids mix of positive and negative cells. | 80-90% of all cells, most organoids fully positive, others only basal cells or mixed. | around 10% of all cells, in groups (part of organoids, no clear distribution). | 49% | <10% | 4 |  |  | 1+ | Luminal B | 1 | 2 | nt | nt |
| ORG-03-C | C | around 40% single- to double-layered tubuloacini, around 10% organoids with multiple tubuli (cribriform), rest solid with vacuolated polygonal cells. | 5-10% of all cells, single layer of positive cells in about 50% of all organoids | around 60% of all cells. 20-25% of organoids fully pos, 50% mixed, rest negative | >95% cells positive. around 70% of organoids completely positive, rest mixed | 50-60%of cells positive. 40-50% of organoids fully positive, few mixed, rest with a single peripheral positive layer. | 40- 50% of cells positive. around 30% organoids fully positive, around 50% with peripheral positive layer, rest negative. | 33.40% | <1% | 2 | 0 | 0 | 2+ | Triple negative basal like | 2 | 3 | nt | nt |
| ORG-05-C | C | around 60% organoids tubuloacinar, around 40% organoids cribriform with comedonecrosis. | around 30% positive for cribriform organoids | tubulo acinar organoids 100% positive, cribriform organoids 50% positive | tubulo acinar organoids <10% positive, cribriform organoids 50% positive | around 10% positive cells for both cell types | all negative | 48.60% | 0.00% | 0 | 0 | 0 | 1+ | Triple negative basal like | 1 | 9 | nt | no |
| ORG-05-A | A | around 75% organoids solid to cribriform, around 25% organoids with squamous differentiation. One organoid with comedonecrosis. | 10% of all cells: single cell layer around all organoids. | around 60-70% of all cells. Organoids with squamous differentiation negative. | around 75% of all cells positive: Organoids with squamous differentiation fully positive, some organoids with acinar differentiation partially to completely positive, other only with single peripheral cell layer positive. | around 60-70% of all cells, distribution comparable to CK5/6. | around 40-50% of all cells positive: organoids with squamous differentiation largely positive, organoids with acinar differentiation luminal cells negative, peripheral cell layer positive | 30.70% | around 20% of "acinar" cells strong signal. | 6 | 26% | 6 | 1+ | Luminal A | 3 | 4 | s | nt |
| ORG-06-C | C | around 40% squamous differentiation, around 40% multilayered sheets with micropapillary projections, around 10% solid, around 10% double-layered tubuloacini. | 30% (basal and luminal cells of tubuli and cell multilayers) | 30% (basal and luminal cells of tubuli and cell multilayers) | 80-90% (cell multilayers and large islands) | 90% (cell multilayers and large islands) | 50% (cell multilayers and large islands partially stained) | 52.60% | 9.30% | 4 | 25.40% | 5 | 1+ | Luminal A | 2 | 3 | nt | nt |
| ORG-06-N | N | single-layered and bi-layered acini, solid "buds" and few squamous organoids | solid "buds" and single-layered acini positive. Bi-layerd acini with peripheral positive layer. | Luminal cells of bi-layered acini positive | 80-90% of all cells positive. around 10-20% of multi-layered acini with luminal negative layer. | 80-90% of all cells positive. around 10-20% of multi-layered acini with luminal negative layer. | 80-90% positive, in some organoids only peripheral cells. | 8.90% | 31.30% | 5 | 6.90% | 4 | 0 |  | 1 | 2 | nt | nt |
| ORG-08-C | C | small single- to double-layered tubuloacini with luminal secretion. | 60% of cells, >90% of organoids. | >98% of cells and organoids. | around 90% of organoids and cells. | around 25% of organoids completely positive, around 40% of organoids partially positive. 40-50% of cells positive. | <1% of cells (often groups of cells / non-tubuloacinar organoids) | 87.70% | 0% | 0 | 0% | 0 | 2+ | Triple negative basal like | 3 | 5 | nt | nt |
| ORG-11-N | N | bi-layered acini | 40% (outer layer) | 90% (weak to moderate staining intensity) | 90% basal and luminal cells (20% of organoids only basal) | 90% basal and luminal cells (20% of organoids only basal) | 40% of all cells, mainly peripheral, few groups/organoids with all cells positive | 53.40% | 47.80% | 5 | 1% | 2 | 0 |  | 1 | 9 | nt | yes |
| ORG-16-C | C | single-layered tubuli, with papillary luminal projections | around 10% of organoids with a peripheral single layer of positive cells. | >95% of all cells positive | >95% of all cells positive | 50% of all cells positive: basal cell layer and intermingled luminal cells. | around 10% of all cells, only peripheral/basal cells. | 4.48% | 1.16% | 3 | 5% | 3 | 0 | Luminal A | 2 | 5 | f | nt |
| ORG-17-C | C | single-layered tubuli and solid organoids with slit-like double-layered lumina | two organoids with a positive peripheral cell layer. | around 80% of cells positive | around 70-80% of cells positive. Some organoids with single peripheral positive cell layer | around 80% of cells positive. Some organoids with single peripheral positive cell layer. | 80% of organoids positive, mostly heterogenous mix of positive and negative cells, around 60% of all cells positive. | 1.98% | 49.50% | 7 | 0.50% | 2 |  | Luminal A | 3 | 11 | s | yes |
| ORG-17-N | N | single- to bi-layered tubuli, some with luminal secretion | around 60% of organoids with 1-3 peripheral layers of positive cells (around 10% of all cells) | around 20% of organoids weakly positive (mostly small organoids). | >95% of organoids and cells positive. Most organoids completely positive, few with only peripheral positive layer. | All organoids, around 50-60% of cells, mainly peripheral layer positive. | around 50% of organoids with a single peripheral positive layer. Few organoids with multiple layers or focal aggregates of positive cells. Small organoids negative. | 37.14% | 5.30% | 5 | 0.50% | 4 | 2+ |  | 2 | 12 | s | yes |
| ORG-25-C1 | C | 40-50% single to double-layered tubuli, 40-50% squamous differentiation, rest solid with small basaloid cells. | around 60% of cells and >90% of organoids positive. Small organoids completely positive, larger organoids (both tubuli and squamous) with single peripheral positive layer. | around 30% of organoids and 25% of cells positive. 50% of ubular organoids completely positive, rest negative. Squamous organoids usually with peripheral positive layer. Small solid organoids variably positive. | All organoids completely positive | All organoids and >90% of cells positive. | Around 80% of organoids and 60-70% of cells positive. Single-layered tubuli usually negative. Double-layered tubuli and squamous organoids with single peripheral positive layer. Small organoids usually completely positive. | 23.40% | 2.20% | 3 | 0% | 0 | 0 | Luminal A | 3 | 8 | s | yes |
| ORG-25-C2 | C | around 70% solid organoids, of those about 10% with squamous differentiation. About 25% of organoids solid with bilayered lumina (duct-like), about 5% composed of bilayered tubuli. | Around 90% of all organoids with one single positive layer in periphery. Desquamated cells around large organoids also positive. | Organoids composed of bilayered tubuli positive (ca. 5%), remaining negative or non-specific peripheric signal | All organoids and >99% of all cells positive. Few peripheral small buds and desquamated cells negative. | All organoids and >99% of all cells positive. Few peripheral small buds and desquamated cells negative. | All organoids at least partially positive. Positive are bilayered tubulie/ducts, 1-2 peripheral cell layers and some scattered cells in the center of larger organoids. Those centers with squamous differentiation are negative. | 68.90% | 5% | 4 | 5% | 4 | 0 | Luminal B | 2 | 12 | s | yes |
| ORG-32-A | A | single- to double-layered acinar structures with secretory activity | 15-20% of organoids with single peripheral positive cell layer (<10% of cells). | around 50% of organoids/cells moderately positive. (all cells per organoid) | All organoids, >95% of cells positive. | >98% of organoids/cells positive. Most organoids completely positive, some only partially. | around 5-10% of organoids with peripheral pos cells (around 2-5% of cells). | 12.58% | 49.50% | 6 | 0% | 0 |  | Luminal A | 3 | 11 | s | yes |
| ORG-32-N | N | mainly single-layered acinar structures | around 30% of organoids positive (completely or peripheral cell layer; mainly small organoids) | around 75% of organoids/cells moderately positive (all cells per organoid) | around 90% of organoids completely positive, few with only peripheral layer positive, 5-10% negative. | around 90% of organoids completely positive, few with only peripheral layer positive, 10% negative | around 50% of organoids with a single peripheral positive layer. Few organoids with multiple layers or focal aggregates of positive cells. Some organoids completely negative | 12.27% | 0.50% | 2 | 0% | 0 | 1+ |  | 1 | 6 | s | nt |
| ORG-33-C | C | uni- and double-layered acinar structures with central necrotic debris, few solid organoids. | negative | 80% of uni- and bilayerd organoids positive (all cells). Solid organoids usually negative. | all organoids negative. | Around 40% of organoids partially or completely positive, both acinar and solid ones. | 46% of positive cells | 20% | <1% | 3 | 3% | 5 | 1+ | Luminal A | 2 | 12 | s | yes |
| ORG-36-C | C | single- to double-layered acinar structures with secretory activity, some squamous differentiation | around 10% of organoids with a peripheral single layer of positive cells. | around 10% of organoids weakly positive | All organoids, >95% of cells positive | around 80% of cells positive. Some organoids with single peripheral positive cell layer. | 20% of positive cells | 22.30% | 3% | 5 | <1% | 4 | 2+ | Luminal A | 3 | 9 | s | yes |
| ORG-36-N | N | single-layered and bi-layered acini, solid "buds" and few squamous organoids | 5% of all cells, single layer of positive cells in about 50% of all organoids | 25% (basal and luminal cells of tubuli and cell multilayers) | around 80% of cells positive | all organoids positive | around 50% of organoids with a single peripheral positive layer. | 27.30% | 14% | 5 | 0% | 0 | 1+ |  | 2 | 10 | s | yes |
| ORG-51-C4 | C | single-layered and bi-layered acini, solid "buds" and few squamous organoids |  |  | around 10% of positive cells |  | all negative | 5% | <1% | 3 | 0% | 0 | 0 | Triple negative basal like | 2 | 12 | s | yes |
| ORG-60-C | C | uni- (rarely bi-)layered acini with luminal secretion |  |  | >99% of organoids, >95% of cells positive. Some luminal cells of bilayered organoids negative. |  | 50% of organoids and 20% of cells positive. | 1% | <1% | 3 | <1% | 3 | 1+ | Triple negative basal like | 1 | 9 | s | yes |
| ORG-60-N | N | Bi-layered > single-layered acini, few solid buds. |  |  | >98% of cells, all organoids positive. <1% organoids with negative luminal layer. |  | 55% of positive cells (basal cells in bilayered organoids, solid buds frequently completely positive) | 62% | 40% | 5 | 0% | 0 | 1+ |  | 2 | 9 | s | yes |
| ORG-63-C | C | Bi-layered > single-layered acini, few solid buds. | 20% of organoids with a positive peripheral cell layer. | 99% of organoids positive | 99% of organoids, 98% of cells positive | Around 80% of organoids with positive peripheral cells, 10% of organoids with also luminal cells positive. | 40% of organoids with mainly peripheral cells positive. | 14% | 8% | 4 | <1% | 3 | 1+ | Luminal A | 2 | 20 | s | yes |
| ORG-63-N | N | 40% organoids with squamous differentiation, remaining organoids small solid buds, few double-layered acini | <10% of organoids with peripheral positive cells. | Squamous organoids negative or with internal positive cell nests, small buds and acinar organoids positive. | Squamous organoids positive, small buds and acinar organoids with only peripheral cells positive. | Squamous organoids positive with internal negative cell nests, small buds and acinar organoids with only peripheral cells positive. | Almost all organoids positive: squamous organoids completely or partially, small buds and acinar organoids only peripheral layer. | 4% | <1% | 3 | <1% | 3 | 1+ |  | 1 | 21 | s | yes |

**Table S4.** Estimation of the number of mutations associated with each Single Base Substitution Signatures (SBS) following signature fit approach and comparison of prevalence of SBS between CMT and HBC

| Associations | SBS | CMT-2-C | CMT-6-C | CMT-10-C | CMT-11-C | CMT-13-C | CMT-14-C | CMT-15-C1 | CMT-15-C2 | CMT-16-C | CMT-17-C | CMT-18-C | CMT-23-C | CMT-24-C | CMT-25-C1 | CMT-25-C2 | CMT-32-C1 | CMT-32-C2 | CMT-33-C | CMT-41-C | CMT-51-C4 | CMT-51-C3 | Prevalence in our CMT cohort | Prevalence in HBC | Prevalence in other human cancer types |
| --- | --- | --- | --- | --- | --- | --- | --- | --- | --- | --- | --- | --- | --- | --- | --- | --- | --- | --- | --- | --- | --- | --- | --- | --- | --- |
| Age of diagnosis. Deamination of methyl cytosines | SBS1 | 0 | 71.05 | 87.72 | 47.94 | 80.41 | 0 | 92.92 | 117.14 | 42.59 | 124.34 | 0 | 164.71 | 100.24 | 470.39 | 230.82 | 0 | 99.420 | 172.703 | 92.88 | 533.66 | 117.241 | 80.95% (17/21) | > 75% | Yes, nearly all |
| APOBEC cytidine deaminases | SBS2 | 0 | 0 | 0 | 0 | 0 | 0 | 0 | 0 | 0 | 0 | 0 | 0 | 27.60 | 0 | 0 | 0 | 0 | 0 | 0 | 130.84 | 0 | 9.52% (2/21) | >75% | > 50% |
|  | SBS13 | 0 | 0 | 0 | 0 | 0 | 0 | 0 | 0 | 0 | 0 | 0 | 0 | 0 | 0 | 0 | 0 | 0 | 0 | 0 | 0 | 0 | 0 | >75% | Yes, many |
| Age of diagnosis? Aetiology unknown | SBS5 | 0 | 0 | 0 | 0 | 0 | 0 | 0 | 0 | 0 | 87.74 | 0 | 101.34 | 125.35 | 0 | 0 | 0 | 94.14 | 0 | 0 | 0 | 0 | 19.05 % (4/21) | >75% | Yes, many |
| Homologous Recombination Repair Deficiency | SBS8 | 0 | 0 | 68.52 | 0 | 0 | 0 | 0 | 0 | 0 | 0 | 0 | 0 | 290.52 | 0 | 0 | 0 | 0 | 0 | 0 | 458.53 | 0 | 14.29% (3/21) | >60% | Yes, many |
|  | SBS3 | 0 | 0 | 0 | 0 | 0 | 0 | 0 | 0 | 0 | 0 | 0 | 0 | 183.63 | 0 | 0 | 0 | 0 | 0 | 0 | 0 | 0 | 4.76% (1/21) | >20% | Yes, many |
| Possibly damage by reactive oxygen species | SBS18 | 0 | 0 | 0 | 0 | 0 | 0 | 0 | 0 | 0 | 0 | 0 | 82.11 | 101.45 | 75.05 | 52.78 | 0 | 46.28 | 221.42 | 0 | 135.92 | 44.93 | 38.1% (8/21) | <20% of samples | Yes: gastric and adrenocortical cancers, neuroblastoma, |
| Unknown | SBS127 | 0 | 0 | 0 | 71.46 | 92.59 | 91.79 | 80.64 | 89.12 | 55.38281 | 122.81 | 0 | 0 | 131.07 | 177.63 | 0 | 0 | 0 | 229.998 | 98.05 | 435.65 | 75.33 | 61.9% (13/21) | 63% | Yes, bone and soft tissue samples |
| Unknown | SBS17 | 0 | 0 | 0 | 0 | 0 | 0 | 0 | 0 | 0 | 0 | 0 | 0 | 0 | 0 | 0 | 0 | 0 | 0 | 0 | 0 | 0 | 4.76% (1/21) | rare <5% of samples | Yes, common in esophageal, liver, gastric and lung cancers, lymphomas |
| Possible artefact | DogUnknownSig | 259.9753 | 256.40 | 319.51 | 171.66 | 318.598 | 274.33 | 289.61 | 355.67 | 274.45 | 320.75 | 260.85 | 337.08 | 295.67 | 325.59 | 267.87 | 272.78 | 306.91 | 406.52 | 382.39 | 458.74 | 327.47 |  |  |  |
|  | unassigned | 28.02 | 126.55 | 135.24 | 100.93 | 23.39 | 58.87 | 132.83 | 178.07 | 71.58 | 100.36 | 79.15 | 204.77 | 4.47 | 74.34 | 120.54 | 38.22 | 118.25 | 198.36 | 85.6751 | 176.66 | 68.02 |  |  |  |

**Table S5.** Genes targeted by the custom canine CRISPR library and gRNAs counts (plasmid DNA pDNA and day 0 D0)

| Construct Barcode | Gene Symbol | pDNA | ORG_63_C_D0_R1 | ORG_63_C_D0_R2 | ORG_63_N_D0_R1 | ORG_63_N_D0_R2 |
| --- | --- | --- | --- | --- | --- | --- |
| ACAGATCTTTACCCGAGATG | ABCB1 | 375 | 1069 | 1974 | 1930 | 2135 |
| ATCGTGGTGGCAAACAACAC | ABCB1 | 480 | 487 | 847 | 763 | 828 |
| ATTGACAGCTATTCGAAGAG | ABCB1 | 364 | 869 | 1563 | 1448 | 1559 |
| CAAACGATGAGCTATCACAA | ABCB1 | 493 | 455 | 651 | 667 | 692 |
| GGTATGCCTATTATTACAGT | ABCB1 | 538 | 386 | 877 | 820 | 769 |
| TGCATAGATCAATAAGAAAG | ABCB1 | 391 | 405 | 817 | 883 | 893 |
| AAACGTACGGACAACCATCA | ABCC2 | 393 | 769 | 1093 | 1147 | 1153 |
| ACTGCTGTCACGTCGAATGG | ABCC2 | 596 | 879 | 1826 | 1789 | 1804 |
| CAGCAATCCTGGATTCAGAA | ABCC2 | 445 | 500 | 610 | 711 | 626 |
| GATTGGTGTATCTAATGGCG | ABCC2 | 521 | 666 | 1312 | 1293 | 1328 |
| TCGAACACTTAGTCGCAGGT | ABCC2 | 483 | 568 | 1196 | 947 | 1017 |
| TGATGAACAGATTACAACCA | ABCC2 | 476 | 510 | 1035 | 930 | 910 |
| AGGAAATCTAGAAACGTCAG | ABCC4 | 405 | 522 | 1160 | 1189 | 1190 |
| TACACAGGTTGCTTTGTCAG | ABCC4 | 600 | 724 | 1570 | 1539 | 1601 |
| TGACGTGAACAAGTTTGACC | ABCC4 | 392 | 456 | 669 | 718 | 711 |
| TGCCTGGGAAAAGTCCTTTG | ABCC4 | 332 | 615 | 893 | 1031 | 1058 |
| TGTTACCTTGTCCCAAAAGG | ABCC4 | 543 | 724 | 1109 | 1022 | 1078 |
| TTAACCGTGATAGGTGATCG | ABCC4 | 349 | 867 | 1553 | 1744 | 1771 |
| AACTTGAAGATAGAATAACG | ABCG2 | 629 | 370 | 609 | 488 | 530 |
| ACATCTCCCGATAATCCATG | ABCG2 | 375 | 498 | 984 | 890 | 917 |
| ACCTGAGCGATAGAGGCCTG | ABCG2 | 479 | 702 | 1319 | 1358 | 1386 |
| GTTGGAACTCAGTTTATACG | ABCG2 | 519 | 547 | 1182 | 1022 | 1109 |
| TATCGAGTAAAAGTAAAGAG | ABCG2 | 424 | 457 | 709 | 665 | 674 |
| TCTGTTTAATACCACAGCAG | ABCG2 | 394 | 714 | 1180 | 1170 | 1257 |
| AACATACCTGGTCCAACAGT | ABI1 | 584 | 402 | 915 | 839 | 739 |
| AGAGTTACCAGAATCTGACC | ABI1 | 341 | 394 | 766 | 592 | 543 |
| AGCTAGGGATTGAGTTGTGT | ABI1 | 541 | 723 | 1312 | 1249 | 1285 |
| CAAATTTAAAATCTAGACCG | ABI1 | 400 | 525 | 862 | 742 | 812 |
| GAAGTCAACATAGTCCAGGC | ABI1 | 444 | 237 | 481 | 474 | 496 |
| GTTGTGTCGAGAAATCTGCC | ABI1 | 475 | 476 | 854 | 718 | 711 |
| CTGCTCCCTCGAAAAGAACG | ABL1 | 511 | 506 | 1050 | 1021 | 1001 |
| GGGCCGAGAAACCAGCAATG | ABL1 | 521 | 835 | 1550 | 1424 | 1443 |
| GGGGAAAAAAGGTGTACGAG | ABL1 | 568 | 561 | 1021 | 943 | 1122 |
| GGTTCATCATCACTCGACTG | ABL1 | 437 | 592 | 1176 | 1055 | 1113 |
| TGGGTAAGGTGACATGCCAT | ABL1 | 604 | 626 | 1148 | 1130 | 1093 |
| TTTAGGATACAATCACAATG | ABL1 | 336 | 422 | 805 | 701 | 777 |
| AAAGACACTGGGCTTACGAG | ABL2 | 533 | 17 | 26 | 12 | 20 |
| CAACATCACAACCATAGGGG | ABL2 | 477 | 485 | 987 | 856 | 924 |
| GTCCCCTATCCATGACAAGT | ABL2 | 466 | 393 | 693 | 694 | 699 |
| GTGACTGAGTACATGCCGTA | ABL2 | 451 | 524 | 1220 | 1198 | 1207 |
| TATCGAATGGAACAACCTGA | ABL2 | 310 | 382 | 565 | 559 | 520 |
| TGCTTCACCAGGGATAGGAA | ABL2 | 536 | 501 | 863 | 804 | 780 |
| AAGTGAAGGAAGCCTACATG | ACAT1 | 636 | 643 | 1190 | 945 | 1010 |
| ACCAAGTTTAGTGGCCGGCA | ACAT1 | 456 | 631 | 1211 | 1367 | 1407 |
| ATTACAGGCAGTCAGATACG | ACAT1 | 673 | 778 | 1076 | 1149 | 1134 |
| CTCCTCTGCTCATGACGTAG | ACAT1 | 566 | 886 | 1526 | 1515 | 1504 |
| GAAGAAGATGAACATTACAC | ACAT1 | 440 | 548 | 589 | 517 | 543 |
| TTTGTTTACAGTGGTGCACG | ACAT1 | 405 | 664 | 1148 | 1253 | 1265 |
| AGGGCATATGCGTTGATCAC | ACER1 | 633 | 800 | 1295 | 1297 | 1343 |
| CCAGGAAGCTAAGCGTCATG | ACER1 | 616 | 713 | 1380 | 1551 | 1575 |
| CTTTCCTAGGAGAGAACAGG | ACER1 | 584 | 553 | 863 | 852 | 900 |
| GGACCAGCAATAAGGAGCTT | ACER1 | 328 | 579 | 1136 | 1072 | 1080 |
| GGGAGCGTTTCTGGGCATAG | ACER1 | 620 | 1076 | 2066 | 2045 | 2054 |
| TGGATGAGATCGCTATCCTG | ACER1 | 588 | 713 | 1492 | 1413 | 1430 |
| AATGTGCGTGTCTTTAAACT | ACER2 | 479 | 506 | 771 | 789 | 748 |
| CATCGCCGAGTTCTACAACA | ACER2 | 636 | 420 | 590 | 655 | 658 |
| CCCCAGAAGGTATCTACCAA | ACER2 | 605 | 684 | 1183 | 1143 | 1162 |
| GCACAGTGCATGGAACCCCC | ACER2 | 408 | 397 | 920 | 786 | 851 |
| GGGCACACATCAGAACCCAA | ACER2 | 386 | 505 | 512 | 465 | 444 |
| TATGCAACATGCTTCAACAG | ACER2 | 429 | 425 | 824 | 838 | 792 |
| ATACAACAGCTGTAAATCAT | ACER3 | 699 | 421 | 635 | 763 | 719 |
| CAGTCCAGTGTGGACGTCGT | ACER3 | 544 | 552 | 968 | 1012 | 1015 |
| GCAATTCAGAGTGTTAGAGA | ACER3 | 524 | 504 | 912 | 724 | 721 |
| TGAGCCATGGATAAACCCTG | ACER3 | 528 | 1056 | 1682 | 1564 | 1593 |
| TTGGAGGTATAATCATAATC | ACER3 | 674 | 532 | 971 | 1029 | 1058 |
| TTTCATATTTCAGAGTCATG | ACER3 | 539 | 747 | 1223 | 968 | 1113 |
| AACACGTGAAATCTTAAGCG | ACSL3 | 513 | 588 | 1040 | 961 | 971 |
| ACGTTGTACGCTACTCTAGG | ACSL3 | 530 | 675 | 997 | 886 | 890 |
| CAGTTATCATGTATACGAGT | ACSL3 | 336 | 530 | 654 | 645 | 627 |
| GATGTTCATGAATCGCTGTG | ACSL3 | 627 | 603 | 1058 | 1124 | 1108 |
| TATGTTGAAACCAACACTGA | ACSL3 | 490 | 392 | 558 | 563 | 581 |
| TGGACGGTAAGCCACCAACG | ACSL3 | 411 | 625 | 1131 | 1301 | 1287 |
| ACATCCGGTTCAGCAATCGA | ACSL6 | 281 | 616 | 850 | 834 | 886 |
| ATGATGATGAGCTTGAGCCC | ACSL6 | 435 | 475 | 926 | 855 | 865 |
| CCAATAAACTGATCCGTACA | ACSL6 | 520 | 642 | 1188 | 1178 | 1144 |
| CTGAGATGCTAAGCCCACGG | ACSL6 | 580 | 686 | 1326 | 1316 | 1264 |
| GATGCTTACCCATGGGAACG | ACSL6 | 424 | 475 | 1167 | 1108 | 1220 |
| TGGCCGGTGAGTGAACCAGT | ACSL6 | 626 | 500 | 1113 | 1067 | 980 |
| AATAGTGATGACCTGCCCGT | ACTB | 379 | 317 | 708 | 650 | 689 |
| ACGCACGATTTCCCGCTCGG | ACTB | 417 | 603 | 1202 | 1000 | 1138 |
| ATGCCCACCATCACGCCCTG | ACTB | 401 | 722 | 1229 | 1110 | 1046 |
| CAACCGTGAGAAGATGACCC | ACTB | 513 | 450 | 903 | 860 | 803 |
| CGTGATGGACTCCGGTGACG | ACTB | 429 | 710 | 1566 | 1449 | 1421 |
| TGACGATGCCGTGCTCAATG | ACTB | 554 | 856 | 1457 | 1271 | 1291 |
| ACGGTGCGCTGGACGAACAG | ACVR1B | 676 | 985 | 1702 | 1803 | 1731 |
| CCACATCACCACCCCTCCAG | ACVR1B | 494 | 461 | 875 | 794 | 818 |
| GAGAGACACATCTCACACGA | ACVR1B | 570 | 307 | 519 | 483 | 572 |
| GATACACGGTGACGATCGAG | ACVR1B | 544 | 762 | 1268 | 1281 | 1297 |
| TCTGTAACAAGATCGACCTG | ACVR1B | 448 | 896 | 1344 | 1456 | 1365 |
| TGTCAGCAGCAATAAATCCA | ACVR1B | 498 | 687 | 1192 | 1328 | 1380 |
| AAAGAGTCTACATTCAGACG | ADAMTSL3 | 632 | 761 | 965 | 939 | 937 |
| AAGATGAAATCAGCCATCAG | ADAMTSL3 | 646 | 464 | 752 | 815 | 814 |
| CAGTCCCCCACAGTTATATG | ADAMTSL3 | 517 | 861 | 1607 | 1504 | 1436 |
| GCAAACACGTAACTCACCGA | ADAMTSL3 | 569 | 418 | 648 | 564 | 605 |
| GCAGCAGGATCATTATACCG | ADAMTSL3 | 504 | 1001 | 1292 | 1316 | 1272 |
| TGAACCCGGCATACTCCCAG | ADAMTSL3 | 608 | 420 | 856 | 776 | 736 |
| CTGGAACACCACTTGGCGTA | ADGRA2 | 469 | 1038 | 1344 | 1428 | 1368 |
| CTTCTCCAGTCTACAACCTG | ADGRA2 | 411 | 769 | 1362 | 1317 | 1250 |
| GCGAGTCGCCAACAATCGCG | ADGRA2 | 338 | 534 | 943 | 835 | 761 |
| GGGAAACCTGACCGAGCCGG | ADGRA2 | 438 | 568 | 1152 | 987 | 1015 |
| TCACCTACATCCTCAACCAC | ADGRA2 | 492 | 339 | 471 | 499 | 463 |
| TCCCCAGAACTCAAGGAACG | ADGRA2 | 654 | 702 | 1258 | 1189 | 1222 |
| AAACAACTCACCTCTAACAG | ADGRL3 | 412 | 428 | 577 | 578 | 585 |
| AACAGAATGGCCGAAACGGT | ADGRL3 | 671 | 422 | 979 | 979 | 951 |
| ACAGGTTATTGGTCAACACA | ADGRL3 | 425 | 829 | 1369 | 1397 | 1527 |
| GTAGTCAATCTTATTCCCTG | ADGRL3 | 287 | 652 | 1160 | 944 | 916 |
| GTCCATGGCACGGACTGAGT | ADGRL3 | 524 | 564 | 1128 | 1145 | 1058 |
| TGGAAGCCCGAGAAATCATG | ADGRL3 | 473 | 602 | 1028 | 1016 | 1055 |
| CAGGTACATGAGCCAGAGTG | ADORA2A | 390 | 773 | 1422 | 1518 | 1659 |
| GAGGGCAGAAACTACTCACA | ADORA2A | 502 | 593 | 1126 | 976 | 1123 |
| GCGGCAGCCGACATTGCCGT | ADORA2A | 664 | 512 | 826 | 892 | 845 |
| GTGACTGGCACAAGGGCCAA | ADORA2A | 384 | 517 | 945 | 1061 | 1106 |
| TGCGGATCTTCCTAGCAGCT | ADORA2A | 465 | 456 | 987 | 806 | 887 |
| TGGCCAGTGACTTGGCAGCG | ADORA2A | 625 | 744 | 1494 | 1352 | 1299 |
| AAACCCCGACCAAAGAGTGA | AFDN | 691 | 734 | 1311 | 1141 | 1139 |
| AAGGATGATAGGCCTTTCCA | AFDN | 508 | 348 | 711 | 545 | 587 |
| AGATCTTAGTCGAATCACAC | AFDN | 361 | 293 | 560 | 386 | 392 |
| AGGGTCCCAGACATAAACCT | AFDN | 365 | 583 | 1072 | 979 | 998 |
| CAGTAACGCTCAACTGAAGG | AFDN | 495 | 577 | 1023 | 958 | 946 |
| GGGAGCACAGTCCCGACTCA | AFDN | 665 | 823 | 1442 | 1268 | 1294 |
| AGCCCACGAAGGACAATGCG | AFF1 | 450 | 640 | 1375 | 1451 | 1361 |
| AGCCGCACACCTCCTGACCA | AFF1 | 424 | 814 | 1427 | 1405 | 1434 |
| GGCATCCAAGCGATGAGGGT | AFF1 | 500 | 633 | 1209 | 1170 | 1102 |
| GTGTGTATTGCTGTCAAAGG | AFF1 | 674 | 714 | 1135 | 1136 | 1128 |
| TGCCACTAAATTGTCTTGAG | AFF1 | 573 | 952 | 1620 | 1656 | 1638 |
| TGCTTCTGGAAGCGACTGCG | AFF1 | 473 | 814 | 1293 | 1227 | 1181 |
| AACCAGCGTCCTCTAACAAG | AFF3 | 496 | 641 | 1000 | 1042 | 1032 |
| ACTTGGATTATTGTGTCCGG | AFF3 | 485 | 801 | 1703 | 1698 | 1613 |
| AGAAACCAACTGCATATGTG | AFF3 | 497 | 634 | 1334 | 1313 | 1280 |
| CAGCAGAACGAGCCACCGCA | AFF3 | 384 | 565 | 1028 | 1074 | 1049 |
| TCTTCCAACTCAGCACAACA | AFF3 | 416 | 363 | 496 | 466 | 467 |
| TCTTTCTCCAGACTAACAAG | AFF3 | 503 | 339 | 641 | 561 | 598 |
| ACCTCAGCGTCTTCTACCAG | AGTRAP | 362 | 663 | 1213 | 1237 | 1265 |
| GACACGCAGCGCTTCGGCGC | AGTRAP | 379 | 532 | 948 | 1005 | 1030 |
| GATGGAGTCCCGCTGAGCCA | AGTRAP | 636 | 635 | 1266 | 1112 | 1138 |
| GGGGCTGCATCACGTTCTCG | AGTRAP | 440 | 761 | 1666 | 1723 | 1646 |
| TCTGTCTTGCAGTTTCTGGG | AGTRAP | 623 | 611 | 1242 | 1210 | 1213 |
| TGAACCAGCAGGATCATCTG | AGTRAP | 559 | 796 | 1522 | 1432 | 1625 |
| ACGTGCATTAGACTGAACCC | AHR | 439 | 650 | 998 | 1128 | 1164 |
| AGGATACTATCACTTCCCAT | AHR | 346 | 351 | 696 | 711 | 795 |
| CTGAAGCAGAGTTGTGCATG | AHR | 607 | 1059 | 1956 | 1949 | 2001 |
| CTTATTCATACTGAAGACCG | AHR | 605 | 735 | 1452 | 1542 | 1327 |
| TAGTGGTAAGGGATCCATCA | AHR | 515 | 827 | 1336 | 1247 | 1237 |
| TGTCTGCTGGATAATTCGTC | AHR | 323 | 439 | 872 | 837 | 814 |
| CCTGCACTCGGAGAAGAACG | AKT1 | 575 | 525 | 1150 | 1037 | 1041 |
| CGTCATGGAGTACGCCAACG | AKT1 | 637 | 861 | 1487 | 1375 | 1461 |
| GCTCCTCGGGTGTCTCCACG | AKT1 | 10 | 724 | 1334 | 1337 | 1353 |
| GCTGTGCAAGGAAGGCATCA | AKT1 | 371 | 856 | 1345 | 1251 | 1321 |
| GGCCAAGCCCAAGCACCGTG | AKT1 | 585 | 715 | 1309 | 1231 | 1204 |
| TACTTCCTCCTCAAGAACGA | AKT1 | 584 | 305 | 520 | 522 | 450 |
| AGACAACGACTACGGCCGCG | AKT2 | 603 | 595 | 1067 | 1033 | 960 |
| AGTGGACCACTGTCATCGAG | AKT2 | 474 | 835 | 1413 | 1578 | 1572 |
| CCTCGAATGCAAATACTCGA | AKT2 | 558 | 419 | 573 | 580 | 626 |
| GACCCCATGGACTATAAGTG | AKT2 | 520 | 757 | 1352 | 1190 | 1198 |
| GGGAAGAACCACTCACGGTG | AKT2 | 481 | 565 | 1336 | 1261 | 1326 |
| TTGGCAAAGTGATTCTGGTG | AKT2 | 554 | 958 | 1597 | 1383 | 1364 |
| ACAAATTGATAATATAGGGG | AKT3 | 455 | 733 | 1199 | 1147 | 1231 |
| AGGATTCGTAAGATGGCGGG | AKT3 | 538 | 1078 | 1509 | 1530 | 1529 |
| ATTTCATGTAGATACTCCAG | AKT3 | 550 | 729 | 1140 | 1167 | 1145 |
| CTGCACCATAGAAACGTGTG | AKT3 | 463 | 774 | 1311 | 1299 | 1300 |
| TATTTGAAACTACTAGGTAA | AKT3 | 629 | 643 | 862 | 910 | 857 |
| TTGCCACTGAAAAGTTGTTG | AKT3 | 339 | 1011 | 1770 | 1833 | 1919 |
| ACCTCGATGATCCTGGACCC | ALK | 413 | 506 | 1161 | 1077 | 1082 |
| ACCTGTAGGGTCAAAGACGG | ALK | 418 | 683 | 1461 | 1323 | 1312 |
| CACTCCTGACACTTACGCTG | ALK | 464 | 391 | 675 | 759 | 746 |
| GCCCTGGAATACATCTCGAG | ALK | 500 | 599 | 1050 | 1012 | 1092 |
| GCTGAACGGCAACTCCGGAG | ALK | 444 | 657 | 1142 | 1249 | 1302 |
| TGGTTGCCAAGGTCGTGTAG | ALK | 313 | 1018 | 1767 | 1922 | 1843 |
| ACCATCCAGATCAACAGCAT | ALOX12B | 399 | 362 | 690 | 681 | 707 |
| CCTTCTGGAGCACCGACAAG | ALOX12B | 596 | 898 | 1623 | 1581 | 1617 |
| GAGAGACTTTGCAACTGGGG | ALOX12B | 467 | 533 | 1166 | 974 | 1078 |
| GTGCAGATCTGTGCACCCGA | ALOX12B | 464 | 593 | 939 | 953 | 968 |
| GTTCCCGGTCACAAATGACA | ALOX12B | 416 | 314 | 471 | 414 | 447 |
| TGGGCATCATGTTCCCTTCG | ALOX12B | 487 | 962 | 1644 | 1686 | 1593 |
| AATGACCTGTTCAAAACCAC | ANPEP | 583 | 259 | 403 | 352 | 328 |
| AGCGCCCAGGTACTCCACGT | ANPEP | 575 | 226 | 742 | 701 | 684 |
| CATTCTCTGTATACTCGCTG | ANPEP | 512 | 682 | 1231 | 1233 | 1278 |
| CCATGGAAATTATGCCCTGA | ANPEP | 397 | 1142 | 1681 | 1511 | 1498 |
| GTACAGGCCATTACTGTTAG | ANPEP | 600 | 693 | 1271 | 1308 | 1337 |
| TCTGGGCAGCATGTTAGACA | ANPEP | 411 | 692 | 1300 | 1238 | 1307 |
| AGAAATACCTGACTTGGGTA | ANXA7 | 533 | 775 | 1393 | 1219 | 1290 |
| CCACCCTCACAGTCTTATGG | ANXA7 | 421 | 576 | 1085 | 1104 | 1115 |
| GACAGATGAACAGGCAATTG | ANXA7 | 454 | 736 | 1255 | 1294 | 1253 |
| GAGTTTACGGAATGCAATGA | ANXA7 | 532 | 770 | 1409 | 1376 | 1485 |
| GGATAAGGATACTGACCCGG | ANXA7 | 518 | 594 | 900 | 889 | 787 |
| TTGGGTTCCCTGAGTCATTG | ANXA7 | 391 | 1174 | 1977 | 1732 | 1748 |
| AGAACAACTAGGTACTTGCC | APC | 691 | 588 | 1111 | 1182 | 1096 |
| CAGCTTGATAGCTACAAACG | APC | 507 | 773 | 1268 | 1328 | 1387 |
| CAGTGCTCTCAGTATATACG | APC | 435 | 635 | 1213 | 1322 | 1311 |
| CCTTCGACACAGTACACCCG | APC | 498 | 709 | 1371 | 1341 | 1355 |
| GCAACGCGAGAAACAAATTG | APC | 430 | 696 | 1180 | 1128 | 1076 |
| TCTACCACTGAGTTACACTG | APC | 328 | 505 | 976 | 956 | 1008 |
| AGTAATAGGGTGAGTCTCCG | APH1A | 443 | 772 | 1392 | 1359 | 1413 |
| CCAGCCAGAAAAATGCCCTG | APH1A | 557 | 773 | 1405 | 1457 | 1472 |
| CTCCCCAAAAGGTATGGAGC | APH1A | 474 | 414 | 820 | 871 | 895 |
| GACACCACTGATGATACCGA | APH1A | 680 | 465 | 684 | 686 | 681 |
| GAGGCGGGCATCTGACCGGT | APH1A | 488 | 502 | 750 | 730 | 777 |
| GGGGTTAGCATCGCTGAGTG | APH1A | 585 | 771 | 1448 | 1500 | 1522 |
| AGATGGTCGAACTGCCACCT | AR | 400 | 694 | 1370 | 1336 | 1262 |
| AGCGGCACAATGCCGCTACG | AR | 466 | 640 | 1042 | 1191 | 1161 |
| CCTGATGTGTGGTATCCGGG | AR | 507 | 643 | 1387 | 1345 | 1368 |
| CGGAGCGCAAGGACGTACGG | AR | 690 | 590 | 1092 | 1180 | 1172 |
| GCTGAGTCATTCTCGTCCGG | AR | 587 | 484 | 1056 | 1120 | 996 |
| TTCATCACCGCAGATCAGAC | AR | 450 | 344 | 435 | 399 | 423 |
| AGTGTCCAGGATTTGTCCGG | ARAF | 525 | 788 | 1486 | 1470 | 1452 |
| CAGCAAGATGTTGACATGAC | ARAF | 481 | 417 | 866 | 810 | 879 |
| GGTGCTGACCATATGGACGT | ARAF | 494 | 548 | 1022 | 1023 | 1039 |
| TGGCACCTCCCAGTAATAGC | ARAF | 421 | 414 | 545 | 558 | 545 |
| TGGTCAGTGGGACGTCTTCG | ARAF | 548 | 1042 | 1739 | 1720 | 1712 |
| TGTCTACGACTCTCTTGACA | ARAF | 445 | 606 | 917 | 943 | 989 |
| AACCACAAAGATGACACCTG | ARF1 | 478 | 709 | 1229 | 1182 | 1060 |
| AGAAATGCGCATTCTCATGG | ARF1 | 469 | 899 | 1328 | 1337 | 1317 |
| AGCTACACGATCTGGGATGT | ARF1 | 459 | 884 | 1347 | 1333 | 1377 |
| GCGTCAAGATCAGGCCTCTG | ARF1 | 569 | 1035 | 1753 | 1848 | 1923 |
| GGCCCATGAGGAGTCTCATG | ARF1 | 550 | 643 | 1080 | 1043 | 1131 |
| TGCTCCCACAGGCTTCAACG | ARF1 | 538 | 465 | 825 | 659 | 756 |
| CGGATCTCAAGCAAGTCAGA | ARFGAP1 | 411 | 362 | 606 | 629 | 637 |
| GAGGACGAGGTCGAGTTCTG | ARFGAP1 | 567 | 739 | 1444 | 1415 | 1462 |
| GCAGGACAAGTACAACAGCA | ARFGAP1 | 452 | 392 | 635 | 616 | 686 |
| GTGGTGCATTCAACCCTCAG | ARFGAP1 | 594 | 539 | 1176 | 1096 | 1072 |
| TGTTCCCGAACCCCACGTAG | ARFGAP1 | 463 | 639 | 1273 | 1163 | 1107 |
| TTAACCCACGCAGGTGAAAG | ARFGAP1 | 441 | 612 | 1120 | 1164 | 1234 |
| AGGTCTTAATGCACCGACGA | ARFGAP3 | 421 | 589 | 1137 | 1137 | 1214 |
| CACTGGCCTCCCAGGCAACG | ARFGAP3 | 537 | 705 | 1465 | 1468 | 1538 |
| CAGCTGCGATGCATGCAGGT | ARFGAP3 | 406 | 535 | 1163 | 1078 | 1009 |
| GAGTCGGAGAGACTTGGCAT | ARFGAP3 | 545 | 602 | 1222 | 1006 | 1054 |
| GATACAAAGAAACACCCCAT | ARFGAP3 | 506 | 417 | 637 | 645 | 612 |
| TTTCTGGGCTTTGCTACGAT | ARFGAP3 | 373 | 692 | 1295 | 1218 | 1211 |
| ACTTCCACAGTGCGGATTGG | ARFRP1 | 366 | 1255 | 2153 | 1888 | 1963 |
| AGAGAAGATGGTGACGAGTG | ARFRP1 | 426 | 797 | 1502 | 1497 | 1520 |
| CCGCTTTAACAAGAACTACA | ARFRP1 | 352 | 306 | 413 | 381 | 393 |
| GACAATGCGGGCAAGACGGT | ARFRP1 | 606 | 405 | 725 | 705 | 730 |
| TGGCACTCCGCATAGTACTG | ARFRP1 | 395 | 480 | 1256 | 1021 | 1025 |
| TTGATTCCACGGATGAGGAG | ARFRP1 | 340 | 806 | 1375 | 1409 | 1303 |
| AATCATTCAGCCACGAGGAG | ARG1 | 538 | 604 | 1164 | 1033 | 960 |
| ACTCCACTGACAACCACAAG | ARG1 | 683 | 326 | 706 | 660 | 626 |
| ATCACTAGGGACATCAACAA | ARG1 | 363 | 322 | 542 | 551 | 596 |
| ATGACTGAAGTGGATAAACT | ARG1 | 580 | 429 | 732 | 676 | 629 |
| GAAGAACGGAAGGACTAGCC | ARG1 | 668 | 187 | 305 | 292 | 271 |
| GTATATTGGCCTAAGAGATG | ARG1 | 419 | 643 | 1169 | 917 | 994 |
| AATAGTGAATCCTCGCTCAG | ARG2 | 473 | 538 | 869 | 1020 | 965 |
| ATATATTGGTCTAAGAGATG | ARG2 | 474 | 883 | 1357 | 1382 | 1326 |
| GACGAAGTCCGTTCACTCGG | ARG2 | 533 | 711 | 1294 | 1223 | 1226 |
| GGGTGTATTGATGTCAGCAT | ARG2 | 388 | 636 | 1059 | 1041 | 1025 |
| GTTTGATCCAGGAAAATCCT | ARG2 | 441 | 495 | 899 | 772 | 754 |
| TATAGCTGTGTCACAATGGG | ARG2 | 494 | 733 | 1535 | 1585 | 1591 |
| AGGGCTCTACCGAATCGTGG | ARHGAP26 | 382 | 833 | 1504 | 1562 | 1703 |
| CCTTGGCCAGTTCATAACCA | ARHGAP26 | 383 | 540 | 979 | 1080 | 1096 |
| GGATTCCAAACAAATCACCA | ARHGAP26 | 322 | 269 | 346 | 293 | 278 |
| TGAAACTGGTACATCATGAG | ARHGAP26 | 504 | 752 | 1395 | 1431 | 1429 |
| TGAAGATGAACGGATACGGA | ARHGAP26 | 452 | 661 | 1118 | 951 | 1093 |
| TGGTGATAACCCCTGGCCTG | ARHGAP26 | 464 | 909 | 1718 | 1633 | 1768 |
| CAATAGACGATCTCCCCATG | ARID1A | 457 | 1056 | 1522 | 1459 | 1586 |
| CCTGTTGACCATACCCACTG | ARID1A | 531 | 674 | 1029 | 1007 | 1039 |
| GGAGCCTGGAATAGGACCCG | ARID1A | 463 | 879 | 1620 | 1692 | 1565 |
| GGGCCGTACTGACTCCCCTG | ARID1A | 461 | 701 | 1444 | 1431 | 1355 |
| GTCCCACCCATAGGATACGG | ARID1A | 575 | 743 | 1055 | 1077 | 1076 |
| TTTGAGTGCAAGATTGAACG | ARID1A | 426 | 1029 | 1504 | 1558 | 1602 |
| CAGATGACCCCGATGCAAGG | ARID1B | 442 | 724 | 1112 | 1139 | 1187 |
| CTGCCCATCCCATACAACTG | ARID1B | 550 | 615 | 965 | 938 | 1001 |
| GTAATTATTAAACTCCGAGA | ARID1B | 378 | 266 | 461 | 466 | 432 |
| GTCCATACTGAGACATCTGA | ARID1B | 374 | 411 | 660 | 577 | 584 |
| TGTGCCTTACGGTTCACGGT | ARID1B | 450 | 330 | 608 | 633 | 637 |
| TTATCCAGGCATGAACCGCA | ARID1B | 528 | 387 | 637 | 749 | 656 |
| ACAGTCTCCATTACACACAG | ARID2 | 608 | 482 | 665 | 645 | 588 |
| CAATATTTCCTAATGTGTCA | ARID2 | 702 | 534 | 880 | 836 | 891 |
| CCTCATGGACCACAAACTGT | ARID2 | 636 | 560 | 1025 | 1041 | 1027 |
| GCTTACCACCCGGTGTCGAA | ARID2 | 569 | 316 | 795 | 772 | 874 |
| GTAAGCCAGCCAGCTCAACA | ARID2 | 561 | 576 | 1011 | 905 | 898 |
| TATGTAAAAACACTAACCTG | ARID2 | 489 | 636 | 1114 | 967 | 1033 |
| ACGCCCCTGCTCTACTCGAG | ARID5B | 444 | 502 | 1192 | 1021 | 1027 |
| AGGCAGTCAACACTCAACAG | ARID5B | 393 | 377 | 799 | 723 | 692 |
| ATAAGTTTACCACCTCGCCG | ARID5B | 379 | 441 | 1018 | 807 | 780 |
| CAGCTTGAAACATAGTCCAA | ARID5B | 521 | 453 | 659 | 672 | 656 |
| GGTACCAGGATTTCGCGGCG | ARID5B | 609 | 779 | 1509 | 1407 | 1437 |
| TGCACGTGCTGAATCACCGA | ARID5B | 550 | 603 | 1016 | 965 | 945 |
| AAGGACCTGGAAAATTGCCA | ASAH1 | 401 | 705 | 1043 | 978 | 959 |
| CAAACCTTTATCAGTGAACC | ASAH1 | 494 | 322 | 726 | 629 | 536 |
| GAACGTTTCAGCATAAATGG | ASAH1 | 462 | 593 | 1202 | 1225 | 1303 |
| GATTATAGTGAATTCCCTGA | ASAH1 | 547 | 519 | 1048 | 931 | 1039 |
| TCAAGATTTATGGTGTACCA | ASAH1 | 556 | 601 | 1022 | 976 | 989 |
| TCTACTACATGGGCGAAACA | ASAH1 | 617 | 525 | 870 | 785 | 827 |
| AAAACTGGCCAGAATGCCCG | ASAH2 | 544 | 711 | 1197 | 1086 | 1093 |
| ACTCTTCGGGATCAACTCCT | ASAH2 | 602 | 738 | 1282 | 1361 | 1550 |
| ATGTGGGCCAAGAATATTGG | ASAH2 | 553 | 800 | 1243 | 1253 | 1244 |
| CTTCATCAAGAAAGGAACCG | ASAH2 | 485 | 682 | 965 | 1020 | 1003 |
| GGTGAGTGTTGAGCCTTGAG | ASAH2 | 279 | 1242 | 1818 | 1886 | 1852 |
| GTTAACCGGACCGCTTGCTG | ASAH2 | 618 | 577 | 1144 | 1002 | 1024 |
| AAACGGCGCCTCAACTTCAG | ASCL1 | 645 | 642 | 1182 | 1096 | 1160 |
| CCAGGTTGACAAGCTTGACG | ASCL1 | 528 | 1029 | 1775 | 1826 | 1786 |
| GCAGCAGCAGCCGGCCGCCG | ASCL1 | 442 | 523 | 950 | 957 | 1044 |
| GTGGCGAAGAAGCAGGCTGC | ASCL1 | 518 | 641 | 861 | 686 | 758 |
| TCTTGTTGGCCGCGCCGTTG | ASCL1 | 485 | 946 | 1398 | 1571 | 1520 |
| TGACTTGTGACCGCCCCCTG | ASCL1 | 477 | 685 | 1424 | 1248 | 1198 |
| ACAAGAAGACTTGACAACTG | ASXL1 | 603 | 578 | 1030 | 1033 | 1097 |
| ACCTGACGCCGATTCCACGT | ASXL1 | 619 | 318 | 616 | 584 | 582 |
| CTAACATCTAAATGCAACGG | ASXL1 | 399 | 576 | 1065 | 1135 | 1178 |
| GAGAATTTATCAGGGCCCGA | ASXL1 | 506 | 286 | 641 | 625 | 705 |
| GAGTCATTGCATCAGAACGT | ASXL1 | 572 | 493 | 907 | 901 | 960 |
| GGCTGACCTTTAACCACCCA | ASXL1 | 539 | 592 | 903 | 940 | 863 |
| AGTAACCCTATTGAGAGCAG | ATIC | 511 | 724 | 1276 | 1276 | 1276 |
| GAATATCTCAAATGCCCTTG | ATIC | 626 | 693 | 1386 | 1394 | 1428 |
| GCACAAGTTTATAAATCCAG | ATIC | 503 | 396 | 545 | 531 | 529 |
| GCACCAATCACTTACCAGCA | ATIC | 607 | 566 | 965 | 960 | 951 |
| GGCATCCCGGAGAGCTGTCG | ATIC | 511 | 515 | 1127 | 1066 | 1041 |
| TGCGATAAAGTCACCAAATG | ATIC | 492 | 996 | 1433 | 1420 | 1311 |
| GCTGTAGATAGATCAGCATG | ATM | 509 | 993 | 1840 | 1632 | 1622 |
| GCTTAACAGGTCACAACACA | ATM | 545 | 472 | 992 | 801 | 806 |
| GTTTAAAAACAAGACACATG | ATM | 320 | 279 | 535 | 439 | 420 |
| TCTACCTCAACGACGACATG | ATM | 598 | 797 | 1339 | 1511 | 1526 |
| TGGCTGTCGTGGACTACATG | ATM | 324 | 921 | 1524 | 1634 | 1597 |
| TTGAGTCCAAATATAAAGCA | ATM | 533 | 456 | 693 | 673 | 704 |
| AAAAATGAGCATCAACGCAG | ATP1A1 | 419 | 291 | 463 | 378 | 419 |
| AATGACAACATACTCGACTC | ATP1A1 | 555 | 347 | 535 | 547 | 467 |
| AGATCGATATGCCAAGATTG | ATP1A1 | 589 | 953 | 1825 | 1849 | 1843 |
| CCTCAAGCCAAGTATACTCA | ATP1A1 | 717 | 605 | 1080 | 1048 | 1148 |
| CTACCTTAATTCCAGCACTT | ATP1A1 | 538 | 297 | 544 | 474 | 482 |
| GCAATATTCCTCGTTTCCAG | ATP1A1 | 474 | 342 | 581 | 541 | 580 |
| CATGATCGACCCACCCCGGG | ATP1A3 | 543 | 594 | 1075 | 1075 | 1138 |
| GACCTGGTAGAGATTAAGGG | ATP1A3 | 681 | 744 | 1161 | 1180 | 1222 |
| GGTACTCACATTGTCACCCG | ATP1A3 | 580 | 810 | 1519 | 1511 | 1498 |
| GGTCTGCCGGAAATACAACA | ATP1A3 | 601 | 138 | 219 | 183 | 206 |
| GTGCGAGCTCTTGTCAAACG | ATP1A3 | 433 | 597 | 969 | 969 | 1012 |
| TCTGGGCCTTACAGTGACAG | ATP1A3 | 553 | 849 | 1525 | 1448 | 1470 |
| AAAATATCACGTAGAACAGA | ATP1B1 | 508 | 350 | 470 | 461 | 422 |
| CTTGGGATCGTTAGGGCGAA | ATP1B1 | 455 | 883 | 1366 | 1378 | 1403 |
| GAGTGGTTGGGCAATTGCTC | ATP1B1 | 472 | 581 | 1067 | 1049 | 1095 |
| GGAGAATTTAACAATGAACG | ATP1B1 | 569 | 532 | 1094 | 963 | 967 |
| GGTGAGCAGCATCACTTGAA | ATP1B1 | 499 | 797 | 1444 | 1510 | 1452 |
| TTGAGCTTTATTAGGACACA | ATP1B1 | 280 | 754 | 1280 | 1308 | 1298 |
| ACTCCAAGCATGCAGTGGTG | ATP1B3 | 418 | 947 | 1765 | 1758 | 1807 |
| ACTCTTCTCATTCACAATGT | ATP1B3 | 683 | 613 | 1107 | 1196 | 1165 |
| CGTGACCAGATTCCTAGTCC | ATP1B3 | 738 | 540 | 897 | 949 | 1039 |
| GGGCGCACCGCCAAGAGCTG | ATP1B3 | 522 | 659 | 1274 | 1271 | 1356 |
| TAAAACCTCAAGGAATGCCA | ATP1B3 | 496 | 322 | 535 | 545 | 491 |
| TTCATTCACTGTATCTGATC | ATP1B3 | 378 | 571 | 731 | 705 | 708 |
| AAGGAGGAGGAAAAGGGTCA | ATP1B4 | 500 | 805 | 1369 | 1331 | 1303 |
| AAGTTAAGGCTATGGGCGAA | ATP1B4 | 563 | 654 | 1180 | 1151 | 1141 |
| CATGCAGAGGGTAATCACAG | ATP1B4 | 514 | 596 | 1116 | 1208 | 1208 |
| GATCGTGAACGAATACCTGT | ATP1B4 | 529 | 261 | 728 | 611 | 688 |
| GGTACCCGATTTAGGAGGGG | ATP1B4 | 475 | 896 | 1498 | 1198 | 1261 |
| TGAATGTAGATTGTCCCCCG | ATP1B4 | 319 | 683 | 1376 | 1258 | 1240 |
| AACAGAAGATCACCGCCAGA | ATP6V0A1 | 593 | 393 | 861 | 730 | 799 |
| CAGTTGAACCCATCTGTCCG | ATP6V0A1 | 340 | 737 | 1220 | 1271 | 1336 |
| GATGGCGGATCCAGACTTGT | ATP6V0A1 | 542 | 571 | 973 | 841 | 835 |
| TCTAAGTCAATCATGTCCCG | ATP6V0A1 | 437 | 545 | 1050 | 985 | 976 |
| TGAAATCGAGAATCCCCTGG | ATP6V0A1 | 527 | 768 | 1278 | 1329 | 1351 |
| TGGAGTCGAGGTCAGTGACA | ATP6V0A1 | 568 | 715 | 1339 | 1303 | 1325 |
| AAGACCACCGAAACAGACTG | ATR | 827 | 618 | 1153 | 1205 | 1227 |
| AAGATGCAAACTCTCAAGCT | ATR | 634 | 360 | 673 | 670 | 649 |
| ATGAAACAGAAATTGAACTG | ATR | 518 | 316 | 761 | 788 | 809 |
| GTGTACAGACAAGTTGACCA | ATR | 376 | 335 | 735 | 603 | 611 |
| TCAGTACTCCAAAAATTGCA | ATR | 520 | 356 | 441 | 412 | 387 |
| TTGTATTCTAGCATCTACGG | ATR | 526 | 619 | 1242 | 1176 | 1280 |
| AAAAAGCATGAGTAAAATTG | ATRX | 687 | 698 | 1029 | 987 | 942 |
| AAGGGTGTGGTCTTTACACG | ATRX | 724 | 870 | 1631 | 1602 | 1549 |
| AGAAGCCAAGATACAGACAT | ATRX | 490 | 500 | 1039 | 981 | 1060 |
| AGAGCAAATAAAAGTGTCAG | ATRX | 501 | 609 | 1204 | 1239 | 1239 |
| GACAAATATAGCAATACCAC | ATRX | 459 | 251 | 366 | 326 | 291 |
| GGCGAAAACGTATTGCTGAG | ATRX | 488 | 681 | 1298 | 1220 | 1290 |
| AATCCATTATCTGCAAATAG | AURKA | 557 | 373 | 457 | 371 | 438 |
| CTCCAGCTGATCCAAGGAGT | AURKA | 491 | 389 | 966 | 768 | 745 |
| GTGACCTAGAGGCAGGACGA | AURKA | 578 | 502 | 916 | 710 | 704 |
| TCAAGAAGTTGATTTACCAG | AURKA | 546 | 568 | 1253 | 911 | 1016 |
| TCAATCTAGAAACTTACCTA | AURKA | 644 | 267 | 475 | 352 | 365 |
| TTAATAGTATATCACAGAGT | AURKA | 565 | 389 | 712 | 602 | 676 |
| ACCACATCCCAGTTACCTGT | AURKB | 582 | 301 | 536 | 379 | 424 |
| ACTATTTCTACGACCGGAGA | AURKB | 456 | 448 | 862 | 669 | 689 |
| CAAGTCTCAGATAGAAAAGG | AURKB | 650 | 351 | 718 | 476 | 590 |
| CCCAATCTCAAAGTCGTCGA | AURKB | 635 | 352 | 663 | 556 | 572 |
| CGACGAGCAGCGAACAGCCA | AURKB | 410 | 478 | 978 | 966 | 925 |
| GCCCCAGAGAGTCCTCCGGA | AURKB | 563 | 446 | 824 | 634 | 707 |
| AATCCAGGCGCATCTACAGT | AURKC | 480 | 473 | 791 | 782 | 744 |
| AGATGAACAGCACACAGCCA | AURKC | 419 | 667 | 952 | 914 | 981 |
| ATTCTGGAATATGCTCCAAG | AURKC | 497 | 359 | 628 | 685 | 688 |
| CAAGGAAAGCCACTTCATTG | AURKC | 504 | 1014 | 1484 | 1472 | 1566 |
| GCACACACCATCTCTGAGGT | AURKC | 575 | 538 | 984 | 891 | 885 |
| GCGGGACGCTGAGCTCCACG | AURKC | 528 | 954 | 1458 | 1617 | 1697 |
| AATGCCAAAGGAAATCCGTG | AXIN1 | 470 | 623 | 911 | 993 | 1096 |
| ATATTTATTTGGAATACACG | AXIN1 | 350 | 476 | 744 | 738 | 697 |
| GCTGTACCGTCTACTTGAGG | AXIN1 | 759 | 767 | 1560 | 1364 | 1432 |
| GGAGCCCTGTGACTCGAACG | AXIN1 | 371 | 727 | 1192 | 1218 | 1247 |
| GTAAGCACGCACCCAAGTCA | AXIN1 | 535 | 646 | 1275 | 1102 | 1131 |
| TCGGCCCACTTCAAGTATGG | AXIN1 | 365 | 623 | 1405 | 1361 | 1373 |
| CACTCCTTACCACATCCGGG | AXL | 399 | 568 | 996 | 1067 | 1049 |
| CCCAGCAATACATACCACCA | AXL | 487 | 640 | 671 | 661 | 693 |
| CGGGTGGTACCAATGTGCCG | AXL | 527 | 747 | 1272 | 1125 | 1218 |
| GGGTACCGGCTGGCCTACCG | AXL | 294 | 762 | 1230 | 1124 | 1312 |
| GTATGAGGTTCACTTACAGG | AXL | 526 | 666 | 978 | 858 | 863 |
| TGAAGCCCATAATGCCAAGG | AXL | 345 | 594 | 1108 | 1090 | 1121 |
| AAATCAATTTCAATCTCTGG | B2M | 607 | 308 | 569 | 531 | 512 |
| ACCCTGACACGTAGCAGTTC | B2M | 612 | 527 | 1077 | 1027 | 995 |
| CATCCAGGCGGCACGCAGCG | B2M | 527 | 867 | 1575 | 1467 | 1540 |
| GAAACCAGCCGTCGCCAGCG | B2M | 541 | 732 | 1047 | 1224 | 1203 |
| GAGTACACTTGAATCTTTGG | B2M | 730 | 453 | 838 | 851 | 871 |
| TCACGACACCCAGCAGAGAA | B2M | 650 | 538 | 899 | 877 | 958 |
| CACGGACGTATCATCCACCA | BAP1 | 372 | 791 | 1417 | 1559 | 1596 |
| CGCATGAAGGACTTCACCAA | BAP1 | 668 | 634 | 1138 | 1167 | 1071 |
| GCCGCTGAACGATGGGAGTG | BAP1 | 443 | 783 | 1525 | 1905 | 1830 |
| GCCTGCATCTAGCGCCAGGG | BAP1 | 496 | 672 | 1469 | 1547 | 1534 |
| GGTCCCTACCATGGTCAATG | BAP1 | 486 | 831 | 1877 | 1614 | 1726 |
| GTCCATCAAGACAAGCAGCG | BAP1 | 487 | 436 | 852 | 905 | 931 |
| ACAGTAACATGGAGTTGCAG | BAX | 448 | 562 | 1256 | 1383 | 1325 |
| ACTCGGAAGAAGACCTCACG | BAX | 470 | 685 | 1484 | 1622 | 1539 |
| AGCGAATGTCTCAAGCGCAT | BAX | 481 | 407 | 540 | 517 | 564 |
| GTTTCATCCAAGATCGAGCA | BAX | 520 | 407 | 655 | 625 | 581 |
| TGAGCAGATCATGAAGACAG | BAX | 577 | 547 | 1091 | 967 | 944 |
| TTCTTGGTGGATGCATCCTG | BAX | 371 | 687 | 1051 | 1013 | 1069 |
| AGGCGGAGATGCCCGGCGCG | BCL2 | 561 | 787 | 1392 | 1346 | 1362 |
| AGGGTCAGGTGGACCACAGG | BCL2 | 685 | 829 | 1386 | 1220 | 1200 |
| CGTGGCAAAGCGTCCCCTCG | BCL2 | 651 | 814 | 1512 | 1420 | 1490 |
| GACCCCACCGAACTCAAAGA | BCL2 | 525 | 287 | 500 | 385 | 361 |
| TCCACTACAAGCTGTCGCAG | BCL2 | 475 | 576 | 836 | 840 | 813 |
| TCGGCGAAGTACGCCGGTAG | BCL2 | 646 | 773 | 1342 | 1284 | 1348 |
| ACGTCAAAACTGTCCAAGCA | BCL2A1 | 508 | 458 | 891 | 890 | 872 |
| CTGCGAGTTTGGCTACACGC | BCL2A1 | 597 | 362 | 659 | 667 | 611 |
| GCTCGGGATGTGACTCAGGG | BCL2A1 | 481 | 914 | 1579 | 1572 | 1640 |
| GGGCCTCAGGCAGCTCACGG | BCL2A1 | 350 | 507 | 909 | 877 | 898 |
| TGATGCCCAACGATGTTCTG | BCL2A1 | 481 | 729 | 1322 | 1308 | 1341 |
| TGCTATGTCCACTCACCCAC | BCL2A1 | 401 | 361 | 624 | 758 | 740 |
| AGTTTGAACTGAGGTACCGG | BCL2L1 | 438 | 588 | 1180 | 1160 | 1151 |
| CAGCGGTGAAACAAGCGCTG | BCL2L1 | 533 | 582 | 842 | 878 | 875 |
| CTGGAGTCAGTTTAGTGATG | BCL2L1 | 447 | 903 | 1404 | 1165 | 1185 |
| GCTCTGATATGCTGTCCCTG | BCL2L1 | 586 | 588 | 1270 | 1307 | 1240 |
| GTTGCCATTGATGGCACTGG | BCL2L1 | 381 | 651 | 1284 | 1114 | 1103 |
| TTCCGGGATGGGGTGAACTG | BCL2L1 | 356 | 614 | 1038 | 803 | 782 |
| CCCAGGTGCCCAGCACCCGG | BCL2L10 | 439 | 850 | 1468 | 1399 | 1402 |
| CTGAAAGGAATGCCCCAACC | BCL2L10 | 572 | 342 | 574 | 506 | 538 |
| GCCTGGCTGGAGGCGCACGA | BCL2L10 | 556 | 556 | 1092 | 1056 | 1064 |
| GGCGAGTGGGAGGCCAGGGT | BCL2L10 | 529 | 350 | 690 | 595 | 610 |
| GGGGAGAGCCAGACTTCCAA | BCL2L10 | 573 | 524 | 1016 | 964 | 966 |
| TAATAAGAGCCTTCTAGCAA | BCL2L10 | 749 | 458 | 923 | 820 | 800 |
| ACGCCACATGCAAATCAGAT | BCL2L12 | 584 | 457 | 655 | 503 | 537 |
| CAGGCTCTGAAACATAGCAT | BCL2L12 | 423 | 562 | 1351 | 1301 | 1278 |
| CCCTAACCCCCAGAGTTACA | BCL2L12 | 551 | 375 | 745 | 820 | 768 |
| CTAGCTGCCTTCTTTAGGCG | BCL2L12 | 415 | 638 | 1460 | 1400 | 1413 |
| GCTCAAGTTCGGTGAAGGAG | BCL2L12 | 543 | 642 | 1218 | 1257 | 1332 |
| GGCTGAAGTCATCAACCAGA | BCL2L12 | 505 | 395 | 553 | 521 | 599 |
| ACTTTGTAGGCTATAAGCTG | BCL2L2 | 512 | 869 | 1641 | 1371 | 1453 |
| CCAAATCAGAGAAGGTGCGC | BCL2L2 | 439 | 423 | 811 | 535 | 596 |
| GAAGAGTTCGTCAGAGACCT | BCL2L2 | 464 | 716 | 958 | 943 | 985 |
| GGCCACAAGACGGCCCCAGT | BCL2L2 | 405 | 388 | 768 | 893 | 916 |
| TCCACTGCACCAAGCCATGC | BCL2L2 | 468 | 494 | 621 | 695 | 659 |
| TTGTGGGACAAGTGCAAGAG | BCL2L2 | 629 | 637 | 1373 | 1083 | 1171 |
| ACACACAGTGACATGCACTA | BCL6 | 585 | 461 | 703 | 650 | 625 |
| ACCAGACCCTTGCGGTTCAG | BCL6 | 635 | 834 | 1378 | 1396 | 1499 |
| CAACCAGAATGCCAAACCGG | BCL6 | 555 | 506 | 1000 | 991 | 1049 |
| CAAGACATCATGGCCTATCG | BCL6 | 556 | 574 | 1408 | 1262 | 1244 |
| CCAACCCCTTCCCTAAGGAG | BCL6 | 437 | 675 | 1083 | 1227 | 1224 |
| GTCTCGGCTCAATCTCCGGG | BCL6 | 456 | 502 | 1162 | 1134 | 1093 |
| ACCACCTAGAGTGCTCGTGG | BCOR | 665 | 672 | 1259 | 1396 | 1429 |
| AGGGCAACGAAATCGCGGTG | BCOR | 764 | 1107 | 2050 | 1877 | 1862 |
| CCCAACAACCAAAGACTCCG | BCOR | 534 | 389 | 625 | 669 | 677 |
| CTGGGTCAATCCTTACATGG | BCOR | 492 | 667 | 1278 | 1250 | 1341 |
| GGCGTGTGAGGTATCAACAT | BCOR | 508 | 582 | 1188 | 1147 | 1191 |
| TAGATCTGAGATCATCAGCA | BCOR | 509 | 551 | 1059 | 1209 | 1146 |
| AGGAATGGTGCTGATCAGTG | BCORL1 | 506 | 1078 | 2212 | 2162 | 2143 |
| CAGAGATTCGGTCACCCCAT | BCORL1 | 484 | 587 | 1170 | 1369 | 1456 |
| GGACGGGAACTAGCCCAGAG | BCORL1 | 398 | 868 | 1626 | 1588 | 1673 |
| GGGGGCCGGAGTAAAGATTG | BCORL1 | 406 | 1006 | 1310 | 1296 | 1343 |
| TGGGTAAGATCGTAGCTCTG | BCORL1 | 389 | 1074 | 1967 | 1952 | 1928 |
| TTCCCATTAACTGAACACCG | BCORL1 | 450 | 342 | 462 | 491 | 469 |
| CGTCGTTTGGAACGCCACCT | BCR | 496 | 640 | 1250 | 1067 | 1077 |
| CTCAAAGTTGGACCTGAGCG | BCR | 552 | 752 | 1362 | 1381 | 1571 |
| CTTCCTGTCCAGTATCAATG | BCR | 651 | 727 | 1115 | 1093 | 1198 |
| TACCAGAGCATCTACGTCGG | BCR | 487 | 722 | 1470 | 1362 | 1411 |
| TCCCAACAGAGGGCAAACAA | BCR | 417 | 339 | 626 | 625 | 609 |
| TTTGTCTCGAAACATGCGGT | BCR | 496 | 494 | 1014 | 979 | 917 |
| ATGACGCGATGTCCGAACAC | BIRC2 | 537 | 444 | 899 | 662 | 776 |
| CTGTAGGGGTTAGTGCTCAG | BIRC2 | 466 | 814 | 1289 | 1316 | 1263 |
| CTGTGAATAAAACTACAGCT | BIRC2 | 541 | 380 | 969 | 823 | 786 |
| GGCCATCTAGTGTCCCCGTG | BIRC2 | 527 | 983 | 1256 | 1082 | 1235 |
| GTCTGGAGATGACCCATGGG | BIRC2 | 521 | 727 | 1638 | 1368 | 1441 |
| TACATACTCCACATTCCCGG | BIRC2 | 430 | 297 | 506 | 429 | 423 |
| ACATACTCGACATTCCCCGC | BIRC3 | 407 | 327 | 401 | 439 | 434 |
| CCTGATGCTAGACAACTGGA | BIRC3 | 622 | 606 | 958 | 865 | 900 |
| CTTCCACAAATGGGCAGTCG | BIRC3 | 431 | 745 | 1416 | 1188 | 1273 |
| GGGCCCCACATAGTAAAAGC | BIRC3 | 492 | 236 | 480 | 457 | 452 |
| TCGGGGTGGCCAGTTACAGA | BIRC3 | 691 | 752 | 1049 | 1011 | 1039 |
| TGAAGTCGACAGGGTCCACA | BIRC3 | 590 | 675 | 1315 | 1304 | 1369 |
| CAAGAACTGGCCGTTCCTGG | BIRC5 | 390 | 527 | 1056 | 926 | 904 |
| CAAGTCTGGCTCGTTCTCAG | BIRC5 | 518 | 415 | 975 | 792 | 755 |
| CGCCTGGCAGCTCTACCTCA | BIRC5 | 511 | 614 | 1262 | 1117 | 1133 |
| CTTGAAGCAGAAGAAACACT | BIRC5 | 379 | 383 | 636 | 579 | 578 |
| GCGCGGTCTCACCCGGTCCG | BIRC5 | 446 | 411 | 1112 | 798 | 828 |
| GGAACGGGACGACTTACATA | BIRC5 | 379 | 532 | 1060 | 806 | 921 |
| CTGTACCCTGGGATTAGCTG | BLM | 391 | 829 | 1662 | 1430 | 1432 |
| GAGACCAGCACAAACTGTGA | BLM | 460 | 756 | 1098 | 1189 | 1212 |
| GAGTCAATTCAGAATTACAC | BLM | 456 | 459 | 686 | 662 | 630 |
| GTATAAGCTCTTCAGGATTG | BLM | 651 | 1047 | 1842 | 1765 | 1705 |
| TCCTTAACAGAATTCCCGGT | BLM | 697 | 251 | 562 | 486 | 491 |
| TCTACTATAAGTGATCTCAA | BLM | 516 | 331 | 596 | 532 | 575 |
| AACCTACCTATAACAACAGG | BMPR1A | 567 | 391 | 712 | 619 | 631 |
| AAGGCCGGTACGGAGAAGTG | BMPR1A | 608 | 721 | 1340 | 1299 | 1260 |
| AGCTCTGTGACTGGTCAATG | BMPR1A | 418 | 824 | 1547 | 1460 | 1531 |
| GCCACGAGAACATTCTTGGT | BMPR1A | 661 | 526 | 1001 | 947 | 1015 |
| TGAATAAGCCAACTTGAGCA | BMPR1A | 502 | 538 | 935 | 824 | 859 |
| TGGAGTAACCTTAGCACCAG | BMPR1A | 571 | 664 | 1196 | 1235 | 1294 |
| ATTTCGCGGGCAGTAACCAG | BMX | 421 | 629 | 1296 | 1208 | 1255 |
| CATCCACGAAGAACCCACTG | BMX | 423 | 600 | 1117 | 1223 | 1079 |
| CTTTGAGTCGCTGTCATACG | BMX | 594 | 749 | 1302 | 1259 | 1355 |
| GTCCGAAATTCCCGCCAGGT | BMX | 499 | 388 | 823 | 759 | 734 |
| TGTGGAGAAAGTAAATCTCG | BMX | 376 | 927 | 1476 | 1353 | 1506 |
| TTGATGGTAGTGAATAAGCT | BMX | 521 | 576 | 1118 | 1101 | 1216 |
| AGAGAAGAAACCGATTGGCT | BRAF | 412 | 572 | 951 | 938 | 1022 |
| AGGACTTGATTAGAGACCAA | BRAF | 393 | 460 | 967 | 895 | 900 |
| ATGATTGGGAGATACCTGAT | BRAF | 403 | 564 | 1015 | 1040 | 994 |
| ATGTTTGACAGCGAAAACCC | BRAF | 568 | 331 | 630 | 608 | 623 |
| TGAACACCACCCAATACCAC | BRAF | 381 | 273 | 393 | 409 | 370 |
| TGGGTAACAATAGCCAGTTG | BRAF | 426 | 760 | 1523 | 1422 | 1466 |
| AAGGAGCCAACAGAGCAGAT | BRCA1 | 644 | 399 | 741 | 758 | 760 |
| ATGTAGACAGACTTATTCTG | BRCA1 | 580 | 584 | 1330 | 1148 | 1049 |
| CAGGCAGTTCATACAAATGC | BRCA1 | 376 | 358 | 695 | 587 | 607 |
| GAAGACCGACTACTTCCCAT | BRCA1 | 576 | 327 | 789 | 765 | 785 |
| TATTTGGAAAAACCTATCGG | BRCA1 | 507 | 382 | 931 | 827 | 915 |
| TGCAACAATTGAGAACCCCA | BRCA1 | 511 | 280 | 496 | 442 | 462 |
| AGCAGTTAACAGCTAATACG | BRCA2 | 644 | 506 | 906 | 716 | 721 |
| ATGTACATCAACTTTCTGAG | BRCA2 | 641 | 488 | 913 | 778 | 779 |
| CAGTTGCTAGTCGGAAGCAA | BRCA2 | 469 | 804 | 1310 | 1307 | 1294 |
| GATATGATATTAATTGACTG | BRCA2 | 575 | 635 | 966 | 1027 | 969 |
| TTCCTCTAGACATCACAACT | BRCA2 | 485 | 418 | 775 | 514 | 518 |
| TTTGTGTAAGAGACTGACTT | BRCA2 | 597 | 332 | 678 | 589 | 651 |
| AGTGGCATCAATGCCACAAG | BRD2 | 519 | 1244 | 1813 | 1690 | 1761 |
| CCAGCTTCAATACCTGCACA | BRD2 | 462 | 539 | 889 | 804 | 863 |
| CTGGCCTTTCTATAAACCAG | BRD2 | 585 | 669 | 1140 | 976 | 1100 |
| GAGTGTAAGGACTTGAGAAG | BRD2 | 531 | 519 | 1187 | 1001 | 1058 |
| GCCACTCTCTCTACGCATAG | BRD2 | 454 | 572 | 953 | 950 | 924 |
| GTGCCATGGCCACAACATCG | BRD2 | 517 | 736 | 1286 | 1172 | 1312 |
| ACACTCACCTTGACGACAGG | BRD3 | 500 | 805 | 1123 | 1228 | 1196 |
| CGCGAGTACCCAGACGCACA | BRD3 | 15 | 498 | 1144 | 1001 | 1051 |
| GAGCGCGAGCGAATGTATGC | BRD3 | 486 | 736 | 1127 | 1078 | 1062 |
| GTCCGAGCTGCCTGAATCCG | BRD3 | 450 | 777 | 1029 | 1102 | 1107 |
| TCAAGGTACGCAGCAAGTGG | BRD3 | 335 | 603 | 1231 | 1126 | 1174 |
| TCTTGGACAGCATTTCCTTG | BRD3 | 567 | 836 | 1765 | 1689 | 1727 |
| CCAGACCCCCGTCATGACAG | BRD4 | 586 | 518 | 1082 | 1149 | 1032 |
| CTGAAGAGGAACGAGCCCAG | BRD4 | 378 | 715 | 1356 | 1229 | 1261 |
| GTACAACCCTCCCGACCATG | BRD4 | 544 | 561 | 1305 | 1071 | 1126 |
| GTGCAAGCCCATGTCATACG | BRD4 | 545 | 480 | 865 | 810 | 822 |
| TCATCTCAACCAGCACTCGG | BRD4 | 493 | 369 | 544 | 505 | 455 |
| TGCCCAAGTTACTCACAGGG | BRD4 | 249 | 644 | 1578 | 1317 | 1197 |
| AACTCCCTGGAGATAAACTT | BRDT | 520 | 338 | 697 | 803 | 716 |
| AATTGTATTTAAATCCATTG | BRDT | 591 | 678 | 1152 | 994 | 1085 |
| CAACTCTACCTCACAAACAG | BRDT | 529 | 298 | 504 | 443 | 404 |
| CAGACAGGCATACTCTCGAT | BRDT | 472 | 543 | 1122 | 886 | 950 |
| GAAAATAACAAAGTCCAAGG | BRDT | 596 | 264 | 448 | 454 | 537 |
| TGACATTGTGAAAAATCCAA | BRDT | 402 | 366 | 560 | 441 | 468 |
| CTATAGCACACAGCTCGTAA | BRIP1 | 536 | 378 | 492 | 454 | 490 |
| GACACTTCATCAACTTGTCA | BRIP1 | 658 | 531 | 681 | 684 | 605 |
| GTGTGTCAGACTGTGAACCA | BRIP1 | 520 | 570 | 1028 | 1126 | 1077 |
| TCCGGAGGACAGCGTACTCA | BRIP1 | 495 | 828 | 1561 | 1337 | 1412 |
| TCTTGGTAGCAGTTTGTCGT | BRIP1 | 399 | 530 | 1005 | 1135 | 1124 |
| TTGGTGCTTAAATCCAGCTG | BRIP1 | 386 | 761 | 1350 | 1329 | 1393 |
| AATGGCGTATCACCCCTTGA | BTK | 565 | 696 | 1335 | 1387 | 1316 |
| ATGAGTATGACTTTGAACGT | BTK | 468 | 483 | 768 | 901 | 878 |
| ATGGAGAGGCCAATATGATG | BTK | 399 | 677 | 1458 | 1310 | 1266 |
| GAGAAAATACCTTACCTGGA | BTK | 415 | 380 | 775 | 620 | 608 |
| TGCCCATCGATCCAGAAGCA | BTK | 504 | 697 | 1168 | 1079 | 1159 |
| TGGAGGAGAGCAACTTACCC | BTK | 365 | 529 | 900 | 928 | 931 |
| ACGAGAATCCAAAGTCACAT | BUB1 | 420 | 317 | 492 | 488 | 482 |
| AGGTTAATCCAGCATCTATG | BUB1 | 592 | 798 | 1430 | 1189 | 1238 |
| CAGCCTAAAAATAAACCCGC | BUB1 | 600 | 199 | 343 | 287 | 268 |
| CATGAGTTCAAGCCACAGAG | BUB1 | 588 | 322 | 598 | 477 | 541 |
| GCAGGTTGTTATGTATTGCA | BUB1 | 669 | 638 | 1249 | 1109 | 1044 |
| TTGGCAAATGCGTTTCGGTG | BUB1 | 542 | 850 | 1617 | 1635 | 1628 |
| AGAAAGCATCAACTCCAATG | BUB1B | 512 | 499 | 937 | 961 | 919 |
| AGCTGAACTACAAATTGCCT | BUB1B | 722 | 507 | 771 | 697 | 635 |
| GAAGAAATTAAAAGCGCGAA | BUB1B | 405 | 413 | 613 | 612 | 629 |
| GCTCCAATCAGCCGTGTAGG | BUB1B | 289 | 985 | 1598 | 1414 | 1396 |
| TCAACATATGGAGTAAAACT | BUB1B | 606 | 271 | 395 | 259 | 358 |
| TGTGACTGTAGTCGCTCCAG | BUB1B | 522 | 465 | 1232 | 994 | 928 |
| CATGGTCCTCATAGTTTACG | CA1 | 568 | 719 | 1383 | 1335 | 1311 |
| CCTAAACGCAGTTAAAACCA | CA1 | 393 | 377 | 694 | 682 | 724 |
| GACACTGATAGGCTTAAGAG | CA1 | 596 | 404 | 606 | 651 | 642 |
| GTCCTCTTTCTGCAAGCTAT | CA1 | 425 | 476 | 585 | 586 | 605 |
| TGGAGTCAAATATTCTGGAG | CA1 | 494 | 576 | 1121 | 1208 | 1096 |
| TTGATAGGTCCTGAACAATG | CA1 | 526 | 752 | 1376 | 1296 | 1319 |
| CAGTTTATCCTGACCAACGA | CA12 | 495 | 406 | 626 | 508 | 638 |
| CGCCGCTAGTAACAAGTCAG | CA12 | 551 | 671 | 1213 | 1164 | 1191 |
| GAAGAGTACTACCGCTACAA | CA12 | 352 | 478 | 961 | 899 | 898 |
| GAGAAGTACCCGTCATGTGG | CA12 | 372 | 457 | 1086 | 947 | 954 |
| GGGAACCAGAATGACCCTCA | CA12 | 586 | 595 | 1098 | 1195 | 1101 |
| TGTTCTTTGCAGGATCCAAG | CA12 | 533 | 390 | 822 | 839 | 870 |
| GAAGGTGAAACTGTCGCAAG | CA13 | 468 | 637 | 1043 | 1055 | 978 |
| GAGCTGCCTCAACAAAACTG | CA13 | 485 | 526 | 873 | 804 | 829 |
| GCATCATACTTGATACTAAG | CA13 | 453 | 254 | 463 | 410 | 439 |
| GGTGGATGGAGTGAGATACG | CA13 | 646 | 956 | 1659 | 1614 | 1539 |
| GTCATCAAAGTCAACGCTGA | CA13 | 566 | 501 | 738 | 785 | 795 |
| GTTGATCCCCATCAGCAATA | CA13 | 511 | 429 | 780 | 778 | 772 |
| CAGAATCATAATGTACAATG | CA14 | 583 | 571 | 672 | 667 | 686 |
| CCCTCCAGATACATGGTAGG | CA14 | 526 | 807 | 1479 | 1574 | 1530 |
| CGGCAGCACTGGACGTACGT | CA14 | 519 | 450 | 730 | 751 | 756 |
| CTCCAAGGATACGACCAGCC | CA14 | 499 | 299 | 682 | 643 | 597 |
| CTTTATGCCTGATTTCATGG | CA14 | 338 | 669 | 1322 | 1261 | 1281 |
| GGGTGAGACAGGATCCCTGA | CA14 | 476 | 728 | 1356 | 1424 | 1376 |
| AGGACCCGAGCACTGGCATA | CA2 | 374 | 844 | 1249 | 1537 | 1513 |
| CAACGGTCACTCTTTCAACG | CA2 | 611 | 718 | 1003 | 989 | 946 |
| GAAGTCAGCACTCTTACCCT | CA2 | 511 | 475 | 955 | 929 | 952 |
| GTGCCCTTGCTACGACCAAG | CA2 | 487 | 604 | 1226 | 1314 | 1244 |
| TCACTGGAACACCAAATATG | CA2 | 480 | 595 | 979 | 1024 | 1040 |
| TTTGGTGTCGATGTCAACGG | CA2 | 396 | 683 | 1281 | 1335 | 1313 |
| AAGCTCTCAAGCAACCTGAC | CA3 | 569 | 379 | 488 | 498 | 534 |
| ATCATCAAACACAACCCTGC | CA3 | 602 | 316 | 382 | 353 | 353 |
| ATTGGACAAAGTTAAGACAA | CA3 | 481 | 663 | 1034 | 949 | 1023 |
| GGATCATAAGAGGCTGACCA | CA3 | 550 | 391 | 808 | 822 | 746 |
| TCTTTAGTATGCAGTTCGAT | CA3 | 466 | 692 | 993 | 992 | 955 |
| TTACAGATAGGACGTGAGAA | CA3 | 642 | 781 | 1328 | 1349 | 1273 |
| CAGTGGAGAGTCCAAAACAA | CA4 | 398 | 568 | 969 | 951 | 860 |
| CATCAACATTGACACTACAG | CA4 | 572 | 13 | 17 | 10 | 10 |
| GAAGACAGTGTGTCCACCAG | CA4 | 520 | 617 | 1025 | 1045 | 1062 |
| GTGCTATGAGATTCAAGCCA | CA4 | 444 | 423 | 728 | 728 | 774 |
| TAGTACACGAGAAAGAGAAG | CA4 | 644 | 723 | 1247 | 1152 | 1107 |
| TGGTCGGAGGAGATGGACGG | CA4 | 470 | 718 | 1268 | 1270 | 1238 |
| ATGATCAACAACGGCCACAC | CA6 | 403 | 483 | 748 | 869 | 867 |
| CATCTGTTGAGCTATGTACT | CA6 | 330 | 515 | 1047 | 1070 | 1132 |
| CTTACCTGAGTAGGTCCACA | CA6 | 657 | 984 | 1516 | 1401 | 1428 |
| GGACAAAGCACAGTTCTGAG | CA6 | 291 | 701 | 1237 | 1240 | 1246 |
| GTTTCAGCCTACCTTAACAA | CA6 | 629 | 267 | 452 | 395 | 405 |
| TAGTGAACGATGTGAATCTG | CA6 | 506 | 1079 | 1605 | 1503 | 1582 |
| ATACAGCTTGTGCCACTGTG | CA7 | 534 | 992 | 1777 | 1897 | 1857 |
| CAATATTGATTGGTGACTGG | CA7 | 393 | 779 | 1478 | 1391 | 1364 |
| CTCCCCACAGCGGTAACTGG | CA7 | 746 | 602 | 1073 | 1121 | 1118 |
| TCTTGGCGTTCCAATGAACA | CA7 | 490 | 628 | 1135 | 1143 | 1269 |
| TGGGGCAAGAAGCACAGTGT | CA7 | 507 | 685 | 1130 | 1127 | 1111 |
| TGTAGAGCGCATCTGTCAGA | CA7 | 428 | 699 | 1343 | 1445 | 1299 |
| AGAGGATCTACCTACTGTTG | CA9 | 615 | 1108 | 1938 | 1610 | 1698 |
| AGAGGATGATCAATTGGATG | CA9 | 458 | 1047 | 1817 | 1674 | 1706 |
| CAGTGCTGATGTGAACCACG | CA9 | 312 | 835 | 1737 | 1586 | 1496 |
| CTCCATAGCGCCAATGACTG | CA9 | 611 | 906 | 1367 | 1459 | 1454 |
| CTGCAGTTGCACCTGCACTG | CA9 | 444 | 896 | 1516 | 1466 | 1535 |
| CTGCGCAACAATGGCCACAC | CA9 | 507 | 551 | 852 | 819 | 834 |
| ATTGCCAGATCCTGACCGAA | CADM2 | 474 | 501 | 801 | 729 | 845 |
| CAGAATGTAACGGTTGTTGA | CADM2 | 477 | 829 | 1472 | 1408 | 1392 |
| CAGCACACTGGATTTCCGAG | CADM2 | 353 | 683 | 1203 | 1261 | 1197 |
| CTTACCCAGAACAGTGAGAT | CADM2 | 519 | 573 | 1042 | 1110 | 1080 |
| GATGGTATAATCTTGACTGA | CADM2 | 538 | 740 | 1140 | 1068 | 1121 |
| GCTGATACTCAATTCATGCC | CADM2 | 467 | 508 | 739 | 775 | 676 |
| ACAGAATGGGCATTGTCCAC | CAMK1D | 452 | 406 | 834 | 775 | 772 |
| AGAGCTATTTGATCGGATAG | CAMK1D | 525 | 1031 | 1569 | 1571 | 1566 |
| ATTCCGAAGAAGGCGCTGAA | CAMK1D | 455 | 489 | 1044 | 1004 | 1094 |
| GGGCCCCACTTACGTTCCGA | CAMK1D | 642 | 622 | 901 | 834 | 918 |
| GTGCTTACCAACGTAGCCCG | CAMK1D | 511 | 661 | 1228 | 1245 | 1288 |
| TGTTGACTGCTGGTCCATCG | CAMK1D | 577 | 677 | 1533 | 1442 | 1392 |
| ATGGTAGAGAATCTCCGGGT | CARD11 | 372 | 437 | 930 | 913 | 843 |
| CCTGGAGCATGACCGCAAGG | CARD11 | 581 | 543 | 852 | 785 | 825 |
| GCATCGGTTGAATAAGATGG | CARD11 | 563 | 855 | 1711 | 1392 | 1421 |
| TGACGACTCATCTACCTCAG | CARD11 | 460 | 706 | 979 | 1013 | 1032 |
| TGCTCGGGGGCAACGCACGA | CARD11 | 651 | 721 | 1063 | 1090 | 1031 |
| TGTGTCTTACAGCACAGTCG | CARD11 | 473 | 719 | 1332 | 1235 | 1236 |
| AGGTGACGTGGTATTGCTGT | CARS | 422 | 743 | 1443 | 1389 | 1314 |
| AGTGTTGCTCGAGTAATCCA | CARS | 577 | 627 | 1065 | 1042 | 992 |
| CCAGCGAGAGACACTCCTAC | CARS | 465 | 502 | 765 | 744 | 770 |
| GCCTCTTCCACGTGGCTCGG | CARS | 433 | 882 | 1548 | 1594 | 1669 |
| TCCGCCATGAATGTCCATCG | CARS | 502 | 492 | 1027 | 920 | 964 |
| TGGCACATATTCACTAACGC | CARS | 404 | 257 | 386 | 335 | 365 |
| AAAATGGATTATCCAGAAAT | CASP3 | 480 | 619 | 963 | 976 | 938 |
| AAAGTAACAGGTTTCTTCCG | CASP3 | 539 | 549 | 690 | 692 | 729 |
| CGAGGTACAGAACTAGACTG | CASP3 | 406 | 539 | 1126 | 1090 | 1062 |
| GGAAGTAAATCAATGGACTC | CASP3 | 405 | 737 | 1253 | 1039 | 1138 |
| TCTACATCTGTACCAGACCG | CASP3 | 551 | 636 | 1497 | 1283 | 1356 |
| TTGCGTGCTTCTAAGCCATG | CASP3 | 498 | 913 | 1710 | 1685 | 1687 |
| ACAGGGATAATCTCGTGCGC | CASP6 | 409 | 520 | 987 | 1156 | 1099 |
| ATCGGCATCTATGTGGCTAG | CASP6 | 612 | 724 | 1288 | 1283 | 1367 |
| CAACAGAGAAATGTTTGACC | CASP6 | 514 | 317 | 654 | 628 | 607 |
| CTTTGATCTTCAATCACGAG | CASP6 | 459 | 386 | 722 | 618 | 592 |
| GATCCACTACATCCAACGGA | CASP6 | 578 | 542 | 925 | 840 | 854 |
| GCTGGAGGACAACGTAACGG | CASP6 | 537 | 635 | 1319 | 1135 | 1100 |
| AGATGACCACATCAAAACCC | CASP7 | 344 | 301 | 519 | 497 | 441 |
| GAAGAGTGATGGGACAAAGG | CASP7 | 450 | 513 | 1051 | 1033 | 1013 |
| GAGATCAATCAAGACTGCCC | CASP7 | 429 | 329 | 491 | 515 | 519 |
| GCCTGCATCTTACTAAGCCA | CASP7 | 546 | 473 | 697 | 797 | 847 |
| TCTATCCCCCCTAAAATGGG | CASP7 | 440 | 926 | 1502 | 1367 | 1382 |
| TTGGTATTTAGGCTTGCCGG | CASP7 | 578 | 757 | 1648 | 1558 | 1580 |
| AGTTAAAAAGAACGGCATTG | CASP8AP2 | 730 | 639 | 1340 | 1204 | 1225 |
| CAAACCTAGGACTTCAACTG | CASP8AP2 | 524 | 345 | 860 | 696 | 731 |
| GAAATCCGTAGTAAAAGACC | CASP8AP2 | 502 | 439 | 651 | 546 | 584 |
| TAACTCAGTCTGCTGAAGTG | CASP8AP2 | 520 | 500 | 1145 | 926 | 881 |
| TCTAACAGTGATACTTCCGA | CASP8AP2 | 266 | 398 | 684 | 529 | 582 |
| TTTGAACAGTCGTCCAAATG | CASP8AP2 | 336 | 657 | 1253 | 1036 | 1026 |
| AAATACGTAGACTCCGAGGT | CAV1 | 448 | 360 | 757 | 742 | 778 |
| ATCCCAATGGCACTCATATG | CAV1 | 756 | 738 | 1196 | 1280 | 1314 |
| CAAGCATCTCAACGACGACG | CAV1 | 502 | 606 | 936 | 928 | 909 |
| CATCTACAAGCCCAACAACA | CAV1 | 438 | 300 | 419 | 366 | 355 |
| GATGTGATTGCAGAACCAGA | CAV1 | 525 | 326 | 558 | 454 | 464 |
| GCACACCAAGGAAATCGACC | CAV1 | 518 | 292 | 478 | 373 | 405 |
| AAGGCTCCCATGATTCTGAA | CBFB | 421 | 587 | 876 | 810 | 796 |
| AAGTCGACATACTCTCGGCT | CBFB | 520 | 597 | 889 | 759 | 807 |
| AGCCCATACCATCCAGTCTC | CBFB | 649 | 642 | 1115 | 1261 | 1207 |
| GAGAAGCAAGTTCGAGAACG | CBFB | 395 | 598 | 955 | 1080 | 1148 |
| GCCGACTTACGATTTCCGAG | CBFB | 575 | 571 | 972 | 1076 | 1053 |
| GGGCTGCCTGGAGTTTGATG | CBFB | 500 | 679 | 1529 | 1473 | 1474 |
| AATATGTGCTGAGAACGATA | CBL | 597 | 334 | 570 | 558 | 550 |
| CCTGGCGGAAACTCTTCCAA | CBL | 740 | 453 | 705 | 741 | 729 |
| GCAGCCATGATTCCCCGAGA | CBL | 594 | 560 | 1322 | 1161 | 1116 |
| GCAGGTCTAAGATATAAGGT | CBL | 546 | 504 | 1100 | 1055 | 995 |
| GTAGATCCGTTTGACCCTCG | CBL | 543 | 1131 | 1633 | 1486 | 1602 |
| TGTTGTAGGAGGTCAAGACG | CBL | 351 | 723 | 1288 | 1464 | 1460 |
| AGGAACAGAGCCCATAATCG | CBLB | 508 | 576 | 1006 | 1134 | 1080 |
| GCTGCACCAGATTGGGACAG | CBLB | 754 | 879 | 1675 | 1604 | 1606 |
| GTTTACCTTTGGTGAACCAG | CBLB | 624 | 691 | 1216 | 1176 | 1222 |
| TAATTTGATGAACCTCATGA | CBLB | 567 | 671 | 876 | 755 | 784 |
| TAGTTCCAAACACATCCCGA | CBLB | 623 | 393 | 483 | 474 | 478 |
| TGACCATATAAAAGTTACGC | CBLB | 661 | 211 | 329 | 328 | 301 |
| AGCAACTCAAACTCAGCCCA | CBLC | 360 | 916 | 1413 | 1455 | 1488 |
| CAAACTCGAAGATGGACACG | CBLC | 501 | 934 | 1331 | 1329 | 1406 |
| CATCTTCAGTTACATGCACG | CBLC | 466 | 733 | 1237 | 1214 | 1105 |
| CGTCTACGTTCACAACCTGG | CBLC | 578 | 558 | 1087 | 917 | 1033 |
| GCTACCTCTACCCAAATGGG | CBLC | 563 | 715 | 1239 | 1330 | 1316 |
| GGCCTTCCTCACATACGATG | CBLC | 659 | 587 | 1022 | 1014 | 1006 |
| CATCTCGCCGTTCCGCCTGG | CCDC6 | 417 | 1041 | 1323 | 1396 | 1421 |
| CTACATACGCTGTAGCTGAG | CCDC6 | 467 | 850 | 1261 | 1345 | 1334 |
| GACCTACAAACTCAAGTGCA | CCDC6 | 495 | 913 | 1046 | 1251 | 1246 |
| GAGATGTCTCTAGGCGACGG | CCDC6 | 460 | 463 | 935 | 989 | 1000 |
| GCTCTCCAGAAAATTGATGC | CCDC6 | 491 | 487 | 660 | 693 | 790 |
| TCCTGCTCGTCGACCTCGGG | CCDC6 | 361 | 520 | 1038 | 1030 | 1040 |
| CAGCTGCCACCATGGAGGGT | CCND1 | 529 | 282 | 591 | 483 | 523 |
| CAGGTGTAGACCTTGCACGG | CCND1 | 512 | 690 | 1440 | 1139 | 1248 |
| CGAGGCCACGAACATGCAGG | CCND1 | 622 | 495 | 1080 | 1027 | 976 |
| GTGCTCGATGAAGTCGTGTG | CCND1 | 500 | 495 | 1011 | 875 | 822 |
| TCTACACTGACAACTCCATC | CCND1 | 307 | 352 | 525 | 524 | 547 |
| TGTGGCACAGAGCGCTACGA | CCND1 | 625 | 605 | 1067 | 901 | 939 |
| GCAGAACTTGCTCACCATCG | CCND2 | 481 | 660 | 1401 | 1229 | 1262 |
| GCTCCCATCTTAGGTCTGTG | CCND2 | 403 | 840 | 1331 | 1417 | 1402 |
| GGTGGCACACAGAGCAATGA | CCND2 | 476 | 714 | 1293 | 1046 | 1073 |
| GTAAATGCACAACTTCTCTG | CCND2 | 312 | 553 | 916 | 944 | 931 |
| GTGCTCGATGAAGTCATGAG | CCND2 | 329 | 594 | 1045 | 1027 | 958 |
| TCGTAGAGCAGGTTGGCGTC | CCND2 | 547 | 434 | 997 | 960 | 968 |
| AATCAGAGCCAGGAAATCAT | CCND3 | 453 | 395 | 570 | 526 | 562 |
| ACAGAGGGCCAAAAAGGTCT | CCND3 | 402 | 493 | 933 | 944 | 1042 |
| CCTCTGCACACACTGGAAGT | CCND3 | 524 | 576 | 987 | 939 | 927 |
| CGCAGGACAAGTAGCGATCC | CCND3 | 563 | 521 | 824 | 842 | 876 |
| GCAGCATGCAGACCGCACCC | CCND3 | 565 | 359 | 954 | 812 | 805 |
| TCCCAACCTCTAGGACTGGG | CCND3 | 522 | 637 | 1162 | 1069 | 1092 |
| AAGATTTGCTGGGGATACTG | CCNE1 | 529 | 570 | 1019 | 1102 | 1110 |
| CATCTGTAACATAAGCAAAC | CCNE1 | 420 | 244 | 394 | 269 | 307 |
| CTGCATATATACATTCAGCC | CCNE1 | 512 | 358 | 548 | 531 | 564 |
| GAAGCAGCAAGTACGCCGTA | CCNE1 | 585 | 700 | 1144 | 1217 | 1266 |
| GGGATAAGCACTTTATGCAG | CCNE1 | 577 | 583 | 1079 | 1040 | 1046 |
| TCTTCTGGATTGGTTGATGG | CCNE1 | 466 | 950 | 1627 | 1715 | 1791 |
| ACTCCCGCAACTCTACAAGG | CCR4 | 507 | 535 | 755 | 822 | 831 |
| AGTGATGACCCCATACGTCA | CCR4 | 412 | 544 | 948 | 958 | 1098 |
| CAGGTGCTGAATAAAAGGCC | CCR4 | 570 | 393 | 718 | 818 | 767 |
| GAAGACCAAGGAGTAGAGAG | CCR4 | 548 | 653 | 1173 | 1016 | 1031 |
| GACCTAGTCCAAAAACCCAC | CCR4 | 672 | 149 | 341 | 326 | 320 |
| GAGCAGGTACACGTCAGTCA | CCR4 | 495 | 777 | 1124 | 1206 | 1181 |
| AAGCTCCCAGCTGTACGTGT | CD19 | 488 | 557 | 936 | 934 | 827 |
| GATAAAGTAATGGCACAGCA | CD19 | 502 | 583 | 1084 | 1060 | 1067 |
| GTTGCCATGGTAACAACGAT | CD19 | 301 | 511 | 966 | 1026 | 969 |
| TCGTGTCATTACCGTCTGGT | CD19 | 449 | 710 | 1366 | 1404 | 1396 |
| TGCTCAAGGGGACCATCTGA | CD19 | 482 | 934 | 1469 | 1577 | 1500 |
| TTAGGGCTGAAGCATTCCAC | CD19 | 362 | 343 | 582 | 488 | 457 |
| AACCCCACGCGGATTCGAGA | CD22 | 357 | 513 | 740 | 660 | 661 |
| AACTCCCAGCATGCCCGATG | CD22 | 554 | 835 | 1460 | 1351 | 1440 |
| CCTGTGCAGCCTGTAACCAG | CD22 | 649 | 691 | 1266 | 1336 | 1358 |
| CGGAGGAGCCAATGTAACAG | CD22 | 431 | 662 | 1175 | 1279 | 1211 |
| TCTGGAATCGTGTAGCAACA | CD22 | 561 | 575 | 1031 | 953 | 926 |
| TTGCCGTGGTGCGTCCACTG | CD22 | 626 | 862 | 1857 | 1728 | 1790 |
| AGAGATGATAGAGGGTCGGT | CD248 | 439 | 378 | 913 | 925 | 957 |
| ATGTGTGTCAACTATGTCGG | CD248 | 468 | 887 | 1690 | 1546 | 1564 |
| CCCCGAGGAAGCCCAACGCG | CD248 | 473 | 573 | 955 | 828 | 903 |
| CTCACACTGTTGCTCACACG | CD248 | 494 | 770 | 1347 | 1320 | 1329 |
| CTTTGATGGTAGCTGGACGG | CD248 | 504 | 714 | 1358 | 1301 | 1333 |
| TGGTCTCCACCGAGTTTGAG | CD248 | 448 | 948 | 1506 | 1531 | 1526 |
| ATGGCGGTGCTGACTACAAG | CD274 | 473 | 664 | 1353 | 1331 | 1248 |
| ATGTGAGATTGCAGGATGCA | CD274 | 555 | 741 | 1281 | 1157 | 1136 |
| CATCAGCCAAAGAATTTCTG | CD274 | 403 | 766 | 1041 | 1075 | 1051 |
| GTATGGTGGCAATGTGACAA | CD274 | 577 | 546 | 1170 | 1079 | 1148 |
| GTTTGCACTAATCGTCTACT | CD274 | 480 | 492 | 817 | 787 | 753 |
| TCAGGCTGAGGGTTACCCTG | CD274 | 585 | 1038 | 1535 | 1521 | 1490 |
| AAACTACCACTCGCGACCGG | CD37 | 388 | 641 | 1052 | 1059 | 1048 |
| ACCCCTCTAGGCGAAGATGT | CD37 | 500 | 633 | 1296 | 1216 | 1244 |
| AGCAGAAAATAAGGCTGCCT | CD37 | 655 | 273 | 593 | 581 | 528 |
| AGTGGAGATGAGTATTCCAA | CD37 | 535 | 737 | 1508 | 1414 | 1447 |
| GAGAATTCCTGAGATTGCCA | CD37 | 562 | 563 | 1007 | 881 | 900 |
| GGGATACCGAACCAGTCCCG | CD37 | 434 | 634 | 1215 | 1138 | 1183 |
| AAGGACGCAGAGACAGATCT | CD38 | 502 | 603 | 1134 | 1117 | 1183 |
| ACGTGTAGCACCGTCCCAGG | CD38 | 443 | 590 | 894 | 899 | 921 |
| AGACACAAGCTCTTCTGGTG | CD38 | 447 | 673 | 1355 | 1284 | 1331 |
| AGACTGCCAAAAGATAGGGA | CD38 | 516 | 644 | 987 | 1087 | 1058 |
| GCTGGCGCATGACTACACGC | CD38 | 366 | 414 | 680 | 642 | 652 |
| TCAAATTGTACCTTGTGACA | CD38 | 615 | 701 | 1367 | 1139 | 1206 |
| CGCACTCTTGACTTTCTTGG | CD52 | 560 | 642 | 1154 | 1114 | 1092 |
| CTTGGTGGTCATTCGAGGAG | CD52 | 440 | 914 | 1579 | 1725 | 1654 |
| GAGGAGTGGTACTGTTCCCC | CD52 | 508 | 621 | 1059 | 1030 | 1046 |
| GCCCAGGCTGCTGAGTGCTG | CD52 | 470 | 849 | 1655 | 1704 | 1701 |
| GGAGACTGATGGTAAGTAGG | CD52 | 384 | 745 | 1616 | 1453 | 1656 |
| TTCTTACCTGAATCATAACC | CD52 | 493 | 334 | 494 | 519 | 510 |
| CAAACTATACGAAGTACTCA | CD58 | 504 | 346 | 674 | 609 | 631 |
| CCGTTCACAGCTCCGAACAC | CD58 | 546 | 350 | 684 | 615 | 648 |
| GGCTGCTGGTGTATTACTGC | CD58 | 562 | 638 | 1142 | 1003 | 1023 |
| GTTGACGTGCAATTTAGTGT | CD58 | 490 | 554 | 989 | 948 | 914 |
| TGAACTCTACCTACAAAAGG | CD58 | 361 | 352 | 689 | 712 | 784 |
| TGATGATGTTAGATTGAAGA | CD58 | 430 | 469 | 760 | 600 | 554 |
| ACTGTGAACAAGAGCCACAT | CD79A | 450 | 593 | 1019 | 1130 | 1025 |
| AGGCTCACCGTCATCGATGG | CD79A | 360 | 920 | 1512 | 1633 | 1631 |
| ATCTTCCTGAGCTACGGCAA | CD79A | 445 | 689 | 1390 | 1253 | 1345 |
| GCAAGCTCAACATCACGTGG | CD79A | 431 | 631 | 1299 | 1108 | 1077 |
| GCTGGCACTCACCGCGCACG | CD79A | 496 | 602 | 1045 | 949 | 1026 |
| TGCGGTTATGCAGGCACTGG | CD79A | 433 | 735 | 1342 | 1320 | 1235 |
| CAACACCACCAGCCAGTTGT | CD79B | 504 | 542 | 1147 | 1197 | 1259 |
| CACACCCATGACTCGCAGCT | CD79B | 449 | 503 | 984 | 935 | 871 |
| GACAGGCACCAGGATCCCAA | CD79B | 561 | 632 | 815 | 887 | 936 |
| GGAAATCAGGTGCCACACGA | CD79B | 532 | 661 | 1248 | 1092 | 1110 |
| GGCATCCAGTTCTCGGACAA | CD79B | 441 | 638 | 1222 | 1231 | 1280 |
| TCTTCCGGGCTATGAAGCGC | CD79B | 567 | 616 | 932 | 1022 | 1108 |
| CAGGACGGGAGGATCTTCTC | CDA | 545 | 501 | 771 | 703 | 686 |
| CCACGGGGAAGTGACTGTAG | CDA | 559 | 694 | 1225 | 1124 | 1150 |
| CCCGTCCTGAGTGAGGATCG | CDA | 599 | 1066 | 1631 | 1628 | 1649 |
| CGCTCTGCACAGACGCCCAG | CDA | 631 | 744 | 1175 | 1289 | 1196 |
| TCAGGGCAATTGCTATCTCC | CDA | 513 | 434 | 914 | 928 | 860 |
| TCCAGAAGGCCATCTCAGAA | CDA | 348 | 626 | 1147 | 1182 | 1212 |
| AAGACCAGACCGATCTCAGG | CDC25A | 469 | 509 | 814 | 671 | 711 |
| AGCACATACCATCCGAGCTG | CDC25A | 539 | 730 | 1416 | 1340 | 1429 |
| CAGACAAAGTGGCCGTCACA | CDC25A | 780 | 556 | 1185 | 1355 | 1372 |
| GAGGCTTGCCATGCATGACG | CDC25A | 550 | 649 | 1316 | 1131 | 1120 |
| GCAGCCTTGAAAATCCCATG | CDC25A | 577 | 587 | 1352 | 1440 | 1374 |
| GCCGAACAGCGCTTTCACGA | CDC25A | 469 | 444 | 873 | 742 | 748 |
| AGACCATGCACAACCTCGCG | CDC25B | 520 | 454 | 1004 | 910 | 881 |
| ATGGGTCGGATCACGCTGCA | CDC25B | 508 | 700 | 1229 | 1086 | 1065 |
| CACAGTGTCTCACCCCTGAG | CDC25B | 547 | 681 | 1157 | 1232 | 1292 |
| GGTACCGCATTATCATCCTG | CDC25B | 492 | 807 | 1263 | 1073 | 1040 |
| TGAAGACAAACCCATCCTGT | CDC25B | 478 | 346 | 783 | 806 | 780 |
| TGGTGATGTTCCGTAGCACA | CDC25B | 475 | 708 | 1321 | 1233 | 1278 |
| AAGATCAAGGCAACATTTGG | CDC25C | 585 | 579 | 1011 | 1045 | 1043 |
| CCAACTAAACGTATTTCCTG | CDC25C | 441 | 625 | 1103 | 1090 | 1136 |
| GTGTGTTGCCAACCGTGTCA | CDC25C | 541 | 588 | 1251 | 940 | 1043 |
| TAATATGATTGACAGGACCA | CDC25C | 307 | 764 | 1188 | 1201 | 1110 |
| TAGCCACAAAGAGCTCGGGA | CDC25C | 568 | 582 | 915 | 964 | 921 |
| TTCAAATTCTCTGGCAACGA | CDC25C | 343 | 424 | 677 | 654 | 659 |
| ACCCAGAGAGTACTACACAT | CDC73 | 417 | 356 | 760 | 456 | 507 |
| AGCAAAGAAACCACGCATTG | CDC73 | 578 | 914 | 1636 | 1438 | 1472 |
| CGGTCAGGTCTTCTAACCAC | CDC73 | 706 | 411 | 864 | 710 | 656 |
| GGCTGTTTGGTACGCAACGT | CDC73 | 263 | 422 | 783 | 612 | 680 |
| GGTTTCAAAATCGACACTAT | CDC73 | 554 | 418 | 672 | 641 | 761 |
| TTTGTCTGAAGCTATGTCAG | CDC73 | 425 | 683 | 1467 | 1493 | 1514 |
| AATCACATCCTACACCGCCG | CDH1 | 460 | 341 | 637 | 612 | 596 |
| ACAGACCCAGTAACTAACGA | CDH1 | 428 | 386 | 515 | 423 | 479 |
| CGTTATAGACCATACCTGGA | CDH1 | 477 | 666 | 872 | 866 | 864 |
| CTCATTATAGCCATTGACAA | CDH1 | 564 | 417 | 606 | 624 | 600 |
| GGGTGTCCCCATGTACACCT | CDH1 | 520 | 384 | 756 | 743 | 748 |
| TGATGACACCCGATTCAAAG | CDH1 | 500 | 608 | 516 | 577 | 616 |
| AGACAAGCGACTAAGCACAA | CDH2 | 507 | 380 | 675 | 666 | 675 |
| CCAGCACCCTCCTCAGTCAA | CDH2 | 621 | 758 | 1106 | 1020 | 995 |
| GTTCCTGTTCCACTCATAGG | CDH2 | 530 | 748 | 1253 | 1268 | 1304 |
| TAATCTAACTGTGACTGACA | CDH2 | 546 | 507 | 847 | 773 | 816 |
| TTGTAAACATGTTGGGCGAA | CDH2 | 330 | 664 | 1164 | 1349 | 1248 |
| TTGTCCCAAGATGTGCATGA | CDH2 | 410 | 592 | 1129 | 1105 | 1109 |
| AACAACAGTGCACATCAGCG | CDH20 | 544 | 557 | 1029 | 975 | 955 |
| AGTCCACAGAGAAATACGGC | CDH20 | 540 | 324 | 525 | 608 | 646 |
| CCACATAGAAGAATCTTCCA | CDH20 | 695 | 303 | 634 | 549 | 542 |
| GACGGAAACACTTCCAACCT | CDH20 | 463 | 306 | 769 | 630 | 723 |
| GGCGCCAGTTAGCTCCACCG | CDH20 | 687 | 602 | 1107 | 993 | 926 |
| GGGCTCAAGCCCTAGACAGG | CDH20 | 614 | 638 | 1428 | 1305 | 1272 |
| ACGGACCCCAACCACAACGA | CDH5 | 628 | 619 | 1177 | 1087 | 1080 |
| CCCTTGGTATAACCTGACAG | CDH5 | 453 | 735 | 1472 | 1491 | 1546 |
| GCAAATCTCGGCAATAGACA | CDH5 | 489 | 490 | 668 | 676 | 745 |
| GCCCACATAATGGGGTAGCG | CDH5 | 580 | 724 | 1380 | 1189 | 1092 |
| TGATGCAGATGATCCCACTG | CDH5 | 465 | 1168 | 1563 | 1526 | 1499 |
| TGCTCCGACGAGCTGCATCG | CDH5 | 517 | 945 | 1648 | 1644 | 1583 |
| AAGGGTAGACACAAAACTAC | CDK1 | 508 | 290 | 500 | 446 | 458 |
| AATCCATGAACTGACCAGGA | CDK1 | 584 | 324 | 662 | 492 | 457 |
| AATCTATTGATTGATGACAA | CDK1 | 532 | 470 | 925 | 789 | 797 |
| ATTTCCCGAATTGCAGTACT | CDK1 | 618 | 544 | 935 | 901 | 846 |
| TATTTGTACCAAATCCTACA | CDK1 | 592 | 321 | 444 | 353 | 370 |
| TGTAACAGGTAGTAACACTG | CDK1 | 489 | 330 | 653 | 463 | 475 |
| AAGACCGGAGTAAAAGTCAT | CDK12 | 524 | 546 | 1039 | 829 | 969 |
| AGGTCTGTGAGTAAGTGACG | CDK12 | 482 | 857 | 1367 | 1409 | 1454 |
| GGGAGAATGATGAACGTCGT | CDK12 | 624 | 415 | 778 | 752 | 770 |
| TAAACCCATAGTATTGAAAG | CDK12 | 436 | 479 | 774 | 733 | 684 |
| TCTTTGTAGATGCATCCTCG | CDK12 | 499 | 765 | 1330 | 1341 | 1425 |
| TGGCCAAGTATATAAAGCCA | CDK12 | 404 | 383 | 781 | 628 | 619 |
| ACTTCTAGATAGGAGCACAG | CDK2 | 561 | 376 | 547 | 564 | 558 |
| ATCTCTCGTATAGCAGTACT | CDK2 | 546 | 464 | 1055 | 1002 | 998 |
| CATGGGTGTAAGTACGAACA | CDK2 | 485 | 862 | 1658 | 1618 | 1541 |
| GAATCTGCTTATCAACGCAG | CDK2 | 556 | 399 | 711 | 667 | 646 |
| GGATGACATCCAATAGCCTA | CDK2 | 338 | 569 | 1070 | 958 | 1037 |
| TGAGAAGGATTACCTTGATG | CDK2 | 590 | 786 | 1370 | 1345 | 1355 |
| AGCGTTAGAGTCCCTAATGG | CDK4 | 517 | 728 | 1455 | 1429 | 1490 |
| CCAGATGGCACTTACGCCTG | CDK4 | 399 | 673 | 1407 | 1256 | 1241 |
| GGCCTTGTACACCGTTCCAT | CDK4 | 577 | 362 | 950 | 853 | 872 |
| GTCTACGTATGCAACACCCG | CDK4 | 556 | 489 | 1061 | 808 | 950 |
| GTCTCGGTGAACGATGCAGT | CDK4 | 407 | 485 | 1005 | 933 | 984 |
| TTCGTGAAGTGGCATTGCTG | CDK4 | 339 | 764 | 1493 | 1420 | 1436 |
| ACAGCGAACAGGGATCCCAA | CDK5 | 515 | 608 | 1001 | 798 | 844 |
| CCGGGAGACCCATGAGATCG | CDK5 | 484 | 971 | 1714 | 1793 | 1788 |
| CCTGGATCCTGAAATTGTGA | CDK5 | 630 | 746 | 1262 | 1090 | 1127 |
| GACCCCAGCTCTATCCCGCG | CDK5 | 568 | 394 | 867 | 782 | 738 |
| TCAGCTTCTTGTCACTATGC | CDK5 | 463 | 507 | 941 | 908 | 986 |
| TGAGTAGGCAGATTTCCCGA | CDK5 | 308 | 605 | 966 | 1037 | 1073 |
| AACACTCCAGAGGTCCACGG | CDK6 | 585 | 312 | 765 | 606 | 620 |
| AGTTCAGATGTCGATCAACT | CDK6 | 441 | 469 | 988 | 888 | 855 |
| ATGATGTTTCAGCTTCTCCG | CDK6 | 495 | 409 | 905 | 799 | 787 |
| CCTTGCAGGTCGTCACACTG | CDK6 | 369 | 400 | 893 | 832 | 820 |
| CTGGTCACTAAAATGTTCTG | CDK6 | 570 | 471 | 1180 | 1111 | 1152 |
| GAAAACTGTAGATGCGGGCA | CDK6 | 364 | 599 | 1041 | 952 | 1031 |
| ATGGTGTAGGTGTCGACATG | CDK7 | 443 | 1052 | 1749 | 1851 | 1827 |
| CAGATTGCCCCGGTTTACAA | CDK7 | 493 | 636 | 963 | 891 | 915 |
| GATGTGTATAAGCTCTATTG | CDK7 | 581 | 505 | 1194 | 1059 | 1125 |
| TCAACACTGGATTCTACATA | CDK7 | 432 | 457 | 758 | 792 | 811 |
| TGTAGGCTTTGATGTGTGAT | CDK7 | 427 | 932 | 1208 | 1277 | 1320 |
| TTTCCATAAAATCAAAGACA | CDK7 | 608 | 277 | 400 | 321 | 382 |
| AAGCCAGTTCAGTTACCTCG | CDK8 | 499 | 651 | 1134 | 1123 | 1093 |
| AATATTTCACTGTCGACAAG | CDK8 | 533 | 742 | 977 | 933 | 1057 |
| ACCGGATCCAAATCTGCTAA | CDK8 | 402 | 724 | 1240 | 1156 | 1218 |
| AGACACTATACCAAAGCTAT | CDK8 | 533 | 415 | 701 | 661 | 599 |
| TATGCTTTAAAACAAATAGA | CDK8 | 463 | 437 | 602 | 614 | 644 |
| TGCTAACTGGGTGTTGCACA | CDK8 | 469 | 657 | 1409 | 1538 | 1419 |
| AAGCTCGCTAAGATCGGCCA | CDK9 | 560 | 417 | 976 | 671 | 738 |
| ACCCGCAGCCCTATCATGCA | CDK9 | 654 | 523 | 942 | 781 | 692 |
| CAATCGCTATACCAACCGTG | CDK9 | 465 | 639 | 1239 | 1111 | 1089 |
| CTATATCCACAGAAACAAGG | CDK9 | 588 | 249 | 427 | 312 | 362 |
| GCTCGCAGAAGTCAAACACT | CDK9 | 432 | 648 | 956 | 934 | 921 |
| GCTTCTGAAACACGAGAATG | CDK9 | 762 | 622 | 1427 | 1400 | 1337 |
| AGGTCATCCCGGCCCCCCCG | CDKN1A | 572 | 506 | 1045 | 962 | 988 |
| CAGGCAAAGTCGCCTTCCAG | CDKN1A | 486 | 968 | 1260 | 1303 | 1294 |
| CAGGCAGGGAGACCTTGGAC | CDKN1A | 629 | 415 | 784 | 750 | 777 |
| CCGGGACTGTGACGCGCTGA | CDKN1A | 5 | 1168 | 1373 | 1412 | 1463 |
| GGCAAGCCTTGCTGCCATGA | CDKN1A | 537 | 1282 | 1946 | 1786 | 1746 |
| GTCAAAGTTCCATCGCTCCC | CDKN1A | 462 | 708 | 878 | 829 | 842 |
| AATCAAAATTCCACTTGCGC | CDKN1B | 537 | 212 | 268 | 225 | 248 |
| AGTTCTACTACAGACCCCCG | CDKN1B | 442 | 813 | 1173 | 1228 | 1217 |
| CAACCACGAAGAGTTGACCC | CDKN1B | 605 | 669 | 680 | 754 | 772 |
| CCCTGGAGGGCAAATACGAG | CDKN1B | 485 | 508 | 1163 | 1160 | 1089 |
| CGAGCGGATGGACGCCAGAC | CDKN1B | 555 | 500 | 777 | 839 | 775 |
| GCCAGGCGGGGCCTTTACTG | CDKN1B | 510 | 721 | 1298 | 1477 | 1514 |
| CACAGTTGGGGTTGGCGCCG | CDKN2A | 472 | 992 | 1622 | 1407 | 1330 |
| CACCCGCCCTGTGCACGACG | CDKN2A | 615 | 1381 | 3621 | 2235 | 2216 |
| CAGGGCGGGTGAGGGTGACG | CDKN2A | 545 | 730 | 1462 | 1307 | 1352 |
| CCGTGGACCTGGCTGAGGAG | CDKN2A | 708 | 1501 | 1994 | 1764 | 1886 |
| CTGGACGTGCGCGATGCCTG | CDKN2A | 435 | 2315 | 3390 | 2775 | 2705 |
| GATGATGGGCAGCACCCGCG | CDKN2A | 381 | 1328 | 1351 | 1260 | 1421 |
| AGGGGCAGGCGGAGACCGTG | CDKN2B | 516 | 884 | 1297 | 1356 | 1422 |
| CACAGTTGGGGTCGGCGCCG | CDKN2B | 528 | 1173 | 1436 | 1579 | 1570 |
| CTTCGGCAGGCGCCCGATCC | CDKN2B | 397 | 566 | 840 | 703 | 724 |
| GAAGCCGGCGCGGATCCCAA | CDKN2B | 413 | 495 | 943 | 879 | 958 |
| GAGGCCACGGGCTTGCTCGG | CDKN2B | 367 | 842 | 1255 | 1175 | 1244 |
| GTGCACAGGGCGGGTGAGGG | CDKN2B | 351 | 615 | 1041 | 989 | 1056 |
| AAGTTGCTCTAGGTCCCCCC | CDKN2C | 567 | 586 | 894 | 913 | 963 |
| ACCTCTAAGTAGTAGTCTCC | CDKN2C | 568 | 457 | 901 | 866 | 886 |
| ATTTCCAAGTTTCATAACCT | CDKN2C | 442 | 406 | 772 | 816 | 775 |
| GAATGACAGCGAAACCAGTT | CDKN2C | 643 | 491 | 848 | 793 | 790 |
| GCAAAGTCTGTAAAGTGTCC | CDKN2C | 624 | 575 | 940 | 920 | 934 |
| TTAACATCGAGGATAATGAA | CDKN2C | 381 | 318 | 766 | 808 | 808 |
| AAAGGCGGACGCCCTACGAG | CDX1 | 528 | 677 | 1270 | 1214 | 1099 |
| AGGACTCCCCGGCCTACCCC | CDX1 | 574 | 434 | 815 | 854 | 784 |
| AGGGGGCCCGAATGCCAGCG | CDX1 | 717 | 987 | 1666 | 1606 | 1653 |
| CTCAGGTAAGACCCGGACCA | CDX1 | 476 | 862 | 1565 | 1479 | 1523 |
| GGGAGTAGCCGGCGAAGTCG | CDX1 | 493 | 828 | 1418 | 1404 | 1459 |
| TCGTCCTTGGGCGCCGAGAA | CDX1 | 486 | 526 | 839 | 874 | 871 |
| AAGCCTTACCTGGTTCAAAG | CEACAM28 | 564 | 419 | 659 | 650 | 658 |
| ACACCAAAATCACTAACCCG | CEACAM28 | 465 | 363 | 638 | 554 | 554 |
| ACACCGGGATCTCTATCCAG | CEACAM28 | 266 | 507 | 1170 | 1030 | 956 |
| CTGCTCCTCAAGTAACACTG | CEACAM28 | 664 | 230 | 465 | 386 | 400 |
| GGGTTGGGACATTGACTCTG | CEACAM28 | 727 | 703 | 1203 | 1204 | 1294 |
| TTTGAAACTGCACTAATACG | CEACAM28 | 334 | 581 | 1111 | 1067 | 1046 |
| ACCTCCGCCCCAGTTCCGGT | CEBPA | 468 | 615 | 1064 | 963 | 995 |
| CAGTTCCAGATCGCGCACTG | CEBPA | 463 | 660 | 1226 | 1190 | 1227 |
| CCGCCGGCCAGCCGACACCG | CEBPA | 414 | 465 | 1114 | 1121 | 1179 |
| CCGGGCGCTAGGACCGCGCG | CEBPA | 582 | 647 | 1161 | 1197 | 1182 |
| GAAAAAGTTGTCCTGCCCCG | CEBPA | 419 | 549 | 1038 | 1079 | 1071 |
| GCGTCCCCCTCCCTGACCTC | CEBPA | 410 | 443 | 991 | 798 | 802 |
| AGTGTGGTAGTGACCCCCCG | CERK | 443 | 1046 | 1881 | 1689 | 1874 |
| CACGGGACGAAGGAACCAAG | CERK | 298 | 547 | 1075 | 1078 | 1067 |
| CGTGCTGTTGTGATGAACGG | CERK | 452 | 895 | 1641 | 1491 | 1592 |
| CGTGGGAAGCACTACGGTAG | CERK | 314 | 1061 | 1934 | 1820 | 1913 |
| GGAGGCTAGCGTAAACAACG | CERK | 515 | 687 | 1012 | 1049 | 1029 |
| GGGATGATGCCGATCCTGAG | CERK | 553 | 646 | 1157 | 1138 | 1146 |
| AAGATGTGAGGGGCCCGTAG | CERS1 | 251 | 901 | 1415 | 1270 | 1284 |
| AATGTCTACTTCAAGTCCCG | CERS1 | 737 | 403 | 806 | 667 | 772 |
| AGGTGAGTATGGGGGCATCG | CERS1 | 378 | 1051 | 1697 | 1652 | 1716 |
| CCCACCAGGTATCACAACGT | CERS1 | 478 | 447 | 800 | 849 | 829 |
| GAGCTGGGGCTATACCATGG | CERS1 | 403 | 696 | 1421 | 1375 | 1364 |
| TACCGTAGAAGACGGAGGGT | CERS1 | 495 | 952 | 1503 | 1284 | 1377 |
| AATCATGTAGTACCAATACT | CERS2 | 513 | 451 | 578 | 656 | 628 |
| CTCTTCCGACTACCTACTAG | CERS2 | 490 | 577 | 760 | 774 | 796 |
| GGATGATAGTAGCCACGTGG | CERS2 | 683 | 745 | 1240 | 1336 | 1326 |
| GGTCCTGGTTGCGACGACGG | CERS2 | 502 | 521 | 1104 | 959 | 1057 |
| GTGGTGTAGCCACATAACTG | CERS2 | 438 | 814 | 1246 | 1150 | 1232 |
| TTCTTCATGTCATAGAACCA | CERS2 | 583 | 649 | 1260 | 1059 | 1042 |
| AATTGTCATGTACAAATGAG | CERS3 | 533 | 822 | 1373 | 1438 | 1475 |
| CAAAATATAGTACCAGTACT | CERS3 | 442 | 667 | 1056 | 932 | 966 |
| CATCATGCACAATCATCACG | CERS3 | 565 | 428 | 729 | 687 | 725 |
| TCATCAGACTAATAGCAGCC | CERS3 | 542 | 426 | 523 | 566 | 564 |
| TCCCAGAGGTCATATACCCA | CERS3 | 655 | 632 | 1058 | 1094 | 1097 |
| TCCGGAACCATCGTTCTACC | CERS3 | 388 | 469 | 982 | 991 | 856 |
| AATACTTCCTCACGAAAGGG | CERS4 | 486 | 504 | 906 | 961 | 884 |
| ACACCTAGCAACCGGCTCAG | CERS4 | 760 | 723 | 1312 | 1311 | 1379 |
| GCTCCCACCCAACAACACGT | CERS4 | 487 | 296 | 455 | 433 | 461 |
| TAGTACAGGGCCGGTTTCAG | CERS4 | 404 | 837 | 1523 | 1458 | 1458 |
| TCACGTGGTAGAGGATCGAC | CERS4 | 407 | 421 | 634 | 664 | 699 |
| TTGAACATCTTACCGGCCTG | CERS4 | 346 | 917 | 1599 | 1523 | 1375 |
| AAACATAAGAGACCAATAGA | CERS5 | 699 | 322 | 509 | 414 | 417 |
| ACAGAATTTAGTGAGCGTTG | CERS5 | 497 | 938 | 1503 | 1408 | 1542 |
| ATGCCAACATGGAGTGCACA | CERS5 | 365 | 685 | 1163 | 1203 | 1255 |
| GAAGGTTATAAGCCCAATGG | CERS5 | 395 | 766 | 1481 | 1342 | 1387 |
| GAGAGGATGTGCCGCGCCCG | CERS5 | 519 | 779 | 1463 | 1340 | 1398 |
| GATAGCTATGCCAGCACTGT | CERS5 | 533 | 500 | 925 | 790 | 922 |
| ACGTTCGCAGCATTCAACGC | CERS6 | 554 | 402 | 658 | 581 | 583 |
| GCCCCGCTTACCTCTCGAAG | CERS6 | 508 | 583 | 1264 | 1164 | 1247 |
| GCTCCCGCACAATGTCACCT | CERS6 | 555 | 421 | 887 | 781 | 833 |
| GGTCTTCACTGCAATTACGA | CERS6 | 535 | 344 | 563 | 478 | 476 |
| GTCAACAATATGGCCCGAGT | CERS6 | 517 | 381 | 744 | 807 | 829 |
| GTCCGGTTCCTAAAAAAGGT | CERS6 | 432 | 441 | 733 | 785 | 710 |
| CGTTTATTGAACAAGATGTG | CHEK1 | 584 | 723 | 1486 | 1319 | 1395 |
| GAGGTTCTTTCACCAACTCA | CHEK1 | 587 | 552 | 998 | 825 | 873 |
| GGGATGGTAATCCTTACTGA | CHEK1 | 514 | 651 | 1085 | 1045 | 1014 |
| GGTGTATCAGAGTCTCCTAG | CHEK1 | 479 | 670 | 1230 | 1100 | 1134 |
| GTTGTCAAGAATATTCTGAT | CHEK1 | 629 | 327 | 615 | 529 | 569 |
| TGCTGTAAGTACTATTCCAC | CHEK1 | 447 | 344 | 429 | 425 | 415 |
| AGCAAGAGGAAGTTCGCTAT | CHEK2 | 461 | 758 | 1008 | 1141 | 1141 |
| AGGAGAGCTGTTTGACAGGG | CHEK2 | 659 | 1033 | 1426 | 1522 | 1640 |
| GCATACATAGAAGATCACAG | CHEK2 | 603 | 407 | 494 | 446 | 448 |
| GGCATCTCCTCGCAGACACA | CHEK2 | 595 | 545 | 740 | 830 | 827 |
| GGCCCATAATCGAGCCCAAG | CHEK2 | 479 | 540 | 996 | 973 | 1032 |
| TCATGTCAAAAACTCTCGGA | CHEK2 | 458 | 299 | 555 | 610 | 600 |
| AAAGCTCCAATAATCAACAG | CHUK | 438 | 341 | 548 | 535 | 489 |
| AAGCAGTGCACTATGTGTCT | CHUK | 551 | 628 | 1080 | 1143 | 1079 |
| AATAAACCAGAAAATTGCTG | CHUK | 442 | 691 | 1175 | 1107 | 1195 |
| AGGCCTTTACAACATTGGCG | CHUK | 533 | 671 | 1191 | 1145 | 1118 |
| TAGTTTAATAGTAGAACCCA | CHUK | 522 | 528 | 1057 | 1011 | 966 |
| TCCATCTAGAACACATTGAG | CHUK | 623 | 554 | 960 | 1046 | 1012 |
| ATCCGTTGGAAGGAACCGGG | CIC | 454 | 683 | 1339 | 1588 | 1527 |
| CCTGGGGGTACAGTTCTCCG | CIC | 386 | 687 | 1187 | 1302 | 1288 |
| CGAAACATCGCGTGACAGTG | CIC | 520 | 605 | 1311 | 1216 | 1268 |
| CGACAGCCTAGACTCTGGCG | CIC | 419 | 620 | 1365 | 1418 | 1412 |
| GAAGCTGATCTTAGACACCT | CIC | 467 | 352 | 715 | 752 | 725 |
| TACTAAGCTTACACCCAGTG | CIC | 517 | 735 | 1408 | 1485 | 1629 |
| ACGCTCACGGGACTCTACGT | CIITA | 466 | 424 | 800 | 747 | 782 |
| CAGGATGAGAAGAACACGGT | CIITA | 570 | 552 | 1046 | 1073 | 988 |
| CCTCGTGAGCTGCCACACAT | CIITA | 522 | 705 | 1075 | 1063 | 953 |
| GGCCACCTCGAACACGACGT | CIITA | 556 | 525 | 805 | 849 | 971 |
| GGCTTATGCCAATATCGGTG | CIITA | 348 | 776 | 1269 | 1112 | 1203 |
| GGGATGCTGACTTACCAACA | CIITA | 656 | 706 | 1224 | 1320 | 1268 |
| CGTTTCGTCTGGATCATATC | CLK1 | 12 | 445 | 817 | 806 | 776 |
| CTTCACACCATCGTTCACAT | CLK1 | 476 | 284 | 466 | 405 | 403 |
| GAGATCACATAGCAGTGCGC | CLK1 | 466 | 389 | 653 | 578 | 541 |
| GATACTTACAATGTACTGCT | CLK1 | 490 | 364 | 509 | 487 | 471 |
| TGTGCAGTCTGACTACACAG | CLK1 | 629 | 516 | 848 | 901 | 888 |
| TTAACTTTCTCTAACAGAGG | CLK1 | 507 | 590 | 938 | 952 | 982 |
| ATGAGAGAGATTATCGAGAT | CLK4 | 410 | 631 | 1102 | 1024 | 914 |
| ATTCTAGCATCTGGACACAT | CLK4 | 387 | 371 | 652 | 617 | 668 |
| GATATGCCATCTGCCTGATG | CLK4 | 421 | 886 | 1416 | 1325 | 1408 |
| GCACTCTACGACTTTGCCAA | CLK4 | 385 | 490 | 918 | 861 | 874 |
| GTAAATCTTCGGTCCGAAGC | CLK4 | 590 | 433 | 637 | 622 | 657 |
| GTTCTCTTGAGTACTACTGT | CLK4 | 756 | 567 | 1085 | 952 | 933 |
| AGAAGAGCAAACGGAACGCT | CLTA | 489 | 545 | 923 | 886 | 850 |
| ATAGCTGCGTAACTGTCTGT | CLTA | 460 | 685 | 1329 | 1255 | 1301 |
| CTCGGAAGCAAGAAGCAGAG | CLTA | 485 | 524 | 893 | 959 | 1028 |
| CTCTCACCCGGAACCCCCGG | CLTA | 460 | 771 | 1352 | 1354 | 1415 |
| GCAGCAGGAGAGCGAGATTG | CLTA | 444 | 881 | 1769 | 1829 | 1832 |
| GCCTTCGCCATCCTGGACGG | CLTA | 493 | 876 | 1554 | 1438 | 1431 |
| AAACACATCTCCATTGATCG | CLTB | 575 | 514 | 1079 | 1020 | 1041 |
| AACCTGAGAGCATCCGCAAG | CLTB | 560 | 640 | 1301 | 1321 | 1296 |
| AGGTGACAGAACAGGAGTGG | CLTB | 475 | 958 | 1719 | 1735 | 1726 |
| CAACTTGTTCACTCTGACGC | CLTB | 650 | 366 | 758 | 763 | 762 |
| GCGGGCATAGAGAACGACGA | CLTB | 436 | 515 | 970 | 964 | 869 |
| TTCTTCTCGTCGTCGGAGAG | CLTB | 442 | 1081 | 1804 | 1646 | 1717 |
| ATAACCACAGGAAGTCGACT | CLTC | 712 | 361 | 776 | 608 | 686 |
| CCTTTCGGCGTACCAAATAG | CLTC | 563 | 542 | 1115 | 1013 | 1034 |
| TATACTCCATAACACGTGTA | CLTC | 594 | 580 | 1379 | 1214 | 1231 |
| TGCCCAGGGGAATTACTCAG | CLTC | 492 | 567 | 1205 | 1122 | 1125 |
| TTTATATGACATAAAGCGTG | CLTC | 605 | 430 | 985 | 865 | 874 |
| TTTGTAGCAAAATCGTGTGG | CLTC | 431 | 820 | 1599 | 1578 | 1525 |
| AAAATAATCGCCCAGCTGAG | CLTCL1 | 526 | 603 | 980 | 872 | 908 |
| CATTAAGGCAGACCGCACAC | CLTCL1 | 521 | 315 | 621 | 596 | 662 |
| GGCTCAGACTGATTTCCCAG | CLTCL1 | 597 | 515 | 883 | 757 | 737 |
| TCAGTACCCTCTTTGCGCAG | CLTCL1 | 459 | 553 | 1092 | 945 | 962 |
| TGCGGTATAAGTACAAGACG | CLTCL1 | 283 | 469 | 1252 | 1090 | 1089 |
| TGCTTACTATGACAGCCGTG | CLTCL1 | 504 | 792 | 1387 | 1518 | 1357 |
| ATTGAAAAGTACATTAAAGA | CMPK1 | 535 | 291 | 464 | 399 | 393 |
| CGGGCCTCACCTCGACGATG | CMPK1 | 447 | 654 | 1588 | 1544 | 1414 |
| CTGGTCGTGTTCGTCCTCGG | CMPK1 | 501 | 247 | 547 | 484 | 534 |
| GTTCACCATACTGTGAAGCT | CMPK1 | 583 | 388 | 838 | 679 | 627 |
| GTTGGAATAAGACCATGGAT | CMPK1 | 518 | 423 | 861 | 828 | 847 |
| TCAGTTTGTTAAAGAGGGTA | CMPK1 | 315 | 653 | 1259 | 1077 | 1073 |
| AAAATTACATTAGTAGCCGG | CNKSR3 | 461 | 551 | 1025 | 984 | 1049 |
| CCTGACCACTACAGTCCAGA | CNKSR3 | 655 | 548 | 804 | 773 | 802 |
| CTTCCTCCAGACACGCACAC | CNKSR3 | 635 | 214 | 383 | 340 | 316 |
| GCACGTGATTACTGGAACCA | CNKSR3 | 510 | 634 | 1310 | 1219 | 1211 |
| GCAGCAATACGACTCCTGTG | CNKSR3 | 518 | 731 | 1370 | 1295 | 1284 |
| GTTTCCAGGCCATAATTCTG | CNKSR3 | 571 | 540 | 1021 | 921 | 1021 |
| AGGAAGTGACACTCACGGCA | COL1A1 | 561 | 678 | 1075 | 968 | 994 |
| CAATGGGGCCAGTCAGACCA | COL1A1 | 765 | 855 | 1278 | 1161 | 1194 |
| GCTCTCCTACAGGGTAACGT | COL1A1 | 589 | 358 | 648 | 548 | 519 |
| GGGACGACCATCCTGACCAG | COL1A1 | 693 | 501 | 970 | 1114 | 1155 |
| TCATAGCCATAAGACATTTG | COL1A1 | 518 | 639 | 1268 | 1213 | 1241 |
| TTTCACCGGGCAGACCACGG | COL1A1 | 536 | 549 | 915 | 793 | 853 |
| ACACTAGACGAAACAGAACA | CPT1A | 578 | 621 | 812 | 714 | 798 |
| AGCGAATGTTTAATACCTCG | CPT1A | 491 | 685 | 1337 | 1442 | 1373 |
| GAGGATTCTGGACGACCCGT | CPT1A | 520 | 379 | 819 | 812 | 824 |
| GCCTCGGACCCAAGTTGCAG | CPT1A | 401 | 890 | 1508 | 1424 | 1468 |
| GTCTGGAAGCTGTATAACAT | CPT1A | 482 | 596 | 898 | 848 | 936 |
| TTGCAATTATTCCCAGCGAG | CPT1A | 526 | 573 | 1255 | 1195 | 1158 |
| ACCAGTATTCATGTAAATGG | CRBN | 404 | 438 | 837 | 765 | 732 |
| ATCAGGATCATCATCACCTG | CRBN | 617 | 794 | 1293 | 1216 | 1193 |
| TGAAAGCAATTGGAAGACAA | CRBN | 384 | 520 | 934 | 860 | 867 |
| TGAACTGCAGACATGGTTGA | CRBN | 606 | 953 | 1618 | 1650 | 1693 |
| TTACGTACTGTATGTGATGT | CRBN | 457 | 972 | 1445 | 1280 | 1322 |
| TTGGAAGTAATCTACAGGAG | CRBN | 367 | 908 | 1483 | 1694 | 1684 |
| AGACTGAATAACTGATGGCT | CREB1 | 569 | 597 | 1130 | 1178 | 1098 |
| AGCTGTACTAAAGTTACGGT | CREB1 | 504 | 415 | 704 | 637 | 629 |
| CTGGCTAACAATGGTACCGA | CREB1 | 514 | 576 | 1002 | 1019 | 1032 |
| GAAGAGAAATTCTTTCAAGG | CREB1 | 730 | 338 | 483 | 524 | 537 |
| GCTGCTAGTTTGGTAAATCG | CREB1 | 482 | 656 | 1329 | 1220 | 1332 |
| TGGGCTAATGTGGCAATCTG | CREB1 | 381 | 681 | 1317 | 1200 | 1281 |
| AGCCCTAAAGGATCGCCGCA | CREBBP | 520 | 714 | 1265 | 1241 | 1291 |
| AGGAATAGTACTTACGTCTG | CREBBP | 386 | 837 | 1462 | 1468 | 1527 |
| CTTCTGGCCTAATGTGCTAG | CREBBP | 562 | 839 | 1401 | 1325 | 1388 |
| GCTGGCTGAGACATTAACAC | CREBBP | 458 | 330 | 530 | 541 | 583 |
| TGCCGGAAAGGTAACGACTC | CREBBP | 365 | 614 | 1085 | 956 | 1026 |
| TTTGTCGTGAAGATGCACAA | CREBBP | 324 | 557 | 986 | 1052 | 1095 |
| CCCTGGGGATTATGTGCTGT | CRKL | 297 | 582 | 1067 | 955 | 976 |
| CCTCACCACATGGAAAGCAC | CRKL | 443 | 332 | 505 | 422 | 445 |
| CGCGAACTAGGAACATTCCG | CRKL | 632 | 431 | 739 | 621 | 661 |
| GAACAGGAATCATCCCAATC | CRKL | 390 | 530 | 775 | 862 | 832 |
| GGGCGCGGGCTCGATTAGGG | CRKL | 488 | 459 | 982 | 912 | 906 |
| GGTCCCCGATCTTAAAACGG | CRKL | 387 | 831 | 1290 | 1475 | 1379 |
| AAGACGTGTTCCATTCCAGA | CRLF2 | 321 | 406 | 686 | 654 | 647 |
| AGAGTATGTTCGACACTGAG | CRLF2 | 525 | 913 | 1340 | 1245 | 1258 |
| AGTCACTTGGGTATGTGTCG | CRLF2 | 557 | 942 | 1620 | 1693 | 1633 |
| CAGAAACTAAAATAAATTTG | CRLF2 | 578 | 504 | 927 | 998 | 1076 |
| GGAATGCAAGCCAATACTCG | CRLF2 | 457 | 787 | 1331 | 1427 | 1418 |
| GTTTCGGGGGACGAGGCACT | CRLF2 | 681 | 734 | 993 | 962 | 999 |
| CAGCCAGAGCCCTTTACAGG | CRTC1 | 388 | 790 | 1268 | 1326 | 1291 |
| CCAAGTGCCAGCTTGACGCG | CRTC1 | 545 | 801 | 1421 | 1246 | 1316 |
| CTGAGACTTACTTGATCCCA | CRTC1 | 657 | 581 | 1099 | 1108 | 1015 |
| TCAGCCCAGCGGATATCTGG | CRTC1 | 622 | 777 | 1694 | 1785 | 1679 |
| TGAAGGACCTGAGCCTGACG | CRTC1 | 433 | 680 | 1328 | 1424 | 1439 |
| TGTCAGTGGACAAACATGGA | CRTC1 | 466 | 779 | 1482 | 1494 | 1351 |
| AATTGCGATTGGCCTACACG | CRTC2 | 369 | 614 | 1223 | 1178 | 1251 |
| ACCCACCTTGGCATCAGCGG | CRTC2 | 684 | 656 | 1188 | 1096 | 1127 |
| CCTTACCCCACATCGACCCC | CRTC2 | 374 | 496 | 961 | 896 | 994 |
| GGACTTCACAGGACCGAGGT | CRTC2 | 385 | 550 | 956 | 848 | 914 |
| GTTCATCACACTGGTGTGAA | CRTC2 | 535 | 795 | 1222 | 1270 | 1225 |
| TGGTGAAGGATATGTGACGG | CRTC2 | 432 | 676 | 1355 | 1302 | 1290 |
| AAGATCATCGAGAGCTACGA | CSF1R | 472 | 477 | 676 | 675 | 704 |
| ACCCTCGAGCTCGATCACGT | CSF1R | 326 | 304 | 673 | 620 | 639 |
| GCACCTTCCAAGGCCGAACA | CSF1R | 623 | 591 | 1048 | 1143 | 999 |
| GGCTGTGGACGATCACCTTG | CSF1R | 611 | 1377 | 1870 | 1852 | 1886 |
| TGGATGGTGAAGCCATACCA | CSF1R | 622 | 855 | 1513 | 1397 | 1415 |
| TTCCTGGCCAGAAACACTAG | CSF1R | 715 | 502 | 676 | 707 | 731 |
| AGTGTTTAGAATCTCCAGTG | CSNK1A1 | 393 | 1199 | 1833 | 1827 | 1919 |
| ATCTTTCTCAGGATTAAACC | CSNK1A1 | 399 | 352 | 450 | 424 | 415 |
| CACATCTTGGTATTGAACAG | CSNK1A1 | 423 | 464 | 1018 | 816 | 768 |
| CGGGTTCTTACCGTATGTGG | CSNK1A1 | 329 | 602 | 1353 | 1009 | 1058 |
| CTGGCGATCAACATTACCAA | CSNK1A1 | 488 | 450 | 685 | 630 | 629 |
| TTGCTCTCGTACAGCAACTG | CSNK1A1 | 654 | 514 | 1305 | 1072 | 1154 |
| AGAGGGAGACTACAATGTCA | CSNK1D | 397 | 821 | 1397 | 1356 | 1528 |
| AGCCAGATAACTTTCTCATG | CSNK1D | 388 | 666 | 1359 | 1300 | 1251 |
| CCAGTCAAACACGTAGTCAT | CSNK1D | 427 | 432 | 849 | 829 | 847 |
| CGTACCGCGCCGTCCCAGTG | CSNK1D | 560 | 673 | 1273 | 1149 | 1077 |
| GGGCTTGTCGTCAAAACGCA | CSNK1D | 705 | 534 | 946 | 1005 | 970 |
| TTTACACAGCACTTCAATGG | CSNK1D | 386 | 428 | 770 | 795 | 756 |
| AGCCTGACAACTTCCTCATG | CSNK1E | 544 | 634 | 1257 | 1327 | 1317 |
| AGGCTTGTCGTCGAACCGCA | CSNK1E | 407 | 702 | 1226 | 1293 | 1362 |
| CATCACATTGTAGTCCCCCT | CSNK1E | 477 | 532 | 936 | 956 | 947 |
| GAAGATGTCGACACCGATTG | CSNK1E | 525 | 764 | 1438 | 1103 | 1268 |
| GTAGGGACTCACCATCTGGT | CSNK1E | 332 | 573 | 1128 | 1092 | 1159 |
| GTGCTCATGTACTTCAACCT | CSNK1E | 583 | 386 | 757 | 657 | 644 |
| CATGCTGTGACAATAATCCA | CSNK2A1 | 676 | 604 | 1316 | 1266 | 1239 |
| CTGGGATTATGAGTCACACG | CSNK2A1 | 445 | 824 | 1303 | 1350 | 1447 |
| CTTGGCTATCCTCACCAACT | CSNK2A1 | 736 | 416 | 905 | 738 | 861 |
| GCTCAGGACCTTTGAAGTAT | CSNK2A1 | 445 | 755 | 1197 | 1318 | 1243 |
| GTTTACGTGTTCAAAAACCA | CSNK2A1 | 543 | 483 | 624 | 566 | 538 |
| TCAGCTACGACTAATAGACT | CSNK2A1 | 428 | 327 | 638 | 623 | 648 |
| AACTGGTTCGAAAACTTGGT | CSNK2A2 | 434 | 475 | 1039 | 858 | 948 |
| ACAATGTCCGTGTAGCATCG | CSNK2A2 | 736 | 863 | 1430 | 1387 | 1440 |
| ACTGTGAAGGACCCTGTGGT | CSNK2A2 | 654 | 622 | 1206 | 1200 | 1312 |
| CCAGATGTACGATTACAGCT | CSNK2A2 | 525 | 697 | 983 | 987 | 852 |
| GAAAAAGAAGATAAAACGAG | CSNK2A2 | 490 | 487 | 694 | 653 | 627 |
| TAGCTGCGGCTCATAGACTG | CSNK2A2 | 424 | 915 | 1451 | 1314 | 1441 |
| AAGTGAACCTCACCATCCAA | CTLA4 | 464 | 303 | 648 | 564 | 587 |
| ACACGAAGCTAGCAACACCC | CTLA4 | 582 | 348 | 612 | 593 | 613 |
| CTGTGCCGCGACATACACAG | CTLA4 | 333 | 422 | 798 | 812 | 794 |
| GTCTTCAGGCAACGCAGCCG | CTLA4 | 524 | 468 | 1025 | 815 | 901 |
| TCCTTTATTAGGGATGCATG | CTLA4 | 430 | 646 | 1272 | 1286 | 1411 |
| TGCCTACATAGTAGGGTGGT | CTLA4 | 439 | 753 | 1517 | 1669 | 1650 |
| AAGAAGGTATCTTGAAACTG | CTNNA1 | 473 | 764 | 1215 | 1060 | 1096 |
| ACATGCTAACAAATTAATAG | CTNNA1 | 439 | 294 | 549 | 646 | 657 |
| ACGCCTGGGATGCAGTATAG | CTNNA1 | 421 | 550 | 1084 | 1184 | 1184 |
| CCACGCTTCACAGAAGAACA | CTNNA1 | 479 | 612 | 987 | 1126 | 1074 |
| TCCACTTGCTCGGTAAAACG | CTNNA1 | 483 | 747 | 826 | 1036 | 991 |
| TGACCGGCGTGAGAGAATTG | CTNNA1 | 442 | 1250 | 1874 | 1896 | 1942 |
| ACACTTACGAGAAACTCCTG | CTNNB1 | 269 | 924 | 1342 | 1166 | 1092 |
| ATGCAATGACTCGAGCTCAG | CTNNB1 | 499 | 655 | 1187 | 1131 | 1092 |
| ATGGCAGTGCGTTTAGCTGG | CTNNB1 | 336 | 864 | 1502 | 1496 | 1546 |
| CAGTGGTAAAGGCAATCCCG | CTNNB1 | 451 | 386 | 916 | 789 | 747 |
| CATCTGAGGAGAACGCATGA | CTNNB1 | 561 | 588 | 1142 | 1010 | 1072 |
| GGGCTCAGATGATATAAATG | CTNNB1 | 582 | 580 | 1285 | 1050 | 1191 |
| ACAGGTCAACTTCACCCATG | CUBN | 523 | 809 | 1352 | 1392 | 1397 |
| AGTAGCGTCCCATCCGCCTG | CUBN | 665 | 921 | 1434 | 1563 | 1460 |
| CAGCCAGGGAGATTACTTGG | CUBN | 513 | 664 | 1217 | 1208 | 1187 |
| CCCACAACTCCAATTACCAA | CUBN | 508 | 251 | 376 | 397 | 371 |
| GAATGCAAGATAAATAATGG | CUBN | 503 | 618 | 951 | 979 | 1004 |
| TGTTACCCCTATATGGACTG | CUBN | 589 | 861 | 1337 | 1376 | 1329 |
| AGTATCGAGCCAGCAACTCA | CUL1 | 530 | 751 | 1395 | 1378 | 1382 |
| CCAGTCAAACCAAGCCCGAG | CUL1 | 600 | 336 | 802 | 891 | 783 |
| GAAGACGATGCATTTGCCAA | CUL1 | 452 | 761 | 1178 | 1122 | 1102 |
| GAGACACACATTCATAATCA | CUL1 | 569 | 354 | 614 | 468 | 524 |
| GCACACAAGATGAACTAGCA | CUL1 | 558 | 608 | 980 | 1128 | 1026 |
| GTTCGCCGTGAATGTGACGA | CUL1 | 665 | 623 | 1340 | 1186 | 1288 |
| ATAGCTGGTACATCTGAGTG | CUL4A | 606 | 847 | 1283 | 1524 | 1507 |
| CAGCTGTTTGTAGAGTGTTG | CUL4A | 431 | 1133 | 1889 | 1862 | 1889 |
| GCTGCCGGACAACTACACTC | CUL4A | 372 | 456 | 715 | 705 | 706 |
| GCTGGAAAGGATTCTTGACA | CUL4A | 501 | 642 | 1324 | 1134 | 1076 |
| GGTCTAAATACGTGATTACT | CUL4A | 629 | 459 | 735 | 665 | 667 |
| TGATAGAAGTGTGCTTCCAG | CUL4A | 524 | 610 | 1399 | 1241 | 1301 |
| AAGACTTGTCATCCACTGGT | CUL4B | 455 | 535 | 906 | 824 | 891 |
| AGAACTACACAGATGAAACA | CUL4B | 515 | 300 | 410 | 404 | 392 |
| CTCTTCAGTAGAGTTCGAGG | CUL4B | 604 | 463 | 1052 | 959 | 941 |
| GAGTATATTAATCTCTTCTG | CUL4B | 513 | 698 | 1066 | 1041 | 1096 |
| GCTTCTTCTGTATCGGTACG | CUL4B | 652 | 920 | 1760 | 1863 | 1767 |
| TTGGCATCCTTACCAAATGG | CUL4B | 349 | 728 | 1424 | 1431 | 1488 |
| AAACGAAGAAATGGGTCCGA | CUX1 | 481 | 389 | 762 | 731 | 772 |
| ACTGCTGAAGAGCTTCCAAG | CUX1 | 623 | 575 | 975 | 985 | 1028 |
| CATCTGTCAAAGAATCTTCG | CUX1 | 490 | 524 | 675 | 611 | 654 |
| GCGGGAGATCGCCCAACTGG | CUX1 | 583 | 706 | 1262 | 1103 | 1157 |
| TAGCTGGACACAGACGACAT | CUX1 | 400 | 554 | 793 | 718 | 775 |
| TCAATTATCCCGCAACACGG | CUX1 | 849 | 403 | 477 | 411 | 395 |
| CAGAACAGCATGATCATCAG | CXCR2 | 444 | 524 | 818 | 725 | 799 |
| CAGGTTCAGCAGATAGACGT | CXCR2 | 447 | 598 | 1200 | 1055 | 1136 |
| CGTGCGTGTGGCATGGACAA | CXCR2 | 549 | 770 | 1357 | 1286 | 1304 |
| CTGTCTCCAAGGTAAAGGGT | CXCR2 | 673 | 695 | 1228 | 1241 | 1203 |
| GGACATCAACAAGTATGCTG | CXCR2 | 336 | 477 | 908 | 802 | 777 |
| TGATAATTATACTTACAACA | CXCR2 | 544 | 392 | 410 | 396 | 345 |
| AAGATGATGGAGTAGACTGT | CXCR4 | 478 | 638 | 1192 | 1049 | 1102 |
| ACAGGATGACGATACCCGGC | CXCR4 | 548 | 557 | 686 | 648 | 682 |
| AGGCGACTATGACTCCATGA | CXCR4 | 563 | 689 | 1191 | 1056 | 1073 |
| CATGGACAATAGCCAAGTAC | CXCR4 | 229 | 570 | 1008 | 876 | 979 |
| GCGGAAAAGGTGGTCTACGT | CXCR4 | 572 | 374 | 614 | 554 | 529 |
| GCTTCTGGTAGCCCTTCGAG | CXCR4 | 514 | 995 | 1388 | 1424 | 1484 |
| ACTGTGGCCAATACTTCCAT | CYLD | 769 | 529 | 1038 | 1128 | 989 |
| CGGATTACCCCATAGAAAGG | CYLD | 399 | 485 | 1043 | 1060 | 1090 |
| CTCCAGTCATAATAAACCAA | CYLD | 468 | 311 | 478 | 420 | 425 |
| GAGGACCCTTATTAGCAGAG | CYLD | 632 | 504 | 887 | 921 | 985 |
| GGATTGGAAAGTGATTATGC | CYLD | 679 | 581 | 952 | 901 | 1126 |
| GTATATCCAAGATCGTTCTG | CYLD | 537 | 799 | 1405 | 1366 | 1453 |
| AGTCCACTTCACCACAGACG | CYP17A1 | 389 | 607 | 1124 | 1201 | 1219 |
| AGTCTTGGTACCCATACGAA | CYP17A1 | 451 | 422 | 908 | 855 | 918 |
| ATATGATAGACATATTCCCC | CYP17A1 | 573 | 359 | 614 | 524 | 562 |
| GATCAGTATGTCTAGCATGT | CYP17A1 | 646 | 670 | 1320 | 1200 | 1217 |
| GGTATCGCTTTCGCAGACCA | CYP17A1 | 580 | 695 | 1220 | 1133 | 1186 |
| TATGGACTTTCCATTCTGGG | CYP17A1 | 637 | 672 | 1280 | 1363 | 1279 |
| AAACTGTTACCATACGCACG | CYP19A1 | 463 | 677 | 1268 | 1187 | 1271 |
| AACTACTACAACAAGATGTA | CYP19A1 | 427 | 751 | 1221 | 1134 | 1187 |
| AGTCACTACAGCTCCCGATT | CYP19A1 | 558 | 538 | 947 | 861 | 844 |
| ATTCTCAAGTACCGTCCAAG | CYP19A1 | 518 | 607 | 896 | 986 | 991 |
| CTCAGCAAAAATCAATTCAG | CYP19A1 | 607 | 354 | 530 | 520 | 482 |
| GAAGACACATCCTCAATACC | CYP19A1 | 548 | 387 | 669 | 762 | 695 |
| ACGCCTTCATCCTCTCGGCG | CYP1B1 | 424 | 518 | 1012 | 903 | 971 |
| ACGCGAAAGGAAGCGAAGCG | CYP1B1 | 545 | 551 | 1084 | 1084 | 989 |
| CTGCTCGAACTCGCGGAAGG | CYP1B1 | 451 | 943 | 1286 | 1437 | 1549 |
| GACGTGTTCCAGATCCGCCT | CYP1B1 | 575 | 327 | 531 | 512 | 508 |
| GCACACGGCGCTCATGACGT | CYP1B1 | 637 | 492 | 1064 | 899 | 943 |
| GCCGCGGCGTTTCCGATCAG | CYP1B1 | 479 | 900 | 1532 | 1545 | 1550 |
| AAACTGTCAGGGAAAAACGC | CYP2C18 | 637 | 247 | 407 | 405 | 458 |
| AGAAGGAAAGTAGGATCACA | CYP2C18 | 395 | 681 | 1330 | 1323 | 1356 |
| CAATAGTCCGTATCTCAGGG | CYP2C18 | 511 | 712 | 1381 | 1404 | 1366 |
| CTCTAAACTTTAGTAACCCT | CYP2C18 | 423 | 387 | 796 | 794 | 704 |
| GGAATCAAAATCGTGCCAAA | CYP2C18 | 513 | 404 | 557 | 460 | 472 |
| TACCCGTACAACACCACGGT | CYP2C18 | 509 | 268 | 609 | 455 | 475 |
| AAACAGACACTCATAAAGGT | CYP3A4 | 467 | 302 | 631 | 507 | 541 |
| AGTATGGAGATGTGTTGGTG | CYP3A4 | 554 | 960 | 1649 | 1686 | 1740 |
| ATCCCACTGGACCAAAAGAC | CYP3A4 | 458 | 462 | 690 | 648 | 761 |
| ATTTAATACTTCAAAAACTG | CYP3A4 | 544 | 329 | 426 | 356 | 376 |
| CAACCCACAAGATCCCTTTG | CYP3A4 | 392 | 801 | 1596 | 1622 | 1575 |
| GTATGGGACCTACACACATG | CYP3A4 | 500 | 817 | 1409 | 1454 | 1549 |
| AATCCAAGGAGGCTTTGAGG | DAXX | 636 | 470 | 929 | 950 | 962 |
| ACGGGTCCTTGGTGCCTCAG | DAXX | 495 | 642 | 1377 | 1377 | 1501 |
| CCCGGGATAGGACCCGAGAG | DAXX | 422 | 745 | 1137 | 1109 | 1079 |
| CGGCAAGAAATGCTACAAGT | DAXX | 479 | 458 | 653 | 489 | 534 |
| GAGCAGCGCATTCCCTATCG | DAXX | 596 | 861 | 1616 | 1510 | 1408 |
| GTAACCTGATGCCCACGTCT | DAXX | 607 | 510 | 889 | 734 | 764 |
| AGAACGGGAACAGCAGATCG | DBN1 | 392 | 884 | 1660 | 1702 | 1693 |
| ATGCAGCTGTCGAAATGAAG | DBN1 | 664 | 624 | 1024 | 932 | 975 |
| ATGTGCTTGTGCTAGCCACG | DBN1 | 474 | 645 | 1455 | 1363 | 1417 |
| CAGCAGCCATTATTGCCCAG | DBN1 | 483 | 558 | 1174 | 1129 | 1159 |
| CTGCCCTTTCATAAAGGCAT | DBN1 | 669 | 729 | 944 | 898 | 879 |
| CTGTTGCTGAACAGATTGAG | DBN1 | 494 | 837 | 1516 | 1457 | 1456 |
| AATTGTGTGTCACAATCGTG | DDB1 | 508 | 27 | 31 | 33 | 43 |
| ATAGTGGCTACTCTCAAAAG | DDB1 | 408 | 602 | 1039 | 1000 | 970 |
| CATTGGAAAAGACCAACTTG | DDB1 | 572 | 777 | 1660 | 1495 | 1555 |
| GATATCATAACGCGAGCCCA | DDB1 | 687 | 440 | 1049 | 821 | 856 |
| GCGGCATGTGAAAACTTACG | DDB1 | 572 | 521 | 977 | 1021 | 1033 |
| TACCAATTTAGGACCCATTG | DDB1 | 296 | 717 | 1372 | 1290 | 1320 |
| ACTGCTCACCCCTTTAATGA | DDB2 | 629 | 430 | 819 | 800 | 666 |
| CCAGGAACCAATCACAGCAT | DDB2 | 566 | 462 | 889 | 959 | 946 |
| GCGCTCCACCACCATAAGCT | DDB2 | 535 | 575 | 1021 | 788 | 819 |
| TCTAGCAGAAGATTTGACTC | DDB2 | 563 | 425 | 822 | 731 | 808 |
| TGGTCACAGGAGACAATGTG | DDB2 | 352 | 726 | 1326 | 1221 | 1262 |
| TGTGGCCCTCCTGTCAAAAG | DDB2 | 522 | 693 | 1082 | 1092 | 1093 |
| ACTGTCCTCAGATGAAAATG | DDIT3 | 575 | 905 | 1540 | 1530 | 1547 |
| AGGAAGACCAAAAAAGACCT | DDIT3 | 501 | 469 | 842 | 781 | 858 |
| ATCAAAAACCTTCACCACTC | DDIT3 | 501 | 228 | 412 | 386 | 420 |
| CTCACCTCCTTGTTTCCAGG | DDIT3 | 543 | 773 | 1359 | 1143 | 1272 |
| CTCTCTGGCTTGGCTGACTG | DDIT3 | 526 | 702 | 1198 | 1180 | 1244 |
| TGACTGGAATCTGGAGAGTG | DDIT3 | 613 | 823 | 1469 | 1375 | 1344 |
| ACTTACGCCCGTCCCCCTCG | DDR1 | 382 | 919 | 1645 | 1719 | 1736 |
| ATTTCACTGAAGAGTAACCA | DDR1 | 436 | 449 | 808 | 909 | 793 |
| CAGGCTATGACTATGTGGGA | DDR1 | 435 | 543 | 1184 | 1227 | 1212 |
| GCAGACAGACACTCATGACC | DDR1 | 599 | 471 | 748 | 648 | 736 |
| TCAGTTCCTCTTCTAACACC | DDR1 | 537 | 464 | 578 | 513 | 481 |
| TGCAGTACTGAGATTCCAGT | DDR1 | 438 | 658 | 900 | 924 | 929 |
| AAGTCGATGACAGCAACACT | DDR2 | 517 | 536 | 824 | 817 | 812 |
| AGACTCACGAATACCACGTG | DDR2 | 682 | 2 | 11 | 11 | 10 |
| CGGACAAATCTGGCCACGAT | DDR2 | 550 | 289 | 815 | 824 | 869 |
| GCTCACTTGGCGATGAGGAG | DDR2 | 652 | 961 | 1510 | 1592 | 1557 |
| GGGGGCGCCACACTACGCAG | DDR2 | 308 | 465 | 907 | 931 | 848 |
| TGGGGGCAAACTCAATCCCA | DDR2 | 408 | 696 | 902 | 886 | 885 |
| AAGAGGTGGAGACATACAGA | DDX5 | 476 | 383 | 858 | 778 | 791 |
| AAGTCCACTTGCATCTATGG | DDX5 | 551 | 573 | 1179 | 1108 | 1112 |
| AGAAATTAGTCAAAAAGAAG | DDX5 | 590 | 437 | 561 | 380 | 364 |
| CTGATAGGCAAACCCTAATG | DDX5 | 429 | 803 | 1518 | 1378 | 1461 |
| GTCACCATGGATACCCATGG | DDX5 | 636 | 508 | 1090 | 1084 | 1155 |
| TGATTGATATGGACGATAGC | DDX5 | 427 | 568 | 1214 | 1058 | 1037 |
| ATAATGAGGCTTGATGACAC | DDX6 | 393 | 245 | 575 | 442 | 499 |
| GAACACTTACTTAATAGTGG | DDX6 | 438 | 463 | 860 | 646 | 658 |
| GACAGCAACTTATCTGCCTA | DDX6 | 531 | 571 | 1230 | 1054 | 1117 |
| GCCGTTCAAGTAAGGGAATG | DDX6 | 577 | 614 | 1465 | 1246 | 1217 |
| TACACTAAGAGGGAAAGTAG | DDX6 | 640 | 478 | 884 | 811 | 852 |
| TTGAAACGGGAGTTACTGAT | DDX6 | 430 | 555 | 884 | 814 | 872 |
| AGCAGCCCGCTCTATAGTCG | DEPDC5 | 515 | 913 | 1887 | 2035 | 1840 |
| AGCGAGACTCTAGCGCACCA | DEPDC5 | 474 | 557 | 1296 | 1231 | 1306 |
| CATCTGCAAACCTTTCATGG | DEPDC5 | 433 | 743 | 1547 | 1589 | 1606 |
| GAGGTAGAATTGTCGCCTTG | DEPDC5 | 457 | 624 | 1243 | 1342 | 1396 |
| GGATCAATATATTGGCCGTG | DEPDC5 | 423 | 652 | 1105 | 1084 | 1168 |
| TCGCCCAGACGAGAATCACG | DEPDC5 | 412 | 430 | 879 | 865 | 873 |
| CTCCTTGTGGAGGTTCCCTA | DHFR | 566 | 655 | 1289 | 1322 | 1228 |
| GACATGGTTTGGATAGTGGG | DHFR | 536 | 560 | 1320 | 1056 | 981 |
| TAAAGCATCATCCAGACTTT | DHFR | 616 | 723 | 1331 | 1248 | 1329 |
| TACCTTAGGGAACCTCCACA | DHFR | 465 | 406 | 685 | 653 | 671 |
| TGAGAAGAATCGACCGTTAA | DHFR | 532 | 695 | 1364 | 1256 | 1255 |
| TTGGTTACAGTACTTACTTG | DHFR | 350 | 729 | 1598 | 1398 | 1337 |
| AAGACCATGGCACAAAACTG | DIABLO | 324 | 535 | 894 | 808 | 816 |
| ACAGCGGTACCTGAACAGGG | DIABLO | 486 | 511 | 731 | 723 | 675 |
| GATCATTGGAGCCAGAGTTG | DIABLO | 519 | 917 | 1414 | 1544 | 1602 |
| TGCTTCAATCAATGCATATG | DIABLO | 457 | 817 | 1697 | 1461 | 1481 |
| TGGGAAGATGAATTCACAGG | DIABLO | 548 | 550 | 942 | 1110 | 1085 |
| TTAGTAACGATGCATTAATG | DIABLO | 488 | 955 | 1512 | 1574 | 1470 |
| CATAAGAAACCGACTACCTG | DICER1 | 612 | 688 | 1228 | 1406 | 1371 |
| CATACCTGTCCAATACCACC | DICER1 | 602 | 231 | 364 | 358 | 367 |
| TAAAATCCACGGGAAGTGAT | DICER1 | 494 | 652 | 923 | 1139 | 1084 |
| TACAAATTTCAGGTCAAGTG | DICER1 | 453 | 621 | 857 | 834 | 903 |
| TCAGCTACATAGAATCGATG | DICER1 | 503 | 467 | 819 | 778 | 805 |
| TGATATACCAAAATGCAACT | DICER1 | 273 | 209 | 346 | 370 | 332 |
| AATTCGAATAGAAACCAGAC | DIS3 | 431 | 210 | 395 | 399 | 339 |
| CCTTCACTAATGAGCACCAT | DIS3 | 669 | 564 | 1228 | 1117 | 1144 |
| GATATTACCACTAGTCTCCG | DIS3 | 354 | 411 | 948 | 722 | 742 |
| GTGTCGGGAAGCAAGTAGTG | DIS3 | 355 | 576 | 1276 | 1065 | 1015 |
| TCATCTATATCCGTACAGCC | DIS3 | 576 | 202 | 428 | 447 | 447 |
| TTTACATAGATAATCTTACA | DIS3 | 406 | 377 | 507 | 567 | 470 |
| CAGTATGCACAGCTTTACGA | DLC1 | 616 | 371 | 581 | 538 | 681 |
| CCAGCACACCAACGAACCTG | DLC1 | 414 | 826 | 1365 | 1398 | 1369 |
| CGTTGTCATAGATACTCAGG | DLC1 | 301 | 625 | 1026 | 962 | 926 |
| CTGATGACCGAGTTGGTCCG | DLC1 | 484 | 974 | 1892 | 1761 | 1779 |
| GAGTCACATGCACTTATGTG | DLC1 | 411 | 831 | 1370 | 1509 | 1492 |
| GTGGAGATCTCCGCGCTCAA | DLC1 | 445 | 627 | 1125 | 931 | 932 |
| AACGAATGCATCCCCCACAA | DLL4 | 287 | 416 | 845 | 789 | 776 |
| AGACTCCCCTTGCTTCAATG | DLL4 | 529 | 820 | 1639 | 1739 | 1661 |
| AGCAACCCCTGTCGCAATGG | DLL4 | 571 | 771 | 1347 | 1288 | 1461 |
| AGCAGACCAGCCCTCCCACG | DLL4 | 431 | 720 | 1226 | 1130 | 1277 |
| GAGGCAACAAGACTCACCAT | DLL4 | 549 | 379 | 688 | 695 | 754 |
| TGTTGCCCCATTCTTGCACG | DLL4 | 558 | 698 | 1434 | 1394 | 1365 |
| ATGGACTTGACATCCCATCA | DMD | 391 | 669 | 1020 | 1042 | 1071 |
| CAGGGTATTGAATGCAACTG | DMD | 614 | 652 | 1247 | 1328 | 1270 |
| CCGAACAACTGAACAGCCGG | DMD | 400 | 534 | 1063 | 1113 | 1038 |
| GCAAACCTGATGGCAAACCG | DMD | 449 | 591 | 1039 | 1152 | 1189 |
| TAAAGGGCAGCATTTGTACA | DMD | 528 | 774 | 1369 | 1476 | 1408 |
| TTCCGACTAATATCAGACAG | DMD | 557 | 757 | 1229 | 1198 | 1095 |
| CCCTTAGGAACCTACCCTGG | DNM1 | 481 | 656 | 1371 | 1203 | 1243 |
| GGACCTTGGTCATTCCGGGT | DNM1 | 618 | 534 | 937 | 856 | 934 |
| GGTTAGACAGTGCACCAAGA | DNM1 | 497 | 604 | 1101 | 972 | 918 |
| GTAGGTAGGGTGTGCCCATG | DNM1 | 511 | 884 | 1917 | 1766 | 1815 |
| GTGACAATGCCTGACCCTCG | DNM1 | 363 | 651 | 1355 | 1400 | 1424 |
| TTGACACCTATGAACTGTCA | DNM1 | 435 | 582 | 1032 | 1071 | 1114 |
| GAACGGCATCCAGTATCGAG | DNMT1 | 445 | 364 | 754 | 606 | 663 |
| GCCTAAACCGAAGTCTAAGG | DNMT1 | 532 | 389 | 585 | 533 | 578 |
| GTCCGCATGGTAACTTGCTG | DNMT1 | 366 | 744 | 1693 | 1574 | 1623 |
| TCACCCAAAAAAATGCACCA | DNMT1 | 347 | 188 | 310 | 235 | 233 |
| TCGGGACTTACTGCGTCAGG | DNMT1 | 403 | 822 | 1603 | 1283 | 1296 |
| TTGGCCCCATAAATGAATGG | DNMT1 | 357 | 578 | 1116 | 1112 | 1082 |
| AGACATGTGGGTCGAACCCG | DNMT3A | 406 | 905 | 1480 | 1452 | 1508 |
| AGCATCCAGGCAATGCATGG | DNMT3A | 520 | 1146 | 1642 | 1875 | 1822 |
| AGTGTGTGGATCTCTTAGTG | DNMT3A | 632 | 1068 | 2172 | 2010 | 2024 |
| CCGGGAACAACTTCCCCGCG | DNMT3A | 665 | 547 | 1099 | 1129 | 1074 |
| CGACGATGAGCCAGAGTACG | DNMT3A | 425 | 811 | 1388 | 1410 | 1324 |
| CTCAGAGCTGTTACCCAACG | DNMT3A | 490 | 682 | 1270 | 1280 | 1278 |
| ACAGCTCAAGGCTCGGTGTG | DOT1L | 665 | 937 | 1986 | 1900 | 2053 |
| CTGTGCGACCAAGTCAAAGG | DOT1L | 617 | 432 | 659 | 594 | 651 |
| GAGAGCGGATTGCCAACACA | DOT1L | 616 | 883 | 1438 | 1384 | 1335 |
| GCTCCCCGATGCCAACAAAG | DOT1L | 692 | 769 | 1089 | 1027 | 904 |
| GGCGGGAAGACCGACGTCCG | DOT1L | 400 | 776 | 1500 | 1537 | 1486 |
| TGTCATCTGTAAGGCCACGG | DOT1L | 569 | 528 | 1046 | 982 | 950 |
| ACAACGAAGAGCTTTACGGT | DPP4 | 318 | 643 | 1266 | 1231 | 1168 |
| CATAGGAGCAGCGACGATAA | DPP4 | 327 | 920 | 1193 | 1156 | 1233 |
| CTTAGAGTACAACTATGTGA | DPP4 | 415 | 539 | 845 | 879 | 790 |
| GAATACTCACAGGTAATCAC | DPP4 | 516 | 222 | 288 | 315 | 294 |
| GTGGATCACATGGTCACCAG | DPP4 | 452 | 591 | 1050 | 1027 | 1021 |
| TGAAATGAGTACCACCGGCT | DPP4 | 380 | 415 | 807 | 758 | 747 |
| ACACGATAAATTTAACTCCA | DPYD | 409 | 328 | 516 | 453 | 460 |
| AGTCCGTCTGAAAGCCGATG | DPYD | 454 | 919 | 1603 | 1535 | 1454 |
| ATTCCACAGGTCAAACCAAG | DPYD | 543 | 370 | 566 | 586 | 605 |
| GCGGGCTCCACAACATAGAG | DPYD | 536 | 790 | 1359 | 1224 | 1210 |
| GTTCAGGCTACCATATGATG | DPYD | 574 | 598 | 1115 | 1120 | 1183 |
| TTTGTCCAGGGCCATACATG | DPYD | 396 | 811 | 1194 | 1307 | 1326 |
| ACCTCCAGGTAGACAACCCA | DRD2 | 454 | 435 | 1137 | 1123 | 1146 |
| ATGGCGTAGTAGTTGTAGTG | DRD2 | 259 | 1026 | 1388 | 1593 | 1605 |
| GGAGGAGTACACCACGAATG | DRD2 | 740 | 834 | 1659 | 1584 | 1512 |
| GGAGTTTCCCAGTGAACAGG | DRD2 | 491 | 624 | 1161 | 989 | 996 |
| GGGTGAGTGCAGTTGCCCTG | DRD2 | 588 | 1081 | 2164 | 1916 | 1888 |
| TAGCGGGTGTTGTAGAGCAT | DRD2 | 397 | 649 | 1319 | 1070 | 1113 |
| AGAAGAGAGAGGATTCCCGC | EDNRB | 389 | 363 | 722 | 643 | 615 |
| CAACACAGTGATTCCCACGG | EDNRB | 512 | 605 | 1113 | 1077 | 1022 |
| CCTCCGCAGATGCCTAAGGG | EDNRB | 864 | 627 | 1266 | 1213 | 1198 |
| GAGTCGAATTAAAGGAATTG | EDNRB | 579 | 738 | 1118 | 1095 | 1037 |
| GGTTATCATATCAAAACCCA | EDNRB | 411 | 350 | 755 | 733 | 724 |
| TTTCAAGTATATCAACACGG | EDNRB | 360 | 412 | 750 | 622 | 622 |
| AAGAGAGTGATCCATACCAC | EED | 630 | 324 | 583 | 579 | 565 |
| ATAACCCAAATAAAACTAAC | EED | 374 | 240 | 300 | 298 | 303 |
| CAAGTGTGTAAATAGTCTCA | EED | 409 | 534 | 1312 | 1308 | 1203 |
| CCGGTTGTTGCAGTCTTATG | EED | 526 | 884 | 1782 | 1579 | 1664 |
| CTGGATATTCCACAATCGTA | EED | 524 | 663 | 996 | 973 | 965 |
| GATGCTGTCAGTATAGAAAG | EED | 579 | 660 | 1081 | 908 | 988 |
| AAAAACTGCACGTCGATCAG | EGFR | 532 | 709 | 1263 | 1118 | 1274 |
| ACCAGGCAGTCGCTCTCGCG | EGFR | 495 | 459 | 916 | 841 | 922 |
| AGCACAATGGATGGCAGACG | EGFR | 628 | 646 | 1211 | 1110 | 1097 |
| AGGCGCGGACACATGAACCG | EGFR | 643 | 434 | 838 | 797 | 842 |
| ATCCGAAAACTTCCAGACCA | EGFR | 382 | 396 | 828 | 763 | 771 |
| GAGAACCTAGAAATCATACG | EGFR | 532 | 467 | 778 | 601 | 657 |
| ACTGACCGAACCCAGAAGGT | EHMT1 | 526 | 508 | 1067 | 1058 | 1092 |
| AGACCAAAGTATTAAAACAG | EHMT1 | 579 | 298 | 537 | 476 | 522 |
| AGGGCAGCGTCCCCTTGACA | EHMT1 | 473 | 787 | 1397 | 1338 | 1485 |
| GCTGGCATTATTGACTCGAG | EHMT1 | 512 | 488 | 923 | 885 | 891 |
| GGCGTCTGAGAAAAAGCACT | EHMT1 | 545 | 443 | 621 | 604 | 625 |
| TGCAGACCTGAATCAGCATG | EHMT1 | 500 | 795 | 1441 | 1333 | 1277 |
| AGTGCAAGTCATCACTCATG | EHMT2 | 508 | 778 | 1694 | 1582 | 1474 |
| CACACGCTCATCCACAGAGT | EHMT2 | 320 | 361 | 776 | 610 | 612 |
| GAAGACCATGCCATTCAGCT | EHMT2 | 310 | 513 | 929 | 838 | 901 |
| GCAATTACCAGCGTGGTTGG | EHMT2 | 545 | 910 | 1471 | 1606 | 1592 |
| TCTTTGGGGGACACCCCTCG | EHMT2 | 465 | 803 | 1559 | 1632 | 1715 |
| TGTAGCGAGCTACCTCCAAG | EHMT2 | 431 | 684 | 1334 | 1291 | 1199 |
| CCTCCAAGGCTGTTGCAAGG | EIF3G | 506 | 579 | 1190 | 1094 | 1030 |
| GAACCTGAGTTAGTGCCGGG | EIF3G | 630 | 341 | 791 | 728 | 696 |
| GACAGAGTACAAGATAGATG | EIF3G | 669 | 555 | 1069 | 932 | 1034 |
| GACCCGATGAACAAGCTCAA | EIF3G | 312 | 305 | 745 | 533 | 498 |
| GGACACGTCGTCACTGACTG | EIF3G | 583 | 660 | 1188 | 1080 | 1029 |
| TCACCAGCGAGCTCCTCAAG | EIF3G | 466 | 409 | 832 | 785 | 842 |
| AAGAACGCTTAAAAAAAGAA | EIF5B | 515 | 227 | 375 | 341 | 309 |
| TACATGTGTGTGACGAAGCT | EIF5B | 496 | 534 | 1152 | 1008 | 1090 |
| TCCGGTATCAAGAGTCACTT | EIF5B | 591 | 697 | 956 | 1059 | 1102 |
| TGCAGAAATCAAATAAAAGG | EIF5B | 375 | 312 | 442 | 358 | 360 |
| TGTCCGATGAGAAGGATGCA | EIF5B | 473 | 505 | 1168 | 914 | 896 |
| TTAGTTGTTGACATTATGCA | EIF5B | 380 | 809 | 1409 | 1409 | 1360 |
| AGAAGGGAGACCCTAAGCAC | ELF3 | 516 | 420 | 792 | 707 | 708 |
| AGACTCCGGTGGAAGTGACG | ELF3 | 598 | 631 | 1192 | 1088 | 1110 |
| CAGTGCTATGTACAGCTCCG | ELF3 | 449 | 806 | 1646 | 1628 | 1641 |
| GGACTGGATCAGCTACCAAG | ELF3 | 813 | 624 | 797 | 753 | 836 |
| GGGGGCGCAGTTGCACAGCG | ELF3 | 649 | 500 | 1104 | 973 | 1068 |
| TGATCCAGCTGAGCTCGTCG | ELF3 | 369 | 508 | 1076 | 947 | 942 |
| AAAGCATATTCATTCTGTGG | ENPP3 | 509 | 540 | 1075 | 966 | 959 |
| AAATCATTATACTATTGTCA | ENPP3 | 355 | 505 | 733 | 642 | 657 |
| ATTACGAGCCCTGATACGAG | ENPP3 | 625 | 493 | 1062 | 861 | 852 |
| GACTACAAGAGTGTCTGCCA | ENPP3 | 473 | 647 | 953 | 916 | 944 |
| GCTTGCATTCCATATCACAC | ENPP3 | 657 | 259 | 404 | 376 | 351 |
| TTGTAATGGATATATTTGGA | ENPP3 | 389 | 761 | 1305 | 1140 | 1170 |
| AAGAACTAGAAGAAAAACGA | EP300 | 556 | 390 | 500 | 382 | 375 |
| AATATGCAGTACCCAAACCC | EP300 | 532 | 301 | 551 | 569 | 558 |
| ATTGACTTCTGTGCTACTAG | EP300 | 536 | 572 | 853 | 731 | 807 |
| CTCCCATAGGACTAGCACCT | EP300 | 594 | 648 | 1112 | 960 | 1063 |
| GCATTATAGGAGAGTTCACG | EP300 | 688 | 692 | 1107 | 1101 | 1150 |
| TTCTTCATCGTGCGGCAGTG | EP300 | 477 | 742 | 1308 | 1178 | 1140 |
| AAATGACCGGCACAAAGTCT | EPCAM | 513 | 474 | 941 | 985 | 980 |
| AGTGTTCACACACCAGCACG | EPCAM | 639 | 647 | 1394 | 1402 | 1487 |
| CCCCACGAGCCTTACACGAG | EPCAM | 511 | 667 | 1200 | 1144 | 1171 |
| CGCAGGCGGTGACTCACGCG | EPCAM | 469 | 456 | 1013 | 888 | 883 |
| GGCAGCACGACTCCGAACCT | EPCAM | 529 | 589 | 1177 | 1051 | 1091 |
| TTCGAATATCGGAAAAGGCG | EPCAM | 411 | 586 | 959 | 949 | 915 |
| ACAGCGCAGCTGAGTCCGAG | EPHA1 | 560 | 831 | 1427 | 1249 | 1260 |
| AGGCCCAGAACGGAGTGTCG | EPHA1 | 336 | 906 | 1453 | 1339 | 1309 |
| CGTGTCGCCGAAGTGAACCG | EPHA1 | 534 | 738 | 1445 | 1402 | 1449 |
| CTCCAACTGGATCTACCGCG | EPHA1 | 590 | 550 | 992 | 1108 | 1072 |
| TCGAACCCTGACAATGTACG | EPHA1 | 414 | 697 | 1143 | 1153 | 1146 |
| TGGCGCTGTGAAACTGAACG | EPHA1 | 361 | 484 | 941 | 976 | 958 |
| CAACCTCTACTACGCCGAGT | EPHA2 | 570 | 808 | 1072 | 935 | 916 |
| CACGCACCCATATGGCAAAG | EPHA2 | 537 | 1104 | 1310 | 1282 | 1262 |
| GAAGCCCCTGAAGACATACG | EPHA2 | 386 | 1017 | 1513 | 1419 | 1388 |
| GCAAAGGTGTAGTTCATGTG | EPHA2 | 745 | 1320 | 1948 | 1883 | 1710 |
| GCACGGGGCACTCCAAACAG | EPHA2 | 344 | 774 | 1074 | 1012 | 1094 |
| TCACGGAGAAGCCCTCAGTG | EPHA2 | 468 | 1368 | 2186 | 2190 | 2124 |
| ACACTTCATGTTACCGTCCG | EPHA3 | 547 | 717 | 1040 | 1023 | 961 |
| CGCCGCGCGAGATCCCACGC | EPHA3 | 484 | 292 | 724 | 511 | 551 |
| CTACAACAAGTCGAAGCCCG | EPHA3 | 397 | 552 | 941 | 1052 | 994 |
| CTCAGCTCAGAAGATCTACG | EPHA3 | 510 | 607 | 992 | 1043 | 995 |
| TGGGATCATATTGGACTACG | EPHA3 | 621 | 681 | 1296 | 1414 | 1342 |
| TTGAGGACTTCTCGCGAAGG | EPHA3 | 317 | 647 | 1348 | 1355 | 1284 |
| AACACCGAGATCCGGGATGT | EPHA4 | 687 | 777 | 1313 | 1166 | 1235 |
| ACATCGTGTACCTGTGACCG | EPHA4 | 609 | 746 | 1345 | 1564 | 1621 |
| CCAGGATATTTCCTACAATG | EPHA4 | 528 | 575 | 998 | 902 | 948 |
| GAATTTGGCGAAGTATGCAG | EPHA4 | 535 | 466 | 878 | 863 | 876 |
| GGAGCCTCGAACCTCCACCA | EPHA4 | 400 | 679 | 1208 | 1245 | 1132 |
| TGACGACCAACCAAGCAGGT | EPHA4 | 566 | 284 | 638 | 668 | 560 |
| ATTAAAATGACTTACACACT | EPHA5 | 622 | 557 | 827 | 759 | 807 |
| CAGCTCCAGTGATTGTGTCA | EPHA5 | 652 | 956 | 1540 | 1577 | 1549 |
| GCAGGTGTGTGTGACGAGTG | EPHA5 | 500 | 828 | 1497 | 1228 | 1261 |
| TCCTCATGGGTATAACTGTG | EPHA5 | 347 | 776 | 1510 | 1468 | 1498 |
| TGAGAACGGGAGAAACATCA | EPHA5 | 347 | 490 | 793 | 840 | 867 |
| TGGATGGGCAACAGCGCACA | EPHA5 | 365 | 921 | 1481 | 1724 | 1657 |
| ACTTAACACAGAAATTCGTG | EPHA6 | 469 | 660 | 992 | 1112 | 1171 |
| AGAACAAAGTTAGAATGACG | EPHA6 | 544 | 453 | 773 | 792 | 868 |
| CTTCCATGTAGGTTAAACTG | EPHA6 | 380 | 701 | 1117 | 1135 | 1152 |
| GAACTTACCATCTTGATCCG | EPHA6 | 470 | 768 | 1044 | 1008 | 1058 |
| TCGCATTGAGAGAGTCATTG | EPHA6 | 403 | 702 | 1266 | 1394 | 1457 |
| TTCACCCACAGGCCTACCCA | EPHA6 | 616 | 707 | 1116 | 1016 | 1028 |
| AGCTGTCCATCAATTCGCCA | EPHA7 | 762 | 585 | 957 | 1076 | 986 |
| CCGCTTACACAATATTCTGT | EPHA7 | 524 | 368 | 609 | 631 | 656 |
| CTCCTCAAGATGTGAATGTG | EPHA7 | 565 | 775 | 1636 | 1680 | 1623 |
| CTGACTTAAGCCGATCCCAG | EPHA7 | 490 | 513 | 966 | 986 | 982 |
| GTGTAGCAACGTCAAGTCTG | EPHA7 | 570 | 778 | 1504 | 1410 | 1408 |
| TTTCGAAAGGCAACGCACAG | EPHA7 | 335 | 580 | 717 | 797 | 784 |
| CCGGGGAAGTCTGCTACGGG | EPHA8 | 535 | 542 | 1074 | 973 | 983 |
| CTTCTACAAGTCGGCCCCTG | EPHA8 | 622 | 993 | 1484 | 1503 | 1543 |
| GAGAAGATGCAGTATCAGAA | EPHA8 | 335 | 476 | 899 | 831 | 949 |
| GCAGTGGGACTCCATCAACG | EPHA8 | 408 | 755 | 1514 | 1399 | 1306 |
| GGCACTTCTTGTAATAGATG | EPHA8 | 498 | 943 | 1727 | 1619 | 1764 |
| GTTGACGGCCTCGATCCAGA | EPHA8 | 374 | 532 | 892 | 847 | 839 |
| ACTCCCGGACAGCTTCGTTG | EPHB1 | 459 | 975 | 1408 | 1480 | 1445 |
| AGAGATGCGCTTCACTGTGA | EPHB1 | 446 | 1074 | 1478 | 1585 | 1556 |
| CTCCGTAATCCTGGAAAGCG | EPHB1 | 397 | 581 | 1013 | 1207 | 1231 |
| GACATGTATCCCCAACGCAG | EPHB1 | 702 | 334 | 656 | 549 | 527 |
| TCGGACCGGCTATTACCGAG | EPHB1 | 440 | 761 | 1398 | 1231 | 1338 |
| TTGACGGAGACATGCTGTGG | EPHB1 | 264 | 1300 | 1861 | 1782 | 1726 |
| AACATCAAGACACTGCGTCT | EPHB4 | 427 | 466 | 923 | 879 | 907 |
| AGCCCACGAACTGCCACCCA | EPHB4 | 598 | 641 | 1146 | 1214 | 1141 |
| GCTGAGTACTCGGACAAGCA | EPHB4 | 361 | 713 | 1401 | 1548 | 1635 |
| GGGCCCTGAAGTACCCAATG | EPHB4 | 710 | 689 | 1178 | 1169 | 1164 |
| TGTCCGACATCCGGGTGACA | EPHB4 | 481 | 683 | 1388 | 1365 | 1385 |
| TTGCACATCACACACCTCGT | EPHB4 | 372 | 544 | 807 | 777 | 783 |
| ACTTCGAACCAAAGCTCCCG | EPHB6 | 641 | 467 | 915 | 874 | 848 |
| CACTTCGACCCCCGCCAGAG | EPHB6 | 578 | 656 | 1303 | 1278 | 1256 |
| CATCCGAGAATTTACCCGGG | EPHB6 | 557 | 472 | 1034 | 1035 | 1076 |
| CCCCAGGCTGCACTGCAACG | EPHB6 | 447 | 695 | 1105 | 1114 | 1227 |
| GACCTTCGAGGCATGCCACG | EPHB6 | 344 | 671 | 1313 | 1081 | 1059 |
| TCAGCTCCGCTACTACGACC | EPHB6 | 487 | 373 | 647 | 614 | 617 |
| ACCATGTCCGAGAACACCGT | ERBB2 | 452 | 396 | 657 | 562 | 556 |
| CAGAACCTGCGAGTAATCCG | ERBB2 | 501 | 396 | 923 | 870 | 900 |
| CAGAGGCTACGAATTGTGCG | ERBB2 | 322 | 703 | 1303 | 1178 | 1170 |
| CTGTCCTTCACATACTCTCG | ERBB2 | 511 | 739 | 1453 | 1386 | 1367 |
| GCATCCTGATCAAGCGAAGG | ERBB2 | 306 | 603 | 990 | 943 | 960 |
| GCCCAGACCGTAGCACACTG | ERBB2 | 583 | 514 | 920 | 882 | 957 |
| CATGGACACGATCGACTGGA | ERBB3 | 670 | 718 | 1416 | 1111 | 1178 |
| CCATACTGATACTTGGTGTG | ERBB3 | 502 | 865 | 1892 | 1428 | 1553 |
| CTGAAGTTGTGCATGTGGGG | ERBB3 | 360 | 781 | 1458 | 1472 | 1477 |
| GTCACTTCTCGGATCCACTG | ERBB3 | 456 | 497 | 1141 | 856 | 890 |
| TAAGCAACTACTACACAGTG | ERBB3 | 342 | 569 | 1159 | 996 | 1002 |
| TCCTGTCGAAACTACAGCCG | ERBB3 | 458 | 377 | 865 | 753 | 770 |
| AATGTCCAGATGGCTTACAG | ERBB4 | 527 | 651 | 1066 | 933 | 1079 |
| AGAATTTACGCATTATCCGT | ERBB4 | 303 | 495 | 738 | 727 | 702 |
| CATAATCCAACAGGCAACCA | ERBB4 | 470 | 551 | 725 | 602 | 630 |
| CATGTTTCTACAGCTTGTGA | ERBB4 | 414 | 672 | 1244 | 1128 | 1155 |
| GTAGAACCCTTAACTCCCAG | ERBB4 | 492 | 504 | 910 | 758 | 877 |
| TAATGGTATGGTAATAACAC | ERBB4 | 420 | 322 | 707 | 658 | 624 |
| AGTGGAAAGATCAAAGACGC | ERBIN | 498 | 523 | 770 | 752 | 757 |
| ATGGAGCTTTCAATTAATGG | ERBIN | 515 | 588 | 1023 | 1176 | 1183 |
| GTGAATTGCTTGATAAGAGG | ERBIN | 410 | 531 | 879 | 830 | 865 |
| TAACTAATATGGAGACTGGA | ERBIN | 636 | 463 | 792 | 848 | 881 |
| TTCTTAAGCTCATCTGGGTA | ERBIN | 586 | 948 | 1604 | 1585 | 1513 |
| TTTACAACAGTTACCCCCAG | ERBIN | 505 | 526 | 895 | 964 | 1000 |
| AGCCCCAATATAGCTAGCAG | ERC1 | 425 | 569 | 1199 | 1172 | 1182 |
| CAGGACTCAAAAAGGACTCA | ERC1 | 660 | 530 | 1100 | 1040 | 1017 |
| CCAGTCGTCTTGTTCTCTCG | ERC1 | 447 | 764 | 1591 | 1498 | 1502 |
| CCTTGACTGCTAAAGAACAA | ERC1 | 539 | 408 | 617 | 600 | 603 |
| GGAGGAACTAAGTTCGAAAG | ERC1 | 418 | 729 | 1167 | 1238 | 1163 |
| TCAGTCAGTTCTGTCACACA | ERC1 | 727 | 620 | 916 | 883 | 953 |
| ACCTCGTAGAAGCGGCAGTG | ERCC2 | 639 | 848 | 1618 | 1287 | 1307 |
| CACTGATCATGGCATACCAG | ERCC2 | 438 | 479 | 867 | 842 | 805 |
| GCACCTCAGGATGAATACAC | ERCC2 | 535 | 238 | 454 | 411 | 425 |
| GTAACTGTAAACCACCACAT | ERCC2 | 693 | 452 | 697 | 566 | 613 |
| TGAGTGTCAACATCACTCGC | ERCC2 | 251 | 427 | 788 | 730 | 659 |
| TGCTGGAATACGTCAAGTGG | ERCC2 | 429 | 699 | 1261 | 1070 | 1165 |
| AAGTCATATTCTGCCAACAG | ERCC3 | 603 | 285 | 560 | 524 | 560 |
| AGTGGATGAGTACGGAGCCA | ERCC3 | 455 | 993 | 1304 | 1393 | 1374 |
| GACAGCTGCATACAGGGAGT | ERCC3 | 495 | 493 | 1040 | 845 | 879 |
| GACTGCAACCCTAGTCCGGG | ERCC3 | 575 | 510 | 938 | 874 | 829 |
| GGGATGTCAGATTTACCCTG | ERCC3 | 455 | 651 | 1484 | 1350 | 1350 |
| TGCAGCTGTCACGCCAACCA | ERCC3 | 441 | 464 | 1035 | 782 | 765 |
| AAAACGAGTTTACAGCTACG | ERCC4 | 472 | 655 | 1217 | 1040 | 1131 |
| AGCACTGACCGAAGTACTGA | ERCC4 | 521 | 784 | 968 | 1077 | 991 |
| CCATAATCCGTCCCTCGAAG | ERCC4 | 402 | 557 | 1003 | 882 | 861 |
| GCACCCTTGTCAGAGCGTAG | ERCC4 | 684 | 490 | 1065 | 1013 | 1069 |
| TATGAGGTCTACACACAAGG | ERCC4 | 468 | 369 | 760 | 695 | 790 |
| TCGAGCATTTACAAACATGG | ERCC4 | 602 | 338 | 649 | 576 | 534 |
| CAGACAATTCACTCAATGAA | ERCC5 | 567 | 391 | 732 | 889 | 862 |
| CGTGAAGTCATCTCCATGTG | ERCC5 | 420 | 657 | 1232 | 1367 | 1443 |
| CTTATTTGAAGCAATGCCAG | ERCC5 | 393 | 811 | 1143 | 927 | 1009 |
| GAATTCATAAGGGGTAGATG | ERCC5 | 482 | 555 | 1091 | 1227 | 1158 |
| TACACAAAGTGTCCCTTTGG | ERCC5 | 473 | 916 | 1651 | 1638 | 1637 |
| TTCGAACTTGAGTAAGACTG | ERCC5 | 444 | 878 | 1535 | 1476 | 1417 |
| AGGTGTGGATGTGTATGACC | ERCC6 | 686 | 604 | 1448 | 1486 | 1485 |
| CATCGGAGTCAACCTGACAG | ERCC6 | 539 | 525 | 1037 | 972 | 1087 |
| CCAGACAAGATGATGCGATG | ERCC6 | 624 | 590 | 1346 | 1368 | 1466 |
| GAAGTTTACAGAATTCTCAA | ERCC6 | 542 | 327 | 461 | 373 | 373 |
| TACAGCAAGATCAGGACTCG | ERCC6 | 566 | 662 | 1312 | 1361 | 1395 |
| TGCCTATTCAGGAGACTGCG | ERCC6 | 511 | 702 | 1150 | 1193 | 1241 |
| AGATCAGCCTGGACTAGTCA | ERG | 648 | 951 | 1746 | 1593 | 1629 |
| CCCTGAAACATACCTAACTG | ERG | 446 | 722 | 1004 | 1061 | 1167 |
| GAATGCAGTATGGCCAAAGG | ERG | 585 | 625 | 1020 | 1009 | 966 |
| GATGACTCTACGCTCGTTCG | ERG | 576 | 831 | 1509 | 1456 | 1524 |
| GGTCTGTCCATAGTCGCCAG | ERG | 461 | 565 | 984 | 860 | 882 |
| TTAATGCATGCTAGAAACAC | ERG | 413 | 280 | 525 | 495 | 485 |
| CGCCGTGTACAACTACCCCG | ESR1 | 418 | 500 | 998 | 923 | 939 |
| GCTGGTTCACATGATCAACT | ESR1 | 461 | 435 | 610 | 541 | 579 |
| GGGATACGGAAAGACCGAAG | ESR1 | 657 | 426 | 1069 | 970 | 979 |
| GTGTGCAATGACTATGCCTC | ESR1 | 599 | 829 | 953 | 1200 | 1116 |
| TAGTCGGAATAGATTATGGG | ESR1 | 331 | 830 | 1281 | 1185 | 1297 |
| TCAGATAATCGACGCCAGGG | ESR1 | 324 | 483 | 1083 | 1039 | 1055 |
| ACTCGTGCATATGATTGGCT | ESR2 | 398 | 847 | 1635 | 1689 | 1573 |
| AGTGATCTTGCTTCACACCA | ESR2 | 662 | 650 | 1117 | 1038 | 933 |
| CGGCTCTCTACGTAGGAGGA | ESR2 | 358 | 659 | 1314 | 1233 | 1275 |
| GGGGCGGCTCACAAGCACGT | ESR2 | 565 | 442 | 719 | 615 | 654 |
| GTGGGTACCGTGTTATGCGA | ESR2 | 484 | 569 | 1307 | 1200 | 1174 |
| TTACTAGTCCAAGTTCAAAG | ESR2 | 311 | 340 | 581 | 501 | 502 |
| ACAGGGACCTAGCATTGCAG | ESRP1 | 417 | 880 | 1514 | 1391 | 1343 |
| ACGAGGACGGGGATTTCGCG | ESRP1 | 420 | 504 | 761 | 754 | 770 |
| GATTTGGGATCAGCCAACCG | ESRP1 | 496 | 608 | 1037 | 1046 | 1049 |
| GCACGAAGTACTAGTCAGAC | ESRP1 | 429 | 556 | 851 | 682 | 765 |
| TCAGGCAATAATACATTCTG | ESRP1 | 586 | 614 | 771 | 758 | 736 |
| TTGTTACCTATCCAGACGGT | ESRP1 | 469 | 571 | 1021 | 930 | 920 |
| AAGGACGTCCTATGTACCAA | ETV1 | 569 | 654 | 1206 | 1110 | 1101 |
| GCAGCTACCCCATGGACCAC | ETV1 | 538 | 571 | 849 | 852 | 844 |
| GGTTCCTGTTTAATCATCAG | ETV1 | 606 | 576 | 943 | 955 | 1007 |
| GTGTCATCATAAAACTGCCT | ETV1 | 565 | 402 | 559 | 537 | 549 |
| TATGATCAGAAGCCACAAGT | ETV1 | 453 | 487 | 944 | 836 | 884 |
| TTCCCTTTGTAGAATCAGCG | ETV1 | 464 | 746 | 1102 | 1002 | 1065 |
| ACGCAGACATCATCTGGGAA | ETV4 | 629 | 725 | 1146 | 1185 | 1188 |
| CCTTCCCCCGGGCAGAACAA | ETV4 | 460 | 608 | 1264 | 1122 | 1191 |
| GAACGAGTGACAAATATCCA | ETV4 | 504 | 552 | 946 | 966 | 992 |
| GGTACATCGACGCACACCCG | ETV4 | 389 | 798 | 1365 | 1363 | 1372 |
| TGATGGCGATTTGTCTAGGG | ETV4 | 270 | 779 | 1285 | 1376 | 1317 |
| TGGCTGGCCTGTTCGTACAG | ETV4 | 426 | 815 | 1725 | 1314 | 1310 |
| ACCAGTATCCATCAGAACAG | ETV5 | 561 | 423 | 839 | 685 | 724 |
| GAATCAACGCAGTAATCCCG | ETV5 | 478 | 476 | 729 | 708 | 751 |
| GGTACTTACAGTTATCAGAC | ETV5 | 500 | 259 | 520 | 633 | 611 |
| GGTTAATGGCTTGAACCCGG | ETV5 | 486 | 425 | 737 | 668 | 710 |
| GTGTCGTCAAAGTACAACCG | ETV5 | 449 | 659 | 1064 | 955 | 990 |
| TGGGTTCTGACATTTGCCGG | ETV5 | 485 | 674 | 1235 | 1113 | 1099 |
| AACGGTGAAAAAAGAATCCG | ETV6 | 528 | 499 | 991 | 877 | 856 |
| ACGGTGCAACAGTTCAATGG | ETV6 | 325 | 942 | 1649 | 1745 | 1680 |
| GCAGCCAGTTTACTGGAGCA | ETV6 | 423 | 725 | 1546 | 1414 | 1429 |
| TCGGTAGCGAAAGTCCTCTT | ETV6 | 568 | 484 | 884 | 825 | 898 |
| TGGAACATGAAGTGGAGTCG | ETV6 | 503 | 902 | 1670 | 1526 | 1595 |
| TTGAAATCCACGGAGTGCCG | ETV6 | 552 | 739 | 1226 | 1087 | 1115 |
| ATACTCAAAAACTTACCACC | EWSR1 | 719 | 278 | 362 | 328 | 326 |
| GATAATACTGACCTGGTAGG | EWSR1 | 497 | 661 | 1079 | 1046 | 1057 |
| GCTGACTTCTTTAAGCAATG | EWSR1 | 554 | 562 | 898 | 902 | 911 |
| GTCCTGTCGGAATGAACCTG | EWSR1 | 415 | 509 | 1286 | 1071 | 1158 |
| GTGGCCTTACCTTGCAGGTG | EWSR1 | 474 | 738 | 1147 | 1274 | 1204 |
| TATCTTAAAAATTACCTGGG | EWSR1 | 589 | 440 | 739 | 730 | 814 |
| AAACATTCTAGCGGCCATCG | EXT1 | 370 | 742 | 1285 | 1275 | 1257 |
| ATATCACGTCCATAACGGAG | EXT1 | 305 | 448 | 794 | 783 | 789 |
| ATTCACGGCAGTCATCCATG | EXT1 | 417 | 588 | 1330 | 1167 | 1268 |
| GATGTCAAACCCCACGTCCT | EXT1 | 432 | 522 | 1079 | 1010 | 1014 |
| GCAGACACAATTCTTGTGGG | EXT1 | 483 | 725 | 1254 | 1192 | 1230 |
| GCCAACTCGAGCATCCACAA | EXT1 | 430 | 420 | 894 | 763 | 849 |
| AAAAAGTACGTGGATGACTT | EXT2 | 531 | 596 | 899 | 926 | 931 |
| ACCTTCCGGAGAAAGGACCG | EXT2 | 338 | 728 | 1347 | 1251 | 1187 |
| AGAGCTGTGTTATAATCTGG | EXT2 | 528 | 791 | 1528 | 1573 | 1487 |
| GATAGTCCCATCCCCGAGCG | EXT2 | 638 | 573 | 1026 | 1023 | 1019 |
| GCGGTGAAACCTTGCGACTG | EXT2 | 414 | 813 | 1535 | 1362 | 1370 |
| TAAATGTACCAACCTCTCAG | EXT2 | 551 | 390 | 736 | 563 | 625 |
| ACATAACATTCCGTATATGG | EZH2 | 511 | 424 | 692 | 627 | 557 |
| ATGTTCGGGGTACATTCAGG | EZH2 | 464 | 665 | 1232 | 1188 | 1113 |
| CTCATTGCGCGGGACCAGGG | EZH2 | 563 | 681 | 1157 | 1110 | 1136 |
| GGACTCCAGCACATTAATGG | EZH2 | 435 | 515 | 1011 | 842 | 931 |
| TGGTACAATAGGCGACTGTG | EZH2 | 381 | 808 | 1489 | 1615 | 1496 |
| TGTGTTATGGAAAATCGTAG | EZH2 | 372 | 687 | 1053 | 1027 | 1137 |
| ATTTGAACCATTCCCACTGG | F2R | 439 | 558 | 912 | 894 | 902 |
| CAGATGGTGGAAATGATCAG | F2R | 461 | 735 | 1450 | 1365 | 1466 |
| GGAGCTGGTCAGATATCCCG | F2R | 636 | 729 | 1489 | 1446 | 1497 |
| GGGTTTCATTGAGCACGTCA | F2R | 445 | 584 | 805 | 742 | 753 |
| TCATCTTCCAGATAAACACG | F2R | 594 | 348 | 745 | 691 | 762 |
| TGACGGTCATCAGCGTTGAC | F2R | 331 | 502 | 975 | 923 | 1001 |
| CCACGTGGCTCTTCCCACGG | FAM46C | 419 | 788 | 1241 | 1274 | 1207 |
| CGCTGTCCAACAAGAACGGG | FAM46C | 567 | 574 | 986 | 986 | 997 |
| CGTGTATGGGGACTTCGAGG | FAM46C | 407 | 682 | 1276 | 1131 | 1170 |
| GACATCGTGCAGACCGTGCG | FAM46C | 438 | 573 | 904 | 1012 | 994 |
| GCAGTCGTAGAAAAAGAGCA | FAM46C | 455 | 939 | 1354 | 1258 | 1334 |
| TCCATGGATGGGCACGACCT | FAM46C | 424 | 599 | 1102 | 1149 | 1098 |
| ACTGCACGACACCCACCCTG | FANCA | 570 | 490 | 1053 | 922 | 947 |
| CAAAGCCTCGAGAGCAACTG | FANCA | 550 | 576 | 1129 | 1087 | 1144 |
| CCTCACAGACTATGTCACAC | FANCA | 460 | 332 | 576 | 594 | 588 |
| CTGCGGTCCTAAGCGTCAGG | FANCA | 566 | 534 | 1133 | 1169 | 1249 |
| GCACATGTACTGACCACTCG | FANCA | 609 | 583 | 1022 | 1237 | 1188 |
| TAATGGTGAACCCACCCACA | FANCA | 494 | 538 | 1258 | 1020 | 1080 |
| ATGAGCTTGATCCATTAACA | FANCB | 569 | 468 | 958 | 1020 | 958 |
| CTGTTATTATAAATGAACTC | FANCB | 565 | 337 | 450 | 505 | 484 |
| CTTTAATAAACCTGGTTCGA | FANCB | 557 | 429 | 871 | 857 | 829 |
| GCAGGGGAGATTGAAAATCT | FANCB | 700 | 596 | 781 | 785 | 806 |
| GTTACTACACTGCTATAAGC | FANCB | 461 | 304 | 451 | 355 | 374 |
| TTCATAAGTTGAACTGCACA | FANCB | 347 | 560 | 836 | 860 | 840 |
| AATAGCGGCTAGTACCTCTG | FANCC | 682 | 794 | 1342 | 1328 | 1443 |
| ATGGATTTAACACTCAAAGG | FANCC | 513 | 364 | 704 | 673 | 700 |
| CAACACCATGACGAGAGATG | FANCC | 611 | 638 | 1311 | 1198 | 1196 |
| CTCAGGAACTCCTGAAATCG | FANCC | 528 | 661 | 1482 | 1473 | 1440 |
| TAAACTTAACTCCTGGACCA | FANCC | 583 | 510 | 903 | 984 | 993 |
| TTCTGTTCCAGGTCCTACGG | FANCC | 548 | 590 | 1184 | 1204 | 1270 |
| AAGGGGAGGAACATAGTCTG | FANCD2 | 547 | 687 | 1690 | 1343 | 1377 |
| CACAACGGAGACACCACTCT | FANCD2 | 446 | 487 | 945 | 867 | 905 |
| GAAATGGCTTGACAGAATTG | FANCD2 | 620 | 630 | 1326 | 1167 | 1206 |
| TGAAAAATAAAGGACGACAA | FANCD2 | 457 | 344 | 598 | 620 | 583 |
| TTTCCGAGAACTGGACATTG | FANCD2 | 488 | 760 | 1601 | 1394 | 1578 |
| TTTGCTAATCTAATCCAAGG | FANCD2 | 452 | 415 | 701 | 630 | 792 |
| AAAAGTTCCGATGTTTGGAA | FANCE | 497 | 628 | 1064 | 1031 | 1002 |
| AAGCTGTAGCTCAACCGGCA | FANCE | 396 | 481 | 800 | 699 | 661 |
| AGATTCCTCTGGCATAACGG | FANCE | 466 | 696 | 1202 | 1097 | 1097 |
| CGGTTGGGGCCGCATCCTCG | FANCE | 419 | 691 | 1228 | 1288 | 1274 |
| GAACGGTGGGGTATCGACTG | FANCE | 472 | 565 | 1191 | 1109 | 1151 |
| GCAACTCCAAACACTCAGGT | FANCE | 643 | 340 | 656 | 584 | 667 |
| CGCTACCTGCGCCGCGCGCA | FANCF | 344 | 654 | 1038 | 1066 | 1096 |
| CTGCGCCAGCTCGCACGCTG | FANCF | 548 | 582 | 1210 | 1082 | 1133 |
| GCGCCCACAGCCCGTCCAGG | FANCF | 596 | 572 | 1085 | 960 | 968 |
| GCTGGCCGTGTCCCGCACGG | FANCF | 574 | 606 | 1190 | 1198 | 1283 |
| TGCTGCGCCTGCTCTTCCCG | FANCF | 456 | 1141 | 1775 | 1944 | 1942 |
| TGCTGTCGGCGCGCCTGCTG | FANCF | 497 | 740 | 1709 | 1734 | 1770 |
| CAGCTGCCTAGACTTGTGGA | FANCG | 653 | 639 | 1271 | 1350 | 1264 |
| CAGGTAACAGTCAATTCCAA | FANCG | 425 | 360 | 610 | 551 | 616 |
| CTGAAAGAGGGATCAACCTG | FANCG | 463 | 738 | 1387 | 1303 | 1324 |
| GCGGACAAAACTCTCTGCTG | FANCG | 538 | 851 | 1288 | 1388 | 1438 |
| GCTAAGCACTGACCATCTTG | FANCG | 403 | 995 | 1831 | 1904 | 1811 |
| GGGAGCAAAAGAGTCGCGGG | FANCG | 435 | 610 | 890 | 930 | 973 |
| AGTTGGGATCATGTTACTCA | FANCI | 562 | 582 | 1191 | 1053 | 1121 |
| ATACAAGCTTCTAACTTCAG | FANCI | 434 | 489 | 963 | 867 | 873 |
| CATAGGATTACTAATGCTAG | FANCI | 464 | 368 | 811 | 888 | 795 |
| TAATGATACTTACTTGAACA | FANCI | 452 | 407 | 673 | 537 | 527 |
| TGATGGCAAACACGATGTGT | FANCI | 467 | 616 | 1136 | 1162 | 1108 |
| TTCCCTGATATGTGCAACGA | FANCI | 479 | 518 | 1053 | 798 | 909 |
| ACAACACGCAGAATTGCACT | FANCL | 464 | 435 | 933 | 888 | 856 |
| ACTCACCAGACTTGTATACG | FANCL | 351 | 713 | 1402 | 1366 | 1352 |
| AGAATCACCAGATTGTGTTG | FANCL | 466 | 960 | 1669 | 1892 | 1738 |
| AGGTGAAAATCACGTCCCTG | FANCL | 416 | 791 | 1628 | 1775 | 1678 |
| CGAAATGAATCCCTCGTACA | FANCL | 715 | 633 | 1018 | 944 | 1005 |
| TTCATCCATAACATCCCAGA | FANCL | 556 | 273 | 436 | 411 | 406 |
| AATTGTGATACGGGTAACAG | FANCM | 509 | 555 | 957 | 860 | 936 |
| AGTCCACGAATGGTTCCTGA | FANCM | 518 | 554 | 1142 | 914 | 892 |
| ATAAAGAATTTATGTCTCCA | FANCM | 433 | 410 | 935 | 851 | 918 |
| ATACAACAAGGCATAATCGA | FANCM | 530 | 284 | 584 | 454 | 492 |
| GAAATTGTACATGACCACGG | FANCM | 543 | 394 | 685 | 625 | 599 |
| GAATGGTCAACTTGATACGT | FANCM | 619 | 335 | 675 | 633 | 691 |
| AAAGAAACTGTACTCAGACC | FAS | 474 | 374 | 651 | 620 | 648 |
| ACAAAAGGGAATGTCACCCT | FAS | 659 | 418 | 1000 | 903 | 845 |
| ACAAAGTGTTGAGATCGGGT | FAS | 508 | 600 | 1253 | 1161 | 1162 |
| AGTGACTGCACGTCTTATGC | FAS | 544 | 447 | 866 | 898 | 886 |
| CGGACGTCAGAATCGGGATG | FAS | 572 | 1189 | 2177 | 2142 | 2116 |
| TGAAAACTTACGTGCTACAC | FAS | 359 | 412 | 769 | 828 | 756 |
| AGGCGACGGTCTCTCCCAGG | FASN | 691 | 543 | 895 | 883 | 874 |
| CCAGAGGGGATACCTACCCA | FASN | 488 | 436 | 1010 | 810 | 846 |
| CTGAACAGCACGGTAGCCGG | FASN | 439 | 399 | 1091 | 889 | 935 |
| GATGTCTCGGAAGTTAAGGG | FASN | 631 | 416 | 897 | 717 | 784 |
| GCCCAGCGTCGCTCTAGACA | FASN | 370 | 617 | 1339 | 1158 | 1148 |
| GGACCATGGCCCTAGAAGTG | FASN | 603 | 582 | 1294 | 1172 | 1206 |
| CCGGATGACCTCATCAGACC | FBXO31 | 524 | 315 | 586 | 572 | 576 |
| CTACGGAGGATTGTTGAATG | FBXO31 | 534 | 809 | 1477 | 1510 | 1525 |
| CTTCCATGGAAAGTGTGCCA | FBXO31 | 518 | 725 | 1473 | 1366 | 1364 |
| GCAGCTGAATACGGTGCATG | FBXO31 | 477 | 716 | 1627 | 1456 | 1397 |
| GCCATACATGCACTCCACCG | FBXO31 | 452 | 615 | 1221 | 1208 | 1263 |
| GGAATTCCGAACATGGCTGA | FBXO31 | 580 | 895 | 1285 | 1252 | 1265 |
| ACAGCTCAGACAACAAGAGG | FBXW7 | 449 | 612 | 1111 | 1211 | 1097 |
| AGGCAGGTCCACGATACTAG | FBXW7 | 466 | 652 | 1113 | 979 | 1012 |
| ATAGAATTGATACTAACTGG | FBXW7 | 543 | 544 | 744 | 708 | 666 |
| ATGAAGTCTCGTTGAAACTG | FBXW7 | 505 | 660 | 1158 | 1184 | 1153 |
| GGAGACCTGAGAGCAGCCAA | FBXW7 | 505 | 725 | 1205 | 1150 | 1068 |
| TCATTAGTGGATCTACAGAT | FBXW7 | 470 | 460 | 669 | 664 | 674 |
| AGGCCCCAGGTATAGAACAG | FCHSD1 | 365 | 678 | 1389 | 1322 | 1225 |
| CATTGAACGGGAATATGGAC | FCHSD1 | 384 | 327 | 655 | 599 | 548 |
| GACTGAACCGAAGTGACCAT | FCHSD1 | 520 | 587 | 1085 | 1050 | 1061 |
| GCAGGAGCGACGTCTTAGTG | FCHSD1 | 594 | 958 | 1832 | 1496 | 1515 |
| GTCTGACCGATACCGTGACC | FCHSD1 | 502 | 376 | 675 | 678 | 723 |
| GTGCGGCTTAGCAAGGTCAA | FCHSD1 | 478 | 882 | 1240 | 1177 | 1322 |
| CAAGCAAGTTTGAACAACAG | FCRL4 | 418 | 432 | 767 | 743 | 740 |
| CGTGTTTGAAGGTGACACGC | FCRL4 | 639 | 477 | 886 | 813 | 809 |
| GAGAAGACTCGGGATCGTAT | FCRL4 | 383 | 475 | 911 | 927 | 954 |
| GCACCTGTCGTACTTCGGAA | FCRL4 | 568 | 613 | 978 | 1070 | 1004 |
| TTGAAGACGACGGTCCATGG | FCRL4 | 468 | 835 | 1394 | 1564 | 1505 |
| TTTACAGAGCTGCCTTCCGT | FCRL4 | 409 | 408 | 773 | 709 | 702 |
| AAGTACCAGGAGGCCAGCAA | FES | 514 | 434 | 791 | 864 | 924 |
| AGAGTGACAGGGAATACGCA | FES | 510 | 627 | 1184 | 1218 | 1109 |
| GAAAGCCATCCCCTTCAAGT | FES | 561 | 362 | 529 | 511 | 496 |
| GCAGCGACTGGAGTAGACCA | FES | 475 | 777 | 1406 | 1362 | 1406 |
| TACTCACCGAGAACTTAGGG | FES | 530 | 582 | 1394 | 1214 | 1194 |
| TAGGCCTACTGATCCGCGAG | FES | 415 | 552 | 1099 | 1136 | 1162 |
| CCAGTGGCTCAAGCACATCG | FGFR1 | 586 | 440 | 937 | 881 | 915 |
| GAAACAGTGTCTCACGCATA | FGFR1 | 520 | 881 | 1202 | 1223 | 1293 |
| GAGCATCATAATGGACTCCG | FGFR1 | 373 | 653 | 1196 | 1086 | 1055 |
| GATCATCATCTACTGCACAG | FGFR1 | 623 | 324 | 526 | 484 | 421 |
| GCTTCCCGAAGACCCTCGCT | FGFR1 | 507 | 528 | 777 | 808 | 877 |
| TGCTGGTCACACAAGCGTAG | FGFR1 | 341 | 694 | 1098 | 1015 | 1055 |
| ACAGCGAAACTTGACAGTAT | FGFR2 | 559 | 492 | 971 | 944 | 896 |
| ACTGCCGGAAGATCCAAAGT | FGFR2 | 451 | 459 | 824 | 782 | 807 |
| AGCGTGGTCCCATCCGACAA | FGFR2 | 480 | 688 | 1507 | 1410 | 1393 |
| AGGCCCACTCTACGTCATTG | FGFR2 | 333 | 1124 | 1806 | 1535 | 1786 |
| GATAGCCATTTACTGCATAG | FGFR2 | 661 | 796 | 1545 | 1311 | 1352 |
| GGAGCAAATACGGGCCCGAT | FGFR2 | 595 | 372 | 790 | 743 | 748 |
| AATGTGTCCGAGCGTGACGG | FGFR3 | 501 | 622 | 1122 | 959 | 1098 |
| AGAGGCCATCGGTATTGACA | FGFR3 | 483 | 846 | 1442 | 1349 | 1341 |
| CACCTATAGACAGTCCCACA | FGFR3 | 422 | 418 | 870 | 870 | 821 |
| CCAGTGGCTCAAGCATGTGG | FGFR3 | 513 | 648 | 1143 | 1099 | 1120 |
| CCTGTCGCTTGAGTGGAAAG | FGFR3 | 532 | 895 | 1691 | 1721 | 1695 |
| GTGTAGTTCCCACGGTCCGA | FGFR3 | 600 | 557 | 1125 | 1018 | 1073 |
| ACAACCCAGCTAGTAGCAGG | FGFR4 | 470 | 649 | 1117 | 1255 | 1210 |
| ATGCAGCATACCTGTCCCGG | FGFR4 | 495 | 579 | 1111 | 980 | 1030 |
| CCTCAGCTGACACGTTCCGA | FGFR4 | 546 | 6 | 14 | 28 | 26 |
| GACCTCTAGCAATGGCAACG | FGFR4 | 452 | 538 | 1042 | 1084 | 1054 |
| TGCAGACAATGCCTCCGACA | FGFR4 | 522 | 641 | 1157 | 1170 | 1153 |
| TGCCCATAGAGTTCTCCACG | FGFR4 | 603 | 912 | 1600 | 1770 | 1690 |
| AATTGGACGCACTCACACAC | FH | 360 | 346 | 475 | 567 | 514 |
| ACCATTCGAGCGGCGTCCGG | FH | 443 | 638 | 1147 | 1116 | 1223 |
| CAGCGGAGCTATGAACACTA | FH | 552 | 464 | 656 | 545 | 552 |
| GCAGCAATGTGCATTGCCGT | FH | 531 | 547 | 975 | 956 | 977 |
| GCTGCGAGCTCATAGATCCT | FH | 562 | 563 | 1190 | 1101 | 1109 |
| TCTTGATTCAAAGATTGCTG | FH | 343 | 609 | 1143 | 1119 | 1185 |
| AAACCACAGAGGGCTTGATG | FHIT | 575 | 559 | 1099 | 896 | 846 |
| AGGTCTCGGAAACGCTCCAC | FHIT | 400 | 424 | 836 | 910 | 907 |
| CATCTGTACACTCACCTGCA | FHIT | 540 | 528 | 993 | 1079 | 1028 |
| GGCGACCCAGAGAGTTGGGA | FHIT | 447 | 850 | 1482 | 1590 | 1635 |
| TCTTGTGAACAGGAAACCTG | FHIT | 326 | 495 | 1003 | 858 | 818 |
| TGAAGCCGGACAGACTGTGA | FHIT | 458 | 818 | 1293 | 1157 | 1294 |
| CAGATACCTGGATGGTCCGA | FKBP9 | 479 | 576 | 1089 | 1042 | 995 |
| GAAAACATATGACACATATG | FKBP9 | 389 | 607 | 961 | 965 | 969 |
| GGTTGTGGAAGTCAATCACG | FKBP9 | 602 | 628 | 1259 | 1152 | 1218 |
| TGAGCGGCGAAGTCAAAGTG | FKBP9 | 663 | 740 | 1414 | 1362 | 1318 |
| TTGGGATGTGTGTAAACGAG | FKBP9 | 460 | 768 | 1215 | 1227 | 1163 |
| TTTGACACATACATCGGGCA | FKBP9 | 528 | 424 | 971 | 875 | 914 |
| AGATAGGGCCTTCACGACCA | FLCN | 478 | 699 | 1568 | 1484 | 1441 |
| ATAAATTCGGTCCATCATGA | FLCN | 590 | 696 | 1229 | 1146 | 1091 |
| CAAAAAGTCAGACATGTGCG | FLCN | 644 | 847 | 1654 | 1647 | 1642 |
| GCGTGCCCAGAGAATGAACA | FLCN | 472 | 557 | 1046 | 913 | 960 |
| GCTGACTGAGAAACTCCTAG | FLCN | 491 | 496 | 917 | 936 | 979 |
| GGCCAACATACGAAAAGATG | FLCN | 545 | 396 | 796 | 871 | 876 |
| AGAACTGTGCAAAATGAACA | FLI1 | 370 | 432 | 703 | 678 | 668 |
| AGCAGTCGCCTAGCCAACCC | FLI1 | 557 | 570 | 899 | 916 | 889 |
| CATGACCACCAACGAGAGGA | FLI1 | 619 | 855 | 1384 | 1444 | 1474 |
| CGGTGAGGGTCAATGTCAAG | FLI1 | 473 | 739 | 1221 | 1135 | 1219 |
| CTCCGTAGGCAGAGTCGAAG | FLI1 | 540 | 657 | 1255 | 1150 | 1242 |
| TCTTATGACTCAGTCCGGAG | FLI1 | 616 | 400 | 818 | 988 | 904 |
| GACCGGACGCTATATGATTG | FLNB | 593 | 906 | 1595 | 1627 | 1811 |
| GATGTTAACATTACATACGG | FLNB | 443 | 236 | 400 | 359 | 434 |
| GCCCGACAATCCCACCATGA | FLNB | 620 | 610 | 1141 | 1113 | 1100 |
| GCCTGGAAGCCCCTACACAG | FLNB | 434 | 453 | 832 | 773 | 825 |
| GTGCTTGATGGCCTTAACCG | FLNB | 486 | 458 | 1093 | 998 | 1063 |
| TGAATCAACTGTGAAGTCGG | FLNB | 494 | 697 | 1211 | 1313 | 1350 |
| AAGAACGCTGTAGAACACAT | FLT1 | 505 | 4 | 18 | 20 | 15 |
| AGAGCCAGGAACATATACAC | FLT1 | 662 | 346 | 563 | 559 | 580 |
| ATCACCTACATGCCGGACTG | FLT1 | 526 | 994 | 1475 | 1519 | 1508 |
| ATGGCTCAGGCAAACCACAC | FLT1 | 539 | 359 | 665 | 652 | 660 |
| CTTGGAAAAAATGCCTACGG | FLT1 | 456 | 744 | 999 | 911 | 871 |
| TGATGGGAAGCGAATAACCT | FLT1 | 429 | 499 | 896 | 1203 | 1120 |
| AAAGCTGTTCACATAAACCA | FLT3 | 538 | 330 | 525 | 618 | 670 |
| AAGTACTGGTCAACACCCCG | FLT3 | 500 | 531 | 943 | 912 | 940 |
| CTCACCTGTACCCATCATCG | FLT3 | 658 | 604 | 1131 | 1125 | 1175 |
| GAGCATTCCAGAACCGACTG | FLT3 | 449 | 606 | 961 | 881 | 1041 |
| GTACTTGTGACAAATAAGCA | FLT3 | 556 | 373 | 932 | 741 | 835 |
| TGGTAATGGGTATCCATCAG | FLT3 | 512 | 612 | 1293 | 1363 | 1396 |
| ATCAGCGTCGAATGGCTCAA | FLT4 | 406 | 876 | 1357 | 1238 | 1316 |
| CCCTACCTGTGATATGCACG | FLT4 | 494 | 9 | 11 | 8 | 13 |
| CGAAGGCGACTACGTATGCG | FLT4 | 307 | 781 | 1493 | 1431 | 1437 |
| CGTGATCCTTGTCGGCACTG | FLT4 | 455 | 773 | 1339 | 1393 | 1397 |
| CTCACCTCTCACAAACACGT | FLT4 | 488 | 276 | 334 | 356 | 369 |
| TCAGTGTCCAGTGGCACTGG | FLT4 | 378 | 774 | 1471 | 1409 | 1443 |
| ATCATAAACATCTCTAACTG | FNTA | 503 | 414 | 628 | 563 | 544 |
| ATGCAAAGAATTATCATGCC | FNTA | 492 | 380 | 593 | 592 | 543 |
| ATGCAGCCAATTACACAGTG | FNTA | 464 | 421 | 1068 | 861 | 905 |
| CTACATCACTGCGATAATTG | FNTA | 460 | 686 | 1021 | 893 | 815 |
| GCAGCAAGAGGAAGCCATGG | FNTA | 390 | 603 | 1185 | 1129 | 1195 |
| GCGTTACCTGTATAGGACAT | FNTA | 701 | 595 | 1049 | 924 | 930 |
| ACAGAAAGTGTAGCCACCAT | FNTB | 544 | 587 | 933 | 1021 | 1018 |
| CAGCTTACCTTGGAACAAGG | FNTB | 569 | 337 | 585 | 543 | 485 |
| CGGCAGATGCGCTTCGAAGG | FNTB | 715 | 749 | 1183 | 1057 | 1194 |
| CTTGTGCATCATTGGCACCG | FNTB | 488 | 826 | 1361 | 1419 | 1346 |
| GTTGGTCAGAGAAGCTACGG | FNTB | 497 | 418 | 1023 | 936 | 906 |
| TCCAGGAACTGACACACACT | FNTB | 670 | 478 | 982 | 960 | 1083 |
| CAATGTTAGTAGGTTCACTG | FOLH1 | 648 | 891 | 994 | 1085 | 1089 |
| CACCAAACACCCATGAGTCA | FOLH1 | 531 | 578 | 991 | 905 | 864 |
| CTTGGTATACAACCTAACAA | FOLH1 | 611 | 420 | 561 | 358 | 393 |
| GGAGAGAAGGCATTAAAAGG | FOLH1 | 527 | 820 | 1404 | 1412 | 1460 |
| TCTGGATAGGACTTCACCCC | FOLH1 | 465 | 669 | 1087 | 1063 | 1013 |
| TTATAGACGTGAAGTCACAG | FOLH1 | 395 | 678 | 927 | 985 | 1111 |
| AACATGTCCTACGCGAACCC | FOXA1 | 804 | 313 | 731 | 645 | 707 |
| CCCGGCCCGGAGCTGATCGG | FOXA1 | 566 | 550 | 1213 | 1324 | 1253 |
| GAAGGAGAGCGAGTGCCGAA | FOXA1 | 384 | 430 | 804 | 882 | 893 |
| GCACTACAGTACTCGCCCTA | FOXA1 | 637 | 437 | 867 | 713 | 779 |
| GCCAACTCGCCGCTCCACCG | FOXA1 | 487 | 561 | 1218 | 1193 | 1308 |
| TGGACGGCGCGTACCCCATG | FOXA1 | 519 | 673 | 1361 | 1306 | 1305 |
| CCCACTTCCAGCCGGGCAAG | FOXL2 | 547 | 592 | 1310 | 1114 | 1100 |
| CTTCTCGAACATGTCCTCGC | FOXL2 | 581 | 284 | 366 | 417 | 427 |
| GAAGCCCCCGTACTCGTACG | FOXL2 | 521 | 683 | 1202 | 1233 | 1227 |
| GATGATGTACTGGTAGATGC | FOXL2 | 641 | 505 | 1029 | 1034 | 1042 |
| TTCATCAAGGTGCCGCGCGA | FOXL2 | 588 | 614 | 1003 | 1027 | 991 |
| TTCTACGAGAAGAACAAGAA | FOXL2 | 359 | 426 | 588 | 693 | 627 |
| GGAGTTCAGCCAGTCCAACT | FOXO1 | 726 | 510 | 707 | 648 | 707 |
| GGCGGTGCAAATGAGTAGCA | FOXO1 | 536 | 802 | 1272 | 1296 | 1244 |
| TAGCATTTGAGCTAGTTCGA | FOXO1 | 461 | 686 | 1366 | 1301 | 1215 |
| TCAAGAGCGTGCCCTACTCA | FOXO1 | 615 | 1011 | 1596 | 1609 | 1739 |
| TTGCTTATCTCAGACAGGCT | FOXO1 | 382 | 327 | 793 | 730 | 803 |
| TTGGTGGATGCTCAATCCAG | FOXO1 | 461 | 715 | 1286 | 1424 | 1353 |
| AACAAGTATACCAAGAGCCG | FOXO3 | 476 | 417 | 770 | 738 | 805 |
| AGCTTCCCATATACCACCAA | FOXO3 | 578 | 424 | 661 | 633 | 623 |
| CATTAGAATTGGTGCGCGAG | FOXO3 | 426 | 699 | 1201 | 1078 | 1106 |
| GTGCATGGCTTACTCACAGA | FOXO3 | 429 | 603 | 983 | 976 | 890 |
| GTTCATGCGAGTCCAGAATG | FOXO3 | 511 | 731 | 1422 | 1220 | 1318 |
| TTCGGGGATCATCGAGTCGG | FOXO3 | 687 | 709 | 1374 | 1306 | 1296 |
| ACCGCGAAGAAGCAGATGTG | FOXO4 | 461 | 776 | 1560 | 1509 | 1516 |
| CCCCGCATAGCTGCTGACTG | FOXO4 | 528 | 612 | 1061 | 1051 | 1107 |
| GTAACAGGTCCTCGGAAGGG | FOXO4 | 404 | 705 | 1247 | 1236 | 1332 |
| GTGCCCTACTTCAAGGACAA | FOXO4 | 515 | 706 | 1123 | 1170 | 1129 |
| GTTCATCAAGGTTCACAACG | FOXO4 | 525 | 345 | 709 | 658 | 642 |
| TCAGTAGTGAGCTCTGGTCG | FOXO4 | 522 | 730 | 1334 | 1291 | 1432 |
| CATGTGCAGATGAGCCATCA | FOXP4 | 642 | 663 | 1413 | 1317 | 1293 |
| GACATCATGGCCACGGACAC | FOXP4 | 495 | 477 | 898 | 752 | 798 |
| GCAGCAGACTCGTTCCCAGA | FOXP4 | 599 | 787 | 979 | 986 | 998 |
| TACACTCTCCATGTCCGTAG | FOXP4 | 641 | 552 | 909 | 989 | 1041 |
| TGGAAAGCAGCAACCCAAAG | FOXP4 | 258 | 555 | 1068 | 1020 | 999 |
| TTTAGGGGGAGCGGCAAACG | FOXP4 | 622 | 771 | 1016 | 1078 | 1002 |
| AAAGAAGAAATCAGATCAGT | FSCB | 442 | 393 | 655 | 667 | 656 |
| AGAGCCTCTTCACTTAGGGT | FSCB | 511 | 569 | 1397 | 1415 | 1463 |
| AGGAATATTACCAATACTAA | FSCB | 586 | 294 | 447 | 401 | 432 |
| TCTTCAGTCTCTCGTATAGG | FSCB | 551 | 746 | 1196 | 1200 | 1192 |
| TGCAAAAAAGTGAATCCTCA | FSCB | 499 | 465 | 634 | 697 | 694 |
| TGGGGGCTGAACTTCATCAG | FSCB | 492 | 633 | 1130 | 959 | 994 |
| AAACCTGAACACTTCGAAGA | FUBP1 | 425 | 332 | 458 | 465 | 490 |
| AATCATGATTCCAGCTAGCA | FUBP1 | 508 | 657 | 1067 | 877 | 930 |
| ACAGATCTCACGCATACAAC | FUBP1 | 517 | 359 | 472 | 434 | 457 |
| CACAGCAAATAGACTATGCG | FUBP1 | 627 | 548 | 1200 | 1127 | 1174 |
| GAGTTAATTCGTGATCAAGG | FUBP1 | 602 | 732 | 1066 | 1223 | 1184 |
| TCAAATGACTATGGTTATGG | FUBP1 | 416 | 711 | 1176 | 1225 | 1214 |
| AAACGTTACAATTGAGTCTG | FUS | 483 | 742 | 1450 | 1627 | 1639 |
| AGCTCCTATAATCCCCCTCA | FUS | 440 | 512 | 1159 | 1159 | 1238 |
| CAAGATCAGTCCTCCATGAG | FUS | 423 | 821 | 1477 | 1262 | 1339 |
| GGCACTCAGTCAACTCCCCA | FUS | 517 | 431 | 781 | 680 | 680 |
| GGTGGTTACAACCGCAGCAG | FUS | 511 | 703 | 1302 | 1181 | 1223 |
| TGGGTCATCAAACGACACCG | FUS | 441 | 475 | 961 | 888 | 822 |
| AAAATTCGCAAACTTGACAA | FYN | 406 | 262 | 589 | 458 | 451 |
| AAACTACAACAACTTCCACG | FYN | 589 | 326 | 405 | 350 | 357 |
| ATCATAAAGGGCTACAAACA | FYN | 352 | 529 | 902 | 827 | 886 |
| GAAGTTGAAACACGACAAGT | FYN | 472 | 471 | 735 | 662 | 741 |
| GGTACCTCTCGGGTTTCCAA | FYN | 471 | 492 | 758 | 701 | 715 |
| TATGACAGGGAACAACTAGG | FYN | 389 | 441 | 784 | 858 | 853 |
| ATGGTGATGGTCTTGACAGC | FZD7 | 543 | 415 | 892 | 902 | 852 |
| CAGCCCGACAGAAAGATGAT | FZD7 | 384 | 451 | 865 | 936 | 824 |
| CTCAGACGACGGCTACCGCA | FZD7 | 444 | 629 | 1059 | 993 | 1064 |
| CTTAAAGTACATCAGGCCGT | FZD7 | 485 | 601 | 1026 | 993 | 964 |
| GCAGCCGTACCACGGCGAGA | FZD7 | 482 | 808 | 1400 | 1245 | 1330 |
| GTGGGCCAGAACACGTCCGA | FZD7 | 345 | 429 | 924 | 853 | 881 |
| CAGATTGTCAGCAAACGGGC | GATA1 | 439 | 290 | 521 | 449 | 524 |
| CCACCCAAAGGAGTTACCTG | GATA1 | 489 | 524 | 1205 | 1113 | 1167 |
| GAGCAACAGCCACTCCACTG | GATA1 | 339 | 688 | 1286 | 1225 | 1239 |
| GCAGTGTTCCACGAAGCTTG | GATA1 | 450 | 1052 | 1679 | 1559 | 1730 |
| GTGGTGACGATACCAAGGCC | GATA1 | 441 | 272 | 486 | 497 | 450 |
| TGGGGCACACAGTCGAGGCA | GATA1 | 418 | 734 | 1269 | 1354 | 1381 |
| CACCCCGACTCGCACCATCC | GATA2 | 440 | 555 | 712 | 778 | 835 |
| CCTGGACTATGCAACAAGTG | GATA2 | 432 | 775 | 1714 | 1463 | 1519 |
| GGCCGGCACATAGGATGGGT | GATA2 | 395 | 421 | 831 | 774 | 743 |
| TCACGGTCCAGGGATTGTGG | GATA2 | 339 | 715 | 1475 | 1604 | 1441 |
| TGCTATCGGTCAGTGACACC | GATA2 | 427 | 588 | 1018 | 915 | 1036 |
| TTCAACCACCTAGACTCGCA | GATA2 | 386 | 421 | 568 | 440 | 467 |
| ACCGAGTTCCCATAGTACGA | GATA3 | 570 | 722 | 1165 | 1040 | 1140 |
| AGAGTCCGGAGCTGTATTCG | GATA3 | 517 | 976 | 1600 | 1758 | 1688 |
| CCGGCTTCGGGTGTAAGTCG | GATA3 | 581 | 660 | 1498 | 1432 | 1469 |
| CTCGAGTCCACGCACTCGCG | GATA3 | 485 | 705 | 1070 | 973 | 1107 |
| GCCATCCCGTCGCCACAGCG | GATA3 | 396 | 658 | 1070 | 994 | 1031 |
| TCCAGAACGTCCATCCACCA | GATA3 | 509 | 399 | 801 | 903 | 922 |
| AGCATTGCGGCGCCACAGAG | GATA4 | 632 | 800 | 1165 | 1248 | 1288 |
| CCCATCACGCCTCCAGAGCG | GATA4 | 490 | 647 | 1295 | 1326 | 1297 |
| CGGCGCTGAGGCTTAATGAG | GATA4 | 378 | 603 | 1201 | 1032 | 1080 |
| CGTGGGCACGTAGACCGGCG | GATA4 | 562 | 780 | 1091 | 1223 | 1132 |
| GCAGCACCGGGCTGTCGAAG | GATA4 | 490 | 781 | 1246 | 1302 | 1365 |
| GTCCCCGGGAAGGAGAAGCG | GATA4 | 465 | 597 | 1155 | 1141 | 1115 |
| ACGTCCCCCGAGCTTCTCGG | GATA6 | 648 | 748 | 1455 | 1270 | 1243 |
| ATCGATGGAGAGACTACGGC | GATA6 | 585 | 371 | 738 | 714 | 708 |
| CAGATGAGATGATTCGGGCT | GATA6 | 4 | 456 | 834 | 763 | 793 |
| GCTTACGACGGCGCGCCCGG | GATA6 | 579 | 739 | 1313 | 1169 | 1080 |
| GGGCGCCCCGAAGGGATGCG | GATA6 | 461 | 771 | 1308 | 1558 | 1443 |
| TTTGAAACGCTTGGGAACAG | GATA6 | 643 | 675 | 1262 | 1316 | 1313 |
| AGCCGCACAAGTGTACGGTG | GLI1 | 465 | 808 | 1816 | 1553 | 1480 |
| AGGATACAACCCCAATGCAG | GLI1 | 696 | 458 | 645 | 592 | 562 |
| CATGTGACCAAGCGGCACCG | GLI1 | 546 | 710 | 1071 | 969 | 1018 |
| CGGCGGTCAAGAGAAACGGG | GLI1 | 555 | 568 | 1039 | 1126 | 1138 |
| GCTGGTTAGCAAGTGCCCAG | GLI1 | 605 | 574 | 1163 | 1119 | 988 |
| GGATACCAGAGGGCTACGGG | GLI1 | 577 | 761 | 1327 | 1256 | 1292 |
| AAGCATGTGAAAACGGTACA | GLI2 | 500 | 566 | 1228 | 1093 | 1097 |
| CCACGACTACCTCAATCCAG | GLI2 | 537 | 425 | 714 | 744 | 797 |
| CCATCTCCACCGACGCGTCG | GLI2 | 428 | 534 | 1089 | 1139 | 1137 |
| CGTCATTCATAAGCGCAGCA | GLI2 | 495 | 440 | 708 | 627 | 623 |
| CTGCTCGAACCGGTGCATGG | GLI2 | 411 | 912 | 1351 | 1561 | 1560 |
| GGTACACATGCGCCGACACA | GLI2 | 691 | 787 | 1348 | 1229 | 1187 |
| AAACAGGACGCATTCCAATG | GLI3 | 496 | 866 | 1396 | 1431 | 1582 |
| ACAGCCAGAAGGAACAACCC | GLI3 | 567 | 338 | 745 | 579 | 597 |
| GAGCTTCCTCTGACCGACGG | GLI3 | 419 | 601 | 1183 | 1058 | 1089 |
| GCCCTCGGTGCCGTACCGTG | GLI3 | 566 | 702 | 1312 | 1588 | 1515 |
| TCTGATGATAGTATTCCGCT | GLI3 | 458 | 534 | 909 | 857 | 830 |
| TGATTGCACTCGCAGACAAG | GLI3 | 335 | 785 | 922 | 841 | 801 |
| AAGTTCAGTCCTGATTTGTG | GLS | 610 | 787 | 1329 | 1550 | 1446 |
| GACGCGTTCGGCAACAGCGA | GLS | 448 | 577 | 876 | 953 | 903 |
| GAGGTAAAAGACATAAAGTC | GLS | 708 | 320 | 594 | 639 | 674 |
| GCATTTACCATAGGATTGTG | GLS | 771 | 717 | 1362 | 1368 | 1268 |
| TCCATACACTCTTTCAACCT | GLS | 531 | 381 | 830 | 707 | 763 |
| TTGTGAATCAGCCAGCGTGA | GLS | 431 | 681 | 1378 | 1326 | 1391 |
| AAAAAGCTCCCGGACTTGGG | GM2A | 472 | 698 | 1247 | 1215 | 1368 |
| ACAGGCTTCTGAACAACACA | GM2A | 544 | 716 | 899 | 757 | 821 |
| CCTTCTGAGGATCGTTGAGG | GM2A | 405 | 857 | 1359 | 1391 | 1321 |
| CTGGGTCAAAGTCCCATGTG | GM2A | 416 | 804 | 1539 | 1521 | 1514 |
| GGACGCGTGAAGGGTCCGCA | GM2A | 452 | 691 | 1266 | 1273 | 1267 |
| TGTTCCCAGGAAAGAGAACA | GM2A | 544 | 619 | 1033 | 1006 | 1047 |
| AATATTGCTGGATCAAACCA | GMPS | 510 | 405 | 893 | 710 | 783 |
| AGAGAAAAGAAAAATTATTG | GMPS | 709 | 487 | 1164 | 1178 | 1238 |
| AGCGCATCCCTTGTTGCAAG | GMPS | 624 | 720 | 1153 | 1246 | 1361 |
| ATGAGCCGCATTTATCACTG | GMPS | 579 | 696 | 1301 | 1298 | 1208 |
| CACATATTACTCTGATTGCC | GMPS | 564 | 657 | 781 | 837 | 810 |
| TGCACCCAGCTATATCGTAG | GMPS | 539 | 621 | 1148 | 1019 | 1177 |
| AAGTGAATATGACCAAGTGC | GNA11 | 482 | 486 | 744 | 671 | 727 |
| CCCCATAGGGCTCAGCACCA | GNA11 | 700 | 431 | 809 | 863 | 897 |
| CTTCCCACAGAACCGCATGG | GNA11 | 624 | 789 | 1273 | 1449 | 1358 |
| GGACCATCATCACCTACCCC | GNA11 | 473 | 599 | 742 | 815 | 772 |
| TAGGGCTACAAGGAACATGA | GNA11 | 628 | 343 | 528 | 493 | 561 |
| TCCCACATCCACCATCCTAC | GNA11 | 403 | 211 | 387 | 432 | 388 |
| ATTAGTTCGAGAAGTCGACG | GNAQ | 445 | 561 | 1306 | 1328 | 1294 |
| GATGTTCTGATACACCAGCT | GNAQ | 463 | 418 | 714 | 667 | 730 |
| GGGGTATTCGATGATCCCTG | GNAQ | 628 | 853 | 1562 | 1453 | 1392 |
| GGTAGGCGGGGTCAGCAACG | GNAQ | 591 | 633 | 1279 | 1167 | 1242 |
| TGAATATGATCAAGTTCTCG | GNAQ | 275 | 832 | 1516 | 1659 | 1658 |
| TTGCAGAATGGTCGATGTAG | GNAQ | 517 | 1477 | 1806 | 1716 | 1667 |
| ACTGGGCGCAGTCAATCAGC | GNAS | 584 | 444 | 525 | 603 | 602 |
| CCCTGAGAACCAGTTCAGAG | GNAS | 538 | 741 | 1117 | 1201 | 1207 |
| CTTCTGCATACCTGATCACT | GNAS | 553 | 615 | 828 | 859 | 791 |
| CTTTAAAAGGGGCGGCGAAG | GNAS | 496 | 835 | 1248 | 1273 | 1282 |
| GGAGTTCTATGAGCATGCCA | GNAS | 540 | 404 | 739 | 674 | 683 |
| TTCCGCAGCATGTTTGACGT | GNAS | 615 | 592 | 1036 | 1126 | 1196 |
| ATGGTGCATTCCAGGTGCTG | GNRH1 | 544 | 914 | 1765 | 1698 | 1733 |
| CCAATTCCAAAACTTGTAGC | GNRH1 | 681 | 351 | 586 | 615 | 621 |
| GGAGTCCATAGGACCAGTGT | GNRH1 | 611 | 562 | 824 | 877 | 819 |
| GTCCTATGGACTCCGCCCTG | GNRH1 | 447 | 956 | 1582 | 1457 | 1569 |
| TCACCACACAGAAGGTCAAC | GNRH1 | 576 | 341 | 544 | 353 | 376 |
| TTTGATTGATTCATTCCAAG | GNRH1 | 571 | 590 | 949 | 870 | 892 |
| ACTGTGATCAGAGCCTGTAG | GOPC | 415 | 633 | 1377 | 1311 | 1275 |
| ACTTACCAGCTCCTCCACAG | GOPC | 551 | 549 | 1114 | 1007 | 1055 |
| AGGGGGACATCACTTTCGAG | GOPC | 522 | 947 | 1509 | 1603 | 1593 |
| GAACAGCTATATGTCTACGA | GOPC | 625 | 340 | 505 | 560 | 547 |
| GGAACATGGACACCCCGCCG | GOPC | 635 | 651 | 1447 | 1184 | 1265 |
| GTTCCAAAGCTTATCGTGAG | GOPC | 431 | 668 | 1226 | 1009 | 1112 |
| ACTGCAGCCCGGACTCAAGT | GPC3 | 677 | 695 | 1248 | 1351 | 1359 |
| AGATTCAAACCTCGATCACA | GPC3 | 441 | 394 | 740 | 742 | 768 |
| CATCAATGAGTGCCTCCGAG | GPC3 | 682 | 639 | 1070 | 1192 | 1204 |
| CATGTACAGAATCTACGACA | GPC3 | 289 | 349 | 482 | 501 | 555 |
| CCTGGGTTCGGATATCAATG | GPC3 | 391 | 845 | 1581 | 1758 | 1621 |
| TTGTGGGGGCTACTGCAATG | GPC3 | 594 | 1109 | 1915 | 1908 | 1867 |
| AATGGACGAGGTGTAATCAG | GSK3A | 487 | 530 | 1024 | 919 | 905 |
| GGAAGCTAGTGCCCGCACCC | GSK3A | 414 | 477 | 983 | 886 | 856 |
| GGTGGCCCGCCATTTCACCA | GSK3A | 383 | 574 | 961 | 998 | 1000 |
| GTCACGGTGACACACCCCCT | GSK3A | 587 | 547 | 1139 | 1085 | 1060 |
| GTGGTAGCCACTCTAGGCCA | GSK3A | 472 | 860 | 1346 | 1120 | 1289 |
| TCAGCCTTACAATATTGCAG | GSK3A | 669 | 449 | 619 | 568 | 487 |
| ACTGATTATACCTCTAGTAT | GSK3B | 454 | 494 | 963 | 977 | 969 |
| ATACCTTGACATAAATCACA | GSK3B | 574 | 303 | 457 | 342 | 393 |
| CTGTATCAGGATCCAACAAG | GSK3B | 679 | 503 | 906 | 810 | 850 |
| GCAATCGGACTATGTTACAG | GSK3B | 573 | 741 | 1414 | 1408 | 1402 |
| GGTCCTGGGGACACCAACAA | GSK3B | 536 | 611 | 1061 | 1050 | 1039 |
| TCTGTATAGCTGACTTCTTG | GSK3B | 550 | 780 | 1623 | 1520 | 1608 |
| CCTTCCCCCAGGGCGCTGTG | GSTP1 | 608 | 707 | 1254 | 1117 | 1248 |
| CTGGCAGGAGGCAGGAAAGG | GSTP1 | 464 | 513 | 1134 | 950 | 946 |
| GCAGCCTCAAGACTCACCAA | GSTP1 | 476 | 413 | 708 | 698 | 616 |
| GTGCCACCCTCCAGCTGTAC | GSTP1 | 476 | 433 | 848 | 793 | 716 |
| TGGGACAACAGCGTTTCAAA | GSTP1 | 543 | 398 | 677 | 674 | 684 |
| TGGGGTCCCTACCTCGAACA | GSTP1 | 344 | 676 | 1156 | 1036 | 1010 |
| AGAGTCTGCTGTATCGTCTG | GUCY1A2 | 448 | 852 | 1543 | 1528 | 1537 |
| AGTGAGAAATTGTGCTCCGA | GUCY1A2 | 255 | 783 | 1246 | 1160 | 1158 |
| CAGTGCACTGCTAATATACT | GUCY1A2 | 534 | 575 | 851 | 832 | 881 |
| CTTGGTCCTGATCACAAACG | GUCY1A2 | 523 | 704 | 1453 | 1374 | 1302 |
| TGGATGAACTCATGGGCCGA | GUCY1A2 | 520 | 794 | 1279 | 1417 | 1418 |
| TGTGGGGGTCAAACATCAAG | GUCY1A2 | 334 | 736 | 1411 | 1514 | 1529 |
| AATCCATGAATGTACCATCA | GUCY2C | 645 | 664 | 832 | 771 | 799 |
| ACTACAAAGAGACTTTAACC | GUCY2C | 594 | 414 | 640 | 535 | 521 |
| ATACTCATGAAACTAACCTG | GUCY2C | 449 | 513 | 908 | 880 | 875 |
| ATGCTGAATGCGCAGACCTG | GUCY2C | 429 | 628 | 1195 | 1203 | 1124 |
| CCTCCAGTGATTGTTACGTG | GUCY2C | 555 | 789 | 1434 | 1401 | 1345 |
| TCATGCCCTGACACAGTACA | GUCY2C | 389 | 692 | 1333 | 1449 | 1563 |
| CCAGCCAATCAGCTCGCGCA | HDAC1 | 550 | 381 | 695 | 653 | 643 |
| CTTCACAAACTCCACACACT | HDAC1 | 500 | 343 | 486 | 519 | 537 |
| TATGTCTTTCAAGTCAACGT | HDAC1 | 512 | 382 | 339 | 457 | 421 |
| TCATCGATCCCATCTCGGAG | HDAC1 | 525 | 661 | 977 | 1202 | 1089 |
| TGAGTCATACGGATTCGGTG | HDAC1 | 412 | 863 | 1544 | 1372 | 1361 |
| TGATCCTAGGTATCACCAGA | HDAC1 | 291 | 563 | 962 | 822 | 845 |
| AGGAGAAGTAAAGAACGCTG | HDAC10 | 554 | 522 | 1187 | 877 | 870 |
| CCCACCCCGTACCTGCGATG | HDAC10 | 565 | 776 | 1436 | 1330 | 1281 |
| GCATTTCCCATCCCGACCTG | HDAC10 | 551 | 449 | 1031 | 1041 | 1095 |
| GGCCTGGGTTCCTCGCAACA | HDAC10 | 447 | 755 | 1321 | 1173 | 1230 |
| GTTGAACACACAGAATCCGT | HDAC10 | 530 | 218 | 391 | 287 | 301 |
| TCCCCCGGGCAACACAAGTG | HDAC10 | 586 | 734 | 1331 | 1122 | 1191 |
| GCAACAGCGAAGGACCACTG | HDAC11 | 537 | 582 | 1119 | 1077 | 1073 |
| GGAGCTGGAATGGGGTACGG | HDAC11 | 576 | 699 | 1215 | 1314 | 1249 |
| GGCCAGTGTGATGTCCGCGT | HDAC11 | 669 | 414 | 965 | 956 | 912 |
| GTGGTGGAAGCCTCCACCTG | HDAC11 | 488 | 909 | 1584 | 1481 | 1530 |
| TCCCGAACTTCCTTGTGCAG | HDAC11 | 516 | 702 | 1284 | 1257 | 1317 |
| TGACAAGCGTGTGTACATCA | HDAC11 | 246 | 814 | 1411 | 1509 | 1452 |
| GTATAGATGATGAATCCTAT | HDAC2 | 500 | 505 | 753 | 724 | 771 |
| TCAAAGAGTCCATCAAACAC | HDAC2 | 535 | 362 | 710 | 620 | 559 |
| TCACTGTCTGGTGATAGGCT | HDAC2 | 431 | 496 | 1062 | 894 | 1021 |
| TGGGTCATGCGGATTCTATG | HDAC2 | 464 | 748 | 1414 | 1334 | 1332 |
| TTACTTTAGTAATTCAAGGA | HDAC2 | 417 | 322 | 534 | 401 | 431 |
| TTTACCATTACTGATGCTTG | HDAC2 | 563 | 905 | 1639 | 1567 | 1596 |
| AAAGCGGCACATGTCATGCT | HDAC3 | 464 | 495 | 816 | 681 | 716 |
| ACTGTCTCAATGTGCCCTTG | HDAC3 | 690 | 1152 | 2305 | 2029 | 2057 |
| ATTGATATTGACATCCACCA | HDAC3 | 558 | 407 | 797 | 680 | 729 |
| GAGCCTTAATGCCTTCAACG | HDAC3 | 484 | 485 | 1032 | 981 | 1067 |
| GGAGCACAATGCACGTGGGT | HDAC3 | 500 | 434 | 928 | 832 | 830 |
| TGGGTCAATGCCAAGCGATG | HDAC3 | 305 | 676 | 1583 | 1454 | 1516 |
| ACTTACCCGTACCAGTAGCG | HDAC4 | 502 | 579 | 1026 | 1090 | 1079 |
| CAGCTGCACATGAACAAGGT | HDAC4 | 584 | 359 | 718 | 737 | 724 |
| CCAAACCAGAGGTTCACCAC | HDAC4 | 520 | 456 | 615 | 651 | 709 |
| GACCTACCTGTGACGTCCAG | HDAC4 | 407 | 806 | 1349 | 1515 | 1477 |
| GACGCCTACTCACCGCGGAA | HDAC4 | 418 | 537 | 1104 | 1027 | 1130 |
| TGTGCATGGACGGGGACACG | HDAC4 | 394 | 774 | 1622 | 1556 | 1560 |
| AAAGGCAGTTTGTAGGAAGG | HDAC5 | 627 | 720 | 1211 | 1213 | 1210 |
| ACTTACGGTGAGGTGTGAGT | HDAC5 | 621 | 857 | 1424 | 1437 | 1450 |
| CTGAAGCACCAGTGCATGTG | HDAC5 | 410 | 778 | 1219 | 1106 | 1208 |
| GAGGACTGAGTACGACTCAA | HDAC5 | 500 | 612 | 970 | 890 | 871 |
| GCCCTGTGGACCCCGCACTG | HDAC5 | 511 | 710 | 1349 | 1421 | 1422 |
| TGGTGCACTGTTACACACGG | HDAC5 | 666 | 570 | 1018 | 1128 | 1079 |
| ACGACCCAGTACATGAATGA | HDAC6 | 462 | 431 | 771 | 759 | 762 |
| GCAGAGCAAGACCTAATCGT | HDAC6 | 581 | 581 | 966 | 863 | 931 |
| GCAGTACCATTCAAAACCTG | HDAC6 | 451 | 504 | 988 | 951 | 919 |
| GCCATGATCATACCGATGCA | HDAC6 | 393 | 593 | 1223 | 1102 | 1124 |
| GTATTGCATGTTCAACCATG | HDAC6 | 443 | 785 | 1335 | 1390 | 1424 |
| TGGATTTGATGCCCTCCAAG | HDAC6 | 429 | 761 | 1327 | 1323 | 1239 |
| AACACAGGGAGTCAAGCCAT | HDAC7 | 466 | 588 | 1076 | 1118 | 1050 |
| AGCTCCCCATGGACACGCCG | HDAC7 | 353 | 723 | 1179 | 1239 | 1204 |
| AGTCGCCTCAAACTGGACAA | HDAC7 | 566 | 697 | 1364 | 1341 | 1445 |
| CACTGTCGGCAAGGCGAACG | HDAC7 | 386 | 806 | 1353 | 1193 | 1141 |
| GAGAAACGGAGGGATCCAAG | HDAC7 | 548 | 675 | 1254 | 1159 | 1185 |
| GGCAGGGTTGGTCTACGACT | HDAC7 | 472 | 438 | 1024 | 913 | 910 |
| ATGTCGGCTTAGGGAAGGGA | HDAC8 | 486 | 692 | 1126 | 1021 | 1106 |
| CAGGGACACTGTCATGACTT | HDAC8 | 585 | 691 | 1365 | 1408 | 1422 |
| GAGAAGCTGAAAGGCCAGTG | HDAC8 | 373 | 584 | 1059 | 1062 | 1058 |
| GGACTGTAGATGTAAACTGG | HDAC8 | 665 | 661 | 1295 | 1335 | 1338 |
| GGGTTGAGTCCGAAAAAAGC | HDAC8 | 558 | 305 | 490 | 436 | 447 |
| TGGGAATATTACGATTGCGA | HDAC8 | 537 | 526 | 809 | 697 | 657 |
| AACAGCATGAGAACTTGACA | HDAC9 | 517 | 460 | 962 | 912 | 867 |
| AGCTTTGATCCAGTGACGTG | HDAC9 | 576 | 1019 | 1761 | 1677 | 1673 |
| ATTGAACAACTGAAGCAACC | HDAC9 | 496 | 209 | 334 | 345 | 360 |
| GAATACAGAGCATCTGGTCA | HDAC9 | 417 | 704 | 1258 | 1268 | 1248 |
| GCTTTCCCTCTAAAGCAACG | HDAC9 | 433 | 526 | 694 | 538 | 519 |
| TGGAGAACTGCTACTGACTG | HDAC9 | 468 | 713 | 1297 | 1074 | 1188 |
| AATTGCACAGTACTCCCAGG | HGF | 449 | 420 | 956 | 779 | 800 |
| ACTACTGTCGAAATCCTCGA | HGF | 614 | 495 | 856 | 759 | 703 |
| CTGGACTAACATGTTCAATG | HGF | 463 | 856 | 1240 | 1087 | 1157 |
| GGATTTCCCGTGTAACACCA | HGF | 515 | 544 | 659 | 649 | 672 |
| GGGATCATCAGACACCGCAC | HGF | 640 | 343 | 629 | 603 | 658 |
| TTATTGCCGAAATCCAGATG | HGF | 435 | 656 | 1361 | 1281 | 1195 |
| ACTAAAGGACAAGTCACCAC | HIF1A | 303 | 284 | 568 | 533 | 465 |
| AGACAAAGTTCACCTGAGGT | HIF1A | 454 | 513 | 1218 | 1159 | 1183 |
| AGCATTACATCATTATACAA | HIF1A | 364 | 319 | 434 | 510 | 585 |
| AGTGTACCCTAACTAGCCGG | HIF1A | 566 | 722 | 1635 | 1511 | 1484 |
| CAAATACATGGGATTGACTC | HIF1A | 457 | 501 | 1025 | 843 | 783 |
| CCTTACACGCAAATAGCTGA | HIF1A | 450 | 518 | 874 | 799 | 798 |
| AGGAACAGCTAGAACACACC | HIP1 | 533 | 289 | 435 | 443 | 436 |
| CACAGAGCGAGACATGTCCA | HIP1 | 353 | 599 | 1264 | 1177 | 1186 |
| GAGCCCAAGCCAACGAACAG | HIP1 | 426 | 772 | 1143 | 1275 | 1323 |
| GCATAAGTGACCAGGCACAG | HIP1 | 575 | 568 | 1018 | 959 | 1029 |
| GCTGGAAACAGACTTGACCT | HIP1 | 575 | 533 | 1087 | 1003 | 1058 |
| TGAGCCGCTTGAAATATTGG | HIP1 | 278 | 660 | 1377 | 1381 | 1355 |
| AGAGCATCAAGGGTGAACGT | HMGCR | 515 | 358 | 672 | 598 | 552 |
| CACAACAGAAGGTTGTCTCG | HMGCR | 535 | 580 | 1228 | 990 | 1028 |
| CCAGAAGGAGAGCTAAACCT | HMGCR | 660 | 353 | 619 | 523 | 570 |
| CTCCTTACTTGATACTTCGG | HMGCR | 446 | 585 | 1027 | 1063 | 940 |
| CTGATGGAAACCCATGAACG | HMGCR | 320 | 475 | 1075 | 927 | 897 |
| GTTCATGGACATCATACAGA | HMGCR | 464 | 443 | 535 | 514 | 427 |
| AAAGCTCTTTCACCATGCCT | HMGCS1 | 530 | 410 | 552 | 472 | 496 |
| AACAAACTTCTCTTACCCAA | HMGCS1 | 586 | 357 | 524 | 480 | 476 |
| ACAACTAATGCATGCTATGG | HMGCS1 | 382 | 611 | 1156 | 1066 | 1102 |
| AGCAGCGGTCTAGTGCACTG | HMGCS1 | 452 | 653 | 1442 | 1216 | 1192 |
| GCCCAATGCAATCATACGAA | HMGCS1 | 513 | 289 | 576 | 481 | 474 |
| GGAAATGCTAGACCTACAGG | HMGCS1 | 510 | 423 | 955 | 887 | 970 |
| ACAGCCCCAGAATCTCATCA | HNF1A | 507 | 555 | 1046 | 943 | 1047 |
| AGACACGCACCTCTGTGACA | HNF1A | 731 | 766 | 1332 | 1225 | 1200 |
| AGCAGCATAACATCCCACAG | HNF1A | 570 | 677 | 1146 | 1198 | 1127 |
| GCGGAGGAATCGTTTCAAGT | HNF1A | 513 | 630 | 931 | 995 | 989 |
| GTACGTCCGCAAGCAGCGAG | HNF1A | 470 | 539 | 1142 | 939 | 956 |
| TGGACAGCAAGCAACTAGCG | HNF1A | 428 | 581 | 1142 | 1023 | 986 |
| AGGGCAGCACCAACTCGTCG | HNF4A | 563 | 517 | 1078 | 989 | 1004 |
| CGCATTGCCGACTAAACCTG | HNF4A | 474 | 896 | 1742 | 1850 | 1732 |
| GAACACCATGGATCGCTTGG | HNF4A | 455 | 723 | 1517 | 1438 | 1430 |
| GCGACCCAGGCAAGATCAAG | HNF4A | 398 | 393 | 841 | 825 | 733 |
| GGCACCGTAGTGTTTCCCTG | HNF4A | 439 | 717 | 1382 | 1338 | 1366 |
| GGGACCGGATCAGCACTCGC | HNF4A | 644 | 535 | 766 | 766 | 879 |
| AAGTACAAGAAGGATCAGAA | HOXA3 | 119 | 378 | 536 | 615 | 607 |
| CCTACTCAACTCACCTACCG | HOXA3 | 455 | 300 | 594 | 549 | 565 |
| GCGCCGATGGCGAGTACCAC | HOXA3 | 727 | 531 | 826 | 949 | 987 |
| GCGTGTCTGCGCACCCTGAG | HOXA3 | 490 | 913 | 1310 | 1401 | 1480 |
| GGCATTATAAGCGAACCCGT | HOXA3 | 590 | 235 | 389 | 358 | 383 |
| TTGGACGACGCCTGCCCAGG | HOXA3 | 468 | 873 | 1297 | 1252 | 1384 |
| GCTCATCCAGAACCACTTCG | HRAS | 439 | 638 | 1304 | 1223 | 1201 |
| GCTGGACATCCTGGACACAG | HRAS | 572 | 487 | 1014 | 1093 | 1101 |
| GGAGCAGATCAAGCGAGTGA | HRAS | 379 | 568 | 1474 | 1171 | 1226 |
| GGGACCAGTACATGCGCACG | HRAS | 424 | 461 | 1029 | 929 | 999 |
| TGACCTGGCTGCTCGCACCG | HRAS | 371 | 510 | 1244 | 1218 | 1249 |
| TGCTTCCGATAGGAGTCCTG | HRAS | 439 | 682 | 1302 | 1189 | 1257 |
| ATGCAGGATCACTGTCAGCG | HRH2 | 534 | 743 | 1159 | 1132 | 1136 |
| CCCCAGATGAATAGACAGGA | HRH2 | 397 | 658 | 1035 | 1112 | 1155 |
| CTGCAATATCTATACCAGCT | HRH2 | 385 | 620 | 988 | 1000 | 1000 |
| GGACTGGGGTGATAAGCACA | HRH2 | 626 | 764 | 1508 | 1260 | 1254 |
| TGAAGCAGTTAGTCAGACTG | HRH2 | 350 | 765 | 1351 | 1325 | 1301 |
| TTGACCTGGACTTTGCACTT | HRH2 | 562 | 742 | 1426 | 1255 | 1254 |
| ACGTCTTGGCACAAAGAGAA | HSP90AA1 | 632 | 620 | 765 | 821 | 705 |
| CACACTTACTCAGATACTCA | HSP90AA1 | 657 | 288 | 563 | 511 | 515 |
| CACTCACAAAGAGAGTAATG | HSP90AA1 | 623 | 571 | 886 | 862 | 786 |
| CGACGACGAGCAGTATGCCT | HSP90AA1 | 526 | 258 | 506 | 453 | 494 |
| CTACACCTCAGGTGAGACCA | HSP90AA1 | 335 | 637 | 1291 | 1191 | 1159 |
| GCAGTTCTCGAAAAACATCA | HSP90AA1 | 427 | 376 | 449 | 396 | 415 |
| ATACTCACATAAAGAGTGAT | HSP90AB1 | 460 | 523 | 1134 | 1079 | 1107 |
| CATCTCCAGATTGGGAAGTG | HSP90AB1 | 520 | 629 | 1171 | 1049 | 1123 |
| CCCGGCCAATGGGCTCACCT | HSP90AB1 | 339 | 419 | 904 | 846 | 825 |
| GGTCAAAAGGAGCCCGACGA | HSP90AB1 | 474 | 434 | 865 | 754 | 831 |
| GTGAAACAGTACATACCTGA | HSP90AB1 | 502 | 295 | 413 | 430 | 407 |
| TCGATTACAGACTTCATCCG | HSP90AB1 | 526 | 655 | 1158 | 1153 | 1153 |
| CAAGACGTGTTCGATTTGAG | HSP90B1 | 607 | 774 | 1445 | 1339 | 1332 |
| CAATAACCTTATCTGCGACA | HSP90B1 | 594 | 356 | 584 | 474 | 553 |
| GAAGAGTTGGTTAAAAACCT | HSP90B1 | 530 | 522 | 802 | 725 | 771 |
| GTTTCCCGGGAAACATTCAA | HSP90B1 | 555 | 487 | 663 | 566 | 568 |
| TAAACTTCCCTATTTATGTG | HSP90B1 | 766 | 831 | 1405 | 1333 | 1387 |
| TATTCATCAAACAGACCACG | HSP90B1 | 599 | 383 | 610 | 515 | 523 |
| AGTGGTTCATGTCGTCCAAC | ID3 | 389 | 443 | 907 | 794 | 815 |
| CCCGCAAGCGCGAGTAGCAG | ID3 | 531 | 786 | 1374 | 1305 | 1288 |
| GGTGCGCGGCTGCTACGAGG | ID3 | 525 | 698 | 1206 | 1105 | 1099 |
| TGGCCAGGCTGCGTTCGGAC | ID3 | 608 | 442 | 748 | 694 | 699 |
| TGGCTAAGCTGAGTGCCTCG | ID3 | 685 | 1156 | 1834 | 2109 | 2009 |
| TGTAGTCGATGACGCGCTGC | ID3 | 339 | 780 | 1500 | 1392 | 1427 |
| AGATGGCTCTGTCTAAGAGT | IDH1 | 525 | 570 | 1420 | 1267 | 1244 |
| AGTTCAAGTTGAAACAAATG | IDH1 | 523 | 920 | 1442 | 1514 | 1523 |
| CAAGCTATGAAATCCGAGGG | IDH1 | 532 | 863 | 1320 | 1378 | 1431 |
| CCCATCCGCTCACAAGCCGG | IDH1 | 281 | 671 | 1056 | 992 | 1012 |
| GCTATAAAGAAGTACAACGT | IDH1 | 524 | 505 | 666 | 654 | 637 |
| TACCTTCAAAGTTATGTACC | IDH1 | 569 | 513 | 844 | 777 | 776 |
| ACCATTGGCAGACACGCCCA | IDH2 | 531 | 794 | 1483 | 1409 | 1410 |
| AGCCACACTGTACTTCTGGG | IDH2 | 389 | 849 | 1303 | 1279 | 1371 |
| CCATCCGGAACATCCTTGGG | IDH2 | 647 | 766 | 1339 | 1273 | 1388 |
| CTAGCTCATCCTACCCCACG | IDH2 | 540 | 709 | 1329 | 1301 | 1351 |
| GAGGTGTACAACTTCCCCGG | IDH2 | 569 | 376 | 789 | 747 | 728 |
| GGCCTGCAAGAACTACGATG | IDH2 | 456 | 722 | 1249 | 1201 | 1276 |
| AGTTCGTGTGCGGAGACAGG | IGF1 | 552 | 867 | 1653 | 1673 | 1630 |
| CAGCACTTAAATAATTGGGT | IGF1 | 538 | 476 | 1045 | 885 | 995 |
| CAGGTAGAAGAGGTGCGAGG | IGF1 | 834 | 640 | 1146 | 1188 | 1099 |
| GGCACCTCAGACAGGCATCG | IGF1 | 722 | 766 | 1437 | 1452 | 1387 |
| TCCCCAGCCCCGCCACAGCT | IGF1 | 495 | 528 | 942 | 799 | 933 |
| TTTGCACAGACAAGCCCACA | IGF1 | 596 | 633 | 1153 | 1106 | 1105 |
| AAAATATGGATCCCAAATCG | IGF1R | 640 | 571 | 888 | 930 | 910 |
| AATCTGAGGAACATTACTCG | IGF1R | 506 | 746 | 1439 | 1296 | 1283 |
| CCGAAGCAGGAACACCACGG | IGF1R | 509 | 476 | 766 | 689 | 671 |
| CCTTCGCCAGATCTTAGGAG | IGF1R | 463 | 597 | 1162 | 1106 | 1091 |
| GATGATGCGATTCTTCCATG | IGF1R | 397 | 746 | 1184 | 1105 | 1045 |
| TCAGTTACTACATCGTGAGG | IGF1R | 485 | 606 | 1129 | 965 | 1041 |
| AACTTCCCCAGATACCCCGT | IGF2 | 638 | 354 | 624 | 614 | 642 |
| CACCGGAAGCACGGTCGGAG | IGF2 | 694 | 671 | 1482 | 1278 | 1290 |
| CCGCAGCAGCCGTGGCATCG | IGF2 | 420 | 835 | 1245 | 1333 | 1300 |
| CGCCGCACAGAGTCTCACTG | IGF2 | 638 | 593 | 1146 | 1184 | 1177 |
| GGCACAGTAGGTCTCCAGAA | IGF2 | 475 | 466 | 899 | 860 | 792 |
| GGTCCCCACAGACAAACTGG | IGF2 | 484 | 561 | 1107 | 972 | 955 |
| ACTGTGACCGTAGCACCCAG | IGF2R | 543 | 872 | 1512 | 1345 | 1486 |
| AGCTGACTACCTACACCACG | IGF2R | 484 | 624 | 1127 | 1200 | 1117 |
| ATGGCACTTACCTACGCCAG | IGF2R | 497 | 520 | 964 | 1019 | 1052 |
| TATGTCAACGGTGACAAGTG | IGF2R | 435 | 598 | 1102 | 1199 | 1079 |
| TGTCTTAGACAGCACGCAGG | IGF2R | 462 | 649 | 1474 | 1252 | 1338 |
| TTGCCGCTATGAAATCGAGT | IGF2R | 534 | 350 | 672 | 566 | 593 |
| ATTGGGCCCGTACACAGGGT | IKBKB | 637 | 830 | 1299 | 1146 | 1209 |
| CGTGATGCACTCAAAAGCCA | IKBKB | 409 | 342 | 655 | 647 | 610 |
| GCCCGAATCCAGCATGACAC | IKBKB | 528 | 314 | 594 | 495 | 519 |
| TAGGCTGAACCACCCCAATG | IKBKB | 463 | 757 | 1360 | 1353 | 1340 |
| TATTGACCTAGGATATGCCA | IKBKB | 623 | 546 | 1120 | 1016 | 1026 |
| TTCCAACTGAGGAAAGTGTG | IKBKB | 387 | 838 | 1675 | 1772 | 1739 |
| ATCATCATCTAGCTCCCGGG | IKBKE | 500 | 497 | 883 | 874 | 866 |
| GAAGACATTCGGGGTAACCG | IKBKE | 386 | 629 | 1250 | 1340 | 1329 |
| GGCAGATTACACCACCGCCA | IKBKE | 419 | 545 | 896 | 778 | 752 |
| GTGATGGAGTACTGCTCTAG | IKBKE | 509 | 698 | 1402 | 1285 | 1312 |
| TCAACACGGCAAGCTACCTG | IKBKE | 568 | 597 | 1268 | 1259 | 1269 |
| TCTGCTCATATACTGCCTCC | IKBKE | 713 | 529 | 784 | 867 | 814 |
| ACCTGAGGACGCACTCCGGT | IKZF1 | 503 | 391 | 799 | 811 | 847 |
| ATGTGGTGGAAAGGTCTTCG | IKZF1 | 438 | 832 | 1533 | 1548 | 1469 |
| CACAACTACTTGCAAAGCAT | IKZF1 | 496 | 310 | 684 | 705 | 726 |
| GATGGCTTGGTCCATCACGT | IKZF1 | 509 | 637 | 1181 | 1223 | 1192 |
| GCTGAATGGGGAAGAATGTG | IKZF1 | 645 | 649 | 1476 | 1470 | 1427 |
| TGTGGACCATGAGCACATTG | IKZF1 | 500 | 1137 | 1741 | 1701 | 1692 |
| AAGATGAATTGCGATGTGTG | IKZF3 | 539 | 673 | 1150 | 1347 | 1404 |
| ACTGGAACCATCTCTGATGT | IKZF3 | 552 | 675 | 1327 | 1213 | 1282 |
| AGAGTGAGATAATACAGACC | IKZF3 | 351 | 578 | 1038 | 975 | 1039 |
| GGACAGATTAGCAAGCAATG | IKZF3 | 549 | 528 | 1127 | 1061 | 1054 |
| TAAGCAGAGAAGCTCCCTTG | IKZF3 | 546 | 881 | 1719 | 1881 | 1859 |
| TGTGCAGTTTAATGTGGCGG | IKZF3 | 443 | 709 | 1389 | 1356 | 1401 |
| AGTCTGCATCCGGATATGGC | IL12RB1 | 511 | 489 | 926 | 1016 | 987 |
| ATACGGTACCCATTCCCATG | IL12RB1 | 418 | 886 | 1354 | 1305 | 1312 |
| CCCCGCCAGAGTTGGACCCG | IL12RB1 | 551 | 702 | 1116 | 1021 | 1147 |
| CTGCGCTGGCCAATGTATGG | IL12RB1 | 442 | 588 | 1098 | 1224 | 1301 |
| GGTCATAGGCCGCACCAGAG | IL12RB1 | 10 | 884 | 1506 | 1444 | 1559 |
| TAAATACGACCCTCCGCCAG | IL12RB1 | 526 | 607 | 1060 | 947 | 1086 |
| ATTATGTGAATGACTTGGTA | IL21 | 653 | 546 | 1154 | 1088 | 1152 |
| CAACTAAAGGCAGCAAACAC | IL21 | 648 | 427 | 503 | 486 | 539 |
| CTAAAGGCAGCAAACACAGG | IL21 | 536 | 540 | 1041 | 915 | 867 |
| GCCTGATGGTCATCTTCTTG | IL21 | 699 | 1001 | 1472 | 1462 | 1458 |
| TCAAGAGAAGATCTTGCTCT | IL21 | 439 | 935 | 1297 | 1335 | 1269 |
| TCTTTCAGAGGCATTGTGAG | IL21 | 611 | 648 | 1177 | 1169 | 1209 |
| CCTGCGCAGAGCGCCAGCAA | IL21R | 631 | 535 | 1116 | 1071 | 1042 |
| GATGTTATAATGTCCCGAGA | IL21R | 701 | 425 | 783 | 789 | 790 |
| GGGGAGAAAGCTGATCTCCG | IL21R | 537 | 602 | 1213 | 1224 | 1225 |
| TCTTCAGGCCTAAGAAGACG | IL21R | 440 | 789 | 1503 | 1462 | 1553 |
| TGAAGTGGAACACATCCATG | IL21R | 470 | 893 | 1838 | 1534 | 1444 |
| TTCTTCCAGCCCCTCTACGT | IL21R | 317 | 424 | 755 | 836 | 856 |
| CCTACATCTTCCCAAACTGG | IL6 | 518 | 681 | 1008 | 1050 | 1071 |
| GAACTGATTAAGTACATCCT | IL6 | 620 | 488 | 876 | 885 | 966 |
| GAGTGGTAGACTATTTGAAG | IL6 | 399 | 573 | 1140 | 1093 | 1090 |
| GGAAAGCAGTAGCCATCACC | IL6 | 609 | 562 | 967 | 877 | 855 |
| TATCCTCCAGAACAACTATG | IL6 | 630 | 384 | 540 | 541 | 545 |
| TGCTTGACAAGAATCACTAC | IL6 | 625 | 330 | 556 | 457 | 515 |
| AACCTTGAAAGTCATAGGCA | IL6ST | 540 | 638 | 881 | 1003 | 1015 |
| ATCTGACGTAACCTTCCCAA | IL6ST | 608 | 368 | 700 | 731 | 836 |
| ATCTGTACAACTCATATGGA | IL6ST | 337 | 613 | 1193 | 1283 | 1265 |
| CTGTTCGGACAAAAAAAGTG | IL6ST | 668 | 639 | 1054 | 1136 | 1119 |
| GACAGTGAATGAATATCGGG | IL6ST | 375 | 570 | 1150 | 1281 | 1322 |
| GTATACCTCTTAAATAGGTG | IL6ST | 613 | 624 | 1108 | 1205 | 1271 |
| AAAAAATGAAAATGACTGGA | IL7R | 410 | 387 | 722 | 732 | 787 |
| CAATGTATGAGATTAAAGTC | IL7R | 515 | 385 | 929 | 947 | 932 |
| GAATGAATAATCATCCAGCT | IL7R | 387 | 342 | 653 | 660 | 587 |
| GTAAGAGTCATCTATCGAGA | IL7R | 506 | 632 | 931 | 951 | 873 |
| GTCAGGGTCATCAAAAGTGC | IL7R | 558 | 367 | 731 | 664 | 709 |
| TCAGAATTAAGCCTATGGTG | IL7R | 635 | 702 | 1352 | 1321 | 1310 |
| AGAAGGCAGTGACCTGTCGG | INHBA | 597 | 752 | 1340 | 1385 | 1277 |
| CCAGGACCAAAGTCACCATC | INHBA | 460 | 475 | 609 | 617 | 645 |
| CTGAAAAGGTGGTGGACGCA | INHBA | 666 | 673 | 1218 | 1123 | 1190 |
| GCAGGCAATCCGAACGTCCA | INHBA | 636 | 645 | 1388 | 1238 | 1296 |
| GCTGCACGTAGGCAAAGTCG | INHBA | 308 | 750 | 1441 | 1188 | 1212 |
| TGAACTCATGGAGCAGACCT | INHBA | 587 | 661 | 1179 | 1127 | 984 |
| AGACCCGCTCTTTCTGACTG | INPP4B | 702 | 808 | 1400 | 1477 | 1405 |
| ATTAAGCAGTAGGTCTGCAT | INPP4B | 864 | 577 | 844 | 782 | 770 |
| CACGTCACGACAGACGTGCA | INPP4B | 410 | 597 | 925 | 1030 | 965 |
| CTCTACGATGTCATCACCGT | INPP4B | 595 | 553 | 792 | 898 | 787 |
| GAACTGCATTATCGACATGG | INPP4B | 455 | 729 | 1289 | 1095 | 1165 |
| TTCCGTAGATTGTCCCAGTG | INPP4B | 517 | 786 | 1125 | 1217 | 1168 |
| ACACAAGTTTAAGAGAGAGT | INSIG1 | 444 | 504 | 887 | 846 | 830 |
| GAACGTCACGCTGTTCCCCG | INSIG1 | 548 | 573 | 996 | 949 | 935 |
| GGAGGGCTGGCGGCTAAAGT | INSIG1 | 473 | 491 | 1091 | 919 | 893 |
| GGCACCACCACCTGGTGCGA | INSIG1 | 476 | 681 | 1385 | 1317 | 1296 |
| GGGGCTGCGAGGCCCGTGAG | INSIG1 | 513 | 786 | 1705 | 1714 | 1712 |
| TCCTCTGAAGGATACAGCAC | INSIG1 | 566 | 394 | 676 | 672 | 702 |
| AGTCTCTACTTCTGCCTATG | INSL6 | 550 | 1155 | 1706 | 1638 | 1647 |
| CTGATTGATATCTTGTGATG | INSL6 | 828 | 877 | 1651 | 1636 | 1592 |
| GCGACATCAGCAAAGCTAGG | INSL6 | 488 | 358 | 576 | 580 | 679 |
| TGTCAATTGGAGCTACTTCG | INSL6 | 289 | 677 | 1429 | 1258 | 1303 |
| TGTCTTATCAGGTGGCAAGT | INSL6 | 511 | 658 | 1109 | 1132 | 1085 |
| TTGAAACCTTCATCCCGGAC | INSL6 | 482 | 355 | 638 | 578 | 628 |
| AAACCGAGATTGGATCCAGG | INSR | 435 | 534 | 1071 | 874 | 918 |
| AGTTGCCTAAACACTCGCTG | INSR | 20 | 455 | 849 | 810 | 796 |
| CTAATCATCAACATTCGAGG | INSR | 568 | 292 | 509 | 493 | 476 |
| GATGCGGGACCAGTCGATGG | INSR | 502 | 669 | 1246 | 1254 | 1269 |
| TGAGGATTACCTGCACAACG | INSR | 310 | 654 | 1073 | 990 | 930 |
| TTACCTCATCGCCATACCGT | INSR | 593 | 329 | 572 | 598 | 588 |
| AGGTGTCCAAGAGTTGAGCA | INTS4 | 405 | 564 | 1041 | 1022 | 1113 |
| ATACTGCAAAGCATCTGCTG | INTS4 | 736 | 640 | 1287 | 1189 | 1286 |
| CAGTATCAAGCTGATCCTCT | INTS4 | 539 | 503 | 992 | 809 | 775 |
| GGATTGGAAGATGAGATGTA | INTS4 | 667 | 697 | 1372 | 1293 | 1160 |
| TCAGTCAGCTCTATCCTGAA | INTS4 | 697 | 388 | 1080 | 782 | 814 |
| TCCCAGTATTGCACCCCATG | INTS4 | 474 | 594 | 1250 | 1076 | 1150 |
| ACAGCGGGTACATGCGCATG | IRS2 | 604 | 710 | 1616 | 1511 | 1660 |
| AGCCGCTCGCACACCCTGAG | IRS2 | 477 | 921 | 1477 | 1475 | 1384 |
| GCTGTACGGGTACATGACCA | IRS2 | 402 | 812 | 1292 | 1353 | 1379 |
| GGCCTACAACCCGTATCCCG | IRS2 | 582 | 574 | 1058 | 1205 | 1166 |
| GGCGCCCTACACGTGCAACG | IRS2 | 488 | 16 | 24 | 17 | 19 |
| TCGTAGTACTCGAGCCGCGG | IRS2 | 561 | 549 | 960 | 1003 | 998 |
| CTCCCAGGTACTATTCAAGG | ITGA2 | 386 | 611 | 865 | 837 | 900 |
| GAAGACGTGTGGTCCCCTGT | ITGA2 | 461 | 646 | 1092 | 1208 | 1241 |
| GCAACAATGACAATATACTG | ITGA2 | 545 | 348 | 476 | 474 | 495 |
| GTACACCATCAACATACAGG | ITGA2 | 505 | 374 | 512 | 452 | 427 |
| TATATCTACAATGGTCACGA | ITGA2 | 579 | 355 | 517 | 575 | 539 |
| TCCCAGACATACCAATATGG | ITGA2 | 620 | 479 | 807 | 784 | 770 |
| ACTCATGCTACAACGCAATG | ITK | 680 | 1149 | 1860 | 1936 | 1926 |
| ATACTTTGAGAATCGTCACG | ITK | 567 | 717 | 1577 | 1456 | 1579 |
| CAGAAATTAGGATGGTACAT | ITK | 551 | 416 | 1012 | 796 | 842 |
| CAGGTCTGGTCACTAGACTC | ITK | 425 | 592 | 1075 | 1080 | 1050 |
| TCTGGAATCTCGTACCATGA | ITK | 464 | 582 | 1189 | 1171 | 1220 |
| TCTTGGCTACTGGCTTAACA | ITK | 548 | 424 | 824 | 894 | 894 |
| AAACGGCCGTGCGTGCACAG | ITPA | 392 | 597 | 1047 | 1009 | 1029 |
| AGATAAGTTTCCATGCACTT | ITPA | 301 | 543 | 1113 | 1051 | 1064 |
| CAAATCCGGCCGTGCGAGCG | ITPA | 625 | 660 | 1120 | 1170 | 1206 |
| CACTTTCGCATTCAGCACCG | ITPA | 562 | 466 | 821 | 750 | 830 |
| CAGGGACCACTCACATGTAG | ITPA | 301 | 528 | 997 | 988 | 970 |
| GATTTCCATACAGAAGTGTC | ITPA | 582 | 468 | 904 | 854 | 876 |
| ACACGGACACACATCTACTC | JAK1 | 620 | 346 | 600 | 604 | 559 |
| ATGTTGTGGACGATCAATGG | JAK1 | 524 | 771 | 1287 | 1334 | 1394 |
| GAGCGATGTCTTGTCATCAA | JAK1 | 401 | 421 | 954 | 911 | 959 |
| TACACTTACCAAGTAACTCA | JAK1 | 606 | 330 | 540 | 536 | 485 |
| TGAATAAGTCCATCAGACAG | JAK1 | 411 | 769 | 1193 | 1229 | 1239 |
| TGGTTTCATCCGAACGACAG | JAK1 | 450 | 715 | 1302 | 1256 | 1231 |
| AGAGCTATAGATATCCAGTG | JAK2 | 571 | 792 | 1527 | 1539 | 1504 |
| AGTGGCTAAACAGTTGGCAT | JAK2 | 535 | 677 | 1228 | 1149 | 1061 |
| ATATCTGAAGTTTCCAACTG | JAK2 | 575 | 811 | 1432 | 1214 | 1255 |
| CAACATCACCAACATTACAG | JAK2 | 642 | 301 | 417 | 348 | 352 |
| TATCGGCACGGAATATCTCG | JAK2 | 614 | 650 | 1283 | 1348 | 1269 |
| TCTTCAGGAGAGAATACCAT | JAK2 | 392 | 426 | 814 | 758 | 744 |
| ACTCTCCAGGCTTAGCACAG | JAK3 | 548 | 607 | 1099 | 1112 | 1051 |
| AGTCCAACCTGATTGTGGTG | JAK3 | 575 | 1269 | 2172 | 2200 | 2185 |
| AGTCCTTTCGTAGCCCGAAG | JAK3 | 655 | 856 | 1419 | 1460 | 1457 |
| CAAGCACAAAAACTGCATGG | JAK3 | 439 | 559 | 1029 | 1078 | 1049 |
| CCTTGTAATCAGGACCCAAG | JAK3 | 417 | 563 | 1019 | 1026 | 994 |
| TACCCCCACTCGTAGCTACA | JAK3 | 474 | 431 | 777 | 803 | 836 |
| GAAATACTACATCGAGTACA | JMJD6 | 464 | 561 | 1010 | 941 | 1014 |
| GCTCTCGTAGTAGTTGTGAC | JMJD6 | 529 | 435 | 760 | 710 | 699 |
| GCTGTCCGAGGCAATACCTG | JMJD6 | 350 | 504 | 1083 | 990 | 1069 |
| GGAATTCACATCGACCCTCT | JMJD6 | 387 | 556 | 962 | 868 | 846 |
| TATGGCACAAGACGGTAAGA | JMJD6 | 562 | 733 | 1446 | 1310 | 1402 |
| TGAACGACCCTACAAGCCCG | JMJD6 | 644 | 604 | 1078 | 975 | 973 |
| AACACTGCTTCTGCACATAG | JMJD7 | 584 | 393 | 607 | 523 | 554 |
| AGGGCGTTACGGATGATGCA | JMJD7 | 628 | 829 | 1671 | 1726 | 1608 |
| CCCCTACCATAGGGGATGAA | JMJD7 | 543 | 649 | 1536 | 1417 | 1338 |
| CTACGAAAACCTCTACTGTG | JMJD7 | 541 | 775 | 1245 | 1284 | 1321 |
| GTTATGCAGATGCTGTACGA | JMJD7 | 569 | 490 | 1006 | 900 | 882 |
| TGTGAACTTCTGGCTAGGAG | JMJD7 | 460 | 582 | 1019 | 1041 | 1014 |
| AACGTGACGGACGAGCAGGA | JUN | 482 | 901 | 1540 | 1568 | 1593 |
| CAGCTTGAGCAGCCCCACGT | JUN | 502 | 399 | 595 | 526 | 554 |
| CAGTCCGAGAGCGGCGCCTA | JUN | 458 | 699 | 1479 | 1244 | 1271 |
| CTGAACCTGGCCGACCCGGT | JUN | 447 | 436 | 578 | 544 | 550 |
| CTGAGGTTGGCGTAGACCGG | JUN | 395 | 716 | 1297 | 1225 | 1392 |
| GCTGCCCAGCGTCACGTCGG | JUN | 522 | 819 | 1596 | 1555 | 1595 |
| ACCAAGACAGAAACTACAGA | KAT6A | 447 | 344 | 741 | 552 | 575 |
| CAAAACGCTCTATTATGACG | KAT6A | 359 | 685 | 1123 | 1016 | 1057 |
| CATCCTGTGAGGACAAACGA | KAT6A | 498 | 743 | 828 | 863 | 866 |
| CTCGAGGTTTCCGCCCATCG | KAT6A | 564 | 579 | 1426 | 1423 | 1434 |
| GGAAGTCCAGCATTCTTAGG | KAT6A | 513 | 768 | 1235 | 1292 | 1272 |
| TTTGCAGGAAGACTATCACG | KAT6A | 503 | 444 | 1059 | 978 | 935 |
| AACGTGGTCTATCTAAGTGG | KAT6B | 453 | 632 | 1208 | 1071 | 1082 |
| AATGGAACAAGCTAGCTGCT | KAT6B | 564 | 422 | 945 | 785 | 835 |
| CTTACCCACAGGAATATGCA | KAT6B | 636 | 617 | 966 | 953 | 1022 |
| GAAAGATGGACCGCAGTACA | KAT6B | 548 | 758 | 1245 | 1205 | 1232 |
| TGTGGTTTAGTCAATCTCGG | KAT6B | 441 | 541 | 1043 | 1080 | 1101 |
| TTACGACGCATACGTTTGGG | KAT6B | 491 | 949 | 1521 | 1501 | 1451 |
| AGATCAGACTCCCCTCCATG | KCNIP1 | 510 | 1179 | 1745 | 1981 | 2042 |
| AGATGACCATGGTTTGCCAT | KCNIP1 | 404 | 726 | 1022 | 1236 | 1211 |
| CAAGCATACCTCGAACTTCA | KCNIP1 | 426 | 414 | 544 | 547 | 540 |
| CCCAGACCAACTTTACCAAG | KCNIP1 | 518 | 546 | 981 | 851 | 794 |
| CCGTCCATGAGAAGCTAAGG | KCNIP1 | 539 | 613 | 1241 | 1364 | 1363 |
| TGGTATTACCAGTATCAGAG | KCNIP1 | 511 | 622 | 1096 | 951 | 917 |
| AATTGGAAGCCAATCCCCCA | KDM1A | 398 | 440 | 1035 | 906 | 975 |
| AGAGTAGACTTCCTCATGAC | KDM1A | 400 | 516 | 672 | 719 | 708 |
| GAATGACAACTTCCAATGCC | KDM1A | 461 | 418 | 686 | 709 | 722 |
| GCTCTAAATAACTGTGAACT | KDM1A | 691 | 699 | 1074 | 945 | 1119 |
| GGGAATCGCCGAGACTCCGG | KDM1A | 598 | 514 | 1046 | 1001 | 1053 |
| TCTGAAAAACAGGATCGAGT | KDM1A | 465 | 260 | 417 | 443 | 428 |
| CTGTGCAACAATGTTCACAC | KDM3A | 375 | 420 | 579 | 624 | 540 |
| CTTTAACTAAAAACGTTGTG | KDM3A | 527 | 800 | 1314 | 1339 | 1307 |
| TAACAGCCCTAATAGTTGTG | KDM3A | 530 | 740 | 1473 | 1578 | 1583 |
| TGCCTTGGGTCCTTACTAGG | KDM3A | 367 | 674 | 1114 | 1133 | 1045 |
| TGGCTGGCAGACCTAACCAG | KDM3A | 522 | 670 | 1083 | 1101 | 1132 |
| TTGAATTGCCTGTTAGAACA | KDM3A | 317 | 734 | 1198 | 1205 | 1282 |
| ACTGTACACAGCTATGCCAA | KDM4A | 359 | 459 | 995 | 1035 | 977 |
| ATATCATCATAGGACGCTCG | KDM4A | 553 | 687 | 1474 | 1504 | 1550 |
| ATCATACAGACAGCGCAGTG | KDM4A | 587 | 706 | 1281 | 1314 | 1302 |
| GGCCCCGAGAGCCAGAAACG | KDM4A | 382 | 774 | 1277 | 1399 | 1328 |
| TACTGGACAGTACGACATGA | KDM4A | 507 | 991 | 1576 | 1651 | 1520 |
| TCTTCAAACTCACTGTACCG | KDM4A | 606 | 494 | 782 | 764 | 817 |
| AGGACTCACCGATCACTACG | KDM4B | 457 | 486 | 862 | 939 | 891 |
| CAGACACTGGAGAAAGACTG | KDM4B | 458 | 634 | 1077 | 1126 | 1114 |
| CATGCCGAAGTACAGATAGG | KDM4B | 506 | 623 | 1248 | 1209 | 1075 |
| CTACCAACTTCGCTACTCTG | KDM4B | 456 | 504 | 885 | 765 | 778 |
| GGCCGAGAGGAAGTTCAACG | KDM4B | 538 | 633 | 1179 | 1260 | 1112 |
| GGGGGCACGACGTTGAGAGG | KDM4B | 444 | 989 | 1438 | 1414 | 1459 |
| AAATCCCAGCGATCTCCAGT | KDM4C | 533 | 274 | 548 | 478 | 427 |
| CACATTTGCATGGCACACCG | KDM4C | 549 | 609 | 821 | 783 | 883 |
| GATGAAGTGGTTACTCCGGG | KDM4C | 444 | 652 | 1235 | 1198 | 1162 |
| GCCTGAGGTTCCATCGATTG | KDM4C | 628 | 970 | 1761 | 1657 | 1688 |
| TTCAGGCAGTTGGCTAACAG | KDM4C | 351 | 641 | 1112 | 997 | 1040 |
| TTCTACCCTACAGATCACTC | KDM4C | 358 | 270 | 475 | 492 | 496 |
| AACACCCCCTACCTGTACTT | KDM4D | 523 | 482 | 831 | 779 | 832 |
| CAGCGGGATCCCCTTCAGTC | KDM4D | 628 | 513 | 814 | 833 | 787 |
| CTTGTTTGATGAAAACACGA | KDM4D | 620 | 346 | 540 | 391 | 427 |
| GAAGAAGAGAGCCACGACAG | KDM4D | 387 | 380 | 808 | 915 | 884 |
| GCCGTAGTCGATCCAGCGCG | KDM4D | 542 | 551 | 851 | 831 | 809 |
| GTCCCAAGGTGCACACCGAG | KDM4D | 447 | 803 | 1457 | 1521 | 1488 |
| AGTGTGTCAATCATTACCGG | KDM5A | 441 | 751 | 1300 | 1222 | 1269 |
| ATGCAAATGAGACAACGGAA | KDM5A | 622 | 513 | 769 | 643 | 684 |
| ATTCTATCAACTACTTACAC | KDM5A | 429 | 253 | 368 | 282 | 329 |
| CGTGCTCCTGTAATCCCGAG | KDM5A | 599 | 541 | 1060 | 1145 | 1083 |
| GACTCGGACCAAACTGACAG | KDM5A | 472 | 849 | 1093 | 986 | 1010 |
| TATATACCAATGTCAGTCCG | KDM5A | 358 | 583 | 1221 | 1208 | 1255 |
| AGTTAATGGAGTAACTCCAG | KDM5B | 699 | 489 | 1055 | 983 | 1075 |
| GAAGCGCCTCGTTGGGACGA | KDM5B | 631 | 732 | 1484 | 1395 | 1508 |
| GCAGTGGCAACGATGAAGAC | KDM5B | 531 | 418 | 730 | 602 | 571 |
| GCTGTGGGCTCCCATATCAG | KDM5B | 571 | 807 | 1296 | 1318 | 1331 |
| TCACCTTCGCTTAGTCACAC | KDM5B | 608 | 225 | 436 | 475 | 512 |
| TCTTGCAAATCATCTCATCG | KDM5B | 567 | 607 | 934 | 965 | 899 |
| AGACACAGTATCTTCGGAGT | KDM5C | 549 | 511 | 1078 | 1172 | 1102 |
| CGGGCTCTGGGACTAGTCAG | KDM5C | 324 | 867 | 1481 | 1610 | 1647 |
| CTGCAGGTATCGATACACCT | KDM5C | 455 | 409 | 910 | 882 | 987 |
| GCTGGACAGACTGTCGAAGG | KDM5C | 358 | 481 | 954 | 873 | 924 |
| GTGCCCAAAGTGTGTCATGG | KDM5C | 656 | 500 | 1258 | 1103 | 1133 |
| TGACAGGAGTATGCTACCAG | KDM5C | 626 | 688 | 1284 | 1311 | 1266 |
| AAGAACATCTAGTCCAACAA | KDM6A | 382 | 295 | 488 | 516 | 540 |
| AGCCTTAATAGCCCTCACAG | KDM6A | 663 | 506 | 876 | 818 | 854 |
| AGGATTCGTAGAGTGTGCCT | KDM6A | 659 | 462 | 861 | 822 | 835 |
| CAGCATTATCTGCATACCAG | KDM6A | 398 | 329 | 614 | 668 | 647 |
| CTAGCAATTCAGTAACACAG | KDM6A | 482 | 437 | 596 | 530 | 560 |
| TGCCCACTTATATGAAACCC | KDM6A | 381 | 490 | 947 | 783 | 847 |
| AGCAGCAACACCGGTCTCCG | KDM6B | 695 | 785 | 1241 | 1147 | 1169 |
| CAAGCTTCCTCCGTATCGAA | KDM6B | 511 | 381 | 687 | 583 | 565 |
| CCGGCCCACCCGTTCCACGA | KDM6B | 526 | 521 | 936 | 933 | 914 |
| CTCAAACTGGTCCTCCAGTG | KDM6B | 616 | 786 | 1431 | 1360 | 1285 |
| GAAGTTTGGACCCCCTTCCG | KDM6B | 559 | 597 | 956 | 1119 | 1137 |
| GACAGAAGTACTGTTACCGG | KDM6B | 644 | 574 | 1086 | 1089 | 1109 |
| ACAGACAGCTCAACTCCGTG | KDR | 459 | 783 | 1444 | 1563 | 1496 |
| ACCGGAACCTCACCATCCGT | KDR | 699 | 350 | 754 | 679 | 584 |
| ACTTGTGCCATTAGAGAACG | KDR | 526 | 889 | 2049 | 1889 | 1955 |
| GACAGCAACTTGCAAGACAG | KDR | 516 | 731 | 1441 | 1219 | 1320 |
| GCTTCTGTTAGCGACCAACA | KDR | 383 | 534 | 952 | 971 | 936 |
| TAGTGGCATGGAATCTTCGG | KDR | 443 | 819 | 1288 | 1312 | 1465 |
| CACGCTGCAGGTCAAATACG | KEAP1 | 342 | 754 | 1399 | 1355 | 1397 |
| CCAGTCACTCAGCTACCTAG | KEAP1 | 609 | 574 | 692 | 619 | 666 |
| GAGTACATCTATATGCACTT | KEAP1 | 403 | 270 | 582 | 558 | 595 |
| GCTGTCGATCTGGTACATGA | KEAP1 | 499 | 384 | 800 | 630 | 643 |
| GGCCGGAACAACTCGCCAGA | KEAP1 | 399 | 654 | 1084 | 899 | 949 |
| TGACCCAGTTGATACAGGCG | KEAP1 | 483 | 687 | 1442 | 1246 | 1238 |
| AGCCTTCACTCGTCCCTGAG | KIAA1549 | 583 | 1064 | 1564 | 1603 | 1509 |
| CCGCTGGCTGAGGTATCTCG | KIAA1549 | 529 | 842 | 1262 | 1493 | 1454 |
| GAGGATTCACTTCAACCGTG | KIAA1549 | 343 | 792 | 1148 | 1146 | 1230 |
| GCTGCTCAGGAACTTGAGCG | KIAA1549 | 536 | 727 | 1298 | 1399 | 1416 |
| GGGTGACAACCCAGAATAGG | KIAA1549 | 340 | 862 | 1300 | 1329 | 1491 |
| TGTGTCGGTAACGAGCCACG | KIAA1549 | 512 | 874 | 1651 | 1703 | 1569 |
| AAATTGGACACACAACACTT | KIF11 | 449 | 385 | 494 | 513 | 527 |
| AACTGAGTTACCTTGTTACG | KIF11 | 335 | 750 | 1452 | 1362 | 1355 |
| AGTCTTGTGAACCTCTAGCA | KIF11 | 446 | 448 | 1061 | 852 | 896 |
| GATTCTCGATAAGGAACATG | KIF11 | 497 | 407 | 840 | 660 | 717 |
| TGAAAAAATTGGTGCCGTTG | KIF11 | 450 | 919 | 1715 | 1764 | 1694 |
| TTTCAGTATAGACACCACAG | KIF11 | 414 | 371 | 766 | 593 | 669 |
| ACAGAAATACCAATCTACTG | KIT | 482 | 369 | 770 | 711 | 710 |
| ATCACAAAGCCAATCAGCAA | KIT | 480 | 278 | 516 | 496 | 504 |
| CACAGACACAGAGTAATAGC | KIT | 495 | 338 | 548 | 479 | 437 |
| CATACAAGGGAAGGTCAACG | KIT | 627 | 870 | 1209 | 1199 | 1205 |
| TGGCCTGATTAAGTCGGATG | KIT | 748 | 1031 | 1379 | 1477 | 1396 |
| TGTTCATATAGATCCACTGC | KIT | 596 | 318 | 708 | 628 | 622 |
| ATATACTGGTGAACTCGGGG | KLF5 | 353 | 458 | 908 | 922 | 873 |
| GAACTGGGCATATCTAGATC | KLF5 | 703 | 370 | 857 | 843 | 818 |
| GAAGAACTGGTCTACGACTG | KLF5 | 566 | 510 | 1003 | 1011 | 1002 |
| GGCATTGTGTATGTGCAAGG | KLF5 | 475 | 573 | 1188 | 1228 | 1199 |
| GTCTTGATGTGTGTTACGCA | KLF5 | 457 | 534 | 1063 | 1031 | 1028 |
| TGTGTTAAGGCCCGCCATGG | KLF5 | 601 | 536 | 1324 | 1321 | 1286 |
| CTGTTGCCAGTACTCCTCCA | KLF6 | 619 | 548 | 1136 | 1163 | 1159 |
| GAGGATGCTCTGAACAGCCC | KLF6 | 562 | 2 | 12 | 13 | 19 |
| GGACCAAAATCATCCTGGCT | KLF6 | 494 | 357 | 793 | 874 | 874 |
| GGAGGACTCGCTGCTCACGT | KLF6 | 340 | 633 | 891 | 910 | 867 |
| TCGGAGGCTGACACATAGCA | KLF6 | 551 | 512 | 1047 | 924 | 962 |
| TGACTAAGACTTCGCCAATG | KLF6 | 380 | 780 | 1514 | 1493 | 1475 |
| AGGATAGCATGGACATTACG | KNL1 | 592 | 587 | 1263 | 1178 | 1174 |
| AGGGTAAATTCTGTATAGTG | KNL1 | 502 | 583 | 1189 | 1031 | 1007 |
| AGTCAAGCAACTGCTACGTG | KNL1 | 340 | 633 | 1308 | 1242 | 1260 |
| CAATAAGCTAACACTACTGA | KNL1 | 495 | 282 | 433 | 380 | 410 |
| GGAAACAAAGATTGCTCACA | KNL1 | 437 | 690 | 1277 | 1054 | 1200 |
| TAGTCTGTAAAAATGAGCTG | KNL1 | 512 | 901 | 1332 | 1218 | 1190 |
| ACATCAAGAACCCTGATTGG | KPNB1 | 474 | 628 | 1438 | 1165 | 1193 |
| CACACAGTGTCCGGATACAA | KPNB1 | 433 | 368 | 679 | 628 | 617 |
| CTGGTCCAATGTGTGTGATG | KPNB1 | 402 | 751 | 1574 | 1453 | 1469 |
| GCACATGAAAGAGTCGACAT | KPNB1 | 685 | 213 | 503 | 352 | 342 |
| TGCCAAGGATTGTTATCCCG | KPNB1 | 252 | 518 | 1256 | 1006 | 1121 |
| TTGTTCGAGATACAACTGCG | KPNB1 | 571 | 610 | 1244 | 1218 | 1246 |
| AAACTTGTGGTAGTTGGAGC | KRAS | 391 | 473 | 765 | 640 | 696 |
| AAGATATTCACCATTATAGG | KRAS | 551 | 401 | 619 | 551 | 572 |
| AATTACTACTTGTTTCCTGT | KRAS | 424 | 346 | 553 | 515 | 548 |
| AGAGGAGTACAGTGCAATGA | KRAS | 329 | 949 | 1518 | 1475 | 1418 |
| CTGAATTAGCTGTATCGTCA | KRAS | 361 | 527 | 673 | 695 | 777 |
| GATGTACCTATGGTCCTAGT | KRAS | 355 | 375 | 531 | 580 | 602 |
| AACCTGCCCATTAACATCGT | LAP3 | 624 | 469 | 883 | 802 | 843 |
| CTCCGCAGCCGACATGACGA | LAP3 | 479 | 648 | 1111 | 1283 | 1337 |
| GGGGTGCTTCACTTGCGTTG | LAP3 | 645 | 692 | 1302 | 1344 | 1241 |
| TCTGGAACTCTCTTCCGTGG | LAP3 | 600 | 967 | 1507 | 1571 | 1567 |
| TGAATAGGACTTCCCCAGTG | LAP3 | 313 | 795 | 1244 | 1263 | 1373 |
| TGCAAACATGGACCTCATGA | LAP3 | 408 | 537 | 914 | 889 | 888 |
| AAGGCGCAGTCCTTGACCAC | LCK | 588 | 450 | 833 | 851 | 908 |
| ACGTCAGAGCCATTCCGCAT | LCK | 565 | 391 | 797 | 754 | 810 |
| CCCAAGTCTCCATCGTGAGA | LCK | 504 | 773 | 1434 | 1338 | 1348 |
| CGGGACTTCGACCAGAACCA | LCK | 651 | 719 | 1300 | 1166 | 1117 |
| GGGTCTTGAGGAAATCCACC | LCK | 531 | 569 | 789 | 836 | 788 |
| GTGCCCGTTGTAGTACCCTG | LCK | 536 | 717 | 1318 | 1377 | 1427 |
| AAAAAACTGTATCTTACACC | LIFR | 628 | 207 | 353 | 271 | 303 |
| AACTAACAATTTGCGAGTAT | LIFR | 436 | 532 | 905 | 1018 | 1050 |
| ACACCTAAAGTGGAATGACA | LIFR | 545 | 342 | 606 | 686 | 810 |
| CCCTGATCCTCAACACAAGG | LIFR | 629 | 373 | 687 | 685 | 718 |
| GTTGATTCTGATCGACCCAG | LIFR | 492 | 546 | 1258 | 1098 | 1236 |
| TGGTCCCTTTGAAGGAACTA | LIFR | 430 | 660 | 1181 | 1533 | 1609 |
| AGGCCCACCTCAGGTAGTCG | LMO1 | 501 | 600 | 1135 | 1131 | 1168 |
| CCTGGACAAGTACTGGCACG | LMO1 | 517 | 761 | 1435 | 1510 | 1403 |
| GCTGATCCCAGCCTTCGAGA | LMO1 | 509 | 680 | 972 | 958 | 1003 |
| GGGCTCCACCCTCTACACCA | LMO1 | 689 | 410 | 576 | 571 | 634 |
| TGCTCTCCGTCCAGCCCAAG | LMO1 | 541 | 611 | 1441 | 1257 | 1254 |
| TGCTGGACAAGGAGGACGGT | LMO1 | 446 | 392 | 932 | 845 | 825 |
| AGGGATGAGCTTACTACAGG | LMO3 | 661 | 563 | 1197 | 1019 | 1036 |
| ATTAGCTTTAGTGTACAGGG | LMO3 | 341 | 579 | 1161 | 1069 | 1028 |
| CAGCCAGACACCAAGCCGAA | LMO3 | 506 | 571 | 770 | 877 | 831 |
| TCCCAAACGACAGTCACAGC | LMO3 | 532 | 389 | 580 | 601 | 629 |
| TGGCTGCAACCGAAAGATCA | LMO3 | 480 | 729 | 1141 | 1117 | 1184 |
| TTTGTCGCAGAGACTATCTG | LMO3 | 645 | 546 | 1186 | 1150 | 1114 |
| AAGAACCGTATGTCATGTCA | LRP1B | 323 | 654 | 1118 | 1334 | 1323 |
| ACTAGATTTGATAACGAACG | LRP1B | 470 | 918 | 1574 | 1862 | 1798 |
| ATGCAACGACGATTGTAGCA | LRP1B | 560 | 587 | 1103 | 1080 | 1059 |
| CACAACCAGCACTTACCCTG | LRP1B | 472 | 695 | 1100 | 962 | 977 |
| CAGGAAGACTAATAGATGAG | LRP1B | 396 | 826 | 1349 | 1490 | 1518 |
| GCTTATTACGGATAACTGGG | LRP1B | 567 | 600 | 1086 | 959 | 993 |
| AGGAAGACAGTAATCCATGG | LRP2 | 567 | 914 | 1496 | 1410 | 1457 |
| ATACAACGTCCAGACGTACA | LRP2 | 602 | 482 | 1009 | 1046 | 1015 |
| GAAGGCATTGTCACACCACG | LRP2 | 800 | 817 | 1314 | 1332 | 1428 |
| GATGAACTACAATTGATCGG | LRP2 | 521 | 511 | 1038 | 984 | 1006 |
| TCAGGACAACGAAATTGACT | LRP2 | 579 | 357 | 558 | 566 | 595 |
| TTGTATTGGATCCTTGCCGA | LRP2 | 616 | 825 | 1166 | 1199 | 1145 |
| AAAGTAAACCCAAATATGTG | LRP6 | 594 | 661 | 1104 | 1089 | 1147 |
| CAAGGCTCAATGGTACCATG | LRP6 | 589 | 769 | 1343 | 1281 | 1295 |
| CCACCATTATCAATTCCACA | LRP6 | 664 | 480 | 674 | 552 | 596 |
| CCCGGACATTCCGAAGACTG | LRP6 | 534 | 490 | 1106 | 1058 | 975 |
| CTTTCCACACCAAAACCTGT | LRP6 | 462 | 329 | 602 | 556 | 524 |
| GCCGAGAGGAGTGAAAGACG | LRP6 | 492 | 787 | 1514 | 1429 | 1378 |
| AAATCCAATTCAATTAGTGT | LRRK2 | 510 | 427 | 587 | 471 | 483 |
| AAGGAACTCTTGAACAAGCG | LRRK2 | 448 | 605 | 930 | 903 | 928 |
| AGCGGACGCCAATCGAACAA | LRRK2 | 374 | 784 | 1393 | 1376 | 1386 |
| CAGATGTACCCAGATGACCA | LRRK2 | 580 | 806 | 1337 | 1383 | 1316 |
| TAACATTCTGGTATTAAACG | LRRK2 | 432 | 605 | 810 | 829 | 760 |
| TGTCGTTTCGAGAGACATCA | LRRK2 | 429 | 537 | 984 | 860 | 920 |
| AATGTCAATGCCCCACCTAG | LTK | 536 | 438 | 849 | 779 | 795 |
| ACTGGAGCAGAATTAGGCGA | LTK | 497 | 856 | 1514 | 1398 | 1469 |
| AGCAAAATCACTTCATCCGA | LTK | 620 | 567 | 843 | 695 | 759 |
| AGCCTTAGAACAGGCCCGAA | LTK | 393 | 487 | 949 | 929 | 906 |
| TGCAACAAGGCAGACCCAGG | LTK | 500 | 435 | 1019 | 966 | 1063 |
| TGTGTGTGAGATACCTCTGC | LTK | 339 | 503 | 1080 | 838 | 934 |
| AGAGGAGCAAGGAGACATCG | LYN | 485 | 833 | 1541 | 1380 | 1344 |
| CCAACTTGATGGATTCCCGT | LYN | 518 | 463 | 817 | 706 | 767 |
| CTATAACAATAGTACCAAAG | LYN | 579 | 284 | 555 | 523 | 470 |
| TAACCAGGAAAGATGCAGAA | LYN | 624 | 402 | 788 | 749 | 813 |
| TAATAACATCACCATGCACA | LYN | 548 | 440 | 670 | 662 | 659 |
| TTAATCATGTCGCTGATACA | LYN | 485 | 983 | 1219 | 1328 | 1311 |
| AAAAGCTTTGATAGAGCAAG | LYPLA1 | 497 | 800 | 1288 | 1160 | 1199 |
| AAGCCCAAAAATATCAAACC | LYPLA1 | 547 | 150 | 277 | 251 | 286 |
| ACCACACAGCAGAAACTGGC | LYPLA1 | 556 | 439 | 690 | 671 | 695 |
| AGTACCAGTCTGGCTACCTG | LYPLA1 | 593 | 760 | 1544 | 1403 | 1460 |
| ATAAAGATAAAGCTCCTCCC | LYPLA1 | 639 | 547 | 782 | 760 | 783 |
| GCTTTCACCAGATTCACAGG | LYPLA1 | 496 | 387 | 868 | 806 | 758 |
| AGTGTCATTGGGGTAACCAT | MAGI1 | 453 | 611 | 1127 | 1005 | 1050 |
| AGTTCATTCACACAAAGCTG | MAGI1 | 382 | 695 | 1474 | 1535 | 1416 |
| CATAGTAGATACCATAGACA | MAGI1 | 500 | 498 | 855 | 921 | 908 |
| GCCACGTCAACCTCACAGTG | MAGI1 | 417 | 652 | 1420 | 1523 | 1541 |
| GCGACGATGCTACTATTCAC | MAGI1 | 371 | 649 | 978 | 920 | 896 |
| GTGACATTGTTGGTGCAACG | MAGI1 | 473 | 749 | 1215 | 1262 | 1265 |
| AAAAACAAACCACCTCCAGG | MAGI3 | 499 | 505 | 863 | 844 | 897 |
| AATGGACAGTCATTAACCAA | MAGI3 | 367 | 501 | 844 | 887 | 889 |
| ACAAAGACGAGGATCCAACC | MAGI3 | 660 | 359 | 726 | 683 | 615 |
| AGTTTGACACAGAATCCCAA | MAGI3 | 548 | 351 | 504 | 417 | 460 |
| GCCATGCATCACAGACCGAT | MAGI3 | 490 | 6 | 16 | 23 | 24 |
| GTACCGTTGCTTATCTTGAG | MAGI3 | 578 | 791 | 1238 | 1176 | 1242 |
| ACTGTCACGTTAAGGCGTTG | MALT1 | 579 | 1015 | 2036 | 1701 | 1775 |
| CACCAAAGGAGCTTTAAGCT | MALT1 | 479 | 629 | 1031 | 943 | 908 |
| CAGGATTATTATATTATGCG | MALT1 | 558 | 780 | 1331 | 1275 | 1301 |
| GTTTGTGAAACTATGTTGCC | MALT1 | 687 | 358 | 620 | 663 | 683 |
| TGCAAATCTGTGTCGAACCA | MALT1 | 597 | 921 | 1395 | 1413 | 1452 |
| TTTACACCCGGCAAGTACCG | MALT1 | 570 | 546 | 1040 | 1024 | 1023 |
| AGCTCCAGGTCACTCATGGG | MAML2 | 469 | 805 | 1568 | 1485 | 1541 |
| CCCTGCTCAAATCCAAACAC | MAML2 | 569 | 287 | 391 | 360 | 390 |
| GATGGCCAGCTACATGTACA | MAML2 | 609 | 698 | 1188 | 1112 | 1174 |
| GCCGACGGGGGCCTCAACTG | MAML2 | 413 | 565 | 1050 | 1111 | 996 |
| TATCAAAAGAATCCGCGTTG | MAML2 | 499 | 1012 | 1350 | 1282 | 1258 |
| TGGGCTGAAACGGAGCTCGG | MAML2 | 436 | 648 | 1221 | 1437 | 1328 |
| ACATCCTCGTGAACTCCCGT | MAP2K1 | 453 | 461 | 806 | 771 | 743 |
| ATCTCCAGGTGAATTAGCTG | MAP2K1 | 479 | 687 | 1300 | 1264 | 1357 |
| CGCTGCTCATACCGACATGT | MAP2K1 | 518 | 564 | 954 | 1085 | 1061 |
| GCAGCAGCGGAAGCGCCTTG | MAP2K1 | 551 | 918 | 1628 | 1703 | 1742 |
| GGTGAACGGGACCAGCTCGG | MAP2K1 | 430 | 664 | 1378 | 1373 | 1399 |
| TGCAACTCCCCGTACATCGT | MAP2K1 | 264 | 455 | 840 | 780 | 709 |
| AAGCACCAGATCATGCACCG | MAP2K2 | 480 | 340 | 772 | 749 | 704 |
| CTGTGCTGACCCTAGGATGG | MAP2K2 | 617 | 1125 | 1808 | 1769 | 1798 |
| GTAGGACCGAGTTCCCACGA | MAP2K2 | 493 | 512 | 977 | 893 | 910 |
| GTGTAACTCCCCATACATCG | MAP2K2 | 770 | 713 | 1326 | 1413 | 1402 |
| TCAAGGATGACGACTTCGAA | MAP2K2 | 381 | 518 | 1128 | 1070 | 1018 |
| TCACCTTTCTCGCCATGATG | MAP2K2 | 470 | 588 | 887 | 819 | 832 |
| AACTTGAAAATCATTCACAG | MAP2K4 | 433 | 413 | 675 | 617 | 582 |
| ACCAAGTGGGCAAATAATGG | MAP2K4 | 532 | 701 | 1394 | 1533 | 1591 |
| ACCAGGAATACTCACATGTG | MAP2K4 | 600 | 1095 | 1532 | 1588 | 1672 |
| AGTGTGCCAAACATACCAGT | MAP2K4 | 267 | 370 | 692 | 639 | 596 |
| GCAGCACTGGGATTTCACTG | MAP2K4 | 447 | 704 | 1176 | 1094 | 1198 |
| TTATGGTGCACTCTTCAGAG | MAP2K4 | 393 | 773 | 1362 | 1520 | 1431 |
| AATGATTAGGGAGAAACTGA | MAP3K1 | 452 | 432 | 800 | 815 | 840 |
| ACGTTAACTTACAGGCACCG | MAP3K1 | 542 | 622 | 1043 | 1116 | 1098 |
| CAACGTACACTTTGTAGACA | MAP3K1 | 540 | 592 | 1031 | 1216 | 1201 |
| CTTTGGCTGGATCACAACGA | MAP3K1 | 751 | 558 | 1036 | 904 | 918 |
| GCGTGAAACAAACTTCTGGA | MAP3K1 | 471 | 515 | 819 | 666 | 694 |
| GTCCGAGGGCTCCGGCGACG | MAP3K1 | 559 | 641 | 1144 | 1117 | 1177 |
| AAAGCTCGTAATTTATACAC | MAP3K5 | 582 | 207 | 401 | 413 | 412 |
| AACCGTTTACTATTACCATG | MAP3K5 | 635 | 755 | 1035 | 975 | 1033 |
| CTTTGTAGATTCGACCAACT | MAP3K5 | 673 | 388 | 719 | 721 | 615 |
| GCACCAGAGATCATAGACAA | MAP3K5 | 376 | 534 | 802 | 953 | 858 |
| GTTGATCACATAAGCCACCG | MAP3K5 | 359 | 643 | 941 | 871 | 938 |
| TCTAGAAAGGTATACATGAG | MAP3K5 | 542 | 990 | 1539 | 1399 | 1293 |
| AATATTAGGATGGTTCACAC | MAP3K7 | 437 | 308 | 580 | 553 | 590 |
| CACACATGACCAATAACAAG | MAP3K7 | 636 | 259 | 406 | 441 | 381 |
| CGATGTCCATGAATGAGCCT | MAP3K7 | 597 | 555 | 891 | 810 | 769 |
| CTGCTGCCCATGCAATGAGT | MAP3K7 | 448 | 640 | 1114 | 1038 | 1061 |
| TTCTCAGCGCCCTTCAATGG | MAP3K7 | 446 | 770 | 1590 | 1525 | 1559 |
| TTTGGGAAGTGATAACACGT | MAP3K7 | 554 | 453 | 1113 | 1026 | 979 |
| ATCGGTCAGATACGGAACCG | MAP3K8 | 595 | 863 | 1518 | 1584 | 1562 |
| CTCTGATTTCATACCTCGGG | MAP3K8 | 427 | 608 | 1136 | 1161 | 1140 |
| TAGGTAATCACTCCCCAGAA | MAP3K8 | 482 | 396 | 900 | 873 | 916 |
| TCAAGACAATAATCAAAATG | MAP3K8 | 783 | 375 | 683 | 559 | 547 |
| TCAATTTAAGCCATCCGACG | MAP3K8 | 558 | 598 | 1126 | 1152 | 1183 |
| TTAGATTTATATGAGCCCAG | MAP3K8 | 459 | 539 | 806 | 899 | 844 |
| AAGCCATCGATTCCTCGGAA | MAP3K9 | 473 | 449 | 987 | 951 | 1008 |
| GAGCTCAGTCATCCCGAAAG | MAP3K9 | 382 | 477 | 1065 | 1027 | 1053 |
| GTGAAGATTAGACTCACCCC | MAP3K9 | 463 | 403 | 622 | 743 | 777 |
| GTTGGTTGGCGTCAGCTGAG | MAP3K9 | 523 | 760 | 1239 | 1189 | 1226 |
| TATAGAGAATGTTCGCCAGG | MAP3K9 | 663 | 652 | 1182 | 1149 | 1142 |
| TGTTACTGGACTTAAGGTCG | MAP3K9 | 614 | 851 | 1506 | 1660 | 1619 |
| ATATATAGTACAAGACCTCA | MAPK1 | 646 | 308 | 600 | 532 | 566 |
| CGGAGTATGTAGCCACACGT | MAPK1 | 519 | 571 | 984 | 974 | 963 |
| CTGGCTGACCTTGAGATCGC | MAPK1 | 525 | 534 | 836 | 815 | 824 |
| GGTCGAGATAATGCTTCCCC | MAPK1 | 568 | 366 | 738 | 575 | 607 |
| GGTGCAGTACATTAGCTGAA | MAPK1 | 634 | 696 | 1023 | 951 | 926 |
| TAGGAGAGGTTGGTGTAGCG | MAPK1 | 575 | 558 | 1405 | 1362 | 1316 |
| AACAAGGTAATTGAACAACT | MAPK10 | 577 | 356 | 529 | 533 | 561 |
| ACGTTATTATAGAGCCCCTG | MAPK10 | 586 | 930 | 1515 | 1545 | 1494 |
| CACACCCCAGAAAACACTGG | MAPK10 | 462 | 719 | 1059 | 996 | 1031 |
| GCAGGTAAGACATTCGCTCA | MAPK10 | 715 | 556 | 1473 | 1450 | 1404 |
| TCAGGGATTCGATAAACAAG | MAPK10 | 343 | 563 | 944 | 1047 | 997 |
| TTTAAAGCCTATTGGCTCTG | MAPK10 | 387 | 841 | 1715 | 1573 | 1620 |
| AAAGGACCTACAGAGAACTG | MAPK14 | 402 | 716 | 1287 | 1210 | 1288 |
| ATAATGGCCGAACTGTTGAC | MAPK14 | 483 | 550 | 920 | 1000 | 980 |
| CACTCACCACACGGAGCCGT | MAPK14 | 498 | 397 | 758 | 848 | 823 |
| CAGGCTATGTGGCTACCAGG | MAPK14 | 574 | 740 | 1221 | 1220 | 1204 |
| GGTAGATAAGGAACTGAACA | MAPK14 | 540 | 397 | 843 | 850 | 909 |
| TTCAGCTGACATAATTCACA | MAPK14 | 573 | 484 | 813 | 701 | 698 |
| AGCTTACGACCACGTGCGCA | MAPK3 | 424 | 685 | 1374 | 1424 | 1403 |
| CCTACTGCCAGCGCACACTG | MAPK3 | 564 | 458 | 864 | 928 | 931 |
| GGATATACTTGAGACCTCGC | MAPK3 | 454 | 346 | 477 | 406 | 456 |
| TCTGACCTTAAGGTCACAGG | MAPK3 | 497 | 593 | 951 | 1090 | 1077 |
| TGAAGGATACACATCCCTCA | MAPK3 | 472 | 484 | 674 | 675 | 764 |
| TGGAGAGCCTAGGATACCTG | MAPK3 | 357 | 631 | 1262 | 1188 | 1283 |
| AGAAACTACAGCCAACAGTA | MAPK8 | 465 | 442 | 858 | 1012 | 910 |
| TAAGGATACGATCTGTACCT | MAPK8 | 767 | 577 | 1081 | 867 | 973 |
| TCCATGAAGTTACATAGTCA | MAPK8 | 499 | 763 | 1266 | 1302 | 1403 |
| TCGCTACTACAGAGCACCTG | MAPK8 | 379 | 644 | 1166 | 1031 | 1015 |
| TTCAAAAGGCCAATTATCTG | MAPK8 | 352 | 678 | 1103 | 1197 | 1134 |
| TTGCAATCAAGAAACTAAGC | MAPK8 | 435 | 245 | 343 | 328 | 309 |
| AATGGATGCTAACTTATGTC | MAPK9 | 575 | 685 | 1019 | 962 | 1050 |
| ACTTCTTAGATATTGATCAG | MAPK9 | 567 | 775 | 1476 | 1506 | 1487 |
| AGAAACTCCAGCCAACCGTG | MAPK9 | 611 | 1044 | 1400 | 1520 | 1435 |
| CTTCATGATGACACCGTACG | MAPK9 | 670 | 853 | 1301 | 1229 | 1185 |
| GTTGCTAGGTTTCAAATCCT | MAPK9 | 348 | 765 | 1127 | 1148 | 1181 |
| TATCAGCAGTTGAAACCAAT | MAPK9 | 555 | 393 | 517 | 401 | 438 |
| AATACCAAAGTCCCAACTTG | MAPKAP1 | 395 | 683 | 1185 | 1235 | 1330 |
| CCCAATCAAGTCCTGCACCC | MAPKAP1 | 678 | 373 | 751 | 722 | 689 |
| CTCAGCAATATGCAGGCAGT | MAPKAP1 | 399 | 734 | 1249 | 1135 | 1102 |
| CTGTCTCCAGGCATTGAAGG | MAPKAP1 | 351 | 775 | 1292 | 1128 | 1212 |
| GCTTCTCTAGGCGGTACTGA | MAPKAP1 | 421 | 1027 | 1511 | 1480 | 1480 |
| GTTGTGCCTACGTGACCCTG | MAPKAP1 | 441 | 813 | 1217 | 1109 | 1101 |
| AGAGACCCTCCGGCGAGTCG | MCL1 | 749 | 428 | 871 | 661 | 686 |
| GCCAAGGACGCGAAACCACT | MCL1 | 359 | 401 | 705 | 675 | 699 |
| GCTTCCGCCAATCACCGCAC | MCL1 | 593 | 320 | 503 | 440 | 476 |
| GGTACGAGCCGGAACCTTTG | MCL1 | 596 | 599 | 1104 | 1086 | 1103 |
| GTACCGAGAGATAATCTCCA | MCL1 | 478 | 385 | 561 | 408 | 380 |
| TTGGGGCCCTCAGCGCCAAT | MCL1 | 442 | 944 | 1170 | 1141 | 1102 |
| AACAGGGACGCCATCAAATC | MDM2 | 506 | 459 | 943 | 845 | 874 |
| AGACACTTATACTATGAAAG | MDM2 | 473 | 537 | 860 | 687 | 679 |
| AGGATTCATTTCATTGCATG | MDM2 | 325 | 770 | 1500 | 1637 | 1594 |
| GAAAATAGCTGTCACCGTGA | MDM2 | 510 | 451 | 767 | 568 | 602 |
| GAGTAACTGTGTATCAGGCA | MDM2 | 556 | 557 | 910 | 752 | 786 |
| GGGAAATACTATCAGATTTG | MDM2 | 461 | 707 | 1133 | 976 | 1113 |
| AATTTAACTCTCCAAGCAAG | MDM4 | 480 | 288 | 498 | 459 | 505 |
| CAAAAACCACAAGTCATCTG | MDM4 | 582 | 330 | 693 | 612 | 603 |
| CCTTGCAGGTGATTGAAGTG | MDM4 | 480 | 701 | 1368 | 1141 | 1134 |
| GAACAGCATATGGTGTACTG | MDM4 | 532 | 445 | 963 | 746 | 793 |
| GAGACATCTAAGCTGGACCT | MDM4 | 438 | 543 | 995 | 980 | 964 |
| TCCAAGTCAAGACCAACTGA | MDM4 | 485 | 367 | 877 | 785 | 817 |
| AGAACGGCAATATCGCTGCG | MECOM | 566 | 635 | 1073 | 1169 | 1126 |
| CAGCCCCTCGATCTAAGCAT | MECOM | 563 | 427 | 981 | 807 | 873 |
| GATGATGTCAGTACGCCAAG | MECOM | 572 | 710 | 1159 | 1191 | 1112 |
| GATGCTAGTCAACCAGATGT | MECOM | 332 | 533 | 898 | 1016 | 1021 |
| GCATGCGCTTATGACGACAA | MECOM | 393 | 529 | 922 | 887 | 847 |
| GGCATGACTCATATTAACCA | MECOM | 476 | 317 | 528 | 539 | 495 |
| ACTTGCACCCTCATCTCACG | MED12 | 385 | 635 | 961 | 1039 | 1003 |
| CAGACACGAGTTCCTGACCT | MED12 | 366 | 462 | 1011 | 861 | 1002 |
| CTAGGGCACTAGGACAACAC | MED12 | 551 | 352 | 597 | 568 | 655 |
| CTGTTGGAAAACCTCGATTG | MED12 | 493 | 723 | 1515 | 1403 | 1448 |
| GTGGCATACTGCACATGTCG | MED12 | 328 | 550 | 1179 | 1186 | 1156 |
| TGTGTGGCGTAGTGAAGCAT | MED12 | 569 | 575 | 1076 | 1002 | 1029 |
| ACTGCATTGATATTTACTGC | MED14 | 580 | 486 | 870 | 713 | 756 |
| AGACACTCAGAATTACCGCA | MED14 | 491 | 383 | 1010 | 681 | 697 |
| AGGTAGAAATAACCGCACCT | MED14 | 483 | 319 | 748 | 657 | 688 |
| ATACTTACAATGTAGTACTG | MED14 | 509 | 457 | 1000 | 816 | 890 |
| ATGTTAATACGAGAGCGGTG | MED14 | 478 | 618 | 1161 | 1100 | 1103 |
| CTTCAACTCGAAACTTCACC | MED14 | 605 | 307 | 471 | 404 | 384 |
| AAGTCACTTACATAGAGCCG | MEF2B | 436 | 538 | 1006 | 902 | 945 |
| CACCAACACCGACATCCTCG | MEF2B | 522 | 501 | 988 | 910 | 1021 |
| CCGTGTACTTCAGGAGCACG | MEF2B | 449 | 627 | 1408 | 1473 | 1479 |
| GACTCTGTAGGCTACCATAG | MEF2B | 457 | 682 | 1258 | 1238 | 1305 |
| GGGCCTCCCCAAGCCCACTG | MEF2B | 543 | 488 | 993 | 893 | 911 |
| GGTGGCTCGGTGAAAAGAGA | MEF2B | 608 | 864 | 1301 | 1365 | 1361 |
| ACCCTTTCGTAGGTACCCCA | MEN1 | 500 | 587 | 896 | 899 | 872 |
| CAGGGGCCAGACAGTCAACG | MEN1 | 347 | 785 | 1265 | 1126 | 1091 |
| GAACATTGCGATTGCGACAG | MEN1 | 450 | 799 | 1365 | 1352 | 1442 |
| GAGGCTGTTCCAGATGACAT | MEN1 | 452 | 556 | 1137 | 1029 | 1055 |
| TGATGGATAGGTCCGCCACA | MEN1 | 391 | 758 | 1350 | 1225 | 1255 |
| TGTCATCCCCAACTTGCTGA | MEN1 | 369 | 763 | 1168 | 1068 | 1049 |
| ACTGGAAAATACCTCAACAG | MET | 643 | 389 | 570 | 625 | 600 |
| AGGCTAAAGGAAACGCAAGA | MET | 448 | 482 | 715 | 757 | 797 |
| CCAGGTGCACACATCTATCG | MET | 460 | 723 | 1413 | 1445 | 1395 |
| GATGACCAACTCATTAGCTG | MET | 414 | 593 | 997 | 831 | 949 |
| TACCTGCATGAAGCGACCCT | MET | 367 | 679 | 1296 | 1192 | 1226 |
| TCCGTGAGAATACACTCCAG | MET | 335 | 685 | 943 | 934 | 845 |
| ATGAGATAAGGCTGCACGAC | MGMT | 618 | 383 | 433 | 510 | 472 |
| CAGCTTCCGCAACACCCGCG | MGMT | 370 | 487 | 970 | 1063 | 1075 |
| CAGGGCTGCTAACTGCTGGT | MGMT | 363 | 617 | 1103 | 1160 | 1128 |
| GAAAATGGGATGGTGAATCG | MGMT | 541 | 522 | 1185 | 994 | 1018 |
| GAGATCTCGATCTTCCCCAA | MGMT | 393 | 492 | 893 | 848 | 830 |
| GCTCCTCATCGCTCCGCCCA | MGMT | 397 | 554 | 1059 | 1115 | 1046 |
| AAGCAGGATCCATCAATCCC | MITF | 451 | 616 | 1115 | 1075 | 1045 |
| ACCACATACAGCAAGCCCAA | MITF | 350 | 320 | 474 | 454 | 531 |
| ACTCACGGATCATTTGACTT | MITF | 425 | 734 | 1217 | 1192 | 1252 |
| CAACCTTCCCAACATAAAAA | MITF | 472 | 216 | 410 | 426 | 451 |
| GGTGGCATGACATGATCGCC | MITF | 415 | 722 | 838 | 829 | 782 |
| TTCACGAGCATCGTGCATGC | MITF | 380 | 625 | 953 | 898 | 916 |
| ACTGCCCAATGCCACAACGG | MLH1 | 508 | 652 | 1296 | 1388 | 1418 |
| AGAGGTGGTTAAATCCACAA | MLH1 | 659 | 516 | 942 | 877 | 903 |
| CTTGGATCTGAATCAACTTG | MLH1 | 471 | 983 | 1604 | 1620 | 1553 |
| GACAGGACCGATGTTTCCAG | MLH1 | 518 | 352 | 846 | 747 | 751 |
| TAATAGTAACATGCGCCACA | MLH1 | 461 | 516 | 737 | 704 | 745 |
| TCTACATCAGAGCTTTCCCG | MLH1 | 610 | 521 | 817 | 746 | 860 |
| AGAACTGGATCTGTCGGAGC | MLLT11 | 526 | 398 | 637 | 585 | 622 |
| AGCTGTACTGGCTACTCACA | MLLT11 | 528 | 937 | 1385 | 1254 | 1336 |
| GACAGATCCAGTTCTGGAAT | MLLT11 | 504 | 609 | 1288 | 1191 | 1189 |
| GCAAGCAACTGGAGTAGAAC | MLLT11 | 518 | 323 | 634 | 597 | 657 |
| GCAGCACTGGCAAAATGACC | MLLT11 | 598 | 268 | 627 | 682 | 650 |
| GGGCATCCTCCAGAAAAGAA | MLLT11 | 440 | 365 | 551 | 503 | 493 |
| AACAAGAATATCGCGTCCGT | MLST8 | 520 | 479 | 790 | 670 | 772 |
| AATTAATGGGTGCATTCACC | MLST8 | 480 | 449 | 779 | 809 | 881 |
| CGGGCTACGACCACACGGTG | MLST8 | 447 | 661 | 1446 | 1456 | 1414 |
| CTATGTCTGGAATCTGACTG | MLST8 | 514 | 736 | 1536 | 1315 | 1480 |
| GCAGACGTGATAGAGACCTC | MLST8 | 384 | 564 | 1123 | 1154 | 1131 |
| TGCGAATGTGCTGATAACCT | MLST8 | 555 | 438 | 714 | 764 | 685 |
| ACCTGTAGGTCAGATGCGTG | MMP1 | 459 | 648 | 1254 | 1469 | 1395 |
| AGATTACAACTTGTATCGCG | MMP1 | 555 | 550 | 840 | 768 | 811 |
| GAGCTTACCTCCCCAGACAA | MMP1 | 663 | 635 | 959 | 874 | 940 |
| GATGCTATCACTACAATTCG | MMP1 | 478 | 702 | 1328 | 1308 | 1304 |
| GGTGACTGGAAAGGTCGACA | MMP1 | 529 | 559 | 1191 | 1045 | 1017 |
| TGATGTCGTCCTGAGATAGC | MMP1 | 536 | 438 | 651 | 616 | 654 |
| AAGTCTCGTCATAGTCGAAG | MMP11 | 437 | 672 | 1132 | 1166 | 1161 |
| ACGAGTTTGGCCATACGCTG | MMP11 | 412 | 673 | 1310 | 1097 | 1176 |
| AGCACCTCTATGGCCAACCC | MMP11 | 458 | 469 | 893 | 833 | 903 |
| AGCACGACCCTCATGCACCT | MMP11 | 377 | 676 | 851 | 926 | 887 |
| GCTCCCGTACCAGCTGCCAT | MMP11 | 676 | 562 | 969 | 945 | 962 |
| TAACGAGATTGCCCCACTGG | MMP11 | 628 | 832 | 1315 | 1203 | 1201 |
| CCATCAACACTGCCTACGAG | MMP14 | 360 | 495 | 883 | 734 | 768 |
| CCCAGGCCCCAACATTGGAG | MMP14 | 434 | 690 | 1163 | 1086 | 1135 |
| CTGCCTGATGACGACCGCCG | MMP14 | 502 | 757 | 1215 | 1285 | 1408 |
| CTTGCGAATGGCCTCGAATG | MMP14 | 624 | 931 | 1435 | 1571 | 1546 |
| GGGGGCGCCTCATAGCCCTG | MMP14 | 619 | 926 | 1351 | 1332 | 1403 |
| GTGAGTGCGCAGATCCCCCG | MMP14 | 517 | 430 | 1027 | 949 | 960 |
| ACACGTATCGGGTTCCAAAG | MPL | 595 | 675 | 1080 | 937 | 995 |
| CCCATGATGCTCCAACAGGA | MPL | 509 | 513 | 743 | 710 | 693 |
| GCTGCGCAGCCAACCCGATG | MPL | 545 | 599 | 1154 | 1122 | 1139 |
| GGAACTTCAGATCAGCTGGG | MPL | 580 | 703 | 1470 | 1528 | 1548 |
| GTCCACATAGAGTACCCGCT | MPL | 627 | 369 | 633 | 621 | 604 |
| GTTCGAGAGAAACAGTTCAG | MPL | 499 | 634 | 1115 | 1134 | 1114 |
| AAATGATACGTATGTAACAC | MRE11 | 504 | 274 | 583 | 492 | 506 |
| AACCTCCACTATAGTCCACC | MRE11 | 541 | 467 | 700 | 633 | 618 |
| AGATAATCCCAAAGTAACCC | MRE11 | 631 | 286 | 483 | 413 | 457 |
| CCATGGGTGAACTACCAAGA | MRE11 | 422 | 520 | 950 | 1039 | 1040 |
| CTCTCACTTCTAACAGAAAG | MRE11 | 527 | 570 | 882 | 783 | 764 |
| GTTTGCTGCGTATCAAAGGA | MRE11 | 432 | 526 | 1003 | 953 | 1008 |
| CATCATTTCTGGATCACTCC | MS4A1 | 663 | 414 | 639 | 687 | 687 |
| CCATCACAGCAAAAACGCCC | MS4A1 | 583 | 374 | 524 | 412 | 413 |
| GGAGGCTGCCTAGGGCAATG | MS4A1 | 555 | 957 | 1569 | 1618 | 1536 |
| GTACCACATAGTTATACAGA | MS4A1 | 557 | 562 | 832 | 681 | 716 |
| TCCCAAAAGGATGCCTTCAG | MS4A1 | 465 | 570 | 1155 | 1049 | 1111 |
| TGTATGTCAACATATGGCAT | MS4A1 | 526 | 728 | 1103 | 1137 | 1202 |
| AAATAGGTTATTCAGAGAGG | MSH2 | 543 | 770 | 1340 | 1191 | 1177 |
| AACAACTCTAGATATGGATC | MSH2 | 493 | 636 | 1192 | 1018 | 976 |
| AACACCCACGACACCAATGG | MSH2 | 467 | 514 | 1029 | 949 | 906 |
| CAAAAATAAAACCGAGTACG | MSH2 | 545 | 851 | 1471 | 1461 | 1439 |
| CGAGAGGTGTTCAAGACGCA | MSH2 | 536 | 729 | 1434 | 1478 | 1592 |
| GATTTATACCCTGATAGAGT | MSH2 | 512 | 352 | 582 | 611 | 563 |
| AATGCCAGTTGATCGTTTGG | MSH6 | 676 | 908 | 1408 | 1463 | 1463 |
| ATCCACACTCTATGTGCCTG | MSH6 | 401 | 851 | 1539 | 1366 | 1412 |
| ATCTGTAGAGTCATTACCAA | MSH6 | 481 | 416 | 712 | 700 | 627 |
| GACGATCGTTGATAGCGTAA | MSH6 | 415 | 664 | 1123 | 1115 | 1167 |
| GGTCATACCTTTAAGCACCT | MSH6 | 415 | 419 | 678 | 626 | 609 |
| TCAAAGGAAGTACAAAAGGG | MSH6 | 380 | 475 | 1050 | 1101 | 1080 |
| ACTGGAAGAACAGACCCGCA | MSN | 688 | 470 | 869 | 873 | 828 |
| AGAGCAGCATAAACTCAACA | MSN | 564 | 427 | 714 | 747 | 735 |
| ATACTTAGACTGGACAGCAT | MSN | 559 | 520 | 812 | 750 | 806 |
| CCGCAAGCCTGACACCATTG | MSN | 526 | 730 | 1480 | 1550 | 1586 |
| CTTATTGAGTTTCAGCCAAG | MSN | 646 | 800 | 1101 | 1005 | 971 |
| TACAGACTGACTCCCAAGAT | MSN | 439 | 583 | 1108 | 1141 | 1178 |
| AAGAACATCTTGGCGCTGAG | MTHFR | 431 | 230 | 652 | 601 | 574 |
| ACATCTACCGTACACAAGAG | MTHFR | 514 | 474 | 916 | 1004 | 1043 |
| CAGGCCACAGTAGTTCACGG | MTHFR | 497 | 577 | 1173 | 1169 | 1059 |
| CCTCAAAGCTCCCTGCGTCG | MTHFR | 514 | 667 | 1203 | 1088 | 1067 |
| GAGCGGTCAACCTCATCTCG | MTHFR | 442 | 651 | 1230 | 1148 | 1200 |
| TAGTTCGAGATGTTCCACGC | MTHFR | 511 | 614 | 846 | 832 | 878 |
| AATCGGATGAATGATACGGG | MTOR | 492 | 576 | 1048 | 1053 | 963 |
| GATCATCCGCCAGATCTGGT | MTOR | 598 | 534 | 845 | 835 | 787 |
| GCGGGGTAGAACTCGTCCAG | MTOR | 499 | 534 | 1038 | 997 | 966 |
| GTGCTCACCCGATACCACTG | MTOR | 489 | 501 | 1061 | 877 | 910 |
| TCTTGACGCAGCTTAACACA | MTOR | 511 | 528 | 973 | 958 | 965 |
| TTAGGAAATGATCCGCACAG | MTOR | 453 | 497 | 910 | 808 | 773 |
| ATTGCACCCAATCCCAACTG | MTUS2 | 661 | 430 | 862 | 956 | 929 |
| CTGAGACTGAAAGCACCCTG | MTUS2 | 551 | 698 | 1014 | 1076 | 1084 |
| GCTGAAATTTGTCCAAGCCA | MTUS2 | 520 | 543 | 894 | 903 | 883 |
| GGGAGGCGTGAGTATCCGGG | MTUS2 | 441 | 726 | 1283 | 1314 | 1412 |
| TGAAGTGTCCACTTGGCCGA | MTUS2 | 578 | 982 | 1515 | 1607 | 1722 |
| TTGACAATGAGGAATTGCGA | MTUS2 | 470 | 495 | 1057 | 889 | 924 |
| CAGCACCAATGGCTCGGACA | MUTYH | 478 | 599 | 1173 | 1020 | 1111 |
| CTTACCACACTCCTCCACGT | MUTYH | 461 | 321 | 561 | 487 | 499 |
| GCTGCCTGGTTAAAATCCCC | MUTYH | 399 | 257 | 322 | 381 | 364 |
| GGCCTCGAGAGTAGTAACCC | MUTYH | 482 | 503 | 1028 | 982 | 1018 |
| GTTCAGTAGGTTCTCCCGAA | MUTYH | 535 | 689 | 870 | 966 | 990 |
| TGACCTCTGAGACCCACACT | MUTYH | 480 | 487 | 1064 | 1221 | 1173 |
| AGAAATACGGTCCGAAACGT | MYB | 395 | 460 | 870 | 848 | 779 |
| AGGTGCATGATCGTCCACCA | MYB | 496 | 631 | 1239 | 1117 | 1229 |
| GCTGACGTCAGCAAATACGA | MYB | 632 | 369 | 654 | 489 | 513 |
| GGAATAGTCATTGTTAACTG | MYB | 369 | 697 | 1359 | 1120 | 1223 |
| GGGGAAAACTAGGTGGACCA | MYB | 579 | 598 | 907 | 937 | 1014 |
| GGGTGAACAAACTCAAGTTG | MYB | 507 | 934 | 1743 | 1877 | 1842 |
| AAATGGTCCATAGTCGCACA | MYBL1 | 608 | 484 | 961 | 891 | 955 |
| AATTCTCATAAGTTTGACTG | MYBL1 | 557 | 734 | 1010 | 965 | 1004 |
| AGAAATATGGGCCAAAACGA | MYBL1 | 465 | 601 | 920 | 868 | 949 |
| ATAAGCGGTTGGGAAATCGT | MYBL1 | 602 | 442 | 733 | 751 | 795 |
| TAATACTCTAAATAGCCTCG | MYBL1 | 542 | 644 | 974 | 785 | 853 |
| TCCTGTAACATCCCCGAATG | MYBL1 | 329 | 624 | 1299 | 1307 | 1251 |
| AGAAATTGATGTTGTTTCTG | MYC | 478 | 1101 | 1959 | 1966 | 1872 |
| CAGGCTGCGCGCAAAGACAG | MYC | 613 | 445 | 1039 | 954 | 950 |
| CGAGTGCATCGACCCCTCCG | MYC | 505 | 401 | 962 | 905 | 847 |
| CTATGACCTCGACTACGACT | MYC | 470 | 339 | 664 | 568 | 583 |
| GGACGCGACTGCGACGTAGG | MYC | 684 | 437 | 1057 | 994 | 993 |
| TCATCATCCAGGACTGCATG | MYC | 457 | 466 | 1023 | 889 | 897 |
| CGAGGAAGAGGAAATTGACG | MYCN | 473 | 653 | 1215 | 1339 | 1323 |
| CTCTCCGTACGTGGAGAGCG | MYCN | 514 | 903 | 1740 | 1654 | 1856 |
| GAGTCCCCGCAACTCCGACT | MYCN | 534 | 417 | 637 | 657 | 684 |
| GCTGGCGTTCCAGAATGTTG | MYCN | 345 | 1053 | 1490 | 1703 | 1635 |
| GGCCCGGGGAGGGCGCAGTC | MYCN | 546 | 666 | 1070 | 975 | 952 |
| TCAAGCTCTTGGCCTTCGGG | MYCN | 473 | 631 | 1344 | 1229 | 1276 |
| ACCTGAGGGATCATCAAGCA | MYD88 | 558 | 624 | 1071 | 1023 | 938 |
| ACGACGTGTTGGTGGAACTG | MYD88 | 500 | 808 | 1679 | 1742 | 1655 |
| AGCTGGAACAGACAAACTAT | MYD88 | 368 | 535 | 984 | 838 | 855 |
| CGCACGTTGAGAGCAGCCAG | MYD88 | 573 | 731 | 1341 | 1209 | 1247 |
| CTCAAGCAGGCGGCCCACCG | MYD88 | 409 | 705 | 1436 | 1311 | 1419 |
| GAACAAACTGGATATCGCTG | MYD88 | 502 | 584 | 1073 | 1085 | 1090 |
| AGGATGGAGAAGATCCCCAT | MYH1 | 442 | 613 | 1109 | 1139 | 1137 |
| CAGTGCCAACCCCTTACTGG | MYH1 | 538 | 667 | 1234 | 1280 | 1323 |
| CTTTACAGTCGCTGACAAGG | MYH1 | 446 | 532 | 846 | 847 | 952 |
| GACACAGAACAGACCTGAGT | MYH1 | 533 | 471 | 731 | 750 | 763 |
| GGTCCACATCGATGGACCCA | MYH1 | 489 | 509 | 1029 | 1078 | 989 |
| TACCCTGAAGAGAGCAGACA | MYH1 | 440 | 625 | 854 | 1038 | 1074 |
| CAAAATCTTCTTCCGAACGG | MYH11 | 401 | 652 | 1050 | 849 | 908 |
| CTGCTCAAACGAGTTCACCT | MYH11 | 639 | 387 | 650 | 556 | 612 |
| CTTAACAACGAATCTTGCAG | MYH11 | 478 | 455 | 771 | 845 | 876 |
| GACGTCACCGGCTACATCGT | MYH11 | 509 | 404 | 823 | 644 | 655 |
| GCAGTCCAAGTGTAGCGATG | MYH11 | 513 | 972 | 1645 | 1543 | 1506 |
| GGGTGTGGTATTGCGTAGCG | MYH11 | 484 | 680 | 1443 | 1237 | 1334 |
| AGTGTCTGAGACTTACAAAG | MYH2 | 392 | 565 | 1004 | 977 | 1044 |
| GGTCAATGTCGATGGATGCA | MYH2 | 462 | 798 | 1540 | 1473 | 1481 |
| TAAGGTGAAAAACCTCACTG | MYH2 | 472 | 694 | 1175 | 1366 | 1270 |
| TATATTGTAGTCCACAACGC | MYH2 | 583 | 262 | 567 | 482 | 554 |
| TCACTTGGTCCTCCAGAGTG | MYH2 | 541 | 990 | 1796 | 1656 | 1680 |
| TGTGGTGGTTGAAGAACTGT | MYH2 | 382 | 763 | 1537 | 1501 | 1543 |
| AACGTGACGGATTTCACCCG | MYH9 | 419 | 693 | 1321 | 1245 | 1366 |
| CAGGTCCAAACGCGAACAGG | MYH9 | 430 | 548 | 1159 | 1280 | 1287 |
| GAAGCTCAAGAACAAACATG | MYH9 | 530 | 510 | 1029 | 1100 | 1128 |
| GCCTACAGGAGTATGATGCA | MYH9 | 645 | 846 | 1482 | 1287 | 1433 |
| GTGTCAAGATCTCATATCTG | MYH9 | 504 | 839 | 1440 | 1305 | 1320 |
| TGATGACTGGTGCAGAAGCG | MYH9 | 444 | 685 | 1239 | 1448 | 1367 |
| ATGGACTAGTTGGTTATATG | NAE1 | 526 | 1127 | 1776 | 1972 | 1934 |
| CAAGGAAATTTACCTGTTCG | NAE1 | 367 | 934 | 1467 | 1449 | 1383 |
| CACAGCCCTAAATACAACTC | NAE1 | 529 | 420 | 660 | 752 | 742 |
| CTTAAAAACTTAGTACTACC | NAE1 | 608 | 465 | 467 | 459 | 439 |
| GTAAACCTACAGAAAAATGA | NAE1 | 538 | 352 | 523 | 444 | 489 |
| TAGCCAAATATTTAGCACAG | NAE1 | 460 | 334 | 602 | 514 | 593 |
| AACGCGTGTGAAAAAATATG | NAMPT | 540 | 599 | 1136 | 1089 | 1058 |
| AATAAAGAAGGTGAAATACG | NAMPT | 537 | 572 | 1007 | 870 | 959 |
| CAGAGGAGTTTCTTCCCAAG | NAMPT | 634 | 646 | 1083 | 1043 | 1118 |
| GAGGGCTATGTCATTCCCCG | NAMPT | 602 | 698 | 1414 | 1330 | 1307 |
| GGCTACTGTGATTGGATACC | NAMPT | 533 | 661 | 1276 | 1134 | 1070 |
| TCCTAGTACCATAACAGCCT | NAMPT | 431 | 274 | 474 | 492 | 509 |
| AATGACGAAAGAATGCAGCA | NBN | 389 | 395 | 672 | 643 | 655 |
| ATAGCTGGTTCATCAATAGG | NBN | 514 | 723 | 1460 | 1388 | 1445 |
| CACAACGTGTGTAGCTGACA | NBN | 453 | 965 | 1650 | 1427 | 1497 |
| GAGAGTTACTGTAATCCTCG | NBN | 566 | 774 | 1513 | 1345 | 1392 |
| GGAATTTGTGTTGTTGATGT | NBN | 531 | 843 | 1190 | 1212 | 1371 |
| GTCAAAGTTACCATTAAAGT | NBN | 520 | 302 | 472 | 483 | 479 |
| AAAGCGCACACTCGTCGGCG | NCEH1 | 596 | 532 | 1031 | 998 | 1072 |
| ACTGTGTACAGCAATGGCTG | NCEH1 | 354 | 1095 | 1729 | 1847 | 1757 |
| ATCGTGTACATCCACGGAGG | NCEH1 | 460 | 663 | 1449 | 1433 | 1400 |
| TACTTCACCATGACGTAACG | NCEH1 | 604 | 645 | 964 | 974 | 894 |
| TCGTGGCGCGTACGACATCG | NCEH1 | 383 | 779 | 1237 | 1436 | 1506 |
| TGTCACCAGAGATGCCGATC | NCEH1 | 527 | 501 | 930 | 926 | 935 |
| AAATTCTGGTCCGTAGGAGT | NCSTN | 581 | 643 | 911 | 948 | 1028 |
| AAGACGAACATGACATTGCG | NCSTN | 509 | 743 | 1164 | 1301 | 1311 |
| AGAGAACATTGACTCATTCG | NCSTN | 402 | 932 | 1627 | 1745 | 1653 |
| CCAGCAGAACCACGTATGGA | NCSTN | 467 | 789 | 1315 | 1170 | 1289 |
| CCTTCAGCATCAACCCAGGT | NCSTN | 560 | 529 | 904 | 897 | 823 |
| TTACCGGTTATGGAAGACCG | NCSTN | 630 | 764 | 1372 | 1393 | 1340 |
| CAGGTGGAACTCCGAAGTGA | NECTIN4 | 415 | 790 | 1471 | 1374 | 1352 |
| CTCCCTCGTACAACTGGACA | NECTIN4 | 390 | 516 | 943 | 839 | 845 |
| GCACTCGTACTCGCCCTCGT | NECTIN4 | 641 | 449 | 820 | 738 | 751 |
| GCCTTGAAGACCAAAAGCTG | NECTIN4 | 449 | 945 | 1289 | 1302 | 1293 |
| GGGTCTCCTCGGTAGAAGCA | NECTIN4 | 578 | 749 | 1393 | 1130 | 1404 |
| TCCTGGCGTCATTCACAGGT | NECTIN4 | 470 | 604 | 761 | 651 | 669 |
| ATGCTAATTAAAGTGAAGGT | NEDD8 | 511 | 444 | 861 | 881 | 876 |
| CAGCAGCGGCTCATCTATAG | NEDD8 | 478 | 758 | 1205 | 1211 | 1233 |
| GAGCGTGTGGAGGAGAAAGA | NEDD8 | 560 | 471 | 1100 | 977 | 988 |
| GATGAGCCGCTGCTGCTGTG | NEDD8 | 378 | 880 | 1393 | 1591 | 1583 |
| GGAGCGAATCAAGGAGCGTG | NEDD8 | 563 | 514 | 1090 | 991 | 1032 |
| TGACATTGAACCCACAGACA | NEDD8 | 533 | 500 | 996 | 838 | 887 |
| AGAGATGTGCCGCCAAGAGA | NEGR1 | 496 | 907 | 1571 | 1683 | 1726 |
| AGTGATATGACCATCAACGA | NEGR1 | 365 | 371 | 747 | 811 | 847 |
| CAGCCTCCAGATACAGAATG | NEGR1 | 266 | 1064 | 1834 | 1785 | 1806 |
| GGTCAGAAAAGGGGACACGG | NEGR1 | 399 | 666 | 1212 | 1105 | 1199 |
| TGGAGGTGATAAATGGTCAG | NEGR1 | 551 | 476 | 1248 | 1323 | 1421 |
| TTCAGACTCAACATACGCCG | NEGR1 | 429 | 406 | 623 | 639 | 619 |
| AGACATTCGTCCGTCCAGCT | NEUROD1 | 545 | 472 | 1005 | 1048 | 1077 |
| CTGCACGAAGCACACCAGGT | NEUROD1 | 629 | 570 | 960 | 1055 | 1068 |
| GAGCTGTCCATGGTGCCGTA | NEUROD1 | 523 | 731 | 1146 | 1150 | 1154 |
| GCTGCCTGCAGCTCAACCCT | NEUROD1 | 453 | 678 | 1167 | 1439 | 1324 |
| GGAAGCCATGAACGCCGAGG | NEUROD1 | 596 | 626 | 1189 | 967 | 1113 |
| TTCTGCGTCTTGGAGTAGCA | NEUROD1 | 502 | 706 | 1250 | 1423 | 1369 |
| CAACTCGCGCAACTTCCTCA | NEUROD2 | 540 | 422 | 651 | 610 | 591 |
| CCAGGCGCGAGCACGGGTAC | NEUROD2 | 492 | 569 | 1076 | 1049 | 1128 |
| CCGCATGCACGACCTGAACG | NEUROD2 | 288 | 632 | 1101 | 1226 | 1127 |
| GACGAGCCGAGGAGCGACAA | NEUROD2 | 779 | 666 | 1116 | 999 | 1046 |
| GGCACCTCGTCTGCACGGAG | NEUROD2 | 478 | 563 | 1082 | 1057 | 1180 |
| TGCACGTAGGACACCAGGTC | NEUROD2 | 516 | 542 | 1084 | 1131 | 1044 |
| AGCACACCCAGCAATACGAA | NF1 | 603 | 306 | 580 | 471 | 512 |
| AGTACACCAAGTATCACGAG | NF1 | 739 | 638 | 931 | 1298 | 1191 |
| CTGGGAACACGGGACCACGT | NF1 | 525 | 579 | 921 | 943 | 999 |
| GAAACTCTGTCAGTTAGTCG | NF1 | 444 | 744 | 1321 | 1375 | 1388 |
| TCCCACTGCAGGAAATACCG | NF1 | 575 | 400 | 766 | 825 | 771 |
| TGCTTCGTACGGTGAGACAA | NF1 | 366 | 876 | 1530 | 1896 | 1836 |
| CTCGCAATTGAACTCCATCT | NF2 | 408 | 578 | 1639 | 1966 | 1911 |
| GACCCCTCTGTTCACAAACG | NF2 | 625 | 803 | 1607 | 1697 | 1689 |
| GCACAAGGGATGAATTGGAG | NF2 | 626 | 786 | 1572 | 1674 | 1762 |
| GGAATATTTGAAGATAGCTC | NF2 | 441 | 667 | 1298 | 1482 | 1604 |
| TAATATCTCATACAGCGACA | NF2 | 592 | 380 | 741 | 897 | 939 |
| TGGACTGCAGTACACGATCA | NF2 | 430 | 1043 | 2312 | 3107 | 2934 |
| ACTGAACACAAGTCCTGGCA | NFE2L2 | 662 | 486 | 797 | 801 | 716 |
| ATCAGCAATGACTGACTGAA | NFE2L2 | 603 | 415 | 949 | 876 | 892 |
| CAAGTTGGTCGAGACTAGCA | NFE2L2 | 591 | 778 | 1265 | 1373 | 1277 |
| CAGATGCACTGGTTTCCGAT | NFE2L2 | 517 | 579 | 936 | 943 | 1007 |
| CCCGTTCGTAGATGACAATG | NFE2L2 | 457 | 929 | 1635 | 1777 | 1786 |
| TATTTGACTTCAGTCAACGA | NFE2L2 | 332 | 333 | 653 | 644 | 640 |
| AATGACAGAGGCATGTACGA | NFKB1 | 505 | 557 | 947 | 863 | 849 |
| AATTTCTCAGATAGTTTCGG | NFKB1 | 579 | 365 | 667 | 659 | 704 |
| GAACCACTAAATCTAATGCC | NFKB1 | 684 | 384 | 737 | 494 | 605 |
| TAGATGGCATCTGACACTAC | NFKB1 | 352 | 368 | 686 | 704 | 735 |
| TCTTCATGCTCAACTTGTGA | NFKB1 | 571 | 712 | 1285 | 1192 | 1338 |
| TTGTCTATGAACATCTGTGG | NFKB1 | 625 | 607 | 1177 | 1347 | 1266 |
| CACCGAGGACGGAGACTCGT | NFKBIA | 398 | 495 | 947 | 817 | 848 |
| CCACAATGCCCAAATAGCCA | NFKBIA | 654 | 447 | 894 | 751 | 795 |
| CTGGACGACCGCCACGACAG | NFKBIA | 503 | 729 | 1102 | 1060 | 1043 |
| GAAGTGGTCCGCCAAGTGAA | NFKBIA | 514 | 837 | 1323 | 1274 | 1213 |
| GGGTATTTCCTCGAAAGTCT | NFKBIA | 589 | 580 | 991 | 878 | 862 |
| TTTGTAGACTCCACTCCACT | NFKBIA | 450 | 416 | 796 | 669 | 747 |
| CAGTGTCAGGACTTCCACGG | NFKBIZ | 685 | 504 | 1012 | 1024 | 1031 |
| CCTGGGCAACAGCAATATGG | NFKBIZ | 361 | 500 | 871 | 897 | 850 |
| CGGCGCTGAAGTGCTTAGCG | NFKBIZ | 675 | 461 | 932 | 929 | 885 |
| GGTCCATCAGACAACGAATC | NFKBIZ | 530 | 562 | 717 | 796 | 751 |
| GTGGTTCTTGACCCTCAAAG | NFKBIZ | 551 | 604 | 1058 | 1002 | 1048 |
| TATCCGAAGTCATAAACAGA | NFKBIZ | 359 | 407 | 509 | 437 | 501 |
| AAACGAGCGATTCGGCGAAT | NO_SITE_10 | 465 | 702 | 1130 | 1055 | 1085 |
| ACCCTAGCGCGCGTATCGAA | NO_SITE_100 | 961 | 443 | 722 | 710 | 735 |
| ACCCTATCGCGACGAACGGT | NO_SITE_101 | 799 | 665 | 1130 | 1230 | 1140 |
| ACCGGACGGCCGATTCGCGT | NO_SITE_105 | 536 | 581 | 1023 | 1051 | 1026 |
| ACCTCGCACGCCGTCGGAAT | NO_SITE_106 | 405 | 637 | 874 | 901 | 855 |
| ACCTTACGTCGCGGCGATAT | NO_SITE_109 | 543 | 374 | 959 | 977 | 988 |
| ACGACCCCGTGCGTCGCGTA | NO_SITE_110 | 533 | 555 | 1068 | 1109 | 1053 |
| ACGACGATTACCGTCCGACC | NO_SITE_111 | 629 | 592 | 1029 | 919 | 897 |
| ACGAGTCGACGCGTCGTTAC | NO_SITE_112 | 613 | 509 | 789 | 782 | 782 |
| ACGATTAACGCTCGCGCCCG | NO_SITE_113 | 433 | 851 | 1441 | 1620 | 1686 |
| ACGCGCCCTTACGCCGATAG | NO_SITE_114 | 619 | 765 | 1198 | 1113 | 1112 |
| ACGGCGTTACGGACGCGAAT | NO_SITE_116 | 584 | 785 | 1534 | 1336 | 1426 |
| ACGGTCGCGCGTCGTATAGA | NO_SITE_117 | 554 | 897 | 1384 | 1365 | 1313 |
| ACGTAACTAATACGCGAACG | NO_SITE_118 | 583 | 643 | 899 | 890 | 852 |
| ACTAATCTCGCGCTCGTATG | NO_SITE_119 | 646 | 1036 | 1627 | 1796 | 1886 |
| ACTGAACGTTCGACGTAACG | NO_SITE_124 | 526 | 1043 | 1610 | 1428 | 1486 |
| AGAACGCCCGATCACGATTA | NO_SITE_129 | 449 | 767 | 1574 | 1557 | 1502 |
| AAACGCGACTAGTCGCTGAT | NO_SITE_13 | 415 | 729 | 1164 | 1174 | 1215 |
| AGAATATGTACGCGCGACGC | NO_SITE_132 | 591 | 882 | 1320 | 1157 | 1158 |
| AGACCCGTATACGCCGTTAA | NO_SITE_135 | 440 | 664 | 1231 | 1285 | 1304 |
| AGATACTGTCGCGTCGCGAC | NO_SITE_138 | 346 | 388 | 716 | 958 | 896 |
| AGATCGCGCGACCGCAATTT | NO_SITE_139 | 304 | 541 | 962 | 1133 | 1128 |
| AAACGGCGATAACGCCGTAA | NO_SITE_14 | 710 | 578 | 1018 | 972 | 995 |
| AGATCGCGTGTACCGTAATA | NO_SITE_140 | 424 | 803 | 1433 | 1397 | 1540 |
| AGCCCGATGCGTGCCGATTA | NO_SITE_144 | 607 | 863 | 1455 | 1658 | 1532 |
| AGCCGTCGATATCGGCGGTT | NO_SITE_145 | 522 | 810 | 1470 | 1455 | 1339 |
| AGCGGCTAAAATATCGCGCT | NO_SITE_147 | 446 | 316 | 659 | 688 | 646 |
| AGCGTATAACCGGCCGATCG | NO_SITE_149 | 548 | 897 | 1548 | 1426 | 1336 |
| AGCGTCACGACGTTCGTATC | NO_SITE_150 | 643 | 637 | 970 | 972 | 920 |
| AGCTACGCCGCATTCGACCG | NO_SITE_151 | 617 | 848 | 1563 | 1480 | 1559 |
| AGCTGCTCGCTACGCCGAAT | NO_SITE_153 | 618 | 828 | 1273 | 1426 | 1415 |
| AGGACCCGATAGTTCGAACG | NO_SITE_157 | 606 | 705 | 1147 | 1172 | 1282 |
| AGGATCGTACGTTCCGCATT | NO_SITE_160 | 568 | 739 | 1074 | 1141 | 1202 |
| AGGCGATAGTCGGCCCGAAC | NO_SITE_162 | 341 | 457 | 932 | 918 | 887 |
| AGGCGCGCTTTACGTTCGAT | NO_SITE_164 | 610 | 681 | 1234 | 1177 | 1107 |
| AGGCGTTTCGACGCGAGGTA | NO_SITE_165 | 513 | 727 | 1395 | 1395 | 1376 |
| AAAGCGCGACCGCGTATAGT | NO_SITE_17 | 552 | 526 | 1010 | 984 | 1058 |
| AGGGCGACAACGATCGAAGT | NO_SITE_170 | 489 | 598 | 963 | 917 | 928 |
| AGTAACCGCGATACGTCGAT | NO_SITE_174 | 416 | 546 | 1303 | 1257 | 1321 |
| AGTATAAGCGCTCGCCGTAC | NO_SITE_175 | 540 | 816 | 1162 | 1324 | 1300 |
| AGTATAGTATCGTCCGCGAT | NO_SITE_176 | 512 | 611 | 1320 | 1322 | 1271 |
| AGTCCGTTCTAATCGCGTAG | NO_SITE_178 | 455 | 925 | 1510 | 1428 | 1405 |
| AGTCGGTCCACGTATCGATT | NO_SITE_180 | 549 | 799 | 1557 | 1535 | 1627 |
| AGTCGTACGCTCATCGGACC | NO_SITE_181 | 417 | 591 | 1040 | 1088 | 1075 |
| AGTGTCGCTTATAGCGCGTT | NO_SITE_183 | 530 | 617 | 1127 | 1042 | 1079 |
| AGTTAACTGTACCGCGCATT | NO_SITE_184 | 372 | 825 | 1106 | 1038 | 1091 |
| ATAACGAATTCGAACGCGTA | NO_SITE_185 | 534 | 819 | 1444 | 1539 | 1582 |
| ATAACGTAGTACGCGCACGG | NO_SITE_187 | 432 | 647 | 1268 | 1196 | 1196 |
| ATAAGCCGCTACGCGTCGAT | NO_SITE_188 | 408 | 645 | 1044 | 993 | 936 |
| ATAATCCGAACGGTCGTAGT | NO_SITE_189 | 7 | 637 | 838 | 861 | 858 |
| ATACGTTTGTCGCCGTAATA | NO_SITE_192 | 598 | 1005 | 1362 | 1448 | 1420 |
| ATACTAAATTCGCGCGATCC | NO_SITE_193 | 593 | 475 | 878 | 1018 | 1042 |
| ATATCGAATTAACGCGTCTA | NO_SITE_195 | 557 | 780 | 1459 | 1427 | 1408 |
| ATATCGTTCCGATACGTCTA | NO_SITE_196 | 529 | 774 | 1544 | 1359 | 1439 |
| ATCAACGACCGGCGTTTCGC | NO_SITE_197 | 449 | 163 | 291 | 223 | 226 |
| ATCAACGTACGTCGCACGAG | NO_SITE_198 | 501 | 790 | 1335 | 1239 | 1212 |
| ATCACTATGCCGGTTCGACG | NO_SITE_200 | 680 | 607 | 1238 | 1320 | 1387 |
| ATCAGCGTACTATCGCGCTT | NO_SITE_201 | 434 | 628 | 1223 | 1233 | 1223 |
| ATCATACGGCGCAACGTACC | NO_SITE_202 | 788 | 697 | 986 | 1123 | 1135 |
| ATCATCGACGTATATCGACG | NO_SITE_203 | 387 | 666 | 1209 | 1201 | 1170 |
| ATCCGAACGATAACGTACGT | NO_SITE_205 | 349 | 597 | 1043 | 1031 | 974 |
| ATCCGACGGAATCGTCCGTA | NO_SITE_206 | 609 | 812 | 1301 | 1354 | 1390 |
| ATCCGGTAATACGGCGCTTG | NO_SITE_207 | 574 | 1032 | 1648 | 1630 | 1699 |
| ATCGATCGTTCAATCGAGCG | NO_SITE_208 | 512 | 657 | 1175 | 1080 | 1038 |
| ATCGCTTACGTATCGTAACG | NO_SITE_209 | 526 | 811 | 1517 | 1456 | 1465 |
| ATCGGCTATCGTAACGCGAC | NO_SITE_210 | 494 | 359 | 631 | 599 | 567 |
| ATCTGTTACGCGTCGCGACT | NO_SITE_213 | 432 | 686 | 958 | 989 | 975 |
| ATGAATTATGCGGACGCGAG | NO_SITE_215 | 555 | 617 | 994 | 982 | 1152 |
| ATGCCGGATCGCGCAACTAA | NO_SITE_217 | 503 | 622 | 1218 | 1187 | 1156 |
| ATGCGACCATCGGGCGATAT | NO_SITE_218 | 660 | 675 | 993 | 1105 | 1123 |
| ATGCGATCGGACATCGCGTA | NO_SITE_220 | 460 | 607 | 996 | 939 | 988 |
| ATGCGCGATACCGTCTATAC | NO_SITE_221 | 543 | 416 | 719 | 716 | 738 |
| ATGCGCTGCGCGCGAATAAT | NO_SITE_222 | 439 | 721 | 1124 | 1079 | 1011 |
| ATGCTAGCGATCGCGCGTAA | NO_SITE_224 | 463 | 907 | 1520 | 1562 | 1517 |
| ATGCTTTAGCCGCGCAATCG | NO_SITE_225 | 593 | 998 | 1447 | 1468 | 1360 |
| ATGGGACGGCACTACGATAC | NO_SITE_226 | 648 | 479 | 803 | 798 | 748 |
| ATGGGATCGCCGATCGATTA | NO_SITE_227 | 488 | 686 | 1210 | 1197 | 1148 |
| ATGGTCACGCGTATCGCGAG | NO_SITE_229 | 581 | 935 | 1426 | 1551 | 1479 |
| ATGGTTCTAGCCGCCGATAA | NO_SITE_231 | 374 | 793 | 1375 | 1402 | 1416 |
| ATGTAGCGACATTCCGCGTT | NO_SITE_233 | 422 | 417 | 953 | 961 | 954 |
| ATGTCGTTACGGCTTGCGAC | NO_SITE_235 | 707 | 560 | 911 | 763 | 930 |
| AAATATAACCGATCGATCGC | NO_SITE_24 | 581 | 333 | 417 | 461 | 480 |
| ATTATATTACGCGCGCGAAA | NO_SITE_242 | 519 | 431 | 482 | 505 | 535 |
| ATTATTGTGCGACGCGGAAT | NO_SITE_243 | 492 | 835 | 1474 | 1338 | 1369 |
| ATTGACCGTTCGTCGCATAA | NO_SITE_246 | 611 | 592 | 1231 | 1248 | 1233 |
| ATTGCGTACTAGCGCGAATT | NO_SITE_247 | 516 | 541 | 1122 | 1144 | 1203 |
| ATTTAATTCGCGCGATCCAC | NO_SITE_251 | 516 | 485 | 541 | 495 | 545 |
| ATTTCGCCATACGCGACGAT | NO_SITE_252 | 412 | 491 | 996 | 1061 | 1118 |
| CAAACGAACTCGTCCGACCA | NO_SITE_253 | 445 | 466 | 1039 | 923 | 941 |
| CAAATCGTCGCGCGCGTATT | NO_SITE_255 | 526 | 589 | 884 | 914 | 914 |
| CAACCGCGTAATACCGCCGT | NO_SITE_256 | 814 | 436 | 906 | 910 | 841 |
| CAACGCGTACCGATAACTCT | NO_SITE_257 | 459 | 509 | 786 | 823 | 880 |
| CAAGCGTCAACCGTCCGTTA | NO_SITE_260 | 349 | 820 | 1248 | 1238 | 1177 |
| CAAGTCGTACCGTACGCACG | NO_SITE_261 | 478 | 674 | 1506 | 1286 | 1401 |
| CAATATCAATCACGCGTCCG | NO_SITE_264 | 702 | 759 | 1253 | 1321 | 1246 |
| CAATCGTCGTAACGACGAGT | NO_SITE_265 | 476 | 785 | 1282 | 1218 | 1245 |
| CACCGATAGTCGAACGCGAC | NO_SITE_275 | 576 | 381 | 537 | 644 | 578 |
| CACCGCAATCTACGGCGTAA | NO_SITE_276 | 456 | 661 | 1051 | 1052 | 967 |
| CACGTAAGCGCGTTATGCCG | NO_SITE_280 | 542 | 664 | 1278 | 1247 | 1283 |
| CACTCTACGTACGTCGATTC | NO_SITE_284 | 394 | 499 | 898 | 1002 | 997 |
| CACTTATCGTCCAACGAACG | NO_SITE_285 | 647 | 606 | 1183 | 1195 | 1131 |
| CAGAACGCGCGAAGCGCGAT | NO_SITE_289 | 542 | 524 | 1097 | 1102 | 1073 |
| CAGATCGCGTGTAATACGCG | NO_SITE_292 | 504 | 801 | 1447 | 1358 | 1475 |
| CAGCGCTTGCTCGTCGCTAT | NO_SITE_299 | 641 | 492 | 996 | 952 | 946 |
| AAAACGCTAGACGTACGCTA | NO_SITE_3 | 569 | 513 | 702 | 781 | 703 |
| CAGCGGCGATCGACGATCAT | NO_SITE_300 | 401 | 649 | 1088 | 945 | 1039 |
| CAGCTAATCCTCGCGATACG | NO_SITE_301 | 446 | 831 | 1641 | 1600 | 1673 |
| CAGGAGTTGTACGACGGTAC | NO_SITE_303 | 599 | 510 | 1079 | 1000 | 1102 |
| CAGGATCGATCGCGTAAGGT | NO_SITE_304 | 560 | 601 | 1058 | 1096 | 951 |
| CAGGCGCGACGCGATATACG | NO_SITE_306 | 488 | 1057 | 1660 | 1659 | 1598 |
| CAGTACGCGCGTTGTCGAAC | NO_SITE_308 | 354 | 552 | 902 | 910 | 890 |
| CAGTCGAACGGGCGTACTAG | NO_SITE_310 | 491 | 858 | 1196 | 1168 | 1221 |
| CAGTCGATAACGAAGGCGCG | NO_SITE_311 | 479 | 755 | 1570 | 1654 | 1666 |
| CATAACCGACGAATCGCGAT | NO_SITE_312 | 630 | 260 | 541 | 420 | 423 |
| CATAACGACCCGTCGATCGC | NO_SITE_313 | 618 | 530 | 1022 | 838 | 870 |
| CATATCGACCGACAAGCGAC | NO_SITE_316 | 489 | 375 | 418 | 439 | 447 |
| CATCGGCGCGATACCGATAA | NO_SITE_317 | 508 | 657 | 1070 | 1220 | 1189 |
| CATCGTACATCGACCGTTAG | NO_SITE_318 | 444 | 617 | 1250 | 1216 | 1155 |
| CATCGTACGTACGTCGAGTT | NO_SITE_319 | 491 | 420 | 624 | 671 | 587 |
| CATGGCGCGATACGTCGGAT | NO_SITE_322 | 485 | 748 | 1179 | 1265 | 1152 |
| CATTCGCGTCGAATCGACGT | NO_SITE_324 | 747 | 502 | 943 | 1072 | 1017 |
| CATTGTATCGAATCGCCGTA | NO_SITE_326 | 551 | 656 | 1192 | 1233 | 1188 |
| CCAACGCGAAACGTCGCGTA | NO_SITE_330 | 550 | 689 | 1333 | 1386 | 1440 |
| CCAACTCGTACGACGCGCAA | NO_SITE_331 | 612 | 443 | 675 | 686 | 720 |
| CCAAGAACGCGTACTACGCG | NO_SITE_332 | 513 | 708 | 1120 | 1117 | 1067 |
| CCAATCTCGACTAACGCGTT | NO_SITE_335 | 347 | 446 | 773 | 814 | 823 |
| CCACCAACGTCGATCGCGAT | NO_SITE_336 | 848 | 388 | 700 | 557 | 636 |
| CCACGCAACGGACGTTAGCG | NO_SITE_339 | 585 | 999 | 1416 | 1395 | 1446 |
| AACCGCGACTTTCGTACGGT | NO_SITE_34 | 367 | 504 | 1166 | 1190 | 1225 |
| CCACGCTCCGCCGTACGATT | NO_SITE_340 | 573 | 571 | 919 | 956 | 989 |
| CCACGTCGAATTAGCGAACC | NO_SITE_341 | 418 | 417 | 972 | 761 | 812 |
| CCAGTGCGACGTCGTTACGT | NO_SITE_347 | 368 | 618 | 1062 | 1117 | 1097 |
| CCATACGCGACCTGCGTAAC | NO_SITE_348 | 366 | 412 | 853 | 849 | 890 |
| CCATCGACTACGCGTCATTG | NO_SITE_349 | 399 | 995 | 1564 | 1668 | 1652 |
| AACCGCGTACGCTTCTCGTA | NO_SITE_35 | 780 | 563 | 970 | 940 | 885 |
| CCCCATCGCGGTTTCGCGTG | NO_SITE_354 | 648 | 702 | 1305 | 1452 | 1305 |
| CCCCGCCCGTCATCGAATAG | NO_SITE_356 | 503 | 610 | 1181 | 1158 | 1187 |
| CCCCGCCGTTCCGTCGAACT | NO_SITE_357 | 584 | 389 | 577 | 588 | 629 |
| CCCCGGTTACGTCGACGAAT | NO_SITE_358 | 499 | 789 | 1140 | 1091 | 1123 |
| CCCGCGTTGCGGCGCGTAAA | NO_SITE_361 | 420 | 1161 | 1674 | 1721 | 1779 |
| CCCGTATCGGACGTATTCGC | NO_SITE_363 | 412 | 345 | 584 | 694 | 633 |
| CCCGTCGGACTCCGTACGAT | NO_SITE_364 | 321 | 543 | 929 | 994 | 997 |
| CCCTACGAACTTACGCGCGT | NO_SITE_366 | 740 | 387 | 933 | 768 | 873 |
| CCCTACGTAATTCGCGTCGT | NO_SITE_367 | 535 | 520 | 960 | 988 | 1043 |
| CCCTATCGGAACGGTTCGCG | NO_SITE_368 | 366 | 499 | 865 | 879 | 874 |
| CCCTCCGGTCGGCGTACGTA | NO_SITE_369 | 496 | 769 | 1457 | 1317 | 1354 |
| AACTATTGACGGACGTATCG | NO_SITE_37 | 353 | 1102 | 1854 | 1859 | 1834 |
| CCCTTCGCGGTAGCCGTATA | NO_SITE_371 | 669 | 945 | 1442 | 1548 | 1491 |
| CCGAGGTTCGTCTTCGCGAC | NO_SITE_373 | 466 | 319 | 578 | 502 | 498 |
| CCGCGCGCTCGTGCGAAATT | NO_SITE_375 | 443 | 665 | 1007 | 1140 | 1151 |
| CCGCGGGCTTCGTATCGACT | NO_SITE_376 | 334 | 345 | 826 | 818 | 792 |
| CCGCGTCCGGCGACTTATTA | NO_SITE_377 | 451 | 690 | 1387 | 1329 | 1388 |
| CCGCGTCCGTAACGGTACGT | NO_SITE_378 | 696 | 544 | 936 | 1033 | 1022 |
| AACTCGATACGCGGACGATT | NO_SITE_38 | 562 | 714 | 1139 | 1138 | 1081 |
| CCGGCGATGTTACTTCGCGA | NO_SITE_380 | 439 | 543 | 1163 | 1154 | 1114 |
| CCGTACGATCGCCGCGCAAC | NO_SITE_381 | 425 | 409 | 706 | 898 | 913 |
| CCGTGAACGTAACTCGCATC | NO_SITE_382 | 583 | 380 | 937 | 912 | 895 |
| CCTACGCGAACCGCGACGAA | NO_SITE_383 | 456 | 752 | 1237 | 1204 | 1248 |
| CCTATCGCCGTACGATGGTT | NO_SITE_384 | 554 | 804 | 1250 | 1326 | 1344 |
| CCTCGCCGAGTATTACGGTC | NO_SITE_388 | 594 | 571 | 876 | 868 | 868 |
| AACTCGTACGCGCGCGAGAT | NO_SITE_39 | 407 | 538 | 1068 | 1196 | 1239 |
| CCTCGGGCACGCGATAAGTT | NO_SITE_390 | 502 | 523 | 1159 | 1131 | 1095 |
| CCTGAGTATCACGTCGCGAC | NO_SITE_391 | 581 | 214 | 405 | 368 | 389 |
| CCTTAGACGTACGCCGGTAA | NO_SITE_395 | 437 | 996 | 1601 | 1479 | 1499 |
| CCTTAGCCGAAATGCGCGAC | NO_SITE_396 | 638 | 401 | 705 | 624 | 607 |
| CCTTATTAGGCGATTCGCGC | NO_SITE_397 | 500 | 411 | 702 | 687 | 747 |
| CCTTCGACGGGCCTTATACG | NO_SITE_398 | 576 | 598 | 1200 | 1103 | 1189 |
| CCTTTCCGCGATTGCGATTA | NO_SITE_399 | 593 | 836 | 1368 | 1344 | 1354 |
| AAAACGTATCGTCCGAATCG | NO_SITE_4 | 486 | 846 | 1457 | 1632 | 1665 |
| AACTGTAATGCGACGATCGG | NO_SITE_40 | 488 | 725 | 1243 | 1262 | 1290 |
| CGAAACCTCGCGACGATCCG | NO_SITE_400 | 416 | 991 | 1494 | 1670 | 1545 |
| CGAAATTCGATCCGCGATTA | NO_SITE_401 | 514 | 639 | 1185 | 1325 | 1347 |
| CGAACGAGCGTACCTCGATA | NO_SITE_402 | 504 | 921 | 1526 | 1478 | 1527 |
| CGAATCGTATTCGCGCGATG | NO_SITE_406 | 527 | 780 | 1630 | 1435 | 1697 |
| CGACGATCTACTTCGATCGG | NO_SITE_408 | 396 | 600 | 1198 | 1391 | 1447 |
| CGAGCGCGTCTAATCGGTTA | NO_SITE_409 | 443 | 729 | 1175 | 1126 | 1140 |
| AACTTACCGGCGATCGTAAT | NO_SITE_41 | 587 | 416 | 898 | 890 | 963 |
| CGCAAACGCATACGCGATCG | NO_SITE_410 | 483 | 896 | 1377 | 1343 | 1350 |
| CGCACGTTCGTAGGCGTATA | NO_SITE_412 | 506 | 789 | 1463 | 1465 | 1483 |
| CGCCATCGGATCGCTACGAC | NO_SITE_414 | 414 | 407 | 503 | 539 | 505 |
| CGCCGATCGCGAATCGCAAA | NO_SITE_415 | 380 | 373 | 470 | 390 | 398 |
| CGCGTGATTAGTCGCGCGTA | NO_SITE_418 | 408 | 627 | 1204 | 1264 | 1439 |
| CGCTCGCGCCTCGTCCGATA | NO_SITE_419 | 687 | 863 | 1363 | 1271 | 1261 |
| AAGAACCGCGCCGATATGCG | NO_SITE_42 | 526 | 750 | 1259 | 1200 | 1282 |
| CGGACGGCGCAATACGACGT | NO_SITE_421 | 306 | 667 | 1075 | 1153 | 1200 |
| CGGAGCGAATCGAACGTCGT | NO_SITE_422 | 645 | 462 | 950 | 897 | 929 |
| CGGCGCGCGCTATGATCGAA | NO_SITE_424 | 348 | 526 | 896 | 911 | 862 |
| CGGCTCGTAGGACGTCCGAC | NO_SITE_426 | 521 | 424 | 743 | 677 | 713 |
| CGGTAGAAGATACGCGACGC | NO_SITE_431 | 472 | 581 | 1005 | 1185 | 1087 |
| CGTAACGGTTCGACGTACGT | NO_SITE_433 | 674 | 532 | 1090 | 1073 | 1065 |
| CGTACGGAACGGCGTTTCGT | NO_SITE_434 | 424 | 271 | 471 | 381 | 374 |
| CGTCGAACGCGACTTACCGC | NO_SITE_435 | 528 | 541 | 885 | 1035 | 1045 |
| CGTCGCGTAACGTTCGATCG | NO_SITE_436 | 460 | 742 | 1363 | 1365 | 1409 |
| CGTCGTACGTCCGATACGAG | NO_SITE_437 | 640 | 781 | 1324 | 1278 | 1281 |
| CGTGTGTAGTCGCGCAACGG | NO_SITE_439 | 532 | 871 | 1506 | 1387 | 1472 |
| CGTGTTAGGTCGTCGCGTAT | NO_SITE_440 | 558 | 645 | 1172 | 1381 | 1343 |
| CGTTCAATAGCGCGCGTCGA | NO_SITE_441 | 755 | 828 | 1388 | 1333 | 1368 |
| CGTTTACGTCTTATCCGTCG | NO_SITE_442 | 529 | 523 | 1248 | 1088 | 1096 |
| CGTTTGACGCCGTACGACGT | NO_SITE_443 | 726 | 595 | 975 | 878 | 902 |
| CTAAACGACGAACGCCGATC | NO_SITE_444 | 438 | 873 | 1464 | 1365 | 1466 |
| CTAAATCGCACGCGCGTTCT | NO_SITE_445 | 513 | 777 | 1293 | 1325 | 1307 |
| CTACGACAACGTATCGGCAA | NO_SITE_449 | 697 | 508 | 892 | 970 | 987 |
| CTACGACGGTCGATCGTTAC | NO_SITE_450 | 509 | 399 | 896 | 921 | 900 |
| CTACGACTATTTCGCGTAGT | NO_SITE_451 | 617 | 476 | 891 | 827 | 824 |
| CTACGCCGTAACGCGAAGTA | NO_SITE_452 | 570 | 560 | 1035 | 1030 | 1046 |
| CTACGTTGCGCTACGATCGT | NO_SITE_453 | 587 | 746 | 1321 | 1444 | 1400 |
| CTAGGGCCGCAACTATCGTA | NO_SITE_456 | 378 | 583 | 1222 | 1068 | 996 |
| CTAGTCATGACGCGCGATTT | NO_SITE_457 | 362 | 660 | 1111 | 1162 | 1213 |
| CTATAGCTCGACGACGGGTC | NO_SITE_460 | 421 | 587 | 1072 | 941 | 928 |
| CTCACCGCGCGATTACGTAA | NO_SITE_464 | 421 | 544 | 706 | 809 | 801 |
| CTCACGTTAGTACTCGCGAC | NO_SITE_465 | 568 | 353 | 418 | 404 | 405 |
| CTCACTACTCGTACGCGTAG | NO_SITE_466 | 469 | 810 | 1246 | 1420 | 1507 |
| CTCATAACGATCCGCGGACT | NO_SITE_467 | 449 | 546 | 989 | 903 | 878 |
| CTCCATCGATTAACCGACGC | NO_SITE_468 | 545 | 358 | 682 | 693 | 724 |
| CTCCCGCGTCGCTATATCGC | NO_SITE_469 | 537 | 584 | 997 | 950 | 1000 |
| AAGCCGTCGGTACGCAACGG | NO_SITE_47 | 550 | 704 | 1430 | 1279 | 1186 |
| CTCCCGTATTCGCGACGTAA | NO_SITE_471 | 508 | 446 | 957 | 913 | 871 |
| CTCCCTATCGCGCGGTCAAT | NO_SITE_472 | 663 | 803 | 1237 | 1275 | 1248 |
| CTCCGCCTACGTATCGGGAT | NO_SITE_476 | 752 | 576 | 1068 | 1083 | 1109 |
| CTCCGGATTACGCGCGTCAA | NO_SITE_477 | 584 | 854 | 1121 | 1335 | 1277 |
| CTCCGTCGATCGCGCCCGGT | NO_SITE_478 | 507 | 656 | 892 | 880 | 956 |
| CTCCGTTACATACGTCGCGC | NO_SITE_479 | 514 | 235 | 463 | 542 | 525 |
| CTCGTCGGCGCGATCTCGTA | NO_SITE_482 | 811 | 936 | 1325 | 1275 | 1271 |
| CTCTCGAGCATGCGCGACTA | NO_SITE_483 | 652 | 927 | 1436 | 1555 | 1524 |
| CTGAACGGTTCTACGCGACG | NO_SITE_486 | 505 | 652 | 1417 | 1448 | 1358 |
| CTGCGACGGGTCGACACTAT | NO_SITE_491 | 562 | 531 | 777 | 778 | 862 |
| CTGCGGTACGCGTCGATCGA | NO_SITE_492 | 393 | 1065 | 1344 | 1505 | 1504 |
| CTGCGGTATTACTCCGGTCG | NO_SITE_493 | 255 | 1138 | 1645 | 1776 | 1744 |
| CTGGTACCCGTACGATACGA | NO_SITE_496 | 338 | 596 | 895 | 827 | 786 |
| AAAACGTGCGCGAGTCGTAT | NO_SITE_5 | 579 | 552 | 1086 | 1102 | 1135 |
| AAGCGCCGTACGAGTAATCG | NO_SITE_50 | 528 | 604 | 1459 | 1315 | 1375 |
| CTTCCGCAAGTCGCGTTATA | NO_SITE_503 | 308 | 911 | 1506 | 1498 | 1495 |
| CTTCGATCAACGTCGTCTAC | NO_SITE_505 | 472 | 662 | 1131 | 1100 | 1146 |
| CTTCGCTACATCGTGCGATT | NO_SITE_507 | 662 | 470 | 897 | 860 | 907 |
| AAGCGCGCACCGTATCGACG | NO_SITE_51 | 607 | 748 | 1220 | 1171 | 1219 |
| CTTTCGTACGTCACGACTAT | NO_SITE_511 | 489 | 910 | 1249 | 1207 | 1264 |
| GAAAACGTCGAACGGCGTTA | NO_SITE_514 | 406 | 969 | 1502 | 1608 | 1557 |
| GAAACCCCGCGCGATCGATT | NO_SITE_517 | 559 | 456 | 911 | 808 | 846 |
| GAACGCCGTAAATTGGTCGT | NO_SITE_519 | 490 | 463 | 1021 | 970 | 942 |
| GAAGATACCGTCGCGCGATA | NO_SITE_521 | 619 | 660 | 1125 | 1098 | 1194 |
| GAAGCGCTCGATCGAACTAC | NO_SITE_522 | 348 | 430 | 830 | 849 | 814 |
| GAATAACGCTCGATCGTACC | NO_SITE_525 | 418 | 587 | 788 | 727 | 700 |
| GAATACACGCGATTCGTAAC | NO_SITE_526 | 558 | 342 | 536 | 560 | 591 |
| GAATCATACGTACGCGATTG | NO_SITE_527 | 619 | 740 | 1280 | 1463 | 1419 |
| GAATCGAGTAGGGTCGATCG | NO_SITE_528 | 482 | 768 | 1356 | 1298 | 1316 |
| GAATCTACGCCGTTCGCCGA | NO_SITE_529 | 325 | 536 | 967 | 933 | 942 |
| AAGCGTCGCGTAACGGCGAA | NO_SITE_53 | 587 | 603 | 1100 | 1104 | 1058 |
| GACAACCGTACGTCGACGAA | NO_SITE_530 | 567 | 713 | 1365 | 1389 | 1366 |
| GACAGCGTAGTCGCGTACGT | NO_SITE_532 | 451 | 403 | 842 | 790 | 872 |
| GACCGACCGACGATCAACGG | NO_SITE_535 | 532 | 436 | 1005 | 1029 | 1055 |
| GACCGCTATGCGTTACGCTA | NO_SITE_536 | 556 | 623 | 1183 | 1172 | 1127 |
| GACGTAACGCGACGCTACTA | NO_SITE_537 | 442 | 635 | 1145 | 1078 | 1032 |
| GACTACGGTATTACGTCGCG | NO_SITE_538 | 470 | 983 | 1450 | 1573 | 1489 |
| GACTCGATGCTCGTCGATAT | NO_SITE_539 | 528 | 717 | 1150 | 1167 | 1246 |
| AAGCGTCGTTCGCGCAATTT | NO_SITE_54 | 604 | 873 | 1225 | 991 | 1072 |
| GACTCGGGAATCGATACGCG | NO_SITE_540 | 459 | 597 | 1181 | 1117 | 1148 |
| GACTTACGCGTTATCGCCAT | NO_SITE_543 | 377 | 588 | 1170 | 1121 | 1215 |
| GAGCGACGTGTATACGAACT | NO_SITE_549 | 679 | 692 | 977 | 1026 | 1078 |
| GAGCGGAAGTACGCGACGAC | NO_SITE_551 | 486 | 429 | 612 | 621 | 631 |
| GAGGCGAAATCTAGCGACGT | NO_SITE_556 | 564 | 523 | 803 | 769 | 727 |
| GAGTTCGCGAACGTAGTGCG | NO_SITE_559 | 575 | 679 | 1209 | 1104 | 1161 |
| GATACGTACCCTATCGACGT | NO_SITE_562 | 573 | 365 | 742 | 715 | 678 |
| GATATCGCACCGTTCGCGTT | NO_SITE_564 | 581 | 310 | 585 | 661 | 661 |
| GATCGGCATTACGTCGCGTC | NO_SITE_565 | 422 | 354 | 659 | 571 | 617 |
| GATGCTTAGTCGTCGACCGT | NO_SITE_567 | 356 | 676 | 1050 | 1017 | 1132 |
| GATGTAAACGTATCGTCGCG | NO_SITE_568 | 651 | 448 | 903 | 959 | 903 |
| AAGTCGTCATAGTCGCGATC | NO_SITE_57 | 528 | 488 | 947 | 816 | 801 |
| GATGTTACGCGTTTCGGGAT | NO_SITE_570 | 522 | 543 | 956 | 812 | 839 |
| GCAAACGGCGATCGTACGCT | NO_SITE_573 | 493 | 485 | 914 | 881 | 838 |
| GCACCGCGCGAATTAAACGT | NO_SITE_574 | 449 | 454 | 783 | 789 | 824 |
| GCACCGCGGTATAACGGTTT | NO_SITE_575 | 410 | 918 | 1489 | 1380 | 1429 |
| GCACGGTACGCGTCGATGAC | NO_SITE_576 | 386 | 706 | 1032 | 952 | 1072 |
| GCAGACTCGTATCGCGCAAC | NO_SITE_578 | 528 | 603 | 1056 | 1041 | 1078 |
| GCAGCGCGATACGATCTAAC | NO_SITE_579 | 390 | 320 | 645 | 604 | 592 |
| GCAGTATACGCGGTTGATCG | NO_SITE_582 | 502 | 897 | 1709 | 1633 | 1685 |
| GCATAAACGCGCGCCGATGG | NO_SITE_584 | 469 | 786 | 1112 | 1011 | 1033 |
| GCATTCATAGCGCGCGCAAC | NO_SITE_588 | 515 | 380 | 780 | 813 | 697 |
| GCATTGGTATGCGTCGACGC | NO_SITE_589 | 535 | 394 | 722 | 753 | 774 |
| GCCACAACCGTCGCGACGTT | NO_SITE_591 | 581 | 534 | 1088 | 981 | 971 |
| GCCACGAGCGCGCGTTAATC | NO_SITE_592 | 441 | 767 | 1310 | 934 | 966 |
| GCCATCGGGCTACGCGACGA | NO_SITE_593 | 454 | 755 | 1383 | 1316 | 1338 |
| GCCCTATACGCGCATTAGCG | NO_SITE_594 | 665 | 938 | 1793 | 1838 | 1891 |
| GCCTGCGTAATACGACGGTC | NO_SITE_598 | 386 | 678 | 1070 | 1139 | 1131 |
| GCCTTAATCGAGTCGCGACG | NO_SITE_599 | 512 | 619 | 1095 | 1192 | 1195 |
| GCGACACGCCGTCGCGTTAT | NO_SITE_600 | 428 | 599 | 992 | 1068 | 1069 |
| GCGACGACGACGGTTGTAGC | NO_SITE_601 | 533 | 475 | 891 | 783 | 714 |
| GCGATCGCGGATATCGTACG | NO_SITE_602 | 541 | 739 | 1493 | 1419 | 1545 |
| GCGCAACGCTAATCCGTACG | NO_SITE_603 | 506 | 390 | 725 | 687 | 779 |
| GCGCATCCTCGTCGCGATAT | NO_SITE_604 | 395 | 552 | 1025 | 900 | 975 |
| GCGCGTACGCGCTTCGACGT | NO_SITE_605 | 341 | 344 | 849 | 856 | 932 |
| GCGCGTCGACGATTAATGCG | NO_SITE_606 | 410 | 633 | 1156 | 1173 | 1163 |
| GCTCCTATACCGGCGCGATC | NO_SITE_609 | 532 | 718 | 990 | 925 | 937 |
| GCTCGAACGGTCGACGCGAT | NO_SITE_611 | 586 | 448 | 943 | 902 | 836 |
| GCTCGCGCCGTTAGTCGTTT | NO_SITE_612 | 470 | 895 | 1576 | 1558 | 1578 |
| GCTCGGGGGATCGTCGACTA | NO_SITE_613 | 429 | 939 | 1675 | 1704 | 1655 |
| GCTGACTAGTCGCGTATCGG | NO_SITE_615 | 443 | 677 | 1368 | 1183 | 1178 |
| GCTTCGCGCGCAAATCCGTT | NO_SITE_624 | 384 | 528 | 914 | 935 | 891 |
| GCTTGATCCGTTCGCGACGG | NO_SITE_625 | 513 | 807 | 1125 | 1274 | 1242 |
| GGACGAACGTAACTCGACGA | NO_SITE_628 | 587 | 619 | 1269 | 1233 | 1326 |
| GGACGATATCCGATCGCGTA | NO_SITE_629 | 371 | 683 | 1087 | 1211 | 1127 |
| AATATATCGCGACGCGATAC | NO_SITE_63 | 599 | 414 | 816 | 763 | 767 |
| GGACGCGTTATACGACGACT | NO_SITE_630 | 508 | 420 | 868 | 874 | 898 |
| GGACGCTAGTACGGACGTAC | NO_SITE_631 | 599 | 776 | 1098 | 1079 | 1041 |
| GGACGGTGTTAACGCGAATT | NO_SITE_633 | 540 | 775 | 1230 | 1320 | 1389 |
| GGACTCGCCGTATACGGCAT | NO_SITE_634 | 631 | 738 | 1417 | 1520 | 1556 |
| AATCCACGATCGACGCGACT | NO_SITE_64 | 467 | 552 | 828 | 959 | 947 |
| GGAGTCGCGCAAACGCTAAT | NO_SITE_643 | 556 | 496 | 834 | 843 | 909 |
| GGAGTCGCGGTAAACCGTAT | NO_SITE_644 | 506 | 693 | 1172 | 1108 | 1101 |
| GGATACCGCTATCGAATCGT | NO_SITE_646 | 480 | 771 | 1220 | 1284 | 1215 |
| GGATGCGATTTACGCGCGAC | NO_SITE_649 | 307 | 457 | 679 | 668 | 622 |
| AATCCGTAACGTTACGTCGG | NO_SITE_65 | 549 | 732 | 1200 | 1137 | 1221 |
| GGCAGCGACGCGTACGCTAT | NO_SITE_654 | 354 | 688 | 1307 | 1369 | 1443 |
| GGCCGCGGTATTATCGTCGT | NO_SITE_657 | 378 | 563 | 942 | 998 | 945 |
| GGCCGCGTAGCGACAATCGA | NO_SITE_658 | 389 | 680 | 1340 | 1363 | 1393 |
| AATCGCGCACGTATAACGAT | NO_SITE_66 | 492 | 579 | 867 | 926 | 949 |
| GGCGTGTACGTCGACGCGTA | NO_SITE_660 | 612 | 798 | 1173 | 1237 | 1234 |
| GGCTCGCGTAATCGGCGTTA | NO_SITE_663 | 448 | 1165 | 1648 | 1896 | 1880 |
| GGCTCGTCGACCCGAACGAT | NO_SITE_664 | 481 | 673 | 1130 | 1128 | 1048 |
| GGCTTCGTATAACGCCGCGA | NO_SITE_665 | 503 | 698 | 1068 | 1320 | 1260 |
| GGGATATACGGTTCGCGCGT | NO_SITE_667 | 524 | 650 | 1147 | 1085 | 1104 |
| GGGATTCGGCGTCGTCGATT | NO_SITE_668 | 551 | 526 | 1160 | 1138 | 1214 |
| AATCGCGCGTCCGATACATT | NO_SITE_67 | 486 | 450 | 852 | 887 | 850 |
| GGGCGTTCAACGCCGATCGA | NO_SITE_674 | 487 | 659 | 1296 | 1216 | 1236 |
| GGGGCGACGTACCGTTCGAC | NO_SITE_676 | 524 | 467 | 668 | 745 | 689 |
| AATCGCGCGTCGTCAATCAT | NO_SITE_68 | 424 | 662 | 1029 | 1032 | 1108 |
| GGGTACTTAACGCGCGCCCA | NO_SITE_684 | 542 | 644 | 1224 | 1344 | 1317 |
| GGGTATTGCGTCGCCGCGAT | NO_SITE_685 | 585 | 455 | 794 | 831 | 679 |
| GGGTCTACGAAGCGCGTACG | NO_SITE_687 | 695 | 958 | 1465 | 1632 | 1670 |
| GGGTGTCGATATCGCGACGA | NO_SITE_689 | 440 | 853 | 1318 | 1565 | 1556 |
| AATCGGGTAATACGCGTTGT | NO_SITE_69 | 637 | 616 | 1151 | 1128 | 980 |
| GGGTTGTCGCTCGACGATTT | NO_SITE_692 | 515 | 594 | 1151 | 1257 | 1263 |
| GGTACGCCGGCGATTAGTAT | NO_SITE_693 | 336 | 822 | 1848 | 1710 | 1722 |
| GGTCACTAGATCGTCGATTA | NO_SITE_695 | 574 | 703 | 1486 | 1244 | 1296 |
| GGTCGGCCTAAACTTCGACG | NO_SITE_697 | 564 | 639 | 1345 | 1347 | 1368 |
| GGTCGGGTTTAACTCGATAC | NO_SITE_698 | 562 | 579 | 928 | 984 | 896 |
| GGTGGGTTCGCAATCGAACG | NO_SITE_700 | 418 | 837 | 1577 | 1615 | 1666 |
| GGTTGAACGATCGTCGCGAA | NO_SITE_703 | 469 | 553 | 946 | 976 | 940 |
| GTAAGGATATTCGTCGACGC | NO_SITE_708 | 642 | 327 | 705 | 796 | 802 |
| AATCGTTCGTTCGTATACGA | NO_SITE_71 | 626 | 495 | 748 | 665 | 676 |
| GTAATCCACGACGGTCGTAG | NO_SITE_710 | 710 | 687 | 1346 | 1313 | 1322 |
| GTAATTATACGCTTCGCTCG | NO_SITE_711 | 499 | 723 | 1047 | 1126 | 1079 |
| GTAGTGCGCTCACGCGATAA | NO_SITE_714 | 531 | 753 | 1354 | 1323 | 1422 |
| GTATAACTCGCGTCACGAAT | NO_SITE_715 | 474 | 360 | 552 | 547 | 520 |
| GTATAAGGTTCGTCGTACGA | NO_SITE_716 | 502 | 770 | 1389 | 1222 | 1376 |
| GTATGTTAACCGCGCGGATC | NO_SITE_718 | 539 | 477 | 936 | 964 | 959 |
| GTCCGATTTCGCAACGTTAG | NO_SITE_723 | 442 | 760 | 1202 | 1441 | 1342 |
| GTCGCATTCTAGCGCGTCGT | NO_SITE_725 | 565 | 507 | 891 | 878 | 926 |
| GTCGCGTAGACGTACGCTTA | NO_SITE_726 | 642 | 854 | 1400 | 1332 | 1330 |
| GTCTACCGTACGCGAGACAA | NO_SITE_728 | 693 | 758 | 1104 | 1053 | 1066 |
| GTCTATAACGGACGCGAGTT | NO_SITE_729 | 605 | 705 | 1057 | 912 | 995 |
| AATGCGGCAAACTTCGATAT | NO_SITE_73 | 445 | 463 | 867 | 944 | 954 |
| GTGCCTAACGCGTCCGTTAA | NO_SITE_738 | 467 | 671 | 854 | 799 | 832 |
| GTGGACCGATTAGCCGATCG | NO_SITE_741 | 456 | 811 | 1312 | 1335 | 1270 |
| GTGGTCCGCAATCGCGTATG | NO_SITE_744 | 624 | 955 | 1264 | 1421 | 1458 |
| GTGTCCGAACGCGTACGAAC | NO_SITE_745 | 562 | 268 | 405 | 419 | 418 |
| GTTCTCGCGACGCATATAAG | NO_SITE_748 | 463 | 562 | 898 | 923 | 879 |
| AATTCGCGGTCATTAACGCG | NO_SITE_75 | 442 | 551 | 1010 | 1003 | 1066 |
| GTTTCGCTTAACGTCGCGTA | NO_SITE_752 | 574 | 607 | 1172 | 1016 | 1003 |
| TAAACTCCGTAGCGTTACGA | NO_SITE_754 | 471 | 835 | 1214 | 1332 | 1340 |
| TAACGCGAACGACGTATAGA | NO_SITE_758 | 443 | 644 | 893 | 979 | 881 |
| TAACGCTCGAATCGTTACGC | NO_SITE_759 | 553 | 608 | 921 | 988 | 981 |
| TAAGCGCACGCGATACGGTC | NO_SITE_761 | 483 | 384 | 769 | 914 | 839 |
| TAAGTCCGACGATTTACGCT | NO_SITE_764 | 695 | 522 | 844 | 900 | 941 |
| TAATGACGCTACGCGCTATC | NO_SITE_766 | 350 | 437 | 771 | 710 | 766 |
| TAATGTCGTACGCTCGGAAC | NO_SITE_769 | 491 | 476 | 888 | 797 | 817 |
| ACAAATCGGTTACGCGCTAC | NO_SITE_77 | 470 | 652 | 1133 | 1068 | 1102 |
| TAATTCCGATCGTCGCGAGT | NO_SITE_770 | 541 | 748 | 1170 | 1109 | 1111 |
| TACAGCGCTACCGCTCGATA | NO_SITE_773 | 451 | 857 | 1312 | 1325 | 1490 |
| TACAGTCTACCATCGCGTAC | NO_SITE_775 | 293 | 502 | 740 | 857 | 815 |
| TACATGACGTGCCGCGCTAT | NO_SITE_776 | 616 | 639 | 1067 | 1074 | 1116 |
| TACCGTGAGAACGTCGTTAC | NO_SITE_777 | 549 | 609 | 1026 | 867 | 912 |
| TACCTAGCGACTACGAATCG | NO_SITE_778 | 480 | 990 | 1732 | 1562 | 1781 |
| TACGCGTAAGTAGTCGCGTG | NO_SITE_779 | 542 | 543 | 1186 | 1064 | 1246 |
| TACGCTTAACGCGCGATTAT | NO_SITE_780 | 583 | 906 | 1092 | 1200 | 1152 |
| TACGTCGCAATGTCGCGCTA | NO_SITE_781 | 554 | 888 | 1439 | 1552 | 1526 |
| TACTATTGGTCGGCGGATTA | NO_SITE_783 | 510 | 1046 | 1685 | 1656 | 1690 |
| TACTCGGCGGACGATTATCG | NO_SITE_784 | 403 | 882 | 1640 | 1803 | 1751 |
| TACTTTCGTACGCGATTCGC | NO_SITE_785 | 454 | 397 | 719 | 739 | 731 |
| TAGATTAGGTAGCGCGTCGA | NO_SITE_789 | 520 | 533 | 1222 | 1027 | 1161 |
| TAGCCGATATTATCGCGCTT | NO_SITE_790 | 393 | 722 | 1152 | 1075 | 1076 |
| TAGCGAATATTACGCGGGGT | NO_SITE_791 | 467 | 593 | 992 | 1112 | 1118 |
| TAGGACACGCTGCGCGAATT | NO_SITE_792 | 586 | 602 | 927 | 927 | 940 |
| TAGGATCGGATACGCAACGG | NO_SITE_793 | 744 | 783 | 1243 | 1386 | 1362 |
| TAGGTCGCGCTAGTAATCGT | NO_SITE_797 | 533 | 515 | 1129 | 1113 | 1053 |
| TAGTGCGCGCAACGATCGTT | NO_SITE_800 | 421 | 491 | 952 | 894 | 865 |
| TATACGCGTCCGATTAGCCG | NO_SITE_802 | 435 | 591 | 706 | 781 | 791 |
| TATCGTTACGCGTGTCGAAT | NO_SITE_803 | 457 | 835 | 1145 | 1313 | 1252 |
| TATGCTACCGTAATCGCGTA | NO_SITE_807 | 448 | 735 | 1169 | 1277 | 1225 |
| TCAACGATGCGACGCCGATT | NO_SITE_812 | 622 | 574 | 1083 | 988 | 914 |
| TCACGCACGCGCCCGAATAT | NO_SITE_815 | 520 | 834 | 1165 | 1257 | 1247 |
| TCACGGACTACGGCCGAAAT | NO_SITE_816 | 540 | 540 | 891 | 795 | 817 |
| ACACGGCGATTTCGTACGTC | NO_SITE_82 | 694 | 637 | 1159 | 1159 | 1159 |
| TCCACTCGACGCGTCGAATA | NO_SITE_823 | 577 | 598 | 971 | 1022 | 1024 |
| TCCGAATCGCCGCGCCCGAT | NO_SITE_827 | 681 | 482 | 1008 | 1109 | 1147 |
| TCCGACTATACGTCGTAACG | NO_SITE_829 | 542 | 900 | 1414 | 1504 | 1513 |
| ACACGTTATCGTACGCGAAA | NO_SITE_83 | 427 | 488 | 751 | 669 | 768 |
| TCCGCTTACCGGGTCGAATA | NO_SITE_830 | 458 | 745 | 1266 | 1131 | 1320 |
| TCCGGGCGACTATTCGACGG | NO_SITE_831 | 484 | 627 | 1127 | 995 | 1087 |
| TCCGGTAACGCGCGTCGATA | NO_SITE_832 | 463 | 741 | 1426 | 1481 | 1499 |
| TCCGTTCGCTAGATCGTACG | NO_SITE_833 | 511 | 823 | 1424 | 1296 | 1396 |
| TCCTTACGCGCGCGGATATG | NO_SITE_836 | 437 | 917 | 1464 | 1411 | 1457 |
| TCCTTGACGTCGTACGGTCG | NO_SITE_837 | 520 | 772 | 1440 | 1591 | 1583 |
| TCCTTTAATCGCGTCGGTCG | NO_SITE_839 | 503 | 513 | 1067 | 1046 | 1053 |
| TCGCGACGGTTTACGTTCGA | NO_SITE_840 | 347 | 550 | 1129 | 1165 | 1193 |
| TCGCGCATTAACGCGGCATA | NO_SITE_841 | 377 | 1085 | 1640 | 1718 | 1676 |
| TCGCGCGGGATACTCGAATT | NO_SITE_842 | 375 | 314 | 743 | 675 | 718 |
| TCGCGTTGATTCCGCGATAT | NO_SITE_843 | 480 | 562 | 1185 | 1098 | 1062 |
| TCGGCGTAACGCGCTACCGT | NO_SITE_844 | 455 | 381 | 910 | 958 | 947 |
| TCGTACGGAACGCATCGCGT | NO_SITE_845 | 675 | 583 | 1024 | 1015 | 1007 |
| TCGTATCGAAGGTACGTGCG | NO_SITE_847 | 666 | 1003 | 1433 | 1441 | 1487 |
| TCGTCGCGACGGCGTCAATT | NO_SITE_848 | 614 | 432 | 889 | 830 | 899 |
| TCGTTCAACGCGCGGCGAAT | NO_SITE_849 | 435 | 533 | 1179 | 1192 | 1109 |
| TCTGATTACGATCGCGATAC | NO_SITE_851 | 595 | 545 | 866 | 887 | 829 |
| TCTTAGCGGCGTTTCGTACG | NO_SITE_855 | 457 | 221 | 457 | 320 | 365 |
| TCTTGCCGTCTATCGTACGA | NO_SITE_857 | 681 | 721 | 1109 | 1119 | 1158 |
| TGAAAGTCGACGATCGACTA | NO_SITE_860 | 573 | 648 | 850 | 926 | 940 |
| TGAATACGTCGCCTTCGATA | NO_SITE_863 | 491 | 587 | 1199 | 1103 | 1193 |
| TGACGACGCGACGTTTCGCA | NO_SITE_864 | 578 | 417 | 747 | 753 | 735 |
| TGACGCTATTAAAGCGCGTC | NO_SITE_865 | 661 | 451 | 697 | 716 | 757 |
| TGACGGACGACGTATCGTAG | NO_SITE_866 | 357 | 1017 | 1461 | 1548 | 1538 |
| TGACGGCGGTTATCCGTCGT | NO_SITE_867 | 583 | 478 | 1067 | 973 | 985 |
| TGAGCGAATCGATACGCGAC | NO_SITE_868 | 474 | 398 | 620 | 553 | 490 |
| TGAGCGTGTGTCGACGAATA | NO_SITE_869 | 395 | 822 | 1406 | 1593 | 1689 |
| TGAGTATCGTCTATCGGACG | NO_SITE_870 | 423 | 766 | 1346 | 1368 | 1391 |
| TGAGTCCGAGCGTATCGACT | NO_SITE_872 | 409 | 603 | 1110 | 1070 | 1067 |
| TGAGTGCCGCTACGCGTATA | NO_SITE_873 | 485 | 880 | 1404 | 1513 | 1571 |
| TGATCGGATCTCGTCGTTAA | NO_SITE_875 | 456 | 692 | 1404 | 1381 | 1409 |
| TGATGAGCCGCGCGCGAATT | NO_SITE_876 | 337 | 620 | 940 | 898 | 908 |
| TGATTCACCCGAATCGATCG | NO_SITE_877 | 381 | 990 | 1325 | 1303 | 1303 |
| TGCAACGGATCGCAATCGCG | NO_SITE_878 | 550 | 598 | 1054 | 1030 | 988 |
| TGCAATACGCCGGTACGTCG | NO_SITE_879 | 507 | 683 | 1490 | 1487 | 1493 |
| TGCACGCCGACGCACGTTTA | NO_SITE_881 | 344 | 799 | 1782 | 1762 | 1835 |
| TGCCCGCGAATTACTCGCGT | NO_SITE_882 | 672 | 804 | 1220 | 1147 | 1166 |
| TGCCGCGCTACGCTATTATG | NO_SITE_883 | 529 | 860 | 1555 | 1434 | 1420 |
| TGCCGCGTTACGTTCGATAG | NO_SITE_884 | 444 | 879 | 1431 | 1412 | 1468 |
| TGCCGGTAAGCGATCGCGAT | NO_SITE_885 | 618 | 377 | 971 | 1107 | 1080 |
| TGCCGTAACCGATCGTAGCG | NO_SITE_886 | 475 | 830 | 1431 | 1495 | 1257 |
| TGCGCCCGCTCGTTACGAAC | NO_SITE_889 | 474 | 422 | 801 | 782 | 736 |
| TGCGCGACATCGATCGTTAT | NO_SITE_890 | 423 | 649 | 1100 | 1266 | 1272 |
| TGCGGAGCGTCGATCGTATA | NO_SITE_891 | 346 | 757 | 1613 | 1564 | 1638 |
| TGCTACCGGCCGTCACGTAA | NO_SITE_892 | 586 | 484 | 1065 | 836 | 842 |
| TGCTCCGATACGTTCGCACG | NO_SITE_895 | 568 | 737 | 1413 | 1437 | 1472 |
| TGCTTACGGTCGATTCGCGA | NO_SITE_897 | 569 | 432 | 633 | 618 | 607 |
| TGGCGACTACCGTCGAACCT | NO_SITE_899 | 358 | 560 | 923 | 1030 | 970 |
| AAACCGCATTACGACTCGAC | NO_SITE_9 | 489 | 243 | 476 | 338 | 358 |
| ACAGGTCGTCCGACCGCGTA | NO_SITE_90 | 636 | 665 | 1318 | 1232 | 1211 |
| TGGCGCACGATGTCGTTACG | NO_SITE_901 | 451 | 921 | 1875 | 1768 | 1851 |
| TGGCGCATAGTGCGTCGATA | NO_SITE_902 | 519 | 592 | 1252 | 1226 | 1254 |
| TGGCGGACTCGGCGTACGAA | NO_SITE_903 | 526 | 820 | 1435 | 1328 | 1362 |
| TGGCGTCACGATCGCTTAAC | NO_SITE_904 | 365 | 493 | 1072 | 1101 | 1149 |
| TGGGTCGATAACGTTCGACC | NO_SITE_908 | 352 | 573 | 938 | 856 | 961 |
| TGGTACCGGCTCGTCGCGAA | NO_SITE_909 | 330 | 642 | 1050 | 1193 | 1149 |
| ACATAGATCGACCGGACGTA | NO_SITE_91 | 491 | 1271 | 1841 | 1818 | 1843 |
| TGGTATACGGCCTTAACGCG | NO_SITE_910 | 462 | 899 | 1340 | 1558 | 1574 |
| TGGTGACCGAATATCGCGCC | NO_SITE_911 | 647 | 363 | 684 | 649 | 573 |
| TGTACGACGTACCGATTCGT | NO_SITE_912 | 390 | 571 | 1055 | 1183 | 1122 |
| TGTACGTGCCTATTCGCGAC | NO_SITE_913 | 481 | 298 | 509 | 568 | 607 |
| TGTACTGCGTATCGCGCAAT | NO_SITE_914 | 402 | 714 | 1117 | 1274 | 1183 |
| TGTCCGCGCATAGTCGCTTT | NO_SITE_916 | 643 | 675 | 1288 | 1262 | 1184 |
| TGTCCGTATTCGCGCGATAT | NO_SITE_917 | 396 | 1069 | 1508 | 1346 | 1423 |
| ACATTACGTCGCGACTCGAA | NO_SITE_92 | 482 | 496 | 715 | 657 | 691 |
| TGTCGGGCGTCTAATCGCGT | NO_SITE_920 | 594 | 468 | 966 | 1023 | 1076 |
| TGTGAACGCGATATAACCCG | NO_SITE_921 | 412 | 905 | 1488 | 1418 | 1383 |
| TGTGCGCGTAATACGTCGTT | NO_SITE_924 | 537 | 619 | 1145 | 1157 | 1131 |
| TGTGGACCTATGTACGATCG | NO_SITE_925 | 584 | 958 | 1720 | 1592 | 1643 |
| TGTGGGCTATCGCGGCGTAT | NO_SITE_926 | 563 | 724 | 1393 | 1294 | 1237 |
| ACCAATCGGCGTTCCGCCAT | NO_SITE_93 | 586 | 321 | 690 | 620 | 706 |
| TGTTTAGCTAGCGCCGATAC | NO_SITE_932 | 502 | 433 | 929 | 937 | 857 |
| TGTTTCTCGCGACTCGCGTA | NO_SITE_933 | 466 | 718 | 1143 | 1067 | 1128 |
| TTACCACGCCATCGCGATTT | NO_SITE_936 | 365 | 657 | 1069 | 1030 | 1074 |
| TTACGTTGCGTACGCGATAC | NO_SITE_939 | 504 | 569 | 1031 | 1084 | 1107 |
| TTAGTACTATTCGCACGCGA | NO_SITE_944 | 434 | 353 | 653 | 556 | 600 |
| TTATACGGCGGTTCGTAGTA | NO_SITE_947 | 407 | 1005 | 1543 | 1637 | 1589 |
| TTATATTACGCGACGAGCGA | NO_SITE_949 | 527 | 599 | 912 | 747 | 846 |
| ACCAGTATTTCGACGTCCGC | NO_SITE_95 | 553 | 418 | 844 | 790 | 852 |
| TTATCAACGGCCGCGCGTAT | NO_SITE_950 | 377 | 559 | 1212 | 1213 | 1204 |
| TTCACGCGACGCATCGACCG | NO_SITE_953 | 481 | 614 | 1428 | 1422 | 1355 |
| TTCAGGTTTACGCGCGAAAT | NO_SITE_954 | 391 | 674 | 1178 | 1183 | 1224 |
| TTCAGTCCGCGATCGAACTC | NO_SITE_955 | 408 | 595 | 1106 | 1001 | 1038 |
| TTCCGAACTATAACGCTCGA | NO_SITE_959 | 422 | 421 | 712 | 631 | 681 |
| TTCCGTACGGACCGTATTCG | NO_SITE_961 | 477 | 767 | 1567 | 1502 | 1542 |
| TTCCGTTATACCGTTACGGG | NO_SITE_962 | 432 | 608 | 1151 | 1140 | 1024 |
| TTCGACTACGTTAGCGACGT | NO_SITE_964 | 338 | 601 | 988 | 1069 | 1018 |
| TTCGCCGGTACCGTCGAACG | NO_SITE_966 | 257 | 586 | 1002 | 990 | 1029 |
| TTCGCGCTTATATCCGTCTA | NO_SITE_967 | 390 | 805 | 1520 | 1376 | 1548 |
| TTCGCGTTATCGCACGGATA | NO_SITE_968 | 298 | 913 | 1715 | 1707 | 1681 |
| TTCGTACGAGCGAACGACGT | NO_SITE_969 | 443 | 593 | 839 | 857 | 838 |
| ACCCCGATTCGCGTCTAGTC | NO_SITE_97 | 745 | 653 | 1011 | 974 | 960 |
| TTCTCCGTATCGACCGGACC | NO_SITE_971 | 389 | 420 | 745 | 754 | 798 |
| TTCTGCGATCGTTCGACCGG | NO_SITE_972 | 606 | 853 | 1364 | 1277 | 1383 |
| TTGCGTAAATCGTGCGTCAA | NO_SITE_977 | 411 | 759 | 1224 | 1175 | 1173 |
| TTGGTCGTCGCGTACCGGTT | NO_SITE_979 | 541 | 765 | 1355 | 1356 | 1362 |
| ACCCGCGCTACGCGTGAGAT | NO_SITE_98 | 531 | 840 | 1340 | 1430 | 1518 |
| TTGTACGTTCGATACTCGCG | NO_SITE_983 | 562 | 806 | 1425 | 1275 | 1225 |
| TTGTATTAACGCGCGACGCG | NO_SITE_984 | 487 | 833 | 1340 | 1357 | 1320 |
| TTGTCCATAGTACGCGCGAC | NO_SITE_985 | 402 | 273 | 572 | 578 | 572 |
| TTGTCGCGGCCGAAACCGTT | NO_SITE_986 | 577 | 327 | 885 | 763 | 733 |
| TTGTCGTATACCGGTCGCAT | NO_SITE_987 | 513 | 601 | 1061 | 1027 | 1108 |
| TTTACAGCGTACCGCGCAAT | NO_SITE_989 | 510 | 547 | 1025 | 1031 | 997 |
| TTTCCGGCGTCGCGTAATGC | NO_SITE_995 | 475 | 657 | 879 | 715 | 665 |
| ACGTGCCTGGATAGGATCGA | NOTCH1 | 439 | 754 | 1472 | 1402 | 1382 |
| CAAACACCCCATCAAACGTG | NOTCH1 | 566 | 689 | 1228 | 1044 | 1146 |
| GCATCTATGTCAGTATGACG | NOTCH1 | 444 | 621 | 1005 | 835 | 886 |
| GCATGCAACAACCATGCGTG | NOTCH1 | 409 | 807 | 1451 | 1288 | 1323 |
| TCCTGCCAGAACACCAACGG | NOTCH1 | 435 | 545 | 941 | 824 | 854 |
| TGTGCACTGTGAGGTCAATG | NOTCH1 | 623 | 873 | 1413 | 1565 | 1603 |
| ACATCGACGAGTGCACTGAG | NOTCH2 | 501 | 654 | 1398 | 1420 | 1389 |
| AGGTGCCACCATTAACACAG | NOTCH2 | 589 | 580 | 997 | 904 | 895 |
| ATCCATGGAAACTGTACCGG | NOTCH2 | 465 | 608 | 1189 | 1262 | 1081 |
| GACAACACGGCACTTACCGG | NOTCH2 | 663 | 536 | 935 | 740 | 810 |
| TGATGAGCTGTGCAACACAG | NOTCH2 | 618 | 483 | 859 | 864 | 862 |
| TTGATGTCCATCTCACAACG | NOTCH2 | 537 | 613 | 1188 | 1218 | 1209 |
| AAGGACCACTATACCCCCTG | NOTCH3 | 444 | 737 | 1441 | 1316 | 1236 |
| AGATCCCACCGTTGACACAT | NOTCH3 | 441 | 334 | 656 | 574 | 557 |
| GAGATTAATGAGGACGACTG | NOTCH3 | 497 | 550 | 1215 | 1077 | 1108 |
| GGGACAGGCACAGTAGAACG | NOTCH3 | 581 | 605 | 1252 | 1154 | 1136 |
| TAAATGTCAGCACACCCCCA | NOTCH3 | 516 | 474 | 1000 | 999 | 991 |
| TTGATCTCCACGTTGCAAAG | NOTCH3 | 549 | 792 | 1196 | 1274 | 1280 |
| CAATGGACAGAACTGCTCAA | NOTCH4 | 382 | 600 | 1039 | 1013 | 999 |
| GCCCACATCACACACACATG | NOTCH4 | 791 | 753 | 1399 | 1276 | 1353 |
| GGGTCCTGTGAGACCACAGC | NOTCH4 | 439 | 545 | 991 | 996 | 1111 |
| GTGGACGAATGTGAGACCCA | NOTCH4 | 478 | 598 | 1164 | 1019 | 1136 |
| TACCTGGGAGACACTCGCAA | NOTCH4 | 475 | 692 | 1188 | 967 | 1031 |
| TCAGTGCTGGTCTGACAGCG | NOTCH4 | 597 | 722 | 1103 | 1107 | 1093 |
| AATGCTCCATGACTACATTG | NPEPPS | 459 | 741 | 1030 | 903 | 869 |
| ACAGCTTCATATGCACCAGA | NPEPPS | 401 | 299 | 548 | 474 | 483 |
| ACGTAGACATGACAGGCGTG | NPEPPS | 596 | 823 | 1296 | 1317 | 1287 |
| CCTGTTCTTCATCACGCCAG | NPEPPS | 732 | 619 | 1140 | 1253 | 1190 |
| GTAATCCAAGTCGATCCACA | NPEPPS | 496 | 331 | 688 | 626 | 610 |
| TGTGATAGGAACCATCCACT | NPEPPS | 560 | 608 | 1035 | 1011 | 1030 |
| AAGTGGTAGCAAGGTTCCAC | NPM1 | 560 | 299 | 624 | 468 | 491 |
| AGTCTCAGTGCCCTTACCGT | NPM1 | 586 | 285 | 747 | 611 | 680 |
| ATTAGTGGACAGCACTTAGT | NPM1 | 687 | 512 | 886 | 909 | 976 |
| ATTTACAGGTTGTGAACTAA | NPM1 | 369 | 434 | 933 | 824 | 857 |
| GATAAGATAACCTATACCTG | NPM1 | 468 | 420 | 916 | 696 | 780 |
| TCACAGGTCAGTTTGGGAGC | NPM1 | 356 | 562 | 831 | 650 | 577 |
| AACAAACTCATCACTGTGTG | NPRL2 | 454 | 923 | 1544 | 1610 | 1736 |
| ACTTGAAGGTGATTGAGCAG | NPRL2 | 522 | 708 | 1259 | 1214 | 1226 |
| ATGGCGGAAACCATCAATGT | NPRL2 | 524 | 349 | 627 | 653 | 623 |
| GAGCAGCTTCGTGTCAACAG | NPRL2 | 531 | 415 | 861 | 805 | 884 |
| GGGCATCACACACAAAGCCT | NPRL2 | 539 | 216 | 450 | 376 | 421 |
| TCACAACTCCATAATACCTG | NPRL2 | 522 | 500 | 674 | 568 | 624 |
| AGGTACTCACAGACACACGC | NPRL3 | 533 | 335 | 380 | 335 | 354 |
| CACATTAAAGAGAATCATGG | NPRL3 | 657 | 316 | 674 | 681 | 706 |
| GATGTGGAGTCGTACCACGC | NPRL3 | 403 | 333 | 809 | 674 | 709 |
| GCTCCGTTCGATGGCCTCGG | NPRL3 | 493 | 483 | 1235 | 1130 | 1060 |
| GGTCTTGATCACACGCACCA | NPRL3 | 465 | 376 | 540 | 446 | 498 |
| GTGATTCTTGCAGCAAATGA | NPRL3 | 621 | 548 | 994 | 911 | 1046 |
| CATGGCGTACAGGTCCGACA | NQO1 | 371 | 493 | 1264 | 1110 | 1176 |
| CCGCCTGAGCCCCGACATCG | NQO1 | 650 | 556 | 1113 | 1084 | 1045 |
| CGGGAACTTTCAGTACCCCG | NQO1 | 523 | 606 | 933 | 856 | 969 |
| GAATGATATTCATGTCTCCG | NQO1 | 430 | 718 | 1404 | 1286 | 1379 |
| GGACTCCAAACCACTGCAGT | NQO1 | 560 | 426 | 758 | 738 | 787 |
| TACGCGGCCATGTATGACAA | NQO1 | 378 | 488 | 1044 | 1074 | 1088 |
| ACCATCTCGAGACATGCCCA | NR1D1 | 597 | 599 | 1209 | 1140 | 1167 |
| ATAGGTGAAGATTTCTCGGT | NR1D1 | 373 | 507 | 947 | 1078 | 1003 |
| CAGGGCTTCATTATGACGCT | NR1D1 | 541 | 517 | 898 | 920 | 897 |
| GGGGCTCCCATTATAGAAGG | NR1D1 | 421 | 598 | 1208 | 1211 | 1300 |
| TGGGTGGGTGAGGTTTCCAA | NR1D1 | 292 | 726 | 1352 | 1361 | 1370 |
| TGTCGCTGTAGAGAGACTCA | NR1D1 | 482 | 625 | 1404 | 1428 | 1537 |
| CAGCAAACAGAGGATGCCTC | NR1H4 | 521 | 558 | 832 | 1098 | 1160 |
| CCAGCTGAGACTGTCTACCA | NR1H4 | 478 | 623 | 1296 | 1340 | 1366 |
| CTCGACAACCAAGTCATGCA | NR1H4 | 464 | 255 | 525 | 500 | 455 |
| GCACACATCAGAATTTGTCA | NR1H4 | 472 | 623 | 811 | 670 | 659 |
| TAGGATGATGACGAAATCTG | NR1H4 | 417 | 848 | 1514 | 1579 | 1495 |
| TGTGTACAAGTGTAAAAACG | NR1H4 | 522 | 769 | 1263 | 1282 | 1132 |
| ACTGTCCAGCATGCCGCTAT | NR3C1 | 522 | 827 | 1454 | 1418 | 1452 |
| ATGCTGTATTCATGTCATAG | NR3C1 | 562 | 777 | 1182 | 1077 | 1131 |
| CATTGAACTCTGCACCCCTG | NR3C1 | 422 | 572 | 1403 | 1316 | 1454 |
| GACCCCGTAATGACACCCCG | NR3C1 | 461 | 478 | 925 | 851 | 920 |
| TAGAAAACACCGACCGACCA | NR3C1 | 352 | 500 | 950 | 953 | 1048 |
| TCTGGAAGGAAACTCGAACG | NR3C1 | 357 | 455 | 983 | 881 | 936 |
| AGCCGGGTTGGAGTCGACAT | NR4A2 | 525 | 579 | 1168 | 1122 | 1245 |
| AGGGGCATTTGGTACAAGCA | NR4A2 | 533 | 672 | 1379 | 1449 | 1446 |
| CCACGCACATGATCGAGCAG | NR4A2 | 340 | 928 | 1440 | 1299 | 1361 |
| GACAGTTTAAAAGGCCGGAG | NR4A2 | 649 | 598 | 1059 | 984 | 954 |
| GGGCTTGTAGTAAACCGACC | NR4A2 | 482 | 462 | 889 | 885 | 908 |
| GGGGCTCCCCCTCCAACGAG | NR4A2 | 569 | 671 | 1136 | 1333 | 1371 |
| AAATGATTTGCTATTATTGA | NRAS | 572 | 685 | 945 | 907 | 956 |
| CCATGAGAGACCAATACATG | NRAS | 714 | 726 | 1255 | 1319 | 1283 |
| GATCATATTCATCTACAAAG | NRAS | 624 | 301 | 399 | 394 | 410 |
| GATGTACCTATGGTGCTAGT | NRAS | 501 | 531 | 677 | 598 | 660 |
| GGATTCTTACCGAAAACAGG | NRAS | 612 | 448 | 599 | 523 | 571 |
| TTGCAGACATTAACCTCTAC | NRAS | 713 | 421 | 565 | 599 | 577 |
| AGAGGATCCCCGTCGACGAG | NRP2 | 664 | 815 | 1488 | 1514 | 1508 |
| ATGGGACCATCGAATCCCCT | NRP2 | 602 | 646 | 874 | 844 | 818 |
| GCACGTACTGCGCCGGAACA | NRP2 | 683 | 784 | 1490 | 1316 | 1291 |
| GCTGGAAGTCAGCACGAATG | NRP2 | 481 | 684 | 1265 | 1247 | 1227 |
| GGTGGACACCTCAACAAAGC | NRP2 | 433 | 425 | 589 | 510 | 504 |
| TGGGACTCCAGAGGTACTCG | NRP2 | 405 | 816 | 1378 | 1521 | 1574 |
| AATCACTTTACACCTAGGCG | NSD1 | 534 | 699 | 860 | 872 | 859 |
| AGAACTGATGACACACCGAA | NSD1 | 588 | 565 | 825 | 706 | 775 |
| CTATCGACAATACTATGTGG | NSD1 | 387 | 803 | 1157 | 1047 | 1109 |
| GAAAAGAAGAAACTCCAACG | NSD1 | 442 | 548 | 1103 | 1105 | 1148 |
| GTGTTACTAGGCGCAACTGT | NSD1 | 551 | 496 | 1093 | 1046 | 1038 |
| TTATCACAAACATTCAACCG | NSD1 | 520 | 403 | 520 | 566 | 561 |
| ACACTCACAGGGTCCTGTCG | NTRK1 | 524 | 980 | 1629 | 1595 | 1702 |
| CACTCAATAAGTGTGGACGG | NTRK1 | 440 | 854 | 1698 | 1627 | 1654 |
| CCAGATGCCCAATGCGTCGG | NTRK1 | 471 | 646 | 1291 | 1302 | 1294 |
| CCCTTTCGAGTTCAACCCCG | NTRK1 | 554 | 763 | 1177 | 1269 | 1263 |
| GCGGAGCCCACTCTTAACGA | NTRK1 | 477 | 648 | 1080 | 1037 | 1149 |
| TAGGAAGAATGTGACGTGCT | NTRK1 | 353 | 584 | 1019 | 1042 | 1068 |
| ACGCCACGATGCCCGGAGAA | NTRK2 | 402 | 864 | 1460 | 1438 | 1394 |
| GCTTCAGTGGTTCTATAACG | NTRK2 | 647 | 658 | 1219 | 1141 | 1202 |
| GGCTGCACCTAACCTCACCG | NTRK2 | 477 | 539 | 826 | 836 | 875 |
| TAGTAGCCAAGAATGAGTAT | NTRK2 | 598 | 762 | 1152 | 1246 | 1228 |
| TGGAAAACAAATCTCCTGTG | NTRK2 | 615 | 632 | 1178 | 1183 | 1242 |
| TTCCAATGATGACGGCATCG | NTRK2 | 560 | 983 | 1633 | 1398 | 1465 |
| AACCCTCAGTACTTCCGTCA | NTRK3 | 508 | 806 | 1060 | 1011 | 1030 |
| ACTCACATATAGCGCAAGTG | NTRK3 | 416 | 1104 | 1839 | 1598 | 1700 |
| ATTCTCCGCAATACATGTCA | NTRK3 | 586 | 382 | 722 | 687 | 803 |
| CTGATGTTCATACGGAACAG | NTRK3 | 599 | 809 | 1347 | 1216 | 1297 |
| GTGATCTTGGACTCCCGTAG | NTRK3 | 409 | 665 | 1158 | 1104 | 1195 |
| TGATCAACAAGTACGGTCGA | NTRK3 | 486 | 592 | 1191 | 1254 | 1272 |
| AAACAAGCAACACTTCTGTG | NUP58 | 20 | 409 | 803 | 774 | 787 |
| ACAGAAAGCACGTTACCTTG | NUP58 | 675 | 678 | 1410 | 1239 | 1263 |
| ACCTAAGGAAAGGCTTGTTG | NUP58 | 485 | 705 | 1428 | 1386 | 1450 |
| ATTACTTCTACCTCAGCTAG | NUP58 | 424 | 562 | 1028 | 901 | 937 |
| GAATGCGTTAGGATTGACTT | NUP58 | 471 | 761 | 1296 | 1359 | 1220 |
| GTAGACGTAGACGTAGTGGC | NUP58 | 428 | 341 | 603 | 660 | 655 |
| AAAAATCGAGAACGTCATAG | ONE_INTERGENIC_SITE_1 | 567 | 724 | 1140 | 1095 | 1062 |
| TTTGTCAATATAGTCGATCT | ONE_INTERGENIC_SITE_1000 | 589 | 403 | 733 | 778 | 789 |
| ACCCTCCCGCGAATATGACA | ONE_INTERGENIC_SITE_102 | 661 | 502 | 680 | 620 | 770 |
| ACCGACGGCCTAGCCGAACA | ONE_INTERGENIC_SITE_103 | 495 | 936 | 1625 | 1792 | 1667 |
| ACCGCTACACACCGCAATAG | ONE_INTERGENIC_SITE_104 | 556 | 630 | 983 | 1070 | 1034 |
| ACCTGGTTGCCAAACGTCGT | ONE_INTERGENIC_SITE_107 | 450 | 655 | 1106 | 1121 | 1148 |
| ACCTTAAACAGCATTCGGCC | ONE_INTERGENIC_SITE_108 | 498 | 330 | 528 | 493 | 447 |
| AAACGCCGGACACTACCGTT | ONE_INTERGENIC_SITE_11 | 453 | 445 | 786 | 824 | 892 |
| ACGGATGCGCCGTGACAACA | ONE_INTERGENIC_SITE_115 | 621 | 688 | 1304 | 1221 | 1206 |
| AAACGCCGTTAAGTCAAAAG | ONE_INTERGENIC_SITE_12 | 361 | 383 | 585 | 536 | 515 |
| ACTATTTGACGTTACGGACC | ONE_INTERGENIC_SITE_120 | 477 | 704 | 1269 | 1095 | 1171 |
| ACTCACTAGCAAAGCGACAG | ONE_INTERGENIC_SITE_121 | 562 | 676 | 1343 | 1192 | 1231 |
| ACTCAGGGGGCAAACGCCAT | ONE_INTERGENIC_SITE_122 | 540 | 677 | 1427 | 1280 | 1297 |
| ACTCTTAGTATATCGGATTG | ONE_INTERGENIC_SITE_123 | 568 | 1307 | 2205 | 2284 | 2164 |
| ACTGAGCATCTGATCGATCC | ONE_INTERGENIC_SITE_125 | 279 | 707 | 1259 | 1242 | 1293 |
| ACTGCGAGTTAACCCTTACA | ONE_INTERGENIC_SITE_126 | 384 | 713 | 1433 | 1530 | 1414 |
| ACTGTCAAGCTGCACGCAAT | ONE_INTERGENIC_SITE_127 | 398 | 654 | 1186 | 1117 | 1110 |
| ACTTGCCCACACGATAATCC | ONE_INTERGENIC_SITE_128 | 464 | 321 | 654 | 551 | 548 |
| AGAACTCTGCAAACGGCGCG | ONE_INTERGENIC_SITE_130 | 734 | 816 | 1251 | 1227 | 1383 |
| AGAATACAACCTACGGTAGC | ONE_INTERGENIC_SITE_131 | 490 | 611 | 915 | 978 | 999 |
| AGAATCGCTGCATGTACTCG | ONE_INTERGENIC_SITE_133 | 529 | 1065 | 1824 | 1924 | 1907 |
| AGACATAGTGCAACGTTGCA | ONE_INTERGENIC_SITE_134 | 453 | 992 | 1656 | 1626 | 1558 |
| AGACCGACGTCTAGTTAGTG | ONE_INTERGENIC_SITE_136 | 555 | 721 | 1425 | 1512 | 1514 |
| AGAGAACGATATCACAATCT | ONE_INTERGENIC_SITE_137 | 548 | 518 | 1102 | 1128 | 1012 |
| AGCAGATGTATTCACGCCAC | ONE_INTERGENIC_SITE_141 | 404 | 583 | 1115 | 1070 | 959 |
| AGCAGCATCGACCGTCACCC | ONE_INTERGENIC_SITE_142 | 321 | 503 | 985 | 936 | 968 |
| AGCAGTGCGGGCCTATACTG | ONE_INTERGENIC_SITE_143 | 535 | 936 | 1499 | 1534 | 1545 |
| AGCCTCCATTGATCGAAGCT | ONE_INTERGENIC_SITE_146 | 498 | 682 | 1105 | 1117 | 1077 |
| AGCGGGTTTGCGAACAAGCG | ONE_INTERGENIC_SITE_148 | 388 | 1042 | 1663 | 1735 | 1735 |
| AAACTACTTAGCTACCGGTA | ONE_INTERGENIC_SITE_15 | 547 | 739 | 1290 | 1272 | 1255 |
| AGCTAGATTTACGCCGGTGG | ONE_INTERGENIC_SITE_152 | 504 | 912 | 1556 | 1540 | 1616 |
| AGCTTGTGTTGCTTAACGCG | ONE_INTERGENIC_SITE_154 | 451 | 691 | 1329 | 1327 | 1338 |
| AGGAACACGCAGGCTACGAA | ONE_INTERGENIC_SITE_155 | 617 | 403 | 539 | 462 | 471 |
| AGGAACTCGGCGAGTGACGA | ONE_INTERGENIC_SITE_156 | 382 | 680 | 1184 | 1201 | 1257 |
| AGGACGCGAAGCAGCACGAT | ONE_INTERGENIC_SITE_158 | 537 | 512 | 1203 | 1067 | 1142 |
| AGGACGGTGAACGTTAGGCA | ONE_INTERGENIC_SITE_159 | 406 | 866 | 1579 | 1548 | 1584 |
| AAAGCAGGTCTACGCTCCAA | ONE_INTERGENIC_SITE_16 | 354 | 711 | 1218 | 1247 | 1179 |
| AGGATGAGGACCGTTCGGAT | ONE_INTERGENIC_SITE_161 | 567 | 525 | 1198 | 1157 | 1044 |
| AGGCGCCAAACCGACTACTA | ONE_INTERGENIC_SITE_163 | 514 | 550 | 1025 | 872 | 923 |
| AGGCTACGGTTTGACCGGTG | ONE_INTERGENIC_SITE_166 | 528 | 1009 | 1543 | 1609 | 1683 |
| AGGCTACTATCACCGAGACC | ONE_INTERGENIC_SITE_167 | 493 | 715 | 1005 | 1085 | 1018 |
| AGGCTCCATTAGAACGTACA | ONE_INTERGENIC_SITE_168 | 502 | 825 | 1267 | 1237 | 1213 |
| AGGCTGACCGTTTACGCTGA | ONE_INTERGENIC_SITE_169 | 501 | 808 | 1571 | 1464 | 1464 |
| AGGGTTGAGGGGGATCGCAT | ONE_INTERGENIC_SITE_171 | 512 | 622 | 1182 | 1242 | 1210 |
| AGGTAACGAGCTCTCGTGAC | ONE_INTERGENIC_SITE_172 | 336 | 543 | 911 | 945 | 938 |
| AGGTCATAGCCGGTCGTGTG | ONE_INTERGENIC_SITE_173 | 480 | 954 | 1708 | 1559 | 1620 |
| AGTCATGGCCCGAACGACCT | ONE_INTERGENIC_SITE_177 | 478 | 611 | 1158 | 1122 | 1073 |
| AGTCCTAACGAACGGCTCGC | ONE_INTERGENIC_SITE_179 | 599 | 547 | 1113 | 1183 | 1264 |
| AAAGTCCGTCAAGTAATCCG | ONE_INTERGENIC_SITE_18 | 637 | 548 | 1052 | 980 | 961 |
| AGTGCGCGTATAGAGTAGTA | ONE_INTERGENIC_SITE_182 | 436 | 824 | 1277 | 1345 | 1427 |
| ATAACGAGCTCCACCGTGCA | ONE_INTERGENIC_SITE_186 | 473 | 672 | 1334 | 1273 | 1315 |
| AAAGTGTTGGGACACGACAC | ONE_INTERGENIC_SITE_19 | 452 | 405 | 810 | 814 | 873 |
| ATACCCGAAACAGCCGAAAG | ONE_INTERGENIC_SITE_190 | 550 | 569 | 962 | 1044 | 1023 |
| ATACGTAAACGCCATGACTC | ONE_INTERGENIC_SITE_191 | 582 | 532 | 879 | 1054 | 1074 |
| ATAGTCGAGTCAACGAACCG | ONE_INTERGENIC_SITE_194 | 439 | 658 | 1064 | 1101 | 1027 |
| ATCACCAGTAAACCTCGTCG | ONE_INTERGENIC_SITE_199 | 554 | 537 | 994 | 804 | 818 |
| AAAACATCCAATGCCGATGT | ONE_INTERGENIC_SITE_2 | 631 | 335 | 792 | 864 | 755 |
| AAATAAACAAAGCATCGCGA | ONE_INTERGENIC_SITE_20 | 323 | 252 | 349 | 286 | 330 |
| ATCATTAAAACGCTCCGCTG | ONE_INTERGENIC_SITE_204 | 523 | 569 | 1149 | 1098 | 1028 |
| AAATAATAGCACGTCCCACG | ONE_INTERGENIC_SITE_21 | 466 | 488 | 823 | 861 | 859 |
| ATCGTCTAACCACCTCGACA | ONE_INTERGENIC_SITE_211 | 574 | 443 | 719 | 700 | 712 |
| ATCTAATCGATTGCGAGTGG | ONE_INTERGENIC_SITE_212 | 528 | 937 | 1516 | 1666 | 1647 |
| ATGAATCCTCCGTCGAATGT | ONE_INTERGENIC_SITE_214 | 333 | 584 | 1205 | 1064 | 1106 |
| ATGACCCCATCTAATTGCGG | ONE_INTERGENIC_SITE_216 | 637 | 496 | 956 | 1136 | 1045 |
| ATGCGAGGTCCAATAGGTAC | ONE_INTERGENIC_SITE_219 | 491 | 591 | 1032 | 953 | 975 |
| AAATACAGCACTCTCGCACA | ONE_INTERGENIC_SITE_22 | 484 | 427 | 770 | 643 | 672 |
| ATGCGTATTAGCTACCGGAT | ONE_INTERGENIC_SITE_223 | 373 | 523 | 957 | 980 | 998 |
| ATGGGTGAACTATACGGTAT | ONE_INTERGENIC_SITE_228 | 559 | 742 | 1377 | 1263 | 1345 |
| AAATACGCATTGTCAACCTT | ONE_INTERGENIC_SITE_23 | 688 | 723 | 1226 | 1183 | 1105 |
| ATGGTGTAGATTGTCGACTC | ONE_INTERGENIC_SITE_230 | 466 | 552 | 969 | 1052 | 935 |
| ATGGTTTAATCGGCTGAACA | ONE_INTERGENIC_SITE_232 | 588 | 780 | 1134 | 960 | 1011 |
| ATGTCGATGCTACGTACCCA | ONE_INTERGENIC_SITE_234 | 435 | 800 | 1328 | 1221 | 1213 |
| ATGTGAAGCGGGTGCGTATG | ONE_INTERGENIC_SITE_236 | 725 | 713 | 1699 | 1556 | 1678 |
| ATGTGATTATTAGACGTGCG | ONE_INTERGENIC_SITE_237 | 475 | 654 | 1233 | 1274 | 1403 |
| ATGTGTCAGACAATTGTACC | ONE_INTERGENIC_SITE_238 | 465 | 466 | 880 | 784 | 832 |
| ATGTTCGATTCAAACGAAGC | ONE_INTERGENIC_SITE_239 | 428 | 363 | 542 | 535 | 526 |
| ATGTTGCGAGTGCCCGTTAG | ONE_INTERGENIC_SITE_240 | 408 | 855 | 1437 | 1642 | 1574 |
| ATTAGCCTGGTCATTCGAGA | ONE_INTERGENIC_SITE_241 | 513 | 508 | 1044 | 844 | 938 |
| ATTCCACAGACCGACGTACT | ONE_INTERGENIC_SITE_244 | 632 | 445 | 822 | 903 | 906 |
| ATTCCCCCAGCTAACTATCG | ONE_INTERGENIC_SITE_245 | 521 | 532 | 1157 | 1142 | 1215 |
| ATTGTAGAGCACAACGCCTG | ONE_INTERGENIC_SITE_248 | 525 | 788 | 1450 | 1282 | 1272 |
| ATTGTCATTTGAATCGCTCG | ONE_INTERGENIC_SITE_249 | 576 | 711 | 1313 | 1385 | 1395 |
| AAATCACGACCCCACGCTAA | ONE_INTERGENIC_SITE_25 | 510 | 576 | 875 | 935 | 1004 |
| ATTTAAACCATGCTCCGACG | ONE_INTERGENIC_SITE_250 | 428 | 491 | 909 | 779 | 741 |
| CAAAGGTCTCGCGTTCTGAC | ONE_INTERGENIC_SITE_254 | 440 | 406 | 737 | 703 | 681 |
| CAACTCACTCGCGTCGGGCG | ONE_INTERGENIC_SITE_258 | 635 | 709 | 1023 | 808 | 898 |
| CAAGACGGTGACTATCGTCT | ONE_INTERGENIC_SITE_259 | 673 | 447 | 672 | 663 | 627 |
| AAATCATTGAGTCGAACCAT | ONE_INTERGENIC_SITE_26 | 561 | 548 | 930 | 975 | 993 |
| CAAGTTGATCACTGCGATGA | ONE_INTERGENIC_SITE_262 | 548 | 879 | 1666 | 1659 | 1623 |
| CAATAATCCACGCGATGCGG | ONE_INTERGENIC_SITE_263 | 509 | 456 | 1108 | 989 | 905 |
| CAATGGACCCCGCCTAACAC | ONE_INTERGENIC_SITE_266 | 630 | 399 | 547 | 524 | 508 |
| CACAACTTGTGCGGGTCCGT | ONE_INTERGENIC_SITE_267 | 559 | 367 | 549 | 601 | 582 |
| CACAAGTAGTGGGACGACTT | ONE_INTERGENIC_SITE_268 | 352 | 729 | 1078 | 1196 | 1196 |
| CACATGCTGGTATCGAGTCG | ONE_INTERGENIC_SITE_269 | 506 | 813 | 1436 | 1652 | 1712 |
| AAATCTGATGCACCCGCATA | ONE_INTERGENIC_SITE_27 | 546 | 554 | 1078 | 1061 | 1124 |
| CACATGTCGCCCGTAAACAC | ONE_INTERGENIC_SITE_270 | 677 | 299 | 576 | 518 | 525 |
| CACCACCTAGTAAGCGAGCG | ONE_INTERGENIC_SITE_271 | 643 | 697 | 1378 | 1149 | 1125 |
| CACCACCTTTGGCACGCCAC | ONE_INTERGENIC_SITE_272 | 501 | 576 | 918 | 879 | 928 |
| CACCCGCGTTCGTCCTACAA | ONE_INTERGENIC_SITE_273 | 569 | 258 | 462 | 395 | 434 |
| CACCCGGTATGATCCCAACG | ONE_INTERGENIC_SITE_274 | 467 | 664 | 1430 | 1309 | 1308 |
| CACCTCTCACACAATTAGCG | ONE_INTERGENIC_SITE_277 | 612 | 472 | 858 | 852 | 787 |
| CACCTGTGTCCATTCGACTG | ONE_INTERGENIC_SITE_278 | 380 | 797 | 1414 | 1236 | 1246 |
| CACGCTTCTATCGCCCGTGA | ONE_INTERGENIC_SITE_279 | 514 | 667 | 1173 | 1196 | 1201 |
| AAATGAGGGACTGGCGATTG | ONE_INTERGENIC_SITE_28 | 568 | 915 | 1646 | 1566 | 1615 |
| CACGTGCATGATCACAACGC | ONE_INTERGENIC_SITE_281 | 577 | 512 | 774 | 730 | 784 |
| CACTACATTGCGGCCGAAGT | ONE_INTERGENIC_SITE_282 | 646 | 464 | 926 | 817 | 803 |
| CACTCGCAGAAGTGCGATGA | ONE_INTERGENIC_SITE_283 | 503 | 756 | 1491 | 1366 | 1340 |
| CACTTCGGTCACTCCGCACG | ONE_INTERGENIC_SITE_286 | 730 | 702 | 1249 | 1294 | 1341 |
| CAGAACAAGCACATTGCGCC | ONE_INTERGENIC_SITE_287 | 488 | 398 | 706 | 639 | 675 |
| CAGAACCCCGCGCCTAAGCA | ONE_INTERGENIC_SITE_288 | 464 | 572 | 1293 | 1270 | 1203 |
| AAATGTAGTCAAATCGTACA | ONE_INTERGENIC_SITE_29 | 300 | 647 | 827 | 800 | 767 |
| CAGACGCATTGTGACCGGTA | ONE_INTERGENIC_SITE_290 | 406 | 591 | 1409 | 1220 | 1195 |
| CAGACGTATTCAAGCGAGGG | ONE_INTERGENIC_SITE_291 | 434 | 917 | 1361 | 1304 | 1293 |
| CAGCAACGATCACATCGTCC | ONE_INTERGENIC_SITE_293 | 323 | 330 | 471 | 477 | 513 |
| CAGCAATCAGCGATGGACGA | ONE_INTERGENIC_SITE_294 | 469 | 811 | 1239 | 1177 | 1250 |
| CAGCATATTGGGTCGAGTAG | ONE_INTERGENIC_SITE_295 | 549 | 780 | 1540 | 1476 | 1525 |
| CAGCATTGGAACCGACCGGT | ONE_INTERGENIC_SITE_296 | 538 | 441 | 883 | 881 | 896 |
| CAGCCACTGTCGGTCGTCTG | ONE_INTERGENIC_SITE_297 | 488 | 1001 | 1805 | 1789 | 1942 |
| CAGCCTAACCCGGCGTACAC | ONE_INTERGENIC_SITE_298 | 551 | 360 | 656 | 617 | 632 |
| AACACGCGTAGGATAACACG | ONE_INTERGENIC_SITE_30 | 562 | 678 | 1255 | 1154 | 1208 |
| CAGCTCTCGCGATACAACCA | ONE_INTERGENIC_SITE_302 | 622 | 681 | 1071 | 1043 | 1074 |
| CAGGCGCCATCCAATACTGT | ONE_INTERGENIC_SITE_305 | 486 | 696 | 1155 | 1070 | 1108 |
| CAGGTGTTTACAAACGCCCG | ONE_INTERGENIC_SITE_307 | 444 | 775 | 1234 | 1280 | 1186 |
| CAGTATTTGCTCGCTCCGAA | ONE_INTERGENIC_SITE_309 | 428 | 575 | 963 | 844 | 981 |
| AACCACAGTCATACGACGAT | ONE_INTERGENIC_SITE_31 | 483 | 612 | 1070 | 1029 | 1001 |
| CATACGCTTACGTATGTTCG | ONE_INTERGENIC_SITE_314 | 620 | 1296 | 1985 | 2061 | 2193 |
| CATATAGGAATGCTTCGACT | ONE_INTERGENIC_SITE_315 | 420 | 287 | 508 | 475 | 450 |
| AACCACTATACGTGGACCAA | ONE_INTERGENIC_SITE_32 | 473 | 659 | 1296 | 1319 | 1354 |
| CATCTCTACGTCCGTACCTG | ONE_INTERGENIC_SITE_320 | 504 | 795 | 1589 | 1435 | 1475 |
| CATGACAAAACCGACGAATC | ONE_INTERGENIC_SITE_321 | 633 | 489 | 737 | 708 | 775 |
| CATGTTGCATTAACCACGCG | ONE_INTERGENIC_SITE_323 | 458 | 618 | 1016 | 1017 | 1081 |
| CATTGGCTGAGAATCGCACT | ONE_INTERGENIC_SITE_325 | 292 | 566 | 906 | 881 | 908 |
| CATTTGCGCCACCGAACCAT | ONE_INTERGENIC_SITE_327 | 533 | 752 | 1057 | 1008 | 1106 |
| CCAACCCTAACGGGTTCGCT | ONE_INTERGENIC_SITE_328 | 522 | 431 | 652 | 637 | 644 |
| CCAACCGCTTTACCTAATAC | ONE_INTERGENIC_SITE_329 | 505 | 276 | 413 | 339 | 384 |
| AACCCTGTCATATTCGCGGG | ONE_INTERGENIC_SITE_33 | 508 | 608 | 1230 | 1316 | 1314 |
| CCAAGTCCGTCCGCCCAACG | ONE_INTERGENIC_SITE_333 | 576 | 805 | 1142 | 933 | 966 |
| CCAATCCTATTGCACGTGTA | ONE_INTERGENIC_SITE_334 | 460 | 750 | 1227 | 1251 | 1266 |
| CCACCGATACTAGCGGCCCA | ONE_INTERGENIC_SITE_337 | 554 | 693 | 1257 | 1275 | 1299 |
| CCACCGCACGGAATTCGGAC | ONE_INTERGENIC_SITE_338 | 576 | 370 | 603 | 611 | 606 |
| CCACGTTGGTACCACGATCA | ONE_INTERGENIC_SITE_342 | 593 | 568 | 1169 | 1008 | 1094 |
| CCAGCAACCAGTACGATATG | ONE_INTERGENIC_SITE_343 | 713 | 947 | 1625 | 1705 | 1724 |
| CCAGCTCTTAGTCGTCAATG | ONE_INTERGENIC_SITE_344 | 573 | 1039 | 1657 | 1524 | 1695 |
| CCAGGTTCTGCGCACGACCT | ONE_INTERGENIC_SITE_345 | 342 | 617 | 1196 | 1274 | 1338 |
| CCAGTGCCACGATGCGAGAA | ONE_INTERGENIC_SITE_346 | 534 | 600 | 1322 | 1425 | 1378 |
| CCATGCATGCGCAATAACGT | ONE_INTERGENIC_SITE_350 | 485 | 443 | 763 | 846 | 778 |
| CCATTTCGGAACCGGGACGT | ONE_INTERGENIC_SITE_351 | 471 | 664 | 1172 | 1224 | 1179 |
| CCCAGCTAGATTTACGCCGG | ONE_INTERGENIC_SITE_352 | 612 | 577 | 1065 | 1113 | 1097 |
| CCCATCACCATACCCGCTAG | ONE_INTERGENIC_SITE_353 | 535 | 548 | 896 | 755 | 760 |
| CCCCCATACCAAAGCCGTAT | ONE_INTERGENIC_SITE_355 | 572 | 418 | 762 | 822 | 790 |
| CCCCGTTAATAAACAGTCCG | ONE_INTERGENIC_SITE_359 | 393 | 859 | 1357 | 1168 | 1159 |
| AACCTCAGTATAAGCGTGAT | ONE_INTERGENIC_SITE_36 | 491 | 587 | 1051 | 995 | 999 |
| CCCGATGAAGCATCGCACGC | ONE_INTERGENIC_SITE_360 | 466 | 506 | 833 | 899 | 880 |
| CCCGGACGTGCCGTAAACCG | ONE_INTERGENIC_SITE_362 | 491 | 616 | 1128 | 1113 | 979 |
| CCCTAATCTGAACCGTGCCG | ONE_INTERGENIC_SITE_365 | 442 | 624 | 1165 | 1123 | 1090 |
| CCCTGGCGTGAGCCTATACG | ONE_INTERGENIC_SITE_370 | 580 | 688 | 1477 | 1495 | 1459 |
| CCGAGACCGTCGTTTCCGAG | ONE_INTERGENIC_SITE_372 | 719 | 509 | 1051 | 1047 | 1084 |
| CCGCATCGAACGAAATCCCG | ONE_INTERGENIC_SITE_374 | 572 | 414 | 646 | 528 | 586 |
| CCGGAGCGAGAACTACGCGA | ONE_INTERGENIC_SITE_379 | 590 | 640 | 935 | 802 | 839 |
| CCTATCTCACCGAATCGAGA | ONE_INTERGENIC_SITE_385 | 391 | 377 | 632 | 511 | 460 |
| CCTCAGCGCACGTCGACCCA | ONE_INTERGENIC_SITE_386 | 586 | 759 | 1400 | 1361 | 1320 |
| CCTCCTGCGCGAATAATGGC | ONE_INTERGENIC_SITE_387 | 421 | 360 | 579 | 549 | 599 |
| CCTCGGGAAGGGTACCGATC | ONE_INTERGENIC_SITE_389 | 594 | 731 | 1264 | 1343 | 1407 |
| CCTGGATGCACGACTATACT | ONE_INTERGENIC_SITE_392 | 543 | 554 | 994 | 844 | 918 |
| CCTGTAGCGCTACATGCAAT | ONE_INTERGENIC_SITE_393 | 553 | 648 | 941 | 947 | 1008 |
| CCTTAACTCGTACTGCACTG | ONE_INTERGENIC_SITE_394 | 494 | 632 | 1046 | 967 | 922 |
| CGAAGGCAAGCGACGAGCGA | ONE_INTERGENIC_SITE_403 | 291 | 1021 | 1494 | 1375 | 1378 |
| CGAAGGCCCGCGTTTATGGT | ONE_INTERGENIC_SITE_404 | 482 | 585 | 1057 | 1158 | 1196 |
| CGAAGTCCTAACTCCCCGAT | ONE_INTERGENIC_SITE_405 | 401 | 459 | 818 | 850 | 829 |
| CGACCATAGGAGTACGAGCG | ONE_INTERGENIC_SITE_407 | 513 | 764 | 1252 | 1223 | 1256 |
| CGCACACCTGCACCCGACGG | ONE_INTERGENIC_SITE_411 | 603 | 532 | 1127 | 1124 | 1153 |
| CGCATCGATATTACTCCCGG | ONE_INTERGENIC_SITE_413 | 613 | 533 | 724 | 814 | 838 |
| CGCCTCCGCCCCTATCGCGG | ONE_INTERGENIC_SITE_416 | 572 | 717 | 1363 | 1319 | 1328 |
| CGCCTCTGTTTACGGTAGCG | ONE_INTERGENIC_SITE_417 | 545 | 765 | 1550 | 1382 | 1398 |
| CGCTGATCCACACACCGCGA | ONE_INTERGENIC_SITE_420 | 535 | 392 | 739 | 700 | 712 |
| CGGCGCCCACGGTATAGTCA | ONE_INTERGENIC_SITE_423 | 360 | 637 | 1067 | 1006 | 952 |
| CGGCGGTCTTACGTAACGAG | ONE_INTERGENIC_SITE_425 | 603 | 555 | 981 | 1000 | 1058 |
| CGGGCATCTGTAGCCTACCG | ONE_INTERGENIC_SITE_427 | 564 | 602 | 1106 | 1145 | 1179 |
| CGGGCATGGCTATGCGTCAG | ONE_INTERGENIC_SITE_428 | 500 | 795 | 1303 | 1426 | 1413 |
| CGGGCGCCGACGGTTCCTTG | ONE_INTERGENIC_SITE_429 | 626 | 876 | 1579 | 1571 | 1572 |
| AAGAAGCGTTTCACGCCCCG | ONE_INTERGENIC_SITE_43 | 613 | 912 | 1304 | 1186 | 1190 |
| CGGTAAGGTACATGTGCGTC | ONE_INTERGENIC_SITE_430 | 469 | 574 | 1029 | 886 | 900 |
| CGGTCAGAAGCGAACGCCTG | ONE_INTERGENIC_SITE_432 | 588 | 753 | 1075 | 1222 | 1039 |
| CGTGGTCATTCGGGTACGCA | ONE_INTERGENIC_SITE_438 | 482 | 525 | 1297 | 1208 | 1158 |
| AAGAGATGTCTAATCGCAGG | ONE_INTERGENIC_SITE_44 | 504 | 774 | 1211 | 1183 | 1197 |
| CTAACTCACAATAACGAGTG | ONE_INTERGENIC_SITE_446 | 622 | 683 | 1290 | 1307 | 1310 |
| CTACCATAATCGATGACGGA | ONE_INTERGENIC_SITE_447 | 579 | 626 | 988 | 942 | 912 |
| CTACCCAGCTAGATTTACGC | ONE_INTERGENIC_SITE_448 | 544 | 304 | 454 | 369 | 408 |
| AAGCACCTATCTTGTATGCG | ONE_INTERGENIC_SITE_45 | 337 | 796 | 1489 | 1543 | 1544 |
| CTACTATAACATATCGCGGG | ONE_INTERGENIC_SITE_454 | 529 | 434 | 830 | 879 | 913 |
| CTAGCAGTTAGAATCGGCTA | ONE_INTERGENIC_SITE_455 | 664 | 800 | 1309 | 1366 | 1527 |
| CTAGTCTGGTCTACCATACG | ONE_INTERGENIC_SITE_458 | 485 | 856 | 1420 | 1432 | 1313 |
| CTATAAGGCACCGACCTACG | ONE_INTERGENIC_SITE_459 | 541 | 708 | 1345 | 1374 | 1481 |
| AAGCACTGTTGATCCGCGCG | ONE_INTERGENIC_SITE_46 | 415 | 626 | 992 | 1008 | 1009 |
| CTATTGACGTAACGCCACGG | ONE_INTERGENIC_SITE_461 | 390 | 604 | 973 | 864 | 808 |
| CTCAACGATTATACCGATAG | ONE_INTERGENIC_SITE_462 | 544 | 634 | 1299 | 1225 | 1188 |
| CTCAATCAAGAATGCGACAG | ONE_INTERGENIC_SITE_463 | 536 | 482 | 970 | 946 | 997 |
| CTCCCGGCATAGCATACTCC | ONE_INTERGENIC_SITE_470 | 415 | 517 | 967 | 765 | 751 |
| CTCCCTTGTAGGACGAACGC | ONE_INTERGENIC_SITE_473 | 353 | 507 | 941 | 927 | 897 |
| CTCCGACCGCAAACCAACGG | ONE_INTERGENIC_SITE_474 | 446 | 588 | 1061 | 1056 | 985 |
| CTCCGCCGAGGTCCGGACAT | ONE_INTERGENIC_SITE_475 | 586 | 646 | 1119 | 1002 | 1017 |
| AAGCCTCCGCACATCGCTTA | ONE_INTERGENIC_SITE_48 | 598 | 819 | 1498 | 1355 | 1365 |
| CTCGAGTTAGCTCTCACCGT | ONE_INTERGENIC_SITE_480 | 588 | 496 | 859 | 1004 | 926 |
| CTCGCCGCTCCCGCATATAG | ONE_INTERGENIC_SITE_481 | 581 | 657 | 1098 | 1189 | 1122 |
| CTCTTCTAGGGACTCCGTAT | ONE_INTERGENIC_SITE_484 | 442 | 554 | 1231 | 1195 | 1226 |
| CTCTTGACCTAACCTTATCG | ONE_INTERGENIC_SITE_485 | 512 | 657 | 1253 | 1342 | 1354 |
| CTGACTCCTGTAGACGTATG | ONE_INTERGENIC_SITE_487 | 468 | 701 | 1533 | 1529 | 1420 |
| CTGAGGCCCACACCGATCGA | ONE_INTERGENIC_SITE_488 | 345 | 825 | 1258 | 1117 | 1196 |
| CTGAGGCTTATAGCGCGGTG | ONE_INTERGENIC_SITE_489 | 603 | 768 | 1412 | 1472 | 1367 |
| AAGCGAATCAGTAACGGTCA | ONE_INTERGENIC_SITE_49 | 503 | 609 | 1088 | 1091 | 1189 |
| CTGCACGTATTAACCTCGAC | ONE_INTERGENIC_SITE_490 | 590 | 341 | 479 | 502 | 464 |
| CTGGCTTGCTAGACCGACAA | ONE_INTERGENIC_SITE_494 | 406 | 890 | 1324 | 1121 | 1246 |
| CTGGGGGTGACGGTCGCAAA | ONE_INTERGENIC_SITE_495 | 538 | 421 | 652 | 770 | 702 |
| CTGTACACAACGTGTCTAGC | ONE_INTERGENIC_SITE_497 | 342 | 587 | 803 | 791 | 749 |
| CTGTATAACCCGGGACCGAT | ONE_INTERGENIC_SITE_498 | 444 | 478 | 1009 | 927 | 975 |
| CTGTGCCCGGAATACCAACG | ONE_INTERGENIC_SITE_499 | 477 | 760 | 1009 | 1093 | 1054 |
| CTTAGTGACCGCGCTAACAC | ONE_INTERGENIC_SITE_500 | 502 | 272 | 502 | 389 | 418 |
| CTTAGTTCAGACTATCGCGG | ONE_INTERGENIC_SITE_501 | 361 | 383 | 639 | 547 | 556 |
| CTTCATCTTTAATGCGTGGT | ONE_INTERGENIC_SITE_502 | 427 | 609 | 1135 | 1250 | 1296 |
| CTTCCTTTCCCAATCGGCGC | ONE_INTERGENIC_SITE_504 | 540 | 506 | 840 | 866 | 966 |
| CTTCGCCAGCGGCGTACAGA | ONE_INTERGENIC_SITE_506 | 615 | 724 | 1311 | 1237 | 1244 |
| CTTCGGAAGCACACCGCACT | ONE_INTERGENIC_SITE_508 | 398 | 453 | 842 | 876 | 774 |
| CTTGACCTGTTTGTCGACTG | ONE_INTERGENIC_SITE_509 | 559 | 745 | 1297 | 1315 | 1209 |
| CTTGTTACGTTAATAAATCG | ONE_INTERGENIC_SITE_510 | 765 | 640 | 1041 | 1094 | 1116 |
| CTTTGCCGTCAAGTTAGTCG | ONE_INTERGENIC_SITE_512 | 426 | 747 | 1440 | 1298 | 1333 |
| GAAAACATTCCAAGCGAACG | ONE_INTERGENIC_SITE_513 | 524 | 468 | 822 | 785 | 740 |
| GAAAACGTGCATGCTACCGT | ONE_INTERGENIC_SITE_515 | 506 | 415 | 870 | 730 | 773 |
| GAAAATCGACACGTGCCGCC | ONE_INTERGENIC_SITE_516 | 478 | 282 | 502 | 465 | 502 |
| GAACAATACTACCACGGAAG | ONE_INTERGENIC_SITE_518 | 402 | 584 | 1135 | 1044 | 1038 |
| AAGCGTAATGGTGCAACCCG | ONE_INTERGENIC_SITE_52 | 598 | 712 | 1238 | 1180 | 1133 |
| GAACTTAGAACGGTCGTGTA | ONE_INTERGENIC_SITE_520 | 529 | 547 | 1176 | 1115 | 1048 |
| GAAGTGATCGATTTACGTTG | ONE_INTERGENIC_SITE_523 | 526 | 875 | 1381 | 1372 | 1436 |
| GAAGTTCCGACCATAAACGC | ONE_INTERGENIC_SITE_524 | 386 | 272 | 496 | 354 | 400 |
| GACACCTCAAGGGGTCGACG | ONE_INTERGENIC_SITE_531 | 515 | 471 | 899 | 852 | 913 |
| GACAGCTCCCGCGAACTATG | ONE_INTERGENIC_SITE_533 | 593 | 609 | 1084 | 992 | 990 |
| GACCACCGGACCACCCGATC | ONE_INTERGENIC_SITE_534 | 698 | 511 | 977 | 956 | 935 |
| GACTGACGATCACACCGTAA | ONE_INTERGENIC_SITE_541 | 628 | 613 | 1224 | 1241 | 1183 |
| GACTGCCCGACTAAGGCGTA | ONE_INTERGENIC_SITE_542 | 449 | 932 | 1410 | 1448 | 1343 |
| GAGAAATGATGCACTACGCT | ONE_INTERGENIC_SITE_544 | 613 | 449 | 650 | 791 | 752 |
| GAGACCACGAAAACCGGTAA | ONE_INTERGENIC_SITE_545 | 444 | 554 | 1134 | 1026 | 1068 |
| GAGACCAGTCAAGCGGCGTA | ONE_INTERGENIC_SITE_546 | 517 | 711 | 1152 | 1093 | 1073 |
| GAGAGTCCATCCGTACACCT | ONE_INTERGENIC_SITE_547 | 423 | 513 | 888 | 919 | 868 |
| GAGCCCGCTTAACCGTAGAG | ONE_INTERGENIC_SITE_548 | 530 | 742 | 1188 | 1120 | 1264 |
| AAGGCAATTCGCCCTAAGGG | ONE_INTERGENIC_SITE_55 | 546 | 530 | 1075 | 1008 | 1085 |
| GAGCGCAGGCCTAATAAGCG | ONE_INTERGENIC_SITE_550 | 497 | 772 | 1163 | 1141 | 1056 |
| GAGGAAGTAACGCGTCCGCG | ONE_INTERGENIC_SITE_552 | 533 | 601 | 1098 | 913 | 937 |
| GAGGCACCGACCGTCAACAG | ONE_INTERGENIC_SITE_553 | 523 | 548 | 1004 | 979 | 1010 |
| GAGGCACGCTCGTCATTTCG | ONE_INTERGENIC_SITE_554 | 497 | 593 | 965 | 787 | 768 |
| GAGGCCTGTAATTGGCGTAC | ONE_INTERGENIC_SITE_555 | 639 | 560 | 1169 | 1015 | 1085 |
| GAGGCGTAGACACGAGATAT | ONE_INTERGENIC_SITE_557 | 457 | 645 | 1184 | 999 | 1072 |
| GAGGTTAAGGCCGACGGAAA | ONE_INTERGENIC_SITE_558 | 621 | 510 | 941 | 936 | 963 |
| AAGGCTTGTCGTACTGATCT | ONE_INTERGENIC_SITE_56 | 565 | 600 | 1078 | 1075 | 1009 |
| GAGTTGCATTACTACGCAGT | ONE_INTERGENIC_SITE_560 | 445 | 404 | 664 | 720 | 695 |
| GATAACTGGCCAATTACGCC | ONE_INTERGENIC_SITE_561 | 467 | 362 | 726 | 624 | 667 |
| GATACTCGGAGAGTCGGTCT | ONE_INTERGENIC_SITE_563 | 549 | 646 | 1136 | 1098 | 1205 |
| GATGCCTCGGTACTCGAAGT | ONE_INTERGENIC_SITE_566 | 311 | 644 | 911 | 898 | 955 |
| GATGTGACACCCAACCCGTA | ONE_INTERGENIC_SITE_569 | 538 | 411 | 706 | 734 | 771 |
| GATTACTCGAACAGCGGTAC | ONE_INTERGENIC_SITE_571 | 448 | 588 | 1061 | 983 | 1040 |
| GATTCAACCAGACGTTAGTA | ONE_INTERGENIC_SITE_572 | 627 | 653 | 1021 | 935 | 959 |
| GCACTTGCAACGCTTACGTG | ONE_INTERGENIC_SITE_577 | 479 | 732 | 1146 | 1034 | 1024 |
| AAGTGAAACGTGGTCGTTCG | ONE_INTERGENIC_SITE_58 | 467 | 871 | 1565 | 1483 | 1507 |
| GCAGCGTGGTCAGTCGTGAA | ONE_INTERGENIC_SITE_580 | 467 | 662 | 1130 | 1194 | 1334 |
| GCAGGGTCTACTATCAAACT | ONE_INTERGENIC_SITE_581 | 706 | 383 | 771 | 845 | 789 |
| GCAGTGGGACTATTTATCGG | ONE_INTERGENIC_SITE_583 | 585 | 799 | 1203 | 1219 | 1220 |
| GCATGCTTGTATCCGCGTAA | ONE_INTERGENIC_SITE_585 | 595 | 719 | 1261 | 1101 | 1170 |
| GCATGTCATCACGTTGCGAT | ONE_INTERGENIC_SITE_586 | 417 | 386 | 720 | 658 | 724 |
| GCATTACTGGTACCCGTCAA | ONE_INTERGENIC_SITE_587 | 538 | 767 | 1158 | 1076 | 1016 |
| AAGTGGTTGTATCGTACGGA | ONE_INTERGENIC_SITE_59 | 468 | 654 | 1113 | 1139 | 1146 |
| GCCAACAGGCGGCATACGAC | ONE_INTERGENIC_SITE_590 | 399 | 284 | 522 | 517 | 476 |
| GCCGAATCGGAGCACGATAG | ONE_INTERGENIC_SITE_595 | 302 | 734 | 1428 | 1537 | 1565 |
| GCCGCTCCCGCATATAGCGG | ONE_INTERGENIC_SITE_596 | 458 | 841 | 1571 | 1622 | 1510 |
| GCCTCGGTTACGATCACGAC | ONE_INTERGENIC_SITE_597 | 616 | 276 | 566 | 586 | 602 |
| AAAAGCACGTCGTGTACACA | ONE_INTERGENIC_SITE_6 | 381 | 658 | 1072 | 1156 | 1127 |
| AAGTTTATGACGAGCACGAC | ONE_INTERGENIC_SITE_60 | 529 | 413 | 514 | 480 | 497 |
| GCTAGTTACCGGCTTACTAT | ONE_INTERGENIC_SITE_607 | 564 | 646 | 981 | 870 | 916 |
| GCTCAACGATTATACCGATA | ONE_INTERGENIC_SITE_608 | 602 | 615 | 1180 | 1137 | 1113 |
| AATAATCCCTGATTCGTGTA | ONE_INTERGENIC_SITE_61 | 341 | 839 | 1295 | 1363 | 1410 |
| GCTCCTGGGTCCTACGCGAC | ONE_INTERGENIC_SITE_610 | 513 | 293 | 583 | 494 | 508 |
| GCTCGTCTATGACGCGGCTA | ONE_INTERGENIC_SITE_614 | 410 | 750 | 1538 | 1479 | 1497 |
| GCTGAGTAGTTTGGCAATCG | ONE_INTERGENIC_SITE_616 | 590 | 530 | 1293 | 1250 | 1266 |
| GCTGATAATTTAATCCACGC | ONE_INTERGENIC_SITE_617 | 419 | 285 | 461 | 436 | 459 |
| GCTGCCCGCGCTTCAATACG | ONE_INTERGENIC_SITE_618 | 234 | 863 | 1498 | 1432 | 1432 |
| GCTGCTAATATTGCCGGTCA | ONE_INTERGENIC_SITE_619 | 410 | 714 | 1482 | 1452 | 1427 |
| AATAGCTCATGGATATCGTA | ONE_INTERGENIC_SITE_62 | 648 | 804 | 1251 | 1158 | 1134 |
| GCTGTGACAGCGCGGTTACG | ONE_INTERGENIC_SITE_620 | 453 | 775 | 1405 | 1505 | 1447 |
| GCTTAACGCGAGGGTCGACG | ONE_INTERGENIC_SITE_621 | 634 | 731 | 1078 | 1119 | 1137 |
| GCTTACATGTCACCCGTCCA | ONE_INTERGENIC_SITE_622 | 558 | 750 | 1208 | 1068 | 1091 |
| GCTTCGCCCTGACTATACCG | ONE_INTERGENIC_SITE_623 | 396 | 588 | 1231 | 1027 | 1022 |
| GCTTGGTAGCTTGAATCGCG | ONE_INTERGENIC_SITE_626 | 621 | 1030 | 1596 | 1419 | 1490 |
| GGACCCAGAATCGTCTGACT | ONE_INTERGENIC_SITE_627 | 542 | 540 | 1007 | 1073 | 921 |
| GGACGGACGGAAAGCCGATA | ONE_INTERGENIC_SITE_632 | 519 | 1027 | 1680 | 1793 | 1877 |
| GGACTGACCGTTTCTCGACC | ONE_INTERGENIC_SITE_635 | 521 | 648 | 1160 | 1175 | 1121 |
| GGAGACCAGTCAAGCGGCGT | ONE_INTERGENIC_SITE_636 | 593 | 380 | 779 | 664 | 624 |
| GGAGAGATGCTACTAACGGG | ONE_INTERGENIC_SITE_637 | 617 | 706 | 1151 | 1208 | 1232 |
| GGAGCACAGTGCCGAACGAC | ONE_INTERGENIC_SITE_638 | 468 | 516 | 655 | 640 | 678 |
| GGAGCCGGGGCATTTGACGC | ONE_INTERGENIC_SITE_639 | 386 | 763 | 1125 | 1149 | 1205 |
| GGAGCGGACTGGTGCCCGAT | ONE_INTERGENIC_SITE_640 | 606 | 680 | 1372 | 1346 | 1339 |
| GGAGGTCGGCTAACTACAAG | ONE_INTERGENIC_SITE_641 | 506 | 616 | 1101 | 1186 | 1106 |
| GGAGTCAAGGGTATCGTGGG | ONE_INTERGENIC_SITE_642 | 521 | 925 | 1607 | 1850 | 1714 |
| GGAGTGTCGACCCCCTGATA | ONE_INTERGENIC_SITE_645 | 583 | 923 | 1497 | 1522 | 1512 |
| GGATATAGGGCACCTCGTCA | ONE_INTERGENIC_SITE_647 | 369 | 618 | 974 | 839 | 815 |
| GGATCTAGCACAATTCGGAG | ONE_INTERGENIC_SITE_648 | 570 | 631 | 1179 | 1083 | 1120 |
| GGATTAGTTTCACGCGTAGA | ONE_INTERGENIC_SITE_650 | 501 | 946 | 1554 | 1723 | 1862 |
| GGCACAAGGTGACGCGACAC | ONE_INTERGENIC_SITE_651 | 459 | 375 | 813 | 721 | 745 |
| GGCACCATGTCGTAGCGTTA | ONE_INTERGENIC_SITE_652 | 336 | 764 | 1334 | 1470 | 1618 |
| GGCAGATCGCGGTGCGATCG | ONE_INTERGENIC_SITE_653 | 403 | 907 | 1826 | 1832 | 1746 |
| GGCATTTGGCCAACACGTCG | ONE_INTERGENIC_SITE_655 | 492 | 567 | 1009 | 955 | 939 |
| GGCCAACGAATCCCTATCGC | ONE_INTERGENIC_SITE_656 | 528 | 301 | 554 | 641 | 618 |
| GGCGCAAACCATTGCGCCGA | ONE_INTERGENIC_SITE_659 | 369 | 682 | 1114 | 994 | 1061 |
| GGCTAAGGCTACGGTTTGAC | ONE_INTERGENIC_SITE_661 | 469 | 500 | 830 | 883 | 913 |
| GGCTCCAGTGCATACAACCG | ONE_INTERGENIC_SITE_662 | 479 | 800 | 1302 | 1252 | 1292 |
| GGGAATTCTTATCCGCGTGT | ONE_INTERGENIC_SITE_666 | 575 | 513 | 1101 | 892 | 1002 |
| GGGCCAACGAATCCCTATCG | ONE_INTERGENIC_SITE_669 | 532 | 833 | 1444 | 1445 | 1414 |
| GGGCCAGACGTATTCAAGCG | ONE_INTERGENIC_SITE_670 | 547 | 585 | 987 | 980 | 965 |
| GGGCGCCTCGATCCGCCCAA | ONE_INTERGENIC_SITE_671 | 622 | 560 | 1013 | 1067 | 970 |
| GGGCGCTCGAATATCCTCGG | ONE_INTERGENIC_SITE_672 | 393 | 659 | 898 | 1120 | 1082 |
| GGGCGGCCTCACGTTATCAA | ONE_INTERGENIC_SITE_673 | 476 | 679 | 1206 | 1120 | 1154 |
| GGGGAATAGGTCCGTCCGAA | ONE_INTERGENIC_SITE_675 | 560 | 761 | 1146 | 1134 | 1103 |
| GGGGCGCGTAACATGAGTGT | ONE_INTERGENIC_SITE_677 | 356 | 763 | 1277 | 1406 | 1365 |
| GGGGCGCTCGTTCAAAGAGT | ONE_INTERGENIC_SITE_678 | 646 | 685 | 1387 | 1360 | 1479 |
| GGGGGAACTTGACTCGAGGT | ONE_INTERGENIC_SITE_679 | 440 | 531 | 750 | 768 | 795 |
| GGGGGAGCGTCTGTCAATTG | ONE_INTERGENIC_SITE_680 | 435 | 713 | 1412 | 1503 | 1565 |
| GGGGTCAGTATAAACCGCCG | ONE_INTERGENIC_SITE_681 | 538 | 472 | 953 | 863 | 910 |
| GGGGTCTAACCCAATTCGTA | ONE_INTERGENIC_SITE_682 | 570 | 768 | 1330 | 1260 | 1294 |
| GGGTAATGGATCTGCGGACC | ONE_INTERGENIC_SITE_683 | 499 | 636 | 877 | 989 | 898 |
| GGGTCCGCGTGACGCTTCGG | ONE_INTERGENIC_SITE_686 | 400 | 686 | 1495 | 1382 | 1473 |
| GGGTGCCTCGTGGTTAGTCG | ONE_INTERGENIC_SITE_688 | 611 | 1030 | 1698 | 1616 | 1647 |
| GGGTGTCTCTCGATGACGGG | ONE_INTERGENIC_SITE_690 | 554 | 736 | 1280 | 1272 | 1356 |
| GGGTTGGTGGGCACTCGTCG | ONE_INTERGENIC_SITE_691 | 579 | 513 | 1014 | 1013 | 1027 |
| GGTACTTAAGGACACGCGAG | ONE_INTERGENIC_SITE_694 | 582 | 631 | 1210 | 1245 | 1183 |
| GGTCAGTAATACGCCCCGCT | ONE_INTERGENIC_SITE_696 | 666 | 326 | 554 | 523 | 532 |
| GGTGAGCAGCGTCACGCAAG | ONE_INTERGENIC_SITE_699 | 579 | 739 | 1481 | 1410 | 1514 |
| AAACATCAGGTGCGCACGAA | ONE_INTERGENIC_SITE_7 | 438 | 895 | 1126 | 1196 | 1137 |
| AATCGGTCCATGCCATAAGT | ONE_INTERGENIC_SITE_70 | 498 | 679 | 948 | 827 | 862 |
| GGTGTTCAAAGATGCGTCGG | ONE_INTERGENIC_SITE_701 | 477 | 704 | 1276 | 1286 | 1215 |
| GGTTAGCAGATCGTTGCACA | ONE_INTERGENIC_SITE_702 | 519 | 574 | 1388 | 1404 | 1362 |
| GGTTGCCACTCGAACGGGAA | ONE_INTERGENIC_SITE_704 | 547 | 666 | 1272 | 1516 | 1414 |
| GGTTGGTTGTCATTCGCTAG | ONE_INTERGENIC_SITE_705 | 447 | 698 | 1272 | 1240 | 1242 |
| GTAACGATGGACCTTGCCGG | ONE_INTERGENIC_SITE_706 | 534 | 505 | 883 | 998 | 922 |
| GTAACTAAGGTCACCGCAAG | ONE_INTERGENIC_SITE_707 | 573 | 509 | 1095 | 1000 | 911 |
| GTAATATTCTAGACTACGCG | ONE_INTERGENIC_SITE_709 | 683 | 580 | 796 | 670 | 739 |
| GTACGTTTCACACCAAACGT | ONE_INTERGENIC_SITE_712 | 559 | 356 | 662 | 610 | 589 |
| GTAGAATTGGCGGATCGTAA | ONE_INTERGENIC_SITE_713 | 504 | 755 | 893 | 874 | 922 |
| GTATCGCGGTGCGAATAGTA | ONE_INTERGENIC_SITE_717 | 541 | 507 | 1004 | 935 | 988 |
| GTCAATGGTGACGGTCGATG | ONE_INTERGENIC_SITE_719 | 492 | 781 | 1834 | 1704 | 1726 |
| AATGCACTAGAACTCTCGAC | ONE_INTERGENIC_SITE_72 | 394 | 193 | 384 | 381 | 405 |
| GTCACTATCGCCAATAGTGG | ONE_INTERGENIC_SITE_720 | 553 | 570 | 1033 | 959 | 877 |
| GTCAGAAGCTAAATCGCTCG | ONE_INTERGENIC_SITE_721 | 563 | 871 | 1379 | 1312 | 1374 |
| GTCAGCGCGTCGGGTTACAC | ONE_INTERGENIC_SITE_722 | 383 | 416 | 661 | 575 | 647 |
| GTCCTCACGACTACCGTGCG | ONE_INTERGENIC_SITE_724 | 545 | 727 | 1242 | 1162 | 1243 |
| GTCTACAACACGAGTCGTGC | ONE_INTERGENIC_SITE_727 | 582 | 440 | 548 | 571 | 671 |
| GTCTTCAGCAGGCGTCGACC | ONE_INTERGENIC_SITE_730 | 676 | 499 | 906 | 757 | 788 |
| GTGAAACAGTGCAGCGCGAT | ONE_INTERGENIC_SITE_731 | 552 | 424 | 688 | 697 | 701 |
| GTGAATCCTGTACTCGAGGT | ONE_INTERGENIC_SITE_732 | 419 | 476 | 804 | 924 | 909 |
| GTGACCTTGTTACACGAGCT | ONE_INTERGENIC_SITE_733 | 392 | 393 | 907 | 879 | 932 |
| GTGACTCAAGCTGTCGGTAA | ONE_INTERGENIC_SITE_734 | 354 | 760 | 1409 | 1349 | 1343 |
| GTGAGCGCACCGATTTGCCT | ONE_INTERGENIC_SITE_735 | 688 | 389 | 717 | 619 | 672 |
| GTGCAATAATCCACGCGATG | ONE_INTERGENIC_SITE_736 | 597 | 816 | 1169 | 1012 | 1073 |
| GTGCCAGCATAACGTCACTA | ONE_INTERGENIC_SITE_737 | 439 | 783 | 1429 | 1406 | 1489 |
| GTGCCTACATAACACGTGCT | ONE_INTERGENIC_SITE_739 | 556 | 642 | 991 | 1057 | 1044 |
| AATGCTGTCTTATGCGACTA | ONE_INTERGENIC_SITE_74 | 400 | 605 | 1077 | 1078 | 1024 |
| GTGCGTGTAACCACTCGTTA | ONE_INTERGENIC_SITE_740 | 479 | 784 | 1281 | 1374 | 1347 |
| GTGGCGTATTTGACTCCACG | ONE_INTERGENIC_SITE_742 | 480 | 824 | 1258 | 1417 | 1348 |
| GTGGGCCCATTAACCCACGT | ONE_INTERGENIC_SITE_743 | 541 | 311 | 680 | 631 | 566 |
| GTGTGAACAAGCTTACGGGT | ONE_INTERGENIC_SITE_746 | 618 | 425 | 720 | 581 | 582 |
| GTTACTCAGAACGTCCGCAT | ONE_INTERGENIC_SITE_747 | 579 | 414 | 611 | 588 | 556 |
| GTTTAAAGCTCGACACGCAT | ONE_INTERGENIC_SITE_749 | 424 | 318 | 436 | 410 | 404 |
| GTTTACATGAAACGCACGAG | ONE_INTERGENIC_SITE_750 | 476 | 610 | 1063 | 1124 | 1101 |
| GTTTCAAACAGGGTCGATCC | ONE_INTERGENIC_SITE_751 | 571 | 475 | 705 | 684 | 680 |
| TAAAACGAACCGGTATGACT | ONE_INTERGENIC_SITE_753 | 501 | 492 | 740 | 639 | 595 |
| TAAACTGTGATATGCGATCG | ONE_INTERGENIC_SITE_755 | 620 | 613 | 1207 | 1243 | 1242 |
| TAAAGTCGGTCGATTAGTTG | ONE_INTERGENIC_SITE_756 | 529 | 1038 | 1557 | 1581 | 1643 |
| TAAATGTATGGACCGCGCCA | ONE_INTERGENIC_SITE_757 | 689 | 697 | 1279 | 1251 | 1292 |
| AATTTCTGGTTAATCGATGT | ONE_INTERGENIC_SITE_76 | 552 | 268 | 543 | 549 | 587 |
| TAAGAGAACTCGGCGATATG | ONE_INTERGENIC_SITE_760 | 412 | 661 | 1472 | 1293 | 1326 |
| TAAGGAAGCCCTGTCGACAT | ONE_INTERGENIC_SITE_762 | 551 | 442 | 866 | 903 | 860 |
| TAAGTAGATAACGCCAACGC | ONE_INTERGENIC_SITE_763 | 453 | 436 | 739 | 731 | 667 |
| TAAGTTCTATCGGTTATCCC | ONE_INTERGENIC_SITE_765 | 397 | 453 | 770 | 887 | 994 |
| TAATGCAACCGTAACACGGC | ONE_INTERGENIC_SITE_767 | 530 | 324 | 622 | 525 | 552 |
| TAATGCTTCGAGCGCGTCTG | ONE_INTERGENIC_SITE_768 | 532 | 763 | 1349 | 1266 | 1367 |
| TACAAGCTGTTAACAACGCG | ONE_INTERGENIC_SITE_771 | 507 | 784 | 1101 | 1169 | 1155 |
| TACACTGCGGTACGGCATAG | ONE_INTERGENIC_SITE_772 | 531 | 883 | 1630 | 1409 | 1455 |
| TACAGGACCCCAATCTACAT | ONE_INTERGENIC_SITE_774 | 585 | 370 | 461 | 394 | 452 |
| ACAACAAGTCTACGTACCAC | ONE_INTERGENIC_SITE_78 | 588 | 415 | 612 | 662 | 689 |
| TACTAAAAGTAATGCGCCAT | ONE_INTERGENIC_SITE_782 | 502 | 684 | 989 | 1034 | 1071 |
| TAGACCCGGTCACTCAATCA | ONE_INTERGENIC_SITE_786 | 597 | 460 | 1017 | 1066 | 935 |
| TAGAGTACCCCCGATGCGAA | ONE_INTERGENIC_SITE_787 | 471 | 535 | 892 | 803 | 861 |
| TAGATCTGGCCGTATCGACT | ONE_INTERGENIC_SITE_788 | 583 | 525 | 825 | 843 | 847 |
| ACACAAAATGCGTCGAGCGT | ONE_INTERGENIC_SITE_79 | 322 | 665 | 1006 | 1117 | 1119 |
| TAGGGTACACGACCTGACTA | ONE_INTERGENIC_SITE_794 | 398 | 616 | 938 | 966 | 969 |
| TAGGGTAGGTAACACGATCA | ONE_INTERGENIC_SITE_795 | 514 | 570 | 1218 | 1069 | 1146 |
| TAGGTATCGCTAGCTATCCT | ONE_INTERGENIC_SITE_796 | 573 | 540 | 1027 | 969 | 1063 |
| TAGTCCAACAGAGCGATCGG | ONE_INTERGENIC_SITE_798 | 391 | 686 | 1348 | 1400 | 1325 |
| TAGTCTACCATAATCGATGA | ONE_INTERGENIC_SITE_799 | 560 | 467 | 832 | 758 | 731 |
| AAACCAAGTGCCGCGCATGT | ONE_INTERGENIC_SITE_8 | 501 | 603 | 1086 | 1012 | 1008 |
| ACACACACCGCGGTAGACTA | ONE_INTERGENIC_SITE_80 | 516 | 796 | 1091 | 964 | 1050 |
| TATAATCTACATACCGAGGC | ONE_INTERGENIC_SITE_801 | 405 | 180 | 347 | 274 | 347 |
| TATCTCCGTAGAATCGGTAG | ONE_INTERGENIC_SITE_804 | 654 | 749 | 1349 | 1400 | 1414 |
| TATGAATCCTCCGTCGAATG | ONE_INTERGENIC_SITE_805 | 576 | 579 | 1025 | 1043 | 1211 |
| TATGCACAAGAGACCGTTAA | ONE_INTERGENIC_SITE_806 | 488 | 661 | 1106 | 1257 | 1340 |
| TATGGTCGTACTATCTCTTG | ONE_INTERGENIC_SITE_808 | 534 | 748 | 1213 | 1233 | 1238 |
| TATGTGTCAATACTTAACGG | ONE_INTERGENIC_SITE_809 | 628 | 627 | 1055 | 1036 | 1005 |
| ACACAGACCATTTGTCGCGT | ONE_INTERGENIC_SITE_81 | 506 | 593 | 1302 | 1444 | 1402 |
| TATTTAGTTACACGCGAGGT | ONE_INTERGENIC_SITE_810 | 447 | 661 | 1031 | 1053 | 1121 |
| TATTTATATGCATACGGGAC | ONE_INTERGENIC_SITE_811 | 491 | 377 | 677 | 580 | 635 |
| TCACCACTTATTAACGATAG | ONE_INTERGENIC_SITE_813 | 324 | 610 | 960 | 1065 | 1079 |
| TCACCGACTCGAAGGGTTGT | ONE_INTERGENIC_SITE_814 | 551 | 801 | 1491 | 1584 | 1612 |
| TCACGTGGATCTTATTACGT | ONE_INTERGENIC_SITE_817 | 509 | 558 | 841 | 908 | 880 |
| TCAGAGGAGTCCATAACGAC | ONE_INTERGENIC_SITE_818 | 473 | 269 | 565 | 558 | 493 |
| TCAGCCAAATCGTGAATAGG | ONE_INTERGENIC_SITE_819 | 641 | 717 | 1051 | 1011 | 997 |
| TCAGGTCATATGGCGACTAG | ONE_INTERGENIC_SITE_820 | 354 | 1017 | 1620 | 1573 | 1572 |
| TCCACAGCGAGGTTAAACCG | ONE_INTERGENIC_SITE_821 | 460 | 549 | 1096 | 1060 | 1095 |
| TCCACCCAGTACGATCAACC | ONE_INTERGENIC_SITE_822 | 424 | 283 | 545 | 426 | 394 |
| TCCCAAACAAGCGAGTCGCG | ONE_INTERGENIC_SITE_824 | 715 | 488 | 946 | 919 | 935 |
| TCCCGATGAAGCATCGCACG | ONE_INTERGENIC_SITE_825 | 548 | 524 | 1114 | 1103 | 1123 |
| TCCCTCAGGCAAGACGACTA | ONE_INTERGENIC_SITE_826 | 679 | 486 | 895 | 793 | 814 |
| TCCGACCACGTATTGAAGCG | ONE_INTERGENIC_SITE_828 | 367 | 751 | 1375 | 1341 | 1320 |
| TCCTCAAAGTCTCTCGAACG | ONE_INTERGENIC_SITE_834 | 400 | 552 | 820 | 951 | 975 |
| TCCTCGGGTACCCGACGACA | ONE_INTERGENIC_SITE_835 | 610 | 567 | 978 | 907 | 913 |
| TCCTTGGGGTACGCTACTTG | ONE_INTERGENIC_SITE_838 | 430 | 729 | 1408 | 1511 | 1440 |
| ACAGACAACTCACTCGCGTC | ONE_INTERGENIC_SITE_84 | 477 | 511 | 920 | 860 | 888 |
| TCGTACTCCTATGGTCGGAG | ONE_INTERGENIC_SITE_846 | 621 | 621 | 1214 | 1128 | 1128 |
| ACAGCAAACTAGTCTCGGTT | ONE_INTERGENIC_SITE_85 | 652 | 409 | 986 | 975 | 943 |
| TCTGACGTCTGATAGTCGGA | ONE_INTERGENIC_SITE_850 | 359 | 743 | 1331 | 1263 | 1321 |
| TCTGGGTAATCATCGCATGC | ONE_INTERGENIC_SITE_852 | 603 | 618 | 876 | 973 | 930 |
| TCTGTATTCACGACACACTC | ONE_INTERGENIC_SITE_853 | 379 | 314 | 441 | 356 | 369 |
| TCTGTTTAATGCCGCATGAG | ONE_INTERGENIC_SITE_854 | 427 | 695 | 1466 | 1518 | 1482 |
| TCTTCTAAAATATCCGTACG | ONE_INTERGENIC_SITE_856 | 524 | 575 | 1116 | 940 | 1080 |
| TCTTGCTAAACTGACGCGGT | ONE_INTERGENIC_SITE_858 | 507 | 489 | 1036 | 1076 | 1021 |
| TCTTGGGTCAAAGTTCGGCG | ONE_INTERGENIC_SITE_859 | 428 | 617 | 1014 | 1061 | 991 |
| ACAGCATGAAACGCGCAGAT | ONE_INTERGENIC_SITE_86 | 515 | 484 | 937 | 955 | 932 |
| TGAACATCTTCCGTACGGAG | ONE_INTERGENIC_SITE_861 | 614 | 552 | 1129 | 1105 | 1047 |
| TGAACGTCGCTTACAATCCA | ONE_INTERGENIC_SITE_862 | 556 | 687 | 1379 | 1244 | 1243 |
| ACAGCCTGCAAACCCGTATA | ONE_INTERGENIC_SITE_87 | 595 | 642 | 1145 | 1158 | 1176 |
| TGAGTATGCACTTGACCGAT | ONE_INTERGENIC_SITE_871 | 406 | 487 | 943 | 959 | 915 |
| TGAGTTCTGTCGAGCACCCG | ONE_INTERGENIC_SITE_874 | 308 | 783 | 1264 | 1410 | 1363 |
| ACAGCTCTCGCGATACAACC | ONE_INTERGENIC_SITE_88 | 482 | 425 | 720 | 681 | 660 |
| TGCACACGGTTAACCTGCGT | ONE_INTERGENIC_SITE_880 | 378 | 433 | 851 | 752 | 867 |
| TGCCTAATACAGCTCCGTCG | ONE_INTERGENIC_SITE_887 | 510 | 776 | 1331 | 1170 | 1280 |
| TGCGAGTCGTGCGCTTACAC | ONE_INTERGENIC_SITE_888 | 528 | 685 | 1218 | 1029 | 1000 |
| ACAGCTTACAGCCGCGTCAC | ONE_INTERGENIC_SITE_89 | 586 | 431 | 919 | 966 | 915 |
| TGCTAGCTCAAAGTCGCCAG | ONE_INTERGENIC_SITE_893 | 380 | 651 | 1185 | 1239 | 1243 |
| TGCTCAACGGTGCTAACGAC | ONE_INTERGENIC_SITE_894 | 424 | 384 | 700 | 692 | 712 |
| TGCTGTGCCAGGTCGACACG | ONE_INTERGENIC_SITE_896 | 402 | 917 | 1665 | 1714 | 1772 |
| TGGAGTGATTCGCGCTCACA | ONE_INTERGENIC_SITE_898 | 546 | 952 | 1592 | 1793 | 1797 |
| TGGCGCAAACCATTGCGCCG | ONE_INTERGENIC_SITE_900 | 297 | 212 | 469 | 473 | 520 |
| TGGCTTTAGTCTCACGACGG | ONE_INTERGENIC_SITE_905 | 549 | 844 | 1482 | 1467 | 1556 |
| TGGGAAATATAGTCGAACGG | ONE_INTERGENIC_SITE_906 | 473 | 890 | 1415 | 1552 | 1637 |
| TGGGTCCACACATGCGACTC | ONE_INTERGENIC_SITE_907 | 503 | 495 | 930 | 809 | 853 |
| TGTATTGCTATCGTAGCACG | ONE_INTERGENIC_SITE_915 | 563 | 922 | 1643 | 1574 | 1678 |
| TGTCCTGGGTCGGTTAATCA | ONE_INTERGENIC_SITE_918 | 424 | 921 | 1534 | 1443 | 1506 |
| TGTCGAGGTGGTTAGACGAT | ONE_INTERGENIC_SITE_919 | 470 | 481 | 852 | 915 | 832 |
| TGTGACCACTAACGTGACGG | ONE_INTERGENIC_SITE_922 | 598 | 549 | 1250 | 1290 | 1256 |
| TGTGATATGCGATCGCGGTA | ONE_INTERGENIC_SITE_923 | 329 | 719 | 1321 | 1338 | 1310 |
| TGTGTGACGTAGGTTAGTTG | ONE_INTERGENIC_SITE_927 | 310 | 1059 | 1614 | 1607 | 1490 |
| TGTGTGTTTAACTCGTATGG | ONE_INTERGENIC_SITE_928 | 581 | 847 | 1692 | 1664 | 1632 |
| TGTGTTAACTCGGGTCGCGG | ONE_INTERGENIC_SITE_929 | 373 | 615 | 1208 | 1145 | 1235 |
| TGTTCCAGCTATGCCGCAAG | ONE_INTERGENIC_SITE_930 | 571 | 611 | 1423 | 1528 | 1523 |
| TGTTGCAGACACGCCACTAC | ONE_INTERGENIC_SITE_931 | 234 | 363 | 727 | 688 | 684 |
| TGTTTGCAATGTACACCCGC | ONE_INTERGENIC_SITE_934 | 548 | 308 | 572 | 610 | 582 |
| TGTTTGGCCGTACTTTAGCG | ONE_INTERGENIC_SITE_935 | 495 | 814 | 1407 | 1470 | 1417 |
| TTACGATCACGACCGGATGG | ONE_INTERGENIC_SITE_937 | 391 | 653 | 1038 | 1008 | 1057 |
| TTACGCGAAGATCTAACGAA | ONE_INTERGENIC_SITE_938 | 623 | 488 | 820 | 807 | 729 |
| ACCAGATACTCCCAACGTTA | ONE_INTERGENIC_SITE_94 | 417 | 452 | 665 | 626 | 690 |
| TTACTGCATTCACGTACTAG | ONE_INTERGENIC_SITE_940 | 387 | 655 | 1048 | 1153 | 979 |
| TTAGACGCATTACGTACGTG | ONE_INTERGENIC_SITE_941 | 389 | 771 | 1380 | 1411 | 1457 |
| TTAGGACAGCCCTATTCGAT | ONE_INTERGENIC_SITE_942 | 508 | 605 | 907 | 979 | 990 |
| TTAGGCGTTTCAGACGACTT | ONE_INTERGENIC_SITE_943 | 608 | 507 | 914 | 888 | 914 |
| TTAGTTGGGTGATCGTATGT | ONE_INTERGENIC_SITE_945 | 510 | 858 | 1390 | 1422 | 1576 |
| TTATACACGGGCTCGACGGC | ONE_INTERGENIC_SITE_946 | 321 | 499 | 973 | 1007 | 1061 |
| TTATACTAGTACCCTCAGTC | ONE_INTERGENIC_SITE_948 | 414 | 412 | 671 | 798 | 736 |
| TTATCGAATTTATCGTATGC | ONE_INTERGENIC_SITE_951 | 571 | 472 | 741 | 774 | 785 |
| TTATGACCGACTTGAATCGA | ONE_INTERGENIC_SITE_952 | 621 | 685 | 1218 | 1101 | 1029 |
| TTCCAAATACGACGCGCAGT | ONE_INTERGENIC_SITE_956 | 702 | 523 | 1008 | 995 | 1016 |
| TTCCAACCATCCAACGACCT | ONE_INTERGENIC_SITE_957 | 555 | 393 | 726 | 779 | 743 |
| TTCCCGCCGACGAGTACTAA | ONE_INTERGENIC_SITE_958 | 419 | 685 | 1039 | 1098 | 1046 |
| ACCATCATACCACGCGCCCA | ONE_INTERGENIC_SITE_96 | 474 | 589 | 985 | 1096 | 1031 |
| TTCCGTAACACCGTATAGCA | ONE_INTERGENIC_SITE_960 | 405 | 504 | 957 | 1008 | 936 |
| TTCCTGAGGACATGCGAAGC | ONE_INTERGENIC_SITE_963 | 460 | 491 | 919 | 801 | 856 |
| TTCGCATCATTCGCCCAACC | ONE_INTERGENIC_SITE_965 | 495 | 325 | 745 | 659 | 641 |
| TTCTAAACAGTACCCGCTGT | ONE_INTERGENIC_SITE_970 | 417 | 384 | 831 | 753 | 770 |
| TTCTTCGTGTCCCGTAAACG | ONE_INTERGENIC_SITE_973 | 405 | 625 | 1130 | 1112 | 1146 |
| TTGAGCCTGCGTGTACCGAA | ONE_INTERGENIC_SITE_974 | 536 | 666 | 1316 | 1214 | 1205 |
| TTGAGGCCTGCACCTACGCT | ONE_INTERGENIC_SITE_975 | 595 | 588 | 1064 | 983 | 971 |
| TTGATACTAAACTGCGCTCG | ONE_INTERGENIC_SITE_976 | 381 | 803 | 1526 | 1746 | 1619 |
| TTGGGTAACTAACTCGTCGA | ONE_INTERGENIC_SITE_978 | 458 | 584 | 1223 | 1126 | 1174 |
| TTGGTGAACGAACGTCGTGC | ONE_INTERGENIC_SITE_980 | 331 | 568 | 868 | 877 | 835 |
| TTGGTGGATGAACCCGTCAT | ONE_INTERGENIC_SITE_981 | 378 | 715 | 1113 | 1208 | 1263 |
| TTGGTTAGGCTACATCCAAC | ONE_INTERGENIC_SITE_982 | 496 | 481 | 856 | 843 | 879 |
| TTTACAAACTCCTACGGCAA | ONE_INTERGENIC_SITE_988 | 306 | 416 | 841 | 776 | 796 |
| ACCCGGTCCTAATACTAAGG | ONE_INTERGENIC_SITE_99 | 642 | 680 | 1242 | 1398 | 1359 |
| TTTAGACAGAGTTTCGCTCG | ONE_INTERGENIC_SITE_990 | 557 | 861 | 1602 | 1504 | 1510 |
| TTTAGGCAGTCTAAGTCGCG | ONE_INTERGENIC_SITE_991 | 368 | 597 | 905 | 1041 | 939 |
| TTTAGTTACACGCGAGGTCG | ONE_INTERGENIC_SITE_992 | 556 | 889 | 1556 | 1628 | 1565 |
| TTTCAGGTAAGGTCCGATCC | ONE_INTERGENIC_SITE_993 | 429 | 574 | 956 | 942 | 1145 |
| TTTCCGCTCACATTCGCATG | ONE_INTERGENIC_SITE_994 | 645 | 762 | 1171 | 1104 | 1228 |
| TTTCCGTATAAAGTTACGAG | ONE_INTERGENIC_SITE_996 | 369 | 532 | 633 | 711 | 739 |
| TTTGCACAATCAGGCAACGT | ONE_INTERGENIC_SITE_997 | 427 | 390 | 747 | 766 | 781 |
| TTTGCCTATATAACCCTTCG | ONE_INTERGENIC_SITE_998 | 500 | 779 | 1196 | 1249 | 1200 |
| TTTGCTATGTGTACTACGTG | ONE_INTERGENIC_SITE_999 | 459 | 1061 | 1490 | 1670 | 1626 |
| AGGGTACAAGGACTCAGTGG | P2RX7 | 383 | 701 | 1399 | 1131 | 1263 |
| TATTCACAGCACTTGAAACA | P2RX7 | 495 | 438 | 680 | 714 | 749 |
| TCGCTTTAATCAGTGACAAG | P2RX7 | 574 | 477 | 943 | 962 | 915 |
| TGAGAAAGTTTGTCATCACG | P2RX7 | 382 | 861 | 1596 | 1589 | 1694 |
| TGGATGGACCCACAAAGCAA | P2RX7 | 344 | 630 | 1099 | 950 | 1025 |
| TGGGTTCTGAGTCTTGTGAA | P2RX7 | 312 | 1044 | 1420 | 1490 | 1486 |
| AAAATCCAAGATGACAGACG | PAK3 | 295 | 401 | 826 | 813 | 777 |
| AAACAGTACCTGATGCCCTG | PAK3 | 549 | 721 | 1163 | 1161 | 1130 |
| AAATACATGAGCTTTACATC | PAK3 | 458 | 289 | 564 | 548 | 598 |
| GGTCGCTTCAAGTCAATCAG | PAK3 | 546 | 686 | 1294 | 1226 | 1259 |
| TGACGAACTATGGGTAGTCA | PAK3 | 385 | 577 | 1224 | 1296 | 1356 |
| TGGGAAGATAGAGCGAAGCC | PAK3 | 376 | 755 | 1051 | 953 | 912 |
| AACTTCATCAAGATCGGCGA | PAK4 | 531 | 357 | 590 | 506 | 548 |
| AGCACGAGCAGAAGTTCACG | PAK4 | 521 | 512 | 1258 | 1165 | 1273 |
| AGCGGTGGGTCATACCTTGG | PAK4 | 582 | 834 | 1841 | 1881 | 1951 |
| GCCCTTTAACACGTACCCGA | PAK4 | 563 | 337 | 491 | 455 | 426 |
| GGACTACCAGCACGAGAACG | PAK4 | 388 | 773 | 1445 | 1274 | 1289 |
| TGCTCGTGCGACACTCGCTG | PAK4 | 604 | 790 | 1504 | 1475 | 1514 |
| ACTTTATCAAAATCGGAGAA | PAK5 | 399 | 715 | 1154 | 1029 | 1004 |
| ATTCACGCCTTCTAGAACTG | PAK5 | 565 | 792 | 1534 | 1559 | 1475 |
| CCAGCCCTCAGCCTACCATG | PAK5 | 443 | 712 | 1459 | 1214 | 1202 |
| CTGGCAAGATCAGATTTCCA | PAK5 | 492 | 601 | 735 | 943 | 955 |
| TCGCTCCAACTCCCTACGGA | PAK5 | 312 | 559 | 1285 | 1194 | 1196 |
| TTGTGGGCTCGGACAAGCGA | PAK5 | 446 | 1040 | 1500 | 1551 | 1501 |
| AATATTATGTTAGAACAACA | PALB2 | 520 | 272 | 393 | 349 | 320 |
| ACCAGGGCCACCGACCCAGT | PALB2 | 496 | 393 | 926 | 838 | 904 |
| CTGGAGTTGTGCCTAAGATG | PALB2 | 608 | 821 | 1566 | 1408 | 1502 |
| GCACAGAAGCATCTTAAGGA | PALB2 | 425 | 376 | 674 | 487 | 538 |
| TAAGCCTCCTAAAGATGACT | PALB2 | 478 | 440 | 802 | 707 | 737 |
| TTACCTGAGTGGTTTAGCTG | PALB2 | 594 | 781 | 1231 | 996 | 1116 |
| AAGAAAAGGATAAGGACAGT | PARP1 | 477 | 522 | 714 | 788 | 807 |
| CAGCATCCGAGTGGTATCTG | PARP1 | 517 | 701 | 1388 | 1346 | 1375 |
| GAAAGTCAAAAAGACTGCGG | PARP1 | 513 | 388 | 727 | 656 | 677 |
| GTTTACACTAGCAGGCGCCA | PARP1 | 348 | 380 | 612 | 567 | 617 |
| TCTTAACAAAGCAGTTCGGG | PARP1 | 505 | 640 | 1187 | 1047 | 1063 |
| TGATGCCTACTACTGTACCG | PARP1 | 477 | 829 | 1481 | 1574 | 1473 |
| ACAGCTAGATCTTCGAGTGC | PARP2 | 552 | 639 | 1036 | 1076 | 1044 |
| ACAGTGGCACAAATCAAGGC | PARP2 | 541 | 277 | 483 | 471 | 458 |
| AGAAAACTCGGAGATGCCAG | PARP2 | 402 | 642 | 1315 | 1122 | 1156 |
| AGACAAAGTACAGCTACTGG | PARP2 | 478 | 614 | 1199 | 1121 | 1136 |
| GCTCCCTTCACAGTACACGT | PARP2 | 601 | 299 | 520 | 498 | 433 |
| GGATCGTGAGAAGTTTGAGA | PARP2 | 671 | 682 | 1340 | 1280 | 1246 |
| AGCGCCCAGTTGGGCGACGG | PATZ1 | 612 | 759 | 1303 | 1317 | 1371 |
| CAATGACCGACCTCATCAGT | PATZ1 | 606 | 551 | 932 | 768 | 799 |
| CCCATCTCTGAGGACCCCGA | PATZ1 | 485 | 601 | 818 | 801 | 785 |
| CTTGAATGGACATATCAAGC | PATZ1 | 501 | 395 | 703 | 683 | 683 |
| GACTTCCCCGTTCCCCAATG | PATZ1 | 734 | 667 | 1165 | 1271 | 1277 |
| GAGGCGGGCATCCTTCCATG | PATZ1 | 586 | 739 | 1212 | 1110 | 1114 |
| AATTGGAGGATCCAAACCAA | PAX5 | 534 | 497 | 821 | 831 | 773 |
| ACGTGTGTGTGACAATGACA | PAX5 | 490 | 728 | 1256 | 1162 | 1237 |
| AGCTGCCTTGAGATGTCGCA | PAX5 | 620 | 614 | 979 | 1028 | 1063 |
| AGGAGTGAATCAGCTTGGGG | PAX5 | 552 | 916 | 1583 | 1548 | 1601 |
| AGTATTGTCTCATCAGCACG | PAX5 | 401 | 872 | 1439 | 1408 | 1377 |
| CACCGACGACACCTGCGTCA | PAX5 | 334 | 507 | 1054 | 1027 | 1021 |
| CATACACGTAGTCTCCAACG | PBRM1 | 568 | 652 | 1385 | 1074 | 1107 |
| CCAAGTCAAGCCTTCGAATG | PBRM1 | 656 | 770 | 1474 | 1582 | 1542 |
| GAATGCCAGGCATTACAACG | PBRM1 | 650 | 826 | 1013 | 951 | 997 |
| GGTCTCTCCGAGTTTCAAGG | PBRM1 | 367 | 679 | 1192 | 1226 | 1248 |
| GTGAGATGTTGGGTCTCAGG | PBRM1 | 530 | 621 | 1112 | 1170 | 1167 |
| TCATTAGGGCACCCAAACGA | PBRM1 | 598 | 415 | 825 | 833 | 776 |
| AAATAAACACTCATTTCCTC | PDCD1LG2 | 445 | 240 | 378 | 347 | 343 |
| AGTCAGATATTTGTAGTCCC | PDCD1LG2 | 633 | 802 | 1339 | 1293 | 1333 |
| CCACATCCCTCACGTCCAAG | PDCD1LG2 | 556 | 688 | 1081 | 991 | 941 |
| CCAGGCCGAAGGCTATCCCC | PDCD1LG2 | 437 | 520 | 706 | 637 | 705 |
| GAACTGTACACAGTAGACTC | PDCD1LG2 | 633 | 440 | 802 | 686 | 842 |
| GTGAGAGAGCCACCTTGCTG | PDCD1LG2 | 493 | 914 | 1686 | 1643 | 1640 |
| CATCTCGCTCACGGAACGGT | PDE4A | 589 | 577 | 1000 | 951 | 927 |
| CCGGTACCGTGGAAGGACGA | PDE4A | 502 | 751 | 1372 | 1251 | 1393 |
| GACTTATATGCTGACCCTGG | PDE4A | 562 | 454 | 1027 | 1004 | 1116 |
| GCTCATGTACAACGATGAGT | PDE4A | 536 | 609 | 1014 | 950 | 979 |
| GCTGTCCGAACGGTACAGGA | PDE4A | 578 | 652 | 834 | 783 | 785 |
| TCTGTCTTCACCCCAAAGCG | PDE4A | 596 | 633 | 949 | 1055 | 1140 |
| AAGAAGAACATGGGTTGACT | PDE4B | 541 | 349 | 599 | 582 | 551 |
| AGAGAAAGGACTCTCTACGC | PDE4B | 653 | 542 | 794 | 755 | 779 |
| CAACTCATGACACAGATAAG | PDE4B | 585 | 524 | 1280 | 1104 | 1182 |
| GCATACATGATGCATGTGAG | PDE4B | 411 | 798 | 1393 | 1127 | 1114 |
| GTTCAAAAGAATGCTGAACC | PDE4B | 551 | 589 | 903 | 843 | 848 |
| TTACAACCGTAAGCCAGGAG | PDE4B | 526 | 668 | 873 | 899 | 858 |
| AGAAGAATCATTGTACATCA | PDE4D | 452 | 386 | 698 | 729 | 742 |
| AGACAGGTCATAATCGCTGT | PDE4D | 587 | 581 | 1002 | 1052 | 1034 |
| ATGGATGGTTGGTTGCACAT | PDE4D | 443 | 563 | 1215 | 1110 | 1137 |
| CCCATCTCTCGGAAATGAGT | PDE4D | 644 | 366 | 562 | 503 | 572 |
| CGTGGCCGCTACACCTCGAG | PDE4D | 542 | 544 | 1038 | 819 | 969 |
| TGACGAATTCAAGCATCCCG | PDE4D | 608 | 477 | 812 | 728 | 705 |
| AAACAGGGAATAGCGATCAG | PDE5A | 482 | 665 | 1336 | 1381 | 1365 |
| AAAGAGGTGAATTACCGCCA | PDE5A | 593 | 474 | 887 | 826 | 821 |
| AATTGCCAGATACGTTAACA | PDE5A | 511 | 416 | 864 | 802 | 857 |
| ATCCGCTTAGAATGGAACAA | PDE5A | 617 | 700 | 1201 | 1156 | 1261 |
| CCTGGGCTACACCAACAACC | PDE5A | 393 | 504 | 592 | 647 | 675 |
| TTGACGACAATGGGTCTAAG | PDE5A | 471 | 622 | 1085 | 1110 | 1086 |
| ACAGACGAAAACGAGATCGA | PDGFRA | 453 | 623 | 986 | 1136 | 1140 |
| ACCAAGTCCAGAATAGACGA | PDGFRA | 393 | 373 | 884 | 768 | 900 |
| ATCTCCACTATCTTTCACCG | PDGFRA | 336 | 504 | 809 | 795 | 749 |
| GATATACGGCCCAACGCTGA | PDGFRA | 424 | 687 | 1197 | 1111 | 1102 |
| GCTGCGTTCTGAGCTCACGG | PDGFRA | 468 | 753 | 1364 | 1417 | 1362 |
| TGAAAAACAAGCTCTCATGT | PDGFRA | 471 | 574 | 928 | 905 | 976 |
| AGTGACGGATCCAAGACTGG | PDGFRB | 584 | 796 | 1406 | 1272 | 1254 |
| CCAGCAATACCTCCATGCGT | PDGFRB | 608 | 369 | 617 | 596 | 576 |
| GATGATTGTGAGAACCACCA | PDGFRB | 558 | 519 | 856 | 862 | 845 |
| GTACGTGTCAGAACTGACGT | PDGFRB | 418 | 461 | 978 | 864 | 833 |
| TCCACCATAGGACGTACGCT | PDGFRB | 348 | 559 | 840 | 836 | 839 |
| TCCTCTGAACTGTTCCCTAG | PDGFRB | 392 | 599 | 977 | 894 | 983 |
| CAATTGCAATGTCTCTGAGG | PDK2 | 473 | 604 | 1323 | 1100 | 1139 |
| CTCCAAACAGCCAATTCACA | PDK2 | 432 | 342 | 613 | 769 | 763 |
| GAATGCCATGCGTGCAACTG | PDK2 | 560 | 652 | 1108 | 1137 | 1188 |
| GGAGACTGGGTCGTCGCCAT | PDK2 | 591 | 467 | 778 | 873 | 896 |
| TCAGGGTACGATGGTCCTCG | PDK2 | 533 | 961 | 1867 | 1907 | 1892 |
| TGAGCATGCGGATGGAAATG | PDK2 | 590 | 753 | 1555 | 1517 | 1372 |
| ATGTATAATATACCAGCTCG | PDPK1 | 508 | 709 | 1456 | 1178 | 1275 |
| CACGAAGCGATTAGGCTGTG | PDPK1 | 617 | 1015 | 1602 | 1801 | 1800 |
| CATTCGATGAGACATGCACC | PDPK1 | 486 | 530 | 827 | 757 | 811 |
| GCTCGGTCGAGACTCAGCCG | PDPK1 | 523 | 471 | 988 | 875 | 958 |
| GGCTGTTCCGACAAACGAGT | PDPK1 | 469 | 329 | 556 | 408 | 504 |
| GTTCTCGGGCTAGAACAACC | PDPK1 | 538 | 525 | 863 | 678 | 708 |
| ATGGCTACCTCGCTCGCCAG | PER1 | 555 | 561 | 1124 | 1133 | 1023 |
| CCGTTTCTGCGCCCGCAATG | PER1 | 398 | 1020 | 1452 | 1439 | 1437 |
| CCTCCGACGATGACAAGCAG | PER1 | 472 | 657 | 919 | 966 | 1068 |
| GCAGCTGGGTGTATGCCGAG | PER1 | 375 | 904 | 1438 | 1540 | 1540 |
| GGCCAGAGCATACTGTAGCG | PER1 | 555 | 574 | 1054 | 1142 | 1132 |
| TCCTCCAGTGATAGCAACGG | PER1 | 467 | 581 | 1083 | 1019 | 1106 |
| AAAAGACTCTTGGTACCTGT | PGK1 | 551 | 403 | 704 | 681 | 704 |
| AAGAATACTTGTCTGGCATG | PGK1 | 296 | 961 | 1421 | 1455 | 1536 |
| ACCAGATAACAAACAACCAG | PGK1 | 343 | 206 | 435 | 356 | 334 |
| CTTCCTCCACATGAAAGCGA | PGK1 | 516 | 329 | 540 | 565 | 524 |
| GAAGGTGCTCAACAACATGG | PGK1 | 640 | 588 | 977 | 821 | 915 |
| TCCTTCATACCCGCCCAGGA | PGK1 | 401 | 493 | 821 | 878 | 969 |
| AAGGGCTGTCGGAAGATACC | PHF6 | 438 | 457 | 875 | 968 | 912 |
| AATTAAAAGAGGCACGAAGC | PHF6 | 255 | 338 | 591 | 589 | 578 |
| AGTGACACCAGGCCTAAATG | PHF6 | 511 | 794 | 1426 | 1435 | 1344 |
| CTTTATCATGCAATGCACAG | PHF6 | 452 | 459 | 700 | 609 | 571 |
| GCATATATTTAATGCCAAGA | PHF6 | 364 | 582 | 820 | 784 | 825 |
| GGCAGCACACCATAAGTGCA | PHF6 | 572 | 749 | 1404 | 1320 | 1408 |
| ACATCTTGTCCTGTGCACCG | PHLPP2 | 435 | 774 | 1593 | 1468 | 1407 |
| AGTCTGACAATCCTTCACTG | PHLPP2 | 401 | 570 | 1147 | 965 | 1031 |
| ATTGTCCCGCAAATCCATGT | PHLPP2 | 537 | 418 | 802 | 824 | 799 |
| GTTGTGTCGCAAATTGAGGT | PHLPP2 | 547 | 686 | 1094 | 1059 | 1077 |
| GTTGTTGGAGTGTGCCACAA | PHLPP2 | 628 | 722 | 1337 | 1238 | 1225 |
| GTTTGATGGGGACCGAAACG | PHLPP2 | 582 | 514 | 1003 | 966 | 924 |
| CCTACGAGTCCTGTATGGCT | PHOX2B | 485 | 647 | 1051 | 958 | 972 |
| CCTGAAGATCGACCTCACCG | PHOX2B | 568 | 708 | 1308 | 1187 | 1170 |
| CGGCCAAGAACGGGTCCTCG | PHOX2B | 586 | 799 | 1682 | 1596 | 1576 |
| GATGTCGGGGTAGTGCGTCT | PHOX2B | 619 | 596 | 1053 | 1077 | 1098 |
| GCTTCCAGTATAACCCGATA | PHOX2B | 426 | 627 | 982 | 1041 | 1046 |
| TACAAACTCTTCACCGACCA | PHOX2B | 439 | 547 | 899 | 989 | 1072 |
| AGTTTCTTCCTGCTAGACGG | PI4KB | 364 | 530 | 1128 | 1057 | 1101 |
| ATGTACATGTTAAGCAACTG | PI4KB | 442 | 466 | 1027 | 1113 | 1016 |
| CCACTCAGCGACACTCCCGT | PI4KB | 525 | 354 | 738 | 598 | 565 |
| GTCAGTGATTGACCGTGAGG | PI4KB | 480 | 701 | 1246 | 1103 | 1147 |
| GTGGAGAACCTGCCCGAATG | PI4KB | 547 | 628 | 1277 | 1117 | 1103 |
| TCCACCTACCCAGATAAGGG | PI4KB | 446 | 645 | 1229 | 1144 | 1193 |
| ACAGTTCGTAAAAGGCCCGG | PICK1 | 375 | 720 | 1193 | 1265 | 1376 |
| CAGGGCATGTCCCTTGACAT | PICK1 | 608 | 631 | 1164 | 1129 | 991 |
| CCAGTACCTGGACAATATAG | PICK1 | 350 | 487 | 685 | 772 | 760 |
| CTGAAAACCATCAAGCCGGT | PICK1 | 567 | 338 | 784 | 820 | 799 |
| CTTCACAAAAGCCTCACTCG | PICK1 | 495 | 589 | 1104 | 1067 | 1008 |
| GATTGACCTGCCATTGACAC | PICK1 | 428 | 615 | 758 | 839 | 906 |
| ACTCCAGACATAACCGGTGA | PIK3C2B | 628 | 491 | 945 | 849 | 840 |
| ATACTGCAGCACATTTGATG | PIK3C2B | 586 | 739 | 1358 | 1354 | 1320 |
| GACAATACTGAATGTACTCG | PIK3C2B | 511 | 695 | 1327 | 1303 | 1383 |
| GGCTACTAAACACATAGCGG | PIK3C2B | 412 | 488 | 905 | 849 | 930 |
| GTATCAATGATGCGATCACA | PIK3C2B | 447 | 542 | 1093 | 967 | 955 |
| TCAGGAGAAAGCGCACCAAA | PIK3C2B | 469 | 441 | 621 | 589 | 685 |
| AACCTCGAACCATAGGATCT | PIK3CA | 571 | 586 | 1145 | 1226 | 1120 |
| AGCAAATCTTCTAATCCATG | PIK3CA | 523 | 612 | 1230 | 1134 | 1052 |
| GATCTCGGGTACAAATTGCT | PIK3CA | 572 | 567 | 1003 | 937 | 955 |
| GTGCGCTATTTATAACCCAA | PIK3CA | 325 | 394 | 623 | 621 | 620 |
| TCAAATACATCCCACAAGCA | PIK3CA | 566 | 339 | 579 | 627 | 633 |
| TCATACCAATTTCTCGATTG | PIK3CA | 567 | 829 | 1571 | 1355 | 1574 |
| AAAAATGCGCAAATTCAGTG | PIK3CB | 614 | 465 | 829 | 799 | 755 |
| AAAGAGCGCTTGCTAATCGG | PIK3CB | 430 | 444 | 773 | 740 | 749 |
| CTTGCAACACATAATCATTG | PIK3CB | 400 | 553 | 707 | 734 | 804 |
| GAATCAAACTGCTGTATATG | PIK3CB | 367 | 823 | 1417 | 1539 | 1529 |
| TGATATCTCTGAGCTTACGA | PIK3CB | 302 | 369 | 663 | 509 | 447 |
| TGTGGCATGGGTAAATACGA | PIK3CB | 481 | 458 | 893 | 878 | 907 |
| ATAGTCAAACAGCATGAGGT | PIK3CD | 496 | 543 | 1151 | 972 | 1035 |
| CGAGCATGGGCGCTTCAGTG | PIK3CD | 506 | 777 | 1295 | 1433 | 1509 |
| GATGGAGGAAGAGTGCACCA | PIK3CD | 371 | 696 | 1091 | 1086 | 1004 |
| GCAAAGTGAATGCCGATGAG | PIK3CD | 666 | 663 | 1267 | 1227 | 1300 |
| GCCGCCAGGTCCACTACCGG | PIK3CD | 484 | 509 | 865 | 948 | 1079 |
| GTTGACGCAGGTGAACACAT | PIK3CD | 357 | 426 | 652 | 618 | 624 |
| AGGTTCAGGGTCAAAATCCG | PIK3CG | 513 | 632 | 1279 | 1201 | 1143 |
| AGTCCAGGCACTATCAGCAG | PIK3CG | 423 | 614 | 1236 | 1240 | 1285 |
| AGTGGTACGAGATCTACGAC | PIK3CG | 496 | 307 | 508 | 540 | 506 |
| GGATCCCCTTAACCCTCTCA | PIK3CG | 372 | 715 | 1191 | 1164 | 1249 |
| GGGAGTACGTGCTCCACATG | PIK3CG | 482 | 1021 | 1832 | 1820 | 1728 |
| GTCTTCATCGTCATCCACCG | PIK3CG | 472 | 742 | 1209 | 1077 | 1090 |
| AAAAAAGTCGGGAATACGAT | PIK3R1 | 899 | 434 | 879 | 782 | 759 |
| AACTTCGAGATACAGCCGAT | PIK3R1 | 559 | 429 | 828 | 832 | 814 |
| AGCGTAAGCCAATACTGATG | PIK3R1 | 385 | 765 | 1424 | 1524 | 1532 |
| AGGCGGAGCGATGTCAGGAG | PIK3R1 | 442 | 526 | 1135 | 1095 | 1125 |
| TGGTTATAATGAAACGACGG | PIK3R1 | 376 | 602 | 1103 | 1071 | 1049 |
| TTACGACAGCTCCTTGATTG | PIK3R1 | 522 | 756 | 1378 | 1350 | 1359 |
| AAGGACCGGATTTCCGACTG | PIM1 | 495 | 799 | 1115 | 1124 | 1092 |
| ATTCTCATCGACCTCAATCG | PIM1 | 426 | 458 | 975 | 939 | 962 |
| CAGGATCAAGACGAAACTGT | PIM1 | 510 | 526 | 840 | 788 | 823 |
| CCGGCAAGTTGTCGGCGACA | PIM1 | 603 | 767 | 1164 | 1302 | 1539 |
| CGACTTTATCACGGAAAGGG | PIM1 | 475 | 541 | 1075 | 942 | 945 |
| GTGCGGCACTGCCACAACTG | PIM1 | 481 | 710 | 1277 | 1149 | 1147 |
| AGATGGCGACAACGTCACGG | PKHD1 | 577 | 540 | 894 | 948 | 1012 |
| AGTAGGATGTTAGCCGAAAG | PKHD1 | 553 | 807 | 1365 | 1372 | 1267 |
| CATGTCTTACAGGCAATCGA | PKHD1 | 548 | 336 | 544 | 551 | 491 |
| TGAGATGCTATAGAGACCGG | PKHD1 | 368 | 550 | 1117 | 1124 | 1120 |
| TGGTGGAAGCCTCTTGACCG | PKHD1 | 318 | 779 | 1529 | 1641 | 1633 |
| TTAGATCCTCCATTACAGTG | PKHD1 | 512 | 549 | 1308 | 1238 | 1262 |
| AATGGATGCTAATTATGTCA | PLA2G4A | 391 | 527 | 915 | 796 | 847 |
| ACAGAGCCATACTGAATCGG | PLA2G4A | 585 | 655 | 1109 | 1299 | 1245 |
| ATTACTCACAATATGCTCTG | PLA2G4A | 559 | 589 | 760 | 708 | 808 |
| CAGTCCATATGAGATCGGCA | PLA2G4A | 641 | 601 | 1054 | 968 | 986 |
| TAAAATGTATACTAACCATG | PLA2G4A | 478 | 411 | 767 | 608 | 647 |
| TGACACTCCAGATCCGTATG | PLA2G4A | 482 | 817 | 1262 | 1233 | 1275 |
| CCCAGTATAAACTATCCTGG | PLA2G4B | 688 | 660 | 1320 | 1248 | 1175 |
| CTTCCTTCGCGCCAGACCGA | PLA2G4B | 699 | 602 | 1062 | 1131 | 1113 |
| GGCTTTGGCCAACCTCTACG | PLA2G4B | 576 | 737 | 1395 | 1360 | 1413 |
| GGGAAAGCCAACCTCATAGG | PLA2G4B | 559 | 510 | 991 | 928 | 924 |
| GGTTCACAGGATGCCTCCGT | PLA2G4B | 531 | 309 | 866 | 648 | 607 |
| TCAGTGAGAAGCTCTCTCGG | PLA2G4B | 631 | 827 | 1442 | 1378 | 1513 |
| AAGAGACAGAACTGAGTGTG | PLA2G4D | 635 | 696 | 1226 | 1298 | 1244 |
| ACAGGTCCACAAAAGTTGTG | PLA2G4D | 524 | 1068 | 2022 | 1825 | 1737 |
| AGACTATACCCTTGAAGTCG | PLA2G4D | 493 | 949 | 1519 | 1434 | 1530 |
| CATCACCAACAACGTCCTTG | PLA2G4D | 497 | 643 | 1136 | 1190 | 1189 |
| CTATGATGAGGACTCAGTCA | PLA2G4D | 392 | 1052 | 1752 | 1848 | 1910 |
| GGGCTATGATTTGTGTGCCG | PLA2G4D | 644 | 832 | 1590 | 1452 | 1528 |
| ACTCTTACGGAAGCCGTATG | PLAUR | 325 | 835 | 1626 | 1823 | 1799 |
| AGCCGCAGGAAACACATTCG | PLAUR | 578 | 729 | 1936 | 1943 | 1999 |
| CAACACCACCAAATGCAATG | PLAUR | 361 | 438 | 865 | 781 | 825 |
| CAGGAGCTGCTTCCAGACAT | PLAUR | 469 | 405 | 961 | 964 | 891 |
| GTGGAAGTGGTGACCCACAG | PLAUR | 543 | 816 | 1173 | 1370 | 1312 |
| TTCCCATATGCGCATGATCG | PLAUR | 330 | 770 | 1568 | 1457 | 1476 |
| CCACGGAGCAAATCCCCCAG | PLCG1 | 587 | 614 | 1104 | 873 | 940 |
| GCTGCGCTATCCCATCAATG | PLCG1 | 510 | 796 | 1371 | 1348 | 1259 |
| GGATGCCAGTCATTTACCAT | PLCG1 | 465 | 706 | 1132 | 1142 | 1155 |
| GGGAACCCGGTAGTTGACCT | PLCG1 | 518 | 632 | 1077 | 995 | 950 |
| TCAAATTCGTTGCAGCGCAG | PLCG1 | 388 | 452 | 1018 | 825 | 795 |
| TCGGAGTACATCACTGACGT | PLCG1 | 790 | 531 | 1059 | 999 | 1093 |
| AAGAGCGAAAGAGTTTCAGG | PLD1 | 505 | 459 | 960 | 910 | 982 |
| ACACTATTGAACTAACACAT | PLD1 | 432 | 274 | 324 | 310 | 298 |
| CGGCATATGACCCAAATCGG | PLD1 | 680 | 725 | 1360 | 1314 | 1270 |
| GAACAGTACTTACTGTCGCG | PLD1 | 384 | 658 | 1083 | 1035 | 1037 |
| TGATAACGAACACAGACTCA | PLD1 | 499 | 688 | 1335 | 1317 | 1321 |
| TTGACAGATACTCCACTCCC | PLD1 | 483 | 473 | 713 | 689 | 615 |
| AGGACCTACTCGTGGCAGAG | PLD2 | 509 | 584 | 1246 | 1170 | 1149 |
| AGGTCACCTTTATGTTAGGG | PLD2 | 650 | 774 | 1037 | 996 | 954 |
| GAGATCACGGAGTTGGCCCA | PLD2 | 711 | 913 | 1420 | 1394 | 1382 |
| GGGGTCGAAGAGCTGAACAA | PLD2 | 479 | 633 | 1205 | 1214 | 1257 |
| TAAATCTCAGGACTCAACCT | PLD2 | 604 | 404 | 734 | 696 | 683 |
| TTGACAGGGAGACTACCCCC | PLD2 | 608 | 506 | 947 | 898 | 881 |
| AACCAAAGTGGAATATGACG | PLK1 | 565 | 385 | 898 | 650 | 678 |
| AACTTACATGATACACCCAA | PLK1 | 422 | 307 | 337 | 295 | 324 |
| CCTGTCTCACCATTGCACCC | PLK1 | 592 | 358 | 561 | 626 | 628 |
| CTCTCCTGGAACTCCACAAG | PLK1 | 565 | 379 | 652 | 500 | 491 |
| GCATCCAAACCCTCAGAGCG | PLK1 | 364 | 481 | 859 | 803 | 782 |
| GCCGGCGAAAGAGATCCCGG | PLK1 | 477 | 309 | 579 | 462 | 514 |
| AGGTCAACTGTCTCAAAGGG | PLK3 | 382 | 635 | 1402 | 1339 | 1366 |
| GAGGCCACTGACACGGAGAG | PLK3 | 286 | 628 | 1225 | 1265 | 1255 |
| GGATCTGGTCAATTGAGGGG | PLK3 | 325 | 824 | 1410 | 1279 | 1249 |
| GGCCAATGGACGTCCGAGTG | PLK3 | 689 | 796 | 1381 | 1485 | 1489 |
| GGTGGGAGAAACGCACGATG | PLK3 | 453 | 840 | 1334 | 1433 | 1382 |
| GTCCGGAAAGGATCTGACGC | PLK3 | 343 | 347 | 615 | 549 | 511 |
| CCAAGATACAGATCATCAAC | PLP2 | 494 | 355 | 507 | 419 | 448 |
| CCACTCCAAAATCATCGCGG | PLP2 | 577 | 551 | 926 | 828 | 872 |
| CGATGGAGGTGATCAGGTAG | PLP2 | 583 | 961 | 1746 | 1578 | 1558 |
| GGATGGCTGCTATGAGGGTT | PLP2 | 462 | 711 | 1304 | 1197 | 1149 |
| TCCTTCAACAAGGACAACGA | PLP2 | 574 | 296 | 596 | 605 | 632 |
| TGATCTGTATCTTGGTGTGC | PLP2 | 606 | 552 | 1135 | 1037 | 1031 |
| AAGTCCCATCATGGGAAACG | PML | 626 | 816 | 1221 | 1190 | 1119 |
| ACGGCCGTGTGCAAGTCGTG | PML | 464 | 846 | 1479 | 1561 | 1581 |
| ACTGTACCAAAGCCCCAGTG | PML | 406 | 483 | 1147 | 1200 | 1226 |
| GCTCAAGTGCGACATCAGCG | PML | 446 | 919 | 1471 | 1634 | 1697 |
| GTCGGTTTACAGGCAGATCG | PML | 578 | 780 | 1273 | 1267 | 1256 |
| TGTTTCAGAGGAACGCGTCG | PML | 587 | 1038 | 1368 | 1310 | 1311 |
| ATGTGTGTAATGATAAATCT | PMS1 | 523 | 718 | 1106 | 1117 | 1087 |
| CTGTAGCAAAAAATCAAATG | PMS1 | 563 | 486 | 701 | 724 | 746 |
| GAAAATGGGATACAAACGAC | PMS1 | 656 | 249 | 439 | 407 | 372 |
| TAGAGTATCCGATCGTCAGG | PMS1 | 470 | 638 | 1380 | 1434 | 1401 |
| TGAGGTACGAGACAATGGTG | PMS1 | 287 | 717 | 1438 | 1387 | 1395 |
| TTCTTGTAGTAATTAAAACC | PMS1 | 480 | 324 | 620 | 626 | 638 |
| AAATAGCTGCTGTACGCTAA | PMS2 | 558 | 657 | 1115 | 1125 | 1095 |
| ACAACAGAGAACAGGTCTCG | PMS2 | 536 | 1082 | 1679 | 1674 | 1626 |
| CTGTGGCCGCAGTAACAGGG | PMS2 | 633 | 472 | 864 | 864 | 852 |
| GGAGTTGGTAGAAAATAGTG | PMS2 | 591 | 862 | 1399 | 1229 | 1255 |
| GGTCTCCAGACTCGTGAATG | PMS2 | 350 | 971 | 1816 | 1568 | 1767 |
| TCACAAACGGAGTCACTAGG | PMS2 | 390 | 510 | 986 | 1000 | 997 |
| CCAGCACCTCTGTCCAACTG | PNRC1 | 657 | 501 | 1113 | 1091 | 1080 |
| CCAGCCCTCGTTCAGAAGAG | PNRC1 | 446 | 484 | 1064 | 1141 | 1170 |
| GAGAGCTCCGAAGTAACCGC | PNRC1 | 641 | 341 | 658 | 543 | 546 |
| GCAGCACCGGCCGAGCCTCG | PNRC1 | 441 | 1035 | 1378 | 1451 | 1340 |
| GCCGCCTTTACCCCGAATCC | PNRC1 | 306 | 609 | 930 | 908 | 979 |
| TGCGGCCGCAGCCCAAACTG | PNRC1 | 376 | 771 | 1367 | 1320 | 1245 |
| AAGCACGTACCAAAGACAGG | POLA1 | 577 | 339 | 684 | 596 | 613 |
| ACAGGAACTGGAGTTAATGG | POLA1 | 505 | 422 | 968 | 774 | 823 |
| AGAGACACAGAAAGTTACGG | POLA1 | 583 | 444 | 815 | 694 | 698 |
| ATGCTCTTACAAGTAGGACG | POLA1 | 456 | 573 | 1129 | 1017 | 1061 |
| TGAGAGGGTTGTAATCCCAA | POLA1 | 478 | 586 | 1166 | 942 | 1118 |
| TGCAAAGACACATCAACATG | POLA1 | 346 | 557 | 1050 | 881 | 854 |
| AATGGCAATCAGATCGAGCA | POLA2 | 603 | 441 | 643 | 610 | 537 |
| CATTCTGTATGGACAGAACG | POLA2 | 434 | 713 | 1658 | 1398 | 1391 |
| TACAGCTCAAGAAATAACCG | POLA2 | 456 | 377 | 666 | 538 | 577 |
| TAGCAAGTCAGTGATTCTTG | POLA2 | 580 | 532 | 1446 | 1247 | 1168 |
| TCTTTGTAGAATCGAAGTGG | POLA2 | 509 | 688 | 1500 | 1163 | 1196 |
| TTAGTGATTCTGGACACCCT | POLA2 | 309 | 606 | 1185 | 1119 | 1200 |
| CAAGAAGATACACGTCACCA | POLE | 542 | 408 | 883 | 619 | 662 |
| CCTGGTACATGCTTAGTCCG | POLE | 604 | 590 | 1162 | 1006 | 1065 |
| CTTCGAGATCAAACGCCGAG | POLE | 503 | 446 | 817 | 891 | 804 |
| GAGTACTACATTGAGCGGCT | POLE | 480 | 316 | 636 | 614 | 667 |
| GGTTGACATAGAATACTCGG | POLE | 458 | 373 | 634 | 422 | 454 |
| TGGTACAACATCAGATACCG | POLE | 370 | 363 | 728 | 544 | 596 |
| ACAACTTCCACCATGGACCG | PORCN | 435 | 816 | 1526 | 1498 | 1493 |
| ACATGGTGGATACCGTGACA | PORCN | 496 | 684 | 1261 | 1139 | 1127 |
| AGGAGGCGGTCACCGTCGAG | PORCN | 572 | 721 | 1419 | 1372 | 1428 |
| CCTGAAGCATGCGAGCACCG | PORCN | 231 | 810 | 1175 | 1109 | 1120 |
| GAAGGAGACAGCACTCTCGT | PORCN | 543 | 472 | 894 | 695 | 707 |
| GGAGAGGAAGACGCCTCGAT | PORCN | 501 | 456 | 922 | 810 | 789 |
| AATGGAATGTCTTCATAGTG | PPARG | 469 | 884 | 1336 | 1307 | 1420 |
| ATAGATGTGACCTAAACTGT | PPARG | 592 | 590 | 1219 | 1227 | 1318 |
| CCTGCACGGCTTCCACTGAG | PPARG | 535 | 868 | 1249 | 1395 | 1350 |
| CTCGGCATTCAATTGCCATG | PPARG | 668 | 895 | 1442 | 1515 | 1442 |
| GTACCCAGGTGCAATCAAAG | PPARG | 571 | 555 | 948 | 981 | 1015 |
| TGGCATCTCTGTGTCAACCA | PPARG | 367 | 618 | 1196 | 1155 | 1167 |
| ACATGTTTCCCATCCAAGTG | PPID | 545 | 611 | 1339 | 1178 | 1188 |
| CAATCATCCAGATTTCCCTG | PPID | 481 | 625 | 915 | 953 | 913 |
| CCCCAGAGTCTTCTTTGACG | PPID | 438 | 838 | 1410 | 1322 | 1242 |
| TCAAATCAAAATGGGACAGG | PPID | 530 | 523 | 917 | 844 | 951 |
| TCAGGAGGGTTTATTGAGCA | PPID | 483 | 689 | 1295 | 1338 | 1413 |
| TTACACATACTTCGATGGAA | PPID | 447 | 532 | 802 | 860 | 833 |
| ACATACCTGCCCATCAACCA | PPP2CA | 611 | 247 | 476 | 474 | 474 |
| ACATCGAACCTCTTGCACGT | PPP2CA | 537 | 410 | 744 | 611 | 645 |
| CTCACCATCCATAGATACAC | PPP2CA | 357 | 257 | 345 | 354 | 304 |
| GAACGCATCACCATTCTTCG | PPP2CA | 496 | 513 | 1136 | 1032 | 1021 |
| GTCCAATGTGTGACTTGCTG | PPP2CA | 588 | 992 | 1789 | 1926 | 1812 |
| TGGGCAATTTCATGATCTCA | PPP2CA | 507 | 601 | 1218 | 1295 | 1234 |
| AGAACCCACCTGCTCGTCAG | PPP2R1A | 491 | 827 | 1384 | 1178 | 1201 |
| AGCCGCACCTCTGGACACTG | PPP2R1A | 336 | 682 | 1110 | 1108 | 1089 |
| ATGGCGTGTCCGATACATGG | PPP2R1A | 609 | 784 | 1354 | 1208 | 1239 |
| CCCAGATACCATCTACGATG | PPP2R1A | 599 | 702 | 1158 | 1164 | 1157 |
| CCTCCCACAAGGTCAAAGGT | PPP2R1A | 520 | 439 | 1055 | 1044 | 1078 |
| GTGCTCAGATGACACCCCCA | PPP2R1A | 390 | 405 | 915 | 915 | 843 |
| ACAAACCACCGTGCACACAC | PPP3CA | 528 | 268 | 488 | 446 | 509 |
| CAACATAGTCCCCTAAGAAG | PPP3CA | 430 | 546 | 825 | 682 | 686 |
| GACCTATGTGTGATATCCTG | PPP3CA | 522 | 746 | 1377 | 1208 | 1239 |
| GATATTGATGCACCGGTCAC | PPP3CA | 581 | 459 | 848 | 869 | 936 |
| GTAAGAATTCACATACAGCT | PPP3CA | 289 | 369 | 514 | 530 | 530 |
| TAACGATGGGAAACCTCGTG | PPP3CA | 483 | 470 | 863 | 1131 | 1035 |
| AAGAGTCTATGAAGCGTGTA | PPP3CB | 423 | 684 | 1323 | 1226 | 1191 |
| ACAGAAAAAGTCAAACTACA | PPP3CB | 474 | 329 | 447 | 435 | 407 |
| CACACACTGCTGGATAGCTG | PPP3CB | 522 | 845 | 1422 | 1240 | 1151 |
| GATGTCAAGCGATGTGTTGG | PPP3CB | 464 | 955 | 1575 | 1420 | 1477 |
| TGCACTTAGAATTATCAATG | PPP3CB | 273 | 562 | 1188 | 1033 | 1017 |
| TTCACCAGAAATACACACAC | PPP3CB | 353 | 201 | 333 | 279 | 330 |
| ATACCCACAACACTGTCCGA | PPP3CC | 640 | 406 | 577 | 653 | 643 |
| CAGAGGCTATTTCAGTATAG | PPP3CC | 470 | 794 | 1519 | 1403 | 1284 |
| CTTCCAGGTATCGAATGTAC | PPP3CC | 603 | 522 | 741 | 637 | 669 |
| GAAGTAGAGGCTCCAATCAC | PPP3CC | 364 | 457 | 835 | 701 | 845 |
| TGGTTTAAGAGGGCAGCAAG | PPP3CC | 396 | 843 | 1391 | 1510 | 1333 |
| TTTGGAAGCCAGTAGGGGTG | PPP3CC | 386 | 1049 | 1789 | 1761 | 1674 |
| AAAGGATTCTGTTGTAACTC | PPP3R1 | 473 | 687 | 986 | 985 | 1060 |
| AAATGAGGCAAGTTATCCGT | PPP3R1 | 616 | 546 | 888 | 853 | 833 |
| CAATTCGGGTTCTTTGAGTG | PPP3R1 | 577 | 836 | 1464 | 1391 | 1331 |
| TGCTTTCCGTATCTATGACA | PPP3R1 | 367 | 550 | 830 | 872 | 903 |
| TTCTGTGGTGTGAAACCAGG | PPP3R1 | 319 | 620 | 1194 | 1148 | 1151 |
| TTCTTGTTCAGAATTCATTG | PPP3R1 | 588 | 752 | 1196 | 1237 | 1251 |
| AAAGACGGCTACATCTCCAA | PPP3R2 | 619 | 434 | 851 | 787 | 889 |
| AAATTAAGAGGCTGGGCAAG | PPP3R2 | 618 | 590 | 1040 | 875 | 903 |
| AGTGGTCGACGTCTTCGACA | PPP3R2 | 374 | 711 | 1174 | 1270 | 1272 |
| CAGGAACTCGTCCACGCTCA | PPP3R2 | 639 | 592 | 1246 | 1213 | 1218 |
| CTCCCAGTTCAGCGTCAGGG | PPP3R2 | 562 | 676 | 1303 | 1331 | 1394 |
| GAACGAAGCCAGTTACCCCG | PPP3R2 | 483 | 545 | 926 | 1007 | 1022 |
| AGGCACCTCTGGGTCCAACA | PRAME | 631 | 478 | 892 | 898 | 880 |
| CAGACAATTAGTTATTGACA | PRAME | 401 | 720 | 954 | 926 | 858 |
| CCAAATAGAGTTCCTCGAGG | PRAME | 322 | 638 | 1180 | 1301 | 1296 |
| GTAGGCGAGGGACTCGTCAG | PRAME | 743 | 649 | 1133 | 1302 | 1100 |
| GTGAAGGCCGCTGTGAACAG | PRAME | 489 | 986 | 1191 | 1142 | 1216 |
| GTGAGGAGCAAACTTCCCCA | PRAME | 459 | 434 | 840 | 718 | 764 |
| AGTAGGCGTACGAGTCCCGG | PRDM1 | 626 | 568 | 1171 | 1045 | 1080 |
| ATTGTCGGGACTTTGCAGAG | PRDM1 | 582 | 689 | 1328 | 1376 | 1319 |
| CGGCTGTACCCGGTGTACGG | PRDM1 | 525 | 839 | 1485 | 1418 | 1333 |
| GGGGTGGTCGTTCACAATGT | PRDM1 | 534 | 432 | 1107 | 1110 | 1166 |
| TTTCACCCTTCGCCGCCGAG | PRDM1 | 568 | 1009 | 1269 | 1202 | 1247 |
| TTTGGGCAGATCTATTCCAG | PRDM1 | 334 | 516 | 1142 | 1049 | 1156 |
| CCTAACGCTCAACAAGAAGT | PREX2 | 454 | 615 | 727 | 878 | 874 |
| TCAGCACACAATCACAACCA | PREX2 | 454 | 530 | 851 | 772 | 782 |
| TGGGATAAAGTGTAACGTGG | PREX2 | 409 | 666 | 1295 | 1146 | 1227 |
| TGGGTAATGATCTCCGAACA | PREX2 | 470 | 662 | 1219 | 1198 | 1084 |
| TGTAACATGTGCAATAACGC | PREX2 | 639 | 288 | 496 | 557 | 515 |
| TGTCTGTTCTAACATAAACG | PREX2 | 346 | 594 | 947 | 1042 | 1053 |
| ATTCTGATGAAGGGTTGGCG | PRF1 | 687 | 643 | 1200 | 1174 | 1213 |
| CAATTTCAACACCTCCACAG | PRF1 | 439 | 476 | 596 | 559 | 584 |
| CATCCACAACGACTGGCGCG | PRF1 | 609 | 505 | 1064 | 1055 | 1005 |
| GGGCCAGCTCACACGTACGC | PRF1 | 364 | 471 | 832 | 874 | 887 |
| TCGAGCCAGGCACGAACCAG | PRF1 | 485 | 502 | 1142 | 1073 | 1276 |
| TCTGCCGCAACACCCTACGG | PRF1 | 648 | 556 | 1076 | 1046 | 1015 |
| ATCTTGACGATAATGAGAGA | PRKAR1A | 562 | 553 | 1151 | 1066 | 1029 |
| CAGCTATAGAAGAATCCTCA | PRKAR1A | 526 | 494 | 787 | 828 | 832 |
| CAGTGCCGAAGTCTACACGG | PRKAR1A | 446 | 568 | 1215 | 1129 | 1148 |
| CGAATGGGCAACCAGTGTTG | PRKAR1A | 456 | 899 | 1850 | 1992 | 1947 |
| CTCCCGTGCAGATTCGAGGG | PRKAR1A | 713 | 763 | 1572 | 1524 | 1486 |
| GAGACCCATGGCATTCCTCA | PRKAR1A | 525 | 454 | 1050 | 862 | 874 |
| CAAAATCAGCAATCTTGATG | PRKCB | 357 | 582 | 1190 | 1196 | 1265 |
| GCCACCTACCGACAACGATG | PRKCB | 480 | 872 | 1599 | 1356 | 1472 |
| GGAACTACGGCAGAAATTCG | PRKCB | 552 | 802 | 1477 | 1532 | 1698 |
| GTCATGCTCTCAGAGCGAAA | PRKCB | 472 | 371 | 640 | 611 | 611 |
| GTTTGTACTTACACAGCATG | PRKCB | 467 | 969 | 1611 | 1536 | 1516 |
| TGTCTGTAGAGATTTGGGAT | PRKCB | 580 | 652 | 1200 | 1328 | 1359 |
| ACTGAGATGATACTGTACAC | PRKCI | 589 | 416 | 934 | 886 | 903 |
| CCAGGACGTTCTGGTACACA | PRKCI | 552 | 621 | 1203 | 1334 | 1483 |
| CGGGTGAAAGCCTACTACCG | PRKCI | 613 | 619 | 1119 | 1118 | 1171 |
| TCAAGCCAAGCGTTTCAACA | PRKCI | 483 | 507 | 1118 | 989 | 1001 |
| TCGTCACAATCGAATGTGGG | PRKCI | 738 | 772 | 1614 | 1603 | 1646 |
| TTAAATTATCTTCATGAACG | PRKCI | 599 | 648 | 1136 | 1103 | 1062 |
| AGAGAGAACCCCATCACCAT | PRKCSH | 611 | 375 | 773 | 798 | 784 |
| AGCTCACTGAGCTACAAGCC | PRKCSH | 578 | 371 | 793 | 665 | 724 |
| CAGGGACAAATACCGGTCTG | PRKCSH | 439 | 816 | 1608 | 1668 | 1763 |
| GAAGCTGCCATTAGGACAGG | PRKCSH | 393 | 734 | 1175 | 1127 | 1136 |
| GCACCGATGAGTACAACAGC | PRKCSH | 633 | 428 | 767 | 786 | 764 |
| TGGCAGAGGTCACGCGAGAG | PRKCSH | 403 | 701 | 1308 | 1285 | 1277 |
| AATGTTCTCCTTGACGCCGA | PRKCZ | 331 | 552 | 911 | 844 | 813 |
| ACCTCAAGCCAGTTATCGAT | PRKCZ | 621 | 345 | 630 | 562 | 584 |
| ATTTGGGAGGACACAGTGCA | PRKCZ | 509 | 1006 | 1406 | 1345 | 1469 |
| CCTGCTTCCAGACAACAAGT | PRKCZ | 397 | 272 | 543 | 499 | 536 |
| CTAGAATCCATCTACCGCCG | PRKCZ | 621 | 597 | 1092 | 1036 | 1051 |
| GGTCATTGCTGCATCTAGGC | PRKCZ | 787 | 461 | 671 | 779 | 686 |
| ACAAAGTAAAAAAACGACGT | PRKDC | 830 | 303 | 613 | 574 | 652 |
| AGACTGTATAATCCACCAGT | PRKDC | 475 | 554 | 755 | 849 | 769 |
| AGAGCTAATTCTGTGACCCG | PRKDC | 446 | 690 | 1441 | 1128 | 1213 |
| CAAGACCCTAAATCTACCCA | PRKDC | 478 | 414 | 759 | 664 | 719 |
| GGTTCCTGAGGTGTATACAC | PRKDC | 645 | 356 | 847 | 655 | 738 |
| TGGCTAATCGCTTACAACTG | PRKDC | 322 | 700 | 1072 | 990 | 994 |
| ACGATGTCCTAATCCCAAAC | PRKN | 527 | 476 | 798 | 773 | 836 |
| AGTCTAAGCAAATCACATGG | PRKN | 415 | 628 | 1085 | 1018 | 957 |
| CACACGCAACTGGTCAACCG | PRKN | 475 | 469 | 669 | 624 | 663 |
| CACAGCTAGACCCACCGAGT | PRKN | 476 | 409 | 987 | 876 | 825 |
| CCCTGTCAAAGAATACAGCC | PRKN | 446 | 393 | 640 | 506 | 496 |
| TGTGATCAGGTTCAAAGCTA | PRKN | 708 | 803 | 1422 | 1259 | 1306 |
| AAGAACTACTGAGCGCACTG | PRLR | 386 | 559 | 908 | 899 | 963 |
| ACCAGATCTAACATCAACCA | PRLR | 507 | 229 | 477 | 472 | 475 |
| GCTCTTGGGAGTGTTAAATG | PRLR | 740 | 824 | 1445 | 1365 | 1340 |
| GGCCACAAAGATCCACACGA | PRLR | 411 | 423 | 677 | 594 | 606 |
| TAAGTCACGTCGACATAACG | PRLR | 393 | 457 | 874 | 789 | 761 |
| TGCCCAGACTACATAACCAG | PRLR | 390 | 368 | 737 | 765 | 683 |
| CAAAGCTGGGATTATCCATG | PRPF40B | 526 | 930 | 1581 | 1553 | 1526 |
| CATATTGGCAAAGCGGACAT | PRPF40B | 565 | 715 | 1117 | 1111 | 1006 |
| GAAAGAGTACAAGTCAGACA | PRPF40B | 498 | 327 | 562 | 559 | 531 |
| TCAATGCCTACAAGGCGCAG | PRPF40B | 631 | 530 | 1334 | 1242 | 1147 |
| TGAAGATTGCGATATGTCAG | PRPF40B | 418 | 644 | 1287 | 1284 | 1235 |
| TGGCCTCAGTTGGAGCAACC | PRPF40B | 370 | 447 | 815 | 781 | 756 |
| AAAGTCATTTGGCATCTGAG | PRPF8 | 329 | 581 | 1332 | 988 | 1113 |
| AACCGAGAACGAATCCGCCG | PRPF8 | 573 | 407 | 710 | 637 | 587 |
| ACCTACCTGAGGTCAGACTG | PRPF8 | 464 | 631 | 1149 | 1279 | 1255 |
| ATATTGTCAGCATAGTCCAA | PRPF8 | 814 | 424 | 801 | 661 | 684 |
| GAACCTGAACAACTACCGTA | PRPF8 | 617 | 294 | 544 | 348 | 380 |
| GGCGGGTACGACCAGAGCGT | PRPF8 | 552 | 431 | 905 | 590 | 662 |
| CCACCATACAGAGAGTCACG | PSEN1 | 426 | 532 | 944 | 960 | 972 |
| CGCAACAGTAATGTAGTCCA | PSEN1 | 619 | 690 | 1129 | 1149 | 1196 |
| CGGAAGATACTGAGACCGTG | PSEN1 | 606 | 622 | 1287 | 1230 | 1196 |
| GTTTCAACCAGCATACGAAG | PSEN1 | 442 | 439 | 504 | 470 | 480 |
| TGTATTTATACAGAACCACC | PSEN1 | 449 | 207 | 263 | 310 | 367 |
| TTATCTAACGGACGACCCCA | PSEN1 | 616 | 831 | 1380 | 1359 | 1345 |
| ACCGAACAGAGCCAAATCAA | PSENEN | 409 | 541 | 935 | 921 | 913 |
| CTGTGCCGGAAGTACTACCT | PSENEN | 414 | 469 | 876 | 968 | 1019 |
| CTTGGAGCGGGTGTCTAACG | PSENEN | 544 | 545 | 1135 | 1106 | 1069 |
| GGTGTATGCTGGGACAAGAA | PSENEN | 591 | 794 | 1236 | 1077 | 1122 |
| TGACCAACCAGAGAAAAGGC | PSENEN | 371 | 261 | 425 | 328 | 346 |
| TGAGTACAATCACCCAGAAG | PSENEN | 607 | 455 | 768 | 832 | 782 |
| CAACAAAACCATGACTACAG | PSMB1 | 507 | 283 | 503 | 322 | 353 |
| CGATTGCCAATACAGTACTG | PSMB1 | 575 | 451 | 963 | 855 | 784 |
| CTACGTTTACAACATCATTG | PSMB1 | 469 | 588 | 1003 | 899 | 911 |
| GTGTACAGCTTCGACCCAGT | PSMB1 | 505 | 357 | 742 | 564 | 681 |
| TCTGACACTCGATTGAGTGA | PSMB1 | 418 | 855 | 1644 | 1422 | 1471 |
| TGTCTACAATCCTCTACTCA | PSMB1 | 519 | 349 | 663 | 532 | 521 |
| AGGCGACGAGGACGTAGTCG | PSMB2 | 531 | 593 | 1379 | 1317 | 1212 |
| CATCTGGACAATACTGCTGG | PSMB2 | 504 | 603 | 1333 | 1015 | 1081 |
| CTCCAGCCTCTCCAACACAT | PSMB2 | 290 | 309 | 431 | 399 | 461 |
| GAGGAGGTTGACATGATACG | PSMB2 | 606 | 409 | 1034 | 914 | 1001 |
| GGGTCCAGCGCTCTACTACA | PSMB2 | 509 | 525 | 900 | 818 | 767 |
| TTTATAAGATGCGAAATGGT | PSMB2 | 427 | 476 | 1235 | 1033 | 1015 |
| AATCTATGAGCTCCGAAACA | PSMB5 | 486 | 302 | 662 | 578 | 542 |
| AGCCTCGTCGCGCCCAACTG | PSMB5 | 490 | 535 | 1367 | 1197 | 1231 |
| CTTCACTGTCCACATAGTAG | PSMB5 | 490 | 559 | 1024 | 844 | 823 |
| GCACGGTTATTTCGGACTCG | PSMB5 | 609 | 950 | 1598 | 1600 | 1635 |
| GGTGTATCAATATAAAGGCA | PSMB5 | 538 | 381 | 777 | 550 | 593 |
| TGATCTTAACAGTTTCAACA | PSMB5 | 634 | 272 | 403 | 303 | 307 |
| AAGACTATACTGGAGTCGGT | PSMD1 | 364 | 318 | 621 | 587 | 532 |
| AATGGCGACATATCTTCCCA | PSMD1 | 558 | 459 | 862 | 698 | 692 |
| AGGCTGCAAACTGCCGACTC | PSMD1 | 420 | 489 | 962 | 1013 | 1001 |
| ATGTTAATGACGATGTCCGG | PSMD1 | 402 | 623 | 1096 | 861 | 971 |
| GTACCCGTATTGGTGGATCC | PSMD1 | 514 | 589 | 1142 | 1040 | 1040 |
| TCCCCATGGCAGCTAAACCA | PSMD1 | 494 | 370 | 734 | 530 | 540 |
| ACCCTGTATGCCTTTAACAG | PSMD13 | 562 | 451 | 993 | 796 | 753 |
| AGCTCTAAAATTAAACATCG | PSMD13 | 613 | 449 | 880 | 767 | 763 |
| ATAGAAACGACTGTGAACCG | PSMD13 | 555 | 451 | 1003 | 782 | 928 |
| CACGTAGTTAGACAGATGAC | PSMD13 | 589 | 444 | 780 | 785 | 786 |
| GCAGGAGAGAGCTTTCACGC | PSMD13 | 447 | 459 | 844 | 763 | 754 |
| GTGGCATCAACTGACACTCC | PSMD13 | 548 | 365 | 738 | 649 | 674 |
| ATAGCAGTAGGATCCTGCAA | PSMD2 | 588 | 577 | 1116 | 901 | 907 |
| ATGATGCGGAAAAATCACAA | PSMD2 | 427 | 312 | 458 | 363 | 405 |
| CTAATGTGTTGGCAAAACTG | PSMD2 | 430 | 601 | 1249 | 861 | 901 |
| CTGTGGCATAGTGAACTCTG | PSMD2 | 421 | 449 | 874 | 777 | 797 |
| GTTTACCTTATTCTCCCCGG | PSMD2 | 583 | 315 | 632 | 498 | 522 |
| TCTGCTTCTGTACTACCCTG | PSMD2 | 417 | 400 | 997 | 898 | 892 |
| ACATCCACCATTCAAAACAA | PTCH1 | 786 | 294 | 453 | 402 | 420 |
| AGTGTCCACTTGGTGCAAGG | PTCH1 | 431 | 943 | 1578 | 1466 | 1507 |
| GGCTTGGCAGAGGACATATG | PTCH1 | 299 | 903 | 1677 | 1722 | 1672 |
| GTTATCGTGAGTCTCCGTGT | PTCH1 | 574 | 580 | 884 | 957 | 903 |
| TATACATATACACATGGACC | PTCH1 | 486 | 352 | 702 | 722 | 692 |
| TGATGTCGATGGGTTTATCG | PTCH1 | 409 | 1025 | 1582 | 1512 | 1585 |
| AAAGCCACCCTGCGTCACCA | PTCH2 | 458 | 593 | 1118 | 1215 | 1187 |
| CAGATAGCCTCCGACCACAC | PTCH2 | 526 | 384 | 700 | 608 | 596 |
| GACTACCAGACGCATGACAT | PTCH2 | 454 | 910 | 1173 | 1192 | 1201 |
| GGCAAAGGCTTGAACCGTGG | PTCH2 | 665 | 615 | 1222 | 1064 | 1114 |
| TGATACTTGCACTTTACTGG | PTCH2 | 390 | 681 | 1152 | 1046 | 1081 |
| TTCTAGGTCTGTCTCAACGA | PTCH2 | 428 | 537 | 864 | 874 | 869 |
| ACCGCCAAATTTAACTGCAG | PTEN | 430 | 400 | 684 | 778 | 697 |
| ACTATTCCAATGTTCAGTGG | PTEN | 511 | 580 | 1160 | 1136 | 1191 |
| CCAATTCAGGACCCACACGA | PTEN | 610 | 412 | 619 | 691 | 736 |
| TACACATAGCGCCTCTGACT | PTEN | 481 | 380 | 730 | 689 | 658 |
| TGTGCATATTTATTACATCG | PTEN | 545 | 576 | 836 | 856 | 955 |
| TTTGTGAAGATCTTGACCAA | PTEN | 499 | 641 | 1329 | 1203 | 1278 |
| AGATCTGGTTCAGTGAAGCC | PTGER3 | 526 | 419 | 760 | 799 | 829 |
| CTTAGCAGCAGATAAACCCA | PTGER3 | 757 | 434 | 1002 | 833 | 833 |
| CTTCTTAATAGCTGTTCGCC | PTGER3 | 512 | 546 | 686 | 726 | 652 |
| GAACCAGATCTTGGATCCCT | PTGER3 | 366 | 496 | 861 | 800 | 785 |
| GGCTTCACTGAACCAGATCT | PTGER3 | 499 | 585 | 1110 | 967 | 1003 |
| TCTTAGCAGCAGATAAACCC | PTGER3 | 457 | 334 | 631 | 543 | 586 |
| ACGTGCGGCGCATGAACTGG | PTGER4 | 488 | 823 | 1471 | 1585 | 1544 |
| CAGGTAGCGCTCGATACTCA | PTGER4 | 645 | 608 | 1180 | 1076 | 1007 |
| CCACTGGCCCTTCATGTACG | PTGER4 | 517 | 933 | 1642 | 1777 | 1587 |
| CCGAAGATGAACATCACCGC | PTGER4 | 516 | 423 | 655 | 525 | 621 |
| GCCCGGCCACTCACCACGAG | PTGER4 | 624 | 825 | 959 | 1083 | 1159 |
| GCTGAAGCCCGCGTACATGT | PTGER4 | 386 | 582 | 1114 | 998 | 962 |
| ACCTGAGTATCTTTGACGGT | PTGS2 | 532 | 695 | 1195 | 1356 | 1405 |
| AGTTTAACACACTCTACCAC | PTGS2 | 526 | 214 | 418 | 362 | 407 |
| GACTATGTACAACACTTGAG | PTGS2 | 695 | 859 | 1568 | 1320 | 1356 |
| GATGACTGTCCAACACCCAT | PTGS2 | 498 | 391 | 779 | 815 | 800 |
| GTGGCATACATCATCAGACC | PTGS2 | 679 | 342 | 593 | 579 | 612 |
| TTCAAGACAGATCATAAGCG | PTGS2 | 400 | 777 | 1391 | 1280 | 1409 |
| ATATGGGTGTCTCCAATGTG | PTK2 | 492 | 767 | 1536 | 1491 | 1387 |
| CATCCATACATCACTAGCTG | PTK2 | 513 | 458 | 957 | 806 | 888 |
| CGAGCTTGGACGCTGCATCG | PTK2 | 503 | 524 | 1200 | 1024 | 1055 |
| GAGAGCTGTACTAAGCTGAT | PTK2 | 616 | 542 | 792 | 661 | 698 |
| TCTGATGATAAAAGACTGCG | PTK2 | 493 | 521 | 961 | 810 | 886 |
| TTTATCAAATCTGTATACTG | PTK2 | 424 | 352 | 559 | 372 | 441 |
| ATCATGGAATTGTATCCCTA | PTK2B | 839 | 662 | 1040 | 1196 | 1187 |
| ATTACTGTGGACCTAGTCAT | PTK2B | 474 | 653 | 1287 | 1269 | 1318 |
| CATGCAACGCTATGCCAGCA | PTK2B | 415 | 656 | 1321 | 1228 | 1259 |
| GATGGGAAGACGAGTCACAG | PTK2B | 601 | 426 | 1063 | 956 | 1016 |
| GCCCGCAGTATGGTGTCACA | PTK2B | 416 | 874 | 1273 | 1242 | 1276 |
| TTGCACCCGCAGATGACAGT | PTK2B | 632 | 628 | 894 | 799 | 805 |
| CACCCTCAGCACCTTCATCG | PTP4A3 | 421 | 615 | 1119 | 1109 | 1159 |
| CTTGTAGCTCACCTCCACGG | PTP4A3 | 539 | 618 | 1086 | 1147 | 1054 |
| GCCAAGTTCTGTGACGACCC | PTP4A3 | 413 | 466 | 1016 | 859 | 909 |
| GGACTGGCCGTTTGACGACG | PTP4A3 | 466 | 1172 | 2064 | 2096 | 2110 |
| GTGCGAAGTGACCTACGACA | PTP4A3 | 585 | 526 | 924 | 945 | 973 |
| TTCGCACACTCGCACCACGG | PTP4A3 | 421 | 446 | 834 | 808 | 877 |
| ACCACAGATAAAGTCAAACA | PTPN11 | 533 | 304 | 492 | 395 | 444 |
| CACATCAAGATTCAGAACAC | PTPN11 | 571 | 268 | 455 | 381 | 372 |
| CTGACAACGGATCATGACAT | PTPN11 | 524 | 465 | 1075 | 896 | 893 |
| GTAGGATCTGCACAGTTCAG | PTPN11 | 536 | 682 | 1118 | 920 | 1000 |
| TCGTTGTCATGACAATCACT | PTPN11 | 594 | 539 | 983 | 824 | 858 |
| TGTGCCCAAAGTTTCCACCA | PTPN11 | 570 | 413 | 1011 | 907 | 844 |
| ACAAGTCAGCAGAACTTCCG | PTPN12 | 329 | 684 | 1258 | 1051 | 1050 |
| ACTGTGATGTACAATACCTG | PTPN12 | 639 | 752 | 1597 | 1739 | 1620 |
| AGAAAAAATGTGAACGCTAT | PTPN12 | 602 | 533 | 1061 | 1058 | 1118 |
| CAAATGATGTGGGAACATCA | PTPN12 | 612 | 483 | 1076 | 957 | 1023 |
| TTATGGGCCAAAAGCGTATG | PTPN12 | 459 | 687 | 1395 | 1558 | 1523 |
| TTTGTGCCATAGATTATACA | PTPN12 | 315 | 352 | 432 | 553 | 587 |
| AATAGATGGACTTTCACTGG | PTPN13 | 489 | 552 | 1174 | 972 | 1122 |
| ACAGGTCTGCCAAAGACTGA | PTPN13 | 451 | 742 | 1318 | 1184 | 1305 |
| ATCTCAGCGTGAACTGACAG | PTPN13 | 419 | 581 | 1047 | 1029 | 1025 |
| GCCAGGTTCGGTCACCAACA | PTPN13 | 343 | 778 | 1355 | 1394 | 1407 |
| GTGAATACCAGTGTAAGACA | PTPN13 | 581 | 512 | 909 | 839 | 870 |
| TTACTTACAAGGATAGACCG | PTPN13 | 319 | 838 | 1222 | 1313 | 1239 |
| CACAAGCATCACGTTGACAT | PTPRD | 810 | 369 | 510 | 586 | 517 |
| CAGAGCGAAGAATCTGACCA | PTPRD | 397 | 482 | 1015 | 819 | 920 |
| CCAGATCCATCGATACAACG | PTPRD | 451 | 298 | 499 | 503 | 501 |
| TCAATGGTAACTCGTTGCTG | PTPRD | 512 | 947 | 1670 | 1618 | 1691 |
| TGTGTGGCCTCAAATAACGT | PTPRD | 521 | 680 | 1146 | 1090 | 1058 |
| TTCATCAGTTCGAATCAACA | PTPRD | 522 | 346 | 509 | 403 | 440 |
| AAATAAATCAGCACTTCCAG | QKI | 381 | 344 | 662 | 579 | 600 |
| ATGGAGCTTGCAATTCTGAA | QKI | 437 | 817 | 1339 | 1246 | 1263 |
| CAAACTGAAAAGAGCCGTTG | QKI | 496 | 962 | 1687 | 1562 | 1639 |
| GCAGAATTGCCTGATGCTGT | QKI | 322 | 751 | 1133 | 974 | 982 |
| GGATGTAAAATAATGGTCCG | QKI | 418 | 913 | 1484 | 1544 | 1497 |
| TCTTCCAACAAAATTAAACT | QKI | 439 | 165 | 343 | 239 | 236 |
| AAACAGATCTTGCTGACAAG | RAB6A | 376 | 575 | 985 | 979 | 945 |
| AATTCAAGCTTGTGTTCCTG | RAB6A | 458 | 825 | 1434 | 1360 | 1376 |
| AGGAAAATAAAATACCTGAT | RAB6A | 452 | 387 | 671 | 683 | 655 |
| AGTTTACGATATCACAAGAT | RAB6A | 449 | 498 | 928 | 918 | 954 |
| ATCAAAAACAATGTACTTGG | RAB6A | 648 | 321 | 531 | 504 | 514 |
| CTACGGCTGCAGCAGAATCA | RAB6A | 424 | 844 | 1486 | 1483 | 1472 |
| AACCAGTGAATCTGGGCTTA | RAC1 | 787 | 921 | 1626 | 1679 | 1711 |
| ACAGTGGGGATGTATTCTCC | RAC1 | 572 | 487 | 1167 | 980 | 923 |
| AGTCCCCACCAAGATGATAG | RAC1 | 771 | 8 | 23 | 20 | 27 |
| AGTGTGTGGTGGTGGAAGAC | RAC1 | 487 | 697 | 1133 | 999 | 1083 |
| GACAGTGGTGTCGCACTTCA | RAC1 | 408 | 562 | 1091 | 905 | 991 |
| GATCCTTACTGTTTGCGGAT | RAC1 | 470 | 404 | 900 | 850 | 826 |
| AACAATTTGCCGGTTGTCGG | RACK1 | 547 | 584 | 1057 | 942 | 889 |
| AAGCTGAAGACCAATCACAT | RACK1 | 360 | 356 | 626 | 536 | 583 |
| GACCAACTTGTCCCAACCAC | RACK1 | 546 | 324 | 777 | 674 | 660 |
| TAGATAAGACGATCATCATG | RACK1 | 672 | 578 | 1176 | 945 | 881 |
| TCCTACAAATCTGCGTGTGG | RACK1 | 572 | 546 | 1197 | 1049 | 993 |
| TGTATGCAAGTACACCGTAC | RACK1 | 454 | 519 | 864 | 902 | 860 |
| AAACTTATAAGTAACAATGA | RAD21 | 513 | 300 | 517 | 466 | 485 |
| ATCAATAGGTTCCAATGCAA | RAD21 | 524 | 354 | 578 | 591 | 556 |
| ATGGAATAACAGACTACTGA | RAD21 | 479 | 324 | 598 | 549 | 506 |
| GGAGGGACATACTCTAAGTC | RAD21 | 627 | 579 | 995 | 853 | 904 |
| GTGTAATTTAGAGAGCAGTG | RAD21 | 649 | 356 | 896 | 623 | 623 |
| TCTGTTCAGGCTCTAACAGT | RAD21 | 451 | 541 | 1241 | 1087 | 1201 |
| AAGGTACCTTTAGGCCAGAA | RAD51 | 507 | 463 | 901 | 846 | 798 |
| AGAAGCTGGATTCCATACTG | RAD51 | 522 | 536 | 1138 | 910 | 852 |
| ATAATGTAGCATATGCTCGA | RAD51 | 307 | 439 | 806 | 682 | 778 |
| ATACCTAGATTCTACCATCA | RAD51 | 523 | 298 | 565 | 492 | 455 |
| GTCTAGCTTCCCATTGACCG | RAD51 | 466 | 412 | 956 | 704 | 777 |
| TGTTTGGAGAATTCCGAACT | RAD51 | 421 | 540 | 929 | 822 | 763 |
| CTCCAGTCCACACACATAGT | RAF1 | 641 | 610 | 1043 | 1108 | 1087 |
| GACAAGTACCTGATTCACTG | RAF1 | 584 | 506 | 914 | 833 | 867 |
| GACCATGTGGACATTAGGTG | RAF1 | 582 | 1027 | 1341 | 1340 | 1292 |
| GCCAGCTTCCTACTTACCGA | RAF1 | 513 | 398 | 705 | 621 | 612 |
| GGAGCCTGACCCAATCCGAG | RAF1 | 541 | 594 | 1138 | 1130 | 1060 |
| TTCTTCTCCAATCAACGAAG | RAF1 | 386 | 423 | 594 | 534 | 496 |
| ACGGCTACTACATCCAAGGT | RAN | 551 | 282 | 734 | 518 | 584 |
| CAGGTGACGCTTCACGAACG | RAN | 458 | 564 | 1312 | 1018 | 1061 |
| CTCGTGTTCCACACCAACAG | RAN | 560 | 392 | 913 | 711 | 685 |
| GCCGAACTTCTCCTGGCCGG | RAN | 457 | 378 | 819 | 630 | 610 |
| TAATGTTTGACGTAACGTCA | RAN | 398 | 621 | 1070 | 970 | 1032 |
| TTTATTGCCACAGAGCACAA | RAN | 488 | 239 | 352 | 335 | 336 |
| ACACCATGACCTTCTCAGAT | RARA | 331 | 466 | 924 | 874 | 939 |
| AGAGTGGTCAGCGCGCCTGG | RARA | 558 | 846 | 1623 | 1678 | 1710 |
| CAAGTGCATCATTAAGACTG | RARA | 395 | 498 | 1014 | 1008 | 954 |
| CCAGCTGGGCAAATACACTA | RARA | 427 | 464 | 811 | 762 | 740 |
| GAGCTACACGCTGACGCCCG | RARA | 506 | 596 | 1386 | 1209 | 1246 |
| GTATCATCAATAAGGTGACC | RARA | 378 | 324 | 670 | 678 | 564 |
| CATCGAGTCCGACTGGACCT | RARB | 606 | 675 | 1086 | 1150 | 1117 |
| CTTCACCGGCTTGACCATCG | RARB | 540 | 731 | 1160 | 1072 | 1078 |
| GCTCTGTGTTTCAATCGCTG | RARB | 598 | 758 | 1333 | 1346 | 1411 |
| GTGTTATTAATAAAGTCACC | RARB | 553 | 216 | 420 | 395 | 333 |
| TCTGCCAGGACAAATCATCG | RARB | 663 | 562 | 1141 | 1157 | 1112 |
| TGGTGGAAAGAGCAATGGGT | RARB | 446 | 476 | 898 | 899 | 860 |
| AAGCACGGCTTGTAGACCCG | RARG | 614 | 755 | 1220 | 1168 | 1181 |
| CAAAAACTGCATCATCAACA | RARG | 482 | 345 | 514 | 563 | 542 |
| CAAGTGCATCATCAAGATCG | RARG | 663 | 640 | 1085 | 1196 | 1281 |
| GGTCATGGTGTCCTGTTCTG | RARG | 595 | 783 | 1561 | 1661 | 1636 |
| GTGGGCATGTCCAAAGAAGG | RARG | 511 | 689 | 927 | 1003 | 1065 |
| GTTAGAAGAGCTCATCACCA | RARG | 538 | 668 | 937 | 966 | 1098 |
| AAGCAAAAACCATTCATCTG | RASA1 | 592 | 994 | 1433 | 1541 | 1575 |
| AATCTAAGAACAGATGAACA | RASA1 | 472 | 482 | 825 | 703 | 758 |
| ATTTACGGAAAAGTAGTCCA | RASA1 | 354 | 668 | 1169 | 1043 | 977 |
| GGATGGACCCGAATACGAGG | RASA1 | 470 | 658 | 1039 | 1087 | 1091 |
| GTCACCAGGAGTATTATCTG | RASA1 | 419 | 1253 | 1709 | 1601 | 1591 |
| TGTGGAGATTACTACATTGG | RASA1 | 526 | 1420 | 2370 | 2188 | 2221 |
| AGGGGAAGTGTTACAAATGG | RB1 | 574 | 850 | 1484 | 1592 | 1593 |
| CAAAGGAAAAAGGAGCTATG | RB1 | 516 | 657 | 1249 | 1194 | 1066 |
| GCAGTGTGATTACTCTGGAG | RB1 | 415 | 525 | 1037 | 1020 | 971 |
| TCACCTCGAACACCAAGGCG | RB1 | 505 | 632 | 1092 | 960 | 994 |
| TTCAGATTGATCCTACAGAC | RB1 | 600 | 279 | 456 | 528 | 526 |
| TTTGCTAAAGCTGTGGGACA | RB1 | 450 | 805 | 1286 | 1303 | 1208 |
| AAAATGGAAAACTTCAGACA | RBL2 | 611 | 338 | 455 | 443 | 467 |
| ACAGGGACAGCATTGACGAG | RBL2 | 493 | 609 | 1225 | 1282 | 1359 |
| CAGAGTTTCTCAATGACACA | RBL2 | 545 | 561 | 937 | 933 | 858 |
| CAGGAACAGAGACTGCTGAA | RBL2 | 421 | 714 | 1285 | 1471 | 1413 |
| CATGAGCGAGAGCTACACGC | RBL2 | 674 | 252 | 445 | 495 | 482 |
| GAACAGGAACAGAAAAGACT | RBL2 | 387 | 316 | 372 | 437 | 396 |
| AAGCAGACCCAACTGAACCG | RBM10 | 649 | 486 | 845 | 780 | 802 |
| AGAAAGCCTGCAGTGACACG | RBM10 | 609 | 1134 | 1565 | 1803 | 1655 |
| CAAGTGACTAAACTCGACGA | RBM10 | 682 | 467 | 932 | 1099 | 955 |
| CATACTCTCGGGGATATGAG | RBM10 | 589 | 567 | 1351 | 1435 | 1319 |
| CGTGAGAAGTGCTTCAAATG | RBM10 | 538 | 759 | 1472 | 1315 | 1356 |
| TAGTAGCTGTAGTCGACCGG | RBM10 | 510 | 686 | 1423 | 1269 | 1333 |
| ACATCGGAAGCAAGAATCGT | RECQL4 | 327 | 520 | 1061 | 989 | 964 |
| AGTGGGGTCACCGTGAACCG | RECQL4 | 495 | 935 | 1474 | 1350 | 1440 |
| ATCGTCTACTGCCACAGACG | RECQL4 | 438 | 539 | 1201 | 1059 | 1194 |
| GCTGTGACTCCTCTACACCC | RECQL4 | 644 | 260 | 528 | 442 | 476 |
| TCGGACGAGAGTCTCCGTGG | RECQL4 | 532 | 550 | 1284 | 1049 | 1075 |
| TGCCACCACTCTACCCACCA | RECQL4 | 415 | 506 | 895 | 808 | 956 |
| ACCCTAGTATCAACTGGAGA | REL | 540 | 512 | 880 | 845 | 937 |
| ATGTGACAGTGCATTTGACA | REL | 461 | 815 | 1442 | 1325 | 1308 |
| GAAATATTTCATCACCACCT | REL | 515 | 382 | 604 | 585 | 587 |
| GTGACTATGTACTTACAACA | REL | 400 | 287 | 480 | 428 | 421 |
| GTTGCGGAGACCTTCTGACC | REL | 414 | 418 | 873 | 793 | 757 |
| TGAAGATTGCGACCTCAACG | REL | 447 | 723 | 1156 | 1274 | 1302 |
| CACAAGTACACTACTCCCTG | RET | 441 | 606 | 1274 | 1318 | 1215 |
| CGGCAGAGCTCATCACACAG | RET | 567 | 839 | 1046 | 1162 | 1192 |
| GAACGACTGGATCCGCATCG | RET | 522 | 757 | 1458 | 1402 | 1391 |
| GAGGCCTGTTCTCTCGAATG | RET | 517 | 802 | 1460 | 1294 | 1280 |
| TGCTCTCTACAGCATCACAG | RET | 544 | 634 | 1034 | 1047 | 1057 |
| TTCTCAACTCCAGTACACAG | RET | 365 | 480 | 539 | 541 | 536 |
| GCAAAAGCAACGACTTCGTG | RFWD2 | 472 | 714 | 1139 | 1156 | 1157 |
| GCACTAGTAATTCCAACATG | RFWD2 | 474 | 922 | 1375 | 1406 | 1475 |
| GCTAATGTGCTGTTATACCA | RFWD2 | 502 | 501 | 925 | 858 | 949 |
| GCTGAGCCGAGACTGGCTCG | RFWD2 | 378 | 906 | 1503 | 1575 | 1651 |
| TAGTCGAACTGCAAGCCAGT | RFWD2 | 460 | 499 | 831 | 763 | 857 |
| TCCAAAGTGTAACTATGTTG | RFWD2 | 540 | 899 | 1626 | 1543 | 1411 |
| AACAAACTGAATCGTCAATG | RHEB | 413 | 712 | 1317 | 1414 | 1353 |
| ATATCTATGGAGTATGTCTG | RHEB | 590 | 1078 | 1858 | 1742 | 1733 |
| GAATACCTATTATGTTGGTT | RHEB | 319 | 609 | 1061 | 1058 | 1037 |
| GGCAAATTGTTGGACATGGT | RHEB | 434 | 574 | 985 | 1040 | 1025 |
| TCCCGGAAGATCGCGATCCT | RHEB | 501 | 523 | 947 | 936 | 950 |
| TTCAACTTGTCGACACAGCT | RHEB | 403 | 458 | 807 | 819 | 738 |
| ACAGAAGTGCTTGACTTCTG | RHOA | 319 | 509 | 1106 | 957 | 1001 |
| ACATAGTTCTCAAACACTGT | RHOA | 347 | 388 | 779 | 673 | 704 |
| AGGAAGATTATGATCGCTTG | RHOA | 578 | 778 | 1428 | 1515 | 1409 |
| GAGCAAACAAGTCTTACCAC | RHOA | 636 | 292 | 607 | 529 | 565 |
| GCTACTCACCTAAGCTATCA | RHOA | 537 | 410 | 707 | 599 | 607 |
| TTCGGAATGATGAGCACACA | RHOA | 480 | 1011 | 1413 | 1504 | 1481 |
| AGTGAAGCGCACCAGCAGAG | RHOH | 456 | 719 | 1114 | 1309 | 1383 |
| CATTACCGGCTGTGTCCCAG | RHOH | 418 | 485 | 1276 | 1029 | 1034 |
| CCCACGGTGTATGAGAACAC | RHOH | 451 | 588 | 958 | 1043 | 1047 |
| GAGCAACTTGCCCTGCACCC | RHOH | 379 | 534 | 1011 | 903 | 930 |
| GTAGCACATCAGCACCACGT | RHOH | 470 | 408 | 720 | 696 | 743 |
| TGGCTACCCAGACTGACCAG | RHOH | 319 | 644 | 1127 | 1005 | 1093 |
| AGTGGGAGTCTATCAGAGCA | RHPN2 | 607 | 901 | 1567 | 1281 | 1260 |
| CAGAGTAGCTACGAACCACA | RHPN2 | 494 | 510 | 827 | 797 | 782 |
| CGTGAAGGCCCACCACTACG | RHPN2 | 583 | 816 | 1244 | 1243 | 1229 |
| CTCTACGACCACATGCCAGA | RHPN2 | 500 | 504 | 565 | 505 | 564 |
| GCGGGACCGCTCGTGCGCCA | RHPN2 | 477 | 627 | 972 | 1004 | 1085 |
| GCTCATGAGCAGTTCAACCC | RHPN2 | 683 | 379 | 834 | 784 | 764 |
| AGCACTTCGATTAGTCAGAA | RICTOR | 450 | 715 | 1156 | 1148 | 1041 |
| ATGTTCGTCGGAGTAACCAA | RICTOR | 512 | 656 | 1175 | 1057 | 1113 |
| GCCTGTTATACTCACATCAG | RICTOR | 623 | 699 | 1367 | 1227 | 1239 |
| TACCTGGATCTGGCACGATG | RICTOR | 694 | 783 | 983 | 953 | 1010 |
| TCTGTAGTGAAAATCAGTAT | RICTOR | 430 | 409 | 801 | 715 | 731 |
| TGTGAGGTTCTTTCAATCAG | RICTOR | 556 | 853 | 1418 | 1217 | 1175 |
| AGCAAACCAGGACTCTTCCA | RIPK1 | 497 | 637 | 900 | 849 | 920 |
| CAGCATCATGAAGCAATGCT | RIPK1 | 416 | 498 | 1031 | 982 | 1003 |
| GTGGAGTAAACTGACTAAAG | RIPK1 | 527 | 614 | 913 | 989 | 1064 |
| TATGAAAACATTCAGAACCC | RIPK1 | 732 | 371 | 636 | 594 | 683 |
| TGGAGAAGGCGTAATACACA | RIPK1 | 362 | 787 | 1129 | 1111 | 1170 |
| TTTCAGGATTACGAGTCCAT | RIPK1 | 404 | 616 | 842 | 891 | 1005 |
| ATCTCTTGCTAAGACCGTCG | RNF213 | 614 | 548 | 1279 | 1248 | 1244 |
| ATGAGTCGGATAACTTTCAG | RNF213 | 545 | 342 | 709 | 719 | 678 |
| CAGCTTATACAGCAGCGACG | RNF213 | 543 | 556 | 899 | 736 | 732 |
| CAGGTACACAAGCTGCTCGG | RNF213 | 666 | 650 | 1126 | 1048 | 1073 |
| GAAAGTGGAGCAGCCTACGT | RNF213 | 732 | 264 | 753 | 576 | 563 |
| TAGGGACCAAGAGTACACCC | RNF213 | 497 | 477 | 827 | 847 | 893 |
| CTAACCACTCCGATTCACCA | ROBO1 | 452 | 310 | 568 | 495 | 453 |
| CTGTGCCTACGGTACGATGG | ROBO1 | 588 | 1041 | 1604 | 1607 | 1587 |
| GCATCTGGCTCTATCGACAC | ROBO1 | 690 | 475 | 948 | 913 | 929 |
| GCTGCAGATCCGATACGCTA | ROBO1 | 458 | 388 | 698 | 661 | 621 |
| GGCACGGAATTACCTCGGAG | ROBO1 | 306 | 630 | 1369 | 1321 | 1218 |
| TCGTTATCACATAAACAAGA | ROBO1 | 404 | 389 | 505 | 434 | 361 |
| ACCACGCTCTATACAGTGAG | ROBO2 | 525 | 856 | 1555 | 1563 | 1631 |
| ATTCCATATCAACAAAACTG | ROBO2 | 566 | 373 | 461 | 440 | 371 |
| CAGTTCCAGCCAAATAACAC | ROBO2 | 602 | 203 | 404 | 306 | 382 |
| CGGTAGCCTTGAATAAACTG | ROBO2 | 558 | 763 | 1298 | 1350 | 1377 |
| GAGGACTAATGAGCAATGGG | ROBO2 | 542 | 633 | 1338 | 1366 | 1326 |
| TATCTGCCAGGCATTAACTG | ROBO2 | 468 | 598 | 1029 | 888 | 884 |
| ACCATCTCGAGACATCCCTA | RORA | 11 | 475 | 820 | 843 | 917 |
| AGTTGGGGAAGTCTCTCCGT | RORA | 530 | 467 | 914 | 803 | 886 |
| GACCTCAGCAACTACATCGA | RORA | 430 | 499 | 934 | 891 | 856 |
| GGAGCTCTTCTCTCAAGTAT | RORA | 634 | 753 | 1197 | 1318 | 1271 |
| GGCGGATTCCTGGTTCAGCG | RORA | 537 | 833 | 1211 | 1328 | 1333 |
| TATGGTGTCATTACATGTGA | RORA | 294 | 737 | 1388 | 1396 | 1450 |
| ACTTACAAACATTCTGTACC | RORC | 503 | 327 | 747 | 618 | 562 |
| CACAGGTGATCACCCCGTAG | RORC | 445 | 688 | 1379 | 1334 | 1416 |
| CGAGGCCATTCAGTACGTGG | RORC | 496 | 879 | 1297 | 1322 | 1306 |
| CTCATCACCTGACCTACCCG | RORC | 410 | 538 | 870 | 864 | 762 |
| CTGGAATACAGCCCCCAGCG | RORC | 435 | 680 | 1227 | 1334 | 1288 |
| GACATCGGTTTCGGCTGGTG | RORC | 651 | 777 | 1298 | 1358 | 1351 |
| CACCAGGGTAGTACCCACCG | ROS1 | 659 | 661 | 1372 | 1300 | 1171 |
| CGTTGACATGAAGACGTGAG | ROS1 | 532 | 785 | 1364 | 1390 | 1501 |
| CTGCCAGGACAAATCTCCCG | ROS1 | 384 | 500 | 977 | 932 | 928 |
| GCTCTCGACTGAGTACAGTG | ROS1 | 487 | 830 | 1480 | 1339 | 1509 |
| GCTGAAATGACATCTCCGAG | ROS1 | 708 | 547 | 933 | 764 | 797 |
| TCCATCGTAACGTCACACTG | ROS1 | 656 | 756 | 1187 | 1155 | 1177 |
| AAGTACATGGTGAAAAGCTG | RPL10 | 526 | 414 | 1017 | 770 | 768 |
| AGATTCGCATCTTTGATCTG | RPL10 | 453 | 590 | 1241 | 934 | 926 |
| CATACTCATCTGACACCATG | RPL10 | 420 | 431 | 520 | 445 | 365 |
| CCTCTGCAGAAGCGTGACTT | RPL10 | 572 | 505 | 965 | 780 | 790 |
| GTTGATACGGATAACATGGA | RPL10 | 556 | 478 | 1098 | 810 | 811 |
| TGTCATCACCCTAGCCCTGG | RPL10 | 589 | 461 | 1175 | 847 | 880 |
| AAGGATGACGAAGTGCAAGT | RPL26 | 522 | 291 | 600 | 471 | 499 |
| ACCGGAGCAAGAATCGTAAG | RPL26 | 477 | 395 | 922 | 700 | 692 |
| GCAAATCGGCAAAGTAGTCC | RPL26 | 429 | 407 | 756 | 649 | 634 |
| TGAACGAGTACAGCGAGAGA | RPL26 | 309 | 411 | 865 | 653 | 658 |
| TGAGGCAGAAGTACAACGTG | RPL26 | 498 | 410 | 888 | 769 | 744 |
| TGCCTCAACTCTTTGGAAAG | RPL26 | 483 | 568 | 1456 | 1339 | 1374 |
| CATTTGTCATCAGGATCAGA | RPL7 | 414 | 356 | 567 | 458 | 497 |
| GAAAAAGCGAAGGAACTTCG | RPL7 | 627 | 358 | 761 | 632 | 641 |
| GTGGAACCGTATATAGCATG | RPL7 | 671 | 522 | 1077 | 787 | 815 |
| TCCAGCTTCGAAAGGCAAGG | RPL7 | 477 | 365 | 907 | 622 | 632 |
| TCTTGGTTTCCAGCTTCGAA | RPL7 | 489 | 415 | 759 | 787 | 704 |
| TGTGGAACCGTATATAGCAT | RPL7 | 358 | 394 | 873 | 661 | 653 |
| AGCGCGGCCACTTGACAAAG | RPL7A | 368 | 438 | 938 | 819 | 778 |
| AGGCCCAAGAACTTCGGCAT | RPL7A | 510 | 286 | 674 | 594 | 576 |
| GGTAGTGATAGCCCACGACG | RPL7A | 658 | 517 | 998 | 881 | 853 |
| TGCCATTAACCAGTTCACCC | RPL7A | 602 | 241 | 473 | 299 | 280 |
| TGCCGTGGTGAAGAAGCAGG | RPL7A | 482 | 403 | 810 | 691 | 655 |
| TGGTGACAGTGTTAACCCCT | RPL7A | 315 | 386 | 920 | 816 | 814 |
| AGTGTAGTGCAGCTCATCTG | RPN1 | 468 | 719 | 1602 | 1511 | 1435 |
| CAGCATGACCCGAGTCATTG | RPN1 | 555 | 724 | 1419 | 1295 | 1441 |
| GGCCAAGGTCTCAGTCGTTG | RPN1 | 544 | 862 | 1648 | 1439 | 1505 |
| GGGAAACCCTACGCGCTCTG | RPN1 | 466 | 764 | 1323 | 1368 | 1414 |
| TACCCCATACCTTAAATGAG | RPN1 | 419 | 520 | 1082 | 973 | 943 |
| TCTTCCACCCGCCAAACAGA | RPN1 | 600 | 363 | 583 | 569 | 589 |
| ATTCGCACCACGTAACCCTG | RPS6 | 510 | 394 | 790 | 599 | 568 |
| CAGCAGGCGGACGCGGCCGT | RPS6 | 595 | 336 | 844 | 657 | 618 |
| CTCGTACCTTCTTTGTTTAG | RPS6 | 318 | 1156 | 1917 | 1821 | 1815 |
| CTGGGCGAGGAATGGAAGGT | RPS6 | 695 | 309 | 814 | 567 | 571 |
| GGTTATTGTGAAAAAAGGTG | RPS6 | 567 | 585 | 1170 | 727 | 802 |
| TTGTGTCCCACAGGGTTACG | RPS6 | 570 | 560 | 1420 | 1014 | 1100 |
| AAATGAAAGCATGGACCATG | RPS6KB1 | 491 | 560 | 956 | 950 | 1051 |
| AAGCGGCCACAATCGTGCTG | RPS6KB1 | 483 | 625 | 1186 | 1113 | 1110 |
| AAGTAACAGGAGCAAATACT | RPS6KB1 | 374 | 618 | 1099 | 1092 | 1045 |
| AATATCATGCTTAATCACCA | RPS6KB1 | 556 | 279 | 334 | 320 | 367 |
| ACTGACCGCTGAGATACTCA | RPS6KB1 | 691 | 655 | 1222 | 1041 | 1046 |
| GAGATCTGCTTAAAAAGGTA | RPS6KB1 | 582 | 576 | 882 | 878 | 796 |
| AATGGTGCCACAGAAAGTAT | RPS6KB2 | 492 | 947 | 1600 | 1487 | 1546 |
| ACTGAGACCAGTGTGAACCT | RPS6KB2 | 555 | 807 | 1150 | 1102 | 1099 |
| AGTGAGCATGTCATACATCA | RPS6KB2 | 516 | 586 | 959 | 865 | 871 |
| CCAAGGACACAGCACACACG | RPS6KB2 | 511 | 782 | 1403 | 1241 | 1187 |
| CCGGGCTGTCAGTCTGAGCG | RPS6KB2 | 499 | 678 | 1225 | 1198 | 1350 |
| GGCCCCCACTCATACCACTG | RPS6KB2 | 661 | 703 | 1099 | 1027 | 1028 |
| AGCTGTTGACGATCACAGCG | RPTOR | 354 | 1009 | 1533 | 1458 | 1427 |
| CCACTTACATGCACGCGATG | RPTOR | 493 | 680 | 1369 | 1402 | 1409 |
| CTTCAGATCCTCTGTCGATG | RPTOR | 383 | 885 | 1385 | 1486 | 1427 |
| GACCACCATCGACCACAACG | RPTOR | 564 | 409 | 747 | 648 | 677 |
| GAGGCACGACGTGAACAGGT | RPTOR | 433 | 458 | 745 | 770 | 773 |
| TGCAGGTCATATATAGACAG | RPTOR | 344 | 555 | 1295 | 1292 | 1162 |
| AGATCTGTTCATGAAACGAG | RRM1 | 485 | 354 | 824 | 710 | 702 |
| ATCTTTGTAGAGCATGTACG | RRM1 | 427 | 564 | 1170 | 910 | 954 |
| CTTGTACCCCAATTCCAATG | RRM1 | 677 | 544 | 1028 | 905 | 825 |
| GTACCAGCATTAAACAGAGT | RRM1 | 534 | 307 | 619 | 493 | 487 |
| TATTTGCCTTTAGCGACCTG | RRM1 | 522 | 571 | 1139 | 1009 | 1047 |
| TTGATTTCCAAGTCTGCTGG | RRM1 | 603 | 494 | 1181 | 942 | 1029 |
| ACCCAGATGGACAGGAATGC | RSPO2 | 394 | 583 | 1082 | 1003 | 1001 |
| AGAACAACTTCTGTTGACAT | RSPO2 | 751 | 570 | 1001 | 936 | 929 |
| CTGGACATTCATCAAAGCAA | RSPO2 | 583 | 475 | 558 | 515 | 541 |
| GATGAAACCATGGAATGTGT | RSPO2 | 383 | 515 | 836 | 912 | 866 |
| GCCACTGCCAAGGCAACCGA | RSPO2 | 632 | 532 | 916 | 861 | 772 |
| GTATCAAATCCCATTTGCAA | RSPO2 | 573 | 332 | 422 | 438 | 437 |
| AGGCCAGTGAATGGAGTCCG | RSPO3 | 423 | 791 | 1448 | 1612 | 1433 |
| CAAAAGAGGGACTGAAACAC | RSPO3 | 397 | 386 | 760 | 833 | 821 |
| GGAAGCCAACAACCATACCA | RSPO3 | 538 | 341 | 619 | 501 | 468 |
| GGCCTTATTATCTTACACGT | RSPO3 | 367 | 314 | 483 | 446 | 470 |
| TCCATTGTAATCTGAACACG | RSPO3 | 530 | 865 | 1538 | 1452 | 1435 |
| TGTCTCTCTTCATGTCCAAG | RSPO3 | 436 | 740 | 1253 | 1185 | 1212 |
| AACACCCATCTCACCTGGAC | RUNX1 | 521 | 216 | 407 | 388 | 361 |
| ATTGATCATAGGACCATGGT | RUNX1 | 647 | 529 | 831 | 756 | 872 |
| CACTCCGACCGACAAACCTG | RUNX1 | 408 | 556 | 1211 | 1117 | 1175 |
| CAGAGCCATCAAAATCACCG | RUNX1 | 433 | 430 | 767 | 593 | 672 |
| CAGTGACCAGAGTGCCGTCG | RUNX1 | 452 | 746 | 1390 | 1448 | 1387 |
| GCTGCGGCGAACAGCCATGA | RUNX1 | 510 | 806 | 1320 | 1275 | 1187 |
| AATGCCTACTCACAAGTCGT | RUNX1T1 | 497 | 573 | 960 | 1100 | 1127 |
| ATCCCTTCAGTCTTACCCAT | RUNX1T1 | 356 | 471 | 704 | 735 | 761 |
| CTGTCGCTCCTCTTAGCTGG | RUNX1T1 | 571 | 682 | 1289 | 1258 | 1245 |
| GAGTCCTGTCTGGGTCGCAG | RUNX1T1 | 489 | 520 | 1141 | 1016 | 1077 |
| GGAAGAGGCGAACTCCAGAC | RUNX1T1 | 553 | 424 | 691 | 690 | 718 |
| TACCACCAGTCCTAGTACGA | RUNX1T1 | 538 | 492 | 989 | 880 | 840 |
| AGGACTGCCTGATCGACAAG | RXRA | 476 | 923 | 1522 | 1572 | 1602 |
| CAAGGACCGGAACGAGAATG | RXRA | 515 | 976 | 1736 | 1797 | 1686 |
| CGTACGTGGACGCGAACATG | RXRA | 464 | 660 | 1206 | 1228 | 1220 |
| GCTCTTCACTCTGGTCGAGT | RXRA | 460 | 495 | 904 | 928 | 972 |
| GGACCTTGAGGACACCATTG | RXRA | 587 | 863 | 1601 | 1548 | 1477 |
| TACGGAGTGTATAGCTGTGA | RXRA | 569 | 779 | 1452 | 1435 | 1450 |
| AGCCGGAGAACAACAAACCC | RXRB | 402 | 475 | 1002 | 903 | 925 |
| CCCCGAATCCCCTTTCCCAG | RXRB | 454 | 439 | 759 | 633 | 701 |
| CCCCGGACCCCTAAGACTGG | RXRB | 487 | 539 | 1180 | 1198 | 1228 |
| CCTGACAGATGTTAGTCACA | RXRB | 485 | 805 | 1582 | 1590 | 1601 |
| GTAGGTCAGATCCTTACGGA | RXRB | 461 | 376 | 808 | 665 | 670 |
| GTGCGAAAAGAAATGCATTG | RXRB | 537 | 500 | 1073 | 893 | 927 |
| AAAGTACCTGACGACAAGAC | RXRG | 376 | 358 | 578 | 499 | 554 |
| AGCACCTCCAGGAATCAACC | RXRG | 646 | 256 | 396 | 383 | 389 |
| CACTGCACTCAGAGTCCGCG | RXRG | 562 | 757 | 1207 | 1240 | 1193 |
| GAGGCAGACTGTGCCAACAG | RXRG | 380 | 528 | 1098 | 1115 | 1232 |
| GCCTCATTGACAAGCGCCAG | RXRG | 495 | 454 | 935 | 842 | 846 |
| TACGGTGTGTACAGTTGTGA | RXRG | 394 | 882 | 1603 | 1472 | 1464 |
| ACAAGCTCACCCCCGCTCAG | S1PR1 | 401 | 523 | 1103 | 960 | 1091 |
| CCAGCGACTTCTCGGAGCTG | S1PR1 | 640 | 906 | 1792 | 1820 | 1747 |
| CGATGAAATAGTACATAGGT | S1PR1 | 505 | 430 | 987 | 1023 | 974 |
| CTGAAGATGAAACTCCACAA | S1PR1 | 239 | 767 | 900 | 894 | 868 |
| TCAACTACGATATCATCGTC | S1PR1 | 460 | 372 | 514 | 691 | 677 |
| TGGCATTAAAATGAGCTCGG | S1PR1 | 474 | 451 | 972 | 951 | 1042 |
| AATGGCCAGCAAGCTACAGG | S1PR3 | 505 | 605 | 1045 | 1057 | 1031 |
| CAAGGTCAACATTCTGATGT | S1PR3 | 546 | 588 | 1082 | 1243 | 1215 |
| CACGAGCACTACAACTACGT | S1PR3 | 499 | 252 | 391 | 425 | 406 |
| CCAGCCCCCACAACTCGGAG | S1PR3 | 367 | 457 | 1053 | 1045 | 984 |
| GCGCGCGTACAGGATCACGA | S1PR3 | 372 | 371 | 585 | 523 | 509 |
| TGCCAATCAGAAGGAAGACG | S1PR3 | 405 | 567 | 1242 | 1218 | 1235 |
| AACGTTTACGAACACTGAGT | SBDS | 599 | 366 | 855 | 658 | 735 |
| ACGTTGATGCTTTAGCGACC | SBDS | 524 | 496 | 984 | 1026 | 949 |
| CGGGGAAAGACGGCGTTGAG | SBDS | 442 | 834 | 1449 | 1591 | 1603 |
| GCGGGAACTACACTGCGTCA | SBDS | 364 | 669 | 1186 | 1228 | 1257 |
| TCGTAAAAGCCACAGTGCGC | SBDS | 453 | 418 | 795 | 720 | 705 |
| TTTACAGTGTGTCTCAACTG | SBDS | 563 | 773 | 1234 | 1149 | 1247 |
| ACAGGGACCGAAATCAGCTG | SCARB1 | 490 | 728 | 1252 | 1328 | 1466 |
| ATAGTCGCTCTCTAAGCCAT | SCARB1 | 489 | 715 | 1343 | 1222 | 1185 |
| CATCCTCACCTGTACACGTA | SCARB1 | 454 | 540 | 1193 | 1215 | 1217 |
| CATGAAGGCACGCTCCCCAA | SCARB1 | 585 | 662 | 1098 | 1026 | 1005 |
| CATTCCACTTGTCCACGAGG | SCARB1 | 492 | 420 | 831 | 866 | 855 |
| GGGCCACGAAGCGATAGGTG | SCARB1 | 343 | 624 | 1232 | 1003 | 1080 |
| ATGACTCTGGAAATCCGTGA | SDHA | 735 | 489 | 796 | 734 | 759 |
| CAGGTTCTGAGGCACGTGAA | SDHA | 492 | 1026 | 1830 | 1719 | 1609 |
| CTACCTCTACCACAGAAGCA | SDHA | 420 | 534 | 784 | 808 | 854 |
| GATCTCTACGCAGTACCCAG | SDHA | 569 | 509 | 750 | 771 | 806 |
| GCAACGGTGTGTGATCTGGT | SDHA | 517 | 676 | 1202 | 1174 | 1110 |
| GCTGCACACGTTGTATGGAA | SDHA | 489 | 677 | 1205 | 1019 | 1132 |
| CATTAATCAGGCGATCATAG | SDHAF2 | 504 | 507 | 738 | 725 | 682 |
| CCTCTGTAGAAACGGTTGGA | SDHAF2 | 288 | 762 | 1592 | 1564 | 1478 |
| GCAGATACTCTTTAGCAAAG | SDHAF2 | 532 | 380 | 659 | 579 | 593 |
| TCGGTTCGCTCCTGCCATGG | SDHAF2 | 553 | 838 | 1491 | 1274 | 1345 |
| TGCTCTACGAGAGCAGAAAG | SDHAF2 | 348 | 838 | 1481 | 1340 | 1286 |
| TGCTGAGCAAAGGAGACACT | SDHAF2 | 483 | 384 | 749 | 711 | 835 |
| ATTAAAGCATCCAACACCAT | SDHB | 526 | 253 | 425 | 492 | 459 |
| GAAATTTGCCATCTACCGAT | SDHB | 340 | 284 | 459 | 406 | 410 |
| GAAGAAGGATGAATCCCAAG | SDHB | 351 | 593 | 1042 | 1162 | 1270 |
| TCAATTTCAGGCCTGCCGGG | SDHB | 567 | 685 | 1206 | 1148 | 1230 |
| TCACATACATATGCGGCAGA | SDHB | 489 | 462 | 877 | 771 | 797 |
| TGTGCCATGAACATCAACGG | SDHB | 361 | 409 | 970 | 921 | 949 |
| AAAGTTTCCAGGGACCAACA | SDHC | 479 | 410 | 776 | 808 | 823 |
| ACAGAGCTGAGGACTAAGGT | SDHC | 368 | 647 | 1070 | 942 | 935 |
| CAGCCAAGGAAGAGATGGAG | SDHC | 416 | 683 | 1178 | 1110 | 1155 |
| CATGTACATACCTGCACTCA | SDHC | 582 | 777 | 1298 | 1317 | 1238 |
| TGATATGGGGAGACAGAGGA | SDHC | 427 | 668 | 1176 | 1277 | 1315 |
| TGGCAAATGGACATTGCCAT | SDHC | 281 | 791 | 1124 | 1092 | 1216 |
| AAAAAAAAAACATACAGTAG | SDHD | 546 | 111 | 193 | 230 | 210 |
| CAACCCTCTCACCAGTCCAG | SDHD | 599 | 516 | 1023 | 866 | 973 |
| CTCTGCGATGGACTACTCCC | SDHD | 451 | 572 | 1111 | 1063 | 1012 |
| CTGGGTTCTACACCATCCTG | SDHD | 533 | 664 | 1222 | 1158 | 1135 |
| TCTGTTTCTCCGAATCCCAG | SDHD | 476 | 426 | 640 | 682 | 704 |
| TGGAGGCTGAGCGTCCTGTG | SDHD | 479 | 661 | 1394 | 1458 | 1503 |
| AAAGGAATTCCTCAGCCGAT | SEC31A | 355 | 614 | 956 | 906 | 854 |
| AGTGGTGCTTTGATATTCAG | SEC31A | 430 | 1057 | 1668 | 1871 | 1763 |
| GACACCAGGAGCCAAAACAC | SEC31A | 634 | 613 | 820 | 686 | 754 |
| GAGGCCCCCAAATCAACTTG | SEC31A | 501 | 775 | 1400 | 1322 | 1272 |
| GCTATGGACACCAATACTGT | SEC31A | 550 | 576 | 834 | 864 | 855 |
| TGGTAATCTATCATCCTCAG | SEC31A | 430 | 678 | 941 | 878 | 883 |
| AAGCATTTGACAATACAGAA | SETBP1 | 579 | 258 | 463 | 355 | 372 |
| CACGGTTGAGACAATTCACG | SETBP1 | 585 | 841 | 1426 | 1551 | 1591 |
| CATGACGCCTCCGACATGCA | SETBP1 | 363 | 692 | 1237 | 1099 | 1066 |
| TGAAGATTCAGACCACCAAG | SETBP1 | 455 | 598 | 1075 | 1014 | 1053 |
| TGAGTGACCCAATCTCGCAA | SETBP1 | 530 | 577 | 786 | 664 | 652 |
| TGCTGGGATACGGATTCTCG | SETBP1 | 315 | 1006 | 1599 | 1697 | 1744 |
| AAGAGCTCAGAAACCTAGGG | SETD2 | 613 | 863 | 1294 | 1379 | 1455 |
| ATAATGGGATCGTTCCGACC | SETD2 | 695 | 459 | 973 | 918 | 915 |
| GAAGAATATAAACAGAGTGT | SETD2 | 422 | 492 | 804 | 821 | 858 |
| GCTGGTAACGGTACTACGGG | SETD2 | 518 | 1105 | 1532 | 1501 | 1601 |
| GTCAAGGAGACATTCATATG | SETD2 | 508 | 789 | 1141 | 1129 | 1064 |
| TGACAGTGTGACTGGATCGG | SETD2 | 592 | 954 | 1468 | 1325 | 1370 |
| AATGCCAAGATCATGATCCG | SF1 | 449 | 621 | 1216 | 932 | 934 |
| AGCCCATCTACAATAGCGAG | SF1 | 555 | 440 | 1023 | 904 | 944 |
| AGTCCAGGGGGGATGACTGT | SF1 | 572 | 530 | 1233 | 1149 | 1143 |
| CCTGGTTACTGCTAACACCA | SF1 | 389 | 286 | 565 | 459 | 478 |
| GCTTACTTGTAATCTGCAGG | SF1 | 542 | 703 | 1241 | 989 | 1092 |
| TCCACACTTGGTACACACTG | SF1 | 333 | 369 | 790 | 565 | 648 |
| AGTTCGACCATATGTGCACA | SF3B1 | 626 | 718 | 1375 | 1250 | 1243 |
| CAGTGCTCGTAAAAACAGAT | SF3B1 | 528 | 411 | 556 | 588 | 567 |
| GGACAGAAGAAATTACCGAC | SF3B1 | 650 | 183 | 325 | 245 | 248 |
| GTCAAGTTGGGATCAAGCAG | SF3B1 | 696 | 473 | 1025 | 822 | 838 |
| TCCTGGCAAGCGAGACACAC | SF3B1 | 421 | 367 | 693 | 612 | 594 |
| TGGCGGATACCCTTCCACAG | SF3B1 | 606 | 222 | 448 | 340 | 328 |
| AACTGGATGATACACCCATG | SFPQ | 596 | 514 | 1175 | 959 | 946 |
| AATAGGACCAAATTGGCTAA | SFPQ | 503 | 586 | 1084 | 939 | 997 |
| ATGGGCCTCAATCAGAACCG | SFPQ | 578 | 609 | 1266 | 1065 | 1073 |
| CCGGGCCTGAGTACCCCCGG | SFPQ | 591 | 666 | 944 | 994 | 1055 |
| GCATTTGAAAGATGCAGCGA | SFPQ | 563 | 432 | 923 | 830 | 940 |
| GGAGAAGATCTCCGACTCGG | SFPQ | 503 | 484 | 1111 | 943 | 857 |
| AAAAGCATATTATGTCGGAA | SGK1 | 578 | 640 | 1341 | 1140 | 1140 |
| AACTTACTGTTTGCATGCAT | SGK1 | 661 | 452 | 782 | 815 | 789 |
| AGAAGGAGAATACTTACCGG | SGK1 | 625 | 605 | 1231 | 1186 | 1226 |
| GCAGCATAGAAACGAGCCCG | SGK1 | 571 | 520 | 1156 | 1044 | 1149 |
| GGTTCTTCTAGCAAGACACA | SGK1 | 490 | 579 | 900 | 882 | 884 |
| GTGTGGTCTCCCAGGCACGT | SGK1 | 423 | 442 | 708 | 724 | 701 |
| ACCTCACAGAACATCAACCT | SGK3 | 626 | 347 | 741 | 700 | 668 |
| ACTCCATCAAAATAGTGTAC | SGK3 | 296 | 374 | 558 | 588 | 595 |
| AGGATCGTTCTCTCTGTAAG | SGK3 | 563 | 1078 | 1696 | 1816 | 1695 |
| AGTAGTTCTGAACAGAAAAG | SGK3 | 559 | 979 | 1540 | 1422 | 1407 |
| GTTTGTCTTCAGGAGATATG | SGK3 | 394 | 768 | 1406 | 1314 | 1371 |
| TGAAGAAATGCTCTGACGTC | SGK3 | 576 | 925 | 1472 | 1352 | 1356 |
| AGCACAGATTCCCTGAACCA | SH2B3 | 594 | 578 | 1085 | 1219 | 1263 |
| CAGAACTCGCTCCAGCCCCG | SH2B3 | 744 | 649 | 1275 | 1104 | 1117 |
| CTCCGCCACCTCATGCGCCG | SH2B3 | 654 | 653 | 1022 | 1084 | 1086 |
| GAACTGCAGCGACACGAGCT | SH2B3 | 493 | 546 | 971 | 872 | 930 |
| GACGAGGCGTCCATGGACAG | SH2B3 | 491 | 934 | 1609 | 1642 | 1617 |
| GCAGCAGCTGAATTCCTGGA | SH2B3 | 469 | 661 | 871 | 879 | 828 |
| CAAGCAGTTTATCCCCAACG | SHH | 532 | 712 | 1248 | 1252 | 1351 |
| CCGAAGCAAGTACGGCATGT | SHH | 550 | 650 | 1106 | 1140 | 1064 |
| CCGGCCGCAGAGTCGTTGAG | SHH | 499 | 758 | 1336 | 1280 | 1297 |
| CCTTCTTGGCACCGTCGTCG | SHH | 378 | 839 | 1401 | 1276 | 1256 |
| GATGTCGGGGTTGTAATTGG | SHH | 387 | 764 | 1363 | 1257 | 1282 |
| TCCCCCCGCAGAGAATTCCG | SHH | 508 | 378 | 771 | 735 | 723 |
| AATTGGATGATATGACACTG | SIRT1 | 535 | 650 | 1319 | 1243 | 1212 |
| ATAGCCTTGTCAGATAAGGA | SIRT1 | 613 | 625 | 1212 | 1183 | 1229 |
| GCGTCGGCGATTGGGTACCG | SIRT1 | 364 | 848 | 1595 | 1666 | 1678 |
| TCAATGTCAAACATTGCTTG | SIRT1 | 393 | 1010 | 1611 | 1600 | 1734 |
| TCTGAAACCGTACCTATCCG | SIRT1 | 641 | 746 | 1517 | 1491 | 1531 |
| TCTGGTTTCATGATAGCAAG | SIRT1 | 496 | 639 | 1452 | 1385 | 1382 |
| ATGAAGTAGTGACACACAGT | SIRT2 | 427 | 401 | 712 | 750 | 813 |
| GCTGCTGCGATGCTACACAC | SIRT2 | 439 | 658 | 1046 | 1054 | 1114 |
| GGCACTTACTGACTGCATGC | SIRT2 | 387 | 398 | 544 | 540 | 601 |
| GGGTACTCCCGTCGGCACAG | SIRT2 | 564 | 749 | 1406 | 1329 | 1327 |
| GTGTGAGAAATGTCACAGCG | SIRT2 | 617 | 686 | 1262 | 1125 | 1160 |
| TTGCTGATGAGGGAAGCAAA | SIRT2 | 461 | 714 | 1365 | 1170 | 1160 |
| AATATTGATGGGCTCGAGAG | SIRT3 | 447 | 608 | 1127 | 1058 | 1084 |
| CCAGCTCAAAAACAGCCTCG | SIRT3 | 535 | 552 | 985 | 871 | 894 |
| CTGAAGTCCGGAATGCCGCT | SIRT3 | 413 | 606 | 799 | 1000 | 991 |
| GGGTGGAGGAGTCTCTATAC | SIRT3 | 518 | 634 | 965 | 1072 | 993 |
| TCCCGGAAGAATGGACATGG | SIRT3 | 521 | 727 | 1315 | 1359 | 1334 |
| TCTTCGGCAGACAGTGCATG | SIRT3 | 330 | 909 | 1544 | 1340 | 1322 |
| AAGCAGAAACCCATCCAGCA | SIRT4 | 530 | 527 | 928 | 717 | 835 |
| CAGGAATCTCCACTGAGTCG | SIRT4 | 478 | 813 | 1772 | 1660 | 1573 |
| CTGGCTGGTGACCCAAAATG | SIRT4 | 665 | 725 | 1346 | 1499 | 1452 |
| GATTGGCAGGAGGATCTCTG | SIRT4 | 456 | 599 | 1184 | 1120 | 1175 |
| GTGCACCACAGTCCAAGCAA | SIRT4 | 283 | 411 | 743 | 735 | 741 |
| TGCTCAAAGCCCAATGTGCA | SIRT4 | 427 | 728 | 1234 | 1426 | 1431 |
| AGCGGAGTCCCAACTTTCAG | SIRT5 | 397 | 739 | 1482 | 1544 | 1685 |
| CTCAGCCACAATTCCACAAG | SIRT5 | 669 | 622 | 1206 | 1424 | 1380 |
| GAGTGATGACCATGACCCGT | SIRT5 | 484 | 381 | 665 | 665 | 658 |
| GTTCTACCATTATCGGCGGG | SIRT5 | 329 | 712 | 1058 | 999 | 1083 |
| GTTTCGTGCAAAAGCCTGAG | SIRT5 | 711 | 556 | 1122 | 1149 | 1134 |
| TGGCTCGACCAAGTTCAAGT | SIRT5 | 323 | 355 | 773 | 569 | 672 |
| ACAGATCTTCGACCCGCCTG | SIRT6 | 497 | 871 | 1541 | 1483 | 1504 |
| CCACGGAAACATGTTCGTAG | SIRT6 | 477 | 617 | 1032 | 960 | 977 |
| GGTCTGGCAGTCCTCCAACG | SIRT6 | 423 | 549 | 1095 | 1073 | 1146 |
| GTCACACAGACACCGCCTGG | SIRT6 | 441 | 500 | 1129 | 1000 | 1014 |
| TCCATCGTCCAGACGCCATG | SIRT6 | 508 | 838 | 1316 | 1366 | 1390 |
| TCCCTGCAAATCCGGCCCAG | SIRT6 | 356 | 589 | 1316 | 1168 | 1077 |
| AGCTCCGAGATGGCCGTCCG | SIRT7 | 759 | 681 | 1186 | 1061 | 1074 |
| CAAAGTGCACAATAGTGTCC | SIRT7 | 514 | 517 | 1063 | 1115 | 1089 |
| GCAGCGTCCACACTCCATTG | SIRT7 | 541 | 1064 | 1725 | 1761 | 1846 |
| GCAGGTGTGCGACGACCCGG | SIRT7 | 538 | 922 | 1784 | 1782 | 1828 |
| TAAGCGGGCGATGCTCATGT | SIRT7 | 438 | 614 | 1161 | 1045 | 1067 |
| TACAGCCTGCACACCCAACA | SIRT7 | 643 | 526 | 1105 | 1129 | 1151 |
| AGTTAGCTGAAGAACTCTGG | SIX4 | 527 | 666 | 1200 | 1235 | 1294 |
| ATCCTTAATGGATTAAATGT | SIX4 | 482 | 868 | 1212 | 1133 | 1059 |
| CCACGTAGCAGGTCGCTCTG | SIX4 | 499 | 906 | 1403 | 1521 | 1460 |
| CTATCCGATGAACCGGAGAG | SIX4 | 517 | 887 | 1537 | 1667 | 1596 |
| GAAGCTGCTACCGAAACAGA | SIX4 | 676 | 375 | 664 | 661 | 656 |
| GTACAAGGCGCGCTACACCG | SIX4 | 370 | 590 | 976 | 1087 | 1079 |
| ACTATCAAGACCATGTTGGA | SKP1 | 443 | 422 | 1111 | 904 | 949 |
| ATGATGAGAACAAAGAAAAG | SKP1 | 605 | 441 | 804 | 735 | 767 |
| CATCATCCTCAGGAGGAGGG | SKP1 | 374 | 601 | 1333 | 1251 | 1139 |
| CTTTGTTACAGGTCATTCAG | SKP1 | 516 | 799 | 1284 | 1200 | 1192 |
| TAACATTTGGTAGAGGAACT | SKP1 | 575 | 588 | 848 | 819 | 784 |
| TGTCTAAGTAGTTCGCAGCC | SKP1 | 429 | 470 | 612 | 560 | 584 |
| CACGAGCGAGAAGACGTAGG | SLC19A1 | 386 | 650 | 1260 | 1174 | 1168 |
| CAGCATCTGGCTGTGCTACG | SLC19A1 | 538 | 774 | 1128 | 1098 | 1033 |
| CTGGTGGGTCTTTAACTCGG | SLC19A1 | 523 | 1040 | 1578 | 1760 | 1621 |
| GCGGGGCGTCCGGGTTCATG | SLC19A1 | 617 | 852 | 1313 | 1349 | 1340 |
| GGTGATCGCGAGCGTGACGG | SLC19A1 | 484 | 697 | 1320 | 1157 | 1188 |
| TTGAAGAAGAGGCTGCGCCG | SLC19A1 | 455 | 745 | 1383 | 1285 | 1339 |
| CAGTTACAGAATTTGTCGGG | SLC22A2 | 705 | 598 | 1152 | 1195 | 1185 |
| CGTATGTGCTTCCTCACTGG | SLC22A2 | 506 | 569 | 1022 | 1106 | 1094 |
| GCGTTCACCCCCGTCTACGT | SLC22A2 | 545 | 360 | 670 | 659 | 658 |
| GTAAAAATGCTAAAGCCATG | SLC22A2 | 466 | 589 | 872 | 1028 | 950 |
| TGTCTCCCCAAACTATACAT | SLC22A2 | 421 | 284 | 472 | 451 | 480 |
| TTTCAGTCGGCTGTGAACGT | SLC22A2 | 368 | 528 | 1080 | 960 | 980 |
| AAGATGACGCTCAGCAGCAA | SLC2A1 | 569 | 587 | 1090 | 1110 | 1082 |
| AGGAGTTCTACAACCAGACA | SLC2A1 | 575 | 630 | 1071 | 1140 | 1197 |
| CCTGCTCATCAACCGCAATG | SLC2A1 | 680 | 659 | 1159 | 1153 | 1182 |
| GAGCCGATGGTGGCATACAC | SLC2A1 | 405 | 526 | 748 | 820 | 762 |
| GCCCAGGATCAGCATCTCAA | SLC2A1 | 478 | 541 | 933 | 928 | 944 |
| TCAGCATGGAGTTCCGCCTG | SLC2A1 | 469 | 739 | 1246 | 1244 | 1193 |
| ATCAAGAAGACAATCAACAC | SLC34A2 | 422 | 191 | 342 | 343 | 314 |
| CGACCAACAGAGACTAGGGG | SLC34A2 | 514 | 677 | 1228 | 1090 | 1090 |
| GAGAGAAACTCACCGATCAG | SLC34A2 | 490 | 665 | 1299 | 1129 | 1210 |
| GCTGACGATGATGCTCGACG | SLC34A2 | 506 | 915 | 1398 | 1416 | 1409 |
| TCAGCCGTCCTATTCCACTG | SLC34A2 | 447 | 319 | 532 | 539 | 531 |
| TGGACTATGAGCTTCGTGAA | SLC34A2 | 425 | 774 | 1375 | 1240 | 1195 |
| ACCCATCATAAGTTAAGGTT | SLCO1B3 | 682 | 618 | 1126 | 1085 | 1085 |
| CAGCATCTGAGGCTAACATT | SLCO1B3 | 431 | 600 | 1073 | 1124 | 1014 |
| TAACATATAGGGGATTGGTA | SLCO1B3 | 401 | 610 | 1121 | 1238 | 1201 |
| TAATATGCTCCGTGGAATTG | SLCO1B3 | 513 | 1048 | 1363 | 1396 | 1455 |
| TGTGTTTGCAAAAATGTACG | SLCO1B3 | 512 | 802 | 1502 | 1390 | 1377 |
| TTTATGTGTAAGGCACTAGG | SLCO1B3 | 481 | 513 | 934 | 1040 | 1099 |
| AGGTGGAAAGGTCGTGATGG | SLITRK6 | 445 | 898 | 1704 | 1574 | 1616 |
| CATCTGACCAAGTTAAGCAG | SLITRK6 | 569 | 475 | 864 | 782 | 781 |
| CTAGAAAGCTTGTTCTAGCA | SLITRK6 | 538 | 646 | 954 | 1018 | 956 |
| GTGGTGGTAAAACTTGTAGG | SLITRK6 | 678 | 582 | 968 | 979 | 935 |
| TCCTCGTGTTCTTCATACAC | SLITRK6 | 425 | 425 | 638 | 664 | 620 |
| TGTTCTAAGAAACCAACGTA | SLITRK6 | 611 | 821 | 1270 | 1182 | 1313 |
| AATCGGCAATATATCACGTG | SMAD2 | 391 | 506 | 1142 | 952 | 1048 |
| ACCCAAACTATGATTAACCG | SMAD2 | 414 | 389 | 595 | 584 | 603 |
| AGAAAACACTAATTTCCCAG | SMAD2 | 464 | 435 | 763 | 699 | 654 |
| ATATTATGAATTAAACCAGA | SMAD2 | 492 | 311 | 434 | 438 | 474 |
| CACCAAATACGATAGATCAG | SMAD2 | 673 | 709 | 1198 | 1170 | 1173 |
| TGCCGTGGCACTAATACCGG | SMAD2 | 423 | 600 | 1311 | 1129 | 1091 |
| AGTGCATCACCATCCCCAGG | SMAD3 | 602 | 665 | 1248 | 1115 | 1188 |
| ATGAACCACAGCATGGACGC | SMAD3 | 433 | 552 | 974 | 985 | 1081 |
| CCCAGAGCAATATTCCAGGT | SMAD3 | 500 | 475 | 858 | 891 | 856 |
| GAATGGCTGTAGTCGTCCAG | SMAD3 | 379 | 790 | 1172 | 1092 | 1148 |
| GGCCGCGTTCCTGTTGACGT | SMAD3 | 368 | 675 | 1175 | 1180 | 1220 |
| TGAATCCCTACCACTATCAG | SMAD3 | 346 | 529 | 946 | 1026 | 1093 |
| AAAGCATGTACATACTATGG | SMAD4 | 548 | 598 | 1031 | 1073 | 1050 |
| CAATACTCAGGAGCTACAGT | SMAD4 | 494 | 369 | 606 | 551 | 504 |
| GGTACTGTATGTCTCCGTTG | SMAD4 | 459 | 844 | 1750 | 1585 | 1684 |
| TAACACTGCAGAGTAATGGT | SMAD4 | 569 | 638 | 1095 | 987 | 1083 |
| TGTTACCATACAGAGAACAT | SMAD4 | 318 | 513 | 722 | 585 | 604 |
| TTACAGTAATGTCCGGGATG | SMAD4 | 478 | 899 | 1847 | 1655 | 1794 |
| AATCCAGAAACTCACGAAGG | SMARCA4 | 412 | 620 | 951 | 991 | 1056 |
| CCAAATTATAGCAGGCCTCA | SMARCA4 | 599 | 650 | 1009 | 920 | 1021 |
| CCACCATTGCAAGCTGACGC | SMARCA4 | 626 | 404 | 677 | 722 | 768 |
| CCTGCAGTCCTATTACGCTG | SMARCA4 | 615 | 875 | 1310 | 1292 | 1324 |
| CCTGGCTTGAGATGAACCCG | SMARCA4 | 528 | 648 | 1175 | 1162 | 1158 |
| GCTTTGGGACAACAGAACCG | SMARCA4 | 425 | 878 | 1615 | 1643 | 1590 |
| ACCATCAACAGGAACCGCAT | SMARCB1 | 487 | 427 | 535 | 461 | 515 |
| AGATCATGGATACACAACCC | SMARCB1 | 486 | 368 | 833 | 800 | 739 |
| CAATGACGAGAAGTACAAGG | SMARCB1 | 586 | 432 | 765 | 651 | 606 |
| GAGAACCTCGGAACATACGG | SMARCB1 | 466 | 383 | 694 | 687 | 646 |
| GCAGATAGAGTCCTACCCCA | SMARCB1 | 596 | 530 | 1113 | 884 | 971 |
| TCAAATCCAGATCATCACAG | SMARCB1 | 441 | 307 | 471 | 461 | 465 |
| ACTTTGTAGGCAGTCCCTAG | SMC1A | 633 | 493 | 1028 | 838 | 883 |
| ATTGCCACAAGCATACTGCA | SMC1A | 766 | 482 | 974 | 875 | 839 |
| CCAGCTACATGAGTACAGTG | SMC1A | 521 | 524 | 1195 | 950 | 992 |
| CCATCGCAAAAAAAACATCG | SMC1A | 714 | 232 | 310 | 275 | 256 |
| CTCTTGATATCATTAATGCG | SMC1A | 410 | 733 | 1368 | 1419 | 1497 |
| GAACATGGATGCCATTATCG | SMC1A | 498 | 620 | 1023 | 1022 | 901 |
| ACAAAAGCAGAAATGACACG | SMC3 | 659 | 498 | 1044 | 1124 | 1097 |
| ACAGATCTTTATGCAAAGCA | SMC3 | 558 | 426 | 731 | 729 | 762 |
| AGCAGATAGAGACCCAGCAA | SMC3 | 326 | 16 | 26 | 23 | 29 |
| GCCTGCTTTCTACACGTGCG | SMC3 | 574 | 769 | 1446 | 1260 | 1255 |
| GGAATATACCATTTACAACC | SMC3 | 705 | 313 | 359 | 333 | 333 |
| GGTGACCAAGTTAGTCATCG | SMC3 | 498 | 689 | 1236 | 1184 | 1154 |
| AGATGGCACCATGAGACTTG | SMO | 521 | 808 | 1450 | 1473 | 1463 |
| CCACATTTGTGGCCGACTGG | SMO | 563 | 1045 | 1253 | 1446 | 1306 |
| GGAGGTTACTTCCTCATCCG | SMO | 485 | 898 | 1586 | 1558 | 1511 |
| GGAGTGACCACGTTAGCAGG | SMO | 379 | 735 | 1297 | 1191 | 1228 |
| GGGCAGCCCGATGGTCACAT | SMO | 614 | 476 | 969 | 978 | 980 |
| TGCTCGGCTTCCGTGAACAG | SMO | 455 | 588 | 1052 | 1062 | 991 |
| AAAGATTTAGGGAAATGCTG | SNAP25 | 604 | 533 | 933 | 752 | 853 |
| CCATATCAACCAAGACATGA | SNAP25 | 609 | 347 | 577 | 474 | 418 |
| CTGCTCGTGTGGTGGACGAA | SNAP25 | 314 | 746 | 1185 | 1448 | 1444 |
| GATGCTTACAAAAAAGCCTG | SNAP25 | 483 | 546 | 858 | 838 | 915 |
| GTTATGTTGGATGAACAAGG | SNAP25 | 521 | 426 | 703 | 617 | 590 |
| TCAGTGGTGGCTTCATTCGG | SNAP25 | 457 | 538 | 982 | 951 | 1013 |
| AGAGGCAGTCGAAGCTCTCG | SOCS1 | 577 | 806 | 1707 | 1807 | 1827 |
| ATCCGCGTGCACTTCCAGGC | SOCS1 | 472 | 307 | 690 | 561 | 581 |
| CGGCTTGCGAGCGGAACGTG | SOCS1 | 554 | 708 | 1458 | 1353 | 1326 |
| CTTCGCGCTCAGCGTGAAGA | SOCS1 | 497 | 818 | 1311 | 1136 | 1168 |
| TGCGCCCCGTGCACGTTCAG | SOCS1 | 542 | 741 | 1482 | 1541 | 1571 |
| TGGTGCGCGACAGCCGCCAG | SOCS1 | 333 | 505 | 1105 | 1024 | 1020 |
| AAGTTGACCACTGTATCCGT | SOD2 | 539 | 296 | 734 | 711 | 774 |
| ACAAACCTGAGCCCTAAGGG | SOD2 | 491 | 664 | 1393 | 1165 | 1283 |
| CCTGGGCTCAAGTTCAATGG | SOD2 | 433 | 604 | 1142 | 1207 | 1218 |
| GGTGTTCAGGTTGTTCACGT | SOD2 | 456 | 411 | 1002 | 938 | 972 |
| TCCAGGGCGCCATAATCGTA | SOD2 | 472 | 549 | 1092 | 934 | 986 |
| TTCCAGCACGAGCAGGACGC | SOD2 | 399 | 653 | 935 | 895 | 945 |
| ACATGGGGGAGCCTTCGCCA | SOX10 | 489 | 632 | 1083 | 1048 | 1062 |
| ACGTTGCCGAAGTCGATGTG | SOX10 | 370 | 835 | 1442 | 1551 | 1436 |
| AGAGCAAGCCACACGTCAAG | SOX10 | 694 | 573 | 1089 | 1162 | 1133 |
| AGGACACTCTGACTCCCCCT | SOX10 | 274 | 453 | 1091 | 929 | 981 |
| TGACAAGCGCCCCTTCATTG | SOX10 | 460 | 964 | 1580 | 1480 | 1501 |
| TGGCAGGTACTGGTCCAACT | SOX10 | 433 | 369 | 780 | 786 | 734 |
| CGGATTATAAATACCGGCCC | SOX2 | 508 | 447 | 916 | 812 | 828 |
| CGTTCATGGGCCGCTTGACG | SOX2 | 511 | 604 | 1098 | 974 | 1097 |
| CTCCGACAAAAGTTTCCACT | SOX2 | 601 | 470 | 808 | 712 | 652 |
| GATAAGTACACGCTGCCCGG | SOX2 | 471 | 612 | 989 | 1027 | 1065 |
| GCAGGGCGCTCACGTCGTAG | SOX2 | 552 | 721 | 1663 | 1406 | 1543 |
| GGCCCGCAGCAAACTTCGGG | SOX2 | 526 | 456 | 1030 | 975 | 1022 |
| CACGGCCAGGTCACCTACAC | SOX9 | 440 | 425 | 968 | 924 | 864 |
| CCACGTCGCGGAAGTCGATG | SOX9 | 362 | 906 | 1782 | 1515 | 1523 |
| CGGATTACAAGTACCAGCCG | SOX9 | 720 | 379 | 845 | 799 | 793 |
| GCTCGGACACCGAGAACACG | SOX9 | 682 | 694 | 1603 | 1473 | 1420 |
| GGAGTGCACCTCGCTCATGC | SOX9 | 658 | 470 | 745 | 751 | 804 |
| TTCCCCGTGTGCATACGCGA | SOX9 | 440 | 681 | 1202 | 1058 | 1062 |
| AGAATCGTGCGGTCTCTGGT | SP1 | 411 | 681 | 1466 | 1252 | 1309 |
| CAGCAGGGACCAACTCTCAA | SP1 | 491 | 411 | 895 | 841 | 910 |
| CATCATCCGGACACCAACAG | SP1 | 385 | 470 | 821 | 873 | 899 |
| GTATGTGACCAATGTACCAG | SP1 | 602 | 464 | 1050 | 905 | 917 |
| TGAGGCAATGGGTGTAAGGG | SP1 | 429 | 638 | 1358 | 1269 | 1393 |
| TGTCCCCCTTGAACTAGCTG | SP1 | 499 | 628 | 1394 | 1277 | 1310 |
| CAGAATCTCATACCACAGAC | SP3 | 688 | 259 | 485 | 332 | 327 |
| CCATCTGTTGACTGAATCTG | SP3 | 522 | 697 | 1206 | 1221 | 1323 |
| CCGCCGCCTTACACATGGTG | SP3 | 505 | 709 | 1330 | 1235 | 1302 |
| GGGATGGAGGACGCGTCGAG | SP3 | 632 | 579 | 1048 | 966 | 978 |
| TGCACACCATGGATTGTCTG | SP3 | 405 | 819 | 1341 | 1310 | 1463 |
| TGGCTGACAAAACCTCCCAT | SP3 | 591 | 592 | 974 | 833 | 835 |
| AGCGCATCTCCCCTAGACGC | SPHK1 | 425 | 445 | 981 | 862 | 865 |
| ATCCTGGTAAACCCGCGCGG | SPHK1 | 526 | 472 | 773 | 868 | 859 |
| CTACGAACAGGTGACTAATG | SPHK1 | 554 | 632 | 1362 | 1261 | 1237 |
| GAGTCCATAACCTCGACCCG | SPHK1 | 397 | 514 | 1173 | 1106 | 997 |
| GGGAGACCACCCTTTCCACG | SPHK1 | 493 | 472 | 723 | 761 | 761 |
| TGGTGAACGGGCTCATGGAG | SPHK1 | 314 | 975 | 1677 | 1739 | 1645 |
| CATCTGCAGTCGGCGCAATG | SPTAN1 | 692 | 1137 | 1488 | 1483 | 1507 |
| GGGAGACTCTCTGGATAGCG | SPTAN1 | 528 | 561 | 977 | 880 | 915 |
| GGTAGCAGCTCGTATGAACG | SPTAN1 | 530 | 731 | 1542 | 1611 | 1591 |
| GTCTCGGCCAAAATCGTCAG | SPTAN1 | 430 | 540 | 929 | 1058 | 1132 |
| GTGGAAGTGAACGACCGTCA | SPTAN1 | 477 | 458 | 1155 | 1070 | 1189 |
| TCTTGGATCAATGGAATACG | SPTAN1 | 443 | 476 | 1152 | 1106 | 1057 |
| ACGATCTGATCTTACAGTCA | SPTLC1 | 432 | 650 | 1068 | 960 | 1008 |
| ACGATGTTGTAGTTGAGAGC | SPTLC1 | 626 | 522 | 699 | 674 | 595 |
| ATTCCCGCTTACTCTAAGAG | SPTLC1 | 584 | 516 | 1170 | 1006 | 1045 |
| CCCCCAGGACTCCAAACGAA | SPTLC1 | 280 | 437 | 643 | 649 | 639 |
| TCTTCTAAAGAATCCTCGCA | SPTLC1 | 578 | 400 | 730 | 590 | 641 |
| TGCTACTTACCTTGACCCGA | SPTLC1 | 365 | 357 | 767 | 578 | 649 |
| ACAACTATCTTGGATTTGCA | SPTLC2 | 470 | 606 | 981 | 826 | 823 |
| ATACAAGGAACCTCTACATG | SPTLC2 | 587 | 753 | 1305 | 1193 | 1127 |
| CATCATGTTACACAAAATGG | SPTLC2 | 370 | 686 | 941 | 832 | 793 |
| CCGAGGCTGACCATAAACAA | SPTLC2 | 363 | 399 | 692 | 693 | 714 |
| CTACCACACCTCGACCTGAA | SPTLC2 | 610 | 495 | 685 | 524 | 470 |
| CTCCTAGAACCAGCGATGCA | SPTLC2 | 490 | 704 | 1171 | 1163 | 1294 |
| AGTGAAAACCATAACGTGAA | SPTLC3 | 654 | 633 | 987 | 1023 | 1110 |
| AGTGGAAACTGCGCGAAGAA | SPTLC3 | 467 | 753 | 1479 | 1485 | 1531 |
| GAAGAAGATTCTCATCCTAG | SPTLC3 | 488 | 446 | 892 | 746 | 795 |
| GCTCACAGTATTGGGTCCGT | SPTLC3 | 401 | 594 | 860 | 759 | 904 |
| TACAACTGGACATTTAGGTG | SPTLC3 | 440 | 771 | 1278 | 1324 | 1259 |
| TGAATCTATGAGGACAGTAA | SPTLC3 | 447 | 563 | 989 | 824 | 786 |
| AACCGTCACAGATCACATTG | SQSTM1 | 570 | 696 | 1149 | 1189 | 1111 |
| CGACCCCAACAAACCCGATG | SQSTM1 | 517 | 592 | 985 | 1003 | 972 |
| GAGTCTGATAACTGCTCAGG | SQSTM1 | 619 | 562 | 1007 | 1003 | 998 |
| GTGGACATTGACGTGGAACA | SQSTM1 | 567 | 644 | 1152 | 1042 | 1065 |
| TTCCCTCACAGGCAGCACAC | SQSTM1 | 496 | 336 | 528 | 508 | 573 |
| TTCTTGAGGAAATTCACACT | SQSTM1 | 492 | 350 | 713 | 692 | 678 |
| AACCTGGAACGGAACCACCA | SRC | 481 | 439 | 653 | 690 | 730 |
| AAGATCCGCAAGCTCGACAG | SRC | 586 | 709 | 1075 | 906 | 901 |
| ACTTTGGCAAGATCACCAGA | SRC | 389 | 583 | 814 | 840 | 791 |
| CATCCAAGCCACAGACTCAG | SRC | 312 | 422 | 662 | 625 | 595 |
| GCTGTACGCGGTGGTGTCTG | SRC | 605 | 820 | 1560 | 1697 | 1705 |
| TCCTTAGAGAGGGTGACTGG | SRC | 420 | 562 | 1056 | 960 | 1051 |
| AACCAACTCACCTTCACCCA | SRGAP3 | 516 | 364 | 613 | 650 | 649 |
| ACTCTGTAGATCGACTGTGT | SRGAP3 | 485 | 874 | 1648 | 1423 | 1397 |
| ATGCAGCTCTCGACTACAAG | SRGAP3 | 782 | 582 | 911 | 789 | 876 |
| CCGCCTTTCACAGATCAGCG | SRGAP3 | 730 | 609 | 870 | 956 | 839 |
| CGATTAGGTCAGAGACATCG | SRGAP3 | 601 | 568 | 1027 | 938 | 1029 |
| TGGAACTCAAACTTAAGCGG | SRGAP3 | 323 | 474 | 978 | 1026 | 1022 |
| CCACGACAAGCGCGACGCCG | SRSF2 | 556 | 408 | 948 | 753 | 772 |
| CGGGTGCAGATGGCGCGCTA | SRSF2 | 598 | 548 | 1088 | 1118 | 1050 |
| CTTCTCGAACACGCGCCTCA | SRSF2 | 538 | 467 | 778 | 694 | 718 |
| GAGACTTGGAGCGGCTGTAG | SRSF2 | 458 | 504 | 907 | 927 | 944 |
| GGAGGGCATGACCTCCCTCA | SRSF2 | 614 | 729 | 1283 | 1269 | 1298 |
| TCGGCGACGTGTACATCCCG | SRSF2 | 477 | 429 | 914 | 840 | 821 |
| ATCAGATGACAATGAGTCAA | SS18 | 480 | 604 | 1142 | 1215 | 1192 |
| ATGATGGGTCAAGTTAACCA | SS18 | 605 | 483 | 1108 | 987 | 938 |
| CAACAATACAATATGCCACA | SS18 | 446 | 389 | 482 | 543 | 457 |
| CACCCCCGCCGCGATCCAGA | SS18 | 530 | 417 | 733 | 660 | 683 |
| GCAGATGTTGCATACGAACT | SS18 | 471 | 531 | 804 | 804 | 744 |
| GGCATGTTGTGAGAGCGTGG | SS18 | 476 | 1010 | 1702 | 1610 | 1604 |
| CAGAACGGGACCTTGAGCGA | SSTR1 | 433 | 541 | 1097 | 986 | 994 |
| CAGGATCACGTAGATGACCA | SSTR1 | 465 | 727 | 1140 | 1206 | 1187 |
| CATCGCCAAAATGCGCATGG | SSTR1 | 547 | 521 | 1240 | 1063 | 1085 |
| GGGCCAGTGGCGAAGCAACG | SSTR1 | 529 | 538 | 1008 | 948 | 1009 |
| GGTAGCGTGCCGCCTTGATG | SSTR1 | 477 | 1016 | 1811 | 1830 | 1853 |
| GTGCGCGAGAAGACCACGAT | SSTR1 | 503 | 351 | 672 | 623 | 645 |
| ATGTAGACGTTGGTGACCGA | SSTR3 | 717 | 769 | 1260 | 1283 | 1233 |
| CGTGGACCGCTACCTAGCAG | SSTR3 | 579 | 601 | 1254 | 1334 | 1226 |
| GAGGACGAAGAGCGCCACCA | SSTR3 | 468 | 431 | 984 | 947 | 990 |
| GGGGATCAGAACACCGCTGA | SSTR3 | 538 | 652 | 1196 | 1048 | 1140 |
| TGGCAGGTGCTCATGCCGTG | SSTR3 | 607 | 909 | 1544 | 1681 | 1746 |
| TGGTCAAGGTGCGCTCGGCG | SSTR3 | 312 | 861 | 1607 | 1637 | 1469 |
| ATGTCCGCGAAGACCACCAG | SSTR5 | 519 | 726 | 1258 | 1135 | 1166 |
| CCACGATGAGCAGGTAACAG | SSTR5 | 472 | 576 | 1014 | 1058 | 1140 |
| CGTCCAGCGTCATGACCAGG | SSTR5 | 490 | 628 | 1207 | 1253 | 1280 |
| CTGCCTGACGGTCATGAGCG | SSTR5 | 389 | 793 | 1677 | 1740 | 1717 |
| GCGGCACGCCAAGATGAAGA | SSTR5 | 389 | 626 | 920 | 973 | 919 |
| GGACGTGTAGATGATGAACA | SSTR5 | 472 | 650 | 1401 | 1275 | 1221 |
| AGATCGATTCAATCATTCTG | STAG1 | 561 | 618 | 1058 | 1055 | 1134 |
| AGCCACCAATTAATGACAGG | STAG1 | 331 | 567 | 1017 | 953 | 1040 |
| GATGTGCTGAGTACACCAAG | STAG1 | 614 | 727 | 1322 | 1352 | 1291 |
| TACAATTCGTCAAGCTGCCG | STAG1 | 695 | 734 | 1205 | 1124 | 1043 |
| TGAATCATATAAACAAGACA | STAG1 | 693 | 434 | 467 | 389 | 350 |
| TTTGTTCATAGATACCGGTA | STAG1 | 682 | 574 | 1094 | 950 | 1060 |
| AATGTCTCACTGCTCTACAA | STAG2 | 386 | 313 | 643 | 616 | 552 |
| ATCCCACTTTGAAAGATCAT | STAG2 | 422 | 512 | 843 | 902 | 852 |
| ATTTCGACATACGAGCACCC | STAG2 | 522 | 231 | 449 | 452 | 471 |
| CAGTCGTAGAGATCCAGAGG | STAG2 | 520 | 632 | 1230 | 1195 | 1236 |
| CTTACCTTAGCTAATAACTG | STAG2 | 570 | 662 | 1065 | 1248 | 1309 |
| GCCATCAAATTATGTCAGGA | STAG2 | 502 | 592 | 963 | 915 | 962 |
| AAAACCCTCAAGAGTCAAGG | STAT3 | 496 | 752 | 1274 | 1255 | 1239 |
| AAGTTGTGACTCTGCTAATG | STAT3 | 462 | 723 | 1341 | 1365 | 1395 |
| AGATTGCCCGAATTGTGGCC | STAT3 | 411 | 417 | 777 | 911 | 929 |
| CAGTCTTGATGACTAACGGC | STAT3 | 550 | 547 | 770 | 849 | 909 |
| CGAGATCACCACAACTGGCA | STAT3 | 424 | 522 | 943 | 1093 | 1040 |
| GCATCGTGAGTGAGTTAGCG | STAT3 | 604 | 871 | 1357 | 1272 | 1295 |
| CCGAAGTGGGGACCAATCGG | STAT6 | 474 | 600 | 1170 | 1130 | 1203 |
| CTTGCAACATATCAGCACCC | STAT6 | 520 | 323 | 496 | 492 | 468 |
| GCCGTGGACAATGACCACCA | STAT6 | 715 | 687 | 1279 | 1327 | 1326 |
| GCCTGGCACCACTACAAGAG | STAT6 | 579 | 788 | 1302 | 1211 | 1225 |
| GGAAATCATCAACAACACGG | STAT6 | 346 | 239 | 441 | 434 | 517 |
| TTCGATTCTTACTGGGCCTG | STAT6 | 454 | 869 | 1633 | 1550 | 1439 |
| ACAGTTTATCCTATCCAATG | STEAP1 | 627 | 728 | 1246 | 1221 | 1144 |
| ATCCATCTATCCAACCAATG | STEAP1 | 575 | 516 | 829 | 793 | 877 |
| GCAAATAAACTAGTGCCAAG | STEAP1 | 465 | 356 | 739 | 637 | 670 |
| GTTGGTGGGAAGTTACAAAA | STEAP1 | 380 | 431 | 1007 | 947 | 1006 |
| TGATGCATGTGCAAAAGCAC | STEAP1 | 388 | 452 | 729 | 855 | 837 |
| TTCTTGTTTGTGCTTAAGCT | STEAP1 | 336 | 811 | 1229 | 1024 | 1146 |
| CCAGAGCTGTTGGAATAACG | STIL | 630 | 501 | 1038 | 1073 | 1067 |
| CTGCTTGGAATTCAACGCGG | STIL | 575 | 411 | 889 | 904 | 834 |
| GGCAAGACTACCTTCGCATG | STIL | 598 | 662 | 1330 | 1208 | 1228 |
| TATATATAGTCCTCAAGTAT | STIL | 534 | 406 | 741 | 647 | 696 |
| TCAGAGTGGTTAATCAACAA | STIL | 453 | 439 | 929 | 736 | 852 |
| TGATACCAAGATTTCCAGTG | STIL | 400 | 621 | 1043 | 888 | 874 |
| AGCAGCAAGCACTTACCCGT | STK11 | 509 | 414 | 858 | 755 | 756 |
| CAGGCCTGTACCCGTTCGAG | STK11 | 533 | 657 | 1578 | 1319 | 1342 |
| CTTGATGGGAGACCTGCTAG | STK11 | 603 | 549 | 1287 | 1411 | 1452 |
| GGACGACATCATCTACACTC | STK11 | 623 | 277 | 358 | 363 | 304 |
| GGTGGACGTGTTATACAATG | STK11 | 368 | 822 | 1808 | 1701 | 1642 |
| TCCAGCAGATCAGACAACAC | STK11 | 415 | 297 | 621 | 606 | 600 |
| CAAGATCTCAGAGAACGTGG | STX1A | 417 | 585 | 989 | 940 | 1017 |
| CCTGAGGATCCGAAAGACAC | STX1A | 554 | 466 | 743 | 769 | 682 |
| CGAGTACAACGCCACACAGT | STX1A | 589 | 470 | 1063 | 891 | 945 |
| TCCCTCCCCAGAGACAAAGG | STX1A | 528 | 446 | 666 | 760 | 756 |
| TGGCCGGACCACGACCAGCG | STX1A | 572 | 573 | 1044 | 1044 | 993 |
| TGTACACGTGAGCAAAACTC | STX1A | 533 | 568 | 810 | 807 | 921 |
| AAGTGTCATACACTTACTCA | SUFU | 501 | 534 | 881 | 791 | 888 |
| ACGGATGGCTCCAACCTGAG | SUFU | 437 | 796 | 1425 | 1427 | 1469 |
| ATACCAGTACTTGACGATAG | SUFU | 367 | 735 | 1327 | 1310 | 1376 |
| CCTGGCTGATAACTGACATG | SUFU | 434 | 1067 | 1870 | 1967 | 1918 |
| GATTGTTGGCGTCTGCACCG | SUFU | 573 | 670 | 1120 | 1206 | 1156 |
| TCTGTATCGGCACACAGCCA | SUFU | 426 | 809 | 1546 | 1464 | 1489 |
| ATGCTGATACCAGGACCCAT | SULT1A1 | 344 | 675 | 877 | 894 | 893 |
| CTATTACCACTTCTACCGCA | SULT1A1 | 365 | 414 | 493 | 390 | 437 |
| CTTGACCTTCTGATCCAACA | SULT1A1 | 515 | 478 | 693 | 737 | 759 |
| GCACGTACCCCAAATCTGGT | SULT1A1 | 341 | 612 | 802 | 829 | 841 |
| GTGTGTCTTGATGAGTCGTG | SULT1A1 | 481 | 682 | 1168 | 1236 | 1254 |
| TGATGTAGAGAAGTGTCGCA | SULT1A1 | 498 | 839 | 1115 | 1078 | 1059 |
| AAGCAAGAAAAGAGCAACAT | SUZ12 | 508 | 267 | 518 | 442 | 497 |
| AGAAATCTATAGATCTGCGT | SUZ12 | 566 | 280 | 547 | 458 | 475 |
| AGTTCCTACAGGTAAAAAGC | SUZ12 | 590 | 325 | 599 | 505 | 553 |
| CTATTTAGAGTGACTCGTCC | SUZ12 | 538 | 293 | 569 | 464 | 429 |
| GGCAGTTCAGAGTACACCAC | SUZ12 | 514 | 328 | 822 | 711 | 769 |
| TTCTTACCTATAGTCTGAGT | SUZ12 | 628 | 561 | 701 | 643 | 685 |
| AATCCTTACGAGCCGGACCG | SYK | 390 | 664 | 1008 | 1062 | 1089 |
| CAACTCGTGAGATCTTCCCG | SYK | 438 | 682 | 1606 | 1443 | 1422 |
| GCATCGATAAGGACAAAACG | SYK | 410 | 459 | 959 | 817 | 797 |
| GCTTCACGTACTCCCTGATG | SYK | 463 | 888 | 1418 | 1205 | 1142 |
| GGCACCTACGCCATCGCGGG | SYK | 416 | 566 | 1122 | 1268 | 1213 |
| TTATGGTTTCTAGACATGGT | SYK | 583 | 695 | 1039 | 1137 | 1109 |
| AATCCAGTTTCAGCATCGTG | SYT1 | 540 | 767 | 1124 | 1186 | 1204 |
| ATGGCGATTAAGGCCCACGG | SYT1 | 393 | 674 | 1036 | 982 | 1033 |
| CTGCCCGCCTTAGACATGGG | SYT1 | 583 | 657 | 1223 | 1343 | 1355 |
| GGATTTCGGTCATGTAACTG | SYT1 | 419 | 951 | 1298 | 1159 | 1198 |
| GGGCAAACTTCAATATTCAC | SYT1 | 355 | 249 | 516 | 426 | 418 |
| GTGCCGTACTCAGAACTGGG | SYT1 | 452 | 788 | 1313 | 1144 | 1137 |
| ACACATCGGATATTAAGTCC | TACC3 | 518 | 452 | 763 | 681 | 709 |
| ACATCCACGATGGAGCCCAA | TACC3 | 611 | 484 | 916 | 743 | 743 |
| CAGCACACAGGACAGCACAC | TACC3 | 491 | 317 | 599 | 606 | 558 |
| GAGAAGAGAACTCCTTGCTG | TACC3 | 569 | 595 | 1393 | 1130 | 1066 |
| GGGACAACATACCCTCAGTG | TACC3 | 337 | 481 | 929 | 913 | 899 |
| GGGATCGTGTATCAGGCAAT | TACC3 | 561 | 609 | 1032 | 1040 | 921 |
| AAAGCTGGCGAAGCGGAACG | TARBP2 | 612 | 677 | 1415 | 1480 | 1458 |
| AGGAACACAAGATACCTGGT | TARBP2 | 470 | 430 | 738 | 859 | 866 |
| AGTGCAGCTGGTGTCGCCAA | TARBP2 | 399 | 646 | 1358 | 1333 | 1288 |
| CCCTCAAACACCTCAAAGGG | TARBP2 | 464 | 491 | 1035 | 996 | 957 |
| GCTTTGAGAAGGTCGTACAC | TARBP2 | 376 | 407 | 892 | 716 | 780 |
| GTTTACCATGACCTGTCGCG | TARBP2 | 583 | 414 | 856 | 769 | 770 |
| ATGCAAACGAGCTTGTCAGA | TASP1 | 656 | 786 | 1293 | 1359 | 1387 |
| CTGCATTTAAAAGAAACAAG | TASP1 | 582 | 401 | 553 | 465 | 468 |
| GAACAAAGCAGTTACCAGGG | TASP1 | 739 | 443 | 807 | 813 | 776 |
| GAAGCGAGGAGGCTTTGTGT | TASP1 | 519 | 616 | 1185 | 1170 | 1175 |
| GCAGGCACTTTGGACACAGT | TASP1 | 709 | 538 | 947 | 968 | 945 |
| TGTGATGCCAGCATAATGGA | TASP1 | 295 | 731 | 1197 | 1290 | 1211 |
| CACGGAGCAGAGGTGAACGG | TCF3 | 553 | 672 | 1277 | 1253 | 1168 |
| GAAGTGAGTGCCGTCACCGT | TCF3 | 537 | 378 | 703 | 632 | 688 |
| GTTATTGCTTGAGTGATCCG | TCF3 | 414 | 684 | 1276 | 1350 | 1397 |
| TCCAGCTCAGGTGATGACTA | TCF3 | 402 | 682 | 1042 | 905 | 995 |
| TCCCAGTATTACTCCTACCC | TCF3 | 461 | 479 | 615 | 600 | 656 |
| TTATCGCCCAGCTATGACGG | TCF3 | 517 | 714 | 1322 | 1350 | 1393 |
| AGAGATTGTCATGTGACCCA | TCF4 | 546 | 748 | 1345 | 1364 | 1369 |
| CTAGCAATAACCCCCGAAGG | TCF4 | 712 | 648 | 1219 | 1187 | 1354 |
| GACTACAATAGGGACTCGCC | TCF4 | 614 | 529 | 735 | 756 | 692 |
| GCTGTTTGGTCTAGAAACGG | TCF4 | 380 | 768 | 1156 | 950 | 967 |
| GGGATGAATCAGCCTGGCTA | TCF4 | 556 | 629 | 1208 | 1110 | 1111 |
| TAATGGTTTGTGCCACTACG | TCF4 | 207 | 783 | 1519 | 1449 | 1477 |
| CCAGGAGGAAGAGAGTACAT | TCF7L1 | 484 | 529 | 1113 | 999 | 950 |
| GGACGCGTTGGAACCCCGCG | TCF7L1 | 569 | 784 | 1191 | 1221 | 1204 |
| GTTCTTCGGAAGAAAGTTAG | TCF7L1 | 553 | 649 | 1004 | 904 | 942 |
| TCATCAGGAACGGGTACCCG | TCF7L1 | 569 | 519 | 1098 | 1297 | 1248 |
| TGACCTGGTGTCCTTGACCG | TCF7L1 | 478 | 696 | 1400 | 1447 | 1494 |
| TGTCTTTGGATCGATCTCTG | TCF7L1 | 492 | 750 | 1231 | 1034 | 1167 |
| CATCACACTCTACATACGAC | TCF7L2 | 461 | 168 | 346 | 243 | 252 |
| GGACCTACCTCCAAACACCA | TCF7L2 | 585 | 293 | 514 | 393 | 428 |
| GTGCACGTTGAAAGAAAGTG | TCF7L2 | 491 | 624 | 1100 | 1096 | 1041 |
| GTGTACCCAATCACGACAGG | TCF7L2 | 602 | 696 | 958 | 885 | 892 |
| TAATACGGAGATATATCTGG | TCF7L2 | 379 | 521 | 1225 | 1055 | 1104 |
| TCACTTACGACAATGTGTGC | TCF7L2 | 541 | 504 | 848 | 813 | 832 |
| AAACTTTGTATAAAGGGAGA | TEC | 514 | 669 | 953 | 871 | 856 |
| CTCTGCTCGACCCTCATAGT | TEC | 613 | 578 | 928 | 911 | 874 |
| GCGGTACCCAGTTAGTACGA | TEC | 623 | 596 | 1082 | 1081 | 1121 |
| GGGGAGCGGACTGTTCGGAG | TEC | 548 | 566 | 1112 | 949 | 973 |
| GTACGATTTCCAAGCAACAG | TEC | 497 | 457 | 855 | 845 | 825 |
| TGCAAAAACATAGAGTGTGT | TEC | 451 | 727 | 1117 | 1092 | 1187 |
| ACTTAACAATTTACAACCCA | TEK | 477 | 305 | 475 | 505 | 581 |
| CAGAGCTCGTATTCTGTGCG | TEK | 13 | 673 | 1409 | 1678 | 1618 |
| CGAGCCCATCACTATAGGAA | TEK | 507 | 420 | 841 | 881 | 801 |
| GAGATGTGTGATCGGTTCCA | TEK | 595 | 638 | 1126 | 1118 | 1126 |
| TCCAGTGTCAATCACGTTTG | TEK | 479 | 1106 | 1841 | 1933 | 1948 |
| TTCTCCCCATAAATCCCGGT | TEK | 515 | 532 | 661 | 618 | 591 |
| ACGGTAACCGAGACCCTGCG | TERT | 468 | 603 | 1206 | 1275 | 1279 |
| CTCTCTCCGACGGTGCTCAG | TERT | 416 | 735 | 1238 | 1470 | 1468 |
| CTGCTTCTGACAGTTCTCGG | TERT | 428 | 742 | 1303 | 1294 | 1265 |
| GCCGGAGCTTCGCTGACAGG | TERT | 482 | 469 | 1042 | 886 | 893 |
| GGCCCGTATGCGTAGCACAA | TERT | 581 | 628 | 1131 | 1025 | 1016 |
| GGGGCCTCATTCGCCAGTAG | TERT | 384 | 644 | 1202 | 1076 | 1121 |
| ATAAATATCTCCACTCACAG | TET1 | 699 | 324 | 549 | 484 | 494 |
| CCAGGAATTTCCCATCGATG | TET1 | 562 | 982 | 1935 | 1736 | 1844 |
| CGGTCATTACATTAGCACAC | TET1 | 464 | 231 | 303 | 350 | 362 |
| GAAGAAACGAAAGCGATGTG | TET1 | 434 | 643 | 1200 | 1155 | 1251 |
| GCGAGCTGTGATCATCTCAA | TET1 | 488 | 706 | 1213 | 1099 | 1163 |
| GCTGTACATTATATGCAACG | TET1 | 606 | 327 | 579 | 557 | 580 |
| AAAGTGGAAGAATGTTTCCG | TET2 | 466 | 682 | 1195 | 1228 | 1390 |
| ACATTATCACAACTCGGGGG | TET2 | 474 | 465 | 1102 | 1043 | 1180 |
| AGGGCAGTCTCAAGGTACAG | TET2 | 468 | 654 | 1265 | 1170 | 1184 |
| CAAAATAGAGGGTCAATGTG | TET2 | 609 | 825 | 1403 | 1383 | 1327 |
| TCTGCCCTGAGGTATGCGAT | TET2 | 540 | 508 | 1073 | 867 | 957 |
| TGGAGAAAGAAATAACTTCG | TET2 | 444 | 600 | 1176 | 1089 | 953 |
| AATCAAGATTGAGTCTTCGG | TET3 | 467 | 381 | 884 | 811 | 864 |
| ACATCCAGTCAGTATTCAAG | TET3 | 604 | 496 | 948 | 836 | 844 |
| ACCCCACATACTCATCATTG | TET3 | 453 | 678 | 1037 | 1021 | 996 |
| CAACCGTGAGATGAGTCGCG | TET3 | 538 | 808 | 1406 | 1352 | 1367 |
| CAAGGAAGACAATCGCTGCG | TET3 | 400 | 879 | 1450 | 1359 | 1458 |
| CTGGAACAGGTCCATCAACT | TET3 | 577 | 506 | 867 | 996 | 1056 |
| CATTCCTGAGCCGAATTGGT | TFDP1 | 660 | 478 | 926 | 797 | 760 |
| CCAGCGTGTTCACTGCAGAG | TFDP1 | 579 | 455 | 1096 | 865 | 942 |
| GGGGACCACGTCCTACAACG | TFDP1 | 615 | 361 | 618 | 570 | 595 |
| GGTGCAGAGGAACCGCCAGG | TFDP1 | 538 | 629 | 1199 | 1120 | 1055 |
| TGCAGCGCCCAACACTATCG | TFDP1 | 529 | 648 | 1406 | 1246 | 1211 |
| TTCTTCTGGTCGTAAGCCTG | TFDP1 | 511 | 873 | 1479 | 1534 | 1407 |
| ATGGTTATAAGATCTCCATC | TFG | 609 | 463 | 903 | 796 | 913 |
| CTCCTGCAGAAGACCGTTCA | TFG | 471 | 801 | 1295 | 1248 | 1302 |
| GGGCTGCTGAGCACCATAGC | TFG | 404 | 344 | 676 | 479 | 516 |
| GTCTGAGCTCCTGTATATGG | TFG | 387 | 754 | 1562 | 1399 | 1381 |
| TATTCCTCTCATAGATACTG | TFG | 504 | 448 | 704 | 592 | 569 |
| TCAGTTCTATCAGCTCTCGA | TFG | 490 | 376 | 536 | 586 | 598 |
| AAAATAGAACTCCCAACTGT | TGFBR1 | 483 | 525 | 882 | 821 | 832 |
| ACTCCACAGACAATGGTACA | TGFBR1 | 378 | 870 | 1427 | 1347 | 1397 |
| AGAGCGCTCATGGTTCCGTG | TGFBR1 | 504 | 791 | 1519 | 1388 | 1464 |
| ATGGTCTGTCTCGAGGAATC | TGFBR1 | 491 | 908 | 1333 | 1439 | 1415 |
| GGGTACCAACAATCTCCATG | TGFBR1 | 380 | 654 | 999 | 959 | 968 |
| TGGCACTCGATGGTGAATGA | TGFBR1 | 309 | 813 | 1266 | 1452 | 1504 |
| ACCTGCAGGAGTACCTCACT | TGFBR2 | 537 | 507 | 799 | 840 | 798 |
| ACGGCTATGGCGATTCCCAG | TGFBR2 | 417 | 855 | 1406 | 1203 | 1249 |
| AGGTAGAGCTGATGTCGGAC | TGFBR2 | 568 | 597 | 944 | 1044 | 936 |
| CATCTTGGGCCTCCCACACA | TGFBR2 | 452 | 867 | 1590 | 1299 | 1408 |
| GACCCCAAGGATACCTACCA | TGFBR2 | 543 | 779 | 958 | 969 | 1089 |
| GGGGAAAGGTCGCTTTGCTG | TGFBR2 | 489 | 890 | 1525 | 1547 | 1507 |
| AATTCTCGTACACCCCCTGG | TIAM1 | 667 | 635 | 1177 | 962 | 1095 |
| CCAGATTGAGAAGTCAGATA | TIAM1 | 441 | 584 | 971 | 996 | 978 |
| GCAAAGCATGCCAGACTCGG | TIAM1 | 493 | 503 | 993 | 858 | 865 |
| GGAGGCTGGCTAAGTAACAG | TIAM1 | 475 | 718 | 1261 | 1095 | 1163 |
| GGTTCATCTCATGCAGACTG | TIAM1 | 429 | 813 | 1345 | 1252 | 1237 |
| GTATGCTCAGAAAAACTCCG | TIAM1 | 483 | 361 | 717 | 643 | 660 |
| AGTGTGCGTCAGCACCCTCG | TMPRSS2 | 553 | 769 | 1350 | 1334 | 1359 |
| CAAGGAATAGCGGATGACAG | TMPRSS2 | 491 | 657 | 1356 | 1216 | 1280 |
| CCAGACCTCTACACAACCCG | TMPRSS2 | 495 | 510 | 1044 | 958 | 944 |
| GGACGGGGTGCACCGCAAGT | TMPRSS2 | 566 | 680 | 1187 | 1166 | 1314 |
| GGATAAAGTTCGGTCCGTAG | TMPRSS2 | 598 | 532 | 1323 | 1273 | 1283 |
| TCGGTGTCCGGGATAGAGTG | TMPRSS2 | 555 | 803 | 1482 | 1510 | 1484 |
| ATGAGGTACAACCCATCTGA | TNF | 388 | 621 | 1020 | 799 | 877 |
| CCAGGAGGGCATTGGCACGT | TNF | 391 | 558 | 977 | 1108 | 1141 |
| GAGATACTTACTGACTGTCT | TNF | 345 | 593 | 887 | 940 | 899 |
| GTGGGTGAGGAGCACATGGG | TNF | 610 | 717 | 1229 | 1165 | 1309 |
| GTGTGGTGGCCCCTGCGACG | TNF | 454 | 607 | 1238 | 1031 | 1026 |
| TGGAGTGATCGGCCCCCAGA | TNF | 525 | 786 | 1277 | 1237 | 1358 |
| AACACCCCACATGTACTGAG | TNFAIP3 | 459 | 360 | 626 | 598 | 640 |
| ACACTGAACGTGCAGCACAA | TNFAIP3 | 486 | 473 | 747 | 757 | 775 |
| ACGTCCATGAGAGAAAGCTG | TNFAIP3 | 557 | 567 | 1231 | 1099 | 1134 |
| AGTTATACTGAAGTCCACCT | TNFAIP3 | 422 | 586 | 769 | 751 | 885 |
| CCTCGCTCTCTGTTAACGAG | TNFAIP3 | 368 | 713 | 1499 | 1340 | 1399 |
| TGCAGCATTGATCAAATGGG | TNFAIP3 | 526 | 881 | 1437 | 1410 | 1362 |
| GACAGCAGCAACTAATGCCA | TNFRSF11A | 512 | 439 | 955 | 924 | 870 |
| GAGAGAAGGAAGTACGTCAT | TNFRSF11A | 347 | 715 | 983 | 1040 | 1025 |
| GCTGCTAAAATTGTTGCCTG | TNFRSF11A | 667 | 634 | 1083 | 1127 | 1060 |
| GGCATTACGAGCATTTCGGA | TNFRSF11A | 367 | 430 | 824 | 803 | 669 |
| GGGATCCAATCAGTCCTGCA | TNFRSF11A | 599 | 656 | 1022 | 972 | 1003 |
| TCCCACGGAGGACGAATACG | TNFRSF11A | 465 | 936 | 1374 | 1594 | 1568 |
| AAGAGGAAGAGTATCCGGTG | TNFRSF14 | 673 | 773 | 1621 | 1563 | 1622 |
| CAGTGCCGAATGTGTGACCC | TNFRSF14 | 619 | 515 | 1032 | 1033 | 1017 |
| CGCCCTGCAAACCTGGACAG | TNFRSF14 | 573 | 790 | 1138 | 1265 | 1130 |
| CTTCTGCATCAGTGAGAACG | TNFRSF14 | 407 | 809 | 1413 | 1374 | 1342 |
| GGGTGCAGGTTACCGGGTGA | TNFRSF14 | 512 | 1065 | 1823 | 1741 | 1754 |
| TCTGTGCTCCCTGTGACCCG | TNFRSF14 | 561 | 922 | 1331 | 1286 | 1393 |
| ACCCGGACACTTCTCCCCAG | TNFRSF4 | 533 | 493 | 1134 | 1180 | 1134 |
| CCACGCTTGTAGCCGTCCCG | TNFRSF4 | 495 | 518 | 990 | 1008 | 1086 |
| GAAACAGTTGTGCTCAGCTA | TNFRSF4 | 390 | 569 | 880 | 909 | 952 |
| GAGCTCAGACGCTGTCTGTG | TNFRSF4 | 609 | 674 | 1334 | 1264 | 1331 |
| GAGGAGATGCACACCCACGC | TNFRSF4 | 463 | 395 | 740 | 751 | 763 |
| TCCATCTGGCTTCTACAATG | TNFRSF4 | 513 | 529 | 1076 | 882 | 915 |
| AAGATGATCCCAAAGTGGCG | TNFRSF8 | 470 | 480 | 904 | 861 | 937 |
| CATCAACCCAGACGACAGCA | TNFRSF8 | 337 | 457 | 749 | 721 | 764 |
| CCCACCCCGTCTGCCCACCA | TNFRSF8 | 350 | 537 | 987 | 988 | 1015 |
| CCGACATTCACAGACGCGAG | TNFRSF8 | 642 | 380 | 788 | 927 | 787 |
| GTGCGAGCCTGACTACTACC | TNFRSF8 | 475 | 503 | 877 | 770 | 861 |
| TCGTTACCTGAAACCAGAAC | TNFRSF8 | 562 | 210 | 319 | 276 | 309 |
| AAAATTCATGAGCAACAACA | TNFRSF9 | 518 | 252 | 475 | 410 | 415 |
| AATTATACATACTTTGTCCA | TNFRSF9 | 527 | 418 | 746 | 802 | 794 |
| ACCAGCTGGGCATTTACTAC | TNFRSF9 | 502 | 332 | 722 | 737 | 748 |
| GATCATTAAATGTCCCAAAG | TNFRSF9 | 745 | 415 | 698 | 671 | 643 |
| TCACACTCTGCATTGCTGAT | TNFRSF9 | 493 | 535 | 1196 | 1223 | 1191 |
| TGTGAACAGGATTGTAAACA | TNFRSF9 | 590 | 613 | 1222 | 1223 | 1212 |
| AACAGCACGCTGAGCCTGAG | TNFSF13B | 516 | 805 | 1350 | 1685 | 1620 |
| CGCCCTGTGGAAAGACACCA | TNFSF13B | 368 | 486 | 784 | 928 | 917 |
| GAAACAAGCGTGCCATTCAG | TNFSF13B | 550 | 666 | 1270 | 1127 | 1225 |
| GATGGTGCCAAGATTCCCTA | TNFSF13B | 424 | 626 | 1129 | 1050 | 1035 |
| TCTTTGGAGCATGGGACAGA | TNFSF13B | 371 | 553 | 1155 | 1092 | 1119 |
| TGGCTGCTCAGCTTTAAAAG | TNFSF13B | 460 | 608 | 1159 | 1058 | 1171 |
| AAGGGCTCACAATGACGTCA | TNKS | 462 | 567 | 946 | 863 | 900 |
| CCATGGCAATTGACTAACGT | TNKS | 652 | 310 | 582 | 654 | 540 |
| GTTGGAGGCCTGTCGCAATG | TNKS | 447 | 558 | 979 | 923 | 1001 |
| TCAACATCAGAAGTGCGTAT | TNKS | 583 | 389 | 601 | 610 | 555 |
| TGAATTGCCATGCAAGTGAT | TNKS | 412 | 628 | 1157 | 1098 | 1025 |
| TGGGTTGGGAGCGACCCCCA | TNKS | 349 | 887 | 1401 | 1454 | 1417 |
| AGACCCTACAAAGAAAAACA | TNKS2 | 377 | 239 | 383 | 300 | 330 |
| ATTCTTGACTCCTCTACACG | TNKS2 | 443 | 693 | 1133 | 1030 | 1023 |
| GCGTGGGTCCCGGTTACCTG | TNKS2 | 549 | 702 | 1061 | 937 | 1013 |
| GTGTAAATGCAATGGACTTG | TNKS2 | 505 | 607 | 1244 | 1078 | 1107 |
| TCAACTGCAGAGACATTGAA | TNKS2 | 611 | 507 | 877 | 718 | 788 |
| TGTCAATGCCACGGACAAGT | TNKS2 | 290 | 453 | 888 | 781 | 779 |
| AAGGAACCTTGGCATCTCTG | TOP1 | 338 | 749 | 1328 | 1001 | 1037 |
| ACTCACTCATCCTCTTCTCG | TOP1 | 577 | 1025 | 1554 | 1403 | 1434 |
| AGTTGGCAATCCTCTCTCTG | TOP1 | 685 | 438 | 1144 | 882 | 855 |
| GAAGAAGAGCGCTATCCTGA | TOP1 | 655 | 464 | 916 | 724 | 736 |
| TTACCTTGATTCGTGAACTG | TOP1 | 345 | 648 | 1086 | 1055 | 952 |
| TTTGAAATACTGGCTCATCT | TOP1 | 356 | 487 | 945 | 838 | 894 |
| CAGACGTGGATGGATAACAT | TOP2A | 531 | 437 | 905 | 769 | 754 |
| GAGATCTATCCCATCTATGG | TOP2A | 572 | 467 | 1112 | 844 | 885 |
| TCCATGAACAAGTAAACCCC | TOP2A | 589 | 274 | 738 | 556 | 580 |
| TCCCTATAATCTGTTATGAG | TOP2A | 406 | 332 | 706 | 578 | 619 |
| TGTACACTTATCCTGACTGA | TOP2A | 335 | 497 | 836 | 764 | 819 |
| TTTGGTACTAGGCTGCACGG | TOP2A | 327 | 344 | 854 | 648 | 646 |
| AAAGGAACGATCCAAGAGCT | TOP2B | 443 | 524 | 953 | 918 | 1051 |
| ACATTGTGAAAATATAACGA | TOP2B | 538 | 377 | 603 | 564 | 590 |
| ATGCAGCTGACAATAAACAG | TOP2B | 563 | 550 | 824 | 879 | 809 |
| GTGTACACTGATACTAACAG | TOP2B | 365 | 313 | 719 | 686 | 718 |
| TAGGTTACATGGCTTACCAG | TOP2B | 589 | 561 | 997 | 827 | 904 |
| TTTGGCTGGTTCATGCAAGG | TOP2B | 570 | 628 | 1144 | 1094 | 1179 |
| AGTTCGTGACCGAGGTTGTG | TP53 | 416 | 1379 | 2133 | 2519 | 2460 |
| CGCGGACGCAGGTATTGGGT | TP53 | 505 | 986 | 1369 | 1694 | 1556 |
| GCTGGCAAAACAACTTGTTG | TP53 | 642 | 1796 | 2196 | 2591 | 2656 |
| GGTGCCAGGGTAGGTCTTCG | TP53 | 357 | 2018 | 2382 | 2755 | 2679 |
| GTATAGTCAGAGCCAACCTG | TP53 | 552 | 3289 | 2870 | 3725 | 3894 |
| TCCTCAGCATCTCATCCGAG | TP53 | 478 | 1887 | 2135 | 2397 | 2344 |
| AAACAAGATTGAGATTAGCA | TP63 | 474 | 657 | 1020 | 1039 | 1069 |
| AAGGACACGTCGAAGCTGTG | TP63 | 488 | 641 | 1236 | 1321 | 1276 |
| AGCACACGATCGAGACATAC | TP63 | 657 | 308 | 536 | 429 | 421 |
| CGATGATTAAAATTGGACGG | TP63 | 570 | 826 | 1378 | 1311 | 1230 |
| GCTGAGCCGTGAATTCAATG | TP63 | 455 | 682 | 1245 | 1201 | 1104 |
| GTATAACAGCTCATCATCTG | TP63 | 411 | 658 | 941 | 843 | 966 |
| AAGGTGACGTCGAAGTGGTG | TP73 | 409 | 615 | 1164 | 1193 | 1135 |
| CATGCCCGTCTACAAGAAGG | TP73 | 620 | 592 | 1403 | 1362 | 1451 |
| CTCACGGTAGTGGTCTTCGT | TP73 | 527 | 382 | 516 | 554 | 634 |
| GCTGCTCAGCAGATTGAACT | TP73 | 683 | 551 | 1026 | 868 | 814 |
| GCTGTGTGCTTTCACATGCA | TP73 | 441 | 939 | 1608 | 1584 | 1768 |
| GTATGTGGACGACCCTGTCA | TP73 | 667 | 736 | 1508 | 1361 | 1411 |
| AAACCGGGCCATGAAAGATG | TPM4 | 543 | 518 | 1146 | 1135 | 1115 |
| CAAATACTCCGAGAACCTGA | TPM4 | 402 | 448 | 455 | 378 | 390 |
| CACTCACAGTTCAGACACCT | TPM4 | 7 | 319 | 697 | 582 | 598 |
| CCACTGCGCACGTACGTCGG | TPM4 | 402 | 863 | 1407 | 1391 | 1315 |
| GGAGGCCGACCGCAAGTATG | TPM4 | 515 | 960 | 1445 | 1653 | 1504 |
| GGAGTCGGATAAGAAAGCTG | TPM4 | 606 | 756 | 1589 | 1502 | 1522 |
| ATTGGATTCCATCAAGAGCA | TPMT | 715 | 492 | 858 | 778 | 874 |
| CCTACCTGGGAAGATCAAAA | TPMT | 548 | 371 | 566 | 627 | 625 |
| GAAAAACCGAGTACTGACTC | TPMT | 538 | 461 | 898 | 799 | 796 |
| GCATTAGTTGCTATTAACCC | TPMT | 452 | 542 | 1069 | 862 | 943 |
| GTGTGGAAATCAGTGAACTT | TPMT | 505 | 455 | 682 | 706 | 670 |
| TTACACAGAAGAACCCATTG | TPMT | 518 | 680 | 1346 | 1301 | 1418 |
| AGCTCTACTACATCAGCTTG | TRIM24 | 638 | 1339 | 2271 | 2184 | 2158 |
| ATGTGACAAACTGACATGCC | TRIM24 | 687 | 452 | 783 | 755 | 729 |
| CACTAGGTGAACGTATTGGT | TRIM24 | 589 | 584 | 991 | 986 | 998 |
| CATCACACCTTGCTCGAAGG | TRIM24 | 585 | 554 | 808 | 712 | 741 |
| GATTGACTTGAGCTCACCAG | TRIM24 | 344 | 523 | 1006 | 925 | 954 |
| TTAGGTATGCACAAGCTGTG | TRIM24 | 588 | 653 | 1465 | 1266 | 1270 |
| ACAACATTAGGAGTATAACC | TRIM33 | 436 | 339 | 527 | 538 | 478 |
| ACGATGGCAAATGCAAATCG | TRIM33 | 598 | 973 | 1702 | 1689 | 1715 |
| ATGCAGACAGATAGACCTTG | TRIM33 | 350 | 805 | 1697 | 1438 | 1537 |
| CTGTTGCTGTAACATCTGCG | TRIM33 | 501 | 944 | 1537 | 1315 | 1415 |
| GCCGAGCAGCGCTTAGTCGG | TRIM33 | 559 | 938 | 1565 | 1490 | 1455 |
| TATTGGAACACAAAGAACAT | TRIM33 | 542 | 258 | 539 | 433 | 428 |
| ACATCGAGCAGAAGGTCCAG | TRIM62 | 342 | 706 | 1130 | 1276 | 1325 |
| CACTCACCGCTCGGACAGCG | TRIM62 | 551 | 721 | 1337 | 1303 | 1346 |
| CCATCGGTGAGGCGTTCGAG | TRIM62 | 577 | 1013 | 1689 | 1550 | 1644 |
| GTCCGACGACTGCACCATCG | TRIM62 | 655 | 588 | 1204 | 1168 | 1237 |
| GTGGGGGGTTGAACCGTGCG | TRIM62 | 605 | 701 | 1125 | 1124 | 1167 |
| TTCCCGACCTCCAAGTACAC | TRIM62 | 365 | 234 | 440 | 398 | 430 |
| CATCACCCTTGCTTATTCCG | TRIO | 428 | 675 | 811 | 769 | 789 |
| CTCCAGTATACAGCACACCC | TRIO | 547 | 387 | 524 | 512 | 517 |
| CTGGATAAAGCACATCCGCG | TRIO | 603 | 716 | 1412 | 1264 | 1314 |
| GTCCGCAAGAGCACCGACGC | TRIO | 633 | 277 | 504 | 550 | 546 |
| TGGACAGGCTGCACTCAACA | TRIO | 615 | 793 | 1091 | 1198 | 1127 |
| TTGCGATAGAGATGTCGACA | TRIO | 472 | 646 | 1266 | 1087 | 1134 |
| CAGATGGGGAAGATCATGCG | TRPC6 | 568 | 709 | 1415 | 1344 | 1357 |
| CCGAGCACCCTTCCGCAAGA | TRPC6 | 724 | 465 | 993 | 957 | 925 |
| GGGAGAAGGCTAGCCAACCG | TRPC6 | 315 | 627 | 1189 | 1183 | 1168 |
| GTTCTAAAGCTGTCATGACT | TRPC6 | 569 | 402 | 609 | 600 | 595 |
| TAAAGGTTATGTTCGGATCG | TRPC6 | 410 | 882 | 1532 | 1469 | 1545 |
| TTTCCTAGCCAGGATCAAGT | TRPC6 | 396 | 437 | 596 | 561 | 598 |
| CCAGTGACAACAAGCGGTGG | TRPV4 | 622 | 416 | 1092 | 1152 | 1095 |
| CGGTAGTAGATGTCCCGGAA | TRPV4 | 558 | 564 | 787 | 708 | 726 |
| GTCAGGTAGTTGACGATGTG | TRPV4 | 431 | 938 | 1685 | 1674 | 1658 |
| TAAGAAGCGCCTGACTGACG | TRPV4 | 485 | 596 | 1229 | 1082 | 1044 |
| TACTCACTAGAGTAGTTGGA | TRPV4 | 559 | 698 | 1338 | 1307 | 1361 |
| TCAGCACATTATCCGCCGGG | TRPV4 | 568 | 667 | 1025 | 1221 | 1187 |
| AGAGCTCTCAGAGATCACGA | TSC1 | 650 | 391 | 796 | 715 | 723 |
| AGCATGCTCTCCGAAACAGG | TSC1 | 580 | 663 | 1185 | 1263 | 1253 |
| CCAGCAGTTACCTTGTCGTG | TSC1 | 291 | 797 | 1570 | 1603 | 1614 |
| CCTCAAAAAGATGATCGTAA | TSC1 | 402 | 365 | 449 | 410 | 435 |
| GGAGTCGCCATCTTCGTATG | TSC1 | 483 | 915 | 1577 | 1621 | 1597 |
| GTCATCCGAGTGGCACAATG | TSC1 | 607 | 921 | 1221 | 1259 | 1256 |
| CAAGTCACCTCATAAAACGA | TSC2 | 492 | 341 | 651 | 612 | 581 |
| CATACTTTCAGGTCCCCGGT | TSC2 | 605 | 369 | 661 | 711 | 693 |
| CATGGAGGTCTTCGTTAGCG | TSC2 | 372 | 702 | 1621 | 1596 | 1542 |
| CCCATCATGTCATAGCCATG | TSC2 | 477 | 653 | 1131 | 1144 | 1121 |
| GATGAGCACCTTATAACGCA | TSC2 | 412 | 398 | 830 | 631 | 775 |
| GCTGGCCAGCAAGTCTGACA | TSC2 | 499 | 646 | 1263 | 1244 | 1216 |
| AACCAGCTGTGTTACACCCA | TSHR | 283 | 629 | 1277 | 1278 | 1214 |
| AGTTGCACATGAGAAAGCGT | TSHR | 457 | 447 | 796 | 765 | 722 |
| CAATAGATGCAACTCTGCAG | TSHR | 554 | 628 | 1089 | 1030 | 1071 |
| CTGGTGTAAGAGACATCCCT | TSHR | 520 | 433 | 768 | 779 | 815 |
| CTTCATATTCCTGGTCAAAG | TSHR | 503 | 451 | 812 | 628 | 725 |
| GCAGGAGCTTAAAAACCCAC | TSHR | 437 | 290 | 432 | 415 | 403 |
| CAGCTCTTTCCAGACCATGT | TSPO | 503 | 560 | 941 | 857 | 909 |
| CGTCAGCAGGAGGTCCACCA | TSPO | 510 | 611 | 1263 | 1275 | 1387 |
| CTACCCCATGGCCGAGTACA | TSPO | 444 | 658 | 1406 | 1252 | 1179 |
| GGGCTCGTACTATGTCCGCG | TSPO | 496 | 347 | 969 | 927 | 900 |
| TGTCGGGTGCCAAAGAAGAG | TSPO | 531 | 812 | 1534 | 1557 | 1472 |
| TTCCCCTGGGACTCTATGCT | TSPO | 358 | 608 | 1210 | 984 | 1009 |
| AGTTAACCTTCTAAATAGCC | TTK | 631 | 327 | 571 | 540 | 551 |
| CAAACCTGTAAAGTTTGAAG | TTK | 357 | 541 | 798 | 739 | 719 |
| CATGTAGGTGCTGATATTGG | TTK | 483 | 625 | 1391 | 1176 | 1131 |
| GAACCATTCCCCAATTCACT | TTK | 347 | 381 | 690 | 567 | 599 |
| GATGATGGCAAACAACCCAG | TTK | 500 | 336 | 612 | 546 | 555 |
| GCACAATTTGAACTATCGCA | TTK | 591 | 372 | 524 | 522 | 556 |
| GAGCACGGGATCGACTCAGT | TUBB1 | 372 | 427 | 780 | 791 | 877 |
| GAGCTGAGCCACAACGTAGG | TUBB1 | 285 | 593 | 1424 | 1307 | 1374 |
| GCACTGTCTAGTGACCTCAC | TUBB1 | 447 | 550 | 958 | 980 | 980 |
| GCTGCTGAGCAGGATTCGTG | TUBB1 | 737 | 676 | 1210 | 1216 | 1307 |
| GGGTGCTCACCAGTGCAGGG | TUBB1 | 661 | 795 | 1335 | 1160 | 1289 |
| TCCCTGCGCAGGTAACTCTG | TUBB1 | 640 | 553 | 1193 | 1201 | 1316 |
| CACGGAGGTAGGGCCATCAG | TYMS | 712 | 440 | 1176 | 889 | 1011 |
| CAGGCGCGCTACAGCCTGAG | TYMS | 480 | 557 | 1188 | 1050 | 1099 |
| CCTTAGCGTTTGTGGATCCC | TYMS | 586 | 663 | 1137 | 1240 | 1128 |
| GAATAATTCTTCTGTCGTCA | TYMS | 493 | 553 | 785 | 749 | 762 |
| GGAGCTGCAGTACCTGAGGC | TYMS | 742 | 550 | 847 | 950 | 955 |
| TGGGGCAGAATACAAAGATA | TYMS | 517 | 614 | 1495 | 1232 | 1344 |
| CCCGGTGACTGACTTCCGAG | U2AF1 | 477 | 576 | 1454 | 1140 | 1153 |
| CGTACACGTTCCCAACTAGG | U2AF1 | 471 | 391 | 642 | 453 | 506 |
| CTGCATCTCGACGTCACTCA | U2AF1 | 424 | 545 | 867 | 1025 | 916 |
| GCGGTTGTTCAAGTCAATCA | U2AF1 | 442 | 523 | 856 | 871 | 851 |
| GTAAATGTTCAAGAGGGCAA | U2AF1 | 570 | 692 | 1230 | 1076 | 1020 |
| GTCATGGAGACAGATGCTCT | U2AF1 | 513 | 480 | 1010 | 836 | 913 |
| CTAAGGAAGTCCTCGACCAG | UBA1 | 478 | 625 | 1137 | 992 | 1004 |
| GATCCTCACAGATTCCAATG | UBA1 | 533 | 401 | 878 | 682 | 759 |
| GGCGTTTGGTTACTTACGTG | UBA1 | 649 | 506 | 1216 | 985 | 1036 |
| TCATAACGGTTCTGGCGCTG | UBA1 | 599 | 376 | 1035 | 786 | 837 |
| TGCGGTAGTACACACAACGG | UBA1 | 416 | 590 | 1077 | 848 | 808 |
| TTCTCCGATTATATCCGTGG | UBA1 | 455 | 543 | 1248 | 921 | 963 |
| ATCCAAGGACCATACCTGCT | UCHL5 | 520 | 568 | 1029 | 1112 | 1088 |
| ATGACTGGATCAGCGCAGTA | UCHL5 | 437 | 682 | 1262 | 1318 | 1288 |
| CAGCCAGGAGAAGAACCAGC | UCHL5 | 501 | 419 | 641 | 538 | 502 |
| CTGATAATGTCTCTCCCAAA | UCHL5 | 489 | 622 | 919 | 816 | 814 |
| GAATTAGATGGATTAAGAGA | UCHL5 | 467 | 354 | 676 | 613 | 558 |
| TTTCTTATTTCAGGTTGCCG | UCHL5 | 528 | 698 | 1106 | 1149 | 1098 |
| AACCAAATGACAGAGAAAGT | UGCG | 470 | 379 | 739 | 638 | 676 |
| AGAGATATAGGACCTTGGAT | UGCG | 538 | 457 | 839 | 807 | 846 |
| CATCATGATCTTGTACACAA | UGCG | 674 | 331 | 422 | 587 | 523 |
| GATGTGTTAGATCAAGCAGG | UGCG | 411 | 623 | 1033 | 1159 | 1110 |
| GCTTCAGAAGAGACACACCC | UGCG | 664 | 276 | 494 | 433 | 452 |
| TGGCCAAAGCAATAGCTGAC | UGCG | 550 | 260 | 548 | 529 | 620 |
| AAAGCTTCGATGCAACTGGG | UMPS | 604 | 641 | 1015 | 964 | 981 |
| AAGACAAGTTGCAGGCACAT | UMPS | 519 | 490 | 864 | 705 | 825 |
| ATTGAAGATGTTGTCACCAG | UMPS | 664 | 474 | 1041 | 972 | 1030 |
| CCGGTTTACATCGATCTGCG | UMPS | 477 | 707 | 1300 | 1213 | 1216 |
| GAACAGATAACTGTAGCCAA | UMPS | 457 | 585 | 1099 | 1066 | 1007 |
| GTCTTGAGTATGCAGATGCT | UMPS | 517 | 713 | 1013 | 1014 | 1044 |
| AAGTAACTGTAATATCCTGG | USP10 | 479 | 521 | 929 | 903 | 967 |
| AGAAGTCTGTAAATATACGT | USP10 | 489 | 224 | 540 | 542 | 515 |
| CCTTTATTGATCAGCCCACG | USP10 | 496 | 654 | 1275 | 1131 | 1147 |
| CGGAGTCGAGTTGCACACTG | USP10 | 607 | 559 | 1131 | 1063 | 1112 |
| GCAACTGAGAATAAATTCAG | USP10 | 368 | 452 | 618 | 626 | 627 |
| TGGGTGAGGTGCCCCCATCG | USP10 | 322 | 820 | 1422 | 1378 | 1582 |
| AACCTTACCCAATTGGACAA | USP13 | 537 | 488 | 734 | 728 | 754 |
| ATGGAGGCAGCAACCAACAA | USP13 | 514 | 501 | 876 | 839 | 819 |
| CCTGAAGAGACATGTGCGAG | USP13 | 600 | 472 | 1018 | 1035 | 949 |
| GTCTATTAGAGGAGAATTCG | USP13 | 523 | 627 | 1318 | 1123 | 1131 |
| TCCAGGACTAAGTTAGGACA | USP13 | 413 | 595 | 1101 | 1035 | 1004 |
| TGATATGCTTCATATGCATG | USP13 | 494 | 979 | 1864 | 1867 | 1789 |
| ACAAGGACAATATCTTCAAC | USP14 | 526 | 363 | 513 | 497 | 568 |
| AGAGGAACAGTTAGCATCTG | USP14 | 630 | 769 | 1523 | 1531 | 1467 |
| CTCACCGGAATAAAGCGGCA | USP14 | 404 | 499 | 711 | 597 | 697 |
| GAAGAAGAAGAGGTTACTAA | USP14 | 468 | 443 | 648 | 584 | 572 |
| GGAATGACTCTACTAATGAT | USP14 | 494 | 503 | 929 | 881 | 913 |
| TCAGCACAGTATATTACTGC | USP14 | 343 | 309 | 603 | 503 | 538 |
| CCATGCCGTAGAGCACTACA | USP5 | 476 | 530 | 812 | 686 | 695 |
| GCTTTAGGAACCCGGATCGT | USP5 | 654 | 585 | 1028 | 842 | 813 |
| GGAAAAGAAGCGGCAAGCTG | USP5 | 544 | 545 | 1152 | 971 | 1011 |
| GGACCTGGCTACATACACCT | USP5 | 610 | 453 | 1020 | 805 | 777 |
| GGATAAGACGATGACGGAGT | USP5 | 528 | 403 | 893 | 656 | 689 |
| TTCGCACCCGATCTCTGACG | USP5 | 490 | 861 | 1327 | 1365 | 1377 |
| AAAACTTACGCAACTCCATG | USP7 | 577 | 540 | 821 | 840 | 852 |
| AGATTTCGCACAAAACACGG | USP7 | 392 | 382 | 598 | 527 | 537 |
| GATATCCGATACATACCCGA | USP7 | 512 | 266 | 426 | 324 | 320 |
| GTGTAAATTTGACGACGACG | USP7 | 476 | 551 | 935 | 956 | 976 |
| TGACAATAAATACGACGCCG | USP7 | 554 | 327 | 679 | 582 | 560 |
| TGTCACACATAACCGGAAGC | USP7 | 547 | 498 | 879 | 847 | 940 |
| ACAAGTGCTGTACCTAACAG | USP9X | 519 | 744 | 1419 | 1285 | 1385 |
| ATGTGACAGTGATTATCACG | USP9X | 403 | 748 | 1308 | 1250 | 1248 |
| GAAGTTGATTGATTAAGTCA | USP9X | 612 | 722 | 1196 | 1025 | 1067 |
| TCTACTGGCAAAGTTAGCAT | USP9X | 580 | 542 | 770 | 765 | 766 |
| TGGATTAGGAATGCTCTGAA | USP9X | 600 | 763 | 1257 | 1130 | 1077 |
| TGGCCACACATAGTTCCACC | USP9X | 353 | 288 | 490 | 627 | 598 |
| ACGCATGATGTCCACCACCT | VAMP1 | 405 | 571 | 1130 | 1096 | 1042 |
| CACCCACTGAAGGGGCAGAA | VAMP1 | 304 | 572 | 995 | 1042 | 1105 |
| CAGACGACTACAGCAAACCC | VAMP1 | 640 | 363 | 538 | 555 | 595 |
| CCACCTGCAAGGCATCCGCT | VAMP1 | 517 | 336 | 736 | 722 | 728 |
| CTGTTACTGGTTGTATTAGG | VAMP1 | 405 | 718 | 1537 | 1453 | 1345 |
| GGACATCATGCGTGTCAATG | VAMP1 | 546 | 771 | 1588 | 1526 | 1575 |
| AATGAGACTCATGGGCATGT | VCP | 527 | 480 | 936 | 888 | 837 |
| AGGGGAGCCTATCAAACGAG | VCP | 737 | 669 | 1391 | 1200 | 1215 |
| AGGGTGTCATGCCAAACTTG | VCP | 625 | 916 | 1898 | 1805 | 1832 |
| CCATAGAGCAAGATTCCCCG | VCP | 323 | 307 | 510 | 382 | 403 |
| GCTACCCATCGATGACACGG | VCP | 569 | 480 | 1060 | 759 | 758 |
| GGTCCTTACCAAATCGCCGA | VCP | 617 | 217 | 469 | 450 | 538 |
| ACCCAGAGCAACTTTGCAGT | VDAC1 | 541 | 637 | 1089 | 1122 | 1209 |
| CAAAATCCACATCACAGCCC | VDAC1 | 583 | 649 | 993 | 887 | 867 |
| CGAGACTCTTACCCTGTGTT | VDAC1 | 521 | 600 | 956 | 1022 | 1086 |
| GTCTGGAAACCAAGTACAGA | VDAC1 | 369 | 336 | 595 | 515 | 532 |
| TGGAACACTGACAACACTCT | VDAC1 | 672 | 718 | 913 | 986 | 964 |
| TTGCAGTCTCAAAATTCATC | VDAC1 | 493 | 267 | 592 | 563 | 583 |
| ACAAGAAATAACTTTGCAGT | VDAC2 | 511 | 479 | 798 | 568 | 680 |
| CAAAGTCAACATCACAACCA | VDAC2 | 410 | 349 | 593 | 607 | 646 |
| CCAAGGTCAGCATAGGATGG | VDAC2 | 470 | 860 | 1631 | 1716 | 1596 |
| CCAGCCCTCATAACCAAAGA | VDAC2 | 568 | 391 | 726 | 569 | 598 |
| CTTTCAAAATAGATCTGTCA | VDAC2 | 495 | 258 | 573 | 438 | 502 |
| GTGAAAACAAAGTCATGCAG | VDAC2 | 596 | 327 | 669 | 622 | 667 |
| AAGCCTTGAGGGACAGTCTG | VDR | 323 | 1058 | 1384 | 1508 | 1685 |
| CGGGGACTGTCGTATCACCA | VDR | 489 | 632 | 1094 | 1150 | 1112 |
| CTGACCAGGTCAGCTAGGTG | VDR | 431 | 436 | 951 | 1039 | 1035 |
| GAACGTGCCCCGAATCTGCG | VDR | 677 | 770 | 1109 | 1162 | 1090 |
| GAACTGGGTGAAGTCGGCAT | VDR | 650 | 543 | 931 | 1078 | 1027 |
| GCTGAAATCAAGTGCCATCG | VDR | 406 | 520 | 1281 | 1428 | 1345 |
| AGATGTACTCGATCTCGTCA | VEGFA | 442 | 522 | 820 | 790 | 815 |
| AGCAAATGTGAATGCAGGTG | VEGFA | 406 | 863 | 1672 | 1780 | 1709 |
| ATGCGGATCAAACCTCATCA | VEGFA | 499 | 552 | 981 | 921 | 858 |
| CCTAGAGTGCGTGCCCACTG | VEGFA | 559 | 604 | 1346 | 1281 | 1365 |
| GCAGCCCCCACACCGCATCA | VEGFA | 530 | 623 | 1043 | 1002 | 1037 |
| GGAGAGCACAAACCCCACGA | VEGFA | 456 | 460 | 884 | 869 | 863 |
| CAGAAGCCCATCATATGTCC | VHL | 626 | 324 | 620 | 574 | 539 |
| CCCGGCGGCAGCGTCGAGTA | VHL | 598 | 759 | 1119 | 1408 | 1344 |
| CCGTCGAAGTTGAGCCACAC | VHL | 536 | 429 | 844 | 638 | 744 |
| CGCCGCATCCACAGCTACCG | VHL | 538 | 433 | 856 | 654 | 700 |
| GAGTTCACCGAGCGCAGCAC | VHL | 494 | 310 | 621 | 571 | 578 |
| TGTCCATCAACATTGAGAGA | VHL | 577 | 604 | 1075 | 906 | 882 |
| AGATACAAAAAAGGAATCAG | WAS | 583 | 558 | 706 | 664 | 710 |
| ATCACGAGTTCACGGTACCG | WAS | 567 | 845 | 1537 | 1692 | 1620 |
| CAGGTACAGCTGAACAACAG | WAS | 427 | 529 | 991 | 1129 | 1061 |
| GCAGGTAAACAACCTGGACC | WAS | 643 | 594 | 936 | 946 | 981 |
| GGATGAAATAGGACTTCTGG | WAS | 530 | 645 | 1151 | 1173 | 1199 |
| GTGTGTAGGACTGCCAAGCA | WAS | 644 | 663 | 1034 | 1068 | 1102 |
| AAAGGATTAATATTCACTTG | WEE1 | 477 | 537 | 1106 | 981 | 991 |
| AGCCCCGGCCAACTTGCAAA | WEE1 | 578 | 623 | 1166 | 1286 | 1236 |
| AGTTTACTCTCCAAAGCTCG | WEE1 | 432 | 713 | 1515 | 1313 | 1424 |
| GAGGAGGACATGTTGCTGCG | WEE1 | 524 | 447 | 859 | 707 | 731 |
| TACATACTCATCGACAGAGC | WEE1 | 494 | 264 | 365 | 370 | 356 |
| TCGTTCTCGCCGGTGCGCTG | WEE1 | 668 | 564 | 1124 | 945 | 955 |
| AGGTATCACACTTATTGCTG | WRN | 357 | 945 | 1753 | 1655 | 1645 |
| AGTTCCCATAATACCCACGG | WRN | 650 | 633 | 882 | 842 | 888 |
| CAGATAATATTTAACCTCTG | WRN | 528 | 834 | 1258 | 1125 | 1181 |
| CAGGCTTGGATATTAAATCA | WRN | 407 | 619 | 1317 | 1265 | 1254 |
| CATTACCAAGACCTAATGCA | WRN | 541 | 548 | 597 | 648 | 697 |
| GGTGTTTAACCAAACCATTG | WRN | 380 | 712 | 1282 | 1150 | 1222 |
| CAGCCCGCTATTCGCAACCA | WT1 | 669 | 670 | 1246 | 1173 | 1135 |
| CAGTACAGAATACACACGCA | WT1 | 573 | 542 | 895 | 892 | 982 |
| CCAGGTCATGCATTCAAGCT | WT1 | 489 | 466 | 1171 | 1119 | 1144 |
| GGGGGTGCAAAATCCAGCAC | WT1 | 398 | 452 | 807 | 739 | 755 |
| TCCTGCTTGATGAAGGAGTG | WT1 | 380 | 914 | 1684 | 1620 | 1618 |
| TCGTGCTGAAAGAGTGGTTG | WT1 | 565 | 1025 | 1640 | 1605 | 1749 |
| AAAGTAAAATGCTTTCACTG | XIAP | 360 | 660 | 791 | 861 | 812 |
| AGCACTAGCTAACTCTCTTG | XIAP | 607 | 972 | 1897 | 1990 | 1866 |
| ATAATGCATGGTCAGAACAC | XIAP | 403 | 334 | 712 | 702 | 757 |
| CTGATGTTGCGAGTACTGAT | XIAP | 492 | 655 | 1201 | 1237 | 1226 |
| CTGGTGAAGGAGATACTGTG | XIAP | 400 | 789 | 1191 | 1349 | 1362 |
| TATATCAGACACCACATACC | XIAP | 301 | 292 | 519 | 553 | 531 |
| CCGGCCCTACCCGGCGACGG | XPA | 442 | 548 | 971 | 1061 | 1070 |
| CGGTGCGGGCGAGCATCGAG | XPA | 551 | 738 | 1438 | 1402 | 1387 |
| GCCCCGAAAATCATTGACAC | XPA | 466 | 272 | 591 | 695 | 651 |
| TATGCCATATGTGAAGAATG | XPA | 439 | 596 | 1246 | 1243 | 1261 |
| TGGCAACTTGTGATAACTGC | XPA | 499 | 449 | 1014 | 962 | 982 |
| TTCTTAACAATAAATTTCAG | XPA | 671 | 373 | 560 | 507 | 509 |
| AAGCCCATGACCTACGTGGT | XPC | 489 | 590 | 776 | 743 | 800 |
| ACTGACTCCAGGACCCAAAG | XPC | 410 | 527 | 1189 | 1004 | 971 |
| AGATGGCCTACAGACTACGT | XPC | 503 | 305 | 797 | 670 | 701 |
| AGCTACCTACGGCAAAGGTG | XPC | 379 | 774 | 1311 | 1216 | 1253 |
| AGTTTGAGACTTATCTCCGG | XPC | 297 | 623 | 1133 | 986 | 1081 |
| GGATGAAGCCCTCAGCGACG | XPC | 488 | 649 | 1183 | 1146 | 1151 |
| AAATCTCAGGGAAACACTCA | XPO1 | 534 | 374 | 659 | 601 | 566 |
| ATTGTTACCTTGGATAACAC | XPO1 | 645 | 238 | 345 | 291 | 327 |
| GAACCTGAACAAAATGCCTG | XPO1 | 621 | 539 | 1018 | 915 | 872 |
| GTGACATTGTTGGAGCAAGT | XPO1 | 381 | 330 | 691 | 509 | 560 |
| TCCAAAGTTGCATGTACAAG | XPO1 | 591 | 949 | 1363 | 1468 | 1575 |
| TGTAAGTGTAAGCCAATACG | XPO1 | 590 | 439 | 962 | 776 | 783 |
| ACATCGATCAGACGACGACG | YAP1 | 594 | 431 | 988 | 942 | 1079 |
| ATGGGAACAAGCCATGACTC | YAP1 | 517 | 640 | 1047 | 963 | 1003 |
| CATCATACTCTGCTGCACTG | YAP1 | 501 | 618 | 779 | 703 | 690 |
| GATGAACCTTTACCAAAACG | YAP1 | 442 | 618 | 1038 | 916 | 883 |
| GGATCCCTTTCTCAACAGGT | YAP1 | 538 | 487 | 963 | 752 | 808 |
| TCAGTTCCTGAGACATCCCG | YAP1 | 447 | 437 | 881 | 877 | 871 |
| AAACCAGCAGGATATGAACT | YES1 | 581 | 665 | 1070 | 1105 | 1071 |
| AAGTTGGTAAAACACTACAC | YES1 | 588 | 483 | 823 | 818 | 837 |
| CTAGTCGCAAAGATTCTCGA | YES1 | 382 | 366 | 636 | 616 | 600 |
| CTGAATCCTGGGAATCAACG | YES1 | 530 | 445 | 794 | 811 | 849 |
| GAAGCAAGATCAATTGCTAC | YES1 | 421 | 425 | 643 | 651 | 672 |
| GCAATCAAAACACTAAAGCC | YES1 | 578 | 344 | 454 | 436 | 460 |
| CATCTCGAAGCATTCCAGCG | ZBTB16 | 346 | 901 | 1205 | 1283 | 1290 |
| GATTCTGGAGTACGCGTACA | ZBTB16 | 584 | 833 | 1407 | 1390 | 1449 |
| GGAGCTGCATTACGGGCGTG | ZBTB16 | 531 | 1020 | 1785 | 1635 | 1669 |
| GGCCGGGACGCTGTGCGATG | ZBTB16 | 558 | 1140 | 1903 | 1896 | 1872 |
| GTCGGAGAGCCGGACCATCG | ZBTB16 | 554 | 701 | 1243 | 1202 | 1215 |
| GTGGACAGTTTGATGACCAT | ZBTB16 | 509 | 613 | 1079 | 1084 | 1047 |
| AATGCAGGAATGGGTAATAG | ZMYM2 | 637 | 481 | 947 | 975 | 891 |
| AGAGGTTCGAGACCATATGC | ZMYM2 | 527 | 515 | 851 | 796 | 811 |
| ATAGGTACTGGGATATACAC | ZMYM2 | 330 | 359 | 482 | 573 | 612 |
| ATATTGCGAATACTGTCAAG | ZMYM2 | 408 | 451 | 890 | 930 | 925 |
| ATTTCAGGATTGGTACTACA | ZMYM2 | 703 | 415 | 816 | 705 | 693 |
| TCGTAATCAGAAACAACCAG | ZMYM2 | 374 | 512 | 711 | 744 | 725 |
| AAGTCCAAGACCAGTAAGAG | ZNF217 | 528 | 682 | 1239 | 1008 | 1114 |
| AGGTCCGGAGAACACGTCCG | ZNF217 | 698 | 1043 | 1610 | 1581 | 1635 |
| CAAACAGATCGATCTTGCCT | ZNF217 | 464 | 530 | 780 | 742 | 743 |
| GAGCCCGGTAACCATCAACG | ZNF217 | 540 | 894 | 1249 | 1156 | 1093 |
| GCGTTACAGAAGAGACCGCC | ZNF217 | 477 | 326 | 561 | 509 | 592 |
| GTGGTGAACGGATCGAGCTG | ZNF217 | 375 | 885 | 1610 | 1477 | 1487 |
| AGAGCGCGCCGTAGGCCTTG | ZNF668 | 448 | 1027 | 1764 | 1691 | 1775 |
| AGGACCTACGCAACCACGAG | ZNF668 | 519 | 718 | 1165 | 1199 | 1293 |
| GAGAACATGCGGCCGCAACG | ZNF668 | 445 | 717 | 1211 | 1221 | 1230 |
| GCAGTCTGCAGGACCATGTG | ZNF668 | 509 | 603 | 1257 | 1271 | 1287 |
| GCAGTGTGACAAGACGTTTG | ZNF668 | 624 | 903 | 1693 | 1660 | 1682 |
| GGTGTTTGCGGAACACCGAA | ZNF668 | 597 | 444 | 847 | 819 | 854 |
| ATCGTCCGTTAAACAGAGAG | ZRSR2 | 424 | 837 | 1400 | 1269 | 1272 |
| ATTATCCTTAAGTCGATGGG | ZRSR2 | 397 | 619 | 1465 | 1105 | 1118 |
| GAAACGAAAGAAACGACGGC | ZRSR2 | 628 | 397 | 634 | 610 | 573 |
| GGCAAAAATTACATGAAGCG | ZRSR2 | 568 | 405 | 676 | 677 | 692 |
| TGTACTCTAGGCTCGCGTCT | ZRSR2 | 405 | 621 | 1365 | 1224 | 1209 |
| TTCCGGAGTTCAAGAACGTG | ZRSR2 | 352 | 513 | 1206 | 877 | 893 |

**Table S6.** Canine mammary tumor organoid medium components

|  | Medium components | Company | Catalogue number | Final concentration |
| --- | --- | --- | --- | --- |
| ADDF+++ | GlutaMax 100x | GIBCO | 35050-038 | 1x |
|  | Hepes 1M | GIBCO | 15630-056 | 10 mM |
|  | Penicillin/Streptomycin | GIBCO | 15070-063 | 50 U·ml-1 |
|  | Primocin | Invivogen | Ant-pm-1 | 50 mg·ml^-1^ |
|  | Advanced DMEM/F12 | GIBCO | 12634-010 | 1x |
| Growth factors | R-Spondin 3 (RSPO3-Fc Fusion Protein) conditioned medium | U protein express | R001 | 10% |
|  | Noggin-Fc fusion Protein conditioned medium | U protein express | N002 | 2% |
|  | Neuregulin 1 | Peprotech | 100-03 | 5 nM |
|  | FGF 7 | Peprotech | 100-19 | 5 ng·ml^-1^ |
|  | FGF 10 | Peprotech | 100-26 | 20 ng·ml^-1^ |
|  | EGF | Peprotech | AF-100-15 | 0.5 ng·ml^-1^ |
|  | A83-01 | Tocris | 2939 | 500 nM |
|  | Y-27632 | Abmole | Y-27632 | 5 mM |
|  | SB202190 | Sigma | S7067 | 1 mM |
|  | B27 supplement | Gibco | 17504-44 | 1x |
|  | N-Acetylcysteine | Sigma | A9165-5g | 1.25 mM |
|  | Nicotinamide | Sigma | N0636 | 10 mM |

**Table S7.** Primary antibodies used for immunohistochemistry

| Antibody against | Clone | Source^1)^ | Antibody Dilution  Antibody Concentration^2)^ | Pretreatment of slides^3)^ | ^4)^ |
| --- | --- | --- | --- | --- | --- |
| Cytokeratin 5/6 (CK5/6) | D5/16B4 | Dako (Agilent) | 1:100 0.51 µg/ml | H2(30) | B |
| CAM5.2 (CK7/8) | CAM 5.2 | BD Biosciences | 1:150 0.505 µg/ml | H2(20)95 | B |
| Cytokeratin 14 (CK14) | LL002 | Leica Novocastra | 1:150 0.16 µg/ml | H2(40)95 | B |
| p63 | 4A4 | Biocare Medical | 1:150 | H2(30) | B |
| MIB-1 (Ki-67 Antigen) | MIB-1 | Dako (Agilent) | 1:50 0.92 µg/ml | H2(30) | B |
| Estrogen Receptor (ER) | EP1 | Dako (Agilent) | 1:50 3.00 µg/ml | H2(40) | B |
| Progesteron Receptor (PR) | polyclonal | Abcam | 1:100 | H2(30) | B |
| Pathway HER2/neu (Her2) | 4B5 | Ventana (Roche) | RTU 6 µg/ml | CC1(36) | V |
| Vimentin | V9 | Dako (Agilent) | 1:1000 0.32 µg/ml | H2(10)95 | B |

^1)^	Abcam, Cambridge, UK; BD Biosciences, San Jose CA, USA; Biocare Medical, Pacheco CA,USA; Dako, Glostrup, Denmark; Leica Biosystems (Novocastra), Newcastle-upon-Tyne, UK; Ventana Medical Systems (Roche Diagnostics), Tucson AZ, USA.

^2)^	Dilutions and working concentrations are given where known; dilutions only are given for antibodies where the Ig concentration is not specified by the manufacturer. RTU = Ready to use antibody.

^3)^	H2(30): Preatretment with Epitope Retrieval Buffer Type 2 (Tris-EDTA, pH 9) (Leica Biosystems) for 30 minutes at 100°C on Bond-III immunostainers. H2(40): idem, 40 minutes at 100°C. H2(20)95: idem, 20 minutes at 95°C. H2(40)95: idem, 40 minutes at 95°C. H2(10)95: idem, 10 minutes at 95°C.

CC1(36): Preatretment with Ultra Cell Conditioning Solution CC1 (Tris) (Roche) for (36) minutes at 95°C on Ventana BenchMark ULTRA immunostainers

^4)^	Immunostainer used: B = Bond-III (Leica Biosystems); V = Benchmark Ultra (Ventana Medical Systems).

**Table S8.** Primers sequences used for sanger sequencing and TIDE analysis and screen analysis

| Gene | guide | Primer | Sequence (5´-3´) | Tm(°C) for PCR protocol (*) |
| --- | --- | --- | --- | --- |
| VIM | sgRNA1 | Forward | ACCATGTCCACCAGGTCTGT | 63°C |
|  |  | Reverse | CGGAGGAGCGCGTGGCGTAC |  |
|  | sgRNA2 | Forward | GCTCACCAATTTTTGGCACT | 63°C |
|  |  | Reverse | GAAAAGTGCTGGTGCTCACA |  |
| AKT1 | | Forward | CTGTCCCCTCGATGCCAC | 64°C |
|  |  | Reverse | GCCACGGAGAAGTTGTTGAG |  |
| PIK3CA | | Forward | CCCATTTGCTCCATACTGATCA | 63°C |
|  |  | Reverse | CATGCTGCTTAATGGTGTGGA |  |
| KRAS | | Forward | TCGTCACTGAATTTTCTGAAGCA | 63°C |
|  |  | Reverse | TGGGCCTGCACAAATCAATA |  |
| TP53 | | Forward | TTCCCATCCATTCCTGTCCC | 64°C |
|  |  | Reverse | CCCTCCTTCACCTCCTCTTG |  |
| P5 primer: AATGATACGGCGACCACCGAGATCTACACTCTTTCCCTACACGACGCTCT TCCGATCT[s] TTGTGGAAAGGACGAAAC*A*C*C*G, where [s] is the barcode region. | | | | |
| P7 primer:  CAAGCAGAAGACGGCATACGAGAT [s] GTGACTGGAGTTCAGAC  GTGTGCTCTTCCGATCTCCAATTCCCACTCCTTTCAAG*A*C*C*T, where [s] is the barcode region. | | | | |

(*) Target loci were amplified following a 3-steps protocol: (a) 98°C for 30 s, (b) 35 cycles at 98°C for 10 s, Tm(°C) for 20 s, and 72°C for 20 s, (c) 72°C for 5 min.

**Table S9.** Primers sequences used for gene editing

| Gene | guide | Sequence (5´-3´) | Exon targeted |
| --- | --- | --- | --- |
| Vimentin | sgRNA1 | GTACGCCACGCGCTCCTCCG | 1 |
|  | sgRNA2 | AGACACTATTGGCCGCCTGC | 6 |
| Non-targeting gRNA | | GTGATTGGGGGTCGTTCGCCA | No gene targeted |
